# Mapping and Probing the Substrate Scope of a Ketoamide-Forming Splicease by Computational Design

**DOI:** 10.64898/2026.11.03.763875

**Authors:** Thomas H. Jordan, Rosalie Lipsh-Sokolik, Jörn Piel

**Author notes:** **Corresponding Author:** Jörn Piel: Institute of Microbiology, Department of Biology, Eidgenössische Technische Hochschule (ETH) Zurich, Switzerland.

## Abstract

α-Keto-β-amide warheads are protease-inhibiting pharmacophores in peptidomimetic antiviral drugs and drug leads, but their diversity-oriented synthesis for library-based drug discovery can be challenging. The same moiety is installed enzymatically in gene-encoded peptides by tyramine spliceases from spliceotide natural products. However, their substrate scope remains poorly understood. We mined bacterial spliceotide pathways for multi-precursor spliceases and identified HypX from *Hyella patelloides* that modified 5 of 7 native precursors in *E. coli*. Single-point mutagenesis showed promiscuity for HypX and the previously identified PlpX but different substrate scopes, suggesting that splicease mining can provide enzymes for focused libraries of various substrate types. To evaluate multipoint variants without exhaustive screening, we combined AlphaFold3 models with Rosetta calculations to rank millions of variants. Top-scoring peptides were spliced more efficiently than single-point mutants. Structural modeling rationalized position-specific tolerance, including a cofactor constraint. This work establishes a mining- and structure-guided strategy to streamline library-based splicease applications.

## INTRODUCTION

α-Keto-β-amides (here referred to as ketoamides) are privileged warheads in peptidomimetic drug design due to the potent activity of the ketoamide moiety against various serine and cysteine proteases.^1–5^ The α-keto group acts as an excellent electrophile for the catalytic serine or cysteine residues in protease active sites that normally cleave peptide bonds. Several US Food and Drug Administration-(FDA-) approved drugs contain ketoamide units, such as the hepatitis C virus (HCV) protease inhibitors telaprevir and boceprevir.^2,6^ Recent public health emergencies, such as the SARS-CoV2 pandemic, have refocused efforts on finding drug leads against therapeutically relevant viral proteases. The ketoamide warhead has been identified as a promising lead motif in SARS-CoV2 main protease inhibitors.^1,2,4,5,7^ Ketoamide drug leads are usually derived from synthetic peptidomimetics, but their diversity-oriented synthesis can be challenging.

Recently, our group discovered that the ketoamide warhead can be enzymatically introduced into gene-encoded peptides via a non-canonical post-translational modification (PTM) installed by tyramine spliceases.^8^ These are specialized radical *S*-adenosyl-L-methionine (rSAM) enzymes that catalyze the remarkable excision of a tyramine unit from an XYG motif (X = variable amino acid, Figure 1A) in a ribosomally translated peptide or protein precursor. Reconnection of the up- and downstream parts generates the ketoamide moiety, i.e., a β-amino acid residue. This modification occurs in a family of ribosomally synthesized and post-translationally modified peptides (RiPPs) called spliceotides that are widespread in bacteria, with the first recorded examples being the *plp* and *pcp* loci from *Pleurocapsa* spp. cyanobacteria. Both BGCs encode, in addition to the precursors, the spliceases PlpX and PcpX, respectively, and a small accessory “Y protein” (PlpY, PcpY) that mediates substrate-maturase interaction.^8,9^ Like their synthetic ketoamide counterparts, the introduction of the ketoamide motif in peptides was shown to result in potent protease inhibitory activities, as well as activity against a range of viruses.^10,11^

**Figure 1.**
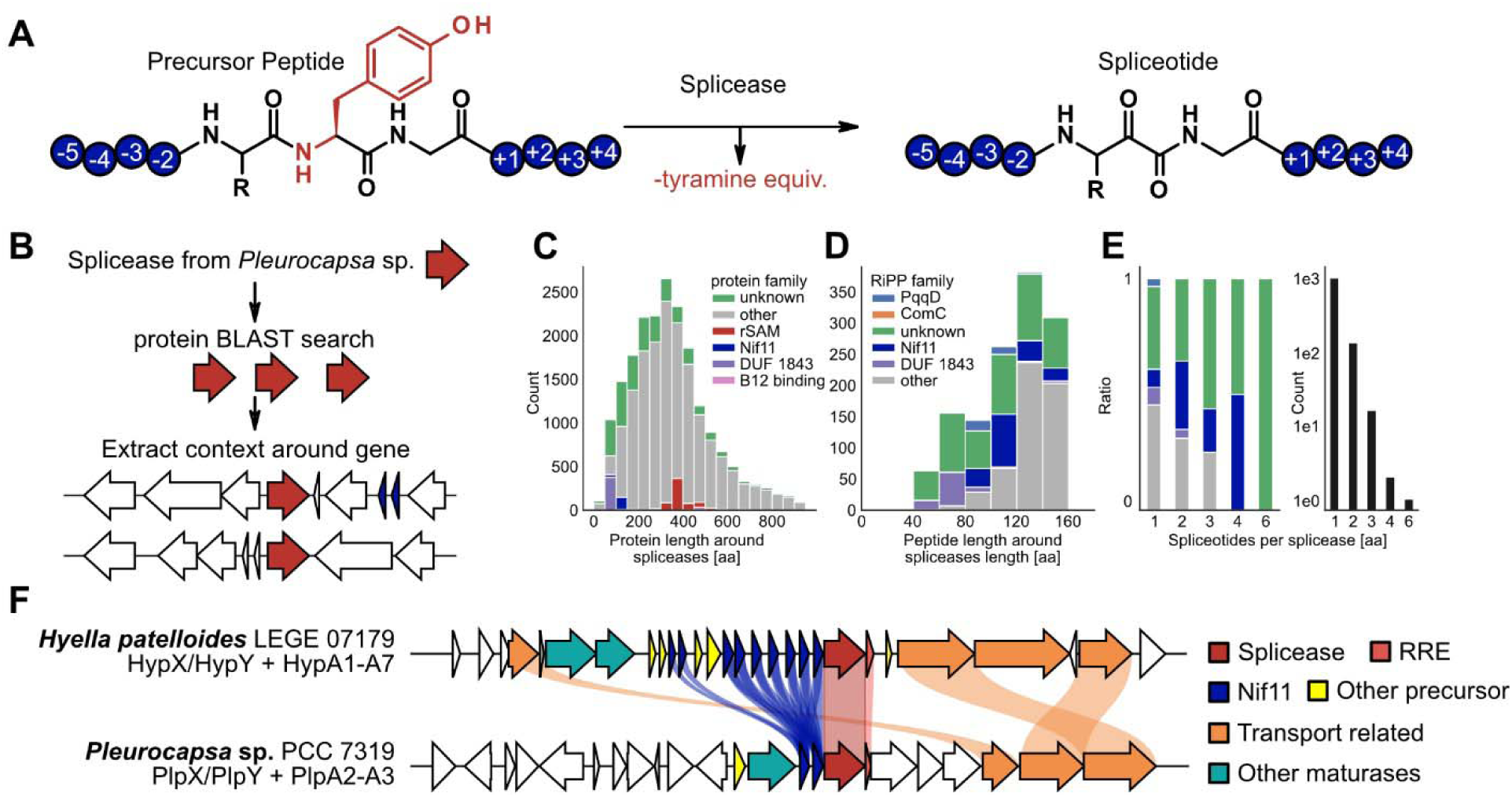
Spliceotides and mining for new spliceases. **(A)** Overview of the splicing reaction catalyzed by spliceases. Precursors produced by the ribosome containing an XYG motif (X = variable amino acid) are modified by spliceases in a radical reaction, leading to the loss of a tyramine equivalent and the formation of an α-keto-β-amide with the X side chain at the splice site. **(B)** From a previously generated BLAST dataset of splicease homologs,^11^ the genomic neighborhood of the encoding genes was extracted and annotated. **(C)** Bar graph representing the most common annotations for proteins encoded in proximity to putative splicease genes. **(D)** Bar plot of putative precursors, including short proteins containing a conserved protease cleavage motif known from RiPPs. **(E)** Stacked bar plot showing the relative distribution of precursor types. If loci contained splicease genes adjacent to multiple putative precursor genes, the precursors predominantly belonged to the Nif11 or an unclassified type **(F)** Clinker alignment of the *Hyella patelloides* and the *Pleurocapsa* sp. spliceotide BGCs.^38^ The alignment shows the prevalence of Nif11 type precursors in the BGC, of which *H. patelloides* has many.

RiPPs are a large peptide natural product superfamily featuring diverse other modifications in addition to tyramine splicing, depending on the pathway type.^12,13^ Bacterial RiPP biosynthetic gene clusters (BGCs) encode one or more genetically encoded precursors that in most cases comprise a C-terminal core and an N-terminal leader, as well as maturases that act on the core to introduce the modifications.^12,13^ As RiPP precursors are gene-encoded, non-natural peptide variants can be generated by changing the coding sequence of the precursor, making RiPP maturases attractive tools to generate chemically diverse libraries of post-translationally modified peptides. Indeed, previous studies showed remarkable substrate promiscuity for several RiPP maturases, such as the cyanobactin heterocyclases, prochlorosin lanthionine synthetases, and proteusin epimerases.^14–16^ However, related maturases can also exhibit marked differences in substrate scope. For example, the cyanobactin heterocyclases PatD and TruD share high sequence identity, yet PatD has a much broader substrate tolerance and forms different heterocyclic rings compared to TruD, which only forms thiazolines.^17,18^ Thus, an initially limited substrate range for a particular RiPP maturase type might be broadened by exploring further enzyme homologs.

The initially discovered PlpX and PcpX spliceases and their homologs have been used to modify various non-natural precursor sequences containing XYG motifs: truncations of the natural core sequences,^8,19,20^ sequences mimicking known protease inhibitors,^8^ precursors fully or partially lacking a leader sequence,^19,21,22^ individual point mutants derived from wild-type sequences,^20^ as well as proteins with introduced splice motifs.^19,21,22^ While these studies establish promiscuity, they mostly offer a qualitative snapshot of splicease activity in contexts that are not necessarily comparable. Although recent progress has been made toward characterizing spliceases within standardized synthetic biology frameworks,^20,23^ a systematic, predictive understanding of their broader substrate range remains an open challenge. Because successful precursor splicing relies on liquid chromatography–mass spectrometry (LC-MS) for verification, investigating the splicease substrate scope is inherently slow and labor-intensive. The primary bottlenecks stem from the limited throughput of peptide co-expression and purification workflows, coupled with extended downstream measurement times.

Despite the considerable biotechnological potential, few studies have investigated the substrate scope of spliceases in depth and assessed their use for producing spliceotide libraries. Here we set out to further investigate the substrate scope of spliceases, characterize a new splicease representative, and explore how combinations of *in-silico* prediction and *in-vitro* validation could help predict whether a given peptide will be modified.

## RESULTS AND DISCUSSION

### Identification and characterization of a substrate-tolerant splicease

Many RiPP BGCs that encode promiscuous maturases also harbor multiple genes for precursors with distinct cores. For example, the prochlorosin lanthionine synthetase ProcM acts on 29 precursors, of which 7 are encoded in the same BGC.^14,24,25^ The presence of diverse precursor genes in the same BGC is therefore suggestive of maturases with broad substrate tolerance that might be suitable for generating artificial peptide libraries.^26–31^ Based on this rationale, we set out to identify further candidates for promiscuous spliceases based on how many putative precursors are in the immediate genetic vicinity of a splicease homolog. Starting from a dataset of putative spliceotides, created by searching for splicease homologs and detecting XYG motifs in translated neighboring genes,^11^ our analysis indicated that most of the BGCs encoded candidates for a single splicease and one or two XYG-containing precursors (Figure 1B-E). Among the BGCs with the highest number of putative precursor genes, one from the cyanobacterium *Hyella patelloides* LEGE07179, termed *hyp* BGC, contains genes for seven putative spliceotide precursors with XYG motifs (HypA1-HypA7), the splicease HypX and its accessory protein HypY, five precursors lacking XYG motifs, as well as YcaO and TfuA homologs predicted to install thioamide bonds.^32^ The five precursors lacking the canonical splice site were not considered further, as they were unlikely susbtrates for splicing. Sequence alignments of the 7 XYG-containing precursors showed that they vary both in the length of the core region as well as amino acid composition (Figure 2A), including the presence of six different residues (AVTCMP) at the X position, suggesting high substrate promiscuity. All share a highly conserved Nif11-like leader sequence also found in the *Pleurocapsa* PlpA2, PlpA3, and PcpA peptides. Since the X residues in the XYG motifs are extended by one carbon from tyrosine during splicing, differences in their side chain result in different ketoamide residues. This suggested that HypXY would be able to generate valine (HypA1 and HypA2), cysteine (HypA3), threonine (HypA4), alanine (HypA5), methionine (HypA6), and proline ketoamides (HypA7), in addition to tolerating variable residues near the XYG motif. We also considered that, based on previous observations,^11^ several spliced ketoamides can undergo further spontaneous modifications. Specifically, serine sidechains can undergo a retro-aldol like reaction, eliminating formaldehyde to produce a glycine ketoamide, whereas cysteine ketoamides can spontaneously form dehydroalanine ketoamides via hydrogen sulfide elimination.^11^ In addition, the TYG motif from HypA4 might similarly produce either a dehydrobutyrine ketoamide via the elimination of a water molecule, or undergo a retro-aldol reaction liberating an acetaldehyde to yield a glycine ketoamide.

**Figure 2:**
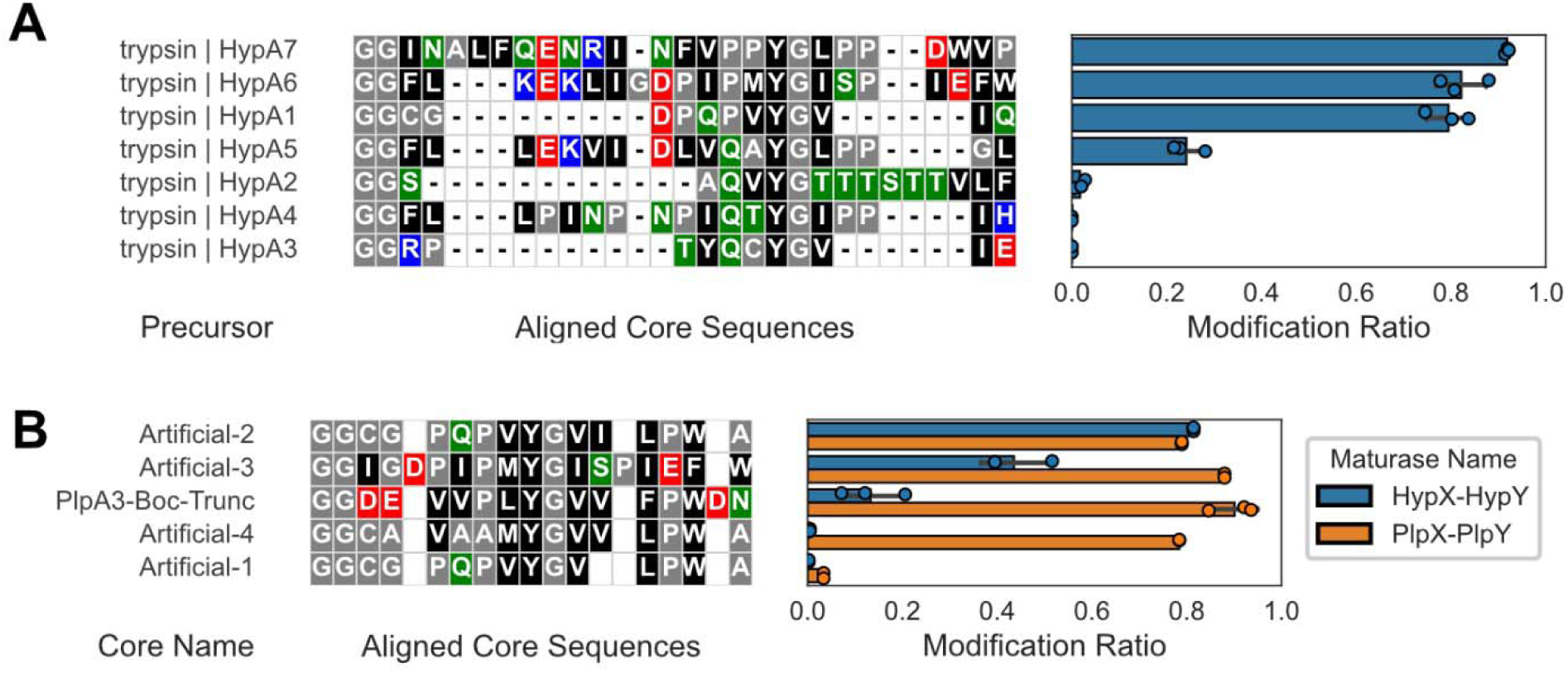
Modification of precursors by the HypXYand PlpXY splicease systems. **(A)** Sequence alignment of the 7 precursor cores, including the conserved leader-core interface motif GG, from the *H. patelloides* BGC, showing the conserved YG motif but different amino acid context of the precursors. The bar plot represents the turnover of the precursor to the mature spliceotide by the HypXY splicease. **(B)** Alignment of several artificial core sequences, with the core sequence “Artificial-2”, a core peptide that is modified efficiently by both HypX-HypY and PlpX-PlpY, and thus serves as the neutral reference point forsubsequent perturbations using point mutations. Bars show the mean ± SD of n = 2-3 biological replicates; points are individual biological replicates.

To assess whether the *hyp* splicease is capable of modifying the diverse putative precursors, high-copy number plasmids encoding the splicease HypX bicistronically alongside HypY under the control of a T7 promoter were constructed. In previous studies from our lab, we used the Duet plasmid system (Novagen) for expression. Of these, we generated Golden-Gate domesticated versions for this work to allow for the rapid, combinatorial cloning of different plasmid variants, speeding up characterization and troubleshooting. We adopted an approach used by Voigt and co-workers of using the RBS calculator to standardize RBS translation strengths for RiPP maturases and refactored the bicistronic transcription unit accordingly.^23^ In our case, precursors were individually produced from low-copy expression plasmids as N-terminally His_6_- and Fh8-solubility-tagged peptides under control of a T7 promoter, and maturase plasmids were then introduced into *E. coli* BL21(DE3). Heterologous production of HypXY alongside HypA1-HypA7 (Figure 2A) and Ni affinity purification of the precursors was followed by digestion of the precursors using trypsin protease. The resulting fragments were then analyzed by liquid chromatography mass spectrometry (LC-MS). Analysis of the LC-MS data confirmed tyramine excision and ketoamide formation for 5 out of the 7 candidate precursors (Figure 2A, Figures S1-7) via detection of ions matching the loss of a tyramine equivalent (C_8_H_9_NO, −135 Da). Tandem MS/MS fragmentation patterns localized this mass loss to the predicted XYG modification sites described in Figure 2A. The enzyme was thus able to modify precursors containing diverse XYG motifs in different amino acid contexts and core positions. No spliced product was observed for HypA3, which contains a CYG motif, nor was the dehydroalanine ketoamide detected that would result from H_2_S loss. Likewise, HypA4 containing a TYG motif remained unmodified based on our inability to detect the canonical, dehydrated, or retro-aldol ketoamide. Digestion of the precursors HypA3 and HypA4 by GluC as an alternative protease showed a similar result, with only unmodified fragments observed (Figures S8-10). In a slightly different setup, a consensus leader sequence derived from a sequence alignment of the HypA1-HypA7 leader sequences (HypAc, Table S4) was fused to the HypA1, A4, A5, A6, and A7 core sequences and co-produced alongside HypXY. After tryptic or GluC digest and LC-MS analysis, a similar modification trend was observed as with the native sequences (Figure S11) indicating that an artificial leader sequence can be substituted for the native leader (Figures S12-16). In the case of the HypA4 core, in contrast to earlier experiments, digestion of the precursor by GluC and analysis via LC-MS showed a very low amount of spliced peptide (−135 Da) as well as possible products corresponding to an additional dehydration (−C_8_H_11_NO_2_) or retro-aldol product (−C_10_H_13_NO_2_), similar to previous observations (Figures S17-18).^11^ Overall, these results indicate that the HypX splicease is a promising enzyme for producing various ketoamides in *E. coli*.

### Comparing the modification efficiency of PlpX and HypX

To compare the splicing efficiency of the spliceases HypX and PlpX, we generated four artificial peptide sequences that mimic the native cores of peptides from either cluster (Figure 2B). To facilitate the analysis of core-derived peptides via LC-MS, we introduced a recognition sequence for Tobacco Etch Virus (TEV) protease in between the leader and core sequences. Specifically, the consensus *Hyella* leader sequence was reused and a TEV protease recognition sequence introduced in front of the conserved GG motif at the leader-core interface. Similarly, we added a TEV protease cleavage site to the PlpA3 leader at the same position (Table S4). To compare the two spliceases, several synthetic core sequences were generated to identify a sequence modified with similar efficiency by both enzymes. The four cores Artificial-1 to Artifical-4 were then fused to the HypAc-TEV and PlpA3-TEV leader sequences for co-production with their respective splicease. Production and purification of the precursor followed by TEV protease digestion and LC-MS analysis indicated that most precursors were modified (Figure 2B, Figures S19-32). The cores Artificial-1 and Artificial-4 were efficiently modified by PlpX but not by HypX. In contrast, Artificial-2 was modified to an equal extent by PlpX and HypX, whereas Artificial-3 was modified by both but with a preference for PlpX (Figure 2B, Figures S19-32). For this reason, we established Artificial-2 as a reference sequence from which to compare the effect of individual point mutations. A previously described^8^ core sequence based on a C-terminally truncated PlpA3 core peptide that mimics boceprevir “PlpA3-boc-trunc” (Figure 2B, Figures S31-52) appeared to be modified more efficiently by PlpX than HypX which is contrary to our hypothesis of HypX being a more substrate tolerant splicease.

To increase experimental throughput, we changed expression conditions from shake flasks to deep 96-well plates. This downscaling generally reduced the modification efficiency of PlpX, and, to a slightly larger extent, HypX. Using the Artificial-2 core as a template, we substituted the three positions immediately preceding the spliced YG motif (positions –3 to –1), and the position immediately succeeding it to a small panel of diverse amino acids (D, F, G, H, I, K, M, P, Q, S, T) to investigate their effect on the peptide modification. Intriguingly, most sequence changes were well-tolerated by both enzymes compared to the parent sequence (Figure 3, Figures S53-133) with a modification efficiency similar or greater than for the Artificial-2 peptide. Similar to other RiPP maturase systems like the heterocyclases PatD and TruD,^18^ the two splicease homologs vary considerably in their amino acid preferences. Furthermore, the PlpX splicease demonstrated greater robustness towards those point mutations than the multi-precursor splicease HypX. For this reason, we set out to characterize PlpX in more detail.

**Figure 3.**
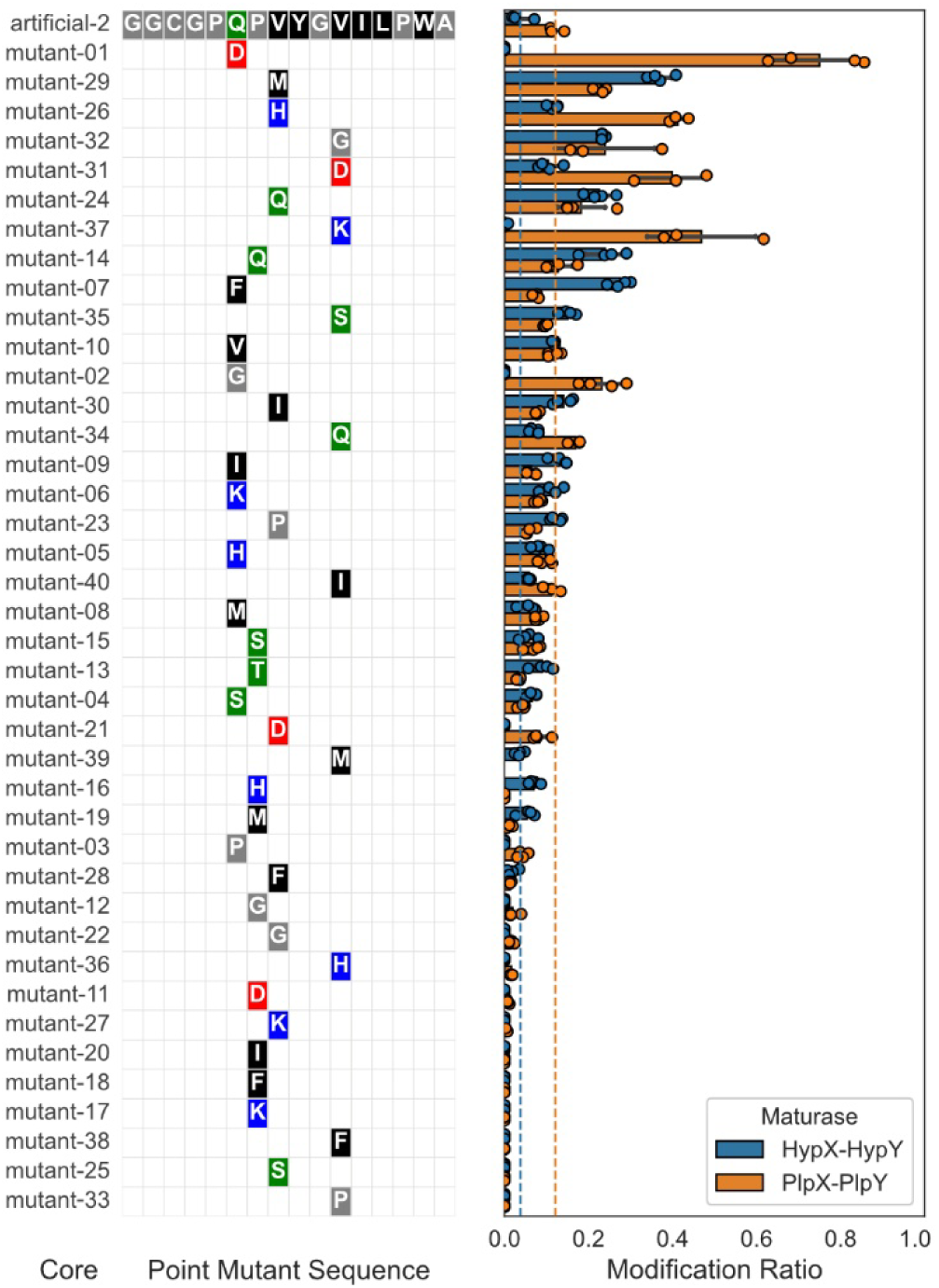
Modification of various point mutations of the “Artificial-02” peptide by the spliceases HypX-HypY and PlpX-PlpY, showing that most mutations are well-tolerated. The amino acid alignment only shows sequence changes from the reference sequence Artificial-02, and the bar plot shows the ratio of spliced peptide over total peptide as measured by LC-MS. The dashed lines represent the modification ratio of the Artificial-02 sequence used as a baseline. Bars show the mean ± SD of n = 3-4 biological replicates; points are individual biological replicates.

### Using *in-silico* modeling to probe the substrate scope of PlpX

Therapeutic and biotechnological applications of spliceotides ideally require that spliceases can modify diverse sets of target precursor peptides. The quality of unbiased spliceotide libraries is dictated by the overall substrate scope of the splicease, as well as the efficiency with which the splicease modifies individual variants, both of which dictate the accessible chemical space. Combinations of several changes in a substrate sequence compared to the wild type increase the chance of loss of modification. Targeted libraries designed to have a high proportion of modified peptides can narrow the search space in a screen and make hits more likely.^33^ We therefore sought a reliable and efficient method for producing such diverse, multipoint mutants that would allow rational, systematic identification of the best candidates without relying on exhaustive screening.

We used AI-based structure prediction together with Rosetta energy calculations^34,35^ to model all multipoint mutant combinations at positions −4 to −1, +1, and +2. Structures of PlpX, PlpY, and the leader-fused Artificial-2 core were first modeled using AlphaFold 3.^36^ The models correctly positioned the YG motif near the homolytic cleavage site of the SAM cofactor. The modeled structures were subsequently used as the basis for modeling all multipoint peptide mutants (−4 to −1 and +1 to +2) (Figure 4A). Initial results showed that the energy calculations were biased toward the negatively charged residues Glu and Asp at position −2, consistent with its predicted proximity to one of the three iron-sulfur clusters. Close inspection of the modeled structures, however, showed that these residues do not fit sterically (Figure 4B). The amino acid choice at this position was therefore restricted to Pro (as in Artificial-2), and the sterically small residues Ala, and Gly, resulting in 9.6 million peptide variants to model. Following an initial, fast round of modeling, we selected the top 200,000 peptides by energy ranking for more detailed modeling (see Methods). The top-ranked designs all harbored Glu (and to a lesser extent Asp and Gln) at position −4, again near the same iron-sulfur cluster, possibly due to Rosetta’s preference for electrostatic interactions. To mitigate the risk of oversampling Glu at this position, we selected 5 designs with Glu at position −4, and the remaining 39 from the set of designs lacking Glu, Asp, and Gln at this position.

**Figure 4.**
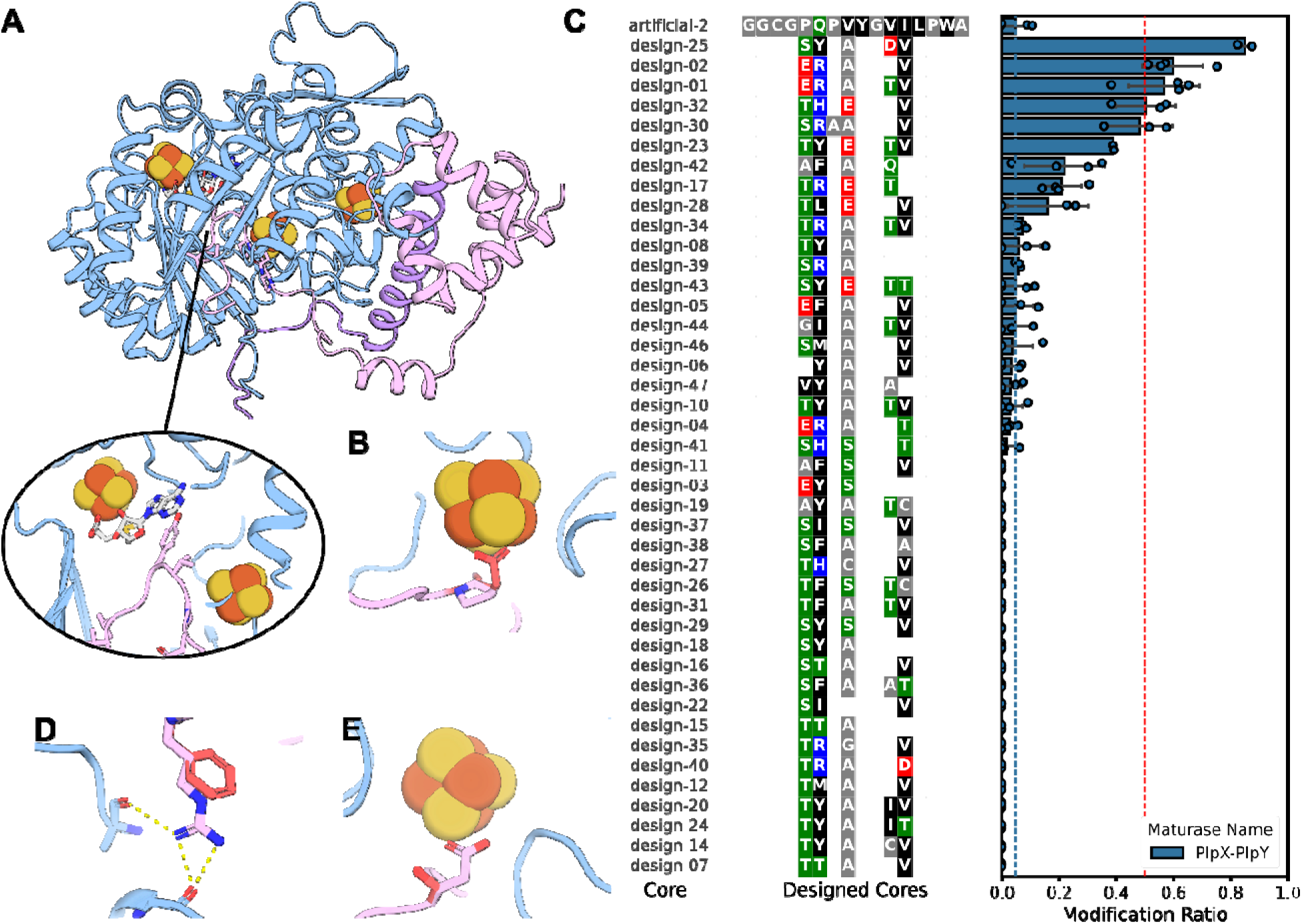
Modeling and validation of Rosetta-selected spliceotide-substrate peptides. **(A)** AF3 model of the Plp-X spliceotide (blue) bound to the Plp-Y peptide (purple), with the Artificial-2 reference peptide (pink), three iron-sulfur clusters (yellow/orange spheres), and SAM cofactor (white sticks). *Inset*: details of the YG motif. **(B)** Position −2 constraint: Glu (red) sterically clashes with an adjacent iron-sulfur cluster, whereas Pro (shown in Artificial-2) accommodates the cluster without conflict. **(C)** Modification ratios for all 47 Rosetta-designed variants, ranked by efficiency. Blue dotted line marks the Artificial-2 reference modification efficiency; red dotted line indicates the 50% efficiency threshold. Bars show the mean ± SD of n = *3-4* biological replicates; points are individual biological replicates. **(D)** Position −3 mechanistic comparison: Arg forms hydrogen bonds with backbone carbonyls (dashed yellow interactions), stabilizing peptide binding, whereas Phe (red) leaves a void without compensatory interactions. **(E)** Position −1 critical interaction: Glu forms favorable electrostatic contacts with the iron-sulfur cluster; Ser (red) disrupts this interaction, with the polar hydroxyl group oriented away from the cluster.

The genes encoding the 47 Rosetta-selected peptides were cloned into the previously used precursor plasmid using standard blunt-end cloning methods. Following expression, purification, and analysis using the plate-based assay described above, replicate datapoints were gathered and summary statistics generated (Figure 4C, Figures S134-177). The single-point spliceotide mutants from the previous experiment yielded only one highly efficiently modified peptide (modification efficiency > 0.5, success rate 2.3%), whereas the multipoint mutants from this Rosetta-selected set showed six highly efficient variants (13.6%).

We analyzed pairs of peptides that differed by a single mutation yet had drastically altered modification efficiencies. Peptide design-02 (modification efficiency 0.58) and peptide design-05 differ by an Arg → Phe substitution at position −3, yet design-05 is not modified at all. Modeling indicates that Arg forms hydrogen bonds with backbone carbonyls in the spliceotide active site, whereas Phe leaves a large void. Consistently, other designs with Phe at position −3 show a modification efficiency averaging 0.04.

Designs d-13 and d-29 differ by a Glu → Ser in position −1. While Glu forms favorable electrostatic interactions with the iron-sulfur cluster, Ser introduces an unsatisfied polar group. Accordingly, peptides bearing Ser at position −1 show modification efficiencies < 0.003, while designs with Glu average 0.42. Similarly, peptide d-32 and d-27 differ by Glu → Cys in the same position, and the modification efficiency drops from 0.586 to zero.

Although 80 designs is a limited sample from which to draw definitive conclusions about spliceotide processivity, we observed an enrichment of highly modified peptides (more than 50% modification ratio) in the Rosetta-selected set (13.6% vs. 2.3% in single-point mutants). This suggests that the energy-ranked population contains peptides compatible for spliceotide modification and capable of targeting diverse protease active-site environments. Energy-ranked selection may therefore enrich for modifiable multipoint variants, although the difference did not reach significance in this pilot set (Fisher’s exact test, one-sided p = 0.055, n=47 per group) and warrants validation in a larger sample. Future efforts can build on this foundation to expand design spaces and screen candidates against a panel of structurally diverse proteases, accelerating the discovery of spliceotide-based inhibitors targeting multiple therapeutic targets. Approaches to gather substrate scope data that are not dependent on LC-MS readout would greatly accelerate data generation, and larger datasets would enable a combination of Rosetta-based scoring and machine-learning based approaches to accurately map the chemical space of various spliceases.

## CONCLUSION

We identified HypX as a new splicease and characterized its substrate range alongside that of the previously reported PlpX, showing that the sequence-related spliceases diverge substantially in substrate tolerance. Contrary to our expectations, HypX was not as promiscuous as PlpX, although the native *hyp* locus encodes more varied and numerous precursors. Building on this result, we evaluated whether Rosetta-based computational ranking could identify precursor variants carrying many mutations relative to the parent sequence that remain good splicease substrates. Compared to single random point mutations, the Rosetta-selected set showed more highly modified precursors, and the structural models provide a rationale for why individual substitutions improve acceptance (Arg→Phe at −3, Glu→Ser/Cys at −1). This study is thus a first proof-of-concept that physics-based modeling can prioritize multipoint variants from a massive chemical space, enriching for functional substrates without exhaustive screening.

The key bottleneck for future follow-up work is data generation, as the current LC-MS readout limits us to dozens of variants per experiment. Higher-throughput techniques such as FACS readout of a spliced phenotype would allow for the screening of up to millions of variants. Coupling such high-throughput data with physics-based energy ranking from AlphaFold3 and Rosetta would test computational predictions at scale and enable machine-learning approaches to uncover non-obvious position-specific. In turn, different splicease families could be compared and contrasted. Together with improved *in silico* modeling, this would support the design of more targeted ketoamide-peptide libraries and, ultimately, establish spliceotides as a generalizable platform for generating diverse protease-inhibitor libraries, with the same workflow applied to new spliceases and targets.

## METHODS AND MATERIALS

### Strains and Materials

Primers for this study were synthesized by Microsynth (Balgach, Switzerland), DNA fragments were ordered from TWIST Bioscience (San Francisco, USA). Q5 DNA polymerase was obtained from New England Biolabs (Ipswich, USA). Amplified DNA was purified using NucleoSpin Gel and PCR Clean-up kits from Macherey-Nagel (Germany), whereas plasmids were isolated using NucleoSpin kits also purchased from Macherey-Nagel. Sanger sequencing and Nanopore sequencing was performed by Microsynth.

Terrific Broth (TB: yeast extract 24 g/L, tryptone 20 g/L, 17 mM KH_2_PO_4_, and 72 mM K_2_HPO_4_) supplemented with 1.2% glycerol) was prepared by the institute staff and supplemented with the appropriate antibiotics before use.

### Small scale shake flask expressions

Cultures of *E. coli* BL21(DE3) tuner containing the corresponding splicease and precursor peptide plasmids were grown overnight in cell culture tubes (Greiner BioOne) containing LB and the appropriate antibiotics. The next day 30 mL of TB medium containing 100 µg/mL Kanamycin and 50 µg/mL of spectinomycin inside 100 mL unbaffled shake flasks were seeded with a 1/100 inoculum of the overnight culture and incubated at 37 °C with 180 rpm orbital shaking until an optical density (OD_600_) between 1.0 and 2.0 was reached. Upon reaching the target OD_600_, expression was initiated by the addition of 1 mM isopropyl β-D-1-thiogalactopyranoside (IPTG), upon which the cultures were transferred to an Infors HT shaker with 180 rpm orbital shaking at either 16, 24, or 30 °C for approximately 22 hours.

### Procedure for purification from shake flasks

The cells from expression cultures were pelleted and then frozen in 50 mL Falcon tubes at −20 °C until use. The cell pellets were thawed and resuspended in 3 mL of Lysis Buffer A (0.3 M NaCl, 50 mM Tris-HCl, 10 mM imidazole, 0.04 mg/mL DNase, 0.5 mg/mL lysozyme, 1X Promega FastBreak, pH 8.0) and incubated at 37 °C for 1 hour to allow for cell lysis and DNA cleavage by the DNase I. To this mixture was added 9 mL of Lysis Buffer B (6.7 M Guanidinium Hydrochloride, 0.3 M NaCl, 50 mM Tris-HCl, 10 mM imidazole, pH 8.0) and the Falcon tube was thoroughly mixed, after which the lysate was centrifuged at 11,000 × g for 30 minutes at 10 °C in a tabletop centrifuge (Eppendorf) to separate the soluble fraction from the pellet. Columns containing 0.2 mL of Ni-NTA resin volume (Protino Ni-NTA, Macherey-Nagel, Germany) were pre-equilibrated with 4 mL of Wash Buffer 1 (5 M guanidinium hydrochloride, 0.3 M NaCl, 50 mM Tris-HCl, 10 mM imidazole, pH 8.0). The soluble fraction from the lysate was then added to the gravity-flow columns, which were capped and rotated at 20 rpm at 4 °C for one hour to allow for binding to the resin. The flowthrough was then collected and one wash with 5 mL of Wash Buffer 1 performed, followed by a wash with 5 mL of Wash Buffer 2 (1 M guanidinium hydrochloride, 0.3 M NaCl, 50 mM Tris-HCl, 20 mM imidazole, pH 8.0), and finally the resin was washed with 5 mL of Wash Buffer 3 (0.3 M NaCl, 50 mM Tris-HCl, 20 mM imidazole, pH 8.0). The bead-bound peptides were eluted thrice using 1 mL of Elution Buffer (0.3 M NaCl, 50 mM Tris-HCl, 250 mM imidazole, pH 8.0) and the resulting eluate stored for further processing.

### Procedure for 96 well plate expressions

Cultures of *E. coli* BL21(DE3) tuner containing the corresponding splicease and precursor peptide plasmids were grown overnight in flat-bottom 96 well plates (TPP, Switzerland) containing 200 µL of LB medium plus 50 µg/mL Kanamycin and 50 µg/mL Spectinomycin at 37 °C. From each well, 10 µL were used to inoculate square-welled deep 96 well plates (Brand) containing TB medium with 100 µg/mL Kanamycin and 50 µg/mL Spectinomycin, upon which the plates were sealed (BreathSeal membranes) and incubated at 37 °C with 900 rpm shaking for 3.5 hours. After this, 0.5 mL of TB medium (3 mM IPTG, 100 µg/mL Kanamycin and 50 ug/mL Spectinomycin) were added to each well for a final IPTG concentration of 1 mM. The plates were again covered with membranes (BreathSeal) and incubated at 24 °C with 900 rpm shaking for 22 hours on an Eppendorf plate shaker. Cells were harvested by centrifugation at 3,200 × g for 30 minutes at 4 °C, and the 96 well plates containing the cell pellet were frozen at −20 °C until use.

### Concentration and cleavage of precursors

Eluates from expressions in shake flasks were concentrated in spin concentrators (Sartorius) at 7,000 × g of rotation and buffer exchanged trypsin buffer (50 mM Tris-HCl, 1 mM CaCl_2_, pH=7.6). Trypsin digestions were performed in a total volume of 50 µL inside 1X trypsin buffer with 5 mM tris(2-carboxyethyl)phosphine (TCEP) and trypsin (sequencing grade modified trypsin, Promega, USA) was added to a final quantity of 0.1 µg. Trypsin digestions were performed overnight at 37 °C, after which the digestion reactions were centrifuged to remove any particles.

### Procedure for 96 well plate purification

From duplicate deep 96 well plates containing the frozen cell pellets from the expression cultures, the plates were thawed and cell pellets were each thoroughly resuspended in 300 µL of Lysis Buffer A (0.3 M NaCl, 50 mM Tris-HCl, 10 mM imidazole, 0.04 mg/mL DNase, 0.5 mg/mL lysozyme, 1X Promega FastBreak, pH 8.0) using a multichannel pipette and shaken at room temperature for 30 minutes on a plate shaker at 900 rpm. To each well was added 900 µL of Lysis Buffer B (6.7 M guanidinium hydrochloride, 0.3 M NaCl, 50 mM Tris-HCl, 10 mM imidazole, pH 8.0) and thoroughly mixed, upon which the plates were placed into an Eppendorf tabletop centrifuge using a swing bucket rotor and centrifuged at 3,200 × g for 90 minutes at 10 °C. After centrifuging, the first 1000 µL of the supernatant were carefully aspirated and placed into new deep 96 well plates. Protino 96 well plates were loaded with 100 uL of PureCube Co-NTA beads (CubeBiotech, Germany) and pre-equilibrated with 0.5 mL of Wash Buffer 1 (5 M guanidinium hydrochloride, 0.3 M NaCl, 50 mM Tris-HCl, 10 mM imidazole, pH 8.0). The first 0.5 mL of supernatant from the cell lysate was added to the purification plates, the plates were centrifuged at 800 × g for 1 minute at 10 °C and the flowthrough collected in deep 96 well plates (Thermo Fisher), after which the flowthrough was loaded once more onto the columns and re-centrifuged. The same was then done for the remaining 0.5 mL of supernatant from the cell lysate to ensure proper loading of the Cobalt beads. The washing steps were then performed by adding 0.5 mL of Wash Buffer 1, 0.5 mL of Wash Buffer 2 (1 M guanidinium hydrochloride, 0.3 M NaCl, 50 mM Tris-HCl, 20 mM imidazole, pH 8.0), and 0,5 mL of Wash Buffer 3 (0.3 M NaCl, 50 mM Tris-HCl, 20 mM imidazole, pH 8.0) which were all performed by centrifuging the plates with the collection plates at 800 × g for 1 minute at 10 °C. The bead-bound peptides were eluted thrice using 0.2 mL of Elution Buffer (0.3 M NaCl, 50 mM Tris-HCl, 250 mM imidazole, pH 8.0) into Nunc 96 well plates (Thermo Fisher) and frozen until use.

### Procedure for 96 well plate TEV digestion

The frozen 96 well plates containing the first and second elutions of the purified protein samples were left to thaw at room temperature. For every sample, 50 µL of the first and second elution were added into 96 well plates (nunc round-bottom, ThermoFisher USA) containing 100 µL per well of cleavage mixture (2X TEV Buffer, 10 mM TCEP, 2.5 units of ProTEVPlus protease, Promega, USA), the plates were then sealed and incubated overnight at 30 °C.

### LC-MS analysis of digested peptides

Liquid chromatography–mass spectrometry (LC-MS) analysis was conducted on a UHPLC system coupled to a Q Exactive Orbitrap mass spectrometer (Thermo Fisher Scientific) equipped with an electrospray ionization (ESI) source. Mobile phase A consisted of LC-MS-grade water containing 0.1% formic acid (v/v), and mobile phase B consisted of acetonitrile (ACN) containing 0.1% formic acid (v/v). Chromatographic separations were evaluated using two distinct gradient elution profiles at a constant flow rate of 0.5 mL/min and column temperature of 50 °C.

*Method A* comprised a 20-min total run time, starting with an initial 2-min isocratic hold at 5% B, followed by a linear gradient increase to 95% B over 12 min (2.0 to 14.0 min). The column was washed at 95% B for 3 min (14.0 to 17.0 min), returned to initial conditions (5% B) at 17.0 min, and re-equilibrated through 20.0 min.

*Method B* comprised a 10-min fast gradient starting with an initial 3-min isocratic hold at 5% B, followed by a rapid step to 20% B over 0.2 min (3.0 to 3.2 min). Mobile phase B was then linearly increased to 98% B over 4.4 min (3.2 to 7.6 min) and held at 98% B for column washing until 9.0 min. The system was returned to 5% B at 9.05 min and re-equilibrated through 10.0 min.

### Mass Spectrometry Acquisition Parameters

Data acquisition on the Q Exactive Orbitrap MS was operated in positive ESI mode utilizing Full MS / Data-Dependent MS^2^ (top-N dd-MS^2^) workflows. For both methods, full MS scans were collected across an *m/z* range of 400–2000 at a resolving power of 70,000 (at *m/z* 200), an Automatic Gain Control (AGC) target of 3 × 10^6^ and a maximum injection time (IT) of 100 ms. Tandem mass spectra (MS^2^) were acquired at a resolution of 17,500, an AGC target of 1 × 10^5^, a maximum IT of 50 ms, and a precursor isolation window of 4.0 *m/z*.

### Automated data analysis

The output files in ThermoFisher .raw format were converted to mzML files using the desktop tool MSConvertGUI3 and saved as described in the supplementary Excel files. A series of custom python scripts were used to automatically search for peptide fragments, hits were then manually validated and processed to extract the peak areas. The full code is freely available on GitHub under <u>jordant/LCMS_RiPP</u>.

### Structure prediction and Rosetta modeling

The structure of the spliceotide complex (Plp-X, Plp-Y, and Artificial-2) was predicted using AF3 (v3.0.1) with sequences in JSON format, specifying ligands as CCD codes (SAM, SF4). Four predictions per structure (seeds: 1, 2, 3, 42) used default parameters on 8 GPUs (25 GB each) in a Singularity container. All models achieved ranking scores > 0.94 with an average pairwise RMSD of 0.21 Å, differing primarily in YG motif backbone conformation (three distinct poses). Model ranks 0, 1, and 2 were refined 10 times each,^37^ representing each YG conformation. Rank 1’s lowest-energy structure was selected as the template for mutant design: its YG Tyr had the lowest Rosetta energy among the three (ΔΔG = −5.04 REU vs. rank 0, −2.6 REU vs. rank 2).

We modeled 9.6 million peptide variants covering all 20 amino acids at positions −4, −3, −1, +1, and +2, with position −2 restricted to Ala, Gly, and Pro. A short refinement protocol screened these to the top 200,000 by Rosetta total_score, which were then subjected to detailed refinement (see Supplementary Files). We selected 47 designs for experimental validation: five containing Glu at position −4, and 39 lacking Glu, Asp, and Gln at this position

## Supporting information

SI_report_plates

Main Supplementary Infromation File

## ASSOCIATED CONTENT

### Data availability statement

The python code for analyzing the LC-MS data is available on GitHub under the link https://github.com/thomashjordan/LCMS_RiPP, Rosetta files are also present in the same repository. Plasmids and strains are available from the authors upon reasonable request.

## AUTHOR INFORMATION

### Author Contributions

Thomas H. Jordan: conceptualization, methodology, investigation, formal analysis, data curation, visualization, writing, editing

Rosalie Lipsh-Sokolik: *in silico* methodology, data curation, visualization, writing, editing Jörn Piel: conceptualization, methodology, funding acquisition, writing, editing

## ACKNOWLEDGEMENTS

This work was financially supported by ETH Zurich (grant no. ETH-21 21-2) and the Monique Dornonville de la Cour Foundation (2026-HS-396). We thank Sarolt Magyari and Daniel Richter for sharing their experience with expressing spliceotides. We would also like to thank Alexander Brachmann and Anna Vagstad for discussion and advice.

