## Supplementary material for "Mapping and Probing the Substrate Scope of a Ketoamide-Forming Splicease by Computational Design": SI_report_plates

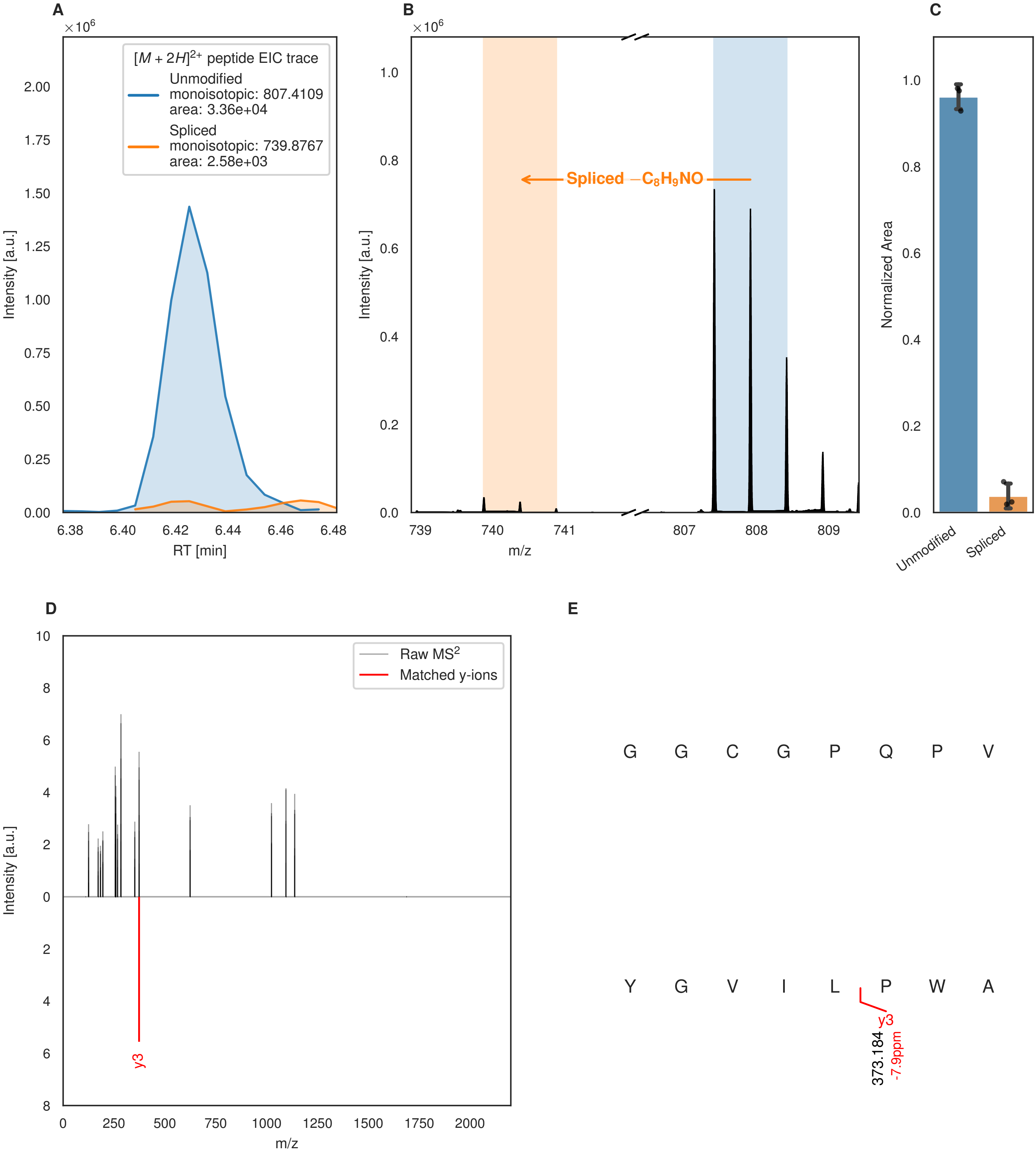
A
$[M + 2H]^{2+}$ peptide EIC trace
Unmodified
monoisotopic: 807.4109
area: 3.36e+04
Spliced
monoisotopic: 739.8767
area: 2.58e+03
Intensity [a.u.]
RT [min]
B
Spliced $-C_8H_9NO$
Intensity [a.u.]
m/z
C
Normalized Area
Unmodified
Spliced
D
Raw MS$^2$
Matched y-ions
y3
Intensity [a.u.]
m/z
E
G G C G P Q P V
Y G V I L P W A
y3
373.184
-7.9ppm

Figure S53 (figure continued from the previous page).

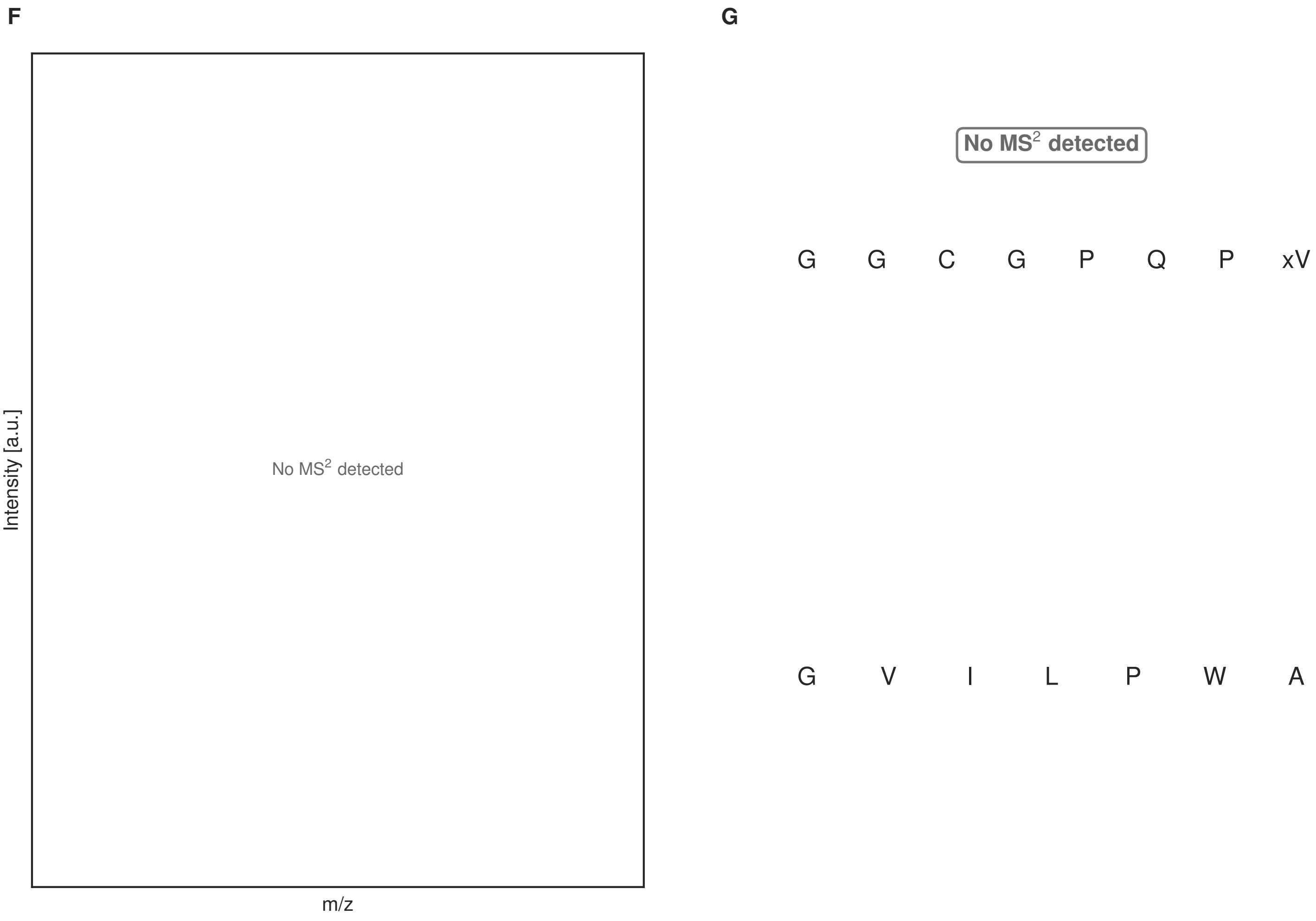


**Figure S53: Modification of the TJ-C-706 precursor peptide by HypXY.** LC-MS analysis of IMAC-purified precursor peptides from *E. coli* BL21(DE3)Tuner containing splicease-encoding plasmid TJ-C-689 (HypXY) and precursor peptide-encoding plasmid TJ-C-706 (RSTN1-HypAc-TEV-Artificial-2) after expression at 24 °C for 22 h in 1.5 mL cultures after digestion with TEV. Data from n=3 independent biological replicates. Spliced amino acids have an x as a prefix. Spliced refers to a single tyramine excision. A representative experimental replicate is shown in panels **A, B.**

**A** Close-up plot of the representative extracted ion chromatogram (EIC) of the various peptide states; the blue trace corresponds to the Unmodified peptide; the orange trace corresponds to the Spliced peptide. **B** Close-up plot of the m/z values of the peptides detected by the mass spectrometer, with the shaded colored areas corresponding to where the expected m/z values for the peptide m/z traces should occur, with the observed spectrum plotted in black. The Unmodified, Spliced peptide(s) were found in the mass spectrum. **C** Bar plot showing the relative abundance of the various peptide states at charge z=2; the bars represent the average relative abundances normalized to total peak areas, with each individual experimental replicate value plotted as a black dot. **D** Mirrored MS$^2$ plot for the Unmodified peptide. In this mirror plot, the full fragmentation spectrum is plotted in black pointing upward, and the matched fragment b ions (blue) and y ions (red) are plotted pointing downward, mirrored against it, so that each matched fragment can be compared directly with the measured spectrum. **E** MS$^2$ spectrum of the fragments corresponding to the Unmodified peptide sequence, with matched b (blue, above) and y ions (red, below) plotted. **F** Mirrored MS$^2$ plot for the Spliced peptide. In this mirror plot, the full fragmentation spectrum is plotted in black pointing upward, and the matched fragment b ions (blue) and y ions (red) are plotted pointing downward, mirrored against it, so that each matched fragment can be compared directly with the measured spectrum. **G** MS$^2$ spectrum of the fragments corresponding to the Spliced peptide sequence, with matched b (blue, above) and y ions (red, below) plotted.

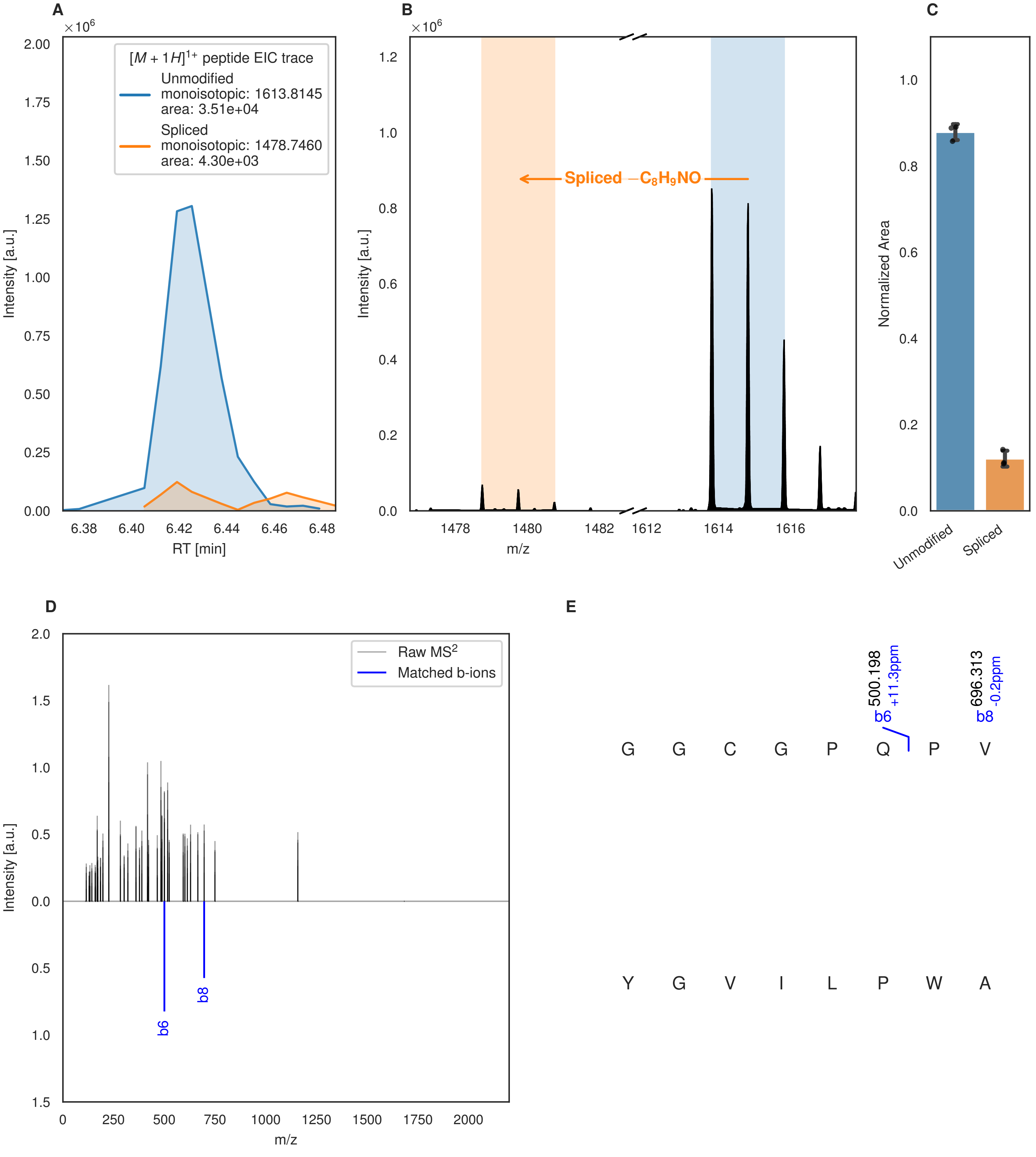

A
$\times 10^6$
$[M+1H]^{1+}$ peptide EIC trace
Unmodified
monoisotopic: 1613.8145
area: 3.51e+04
Spliced
monoisotopic: 1478.7460
area: 4.30e+03
Intensity [a.u.]
RT [min]
B
$\times 10^6$
Spliced $-C_8H_9NO$
Intensity [a.u.]
m/z
C
Normalized Area
Unmodified
Spliced
D
Raw MS$^2$
Matched b-ions
Intensity [a.u.]
m/z
b6
b8
E
500.198
+11.3ppm
b6
696.313
-0.2ppm
b8
G G C G P Q P V
Y G V I L P W A

Figure S54 (figure continued from the previous page).

F

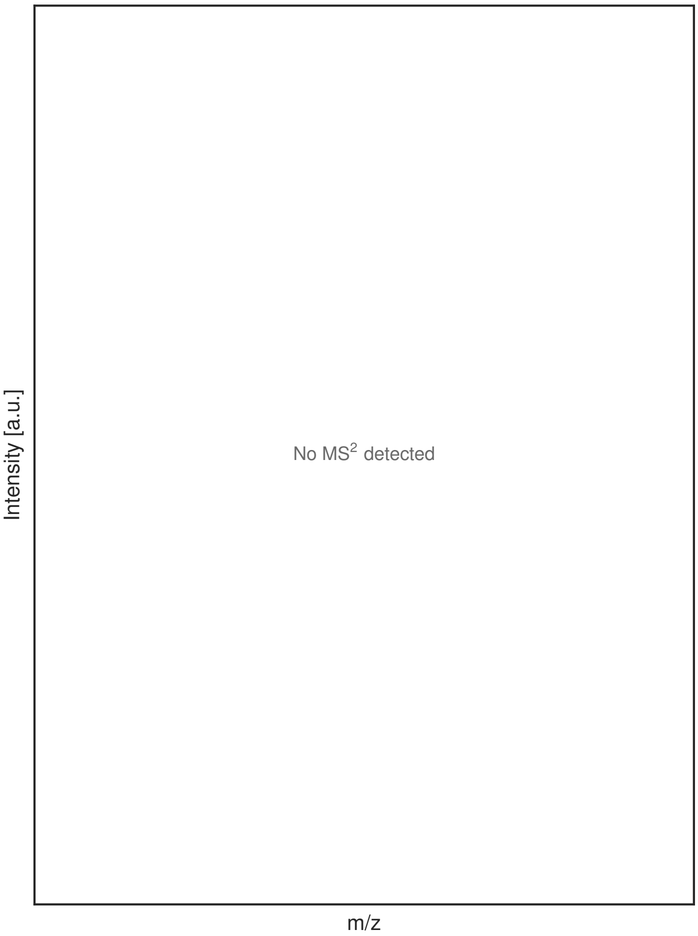


G

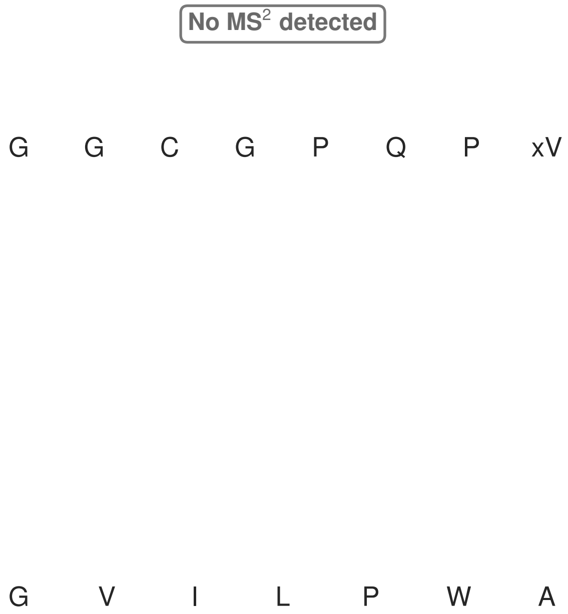


**Figure S54: Modification of the TJ-C-710 precursor peptide by PlpXY.** LC-MS analysis of IMAC-purified precursor peptides from *E. coli* BL21(DE3)Tuner containing splicease-encoding plasmid TJ-C-599 (PlpXY) and precursor peptide-encoding plasmid TJ-C-710 (RSTN1-PlpA3-TEV-Artificial-2) after expression at 24 °C for 22 h in 1.5 mL cultures after digestion with TEV. Data from n=3 independent biological replicates. Spliced amino acids have an x as a prefix. Spliced refers to a single tyramine excision. A representative experimental replicate is shown in panels **A, B.**

**A** Close-up plot of the representative extracted ion chromatogram (EIC) of the various peptide states; the blue trace corresponds to the Unmodified peptide; the orange trace corresponds to the Spliced peptide. **B** Close-up plot of the m/z values of the peptides detected by the mass spectrometer, with the shaded colored areas corresponding to where the expected m/z values for the peptide m/z traces should occur, with the observed spectrum plotted in black. The Unmodified, Spliced peptide(s) were found in the mass spectrum. **C** Bar plot showing the relative abundance of the various peptide states at charge z=1; the bars represent the average relative abundances normalized to total peak areas, with each individual experimental replicate value plotted as a black dot. **D** Mirrored MS$^2$ plot for the Unmodified peptide. In this mirror plot, the full fragmentation spectrum is plotted in black pointing upward, and the matched fragment b ions (blue) and y ions (red) are plotted pointing downward, mirrored against it, so that each matched fragment can be compared directly with the measured spectrum. **E** MS$^2$ spectrum of the fragments corresponding to the Unmodified peptide sequence, with matched b (blue, above) and y ions (red, below) plotted. **F** Mirrored MS$^2$ plot for the Spliced peptide. In this mirror plot, the full fragmentation spectrum is plotted in black pointing upward, and the matched fragment b ions (blue) and y ions (red) are plotted pointing downward, mirrored against it, so that each matched fragment can be compared directly with the measured spectrum. **G** MS$^2$ spectrum of the fragments corresponding to the Spliced peptide sequence, with matched b (blue, above) and y ions (red, below) plotted.

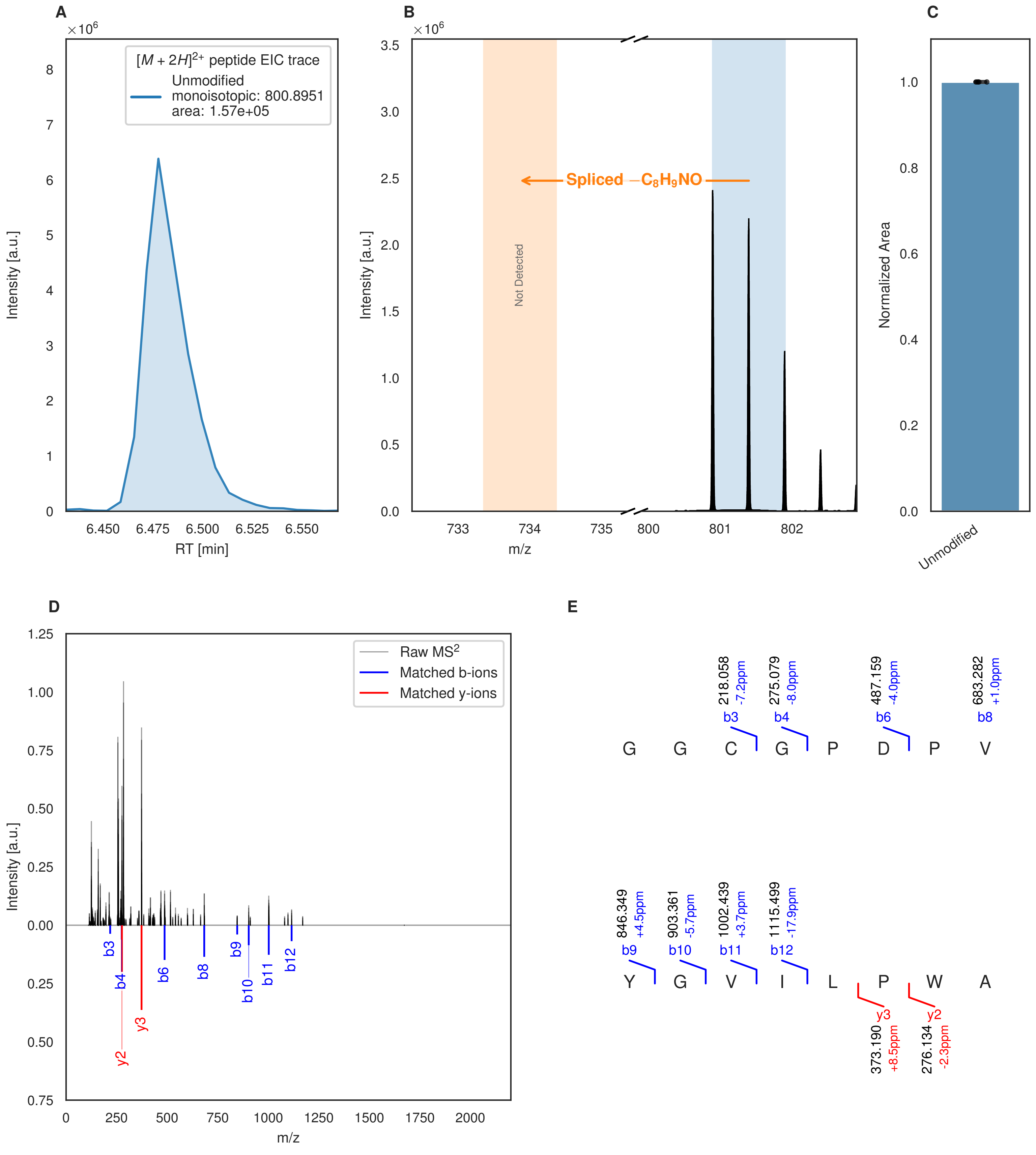


**Figure S55: Modification of the mutant_1 precursor peptide by HypXY.** LC-MS analysis of IMAC-purified precursor peptides from *E. coli* BL21(DE3)Tuner containing splicease-encoding plasmid TJ-C-689 (HypXY) and precursor peptide-encoding plasmid mutant_1 (RSTN1-HypAc-TEV-mutant-1) after expression at 24 °C for 22 h in 1.5 mL cultures after digestion with TEV. Data from n=4 independent biological replicates. Spliced amino acids have an x as a prefix. Spliced refers to a single tyramine excision. A representative experimental replicate is shown in panels **A, B.**

**A** Close-up plot of the representative extracted ion chromatogram (EIC) of the various peptide states; the blue trace corresponds to the Unmodified peptide. **B** Close-up plot of the m/z values of the peptides detected by the mass spectrometer, with the shaded colored areas corresponding to where the expected m/z values for the peptide m/z

(Caption continued on the next page.)

traces should occur, with the observed spectrum plotted in black. The Unmodified peptide(s) were found in the mass spectrum and the Spliced peptide(s) were not detected. **C** Bar plot showing the relative abundance of the various peptide states at charge z=2; the bars represent the average relative abundances normalized to total peak areas, with each individual experimental replicate value plotted as a black dot. **D** Mirrored $MS^2$ plot for the Unmodified peptide. In this mirror plot, the full fragmentation spectrum is plotted in black pointing upward, and the matched fragment b ions (blue) and y ions (red) are plotted pointing downward, mirrored against it, so that each matched fragment can be compared directly with the measured spectrum. **E** $MS^2$ spectrum of the fragments corresponding to the Unmodified peptide sequence, with matched b (blue, above) and y ions (red, below) plotted.

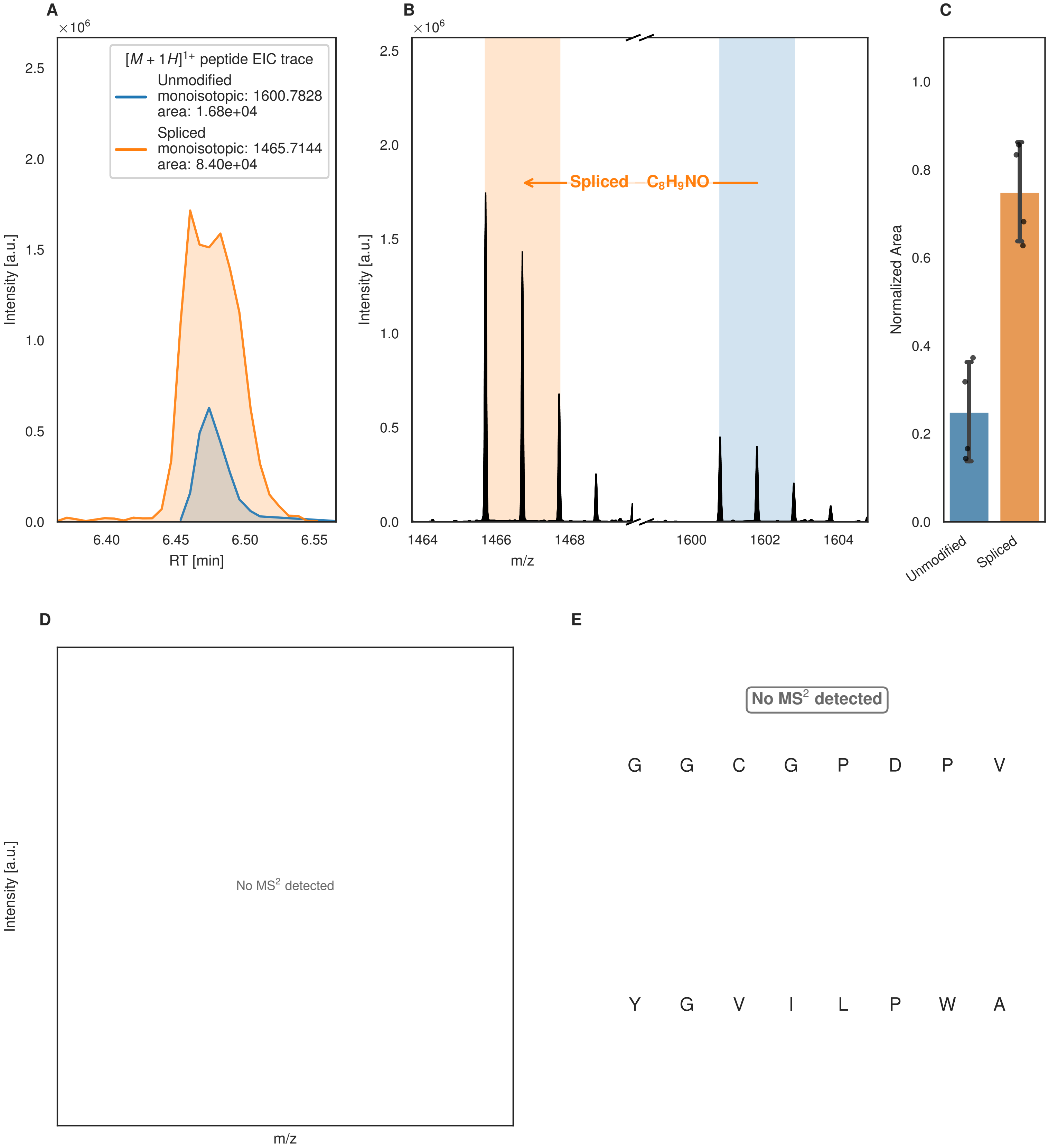
A
×10^6
[M + 1H]^1+ peptide EIC trace
Unmodified
monoisotopic: 1600.7828
area: 1.68e+04
Spliced
monoisotopic: 1465.7144
area: 8.40e+04
Intensity [a.u.]
RT [min]
B
Spliced −C8H9NO
m/z
C
Normalized Area
Unmodified
Spliced
D
No MS2 detected
E
No MS2 detected
G G C G P D P V
Y G V I L P W A

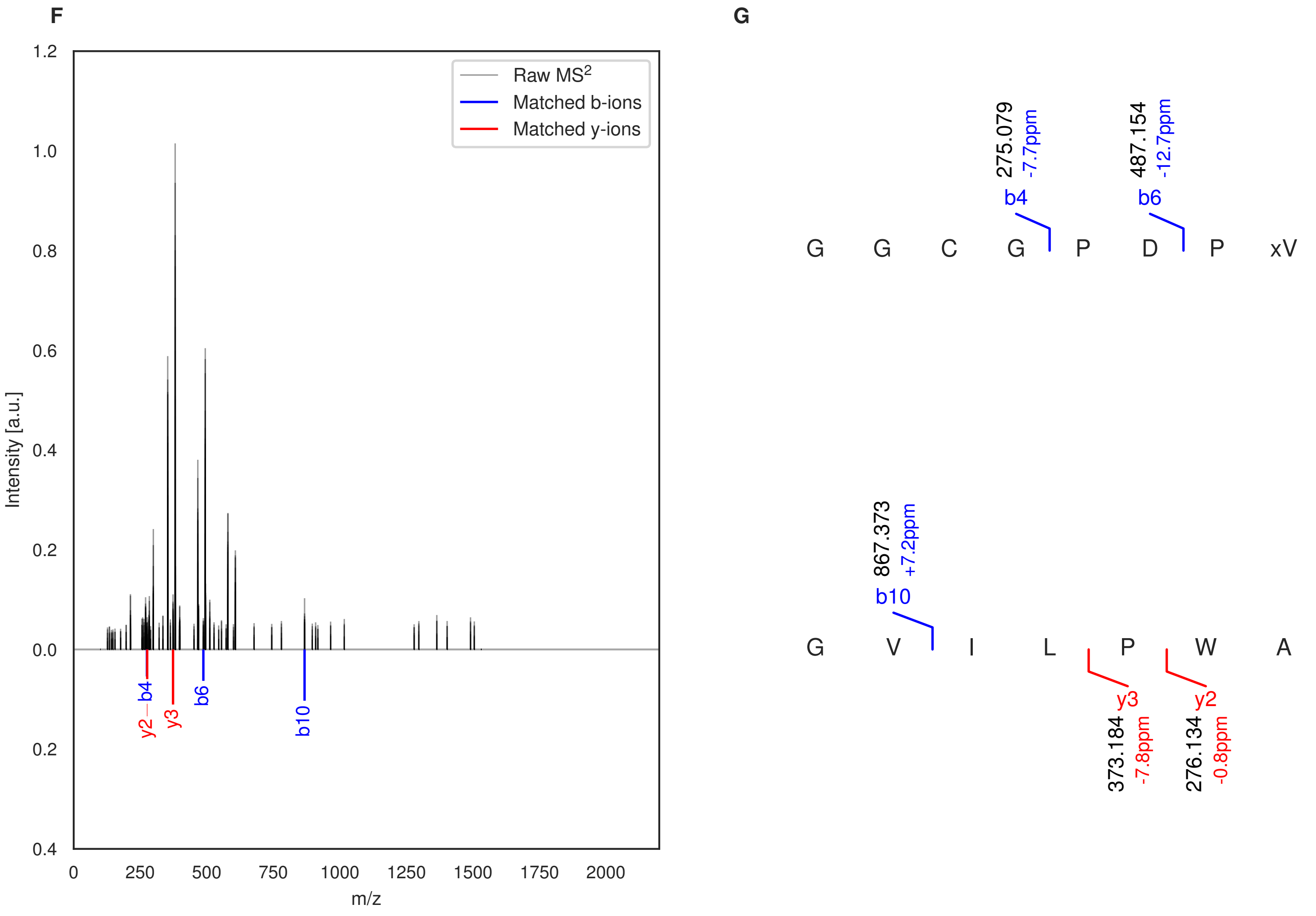


**Figure S56: Modification of the mutant_1 precursor peptide by PlpXY.** LC-MS analysis of IMAC-purified precursor peptides from *E. coli* BL21(DE3)Tuner containing splicease-encoding plasmid TJ-C-559 (PlpXY) and precursor peptide-encoding plasmid mutant_1 (RSTN1-PlpA3-TEV-mutant-1) after expression at 24 °C for 22 h in 1.5 mL cultures after digestion with TEV. Data from n=4 independent biological replicates. Spliced amino acids have an x as a prefix. Spliced refers to a single tyramine excision. A representative experimental replicate is shown in panels **A, B.**

**A** Close-up plot of the representative extracted ion chromatogram (EIC) of the various peptide states; the blue trace corresponds to the Unmodified peptide; the orange trace corresponds to the Spliced peptide. **B** Close-up plot of the m/z values of the peptides detected by the mass spectrometer, with the shaded colored areas corresponding to where the expected m/z values for the peptide m/z traces should occur, with the observed spectrum plotted in black. The Unmodified, Spliced peptide(s) were found in the mass spectrum. **C** Bar plot showing the relative abundance of the various peptide states at charge z=1; the bars represent the average relative abundances normalized to total peak areas, with each individual experimental replicate value plotted as a black dot. **D** Mirrored $MS^2$ plot for the Unmodified peptide. In this mirror plot, the full fragmentation spectrum is plotted in black pointing upward, and the matched fragment b ions (blue) and y ions (red) are plotted pointing downward, mirrored against it, so that each matched fragment can be compared directly with the measured spectrum. **E** $MS^2$ spectrum of the fragments corresponding to the Unmodified peptide sequence, with matched b (blue, above) and y ions (red, below) plotted. **F** Mirrored $MS^2$ plot for the Spliced peptide. In this mirror plot, the full fragmentation spectrum is plotted in black pointing upward, and the matched fragment b ions (blue) and y ions (red) are plotted pointing downward, mirrored against it, so that each matched fragment can be compared directly with the measured spectrum. **G** $MS^2$ spectrum of the fragments corresponding to the Spliced peptide sequence, with matched b (blue, above) and y ions (red, below) plotted.

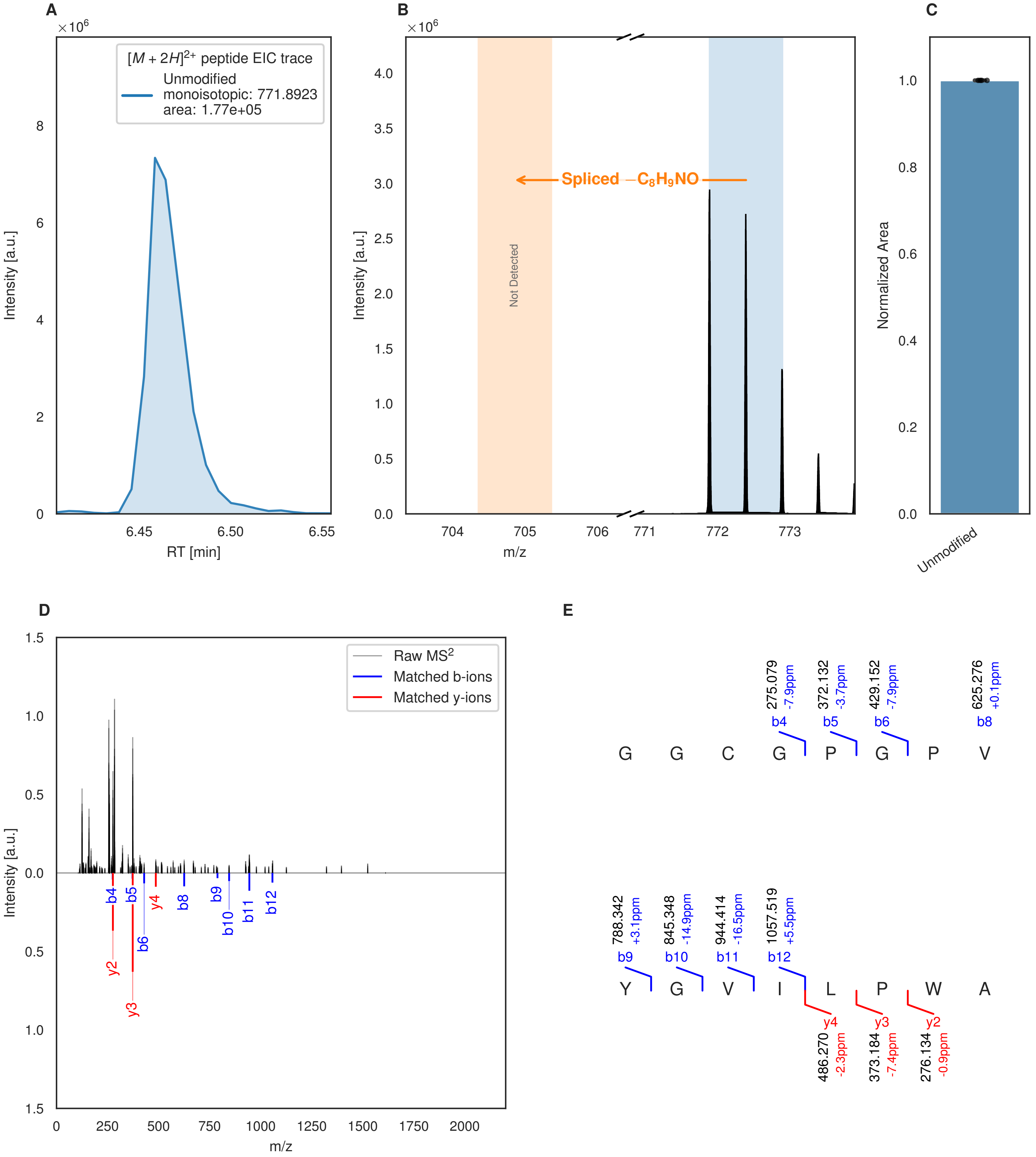


**Figure S57: Modification of the mutant_2 precursor peptide by HypXY.** LC-MS analysis of IMAC-purified precursor peptides from *E. coli* BL21(DE3)Tuner containing splicease-encoding plasmid TJ-C-689 (HypXY) and precursor peptide-encoding plasmid mutant_2 (RSTN1-HypAc-TEV-mutant-2) after expression at 24 °C for 22 h in 1.5 mL cultures after digestion with TEV. Data from n=4 independent biological replicates. Spliced amino acids have an x as a prefix. Spliced refers to a single tyramine excision. A representative experimental replicate is shown in panels **A, B.**

**A** Close-up plot of the representative extracted ion chromatogram (EIC) of the various peptide states; the blue trace corresponds to the Unmodified peptide. **B** Close-up plot of the m/z values of the peptides detected by the mass spectrometer, with the shaded colored areas corresponding to where the expected m/z values for the peptide m/z

(Caption continued on the next page.)

traces should occur, with the observed spectrum plotted in black. The Unmodified peptide(s) were found in the mass spectrum and the Spliced peptide(s) were not detected. **C** Bar plot showing the relative abundance of the various peptide states at charge z=2; the bars represent the average relative abundances normalized to total peak areas, with each individual experimental replicate value plotted as a black dot. **D** Mirrored $MS^2$ plot for the Unmodified peptide. In this mirror plot, the full fragmentation spectrum is plotted in black pointing upward, and the matched fragment b ions (blue) and y ions (red) are plotted pointing downward, mirrored against it, so that each matched fragment can be compared directly with the measured spectrum. **E** $MS^2$ spectrum of the fragments corresponding to the Unmodified peptide sequence, with matched b (blue, above) and y ions (red, below) plotted.

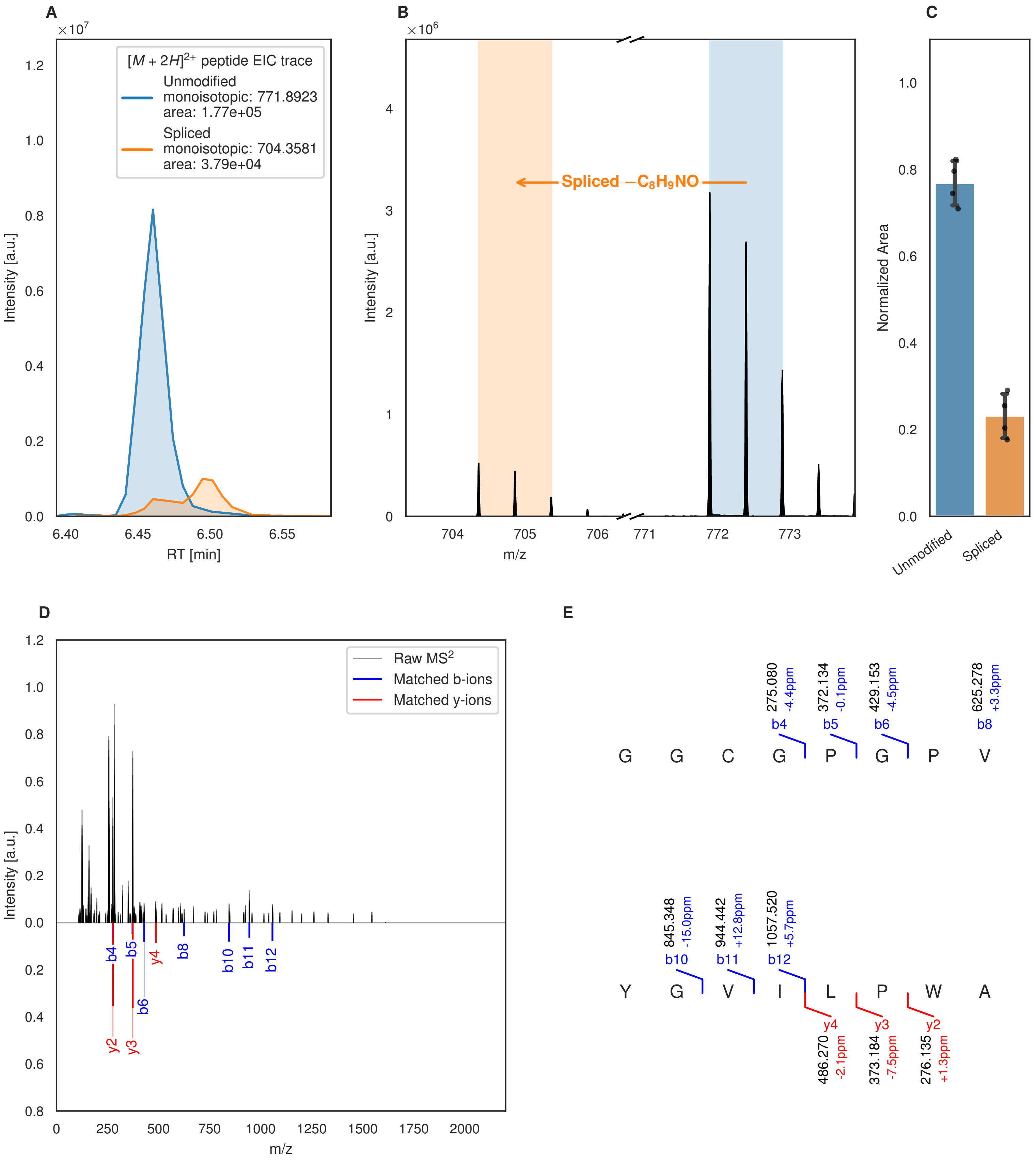
A
×10^7
[M + 2H]2+ peptide EIC trace
Unmodified
monoisotopic: 771.8923
area: 1.77e+05
Spliced
monoisotopic: 704.3581
area: 3.79e+04
Intensity [a.u.]
RT [min]
B
×10^6
Spliced −C8H9NO
Intensity [a.u.]
m/z
C
Normalized Area
Unmodified
Spliced
D
Raw MS2
Matched b-ions
Matched y-ions
Intensity [a.u.]
m/z
E
275.080 -4.4ppm b4
372.134 -0.1ppm b5
429.153 -4.5ppm b6
625.278 +3.3ppm b8
G G C G P G P V
845.348 -15.0ppm b10
944.442 +12.8ppm b11
1057.520 +5.7ppm b12
Y G V I L P W A
y4 486.270 -2.1ppm
y3 373.184 -7.5ppm
y2 276.135 +1.3ppm

Figure S58 (figure continued from the previous page).

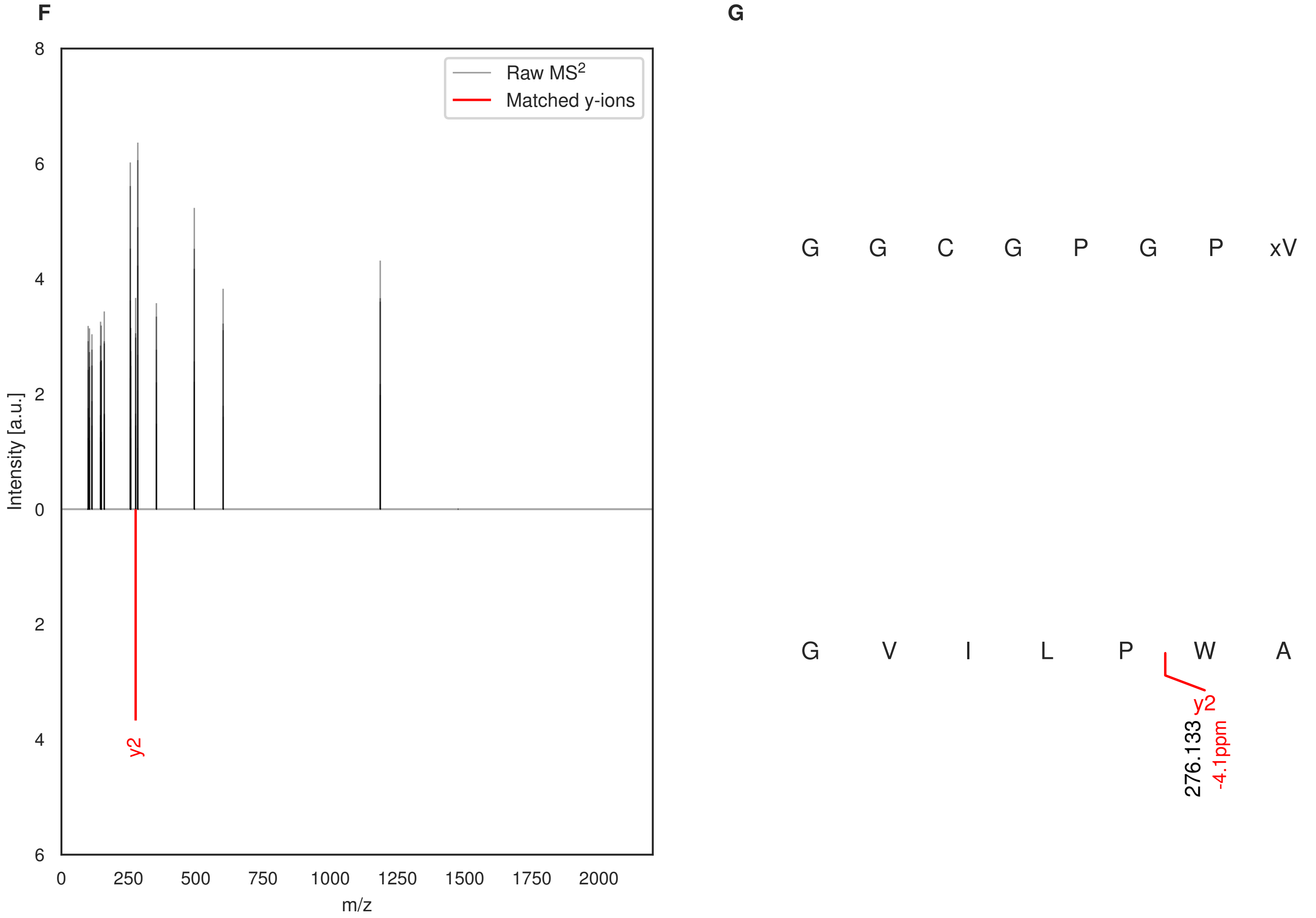


**Figure S58: Modification of the mutant_2 precursor peptide by PlpXY.** LC-MS analysis of IMAC-purified precursor peptides from *E. coli* BL21(DE3)Tuner containing splicease-encoding plasmid TJ-C-559 (PlpXY) and precursor peptide-encoding plasmid mutant_2 (RSTN1-PlpA3-TEV-mutant-2) after expression at 24 °C for 22 h in 1.5 mL cultures after digestion with TEV. Data from n=4 independent biological replicates. Spliced amino acids have an x as a prefix. Spliced refers to a single tyramine excision. A representative experimental replicate is shown in panels **A, B.**

**A** Close-up plot of the representative extracted ion chromatogram (EIC) of the various peptide states; the blue trace corresponds to the Unmodified peptide; the orange trace corresponds to the Spliced peptide. **B** Close-up plot of the m/z values of the peptides detected by the mass spectrometer, with the shaded colored areas corresponding to where the expected m/z values for the peptide m/z traces should occur, with the observed spectrum plotted in black. The Unmodified, Spliced peptide(s) were found in the mass spectrum. **C** Bar plot showing the relative abundance of the various peptide states at charge z=2; the bars represent the average relative abundances normalized to total peak areas, with each individual experimental replicate value plotted as a black dot. **D** Mirrored MS$^2$ plot for the Unmodified peptide. In this mirror plot, the full fragmentation spectrum is plotted in black pointing upward, and the matched fragment b ions (blue) and y ions (red) are plotted pointing downward, mirrored against it, so that each matched fragment can be compared directly with the measured spectrum. **E** MS$^2$ spectrum of the fragments corresponding to the Unmodified peptide sequence, with matched b (blue, above) and y ions (red, below) plotted. **F** Mirrored MS$^2$ plot for the Spliced peptide. In this mirror plot, the full fragmentation spectrum is plotted in black pointing upward, and the matched fragment b ions (blue) and y ions (red) are plotted pointing downward, mirrored against it, so that each matched fragment can be compared directly with the measured spectrum. **G** MS$^2$ spectrum of the fragments corresponding to the Spliced peptide sequence, with matched b (blue, above) and y ions (red, below) plotted.

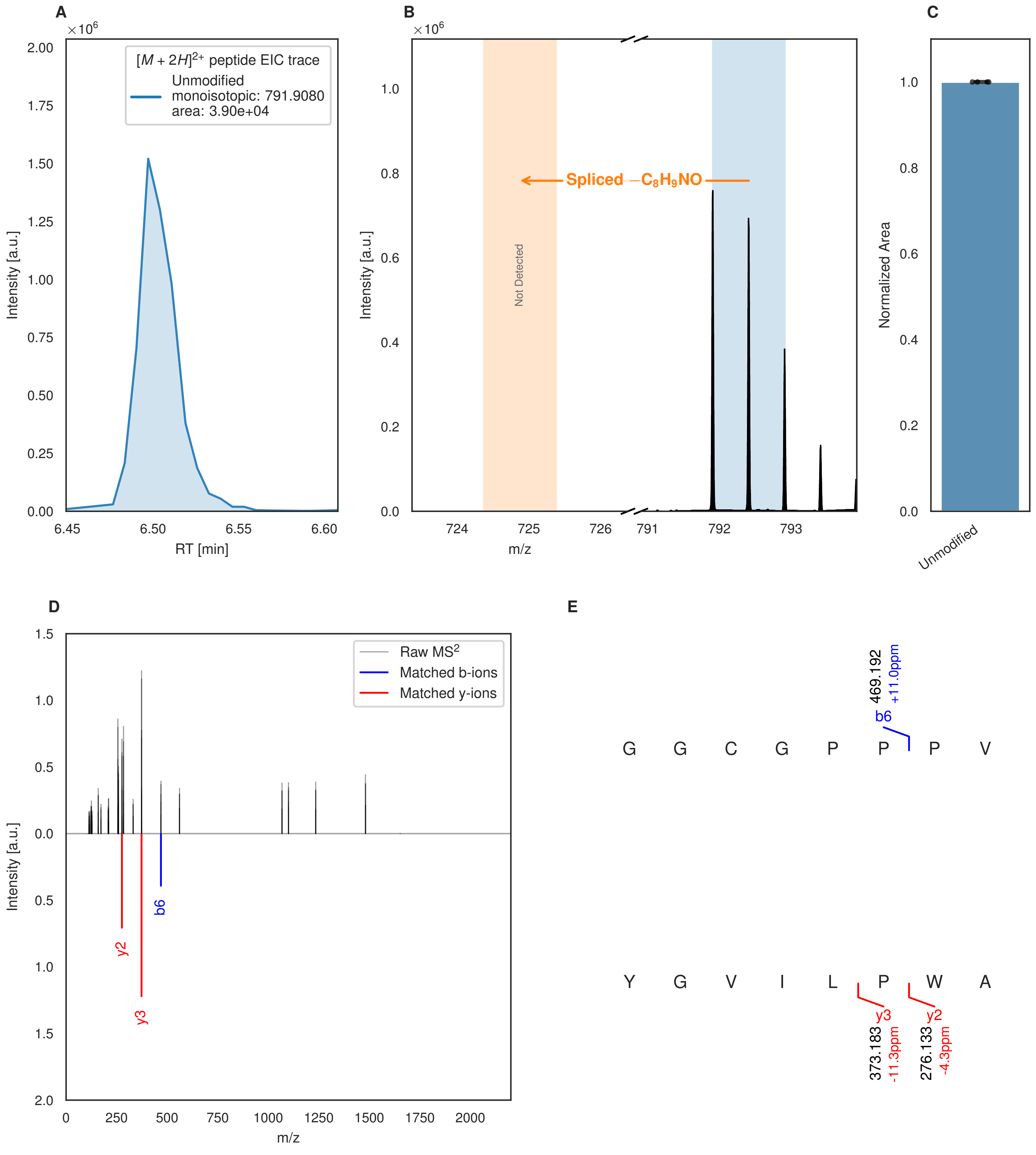


**Figure S59: Modification of the mutant_3 precursor peptide by HypXY.** LC-MS analysis of IMAC-purified precursor peptides from *E. coli* BL21(DE3)Tuner containing splicease-encoding plasmid TJ-C-689 (HypXY) and precursor peptide-encoding plasmid mutant_3 (RSTN1-HypAc-TEV-mutant-3) after expression at 24 °C for 22 h in 1.5 mL cultures after digestion with TEV. Data from n=4 independent biological replicates. Spliced amino acids have an x as a prefix. Spliced refers to a single tyramine excision. A representative experimental replicate is shown in panels **A, B.**

**A** Close-up plot of the representative extracted ion chromatogram (EIC) of the various peptide states; the blue trace corresponds to the Unmodified peptide. **B** Close-up plot of the m/z values of the peptides detected by the mass spectrometer, with the shaded colored areas corresponding to where the expected m/z values for the peptide m/z

(Caption continued on the next page.)

traces should occur, with the observed spectrum plotted in black. The Unmodified peptide(s) were found in the mass spectrum and the Spliced peptide(s) were not detected. **C** Bar plot showing the relative abundance of the various peptide states at charge z=2; the bars represent the average relative abundances normalized to total peak areas, with each individual experimental replicate value plotted as a black dot. **D** Mirrored $MS^2$ plot for the Unmodified peptide. In this mirror plot, the full fragmentation spectrum is plotted in black pointing upward, and the matched fragment b ions (blue) and y ions (red) are plotted pointing downward, mirrored against it, so that each matched fragment can be compared directly with the measured spectrum. **E** $MS^2$ spectrum of the fragments corresponding to the Unmodified peptide sequence, with matched b (blue, above) and y ions (red, below) plotted.

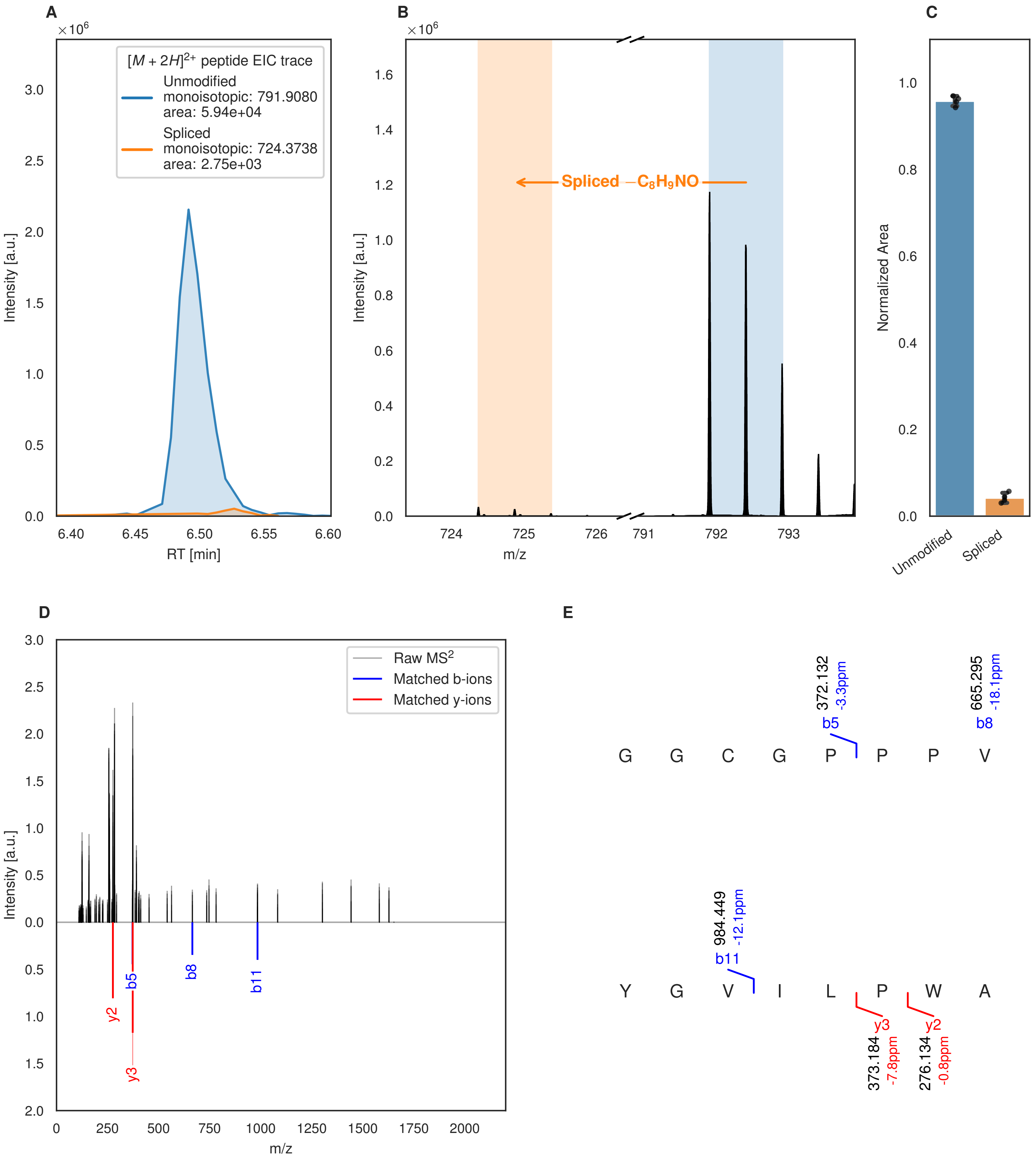
A
B
C
D
E
[M + 2H]2+ peptide EIC trace
Unmodified
monoisotopic: 791.9080
area: 5.94e+04
Spliced
monoisotopic: 724.3738
area: 2.75e+03
Intensity [a.u.]
RT [min]
×10^6
Spliced −C8H9NO
m/z
Normalized Area
Unmodified
Spliced
Raw MS2
Matched b-ions
Matched y-ions
y2
b5
y3
b8
b11
G G C G P P P V
372.132
-3.3ppm
b5
665.295
-18.1ppm
b8
Y G V I L P W A
984.449
-12.1ppm
b11
y3
373.184
-7.8ppm
y2
276.134
-0.8ppm

Figure S60 (figure continued from the previous page).

F

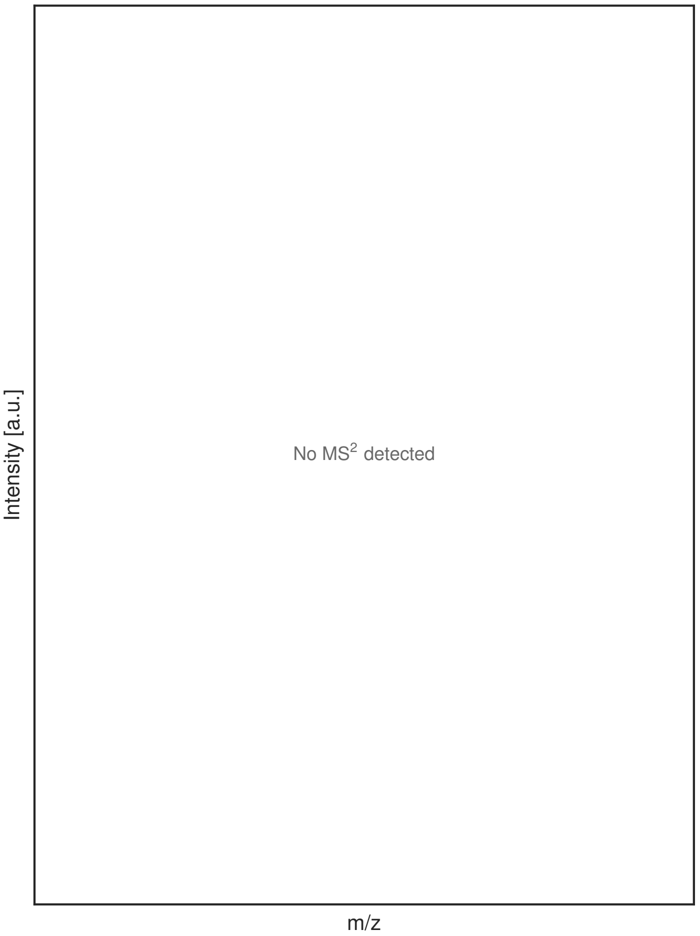


G

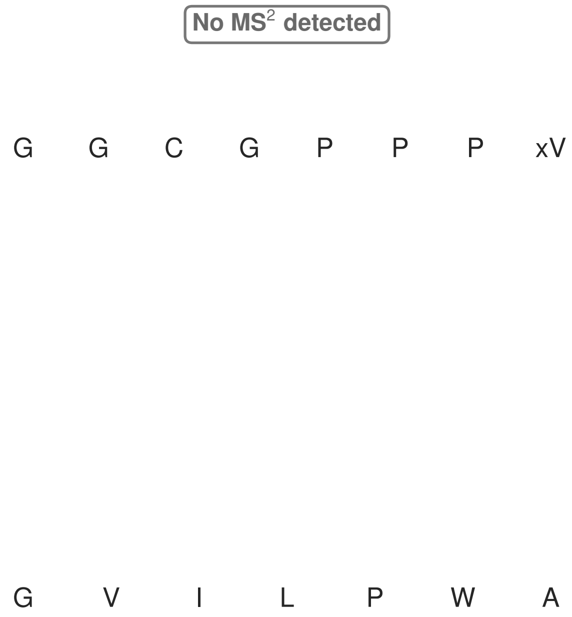


**Figure S60: Modification of the mutant_3 precursor peptide by PlpXY.** LC-MS analysis of IMAC-purified precursor peptides from *E. coli* BL21(DE3)Tuner containing splicease-encoding plasmid TJ-C-559 (PlpXY) and precursor peptide-encoding plasmid mutant_3 (RSTN1-PlpA3-TEV-mutant-3) after expression at 24 °C for 22 h in 1.5 mL cultures after digestion with TEV. Data from n=4 independent biological replicates. Spliced amino acids have an x as a prefix. Spliced refers to a single tyramine excision. A representative experimental replicate is shown in panels **A, B.**

**A** Close-up plot of the representative extracted ion chromatogram (EIC) of the various peptide states; the blue trace corresponds to the Unmodified peptide; the orange trace corresponds to the Spliced peptide. **B** Close-up plot of the m/z values of the peptides detected by the mass spectrometer, with the shaded colored areas corresponding to where the expected m/z values for the peptide m/z traces should occur, with the observed spectrum plotted in black. The Unmodified, Spliced peptide(s) were found in the mass spectrum. **C** Bar plot showing the relative abundance of the various peptide states at charge z=2; the bars represent the average relative abundances normalized to total peak areas, with each individual experimental replicate value plotted as a black dot. **D** Mirrored MS$^2$ plot for the Unmodified peptide. In this mirror plot, the full fragmentation spectrum is plotted in black pointing upward, and the matched fragment b ions (blue) and y ions (red) are plotted pointing downward, mirrored against it, so that each matched fragment can be compared directly with the measured spectrum. **E** MS$^2$ spectrum of the fragments corresponding to the Unmodified peptide sequence, with matched b (blue, above) and y ions (red, below) plotted. **F** Mirrored MS$^2$ plot for the Spliced peptide. In this mirror plot, the full fragmentation spectrum is plotted in black pointing upward, and the matched fragment b ions (blue) and y ions (red) are plotted pointing downward, mirrored against it, so that each matched fragment can be compared directly with the measured spectrum. **G** MS$^2$ spectrum of the fragments corresponding to the Spliced peptide sequence, with matched b (blue, above) and y ions (red, below) plotted.

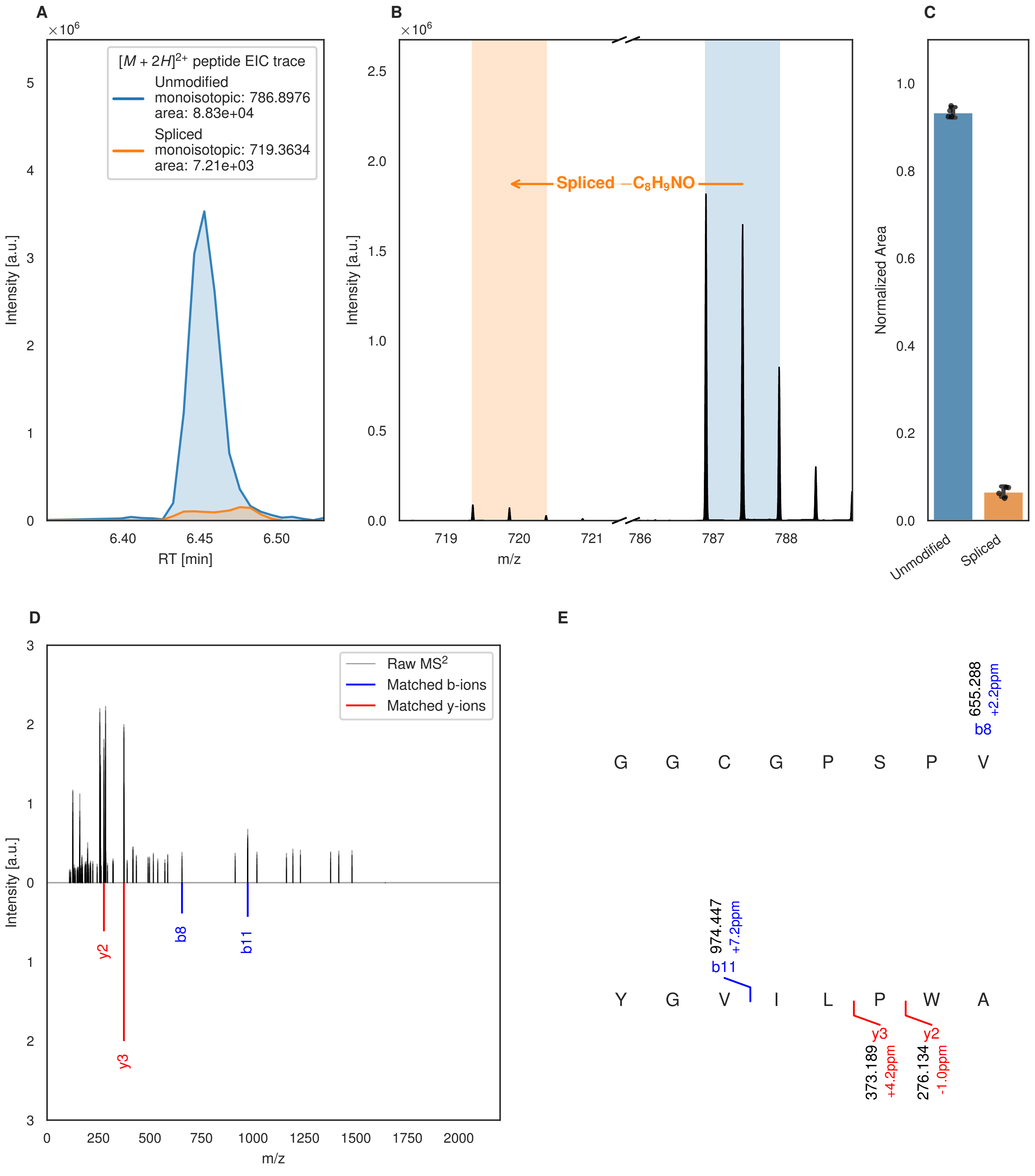

A
×10^6
[M + 2H]^2+ peptide EIC trace
Unmodified
monoisotopic: 786.8976
area: 8.83e+04
Spliced
monoisotopic: 719.3634
area: 7.21e+03
Intensity [a.u.]
RT [min]
B
Spliced −C8H9NO
m/z
C
Normalized Area
Unmodified
Spliced
D
Raw MS2
Matched b-ions
Matched y-ions
y2
y3
b8
b11
E
G G C G P S P V
655.288
+2.2ppm
b8
974.447
+7.2ppm
b11
Y G V I L P W A
y3
373.189
+4.2ppm
y2
276.134
-1.0ppm

Figure S61 (figure continued from the previous page).

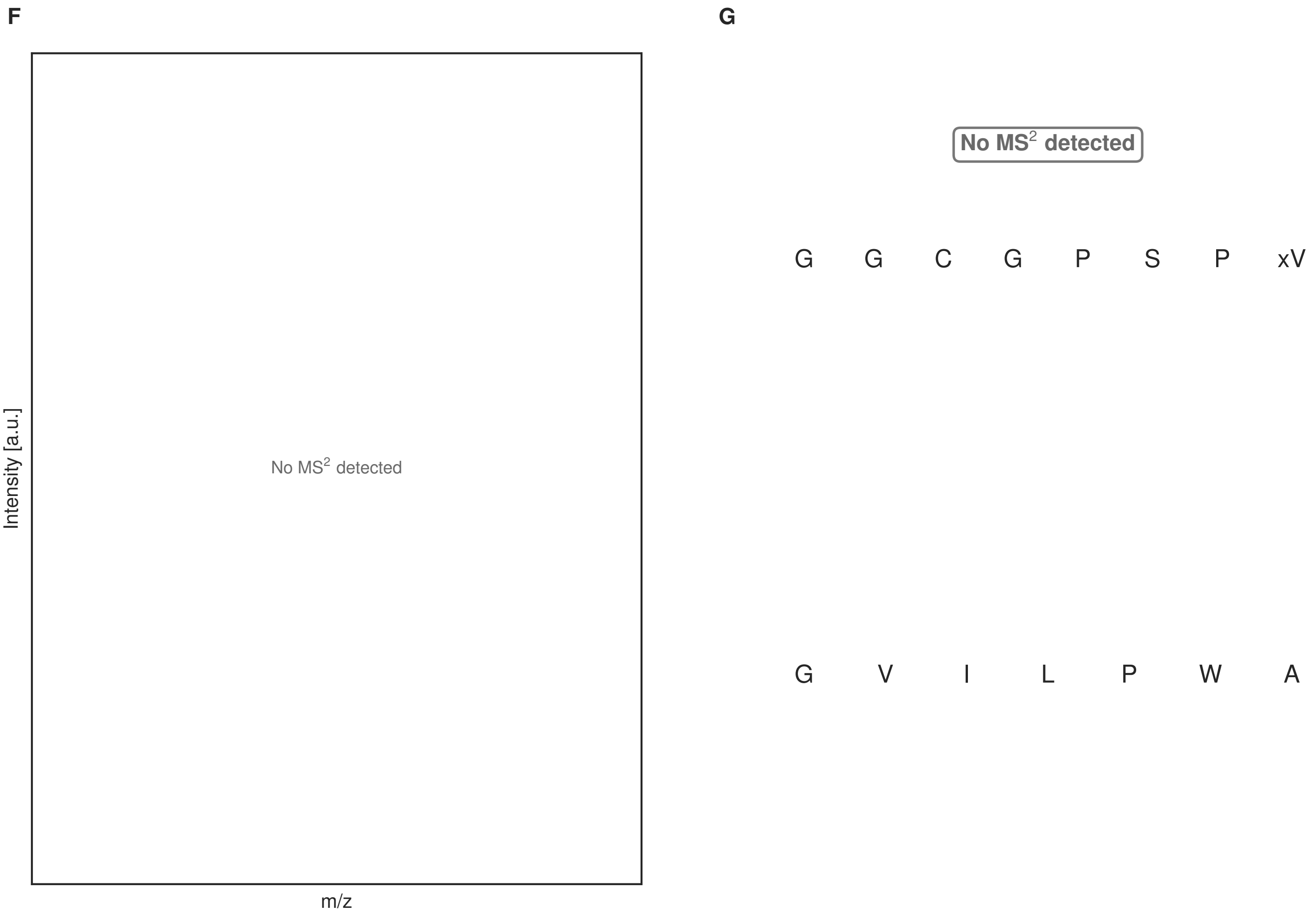


**Figure S61: Modification of the mutant_4 precursor peptide by HypXY.** LC-MS analysis of IMAC-purified precursor peptides from *E. coli* BL21(DE3)Tuner containing splicease-encoding plasmid TJ-C-689 (HypXY) and precursor peptide-encoding plasmid mutant_4 (RSTN1-HypAc-TEV-mutant-4) after expression at 24 °C for 22 h in 1.5 mL cultures after digestion with TEV. Data from n=4 independent biological replicates. Spliced amino acids have an x as a prefix. Spliced refers to a single tyramine excision. A representative experimental replicate is shown in panels **A, B.**

**A** Close-up plot of the representative extracted ion chromatogram (EIC) of the various peptide states; the blue trace corresponds to the Unmodified peptide; the orange trace corresponds to the Spliced peptide. **B** Close-up plot of the m/z values of the peptides detected by the mass spectrometer, with the shaded colored areas corresponding to where the expected m/z values for the peptide m/z traces should occur, with the observed spectrum plotted in black. The Unmodified, Spliced peptide(s) were found in the mass spectrum. **C** Bar plot showing the relative abundance of the various peptide states at charge z=2; the bars represent the average relative abundances normalized to total peak areas, with each individual experimental replicate value plotted as a black dot. **D** Mirrored $MS^2$ plot for the Unmodified peptide. In this mirror plot, the full fragmentation spectrum is plotted in black pointing upward, and the matched fragment b ions (blue) and y ions (red) are plotted pointing downward, mirrored against it, so that each matched fragment can be compared directly with the measured spectrum. **E** $MS^2$ spectrum of the fragments corresponding to the Unmodified peptide sequence, with matched b (blue, above) and y ions (red, below) plotted. **F** Mirrored $MS^2$ plot for the Spliced peptide. In this mirror plot, the full fragmentation spectrum is plotted in black pointing upward, and the matched fragment b ions (blue) and y ions (red) are plotted pointing downward, mirrored against it, so that each matched fragment can be compared directly with the measured spectrum. **G** $MS^2$ spectrum of the fragments corresponding to the Spliced peptide sequence, with matched b (blue, above) and y ions (red, below) plotted.

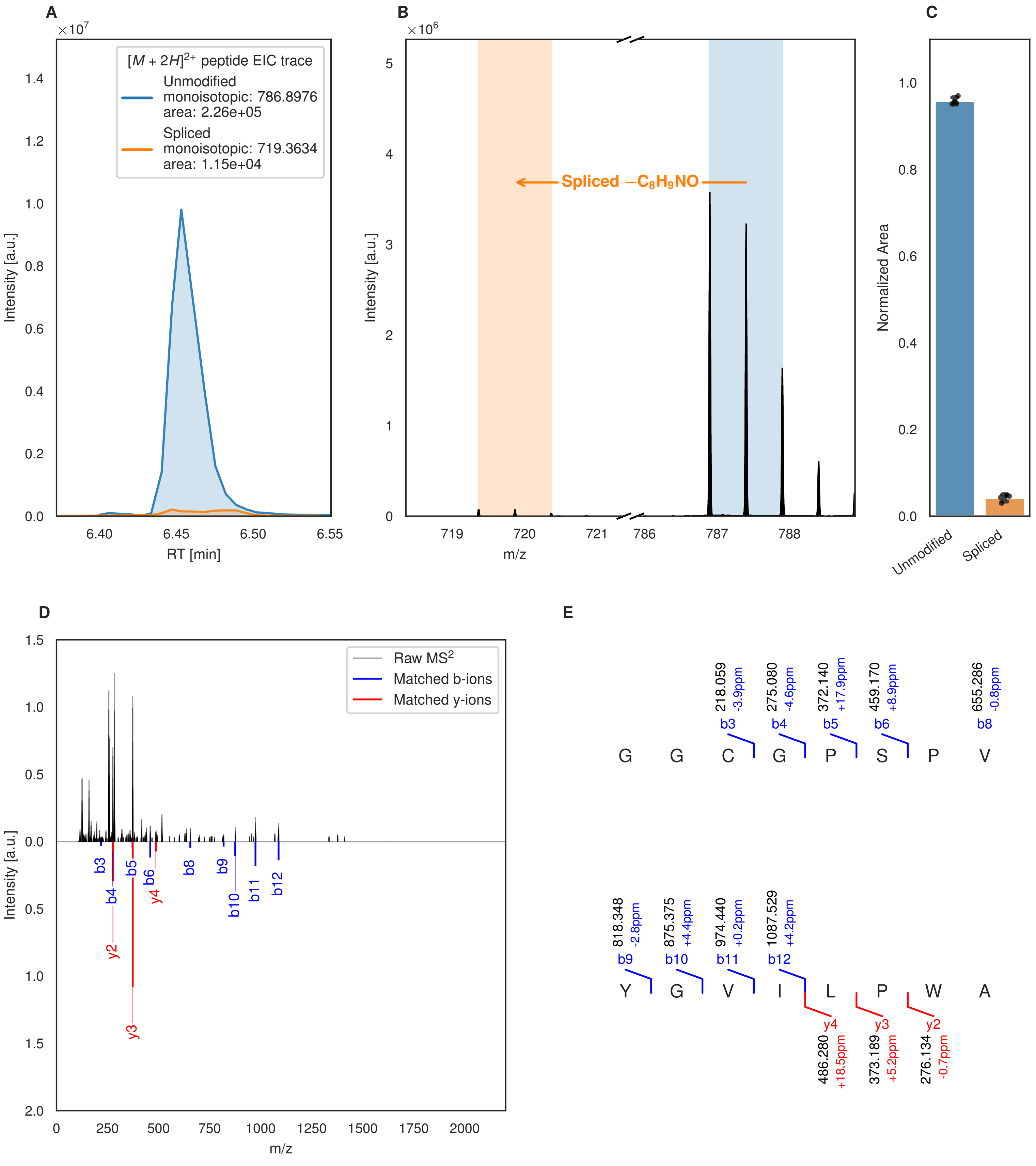
A
×10^7
[M + 2H]^2+ peptide EIC trace
Unmodified
monoisotopic: 786.8976
area: 2.26e+05
Spliced
monoisotopic: 719.3634
area: 1.15e+04
Intensity [a.u.]
RT [min]
B
×10^6
Spliced −C8H9NO
Intensity [a.u.]
m/z
C
Normalized Area
Unmodified
Spliced
D
Raw MS2
Matched b-ions
Matched y-ions
Intensity [a.u.]
m/z
E
218.059 -3.9ppm b3
275.080 -4.6ppm b4
372.140 +17.9ppm b5
459.170 +8.9ppm b6
655.286 -0.8ppm b8
G G C G P S P V
818.348 -2.8ppm b9
875.375 +4.4ppm b10
974.440 +0.2ppm b11
1087.529 +4.2ppm b12
Y G V I L P W A
y4 486.280 +18.5ppm
y3 373.189 +5.2ppm
y2 276.134 -0.7ppm

Figure S62 (figure continued from the previous page).

F

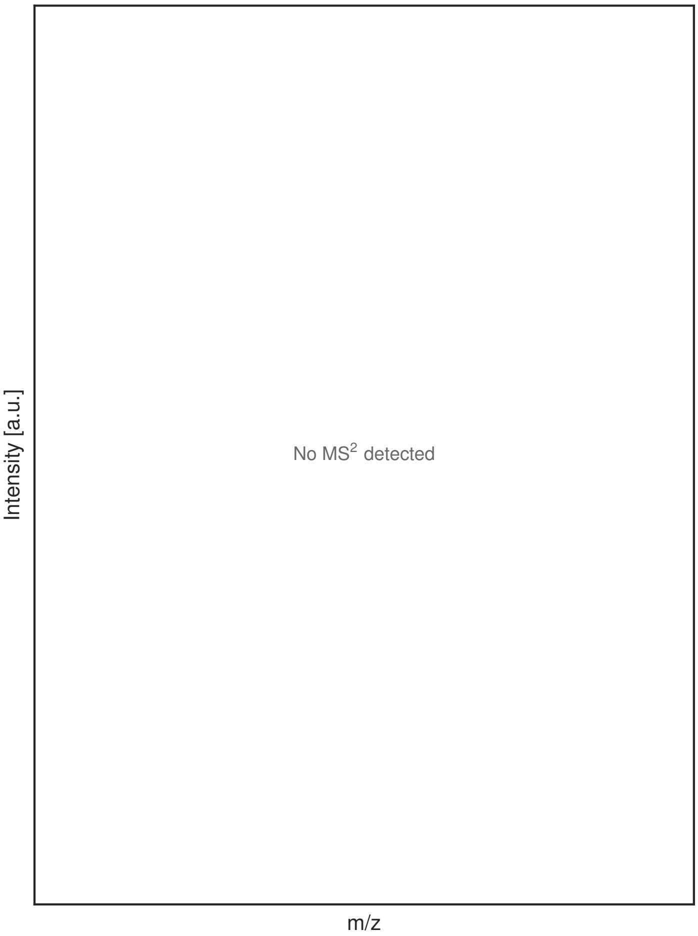


G

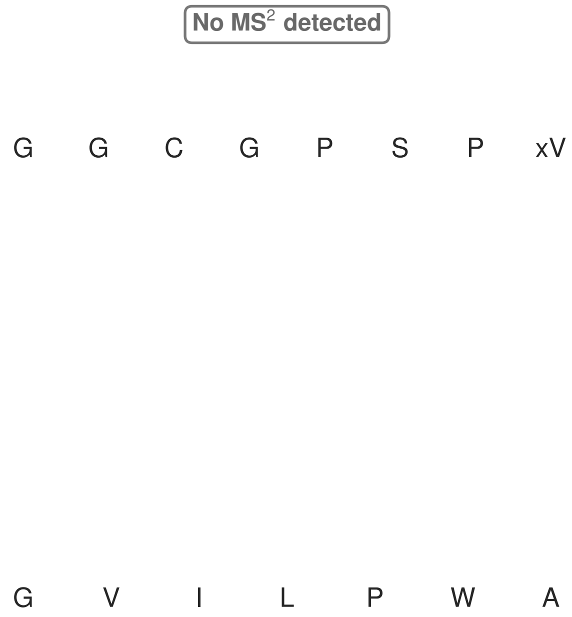


**Figure S62: Modification of the mutant_4 precursor peptide by PlpXY.** LC-MS analysis of IMAC-purified precursor peptides from *E. coli* BL21(DE3)Tuner containing splicease-encoding plasmid TJ-C-559 (PlpXY) and precursor peptide-encoding plasmid mutant_4 (RSTN1-PlpA3-TEV-mutant-4) after expression at 24 °C for 22 h in 1.5 mL cultures after digestion with TEV. Data from n=4 independent biological replicates. Spliced amino acids have an x as a prefix. Spliced refers to a single tyramine excision. A representative experimental replicate is shown in panels **A, B.**

**A** Close-up plot of the representative extracted ion chromatogram (EIC) of the various peptide states; the blue trace corresponds to the Unmodified peptide; the orange trace corresponds to the Spliced peptide. **B** Close-up plot of the m/z values of the peptides detected by the mass spectrometer, with the shaded colored areas corresponding to where the expected m/z values for the peptide m/z traces should occur, with the observed spectrum plotted in black. The Unmodified, Spliced peptide(s) were found in the mass spectrum. **C** Bar plot showing the relative abundance of the various peptide states at charge z=2; the bars represent the average relative abundances normalized to total peak areas, with each individual experimental replicate value plotted as a black dot. **D** Mirrored MS$^2$ plot for the Unmodified peptide. In this mirror plot, the full fragmentation spectrum is plotted in black pointing upward, and the matched fragment b ions (blue) and y ions (red) are plotted pointing downward, mirrored against it, so that each matched fragment can be compared directly with the measured spectrum. **E** MS$^2$ spectrum of the fragments corresponding to the Unmodified peptide sequence, with matched b (blue, above) and y ions (red, below) plotted. **F** Mirrored MS$^2$ plot for the Spliced peptide. In this mirror plot, the full fragmentation spectrum is plotted in black pointing upward, and the matched fragment b ions (blue) and y ions (red) are plotted pointing downward, mirrored against it, so that each matched fragment can be compared directly with the measured spectrum. **G** MS$^2$ spectrum of the fragments corresponding to the Spliced peptide sequence, with matched b (blue, above) and y ions (red, below) plotted.

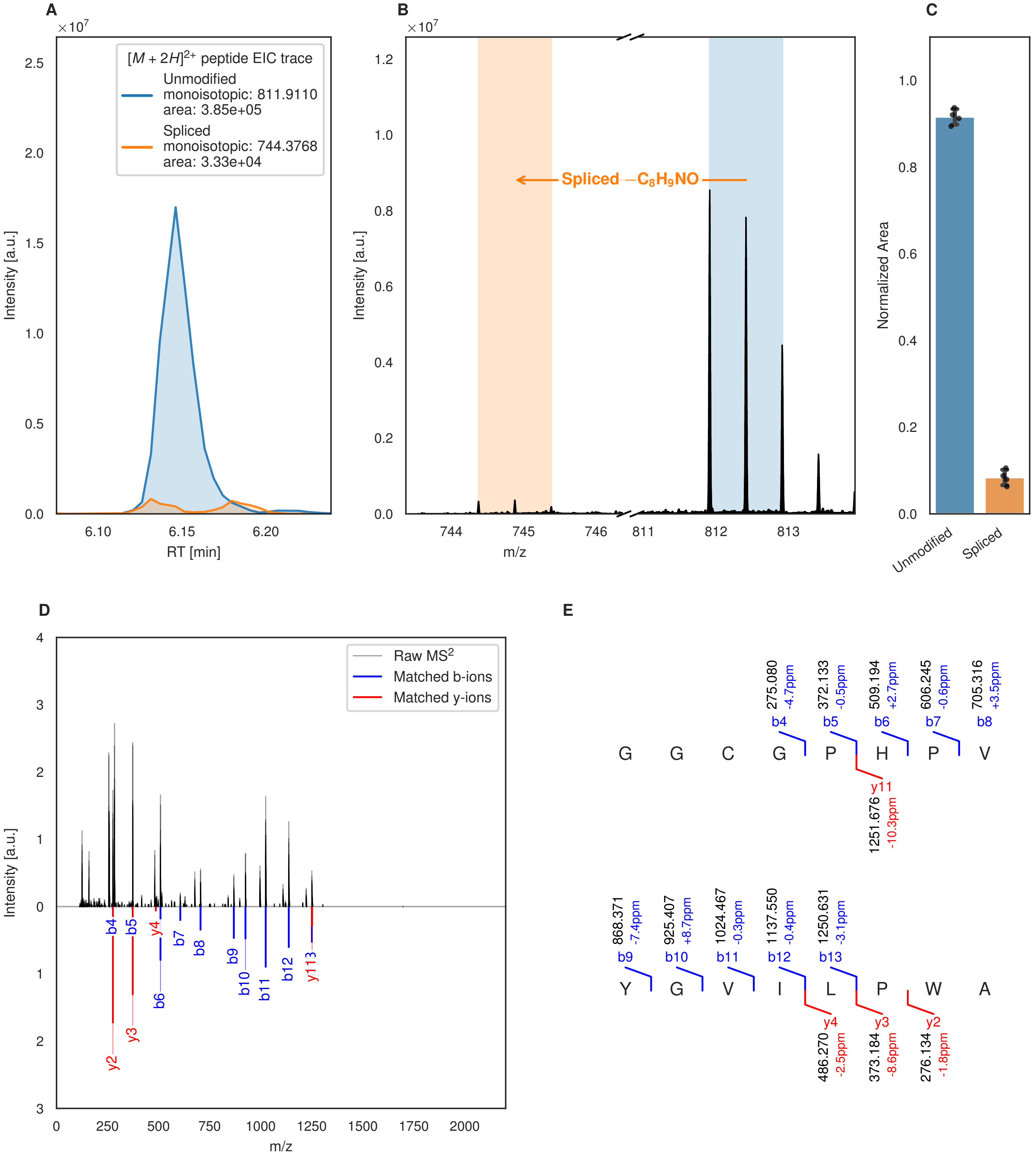
A
[M + 2H]2+ peptide EIC trace
Unmodified
monoisotopic: 811.9110
area: 3.85e+05
Spliced
monoisotopic: 744.3768
area: 3.33e+04
Intensity [a.u.]
RT [min]
B
Spliced −C8H9NO
Intensity [a.u.]
m/z
C
Normalized Area
Unmodified
Spliced
D
Raw MS2
Matched b-ions
Matched y-ions
Intensity [a.u.]
m/z
E
G G C G P H P V
b4 275.080 -4.7ppm
b5 372.133 -0.5ppm
b6 509.194 +2.7ppm
b7 606.245 -0.6ppm
b8 705.316 +3.5ppm
y11 1251.676 -10.3ppm
Y G V I L P W A
b9 868.371 -7.4ppm
b10 925.407 +8.7ppm
b11 1024.467 -0.3ppm
b12 1137.550 -0.4ppm
b13 1250.631 -3.1ppm
y4 486.270 -2.5ppm
y3 373.184 -8.6ppm
y2 276.134 -1.8ppm

Figure S63 (figure continued from the previous page).

F

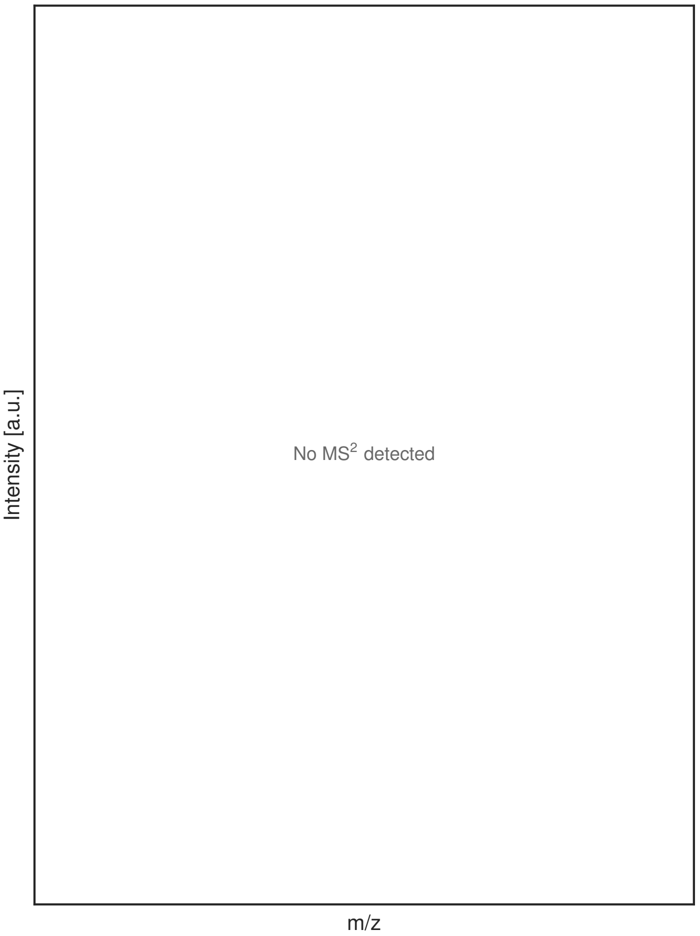


G

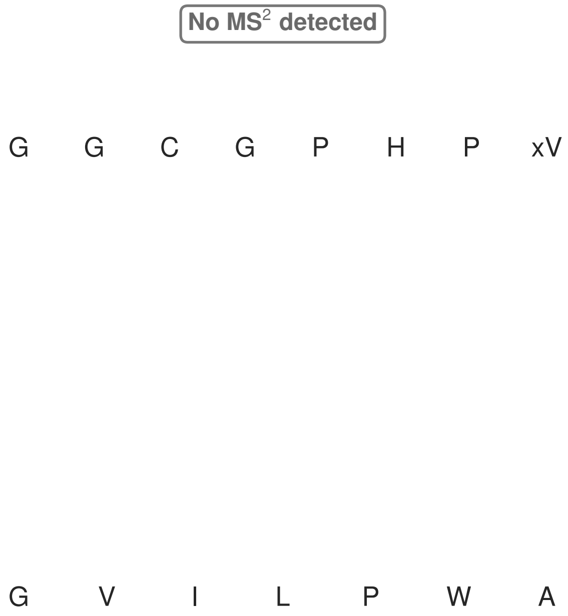


**Figure S63: Modification of the mutant_5 precursor peptide by HypXY.** LC-MS analysis of IMAC-purified precursor peptides from *E. coli* BL21(DE3)Tuner containing splicease-encoding plasmid TJ-C-689 (HypXY) and precursor peptide-encoding plasmid mutant_5 (RSTN1-HypAc-TEV-mutant-5) after expression at 24 °C for 22 h in 1.5 mL cultures after digestion with TEV. Data from n=4 independent biological replicates. Spliced amino acids have an x as a prefix. Spliced refers to a single tyramine excision. A representative experimental replicate is shown in panels **A, B.**

**A** Close-up plot of the representative extracted ion chromatogram (EIC) of the various peptide states; the blue trace corresponds to the Unmodified peptide; the orange trace corresponds to the Spliced peptide. **B** Close-up plot of the m/z values of the peptides detected by the mass spectrometer, with the shaded colored areas corresponding to where the expected m/z values for the peptide m/z traces should occur, with the observed spectrum plotted in black. The Unmodified, Spliced peptide(s) were found in the mass spectrum. **C** Bar plot showing the relative abundance of the various peptide states at charge z=2; the bars represent the average relative abundances normalized to total peak areas, with each individual experimental replicate value plotted as a black dot. **D** Mirrored MS$^2$ plot for the Unmodified peptide. In this mirror plot, the full fragmentation spectrum is plotted in black pointing upward, and the matched fragment b ions (blue) and y ions (red) are plotted pointing downward, mirrored against it, so that each matched fragment can be compared directly with the measured spectrum. **E** MS$^2$ spectrum of the fragments corresponding to the Unmodified peptide sequence, with matched b (blue, above) and y ions (red, below) plotted. **F** Mirrored MS$^2$ plot for the Spliced peptide. In this mirror plot, the full fragmentation spectrum is plotted in black pointing upward, and the matched fragment b ions (blue) and y ions (red) are plotted pointing downward, mirrored against it, so that each matched fragment can be compared directly with the measured spectrum. **G** MS$^2$ spectrum of the fragments corresponding to the Spliced peptide sequence, with matched b (blue, above) and y ions (red, below) plotted.

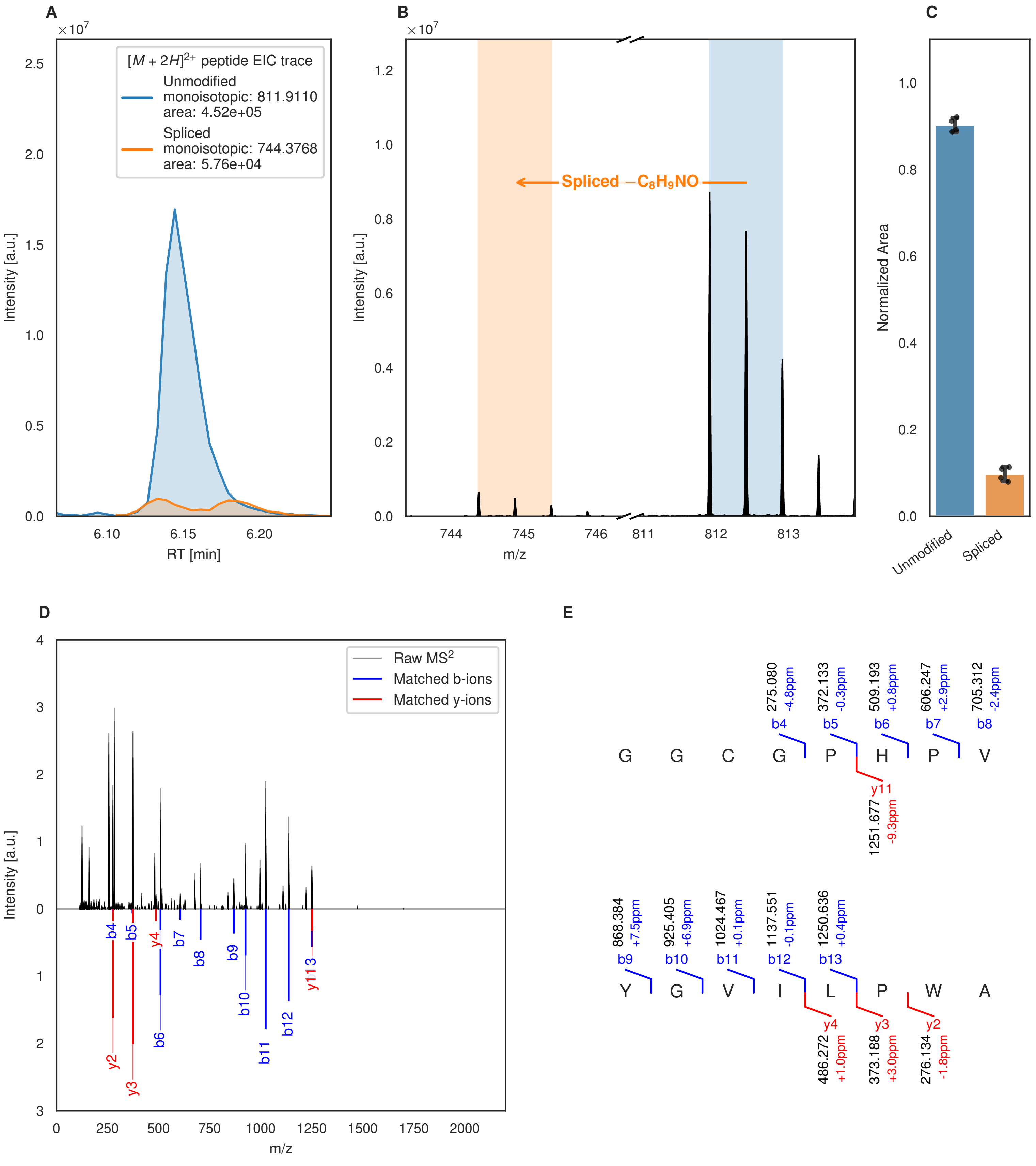
A
B
C
D
E
$[M+2H]^{2+}$ peptide EIC trace
Unmodified
monoisotopic: 811.9110
area: 4.52e+05
Spliced
monoisotopic: 744.3768
area: 5.76e+04
Intensity [a.u.]
RT [min]
m/z
Spliced −$C_8H_9NO$
Normalized Area
Unmodified
Spliced
Raw $MS^2$
Matched b-ions
Matched y-ions
G G C G P H P V
Y G V I L P W A

Figure S64 (figure continued from the previous page).

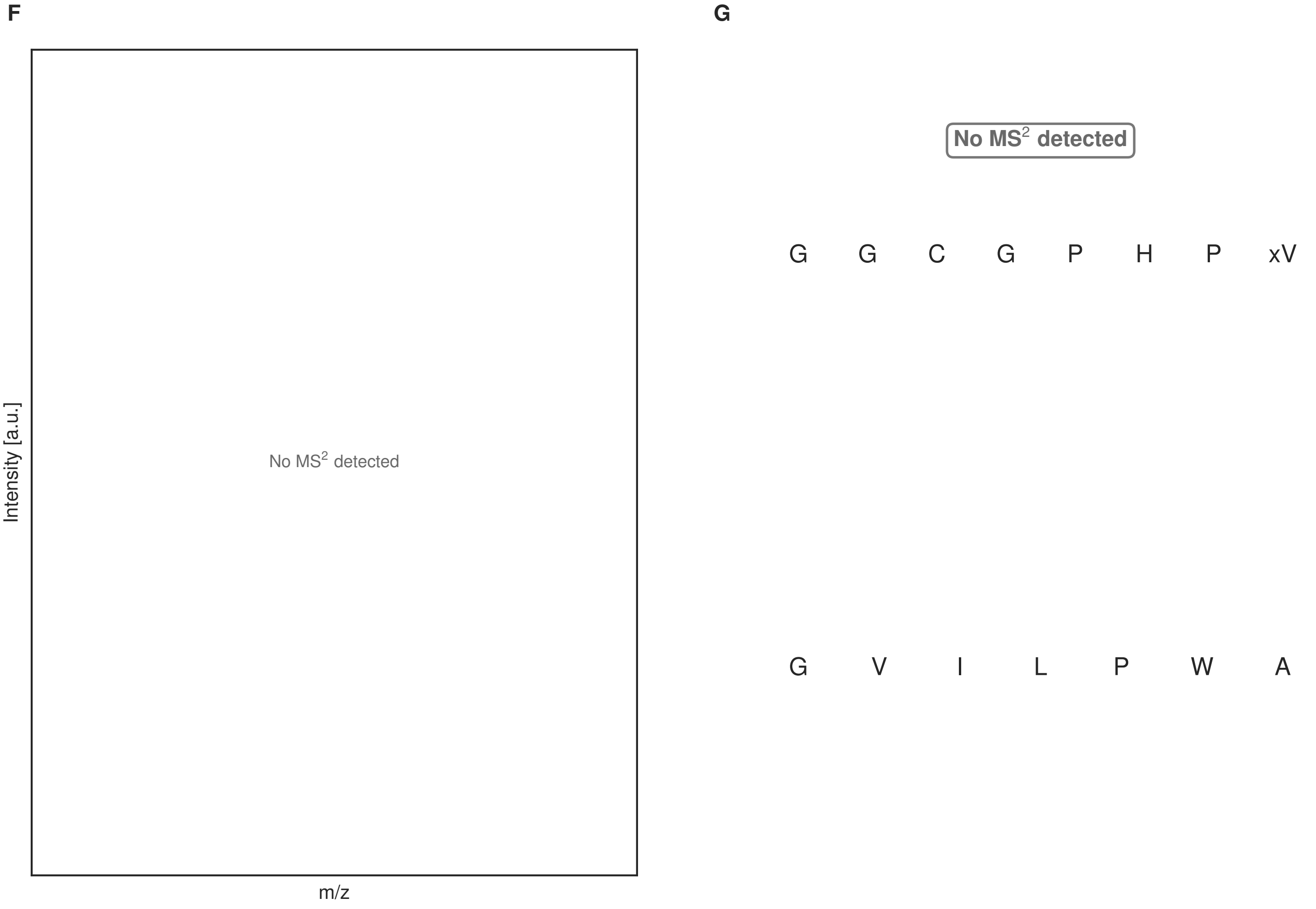


**Figure S64: Modification of the mutant_5 precursor peptide by PlpXY.** LC-MS analysis of IMAC-purified precursor peptides from *E. coli* BL21(DE3)Tuner containing splicease-encoding plasmid TJ-C-559 (PlpXY) and precursor peptide-encoding plasmid mutant_5 (RSTN1-PlpA3-TEV-mutant-5) after expression at 24 °C for 22 h in 1.5 mL cultures after digestion with TEV. Data from n=4 independent biological replicates. Spliced amino acids have an x as a prefix. Spliced refers to a single tyramine excision. A representative experimental replicate is shown in panels **A, B.**

**A** Close-up plot of the representative extracted ion chromatogram (EIC) of the various peptide states; the blue trace corresponds to the Unmodified peptide; the orange trace corresponds to the Spliced peptide. **B** Close-up plot of the m/z values of the peptides detected by the mass spectrometer, with the shaded colored areas corresponding to where the expected m/z values for the peptide m/z traces should occur, with the observed spectrum plotted in black. The Unmodified, Spliced peptide(s) were found in the mass spectrum. **C** Bar plot showing the relative abundance of the various peptide states at charge z=2; the bars represent the average relative abundances normalized to total peak areas, with each individual experimental replicate value plotted as a black dot. **D** Mirrored $MS^2$ plot for the Unmodified peptide. In this mirror plot, the full fragmentation spectrum is plotted in black pointing upward, and the matched fragment b ions (blue) and y ions (red) are plotted pointing downward, mirrored against it, so that each matched fragment can be compared directly with the measured spectrum. **E** $MS^2$ spectrum of the fragments corresponding to the Unmodified peptide sequence, with matched b (blue, above) and y ions (red, below) plotted. **F** Mirrored $MS^2$ plot for the Spliced peptide. In this mirror plot, the full fragmentation spectrum is plotted in black pointing upward, and the matched fragment b ions (blue) and y ions (red) are plotted pointing downward, mirrored against it, so that each matched fragment can be compared directly with the measured spectrum. **G** $MS^2$ spectrum of the fragments corresponding to the Spliced peptide sequence, with matched b (blue, above) and y ions (red, below) plotted.

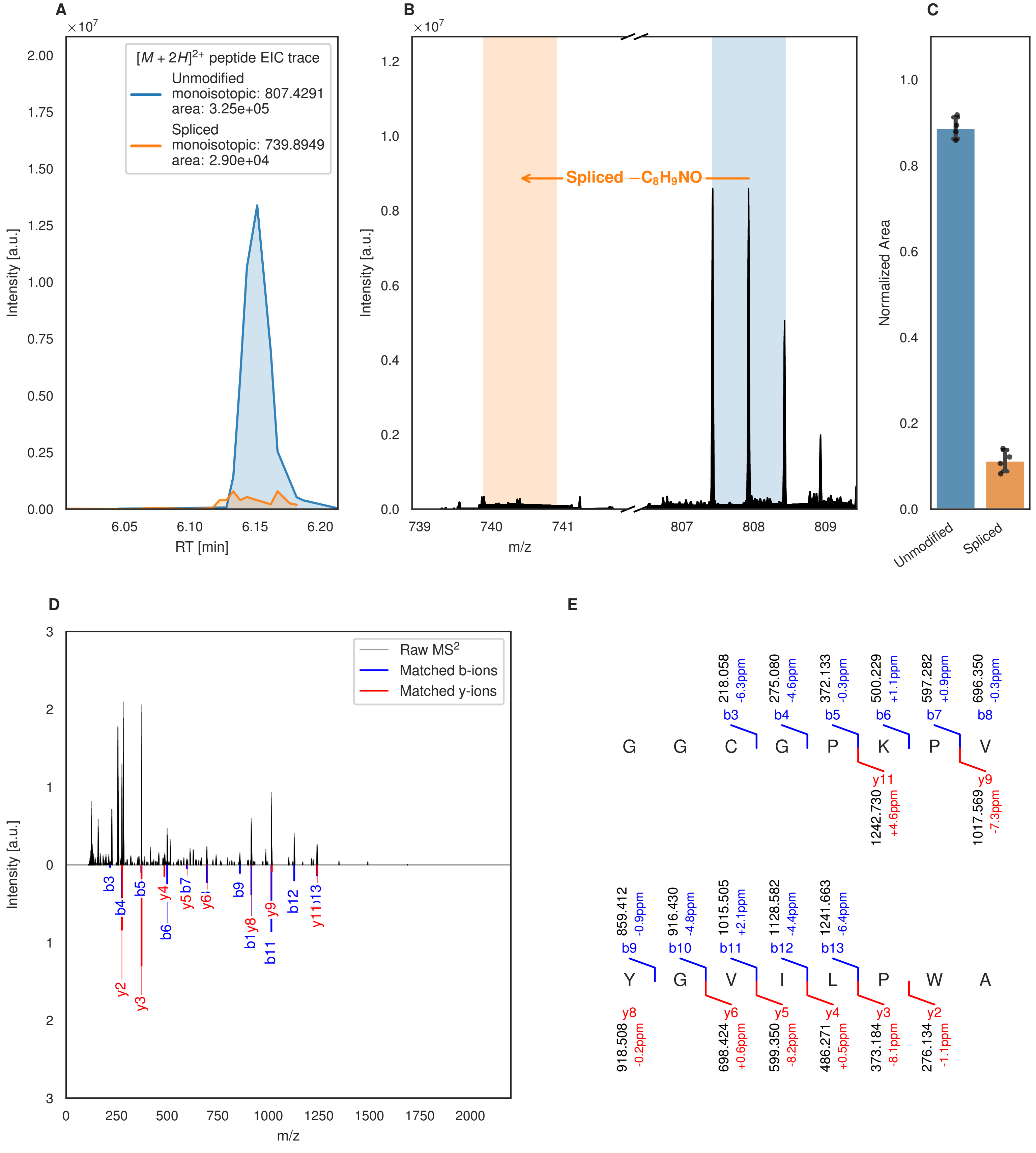
A
B
C
D
E
[M + 2H]2+ peptide EIC trace
Unmodified
monoisotopic: 807.4291
area: 3.25e+05
Spliced
monoisotopic: 739.8949
area: 2.90e+04
Intensity [a.u.]
RT [min]
Spliced −C8H9NO
m/z
Normalized Area
Unmodified
Spliced
Raw MS2
Matched b-ions
Matched y-ions
G G C G P K P V
Y G V I L P W A
218.058 -6.3ppm b3
275.080 -4.6ppm b4
372.133 -0.3ppm b5
500.229 +1.1ppm b6
597.282 +0.9ppm b7
696.350 -0.3ppm b8
y11 1242.730 +4.6ppm
y9 1017.569 -7.3ppm
859.412 -0.9ppm b9
916.430 -4.8ppm b10
1015.505 +2.1ppm b11
1128.582 -4.4ppm b12
1241.663 -6.4ppm b13
y8 918.508 -0.2ppm
y6 698.424 +0.6ppm
y5 599.350 -8.2ppm
y4 486.271 +0.5ppm
y3 373.184 -8.1ppm
y2 276.134 -1.1ppm

Figure S65 (figure continued from the previous page).

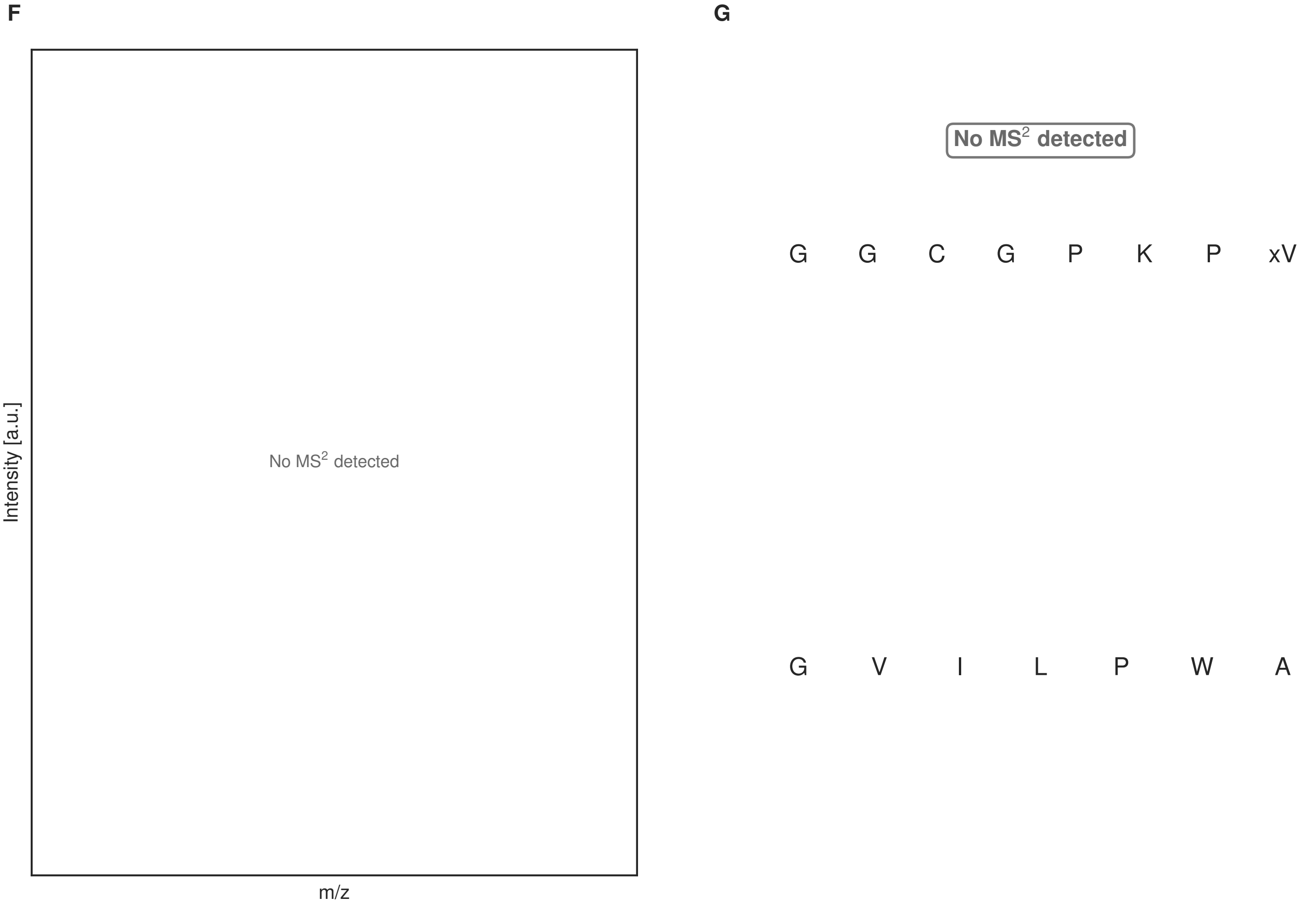


**Figure S65: Modification of the mutant_6 precursor peptide by HypXY.** LC-MS analysis of IMAC-purified precursor peptides from *E. coli* BL21(DE3)Tuner containing splicease-encoding plasmid TJ-C-689 (HypXY) and precursor peptide-encoding plasmid mutant_6 (RSTN1-HypAc-TEV-mutant-6) after expression at 24 °C for 22 h in 1.5 mL cultures after digestion with TEV. Data from n=4 independent biological replicates. Spliced amino acids have an x as a prefix. Spliced refers to a single tyramine excision. A representative experimental replicate is shown in panels **A, B.**

**A** Close-up plot of the representative extracted ion chromatogram (EIC) of the various peptide states; the blue trace corresponds to the Unmodified peptide; the orange trace corresponds to the Spliced peptide. **B** Close-up plot of the m/z values of the peptides detected by the mass spectrometer, with the shaded colored areas corresponding to where the expected m/z values for the peptide m/z traces should occur, with the observed spectrum plotted in black. The Unmodified, Spliced peptide(s) were found in the mass spectrum. **C** Bar plot showing the relative abundance of the various peptide states at charge z=2; the bars represent the average relative abundances normalized to total peak areas, with each individual experimental replicate value plotted as a black dot. **D** Mirrored $MS^2$ plot for the Unmodified peptide. In this mirror plot, the full fragmentation spectrum is plotted in black pointing upward, and the matched fragment b ions (blue) and y ions (red) are plotted pointing downward, mirrored against it, so that each matched fragment can be compared directly with the measured spectrum. **E** $MS^2$ spectrum of the fragments corresponding to the Unmodified peptide sequence, with matched b (blue, above) and y ions (red, below) plotted. **F** Mirrored $MS^2$ plot for the Spliced peptide. In this mirror plot, the full fragmentation spectrum is plotted in black pointing upward, and the matched fragment b ions (blue) and y ions (red) are plotted pointing downward, mirrored against it, so that each matched fragment can be compared directly with the measured spectrum. **G** $MS^2$ spectrum of the fragments corresponding to the Spliced peptide sequence, with matched b (blue, above) and y ions (red, below) plotted.

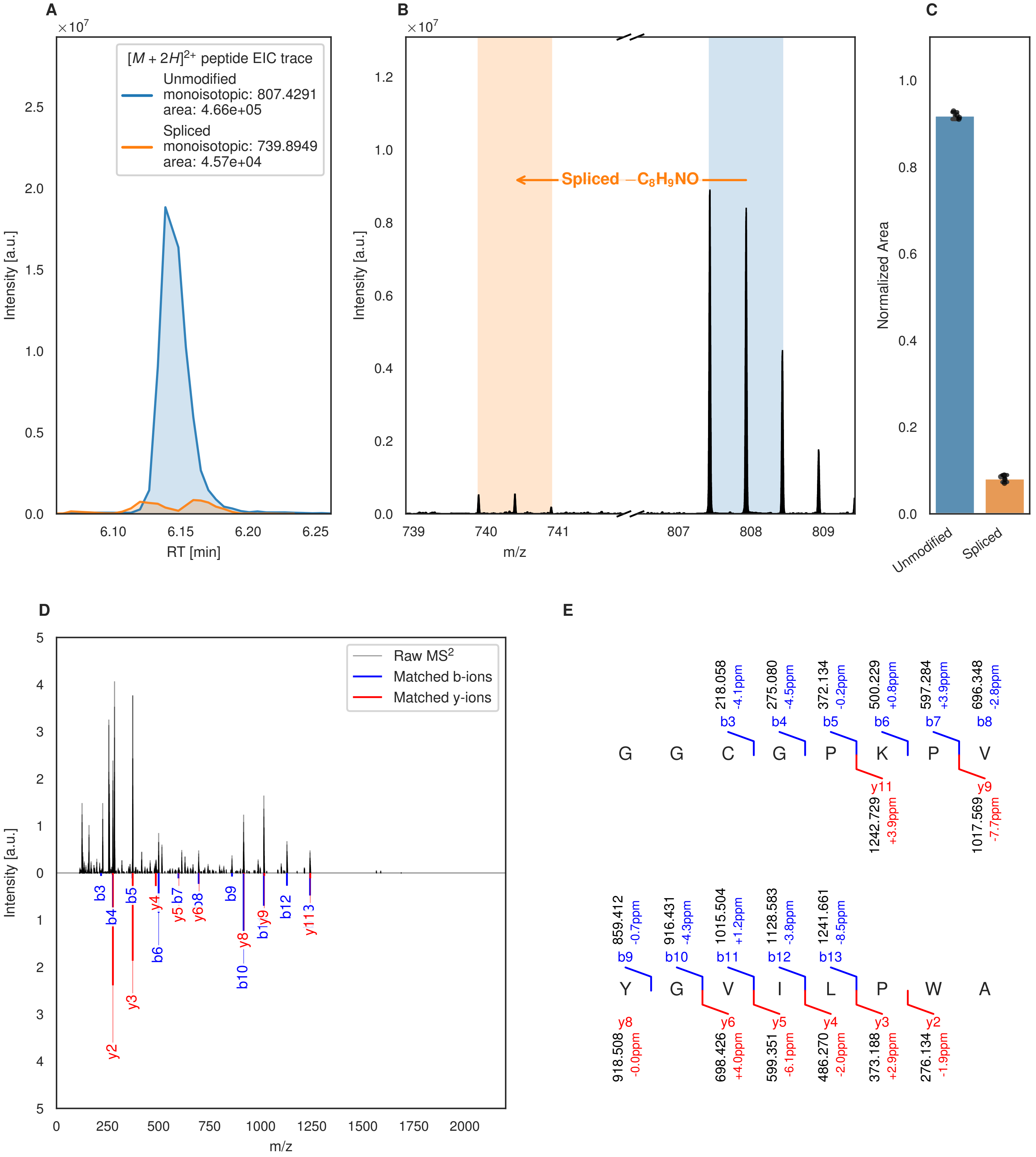
A
B
C
D
E
[M + 2H]2+ peptide EIC trace
Unmodified
monoisotopic: 807.4291
area: 4.66e+05
Spliced
monoisotopic: 739.8949
area: 4.57e+04
Intensity [a.u.]
RT [min]
Spliced −C8H9NO
m/z
Normalized Area
Unmodified
Spliced
Raw MS2
Matched b-ions
Matched y-ions
218.058 -4.1ppm b3
275.080 -4.5ppm b4
372.134 -0.2ppm b5
500.229 +0.8ppm b6
597.284 +3.9ppm b7
696.348 -2.8ppm b8
G G C G P K P V
y11 1242.729 +3.9ppm
y9 1017.569 -7.7ppm
859.412 -0.7ppm b9
916.431 -4.3ppm b10
1015.504 +1.2ppm b11
1128.583 -3.8ppm b12
1241.661 -8.5ppm b13
Y G V I L P W A
y8 918.508 -0.0ppm
y6 698.426 +4.0ppm
y5 599.351 -6.1ppm
y4 486.270 -2.0ppm
y3 373.188 +2.9ppm
y2 276.134 -1.9ppm

F

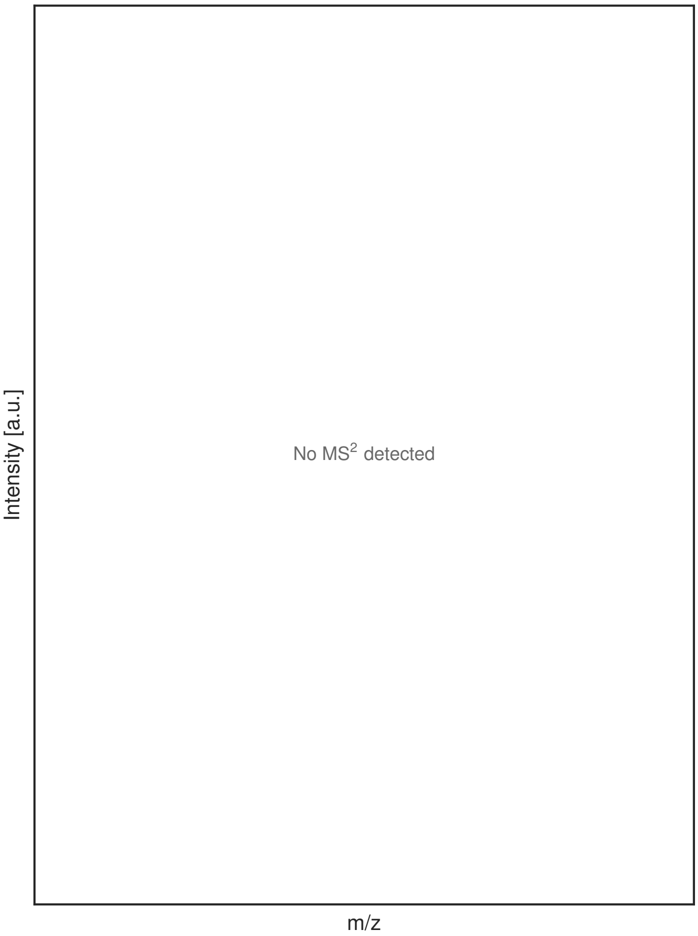


G

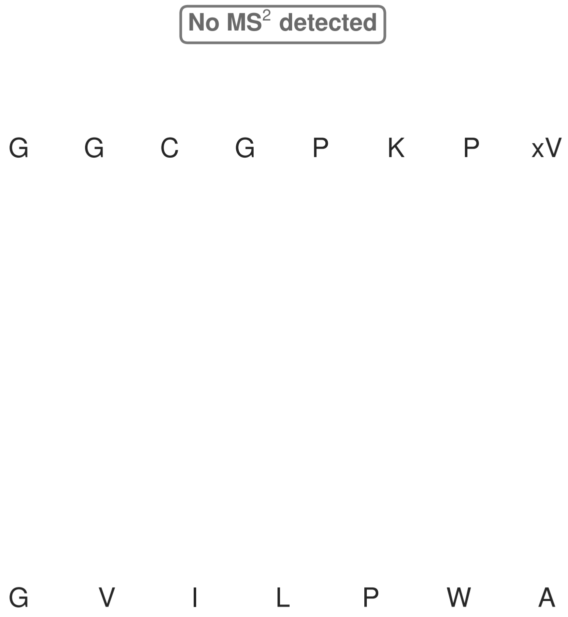


**Figure S66: Modification of the mutant_6 precursor peptide by PlpXY.** LC-MS analysis of IMAC-purified precursor peptides from *E. coli* BL21(DE3)Tuner containing splicease-encoding plasmid TJ-C-559 (PlpXY) and precursor peptide-encoding plasmid mutant_6 (RSTN1-PlpA3-TEV-mutant-6) after expression at 24 °C for 22 h in 1.5 mL cultures after digestion with TEV. Data from n=4 independent biological replicates. Spliced amino acids have an x as a prefix. Spliced refers to a single tyramine excision. A representative experimental replicate is shown in panels **A, B.**

**A** Close-up plot of the representative extracted ion chromatogram (EIC) of the various peptide states; the blue trace corresponds to the Unmodified peptide; the orange trace corresponds to the Spliced peptide. **B** Close-up plot of the m/z values of the peptides detected by the mass spectrometer, with the shaded colored areas corresponding to where the expected m/z values for the peptide m/z traces should occur, with the observed spectrum plotted in black. The Unmodified, Spliced peptide(s) were found in the mass spectrum. **C** Bar plot showing the relative abundance of the various peptide states at charge z=2; the bars represent the average relative abundances normalized to total peak areas, with each individual experimental replicate value plotted as a black dot. **D** Mirrored $MS^2$ plot for the Unmodified peptide. In this mirror plot, the full fragmentation spectrum is plotted in black pointing upward, and the matched fragment b ions (blue) and y ions (red) are plotted pointing downward, mirrored against it, so that each matched fragment can be compared directly with the measured spectrum. **E** $MS^2$ spectrum of the fragments corresponding to the Unmodified peptide sequence, with matched b (blue, above) and y ions (red, below) plotted. **F** Mirrored $MS^2$ plot for the Spliced peptide. In this mirror plot, the full fragmentation spectrum is plotted in black pointing upward, and the matched fragment b ions (blue) and y ions (red) are plotted pointing downward, mirrored against it, so that each matched fragment can be compared directly with the measured spectrum. **G** $MS^2$ spectrum of the fragments corresponding to the Spliced peptide sequence, with matched b (blue, above) and y ions (red, below) plotted.

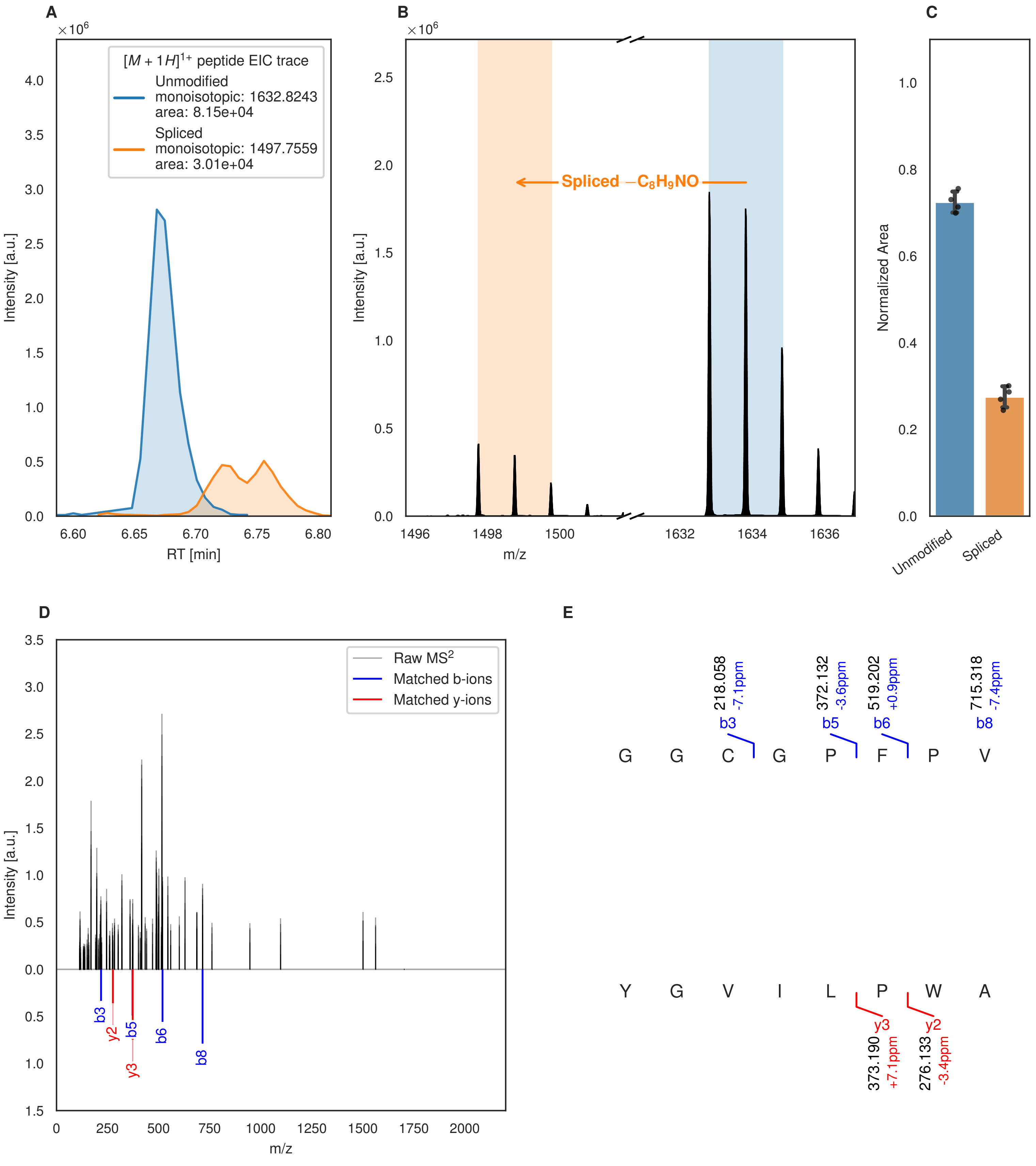
A
B
C
D
E
$\times 10^6$
$[M+1H]^{1+}$ peptide EIC trace
Unmodified
monoisotopic: 1632.8243
area: 8.15e+04
Spliced
monoisotopic: 1497.7559
area: 3.01e+04
Intensity [a.u.]
RT [min]
Spliced $-C_8H_9NO$
m/z
Normalized Area
Unmodified
Spliced
Raw $MS^2$
Matched b-ions
Matched y-ions
b3
y2
b5
y3
b6
b8
218.058 -7.1ppm
372.132 -3.6ppm
519.202 +0.9ppm
715.318 -7.4ppm
G G C G P F P V
Y G V I L P W A
373.190 +7.1ppm
276.133 -3.4ppm

Figure S67 (figure continued from the previous page).

F

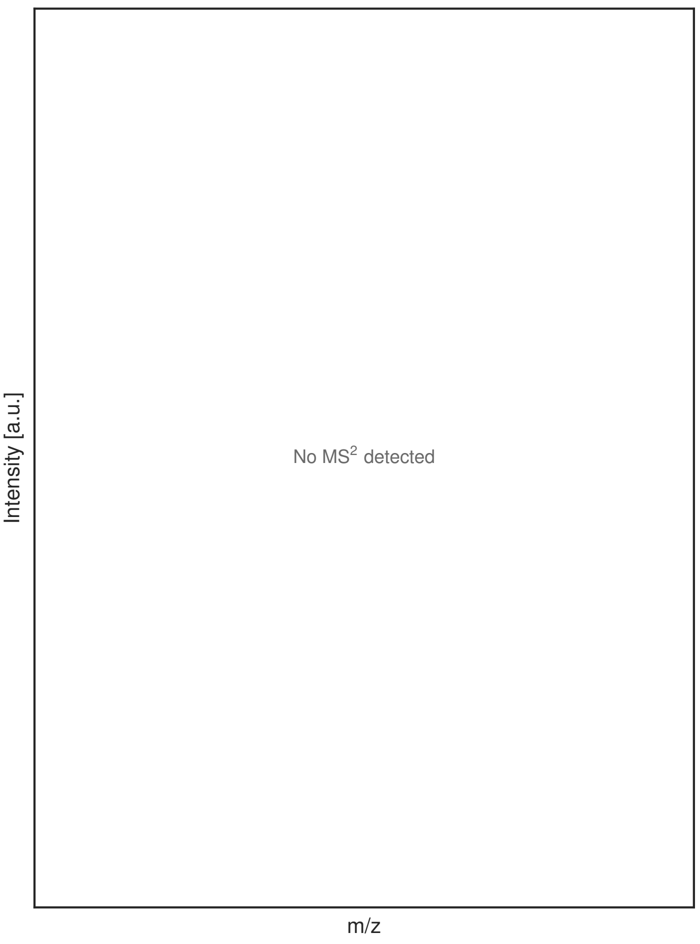


G

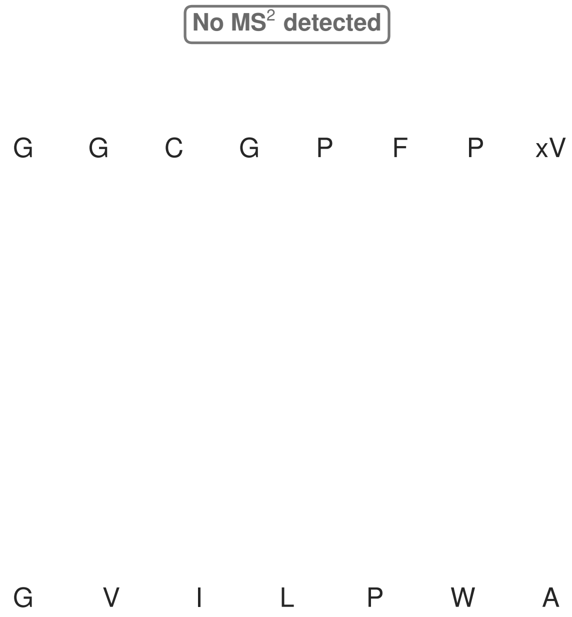


**Figure S67: Modification of the mutant_7 precursor peptide by HypXY.** LC-MS analysis of IMAC-purified precursor peptides from *E. coli* BL21(DE3)Tuner containing splicease-encoding plasmid TJ-C-689 (HypXY) and precursor peptide-encoding plasmid mutant_7 (RSTN1-HypAc-TEV-mutant-7) after expression at 24 °C for 22 h in 1.5 mL cultures after digestion with TEV. Data from n=4 independent biological replicates. Spliced amino acids have an x as a prefix. Spliced refers to a single tyramine excision. A representative experimental replicate is shown in panels **A, B.**

**A** Close-up plot of the representative extracted ion chromatogram (EIC) of the various peptide states; the blue trace corresponds to the Unmodified peptide; the orange trace corresponds to the Spliced peptide. **B** Close-up plot of the m/z values of the peptides detected by the mass spectrometer, with the shaded colored areas corresponding to where the expected m/z values for the peptide m/z traces should occur, with the observed spectrum plotted in black. The Unmodified, Spliced peptide(s) were found in the mass spectrum. **C** Bar plot showing the relative abundance of the various peptide states at charge z=1; the bars represent the average relative abundances normalized to total peak areas, with each individual experimental replicate value plotted as a black dot. **D** Mirrored MS$^2$ plot for the Unmodified peptide. In this mirror plot, the full fragmentation spectrum is plotted in black pointing upward, and the matched fragment b ions (blue) and y ions (red) are plotted pointing downward, mirrored against it, so that each matched fragment can be compared directly with the measured spectrum. **E** MS$^2$ spectrum of the fragments corresponding to the Unmodified peptide sequence, with matched b (blue, above) and y ions (red, below) plotted. **F** Mirrored MS$^2$ plot for the Spliced peptide. In this mirror plot, the full fragmentation spectrum is plotted in black pointing upward, and the matched fragment b ions (blue) and y ions (red) are plotted pointing downward, mirrored against it, so that each matched fragment can be compared directly with the measured spectrum. **G** MS$^2$ spectrum of the fragments corresponding to the Spliced peptide sequence, with matched b (blue, above) and y ions (red, below) plotted.

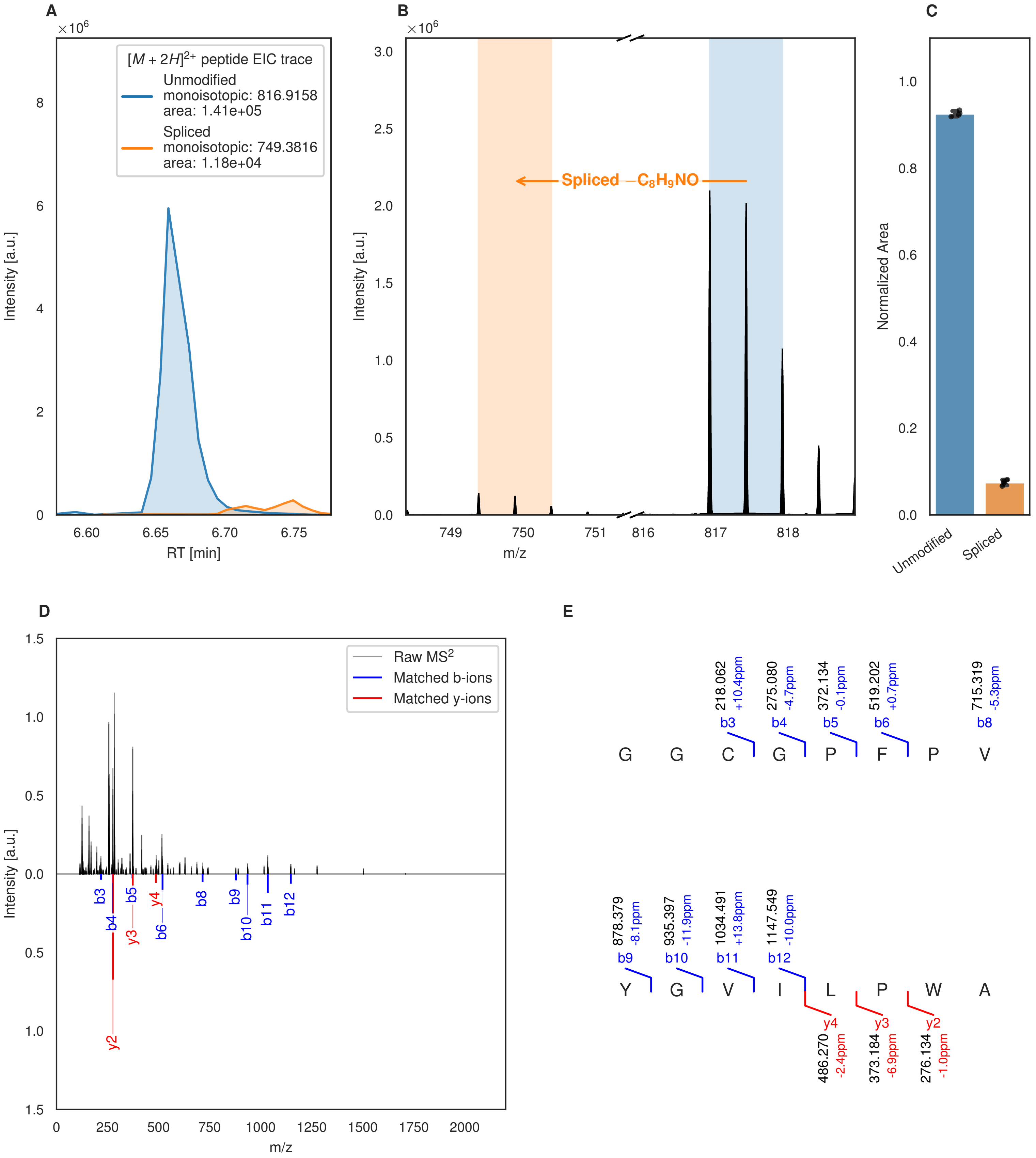
A
[M + 2H]2+ peptide EIC trace
Unmodified
monoisotopic: 816.9158
area: 1.41e+05
Spliced
monoisotopic: 749.3816
area: 1.18e+04
Intensity [a.u.]
RT [min]
B
Spliced −C8H9NO
Intensity [a.u.]
m/z
C
Normalized Area
Unmodified
Spliced
D
Raw MS2
Matched b-ions
Matched y-ions
Intensity [a.u.]
m/z
E
218.062 +10.4ppm b3
275.080 -4.7ppm b4
372.134 -0.1ppm b5
519.202 +0.7ppm b6
715.319 -5.3ppm b8
G G C G P F P V
878.379 -8.1ppm b9
935.397 -11.9ppm b10
1034.491 +13.8ppm b11
1147.549 -10.0ppm b12
Y G V I L P W A
y4 486.270 -2.4ppm
y3 373.184 -6.9ppm
y2 276.134 -1.0ppm

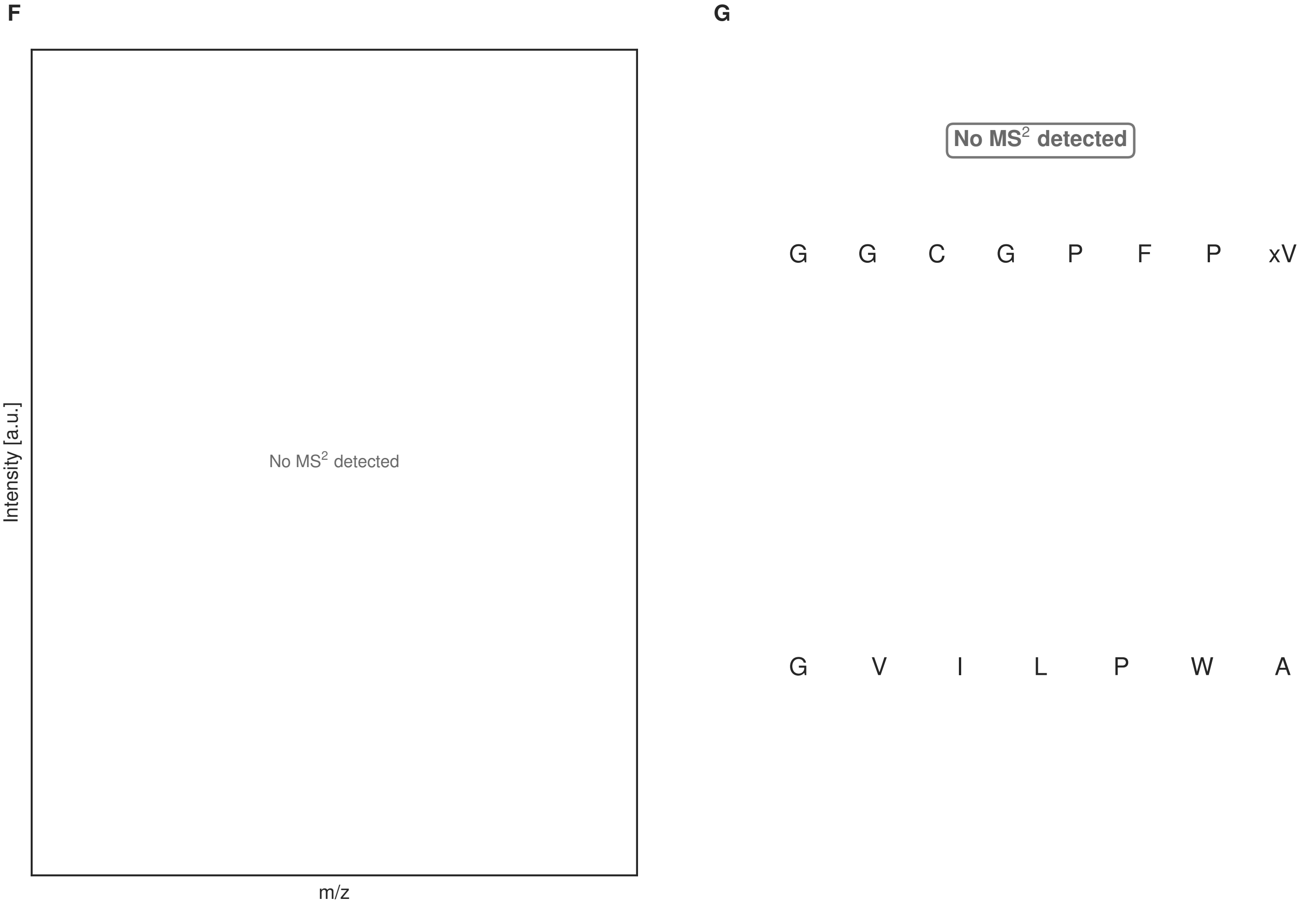


**Figure S68: Modification of the mutant_7 precursor peptide by PlpXY.** LC-MS analysis of IMAC-purified precursor peptides from *E. coli* BL21(DE3)Tuner containing splicease-encoding plasmid TJ-C-559 (PlpXY) and precursor peptide-encoding plasmid mutant_7 (RSTN1-PlpA3-TEV-mutant-7) after expression at 24 °C for 22 h in 1.5 mL cultures after digestion with TEV. Data from n=4 independent biological replicates. Spliced amino acids have an x as a prefix. Spliced refers to a single tyramine excision. A representative experimental replicate is shown in panels **A, B.**

**A** Close-up plot of the representative extracted ion chromatogram (EIC) of the various peptide states; the blue trace corresponds to the Unmodified peptide; the orange trace corresponds to the Spliced peptide. **B** Close-up plot of the m/z values of the peptides detected by the mass spectrometer, with the shaded colored areas corresponding to where the expected m/z values for the peptide m/z traces should occur, with the observed spectrum plotted in black. The Unmodified, Spliced peptide(s) were found in the mass spectrum. **C** Bar plot showing the relative abundance of the various peptide states at charge z=2; the bars represent the average relative abundances normalized to total peak areas, with each individual experimental replicate value plotted as a black dot. **D** Mirrored $MS^2$ plot for the Unmodified peptide. In this mirror plot, the full fragmentation spectrum is plotted in black pointing upward, and the matched fragment b ions (blue) and y ions (red) are plotted pointing downward, mirrored against it, so that each matched fragment can be compared directly with the measured spectrum. **E** $MS^2$ spectrum of the fragments corresponding to the Unmodified peptide sequence, with matched b (blue, above) and y ions (red, below) plotted. **F** Mirrored $MS^2$ plot for the Spliced peptide. In this mirror plot, the full fragmentation spectrum is plotted in black pointing upward, and the matched fragment b ions (blue) and y ions (red) are plotted pointing downward, mirrored against it, so that each matched fragment can be compared directly with the measured spectrum. **G** $MS^2$ spectrum of the fragments corresponding to the Spliced peptide sequence, with matched b (blue, above) and y ions (red, below) plotted.

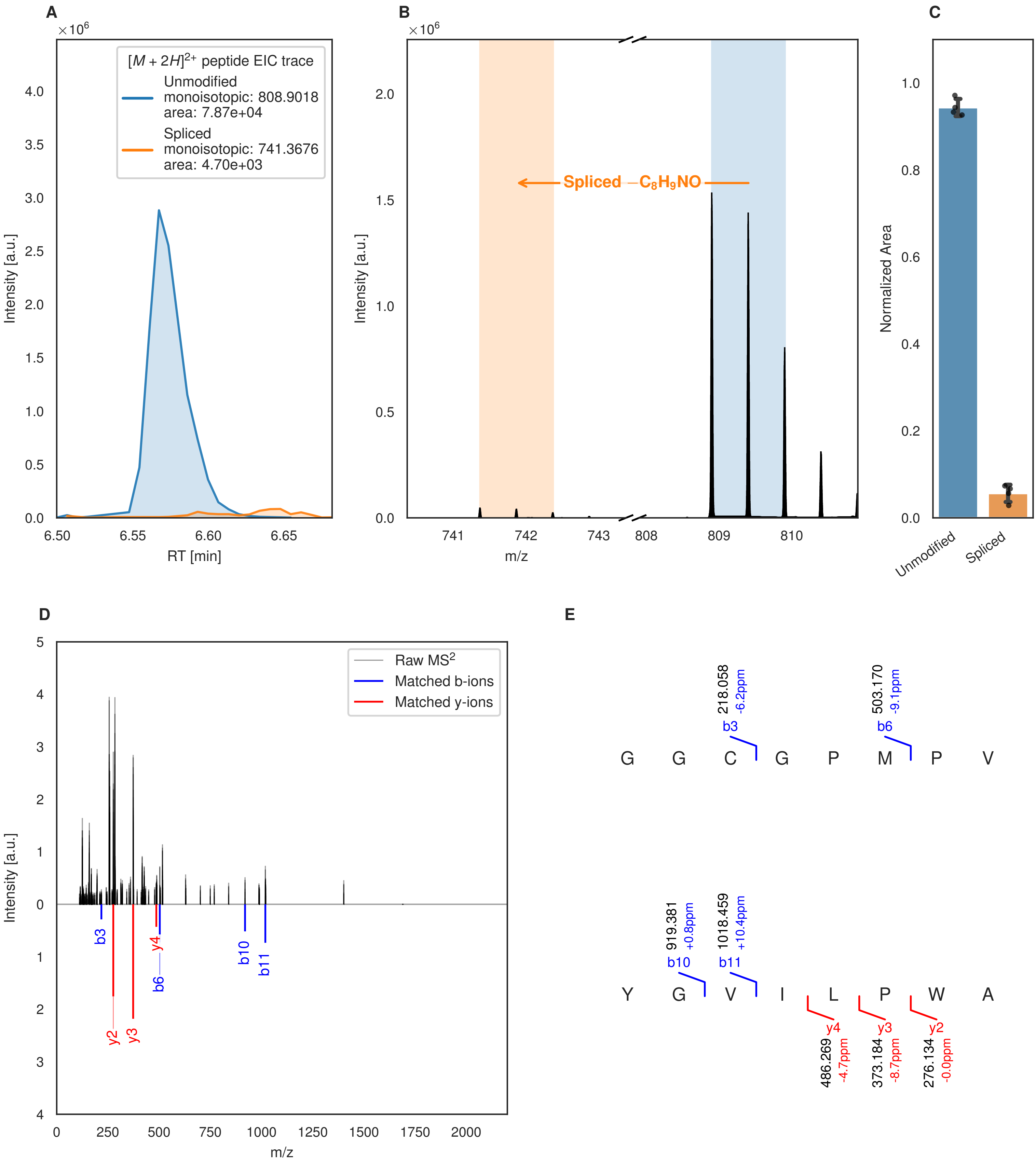
A
$\times10^6$
$[M+2H]^{2+}$ peptide EIC trace
Unmodified
monoisotopic: 808.9018
area: 7.87e+04
Spliced
monoisotopic: 741.3676
area: 4.70e+03
Intensity [a.u.]
RT [min]
B
$\times10^6$
Spliced $-C_8H_9NO$
Intensity [a.u.]
m/z
C
Normalized Area
Unmodified
Spliced
D
Raw MS$^2$
Matched b-ions
Matched y-ions
Intensity [a.u.]
m/z
b3
b6
b10
b11
y2
y3
y4
E
G G C G P M P V
218.058
-6.2ppm
b3
503.170
-9.1ppm
b6
Y G V I L P W A
919.381
+0.8ppm
b10
1018.459
+10.4ppm
b11
y4
486.269
-4.7ppm
y3
373.184
-8.7ppm
y2
276.134
-0.0ppm

F

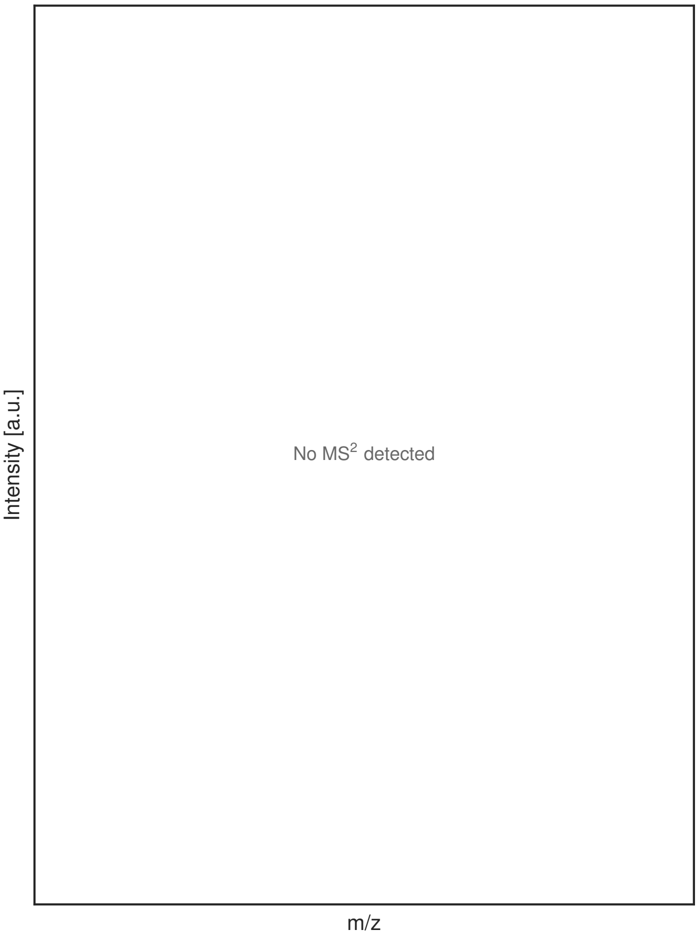


G

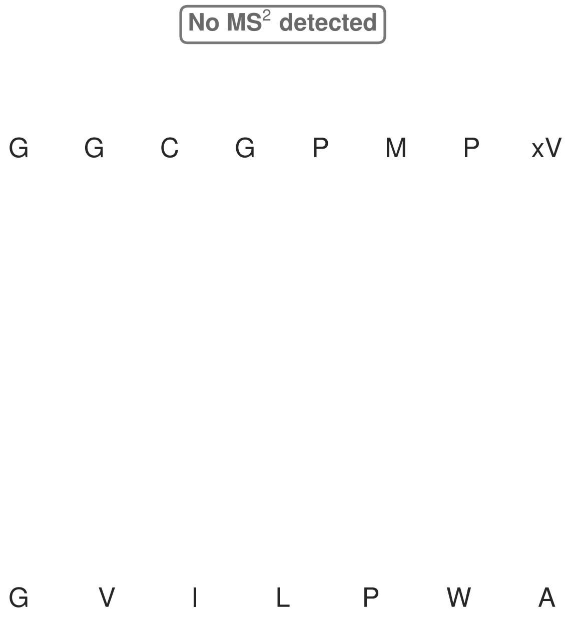


**Figure S69: Modification of the mutant_8 precursor peptide by HypXY.** LC-MS analysis of IMAC-purified precursor peptides from *E. coli* BL21(DE3)Tuner containing splicease-encoding plasmid TJ-C-689 (HypXY) and precursor peptide-encoding plasmid mutant_8 (RSTN1-HypAc-TEV-mutant-8) after expression at 24 °C for 22 h in 1.5 mL cultures after digestion with TEV. Data from n=4 independent biological replicates. Spliced amino acids have an x as a prefix. Spliced refers to a single tyramine excision. A representative experimental replicate is shown in panels **A, B.**

**A** Close-up plot of the representative extracted ion chromatogram (EIC) of the various peptide states; the blue trace corresponds to the Unmodified peptide; the orange trace corresponds to the Spliced peptide. **B** Close-up plot of the m/z values of the peptides detected by the mass spectrometer, with the shaded colored areas corresponding to where the expected m/z values for the peptide m/z traces should occur, with the observed spectrum plotted in black. The Unmodified, Spliced peptide(s) were found in the mass spectrum. **C** Bar plot showing the relative abundance of the various peptide states at charge z=2; the bars represent the average relative abundances normalized to total peak areas, with each individual experimental replicate value plotted as a black dot. **D** Mirrored MS$^2$ plot for the Unmodified peptide. In this mirror plot, the full fragmentation spectrum is plotted in black pointing upward, and the matched fragment b ions (blue) and y ions (red) are plotted pointing downward, mirrored against it, so that each matched fragment can be compared directly with the measured spectrum. **E** MS$^2$ spectrum of the fragments corresponding to the Unmodified peptide sequence, with matched b (blue, above) and y ions (red, below) plotted. **F** Mirrored MS$^2$ plot for the Spliced peptide. In this mirror plot, the full fragmentation spectrum is plotted in black pointing upward, and the matched fragment b ions (blue) and y ions (red) are plotted pointing downward, mirrored against it, so that each matched fragment can be compared directly with the measured spectrum. **G** MS$^2$ spectrum of the fragments corresponding to the Spliced peptide sequence, with matched b (blue, above) and y ions (red, below) plotted.

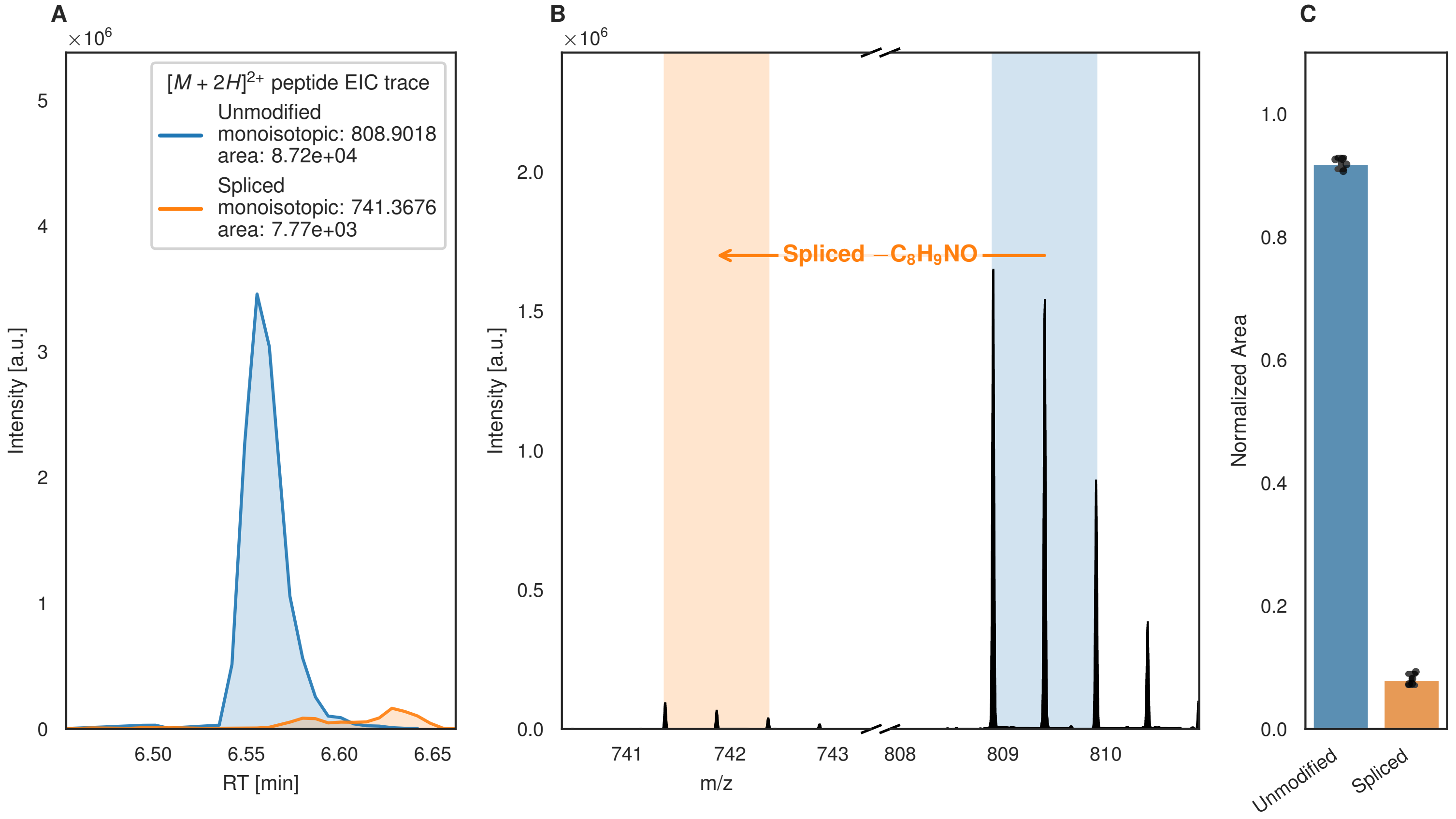
A
B
C
$[M+2H]^{2+}$ peptide EIC trace
Unmodified
monoisotopic: 808.9018
area: 8.72e+04
Spliced
monoisotopic: 741.3676
area: 7.77e+03
Intensity [a.u.]
RT [min]
Spliced −C8H9NO
m/z
Normalized Area
Unmodified
Spliced

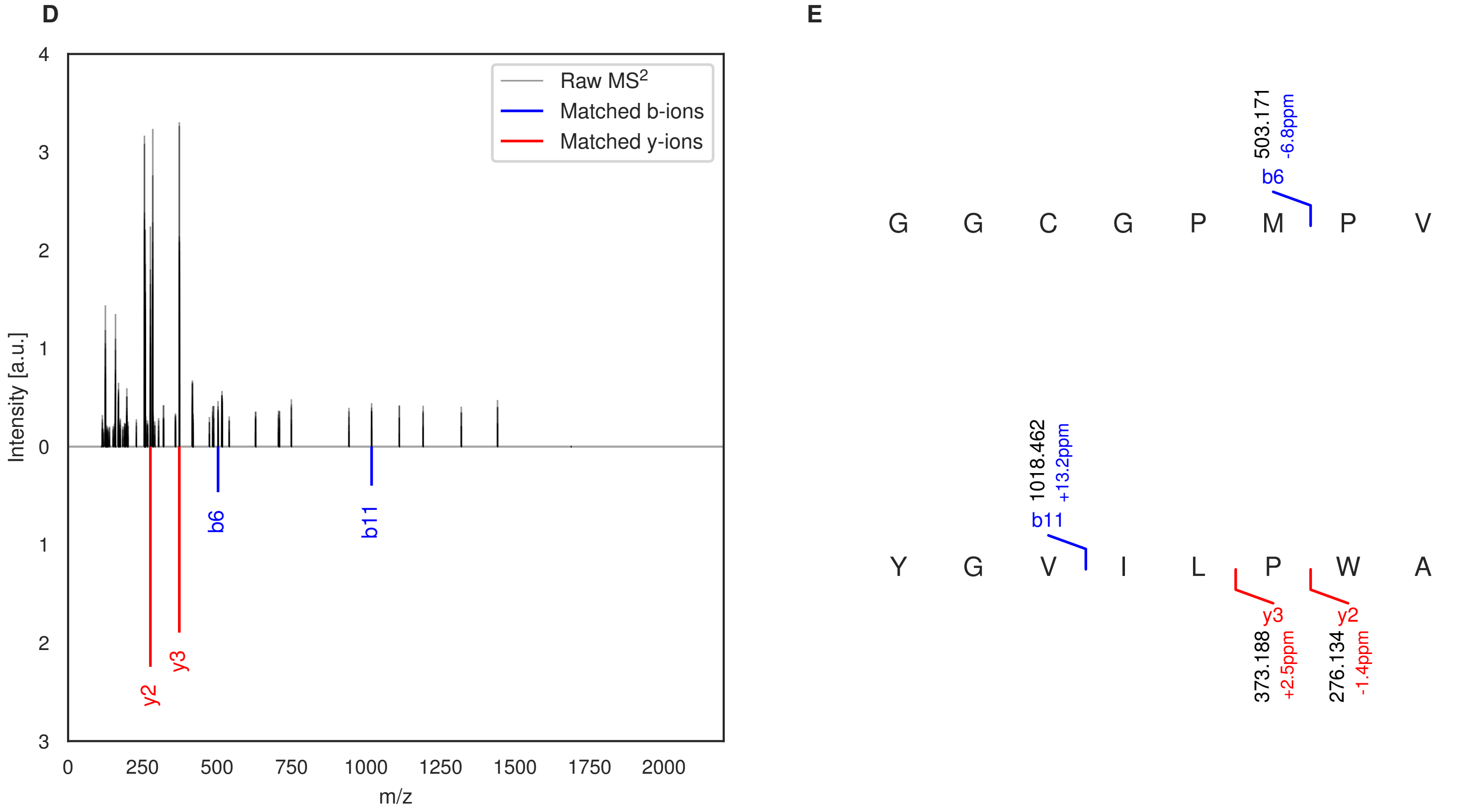
D
E
Raw MS2
Matched b-ions
Matched y-ions
Intensity [a.u.]
m/z
b6
b11
y2
y3
G G C G P M P V
503.171
-6.8ppm
Y G V I L P W A
1018.462
+13.2ppm
373.188
+2.5ppm
276.134
-1.4ppm

Figure S70 (figure continued from the previous page).

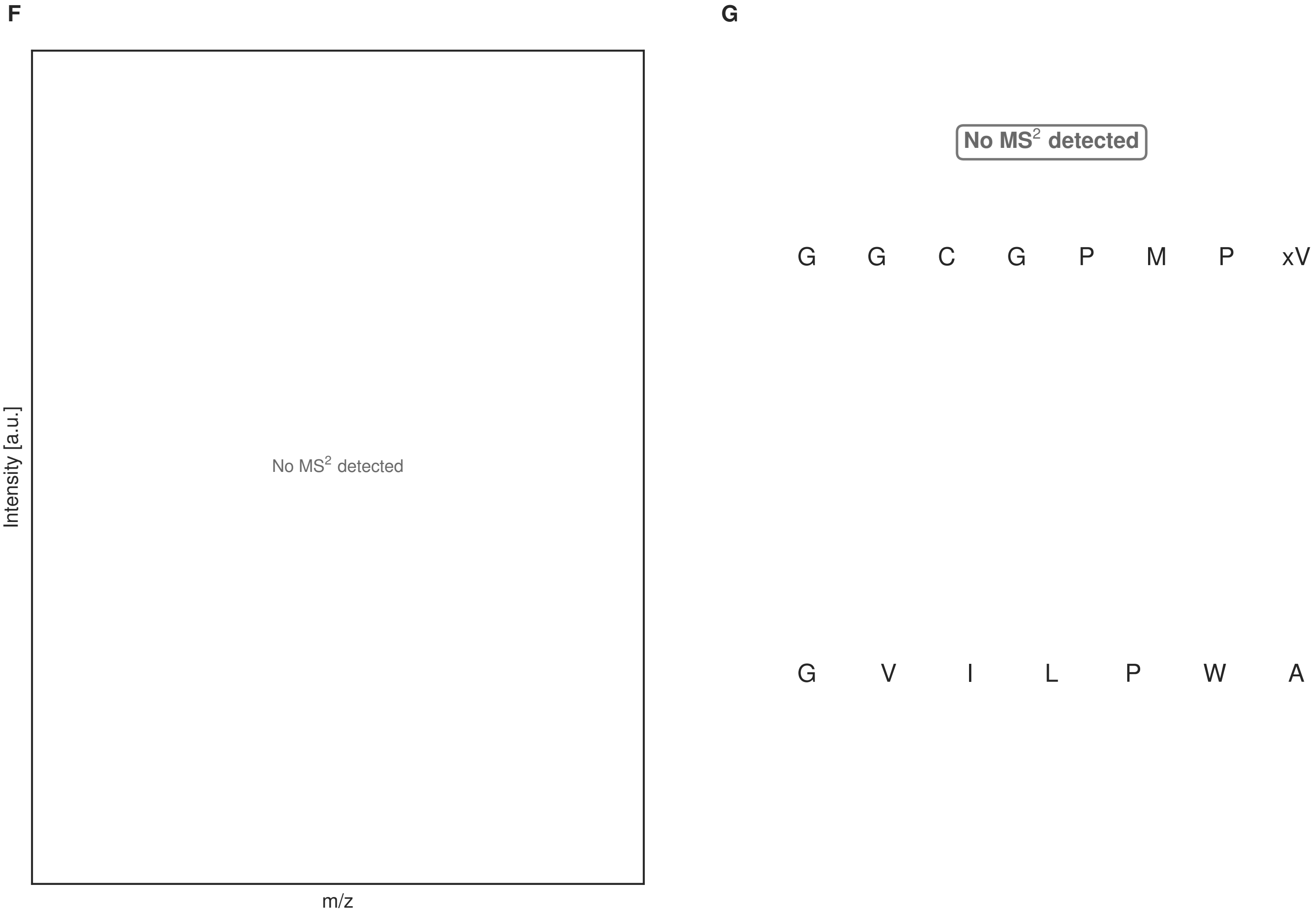


**Figure S70: Modification of the mutant_8 precursor peptide by PlpXY.** LC-MS analysis of IMAC-purified precursor peptides from *E. coli* BL21(DE3)Tuner containing splicease-encoding plasmid TJ-C-559 (PlpXY) and precursor peptide-encoding plasmid mutant_8 (RSTN1-PlpA3-TEV-mutant-8) after expression at 24 °C for 22 h in 1.5 mL cultures after digestion with TEV. Data from n=4 independent biological replicates. Spliced amino acids have an x as a prefix. Spliced refers to a single tyramine excision. A representative experimental replicate is shown in panels **A, B.**

**A** Close-up plot of the representative extracted ion chromatogram (EIC) of the various peptide states; the blue trace corresponds to the Unmodified peptide; the orange trace corresponds to the Spliced peptide. **B** Close-up plot of the m/z values of the peptides detected by the mass spectrometer, with the shaded colored areas corresponding to where the expected m/z values for the peptide m/z traces should occur, with the observed spectrum plotted in black. The Unmodified, Spliced peptide(s) were found in the mass spectrum. **C** Bar plot showing the relative abundance of the various peptide states at charge z=2; the bars represent the average relative abundances normalized to total peak areas, with each individual experimental replicate value plotted as a black dot. **D** Mirrored MS$^2$ plot for the Unmodified peptide. In this mirror plot, the full fragmentation spectrum is plotted in black pointing upward, and the matched fragment b ions (blue) and y ions (red) are plotted pointing downward, mirrored against it, so that each matched fragment can be compared directly with the measured spectrum. **E** MS$^2$ spectrum of the fragments corresponding to the Unmodified peptide sequence, with matched b (blue, above) and y ions (red, below) plotted. **F** Mirrored MS$^2$ plot for the Spliced peptide. In this mirror plot, the full fragmentation spectrum is plotted in black pointing upward, and the matched fragment b ions (blue) and y ions (red) are plotted pointing downward, mirrored against it, so that each matched fragment can be compared directly with the measured spectrum. **G** MS$^2$ spectrum of the fragments corresponding to the Spliced peptide sequence, with matched b (blue, above) and y ions (red, below) plotted.

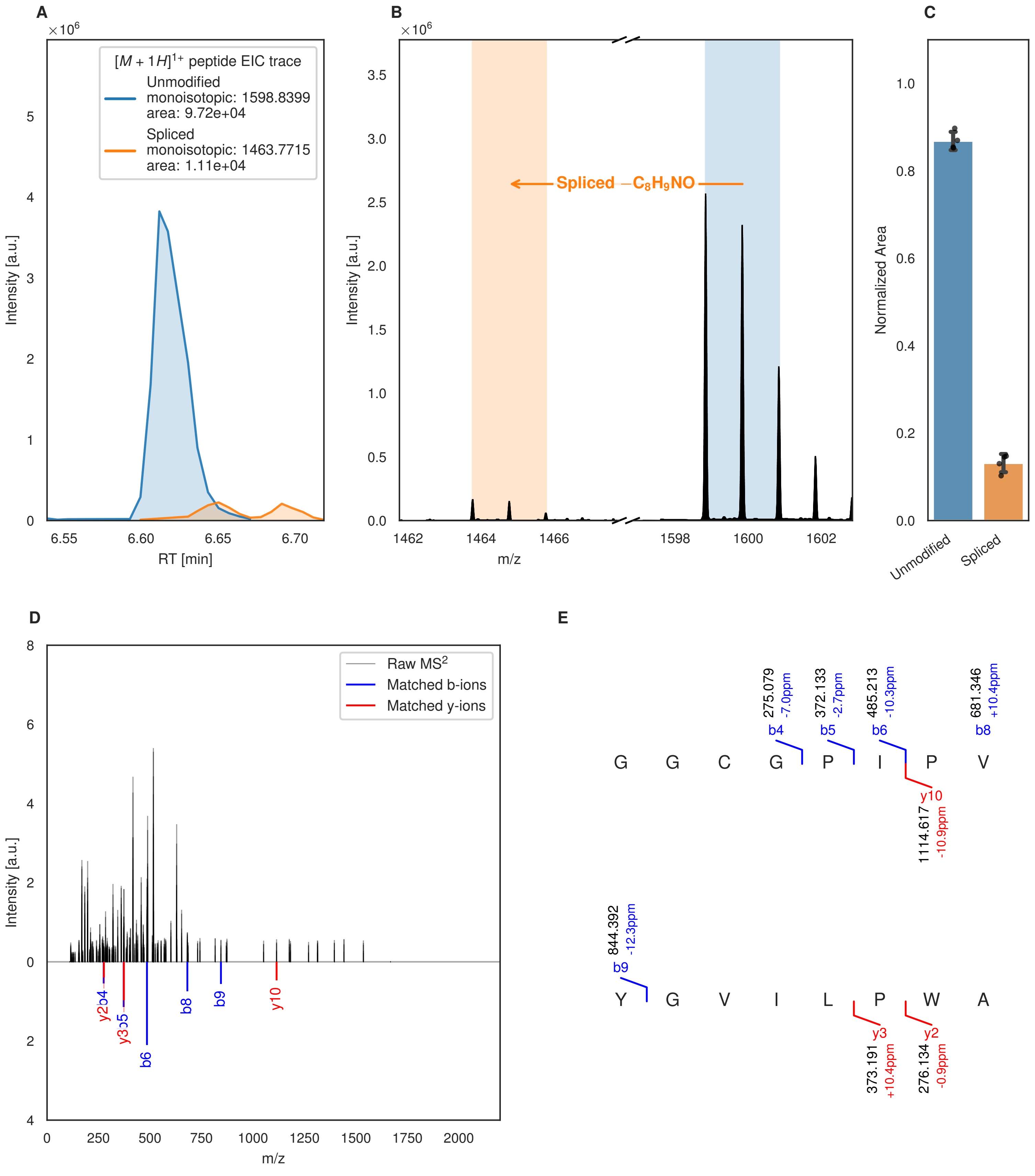

A
$\times 10^6$
$[M+1H]^{1+}$ peptide EIC trace
Unmodified
monoisotopic: 1598.8399
area: 9.72e+04
Spliced
monoisotopic: 1463.7715
area: 1.11e+04
Intensity [a.u.]
RT [min]
B
Spliced $-C_8H_9NO$
Intensity [a.u.]
m/z
C
Normalized Area
Unmodified
Spliced
D
Raw MS$^2$
Matched b-ions
Matched y-ions
Intensity [a.u.]
m/z
E
275.079 -7.0ppm b4
372.133 -2.7ppm b5
485.213 -10.3ppm b6
681.346 +10.4ppm b8
G G C G P I P V
y10 1114.617 -10.9ppm
844.392 -12.3ppm b9
Y G V I L P W A
y3 373.191 +10.4ppm
y2 276.134 -0.9ppm

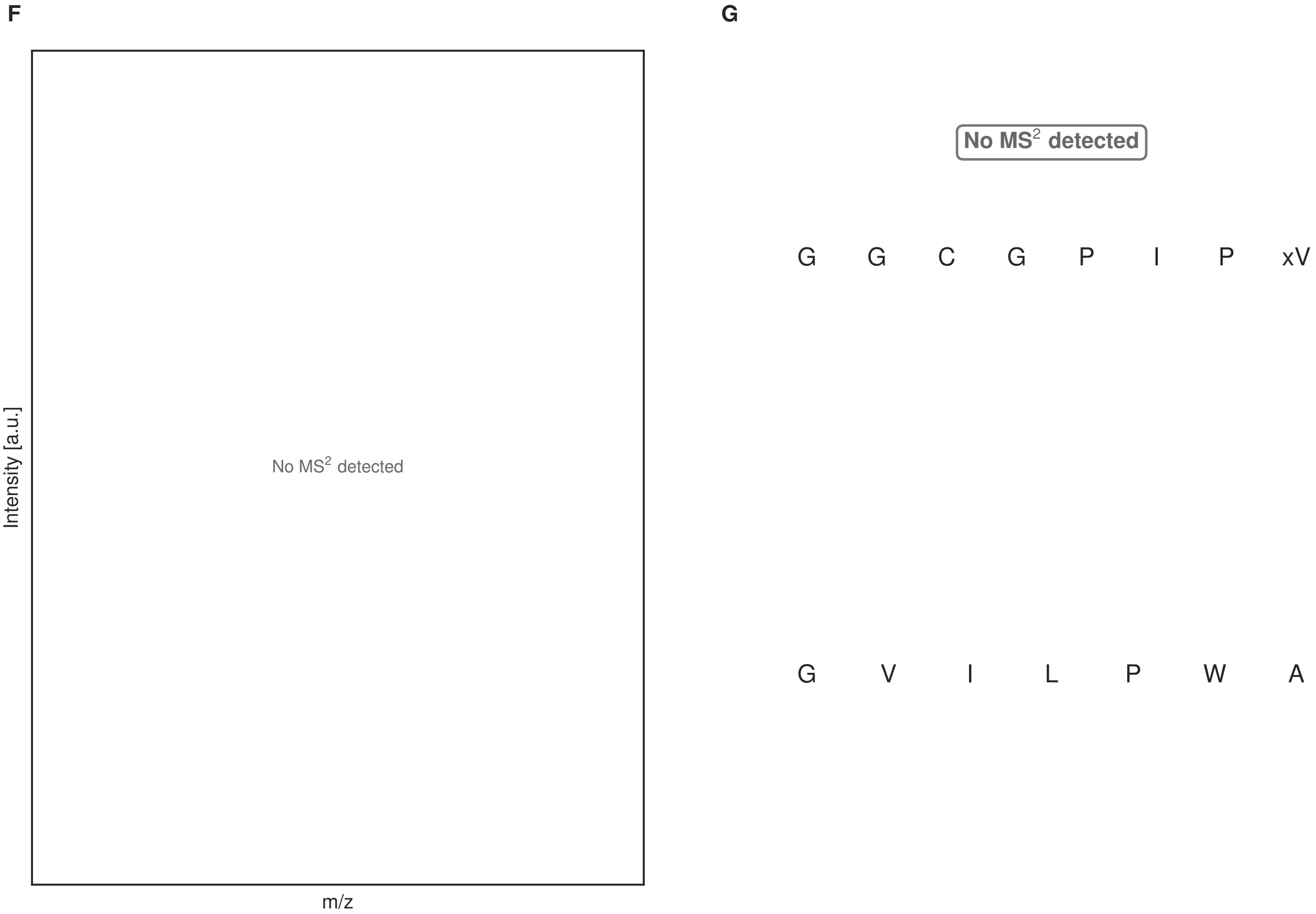


**Figure S71: Modification of the mutant_9 precursor peptide by HypXY.** LC-MS analysis of IMAC-purified precursor peptides from *E. coli* BL21(DE3)Tuner containing splicease-encoding plasmid TJ-C-689 (HypXY) and precursor peptide-encoding plasmid mutant_9 (RSTN1-HypAc-TEV-mutant-9) after expression at 24 °C for 22 h in 1.5 mL cultures after digestion with TEV. Data from n=4 independent biological replicates. Spliced amino acids have an x as a prefix. Spliced refers to a single tyramine excision. A representative experimental replicate is shown in panels **A, B.**

**A** Close-up plot of the representative extracted ion chromatogram (EIC) of the various peptide states; the blue trace corresponds to the Unmodified peptide; the orange trace corresponds to the Spliced peptide. **B** Close-up plot of the m/z values of the peptides detected by the mass spectrometer, with the shaded colored areas corresponding to where the expected m/z values for the peptide m/z traces should occur, with the observed spectrum plotted in black. The Unmodified, Spliced peptide(s) were found in the mass spectrum. **C** Bar plot showing the relative abundance of the various peptide states at charge z=1; the bars represent the average relative abundances normalized to total peak areas, with each individual experimental replicate value plotted as a black dot. **D** Mirrored $MS^2$ plot for the Unmodified peptide. In this mirror plot, the full fragmentation spectrum is plotted in black pointing upward, and the matched fragment b ions (blue) and y ions (red) are plotted pointing downward, mirrored against it, so that each matched fragment can be compared directly with the measured spectrum. **E** $MS^2$ spectrum of the fragments corresponding to the Unmodified peptide sequence, with matched b (blue, above) and y ions (red, below) plotted. **F** Mirrored $MS^2$ plot for the Spliced peptide. In this mirror plot, the full fragmentation spectrum is plotted in black pointing upward, and the matched fragment b ions (blue) and y ions (red) are plotted pointing downward, mirrored against it, so that each matched fragment can be compared directly with the measured spectrum. **G** $MS^2$ spectrum of the fragments corresponding to the Spliced peptide sequence, with matched b (blue, above) and y ions (red, below) plotted.

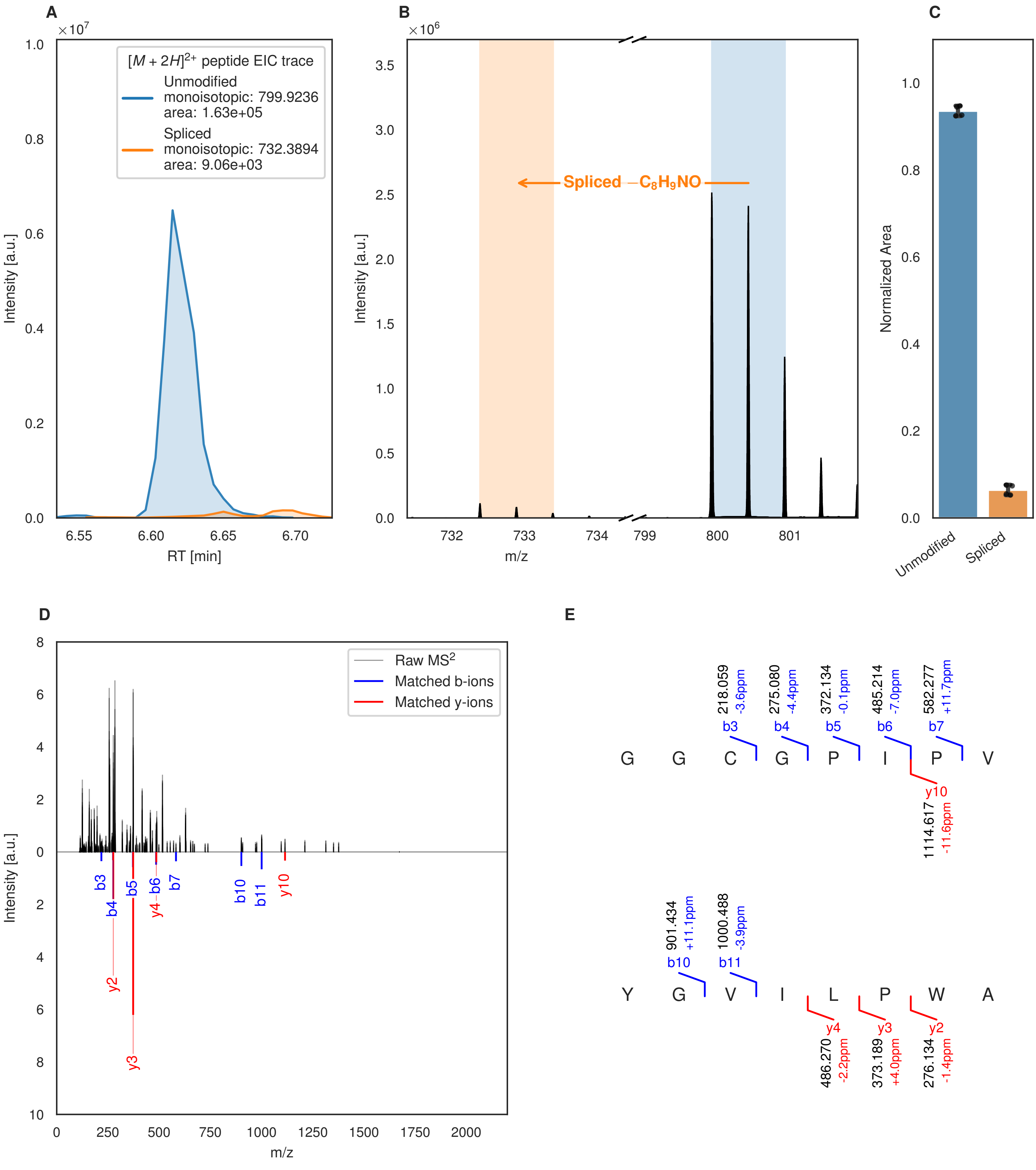
A
×10^7
[M + 2H]2+ peptide EIC trace
Unmodified
monoisotopic: 799.9236
area: 1.63e+05
Spliced
monoisotopic: 732.3894
area: 9.06e+03
Intensity [a.u.]
RT [min]
B
×10^6
Spliced −C8H9NO
Intensity [a.u.]
m/z
C
Normalized Area
Unmodified
Spliced
D
Raw MS2
Matched b-ions
Matched y-ions
Intensity [a.u.]
m/z
b3
b4
b5
y4
b6
b7
b10
b11
y10
y2
y3
E
218.059 -3.6ppm b3
275.080 -4.4ppm b4
372.134 -0.1ppm b5
485.214 -7.0ppm b6
582.277 +11.7ppm b7
G G C G P I P V
y10 1114.617 -11.6ppm
901.434 +11.1ppm b10
1000.488 -3.9ppm b11
Y G V I L P W A
y4 486.270 -2.2ppm
y3 373.189 +4.0ppm
y2 276.134 -1.4ppm

**F**

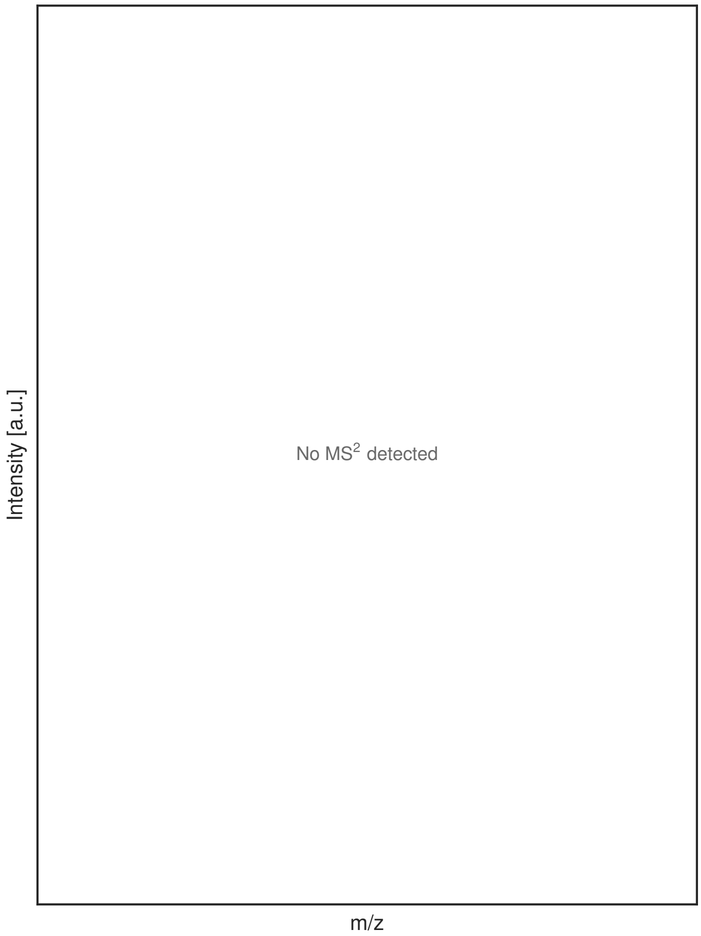


**G**

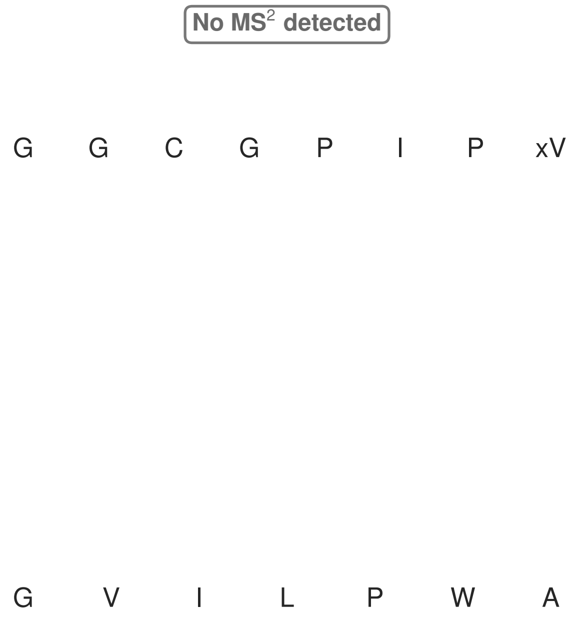


**Figure S72: Modification of the mutant_9 precursor peptide by PlpXY.** LC-MS analysis of IMAC-purified precursor peptides from *E. coli* BL21(DE3)Tuner containing splicease-encoding plasmid TJ-C-559 (PlpXY) and precursor peptide-encoding plasmid mutant_9 (RSTN1-PlpA3-TEV-mutant-9) after expression at 24 °C for 22 h in 1.5 mL cultures after digestion with TEV. Data from n=4 independent biological replicates. Spliced amino acids have an x as a prefix. Spliced refers to a single tyramine excision. A representative experimental replicate is shown in panels **A, B.**

**A** Close-up plot of the representative extracted ion chromatogram (EIC) of the various peptide states; the blue trace corresponds to the Unmodified peptide; the orange trace corresponds to the Spliced peptide. **B** Close-up plot of the m/z values of the peptides detected by the mass spectrometer, with the shaded colored areas corresponding to where the expected m/z values for the peptide m/z traces should occur, with the observed spectrum plotted in black. The Unmodified, Spliced peptide(s) were found in the mass spectrum. **C** Bar plot showing the relative abundance of the various peptide states at charge z=2; the bars represent the average relative abundances normalized to total peak areas, with each individual experimental replicate value plotted as a black dot. **D** Mirrored MS$^2$ plot for the Unmodified peptide. In this mirror plot, the full fragmentation spectrum is plotted in black pointing upward, and the matched fragment b ions (blue) and y ions (red) are plotted pointing downward, mirrored against it, so that each matched fragment can be compared directly with the measured spectrum. **E** MS$^2$ spectrum of the fragments corresponding to the Unmodified peptide sequence, with matched b (blue, above) and y ions (red, below) plotted. **F** Mirrored MS$^2$ plot for the Spliced peptide. In this mirror plot, the full fragmentation spectrum is plotted in black pointing upward, and the matched fragment b ions (blue) and y ions (red) are plotted pointing downward, mirrored against it, so that each matched fragment can be compared directly with the measured spectrum. **G** MS$^2$ spectrum of the fragments corresponding to the Spliced peptide sequence, with matched b (blue, above) and y ions (red, below) plotted.

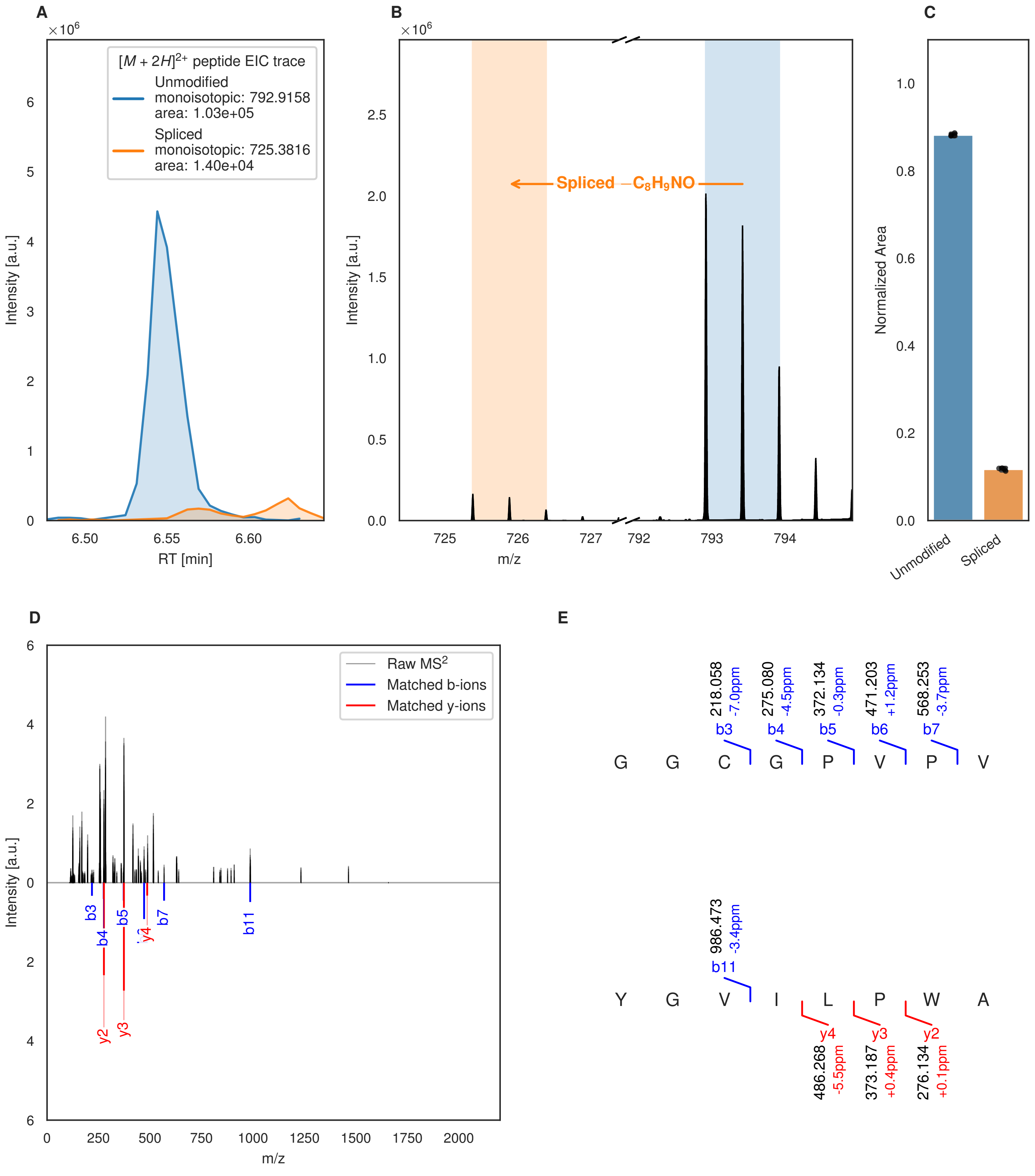
A
B
C
D
E
$[M + 2H]^{2+}$ peptide EIC trace
Unmodified
monoisotopic: 792.9158
area: 1.03e+05
Spliced
monoisotopic: 725.3816
area: 1.40e+04
Intensity [a.u.]
RT [min]
Spliced −$C_8H_9NO$
m/z
Normalized Area
Unmodified
Spliced
Raw $MS^2$
Matched b-ions
Matched y-ions
G G C G P V P V
Y G V I L P W A
218.058 -7.0ppm b3
275.080 -4.5ppm b4
372.134 -0.3ppm b5
471.203 +1.2ppm b6
568.253 -3.7ppm b7
986.473 -3.4ppm b11
y4 486.268 -5.5ppm
y3 373.187 +0.4ppm
y2 276.134 +0.1ppm

Figure S73 (figure continued from the previous page).

F

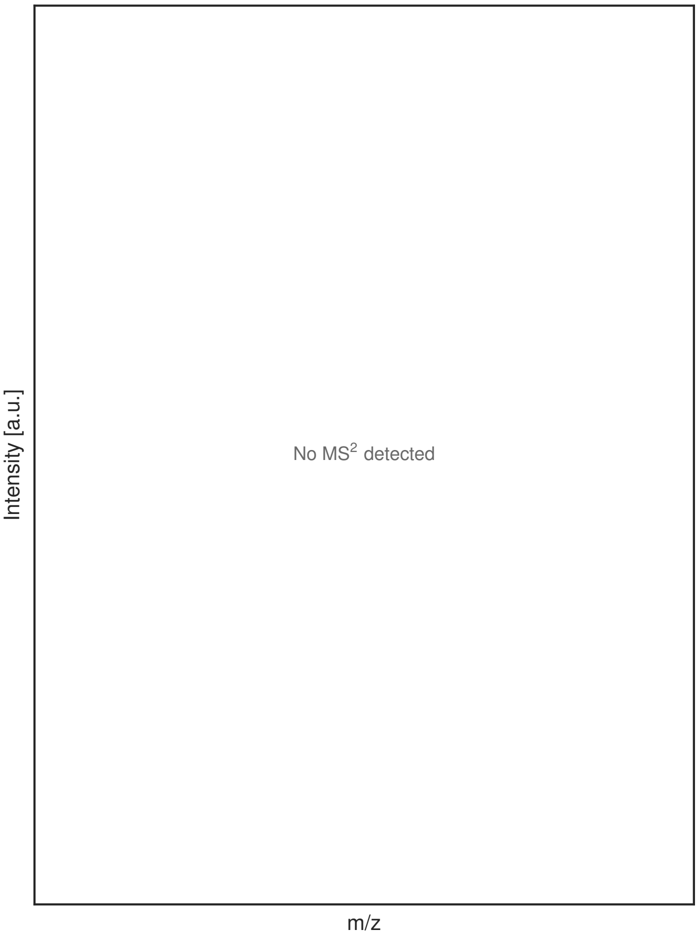


G

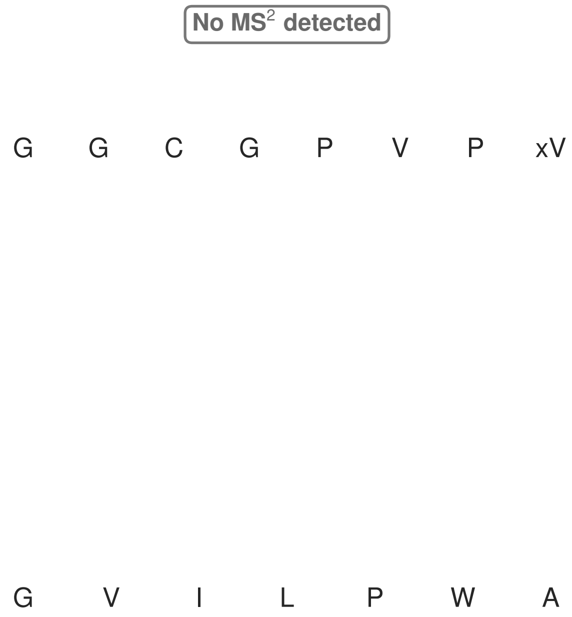


**Figure S73: Modification of the mutant_10 precursor peptide by HypXY.** LC-MS analysis of IMAC-purified precursor peptides from *E. coli* BL21(DE3)Tuner containing splicease-encoding plasmid TJ-C-689 (HypXY) and precursor peptide-encoding plasmid mutant_10 (RSTN1-HypAc-TEV-mutant-10) after expression at 24 °C for 22 h in 1.5 mL cultures after digestion with TEV. Data from n=4 independent biological replicates. Spliced amino acids have an x as a prefix. Spliced refers to a single tyramine excision. A representative experimental replicate is shown in panels **A, B.**

**A** Close-up plot of the representative extracted ion chromatogram (EIC) of the various peptide states; the blue trace corresponds to the Unmodified peptide; the orange trace corresponds to the Spliced peptide. **B** Close-up plot of the m/z values of the peptides detected by the mass spectrometer, with the shaded colored areas corresponding to where the expected m/z values for the peptide m/z traces should occur, with the observed spectrum plotted in black. The Unmodified, Spliced peptide(s) were found in the mass spectrum. **C** Bar plot showing the relative abundance of the various peptide states at charge z=2; the bars represent the average relative abundances normalized to total peak areas, with each individual experimental replicate value plotted as a black dot. **D** Mirrored MS$^2$ plot for the Unmodified peptide. In this mirror plot, the full fragmentation spectrum is plotted in black pointing upward, and the matched fragment b ions (blue) and y ions (red) are plotted pointing downward, mirrored against it, so that each matched fragment can be compared directly with the measured spectrum. **E** MS$^2$ spectrum of the fragments corresponding to the Unmodified peptide sequence, with matched b (blue, above) and y ions (red, below) plotted. **F** Mirrored MS$^2$ plot for the Spliced peptide. In this mirror plot, the full fragmentation spectrum is plotted in black pointing upward, and the matched fragment b ions (blue) and y ions (red) are plotted pointing downward, mirrored against it, so that each matched fragment can be compared directly with the measured spectrum. **G** MS$^2$ spectrum of the fragments corresponding to the Spliced peptide sequence, with matched b (blue, above) and y ions (red, below) plotted.

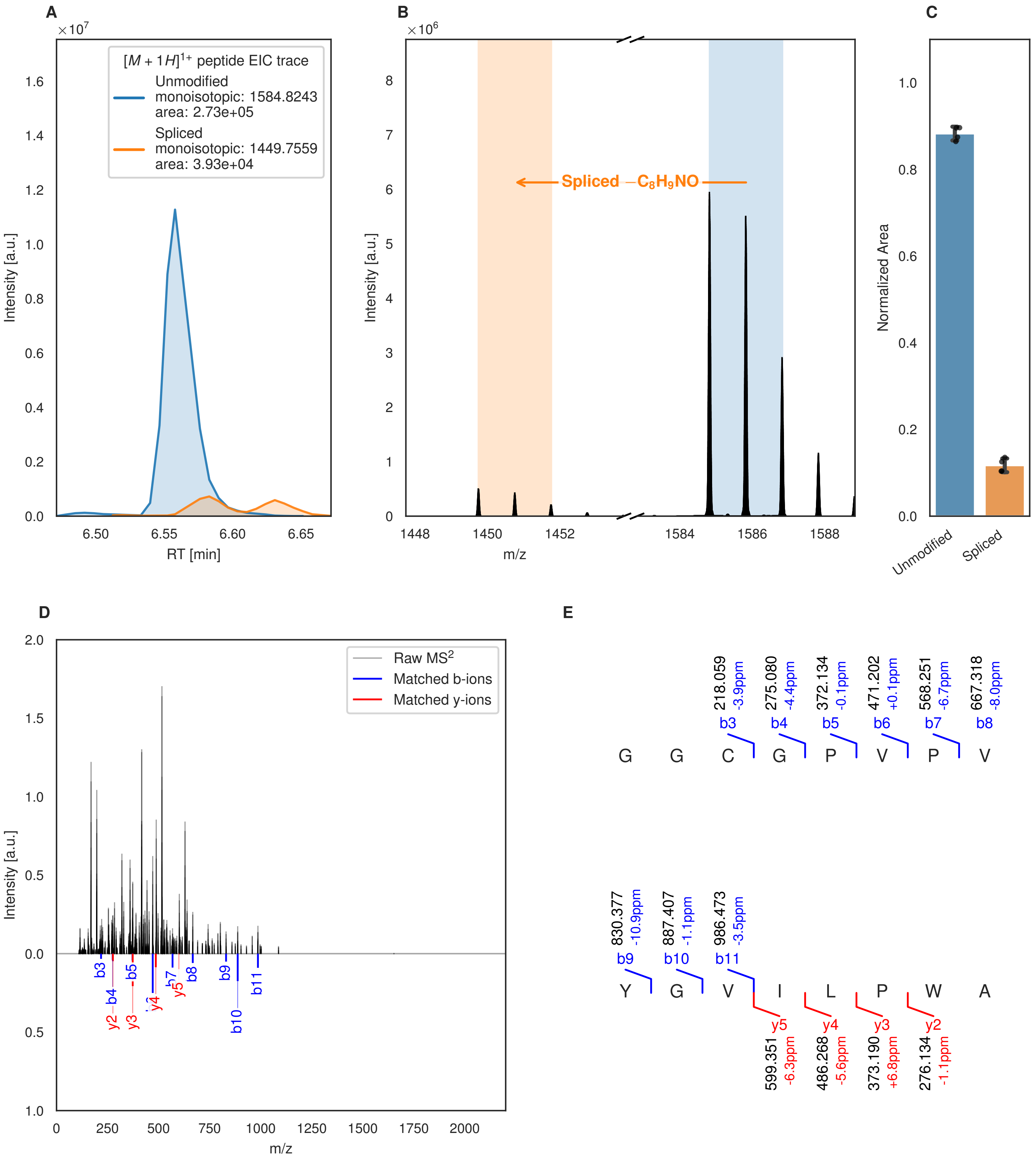
A
[M + 1H]1+ peptide EIC trace
Unmodified
monoisotopic: 1584.8243
area: 2.73e+05
Spliced
monoisotopic: 1449.7559
area: 3.93e+04
Intensity [a.u.]
RT [min]
B
Spliced −C8H9NO
Intensity [a.u.]
m/z
C
Normalized Area
Unmodified
Spliced
D
Raw MS2
Matched b-ions
Matched y-ions
Intensity [a.u.]
m/z
E
G G C G P V P V
b3 218.059 -3.9ppm
b4 275.080 -4.4ppm
b5 372.134 -0.1ppm
b6 471.202 +0.1ppm
b7 568.251 -6.7ppm
b8 667.318 -8.0ppm
Y G V I L P W A
b9 830.377 -10.9ppm
b10 887.407 -1.1ppm
b11 986.473 -3.5ppm
y5 599.351 -6.3ppm
y4 486.268 -5.6ppm
y3 373.190 +6.8ppm
y2 276.134 -1.1ppm

Figure S74 (figure continued from the previous page).

**F**

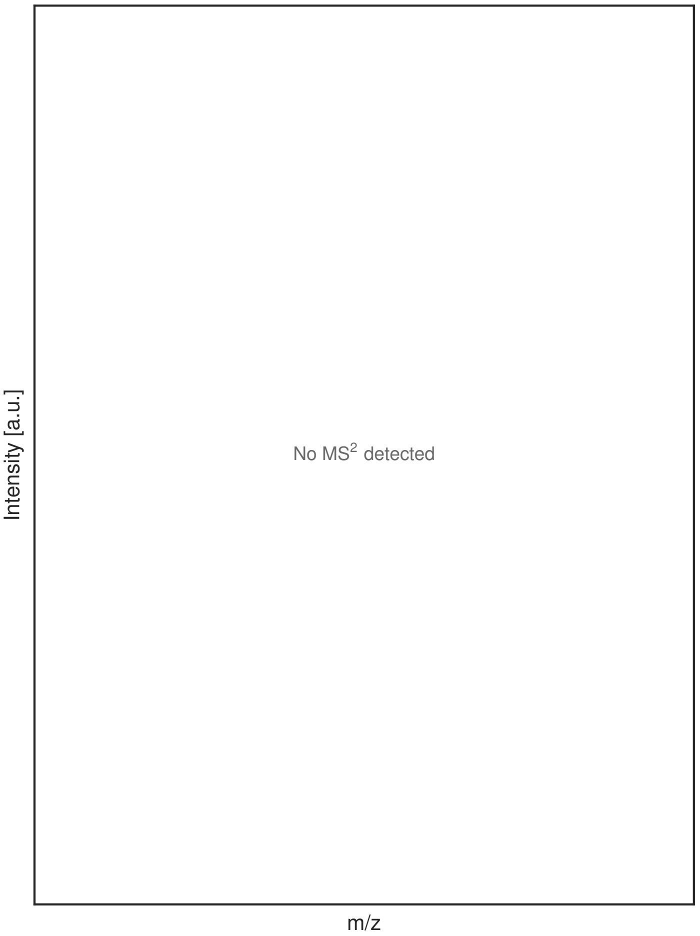


**G**

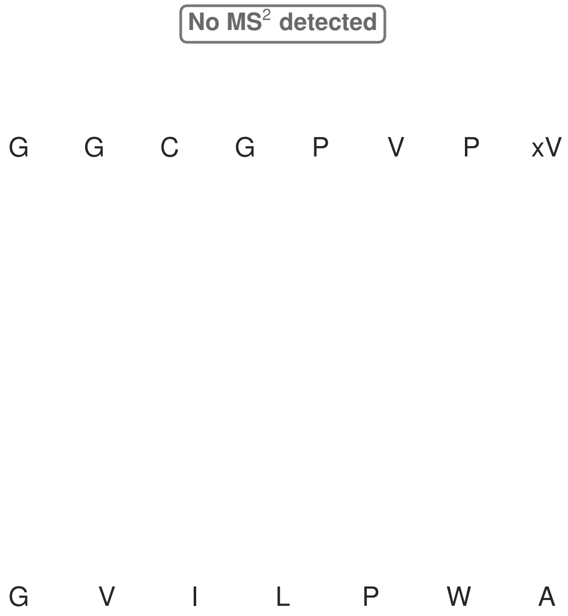


**Figure S74: Modification of the mutant_10 precursor peptide by PlpXY.** LC-MS analysis of IMAC-purified precursor peptides from *E. coli* BL21(DE3)Tuner containing splicease-encoding plasmid TJ-C-559 (PlpXY) and precursor peptide-encoding plasmid mutant_10 (RSTN1-PlpA3-TEV-mutant-10) after expression at 24 °C for 22 h in 1.5 mL cultures after digestion with TEV. Data from n=4 independent biological replicates. Spliced amino acids have an x as a prefix. Spliced refers to a single tyramine excision. A representative experimental replicate is shown in panels **A, B.**

**A** Close-up plot of the representative extracted ion chromatogram (EIC) of the various peptide states; the blue trace corresponds to the Unmodified peptide; the orange trace corresponds to the Spliced peptide. **B** Close-up plot of the m/z values of the peptides detected by the mass spectrometer, with the shaded colored areas corresponding to where the expected m/z values for the peptide m/z traces should occur, with the observed spectrum plotted in black. The Unmodified, Spliced peptide(s) were found in the mass spectrum. **C** Bar plot showing the relative abundance of the various peptide states at charge z=1; the bars represent the average relative abundances normalized to total peak areas, with each individual experimental replicate value plotted as a black dot. **D** Mirrored MS$^2$ plot for the Unmodified peptide. In this mirror plot, the full fragmentation spectrum is plotted in black pointing upward, and the matched fragment b ions (blue) and y ions (red) are plotted pointing downward, mirrored against it, so that each matched fragment can be compared directly with the measured spectrum. **E** MS$^2$ spectrum of the fragments corresponding to the Unmodified peptide sequence, with matched b (blue, above) and y ions (red, below) plotted. **F** Mirrored MS$^2$ plot for the Spliced peptide. In this mirror plot, the full fragmentation spectrum is plotted in black pointing upward, and the matched fragment b ions (blue) and y ions (red) are plotted pointing downward, mirrored against it, so that each matched fragment can be compared directly with the measured spectrum. **G** MS$^2$ spectrum of the fragments corresponding to the Spliced peptide sequence, with matched b (blue, above) and y ions (red, below) plotted.

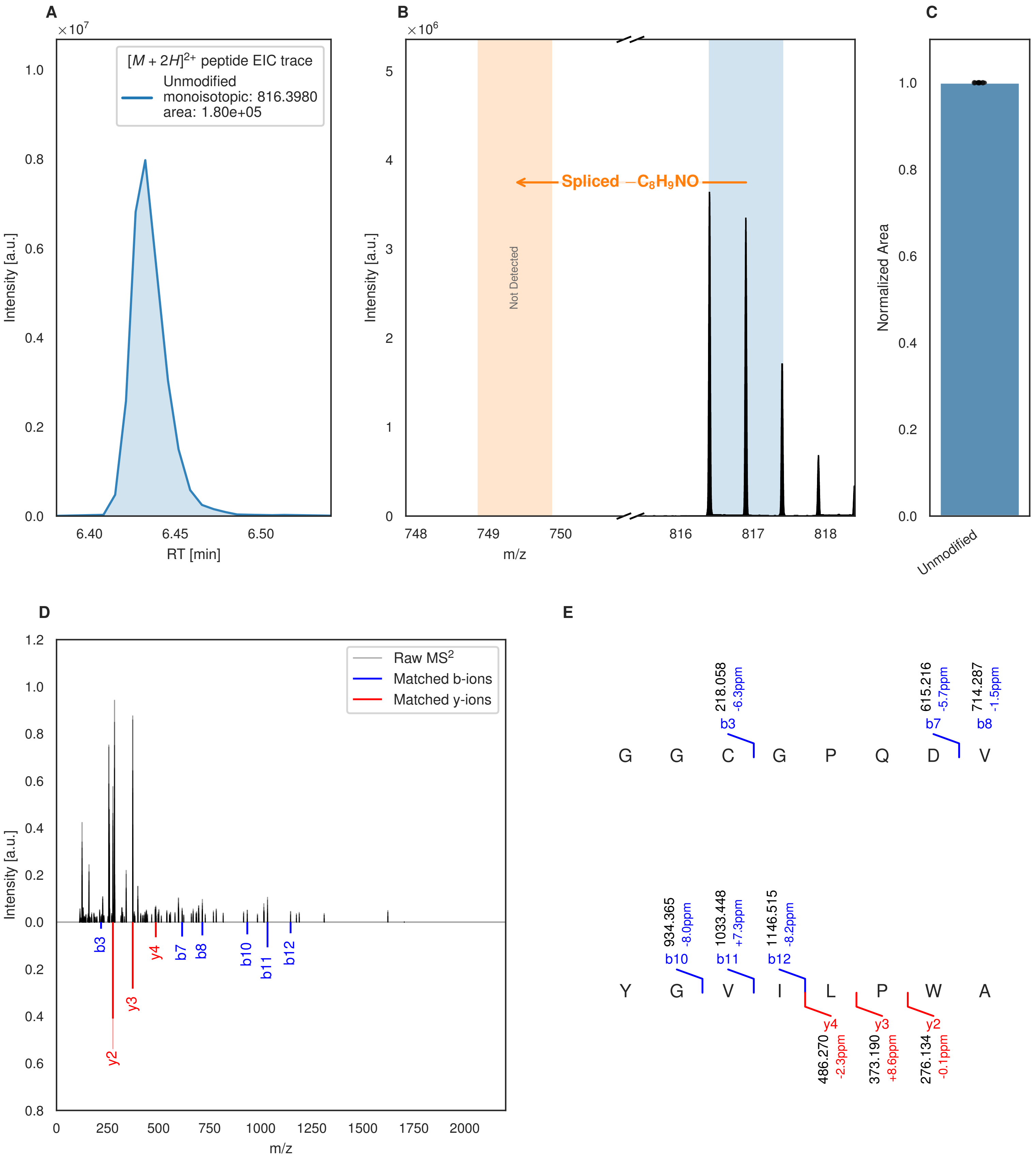


**Figure S75: Modification of the mutant_11 precursor peptide by HypXY.** LC-MS analysis of IMAC-purified precursor peptides from *E. coli* BL21(DE3)Tuner containing splicease-encoding plasmid TJ-C-689 (HypXY) and precursor peptide-encoding plasmid mutant_11 (RSTN1-HypAc-TEV-mutant-11) after expression at 24 °C for 22 h in 1.5 mL cultures after digestion with TEV. Data from n=4 independent biological replicates. Spliced amino acids have an x as a prefix. Spliced refers to a single tyramine excision. A representative experimental replicate is shown in panels **A, B.**

**A** Close-up plot of the representative extracted ion chromatogram (EIC) of the various peptide states; the blue trace corresponds to the Unmodified peptide. **B** Close-up plot of the m/z values of the peptides detected by the mass spectrometer, with the shaded colored areas corresponding to where the expected m/z values for the peptide m/z

(Caption continued on the next page.)

traces should occur, with the observed spectrum plotted in black. The Unmodified peptide(s) were found in the mass spectrum and the Spliced peptide(s) were not detected. **C** Bar plot showing the relative abundance of the various peptide states at charge z=2; the bars represent the average relative abundances normalized to total peak areas, with each individual experimental replicate value plotted as a black dot. **D** Mirrored $MS^2$ plot for the Unmodified peptide. In this mirror plot, the full fragmentation spectrum is plotted in black pointing upward, and the matched fragment b ions (blue) and y ions (red) are plotted pointing downward, mirrored against it, so that each matched fragment can be compared directly with the measured spectrum. **E** $MS^2$ spectrum of the fragments corresponding to the Unmodified peptide sequence, with matched b (blue, above) and y ions (red, below) plotted.

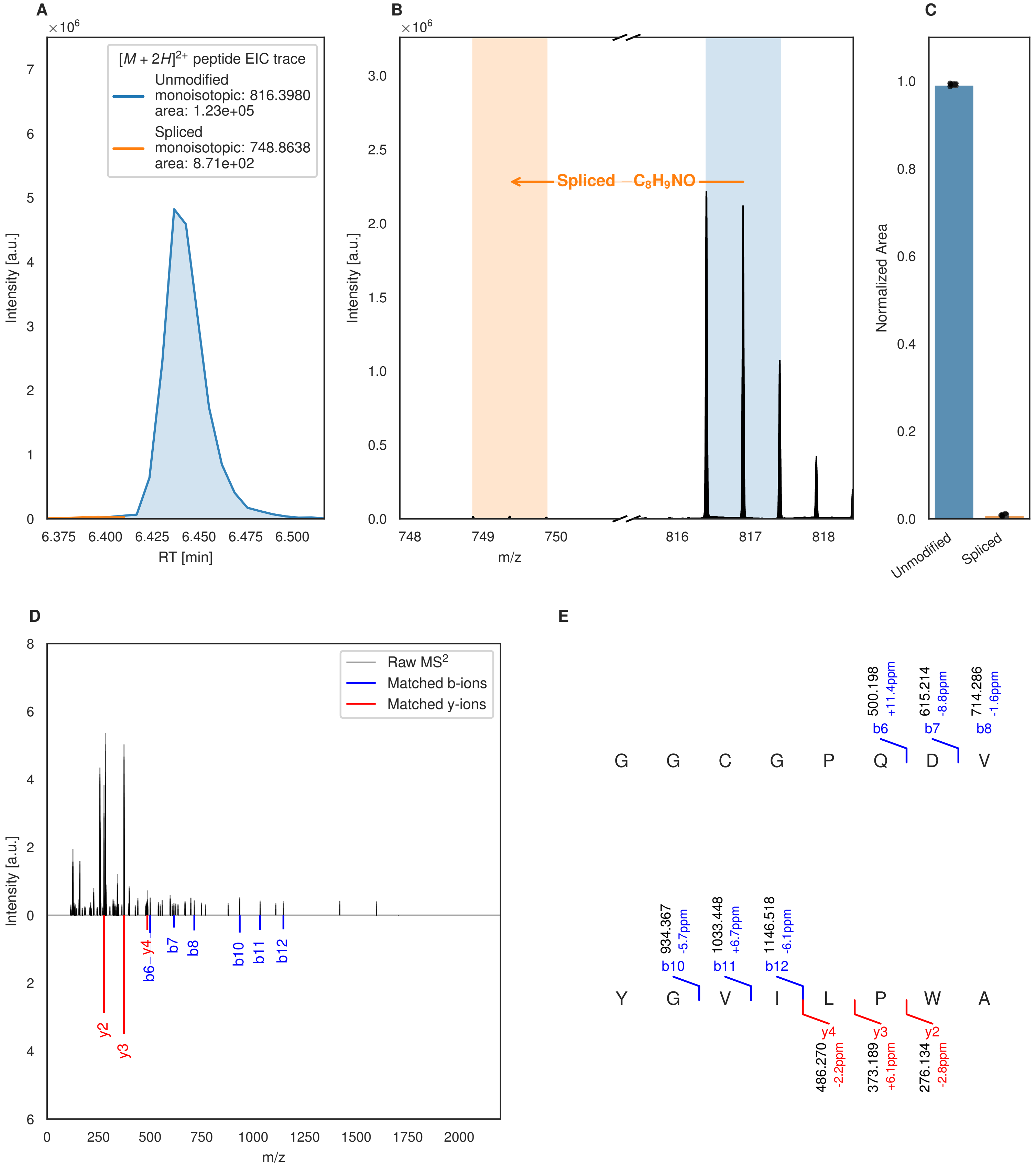
A
×10^6
[M + 2H]^2+ peptide EIC trace
Unmodified
monoisotopic: 816.3980
area: 1.23e+05
Spliced
monoisotopic: 748.8638
area: 8.71e+02
Intensity [a.u.]
RT [min]
B
×10^6
Spliced −C8H9NO
Intensity [a.u.]
m/z
C
Normalized Area
Unmodified
Spliced
D
Raw MS2
Matched b-ions
Matched y-ions
Intensity [a.u.]
m/z
b6−y4
b7
b8
b10
b11
b12
y2
y3
E
500.198
+11.4ppm
b6
615.214
-8.8ppm
b7
714.286
-1.6ppm
b8
G G C G P Q D V
934.367
-5.7ppm
b10
1033.448
+6.7ppm
b11
1146.518
-6.1ppm
b12
Y G V I L P W A
y4
486.270
-2.2ppm
y3
373.189
+6.1ppm
y2
276.134
-2.8ppm

Figure S76 (figure continued from the previous page).

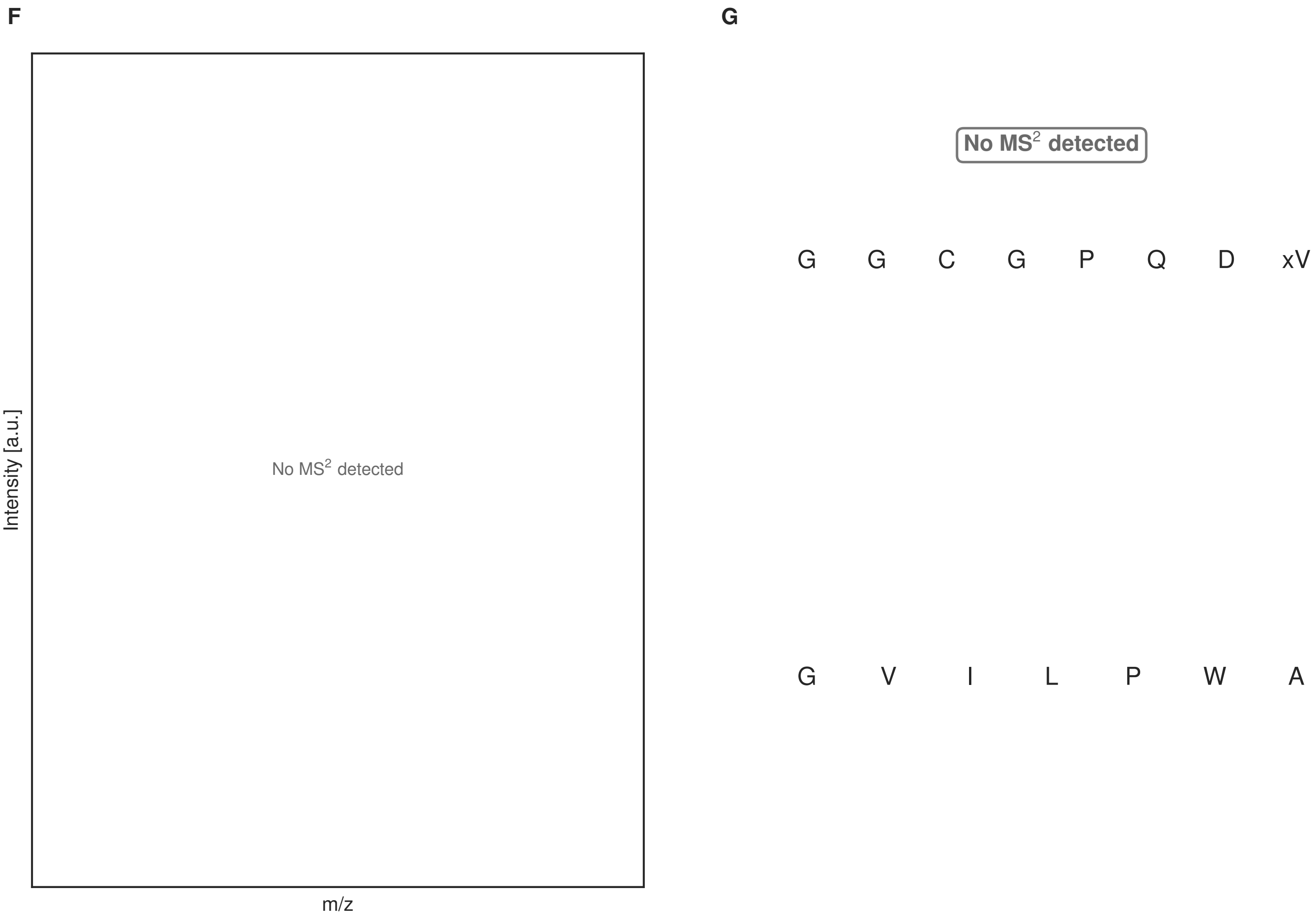


**Figure S76: Modification of the mutant_11 precursor peptide by PlpXY.** LC-MS analysis of IMAC-purified precursor peptides from *E. coli* BL21(DE3)Tuner containing splicease-encoding plasmid TJ-C-559 (PlpXY) and precursor peptide-encoding plasmid mutant_11 (RSTN1-PlpA3-TEV-mutant-11) after expression at 24 °C for 22 h in 1.5 mL cultures after digestion with TEV. Data from n=4 independent biological replicates. Spliced amino acids have an x as a prefix. Spliced refers to a single tyramine excision. A representative experimental replicate is shown in panels **A, B.**

**A** Close-up plot of the representative extracted ion chromatogram (EIC) of the various peptide states; the blue trace corresponds to the Unmodified peptide; the orange trace corresponds to the Spliced peptide. **B** Close-up plot of the m/z values of the peptides detected by the mass spectrometer, with the shaded colored areas corresponding to where the expected m/z values for the peptide m/z traces should occur, with the observed spectrum plotted in black. The Unmodified, Spliced peptide(s) were found in the mass spectrum. **C** Bar plot showing the relative abundance of the various peptide states at charge z=2; the bars represent the average relative abundances normalized to total peak areas, with each individual experimental replicate value plotted as a black dot. **D** Mirrored $MS^2$ plot for the Unmodified peptide. In this mirror plot, the full fragmentation spectrum is plotted in black pointing upward, and the matched fragment b ions (blue) and y ions (red) are plotted pointing downward, mirrored against it, so that each matched fragment can be compared directly with the measured spectrum. **E** $MS^2$ spectrum of the fragments corresponding to the Unmodified peptide sequence, with matched b (blue, above) and y ions (red, below) plotted. **F** Mirrored $MS^2$ plot for the Spliced peptide. In this mirror plot, the full fragmentation spectrum is plotted in black pointing upward, and the matched fragment b ions (blue) and y ions (red) are plotted pointing downward, mirrored against it, so that each matched fragment can be compared directly with the measured spectrum. **G** $MS^2$ spectrum of the fragments corresponding to the Spliced peptide sequence, with matched b (blue, above) and y ions (red, below) plotted.

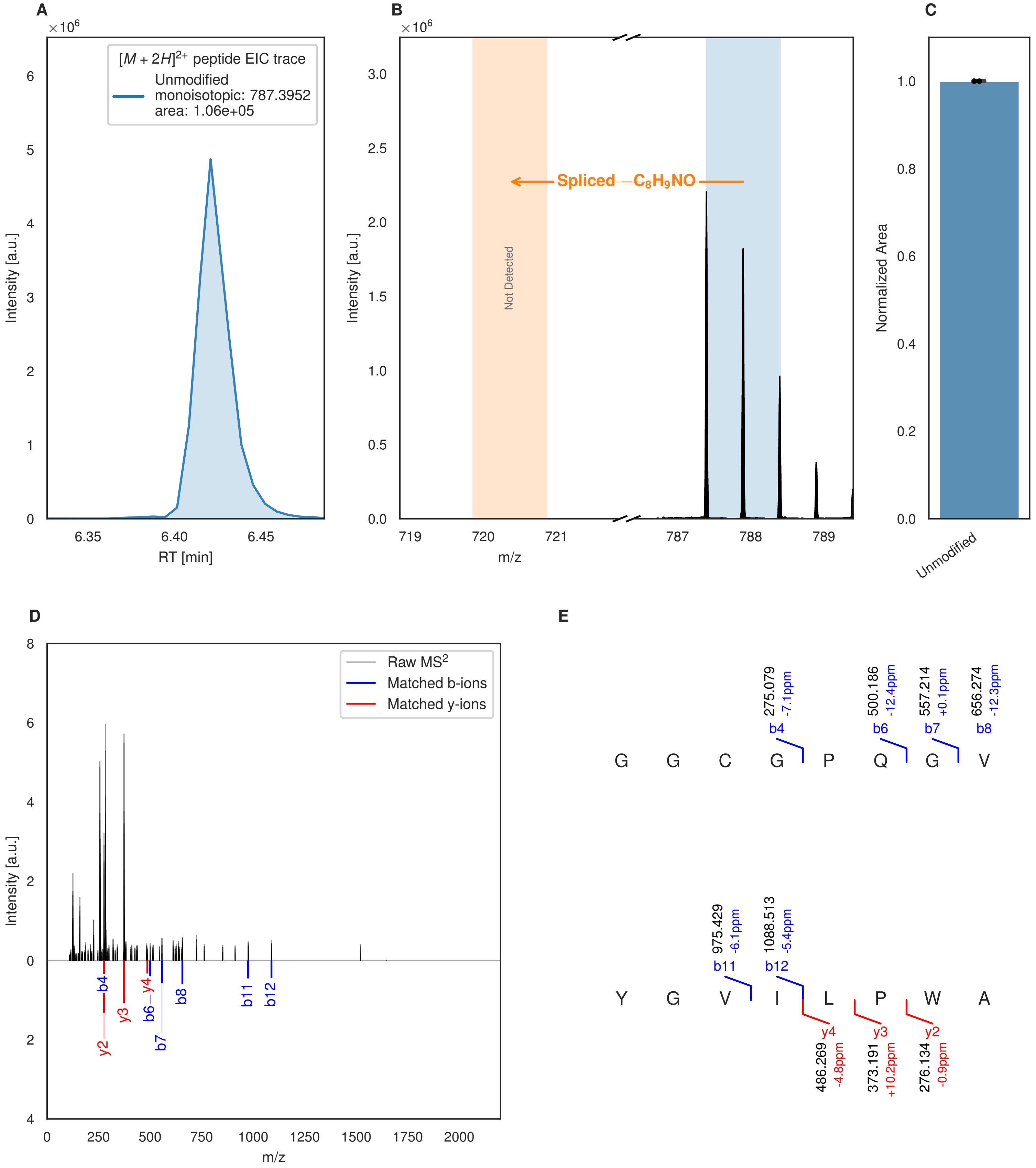


**Figure S77: Modification of the mutant_12 precursor peptide by HypXY.** LC-MS analysis of IMAC-purified precursor peptides from *E. coli* BL21(DE3)Tuner containing splicease-encoding plasmid TJ-C-689 (HypXY) and precursor peptide-encoding plasmid mutant_12 (RSTN1-HypAc-TEV-mutant-12) after expression at 24 °C for 22 h in 1.5 mL cultures after digestion with TEV. Data from n=4 independent biological replicates. Spliced amino acids have an x as a prefix. Spliced refers to a single tyramine excision. A representative experimental replicate is shown in panels **A, B.**

**A** Close-up plot of the representative extracted ion chromatogram (EIC) of the various peptide states; the blue trace corresponds to the Unmodified peptide. **B** Close-up plot of the m/z values of the peptides detected by the mass spectrometer, with the shaded colored areas corresponding to where the expected m/z values for the peptide m/z

(Caption continued on the next page.)

traces should occur, with the observed spectrum plotted in black. The Unmodified peptide(s) were found in the mass spectrum and the Spliced peptide(s) were not detected. **C** Bar plot showing the relative abundance of the various peptide states at charge z=2; the bars represent the average relative abundances normalized to total peak areas, with each individual experimental replicate value plotted as a black dot. **D** Mirrored $MS^2$ plot for the Unmodified peptide. In this mirror plot, the full fragmentation spectrum is plotted in black pointing upward, and the matched fragment b ions (blue) and y ions (red) are plotted pointing downward, mirrored against it, so that each matched fragment can be compared directly with the measured spectrum. **E** $MS^2$ spectrum of the fragments corresponding to the Unmodified peptide sequence, with matched b (blue, above) and y ions (red, below) plotted.

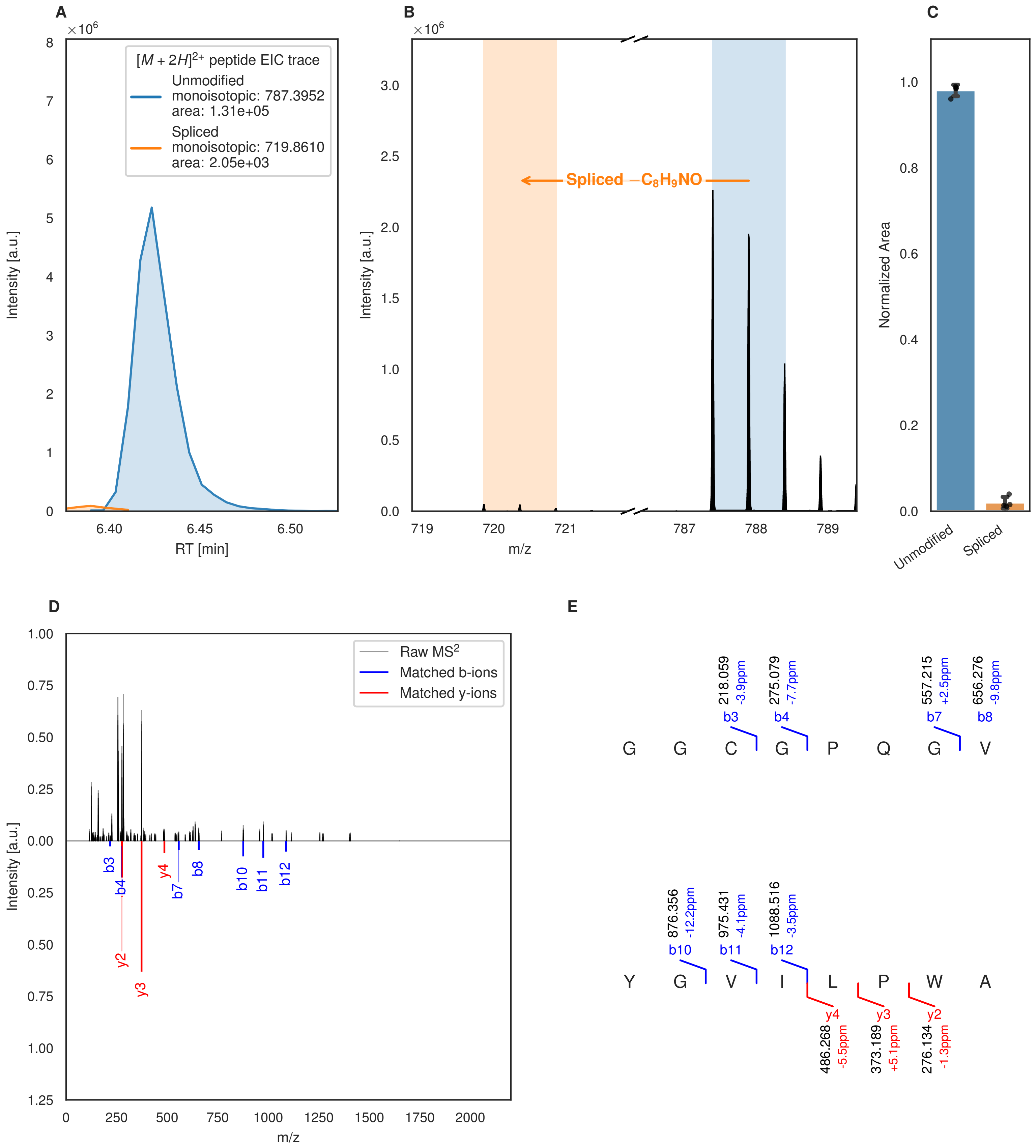
A
B
C
D
E
$[M+2H]^{2+}$ peptide EIC trace
Unmodified
monoisotopic: 787.3952
area: 1.31e+05
Spliced
monoisotopic: 719.8610
area: 2.05e+03
Intensity [a.u.]
RT [min]
Spliced $-C_8H_9NO$
m/z
Normalized Area
Unmodified
Spliced
Raw MS$^2$
Matched b-ions
Matched y-ions
G G C G P Q G V
Y G V I L P W A
218.059 -3.9ppm b3
275.079 -7.7ppm b4
557.215 +2.5ppm b7
656.276 -9.8ppm b8
876.356 -12.2ppm b10
975.431 -4.1ppm b11
1088.516 -3.5ppm b12
y4 486.268 -5.5ppm
y3 373.189 +5.1ppm
y2 276.134 -1.3ppm

Figure S78 (figure continued from the previous page).

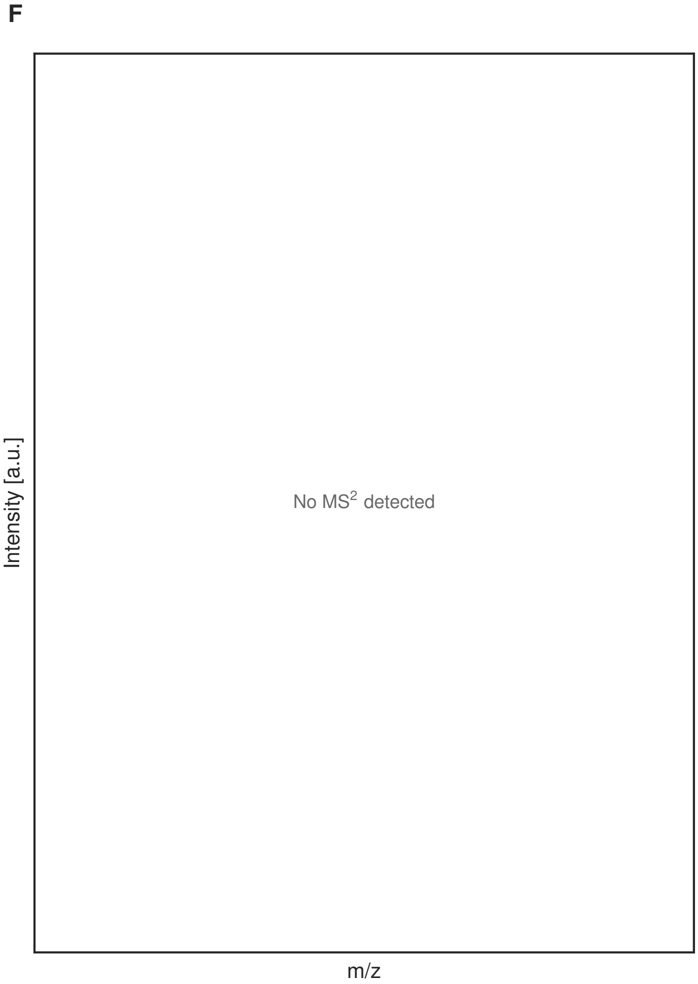


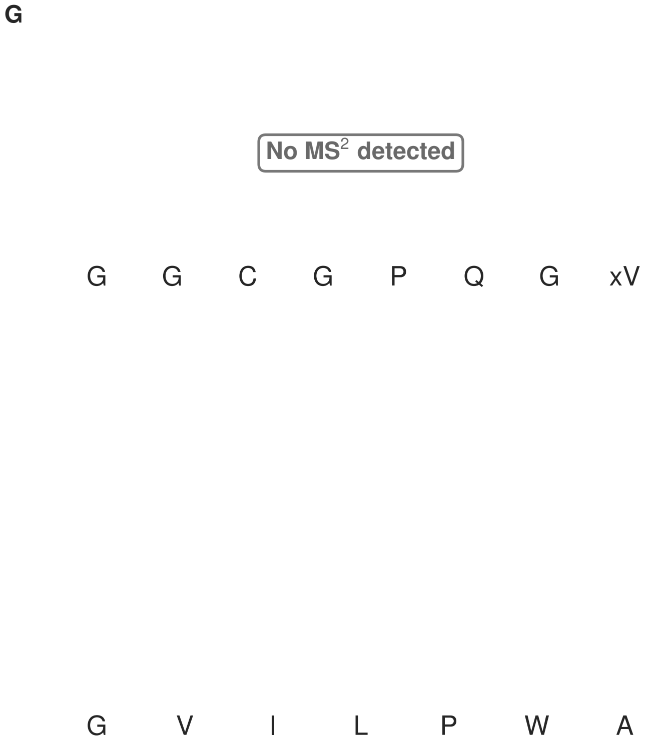


**Figure S78: Modification of the mutant_12 precursor peptide by PlpXY.** LC-MS analysis of IMAC-purified precursor peptides from *E. coli* BL21(DE3)Tuner containing splicease-encoding plasmid TJ-C-559 (PlpXY) and precursor peptide-encoding plasmid mutant_12 (RSTN1-PlpA3-TEV-mutant-12) after expression at 24 °C for 22 h in 1.5 mL cultures after digestion with TEV. Data from n=4 independent biological replicates. Spliced amino acids have an x as a prefix. Spliced refers to a single tyramine excision. A representative experimental replicate is shown in panels **A, B.**

**A** Close-up plot of the representative extracted ion chromatogram (EIC) of the various peptide states; the blue trace corresponds to the Unmodified peptide; the orange trace corresponds to the Spliced peptide. **B** Close-up plot of the m/z values of the peptides detected by the mass spectrometer, with the shaded colored areas corresponding to where the expected m/z values for the peptide m/z traces should occur, with the observed spectrum plotted in black. The Unmodified, Spliced peptide(s) were found in the mass spectrum. **C** Bar plot showing the relative abundance of the various peptide states at charge z=2; the bars represent the average relative abundances normalized to total peak areas, with each individual experimental replicate value plotted as a black dot. **D** Mirrored MS$^2$ plot for the Unmodified peptide. In this mirror plot, the full fragmentation spectrum is plotted in black pointing upward, and the matched fragment b ions (blue) and y ions (red) are plotted pointing downward, mirrored against it, so that each matched fragment can be compared directly with the measured spectrum. **E** MS$^2$ spectrum of the fragments corresponding to the Unmodified peptide sequence, with matched b (blue, above) and y ions (red, below) plotted. **F** Mirrored MS$^2$ plot for the Spliced peptide. In this mirror plot, the full fragmentation spectrum is plotted in black pointing upward, and the matched fragment b ions (blue) and y ions (red) are plotted pointing downward, mirrored against it, so that each matched fragment can be compared directly with the measured spectrum. **G** MS$^2$ spectrum of the fragments corresponding to the Spliced peptide sequence, with matched b (blue, above) and y ions (red, below) plotted.

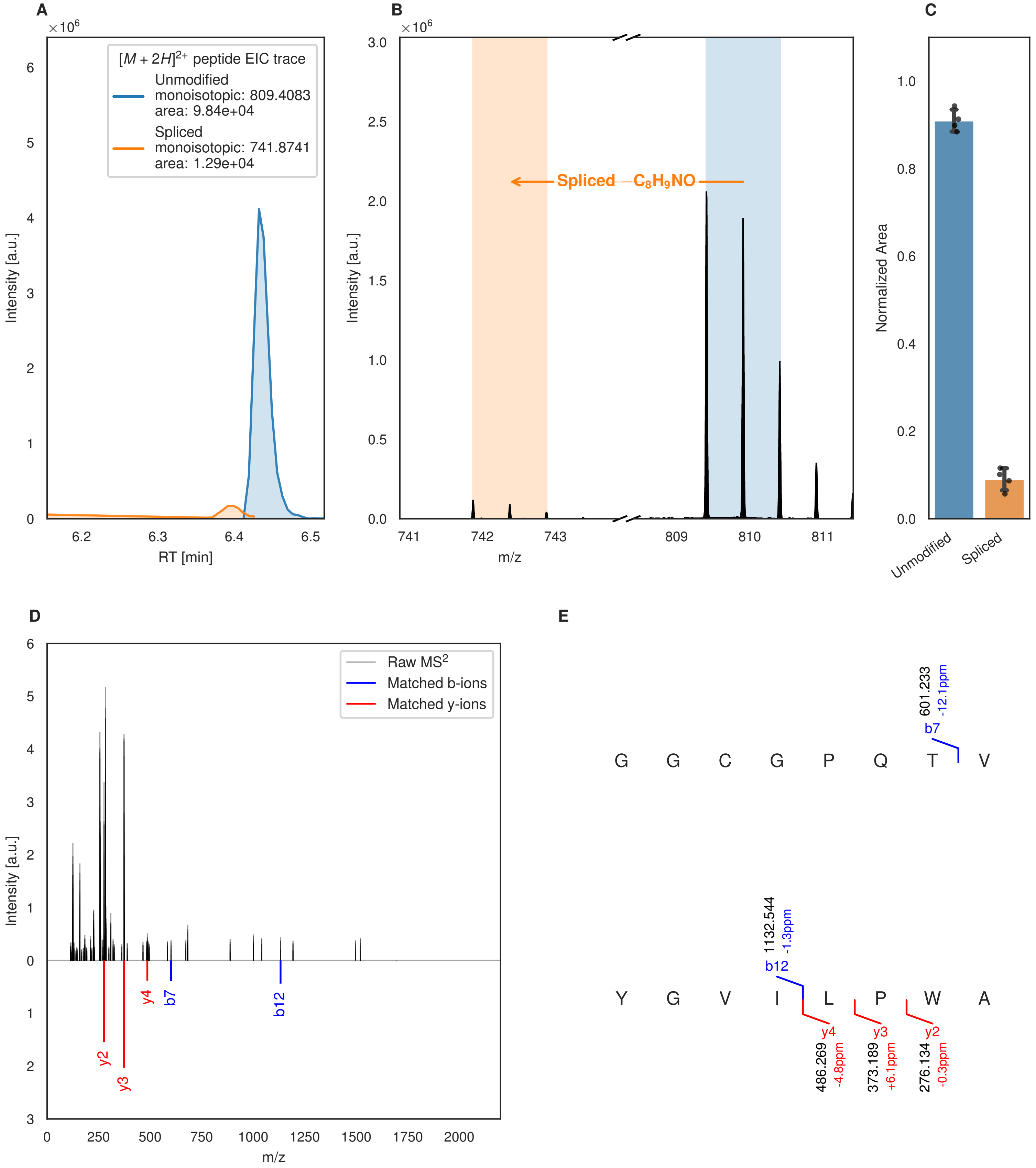

A
B
C
D
E
$[M+2H]^{2+}$ peptide EIC trace
Unmodified
monoisotopic: 809.4083
area: 9.84e+04
Spliced
monoisotopic: 741.8741
area: 1.29e+04
Intensity [a.u.]
RT [min]
Spliced −C8H9NO
m/z
Normalized Area
Unmodified
Spliced
Raw MS2
Matched b-ions
Matched y-ions
G G C G P Q T V
b7
601.233
-12.1ppm
Y G V I L P W A
b12
1132.544
-1.3ppm
y4
486.269
-4.8ppm
y3
373.189
+6.1ppm
y2
276.134
-0.3ppm

Figure S79 (figure continued from the previous page).

**F**

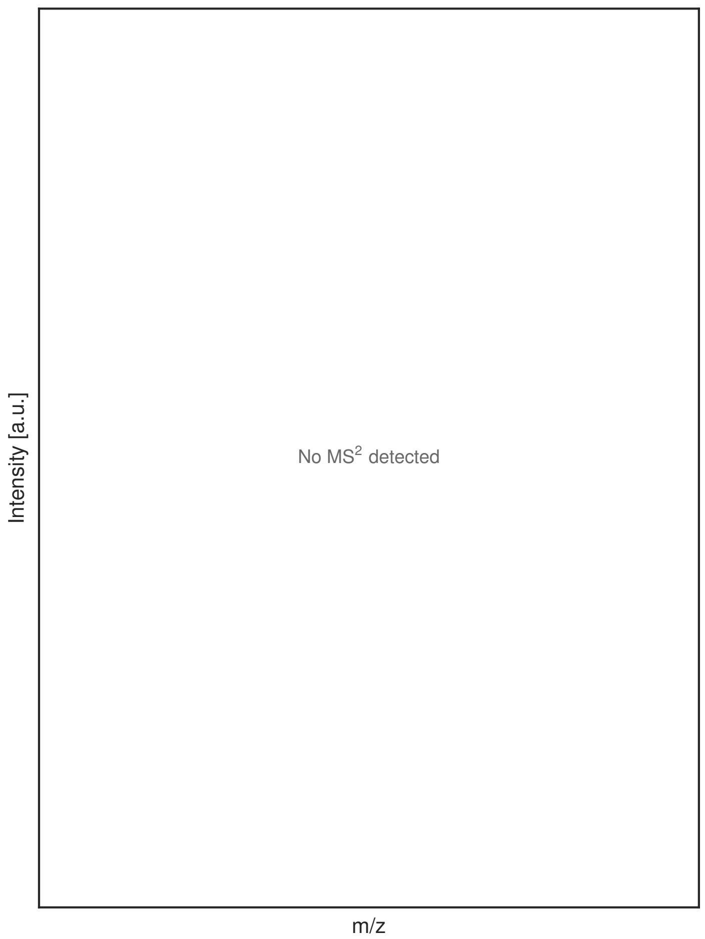


**G**

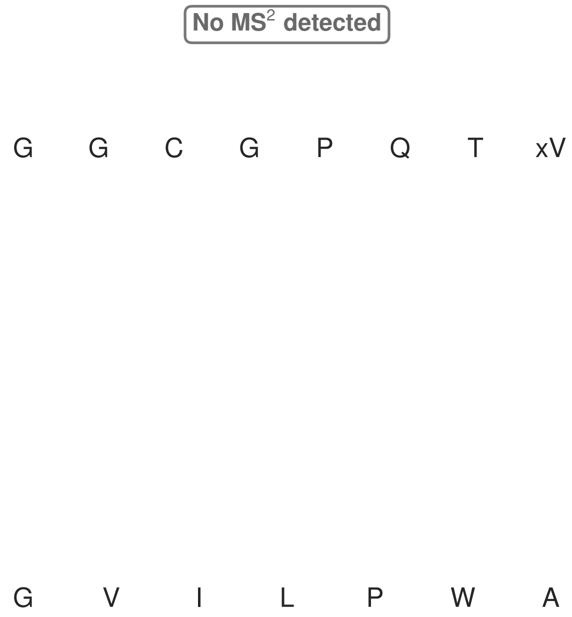


**Figure S79: Modification of the mutant_13 precursor peptide by HypXY.** LC-MS analysis of IMAC-purified precursor peptides from *E. coli* BL21(DE3)Tuner containing splicease-encoding plasmid TJ-C-689 (HypXY) and precursor peptide-encoding plasmid mutant_13 (RSTN1-HypAc-TEV-mutant-13) after expression at 24 °C for 22 h in 1.5 mL cultures after digestion with TEV. Data from n=4 independent biological replicates. Spliced amino acids have an x as a prefix. Spliced refers to a single tyramine excision. A representative experimental replicate is shown in panels **A, B.**

**A** Close-up plot of the representative extracted ion chromatogram (EIC) of the various peptide states; the blue trace corresponds to the Unmodified peptide; the orange trace corresponds to the Spliced peptide. **B** Close-up plot of the m/z values of the peptides detected by the mass spectrometer, with the shaded colored areas corresponding to where the expected m/z values for the peptide m/z traces should occur, with the observed spectrum plotted in black. The Unmodified, Spliced peptide(s) were found in the mass spectrum. **C** Bar plot showing the relative abundance of the various peptide states at charge z=2; the bars represent the average relative abundances normalized to total peak areas, with each individual experimental replicate value plotted as a black dot. **D** Mirrored $MS^2$ plot for the Unmodified peptide. In this mirror plot, the full fragmentation spectrum is plotted in black pointing upward, and the matched fragment b ions (blue) and y ions (red) are plotted pointing downward, mirrored against it, so that each matched fragment can be compared directly with the measured spectrum. **E** $MS^2$ spectrum of the fragments corresponding to the Unmodified peptide sequence, with matched b (blue, above) and y ions (red, below) plotted. **F** Mirrored $MS^2$ plot for the Spliced peptide. In this mirror plot, the full fragmentation spectrum is plotted in black pointing upward, and the matched fragment b ions (blue) and y ions (red) are plotted pointing downward, mirrored against it, so that each matched fragment can be compared directly with the measured spectrum. **G** $MS^2$ spectrum of the fragments corresponding to the Spliced peptide sequence, with matched b (blue, above) and y ions (red, below) plotted.

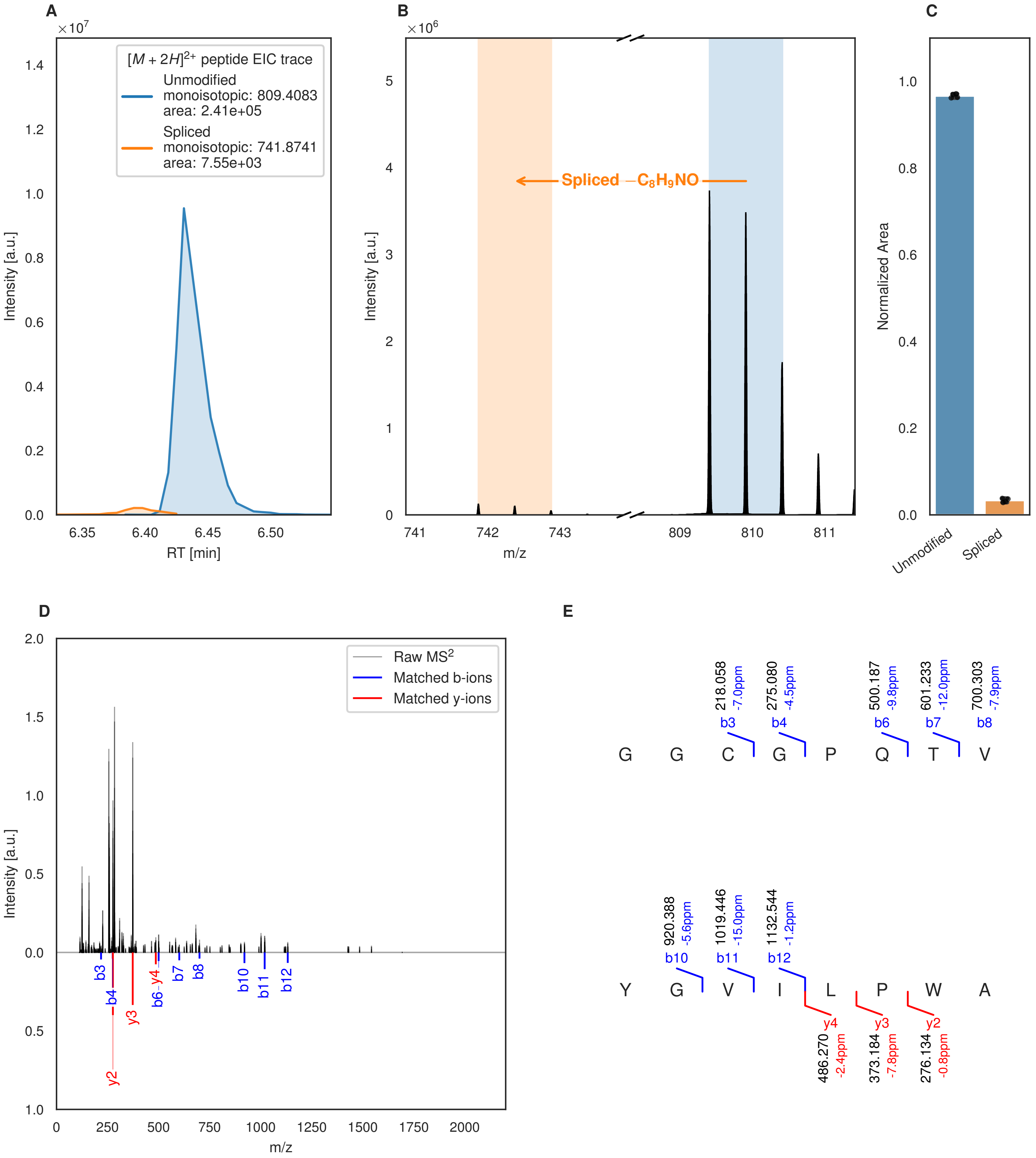
A
[M + 2H]2+ peptide EIC trace
Unmodified
monoisotopic: 809.4083
area: 2.41e+05
Spliced
monoisotopic: 741.8741
area: 7.55e+03
Intensity [a.u.]
RT [min]
B
Spliced −C8H9NO
m/z
C
Normalized Area
Unmodified
Spliced
D
Raw MS2
Matched b-ions
Matched y-ions
m/z
E
G G C G P Q T V
Y G V I L P W A
218.058 -7.0ppm b3
275.080 -4.5ppm b4
500.187 -9.8ppm b6
601.233 -12.0ppm b7
700.303 -7.9ppm b8
920.388 -5.6ppm b10
1019.446 -15.0ppm b11
1132.544 -1.2ppm b12
y4 486.270 -2.4ppm
y3 373.184 -7.8ppm
y2 276.134 -0.8ppm

**F**

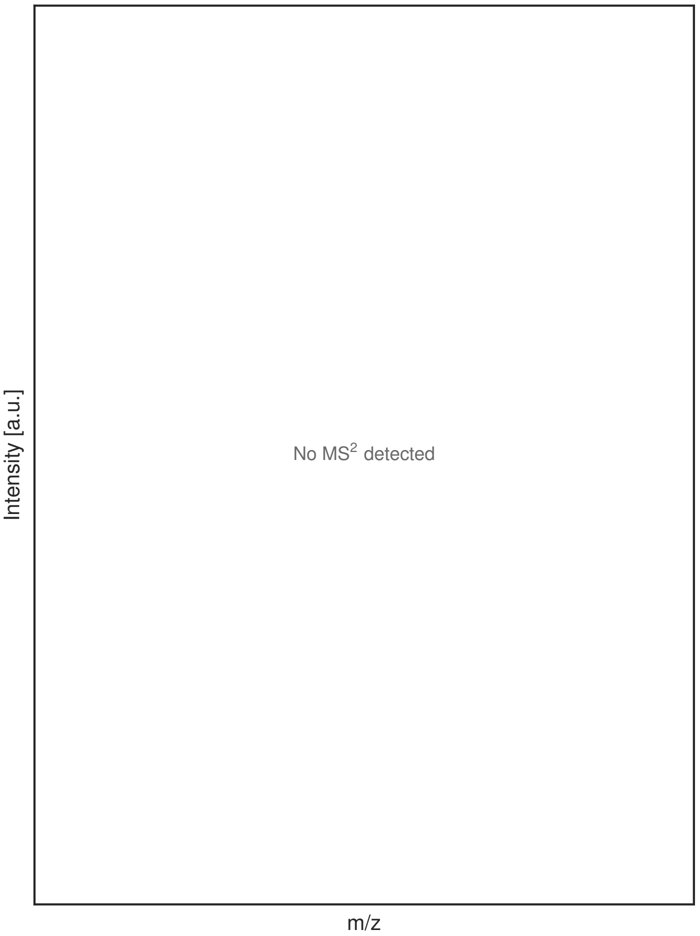


**G**

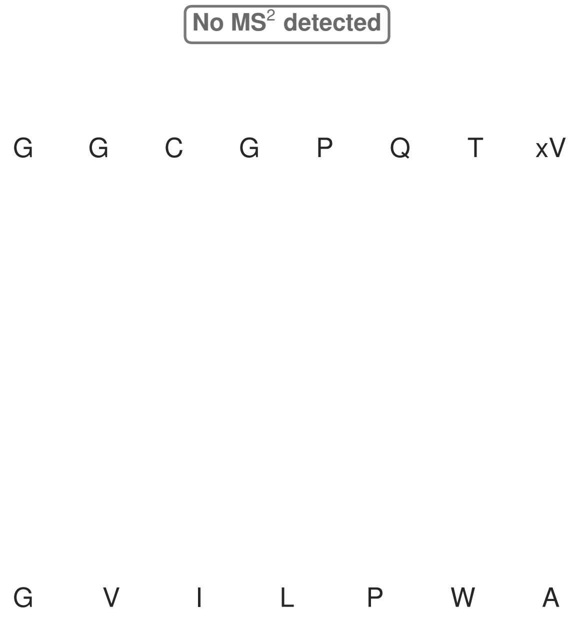


**Figure S80: Modification of the mutant_13 precursor peptide by PlpXY.** LC-MS analysis of IMAC-purified precursor peptides from *E. coli* BL21(DE3)Tuner containing splicease-encoding plasmid TJ-C-559 (PlpXY) and precursor peptide-encoding plasmid mutant_13 (RSTN1-PlpA3-TEV-mutant-13) after expression at 24 °C for 22 h in 1.5 mL cultures after digestion with TEV. Data from n=4 independent biological replicates. Spliced amino acids have an x as a prefix. Spliced refers to a single tyramine excision. A representative experimental replicate is shown in panels **A, B.**

**A** Close-up plot of the representative extracted ion chromatogram (EIC) of the various peptide states; the blue trace corresponds to the Unmodified peptide; the orange trace corresponds to the Spliced peptide. **B** Close-up plot of the m/z values of the peptides detected by the mass spectrometer, with the shaded colored areas corresponding to where the expected m/z values for the peptide m/z traces should occur, with the observed spectrum plotted in black. The Unmodified, Spliced peptide(s) were found in the mass spectrum. **C** Bar plot showing the relative abundance of the various peptide states at charge z=2; the bars represent the average relative abundances normalized to total peak areas, with each individual experimental replicate value plotted as a black dot. **D** Mirrored MS$^2$ plot for the Unmodified peptide. In this mirror plot, the full fragmentation spectrum is plotted in black pointing upward, and the matched fragment b ions (blue) and y ions (red) are plotted pointing downward, mirrored against it, so that each matched fragment can be compared directly with the measured spectrum. **E** MS$^2$ spectrum of the fragments corresponding to the Unmodified peptide sequence, with matched b (blue, above) and y ions (red, below) plotted. **F** Mirrored MS$^2$ plot for the Spliced peptide. In this mirror plot, the full fragmentation spectrum is plotted in black pointing upward, and the matched fragment b ions (blue) and y ions (red) are plotted pointing downward, mirrored against it, so that each matched fragment can be compared directly with the measured spectrum. **G** MS$^2$ spectrum of the fragments corresponding to the Spliced peptide sequence, with matched b (blue, above) and y ions (red, below) plotted.

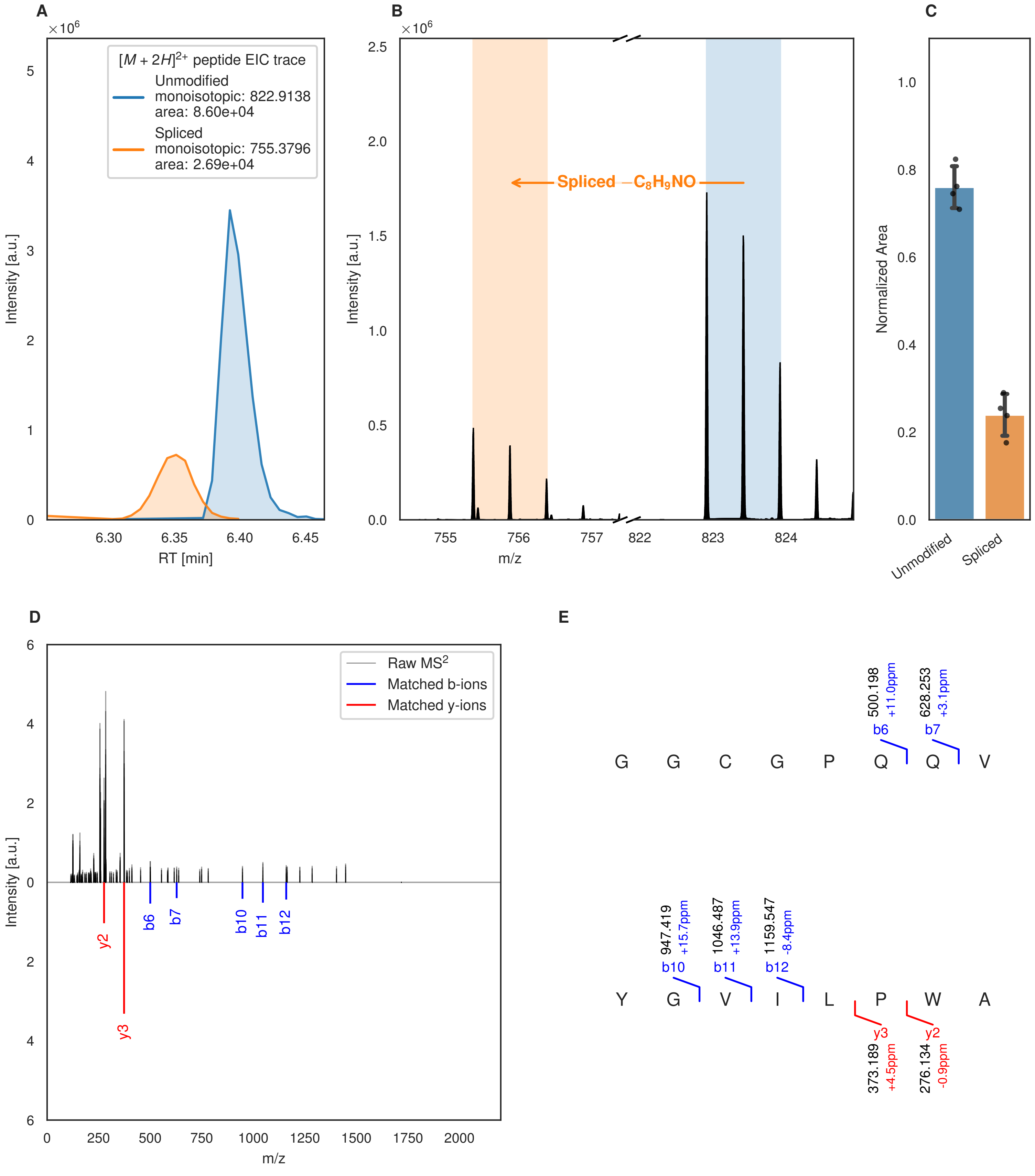
A
B
C
D
E
[M + 2H]2+ peptide EIC trace
Unmodified
monoisotopic: 822.9138
area: 8.60e+04
Spliced
monoisotopic: 755.3796
area: 2.69e+04
Intensity [a.u.]
RT [min]
Spliced −C8H9NO
m/z
Normalized Area
Unmodified
Spliced
Raw MS2
Matched b-ions
Matched y-ions
y2
y3
b6
b7
b10
b11
b12
G G C G P Q Q V
500.198 +11.0ppm
628.253 +3.1ppm
Y G V I L P W A
947.419 +15.7ppm
1046.487 +13.9ppm
1159.547 -8.4ppm
373.189 +4.5ppm
276.134 -0.9ppm

Figure S81 (figure continued from the previous page).

F

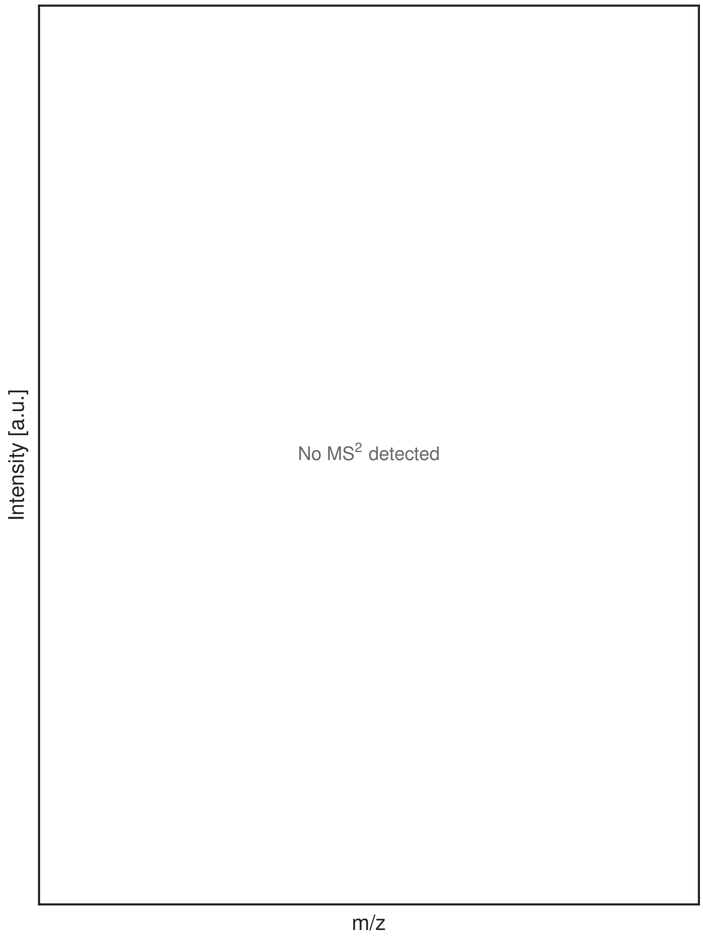


G

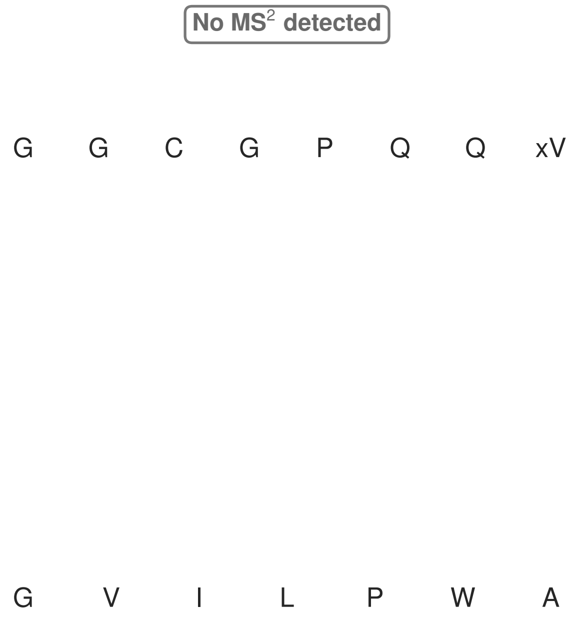


**Figure S81: Modification of the mutant_14 precursor peptide by HypXY.** LC-MS analysis of IMAC-purified precursor peptides from *E. coli* BL21(DE3)Tuner containing splicease-encoding plasmid TJ-C-689 (HypXY) and precursor peptide-encoding plasmid mutant_14 (RSTN1-HypAc-TEV-mutant-14) after expression at 24 °C for 22 h in 1.5 mL cultures after digestion with TEV. Data from n=4 independent biological replicates. Spliced amino acids have an x as a prefix. Spliced refers to a single tyramine excision. A representative experimental replicate is shown in panels **A, B.**

**A** Close-up plot of the representative extracted ion chromatogram (EIC) of the various peptide states; the blue trace corresponds to the Unmodified peptide; the orange trace corresponds to the Spliced peptide. **B** Close-up plot of the m/z values of the peptides detected by the mass spectrometer, with the shaded colored areas corresponding to where the expected m/z values for the peptide m/z traces should occur, with the observed spectrum plotted in black. The Unmodified, Spliced peptide(s) were found in the mass spectrum. **C** Bar plot showing the relative abundance of the various peptide states at charge z=2; the bars represent the average relative abundances normalized to total peak areas, with each individual experimental replicate value plotted as a black dot. **D** Mirrored MS$^2$ plot for the Unmodified peptide. In this mirror plot, the full fragmentation spectrum is plotted in black pointing upward, and the matched fragment b ions (blue) and y ions (red) are plotted pointing downward, mirrored against it, so that each matched fragment can be compared directly with the measured spectrum. **E** MS$^2$ spectrum of the fragments corresponding to the Unmodified peptide sequence, with matched b (blue, above) and y ions (red, below) plotted. **F** Mirrored MS$^2$ plot for the Spliced peptide. In this mirror plot, the full fragmentation spectrum is plotted in black pointing upward, and the matched fragment b ions (blue) and y ions (red) are plotted pointing downward, mirrored against it, so that each matched fragment can be compared directly with the measured spectrum. **G** MS$^2$ spectrum of the fragments corresponding to the Spliced peptide sequence, with matched b (blue, above) and y ions (red, below) plotted.

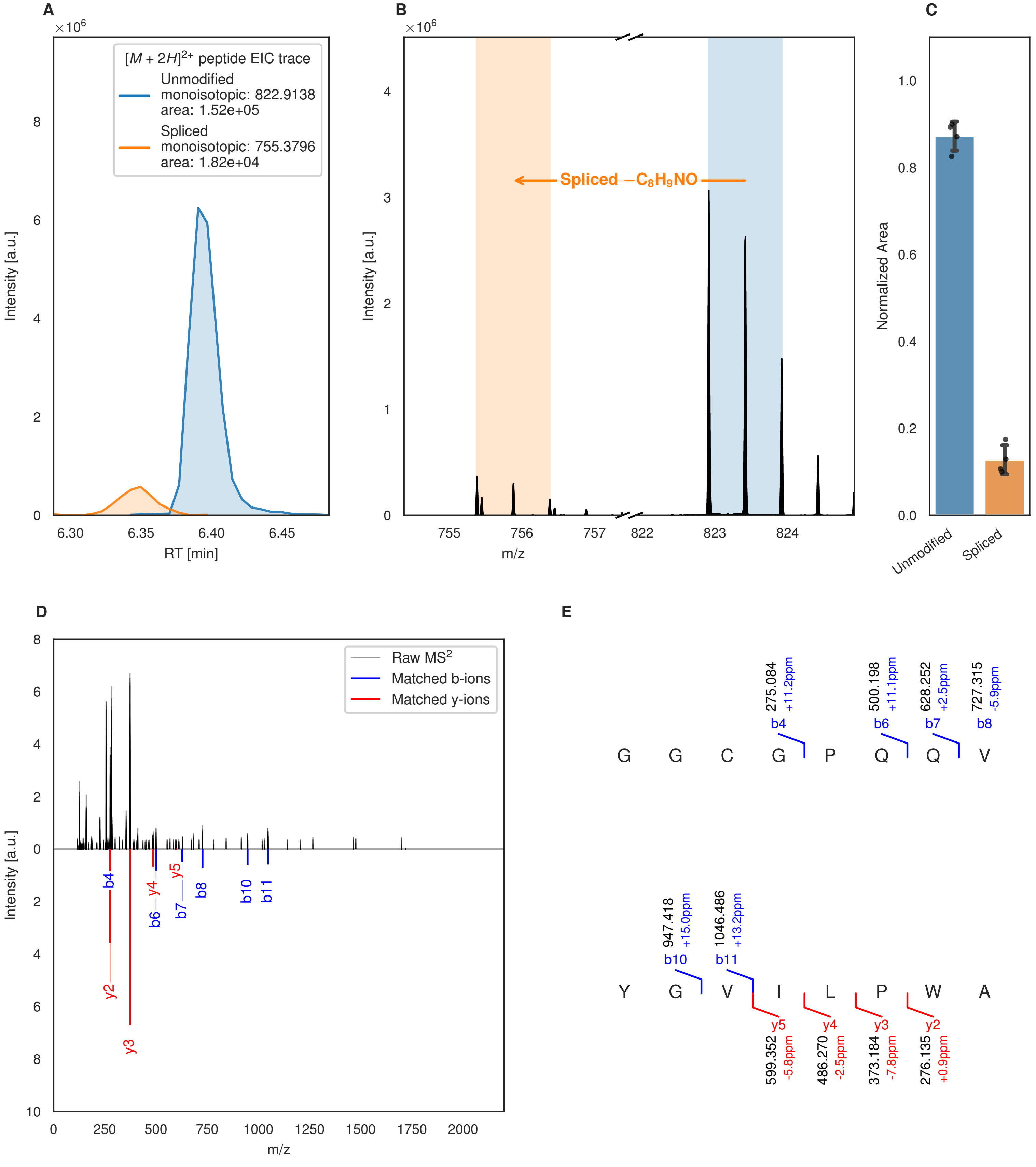
A
B
C
D
E
[M + 2H]2+ peptide EIC trace
Unmodified
monoisotopic: 822.9138
area: 1.52e+05
Spliced
monoisotopic: 755.3796
area: 1.82e+04
Intensity [a.u.]
RT [min]
Spliced −C8H9NO
m/z
Normalized Area
Unmodified
Spliced
Raw MS2
Matched b-ions
Matched y-ions
G G C G P Q Q V
Y G V I L P W A
b4 275.084 +11.2ppm
b6 500.198 +11.1ppm
b7 628.252 +2.5ppm
b8 727.315 -5.9ppm
b10 947.418 +15.0ppm
b11 1046.486 +13.2ppm
y5 599.352 -5.8ppm
y4 486.270 -2.5ppm
y3 373.184 -7.8ppm
y2 276.135 +0.9ppm

Figure S82 (figure continued from the previous page).

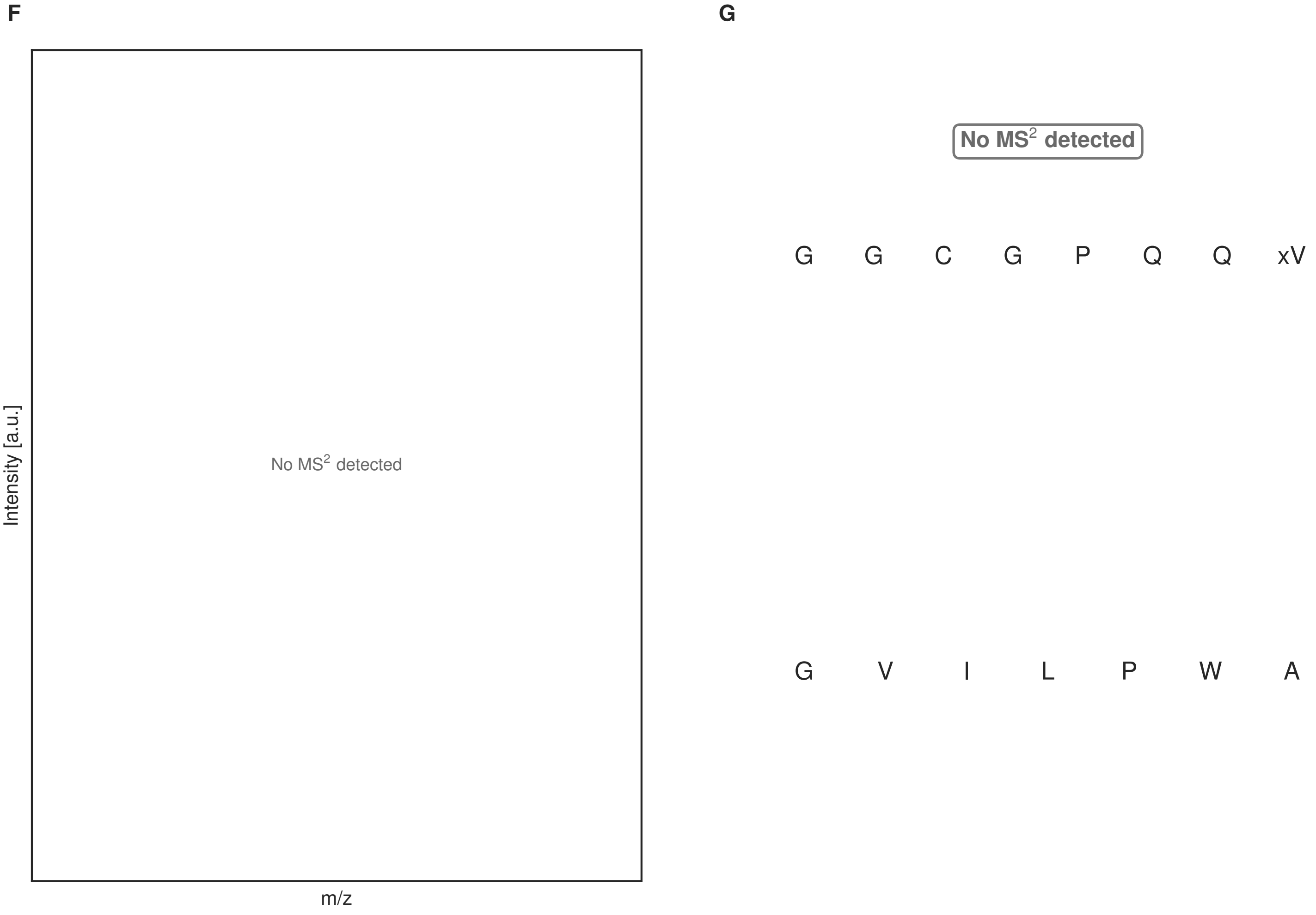


**Figure S82: Modification of the mutant_14 precursor peptide by PlpXY.** LC-MS analysis of IMAC-purified precursor peptides from *E. coli* BL21(DE3)Tuner containing splicease-encoding plasmid TJ-C-559 (PlpXY) and precursor peptide-encoding plasmid mutant_14 (RSTN1-PlpA3-TEV-mutant-14) after expression at 24 °C for 22 h in 1.5 mL cultures after digestion with TEV. Data from n=4 independent biological replicates. Spliced amino acids have an x as a prefix. Spliced refers to a single tyramine excision. A representative experimental replicate is shown in panels **A, B.**

**A** Close-up plot of the representative extracted ion chromatogram (EIC) of the various peptide states; the blue trace corresponds to the Unmodified peptide; the orange trace corresponds to the Spliced peptide. **B** Close-up plot of the m/z values of the peptides detected by the mass spectrometer, with the shaded colored areas corresponding to where the expected m/z values for the peptide m/z traces should occur, with the observed spectrum plotted in black. The Unmodified, Spliced peptide(s) were found in the mass spectrum. **C** Bar plot showing the relative abundance of the various peptide states at charge z=2; the bars represent the average relative abundances normalized to total peak areas, with each individual experimental replicate value plotted as a black dot. **D** Mirrored $MS^2$ plot for the Unmodified peptide. In this mirror plot, the full fragmentation spectrum is plotted in black pointing upward, and the matched fragment b ions (blue) and y ions (red) are plotted pointing downward, mirrored against it, so that each matched fragment can be compared directly with the measured spectrum. **E** $MS^2$ spectrum of the fragments corresponding to the Unmodified peptide sequence, with matched b (blue, above) and y ions (red, below) plotted. **F** Mirrored $MS^2$ plot for the Spliced peptide. In this mirror plot, the full fragmentation spectrum is plotted in black pointing upward, and the matched fragment b ions (blue) and y ions (red) are plotted pointing downward, mirrored against it, so that each matched fragment can be compared directly with the measured spectrum. **G** $MS^2$ spectrum of the fragments corresponding to the Spliced peptide sequence, with matched b (blue, above) and y ions (red, below) plotted.

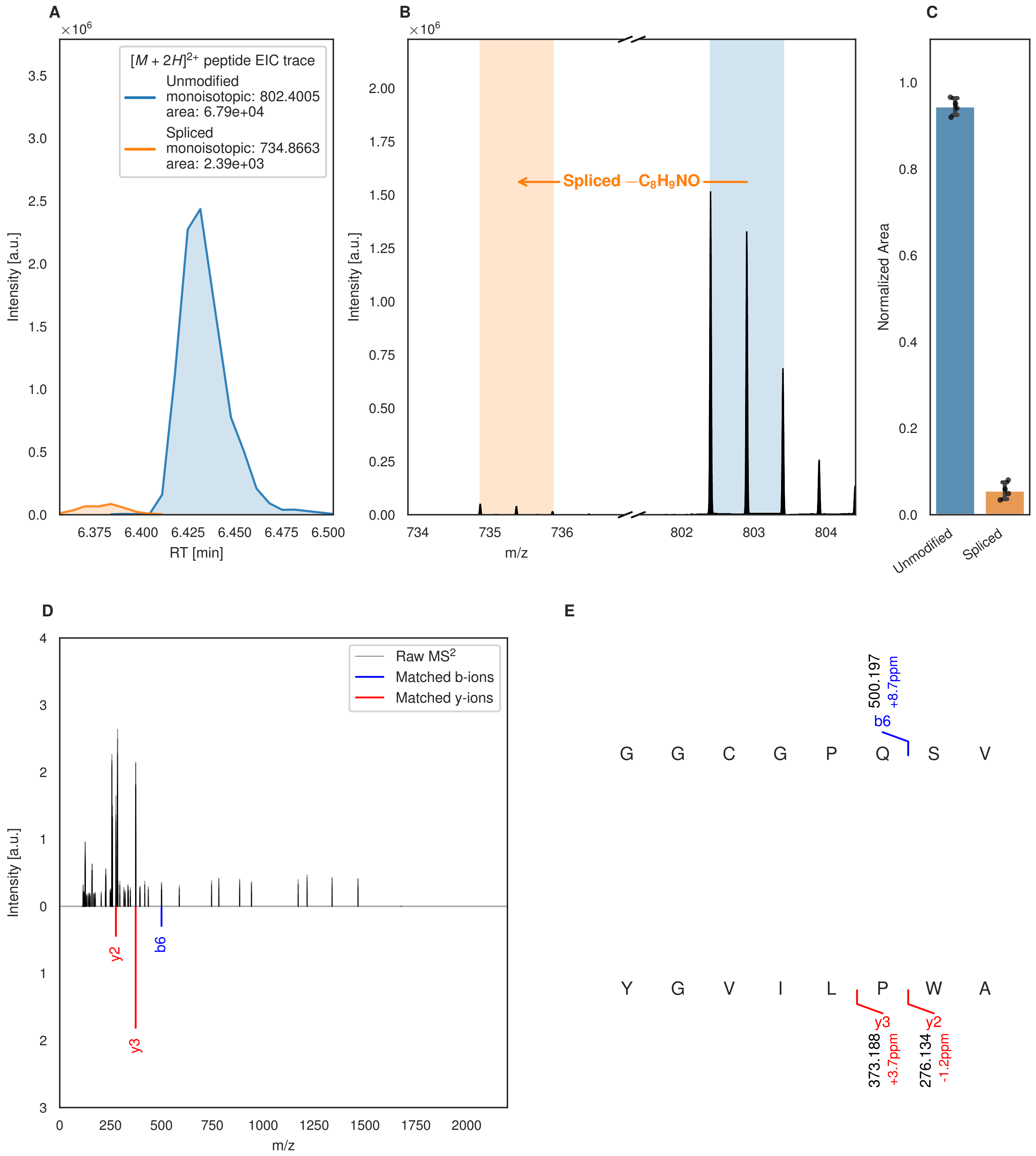
A
×10^6
[M + 2H]2+ peptide EIC trace
Unmodified
monoisotopic: 802.4005
area: 6.79e+04
Spliced
monoisotopic: 734.8663
area: 2.39e+03
Intensity [a.u.]
RT [min]
B
×10^6
Spliced −C8H9NO
Intensity [a.u.]
m/z
C
Normalized Area
Unmodified
Spliced
D
Raw MS2
Matched b-ions
Matched y-ions
Intensity [a.u.]
m/z
y2
y3
b6
E
500.197
+8.7ppm
b6
G G C G P Q S V
Y G V I L P W A
y3
373.188
+3.7ppm
y2
276.134
-1.2ppm

Figure S83 (figure continued from the previous page).

F

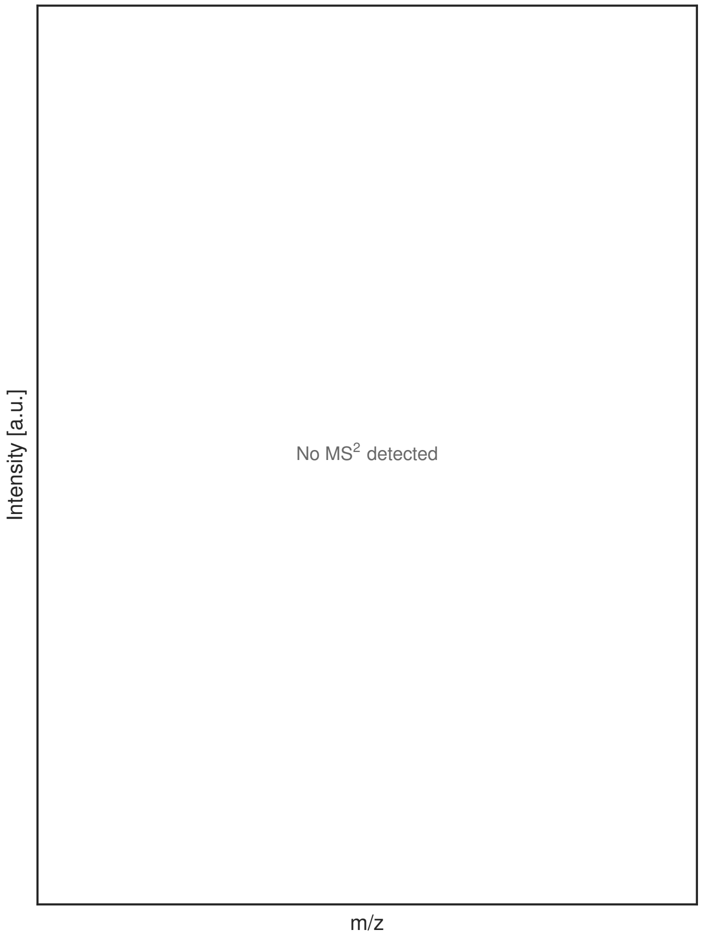


G

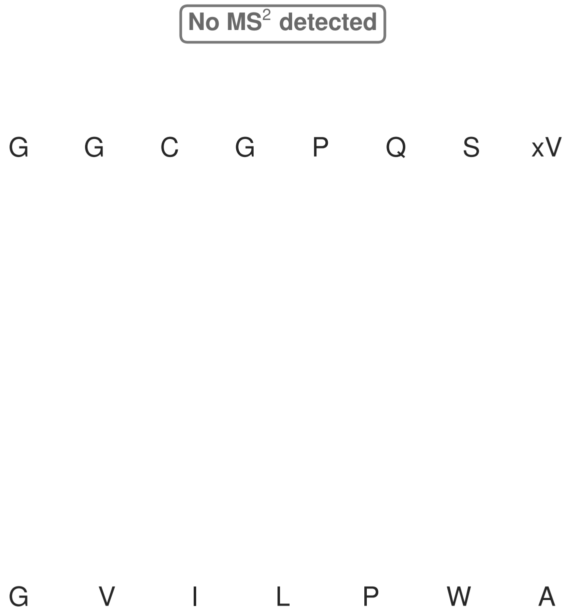


**Figure S83: Modification of the mutant_15 precursor peptide by HypXY.** LC-MS analysis of IMAC-purified precursor peptides from *E. coli* BL21(DE3)Tuner containing splicease-encoding plasmid TJ-C-689 (HypXY) and precursor peptide-encoding plasmid mutant_15 (RSTN1-HypAc-TEV-mutant-15) after expression at 24 °C for 22 h in 1.5 mL cultures after digestion with TEV. Data from n=4 independent biological replicates. Spliced amino acids have an x as a prefix. Spliced refers to a single tyramine excision. A representative experimental replicate is shown in panels **A, B.**

**A** Close-up plot of the representative extracted ion chromatogram (EIC) of the various peptide states; the blue trace corresponds to the Unmodified peptide; the orange trace corresponds to the Spliced peptide. **B** Close-up plot of the m/z values of the peptides detected by the mass spectrometer, with the shaded colored areas corresponding to where the expected m/z values for the peptide m/z traces should occur, with the observed spectrum plotted in black. The Unmodified, Spliced peptide(s) were found in the mass spectrum. **C** Bar plot showing the relative abundance of the various peptide states at charge z=2; the bars represent the average relative abundances normalized to total peak areas, with each individual experimental replicate value plotted as a black dot. **D** Mirrored $MS^2$ plot for the Unmodified peptide. In this mirror plot, the full fragmentation spectrum is plotted in black pointing upward, and the matched fragment b ions (blue) and y ions (red) are plotted pointing downward, mirrored against it, so that each matched fragment can be compared directly with the measured spectrum. **E** $MS^2$ spectrum of the fragments corresponding to the Unmodified peptide sequence, with matched b (blue, above) and y ions (red, below) plotted. **F** Mirrored $MS^2$ plot for the Spliced peptide. In this mirror plot, the full fragmentation spectrum is plotted in black pointing upward, and the matched fragment b ions (blue) and y ions (red) are plotted pointing downward, mirrored against it, so that each matched fragment can be compared directly with the measured spectrum. **G** $MS^2$ spectrum of the fragments corresponding to the Spliced peptide sequence, with matched b (blue, above) and y ions (red, below) plotted.

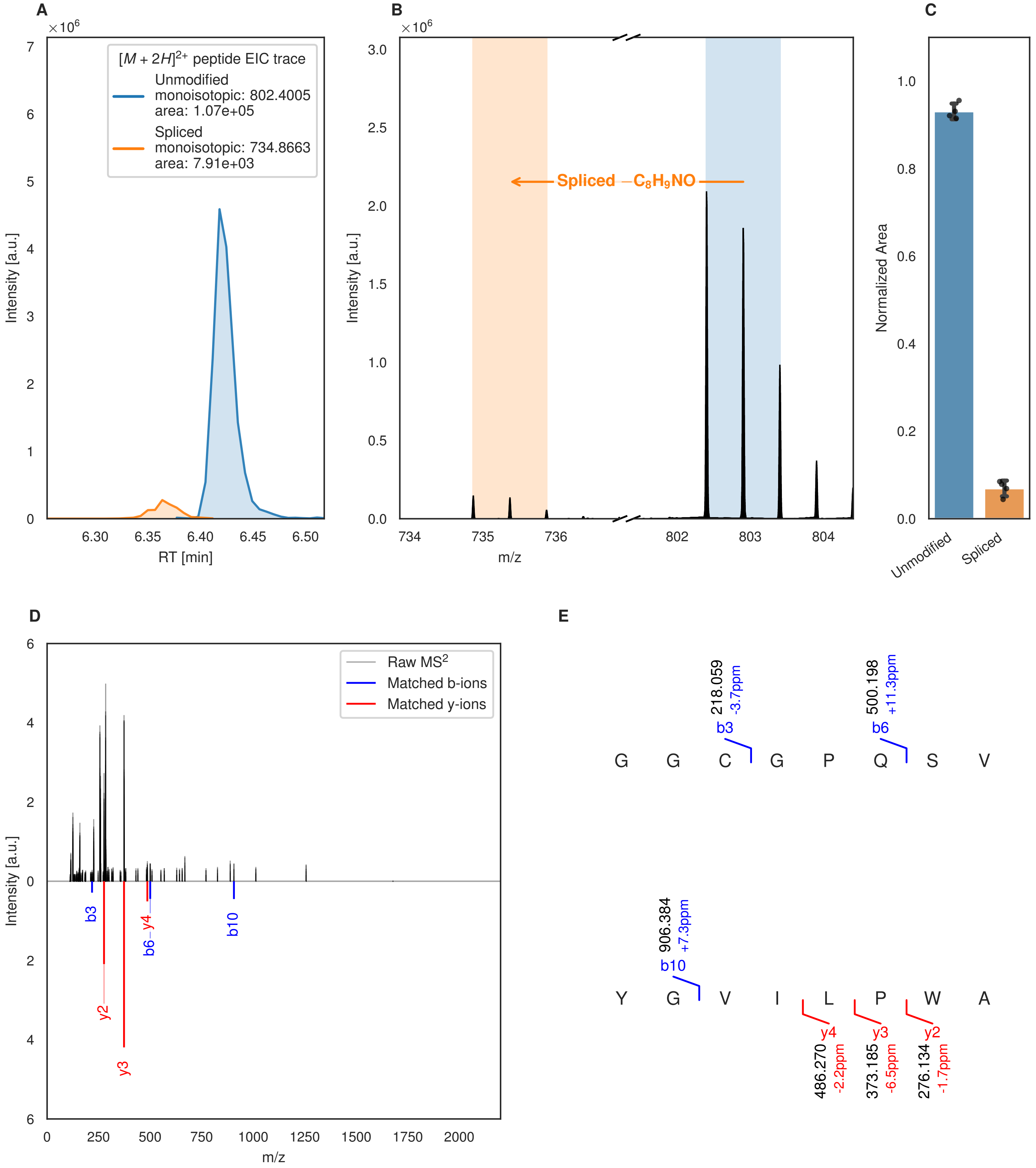
A
B
C
$[M + 2H]^{2+}$ peptide EIC trace
Unmodified
monoisotopic: 802.4005
area: 1.07e+05
Spliced
monoisotopic: 734.8663
area: 7.91e+03
Intensity [a.u.]
RT [min]
Spliced $-C_8H_9NO$
m/z
Normalized Area
Unmodified
Spliced
D
E
Raw $MS^2$
Matched b-ions
Matched y-ions
b3
y2
y3
b6
y4
b10
218.059
-3.7ppm
500.198
+11.3ppm
906.384
+7.3ppm
486.270
-2.2ppm
373.185
-6.5ppm
276.134
-1.7ppm
G G C G P Q S V
Y G V I L P W A

Figure S84 (figure continued from the previous page).

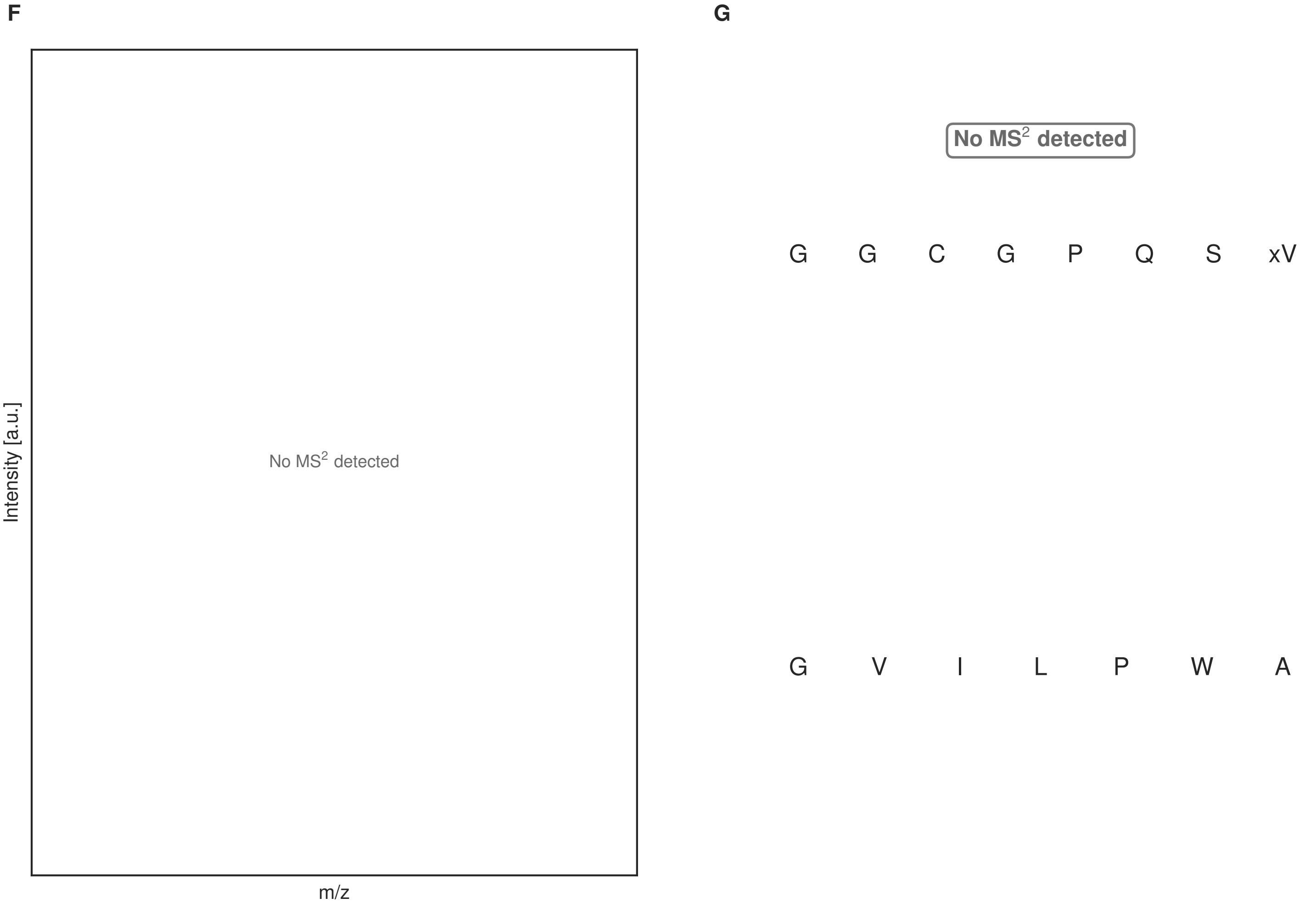


**Figure S84: Modification of the mutant_15 precursor peptide by PlpXY.** LC-MS analysis of IMAC-purified precursor peptides from *E. coli* BL21(DE3)Tuner containing splicease-encoding plasmid TJ-C-559 (PlpXY) and precursor peptide-encoding plasmid mutant_15 (RSTN1-PlpA3-TEV-mutant-15) after expression at 24 °C for 22 h in 1.5 mL cultures after digestion with TEV. Data from n=4 independent biological replicates. Spliced amino acids have an x as a prefix. Spliced refers to a single tyramine excision. A representative experimental replicate is shown in panels **A, B.**

**A** Close-up plot of the representative extracted ion chromatogram (EIC) of the various peptide states; the blue trace corresponds to the Unmodified peptide; the orange trace corresponds to the Spliced peptide. **B** Close-up plot of the m/z values of the peptides detected by the mass spectrometer, with the shaded colored areas corresponding to where the expected m/z values for the peptide m/z traces should occur, with the observed spectrum plotted in black. The Unmodified, Spliced peptide(s) were found in the mass spectrum. **C** Bar plot showing the relative abundance of the various peptide states at charge z=2; the bars represent the average relative abundances normalized to total peak areas, with each individual experimental replicate value plotted as a black dot. **D** Mirrored MS$^2$ plot for the Unmodified peptide. In this mirror plot, the full fragmentation spectrum is plotted in black pointing upward, and the matched fragment b ions (blue) and y ions (red) are plotted pointing downward, mirrored against it, so that each matched fragment can be compared directly with the measured spectrum. **E** MS$^2$ spectrum of the fragments corresponding to the Unmodified peptide sequence, with matched b (blue, above) and y ions (red, below) plotted. **F** Mirrored MS$^2$ plot for the Spliced peptide. In this mirror plot, the full fragmentation spectrum is plotted in black pointing upward, and the matched fragment b ions (blue) and y ions (red) are plotted pointing downward, mirrored against it, so that each matched fragment can be compared directly with the measured spectrum. **G** MS$^2$ spectrum of the fragments corresponding to the Spliced peptide sequence, with matched b (blue, above) and y ions (red, below) plotted.

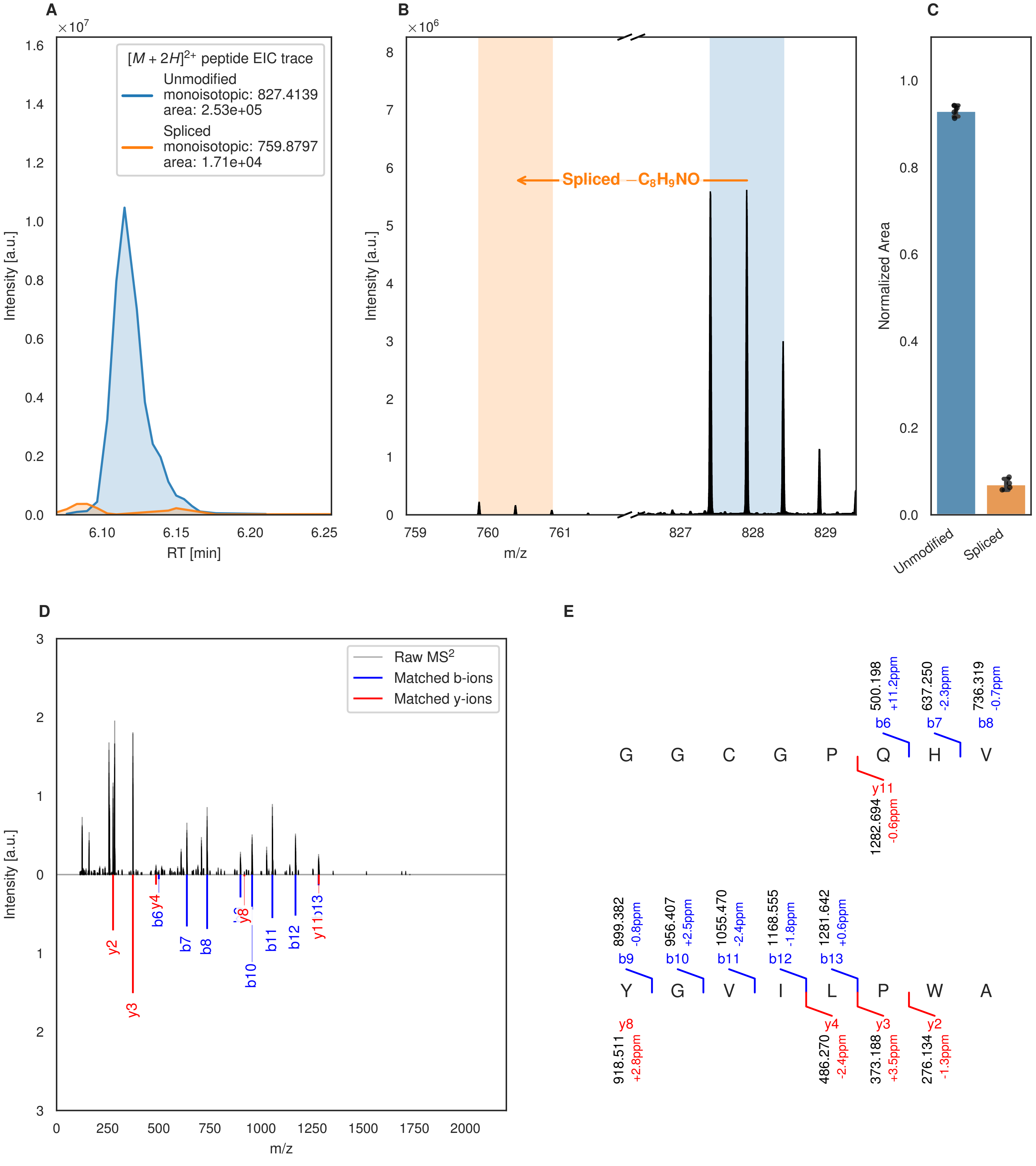
A
B
C
D
E
[M + 2H]2+ peptide EIC trace
Unmodified
monoisotopic: 827.4139
area: 2.53e+05
Spliced
monoisotopic: 759.8797
area: 1.71e+04
Intensity [a.u.]
RT [min]
Spliced −C8H9NO
m/z
Normalized Area
Unmodified
Spliced
Raw MS2
Matched b-ions
Matched y-ions
G G C G P Q H V
Y G V I L P W A
b6 500.198 +11.2ppm
b7 637.250 -2.3ppm
b8 736.319 -0.7ppm
y11 1282.694 -0.6ppm
b9 899.382 -0.8ppm
b10 956.407 +2.5ppm
b11 1055.470 -2.4ppm
b12 1168.555 -1.8ppm
b13 1281.642 +0.6ppm
y8 918.511 +2.8ppm
y4 486.270 -2.4ppm
y3 373.188 +3.5ppm
y2 276.134 -1.3ppm

F

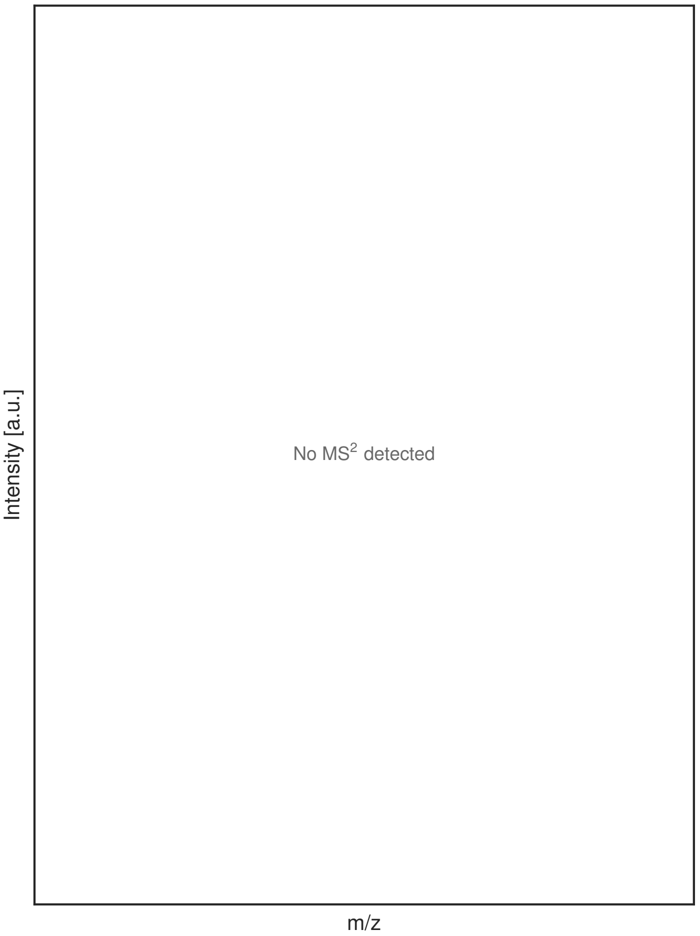


G

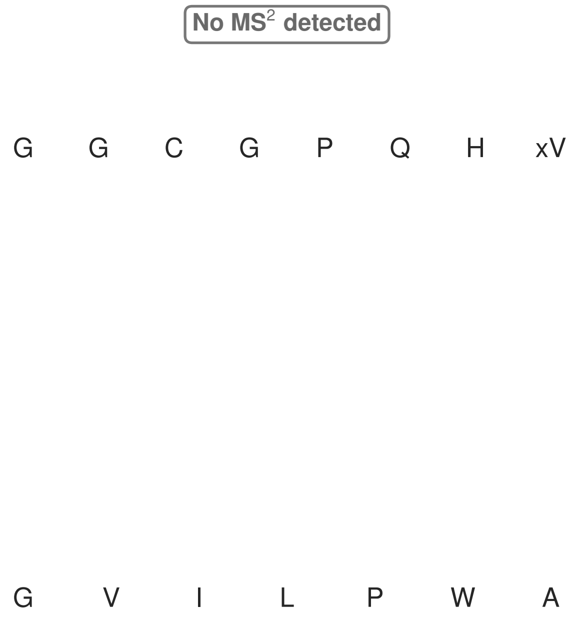


**Figure S85: Modification of the mutant_16 precursor peptide by HypXY.** LC-MS analysis of IMAC-purified precursor peptides from *E. coli* BL21(DE3)Tuner containing splicease-encoding plasmid TJ-C-689 (HypXY) and precursor peptide-encoding plasmid mutant_16 (RSTN1-HypAc-TEV-mutant-16) after expression at 24 °C for 22 h in 1.5 mL cultures after digestion with TEV. Data from n=4 independent biological replicates. Spliced amino acids have an x as a prefix. Spliced refers to a single tyramine excision. A representative experimental replicate is shown in panels **A, B.**

**A** Close-up plot of the representative extracted ion chromatogram (EIC) of the various peptide states; the blue trace corresponds to the Unmodified peptide; the orange trace corresponds to the Spliced peptide. **B** Close-up plot of the m/z values of the peptides detected by the mass spectrometer, with the shaded colored areas corresponding to where the expected m/z values for the peptide m/z traces should occur, with the observed spectrum plotted in black. The Unmodified, Spliced peptide(s) were found in the mass spectrum. **C** Bar plot showing the relative abundance of the various peptide states at charge z=2; the bars represent the average relative abundances normalized to total peak areas, with each individual experimental replicate value plotted as a black dot. **D** Mirrored $MS^2$ plot for the Unmodified peptide. In this mirror plot, the full fragmentation spectrum is plotted in black pointing upward, and the matched fragment b ions (blue) and y ions (red) are plotted pointing downward, mirrored against it, so that each matched fragment can be compared directly with the measured spectrum. **E** $MS^2$ spectrum of the fragments corresponding to the Unmodified peptide sequence, with matched b (blue, above) and y ions (red, below) plotted. **F** Mirrored $MS^2$ plot for the Spliced peptide. In this mirror plot, the full fragmentation spectrum is plotted in black pointing upward, and the matched fragment b ions (blue) and y ions (red) are plotted pointing downward, mirrored against it, so that each matched fragment can be compared directly with the measured spectrum. **G** $MS^2$ spectrum of the fragments corresponding to the Spliced peptide sequence, with matched b (blue, above) and y ions (red, below) plotted.

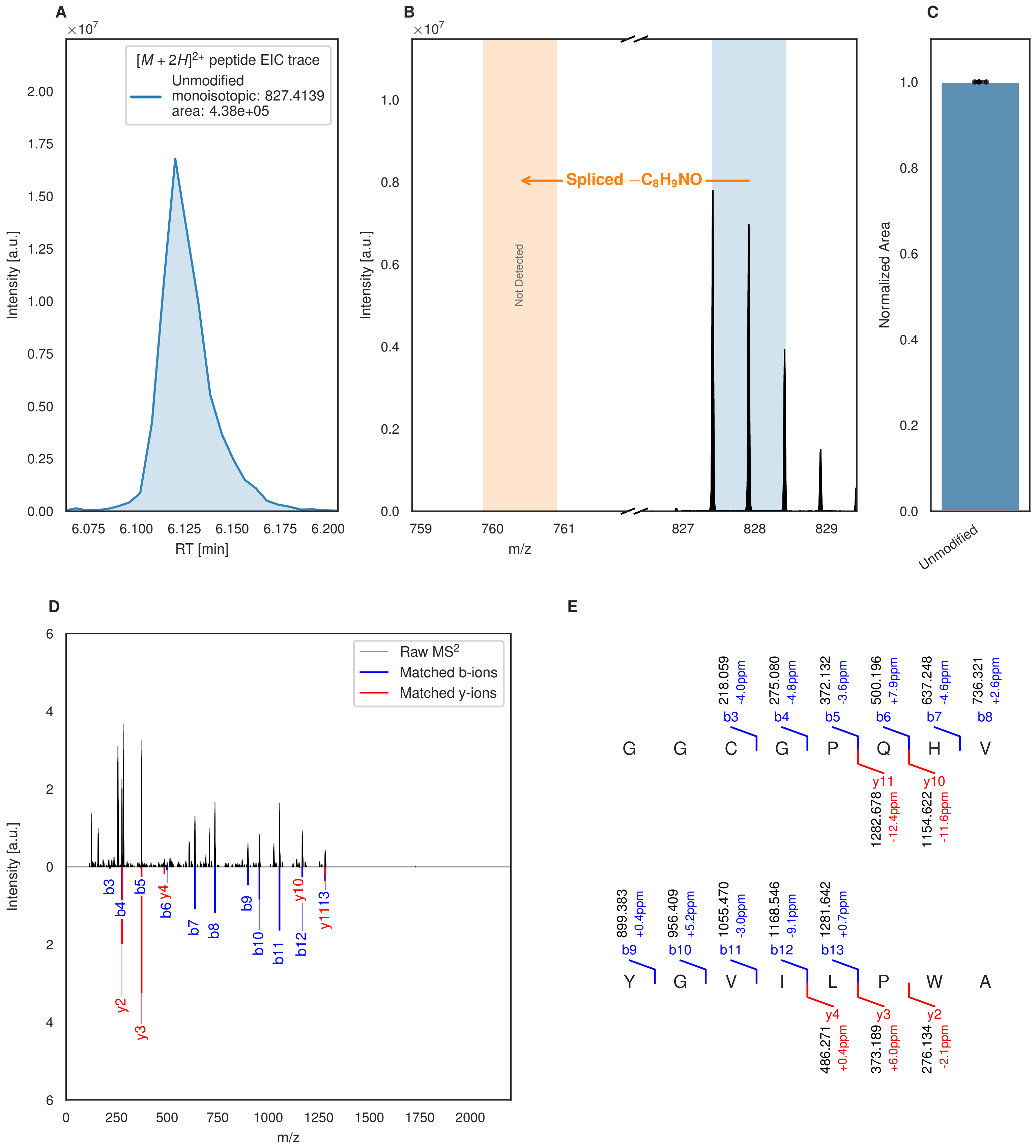


**Figure S86: Modification of the mutant_16 precursor peptide by PlpXY.** LC-MS analysis of IMAC-purified precursor peptides from *E. coli* BL21(DE3)Tuner containing splicease-encoding plasmid TJ-C-559 (PlpXY) and precursor peptide-encoding plasmid mutant_16 (RSTN1-PlpA3-TEV-mutant-16) after expression at 24 °C for 22 h in 1.5 mL cultures after digestion with TEV. Data from n=4 independent biological replicates. Spliced amino acids have an x as a prefix. Spliced refers to a single tyramine excision. A representative experimental replicate is shown in panels **A, B.**

**A** Close-up plot of the representative extracted ion chromatogram (EIC) of the various peptide states; the blue trace corresponds to the Unmodified peptide. **B** Close-up plot of the m/z values of the peptides detected by the mass spectrometer, with the shaded colored areas corresponding to where the expected m/z values for the peptide m/z

(Caption continued on the next page.)

traces should occur, with the observed spectrum plotted in black. The Unmodified peptide(s) were found in the mass spectrum and the Spliced peptide(s) were not detected. **C** Bar plot showing the relative abundance of the various peptide states at charge z=2; the bars represent the average relative abundances normalized to total peak areas, with each individual experimental replicate value plotted as a black dot. **D** Mirrored $MS^2$ plot for the Unmodified peptide. In this mirror plot, the full fragmentation spectrum is plotted in black pointing upward, and the matched fragment b ions (blue) and y ions (red) are plotted pointing downward, mirrored against it, so that each matched fragment can be compared directly with the measured spectrum. **E** $MS^2$ spectrum of the fragments corresponding to the Unmodified peptide sequence, with matched b (blue, above) and y ions (red, below) plotted.

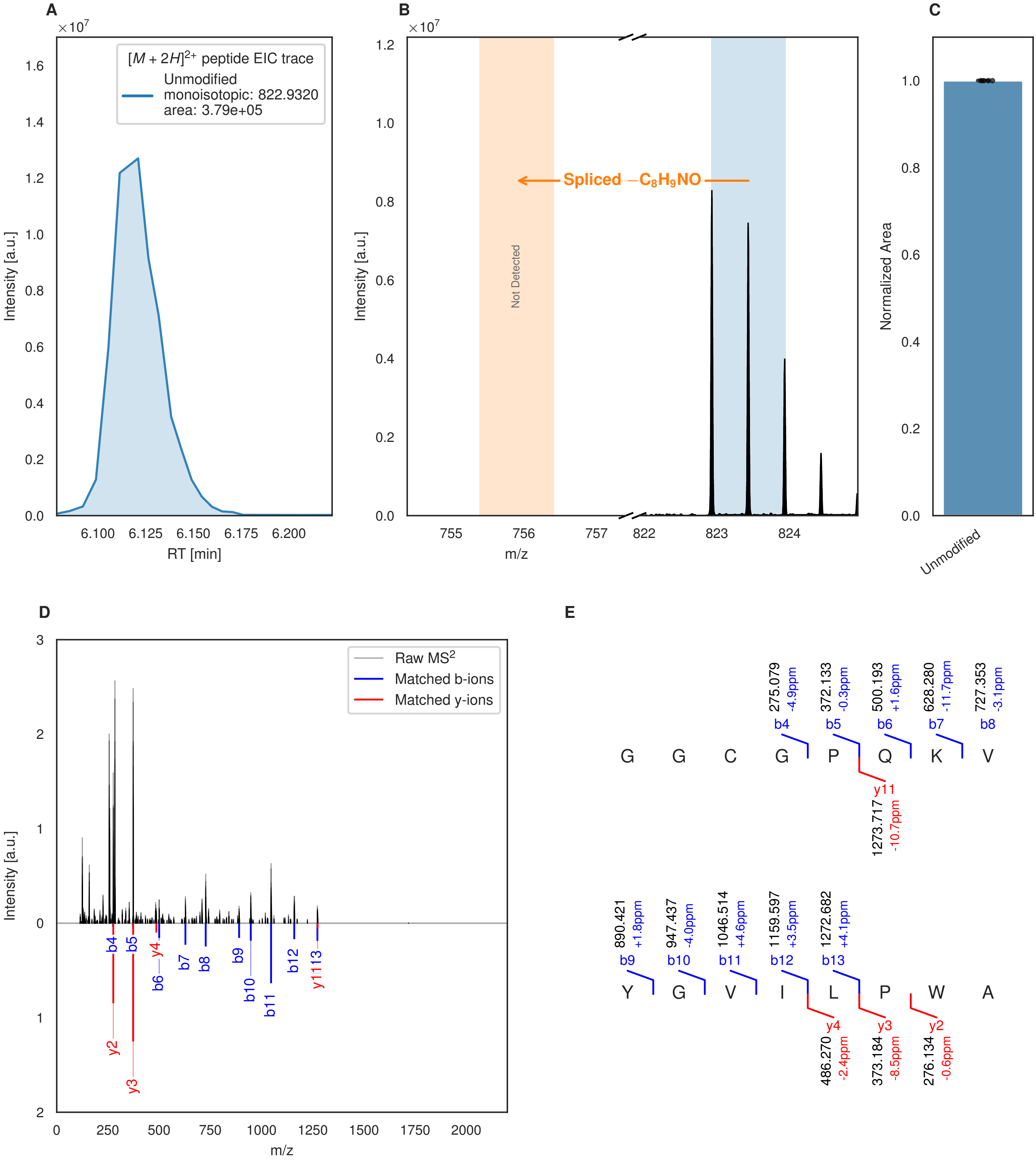


**Figure S87: Modification of the mutant_17 precursor peptide by HypXY.** LC-MS analysis of IMAC-purified precursor peptides from *E. coli* BL21(DE3)Tuner containing splicease-encoding plasmid TJ-C-689 (HypXY) and precursor peptide-encoding plasmid mutant_17 (RSTN1-HypAc-TEV-mutant-17) after expression at 24 °C for 22 h in 1.5 mL cultures after digestion with TEV. Data from n=4 independent biological replicates. Spliced amino acids have an x as a prefix. Spliced refers to a single tyramine excision. A representative experimental replicate is shown in panels **A, B.**

**A** Close-up plot of the representative extracted ion chromatogram (EIC) of the various peptide states; the blue trace corresponds to the Unmodified peptide. **B** Close-up plot of the m/z values of the peptides detected by the mass spectrometer, with the shaded colored areas corresponding to where the expected m/z values for the peptide m/z

(Caption continued on the next page.)

traces should occur, with the observed spectrum plotted in black. The Unmodified peptide(s) were found in the mass spectrum and the Spliced peptide(s) were not detected. **C** Bar plot showing the relative abundance of the various peptide states at charge z=2; the bars represent the average relative abundances normalized to total peak areas, with each individual experimental replicate value plotted as a black dot. **D** Mirrored MS$^2$ plot for the Unmodified peptide. In this mirror plot, the full fragmentation spectrum is plotted in black pointing upward, and the matched fragment b ions (blue) and y ions (red) are plotted pointing downward, mirrored against it, so that each matched fragment can be compared directly with the measured spectrum. **E** MS$^2$ spectrum of the fragments corresponding to the Unmodified peptide sequence, with matched b (blue, above) and y ions (red, below) plotted.

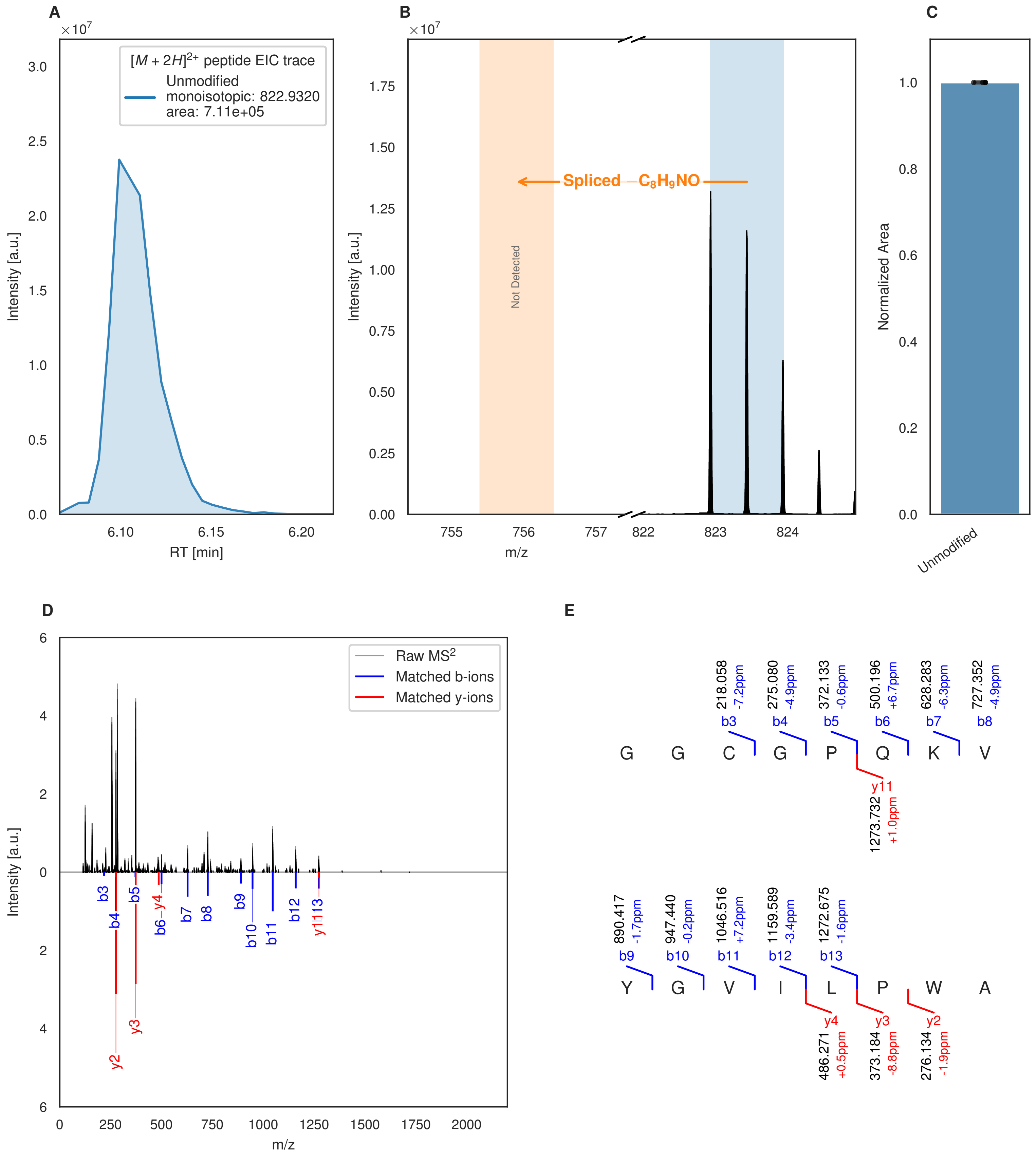


**Figure S88: Modification of the mutant_17 precursor peptide by PlpXY.** LC-MS analysis of IMAC-purified precursor peptides from *E. coli* BL21(DE3)Tuner containing splicease-encoding plasmid TJ-C-559 (PlpXY) and precursor peptide-encoding plasmid mutant_17 (RSTN1-PlpA3-TEV-mutant-17) after expression at 24 °C for 22 h in 1.5 mL cultures after digestion with TEV. Data from n=4 independent biological replicates. Spliced amino acids have an x as a prefix. Spliced refers to a single tyramine excision. A representative experimental replicate is shown in panels **A, B.**

**A** Close-up plot of the representative extracted ion chromatogram (EIC) of the various peptide states; the blue trace corresponds to the Unmodified peptide. **B** Close-up plot of the m/z values of the peptides detected by the mass spectrometer, with the shaded colored areas corresponding to where the expected m/z values for the peptide m/z

(Caption continued on the next page.)

traces should occur, with the observed spectrum plotted in black. The Unmodified peptide(s) were found in the mass spectrum and the Spliced peptide(s) were not detected. **C** Bar plot showing the relative abundance of the various peptide states at charge z=2; the bars represent the average relative abundances normalized to total peak areas, with each individual experimental replicate value plotted as a black dot. **D** Mirrored $MS^2$ plot for the Unmodified peptide. In this mirror plot, the full fragmentation spectrum is plotted in black pointing upward, and the matched fragment b ions (blue) and y ions (red) are plotted pointing downward, mirrored against it, so that each matched fragment can be compared directly with the measured spectrum. **E** $MS^2$ spectrum of the fragments corresponding to the Unmodified peptide sequence, with matched b (blue, above) and y ions (red, below) plotted.

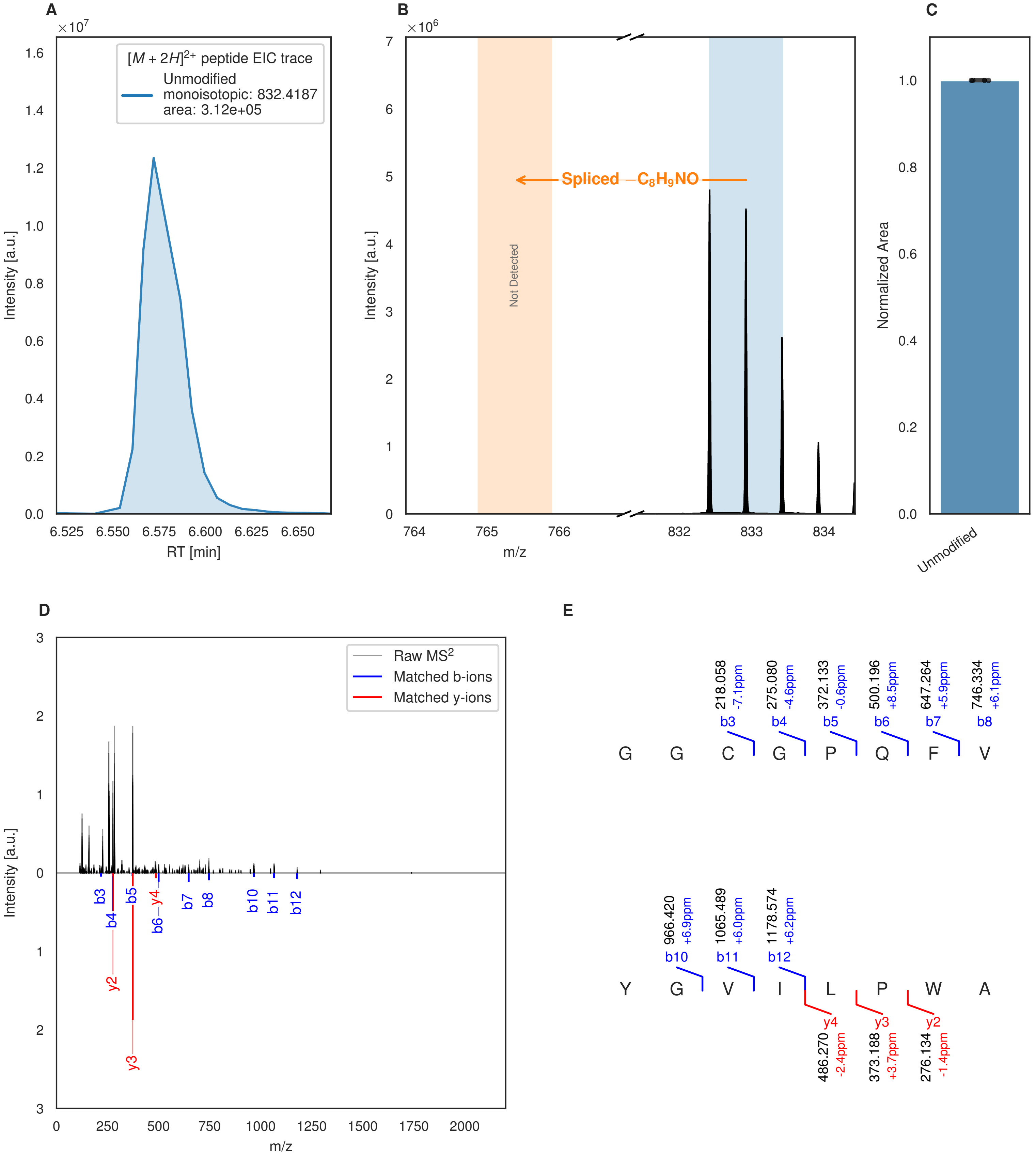


**Figure S89: Modification of the mutant_18 precursor peptide by HypXY.** LC-MS analysis of IMAC-purified precursor peptides from *E. coli* BL21(DE3)Tuner containing splicease-encoding plasmid TJ-C-689 (HypXY) and precursor peptide-encoding plasmid mutant_18 (RSTN1-HypAc-TEV-mutant-18) after expression at 24 °C for 22 h in 1.5 mL cultures after digestion with TEV. Data from n=4 independent biological replicates. Spliced amino acids have an x as a prefix. Spliced refers to a single tyramine excision. A representative experimental replicate is shown in panels **A, B.**

**A** Close-up plot of the representative extracted ion chromatogram (EIC) of the various peptide states; the blue trace corresponds to the Unmodified peptide. **B** Close-up plot of the m/z values of the peptides detected by the mass spectrometer, with the shaded colored areas corresponding to where the expected m/z values for the peptide m/z

(Caption continued on the next page.)

traces should occur, with the observed spectrum plotted in black. The Unmodified peptide(s) were found in the mass spectrum and the Spliced peptide(s) were not detected. **C** Bar plot showing the relative abundance of the various peptide states at charge z=2; the bars represent the average relative abundances normalized to total peak areas, with each individual experimental replicate value plotted as a black dot. **D** Mirrored $MS^2$ plot for the Unmodified peptide. In this mirror plot, the full fragmentation spectrum is plotted in black pointing upward, and the matched fragment b ions (blue) and y ions (red) are plotted pointing downward, mirrored against it, so that each matched fragment can be compared directly with the measured spectrum. **E** $MS^2$ spectrum of the fragments corresponding to the Unmodified peptide sequence, with matched b (blue, above) and y ions (red, below) plotted.

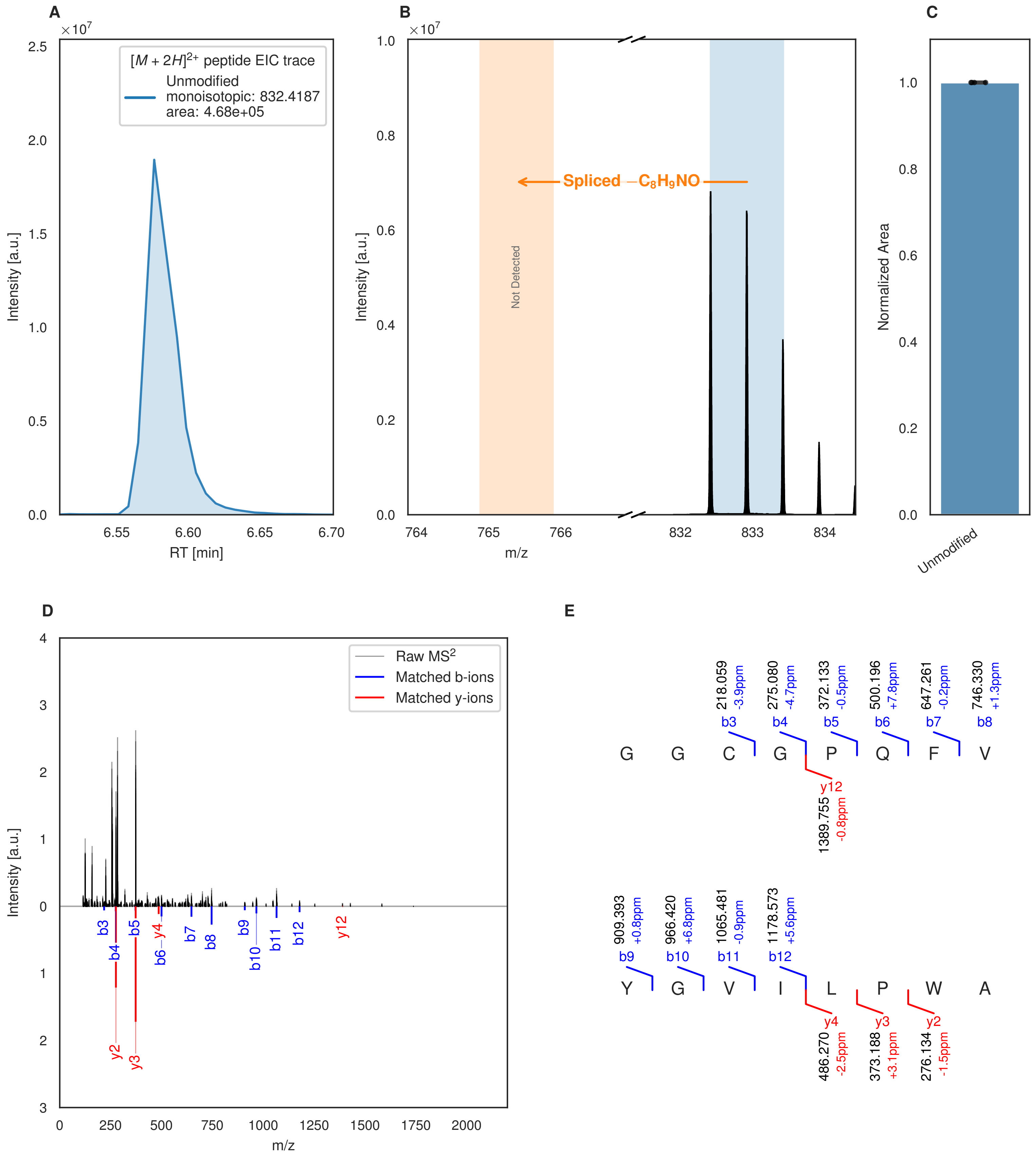


**Figure S90: Modification of the mutant_18 precursor peptide by PlpXY.** LC-MS analysis of IMAC-purified precursor peptides from *E. coli* BL21(DE3)Tuner containing splicease-encoding plasmid TJ-C-559 (PlpXY) and precursor peptide-encoding plasmid mutant_18 (RSTN1-PlpA3-TEV-mutant-18) after expression at 24 °C for 22 h in 1.5 mL cultures after digestion with TEV. Data from n=4 independent biological replicates. Spliced amino acids have an x as a prefix. Spliced refers to a single tyramine excision. A representative experimental replicate is shown in panels **A, B.**

**A** Close-up plot of the representative extracted ion chromatogram (EIC) of the various peptide states; the blue trace corresponds to the Unmodified peptide. **B** Close-up plot of the m/z values of the peptides detected by the mass spectrometer, with the shaded colored areas corresponding to where the expected m/z values for the peptide m/z

(Caption continued on the next page.)

traces should occur, with the observed spectrum plotted in black. The Unmodified peptide(s) were found in the mass spectrum and the Spliced peptide(s) were not detected. **C** Bar plot showing the relative abundance of the various peptide states at charge z=2; the bars represent the average relative abundances normalized to total peak areas, with each individual experimental replicate value plotted as a black dot. **D** Mirrored $MS^2$ plot for the Unmodified peptide. In this mirror plot, the full fragmentation spectrum is plotted in black pointing upward, and the matched fragment b ions (blue) and y ions (red) are plotted pointing downward, mirrored against it, so that each matched fragment can be compared directly with the measured spectrum. **E** $MS^2$ spectrum of the fragments corresponding to the Unmodified peptide sequence, with matched b (blue, above) and y ions (red, below) plotted.

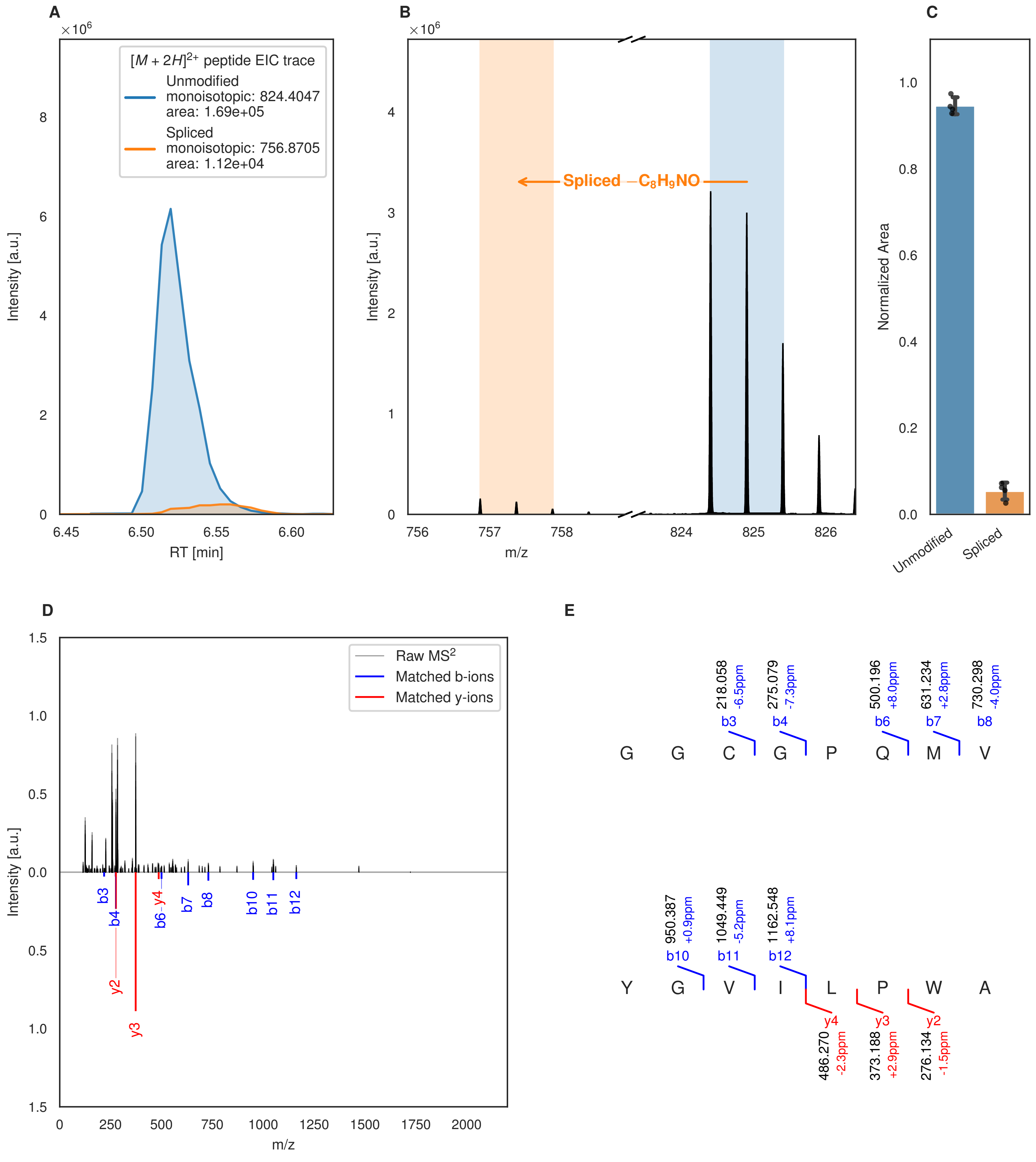

A
×10^6
[M + 2H]2+ peptide EIC trace
Unmodified
monoisotopic: 824.4047
area: 1.69e+05
Spliced
monoisotopic: 756.8705
area: 1.12e+04
Intensity [a.u.]
RT [min]
B
×10^6
Spliced −C8H9NO
Intensity [a.u.]
m/z
C
Normalized Area
Unmodified
Spliced
D
Raw MS2
Matched b-ions
Matched y-ions
Intensity [a.u.]
m/z
b3
b4
b6
y4
b7
b8
b10
b11
b12
y2
y3
E
218.058
-6.5ppm
b3
275.079
-7.3ppm
b4
500.196
+8.0ppm
b6
631.234
+2.8ppm
b7
730.298
-4.0ppm
b8
G G C G P Q M V
950.387
+0.9ppm
b10
1049.449
-5.2ppm
b11
1162.548
+8.1ppm
b12
Y G V I L P W A
y4
486.270
-2.3ppm
y3
373.188
+2.9ppm
y2
276.134
-1.5ppm

Figure S91 (figure continued from the previous page).

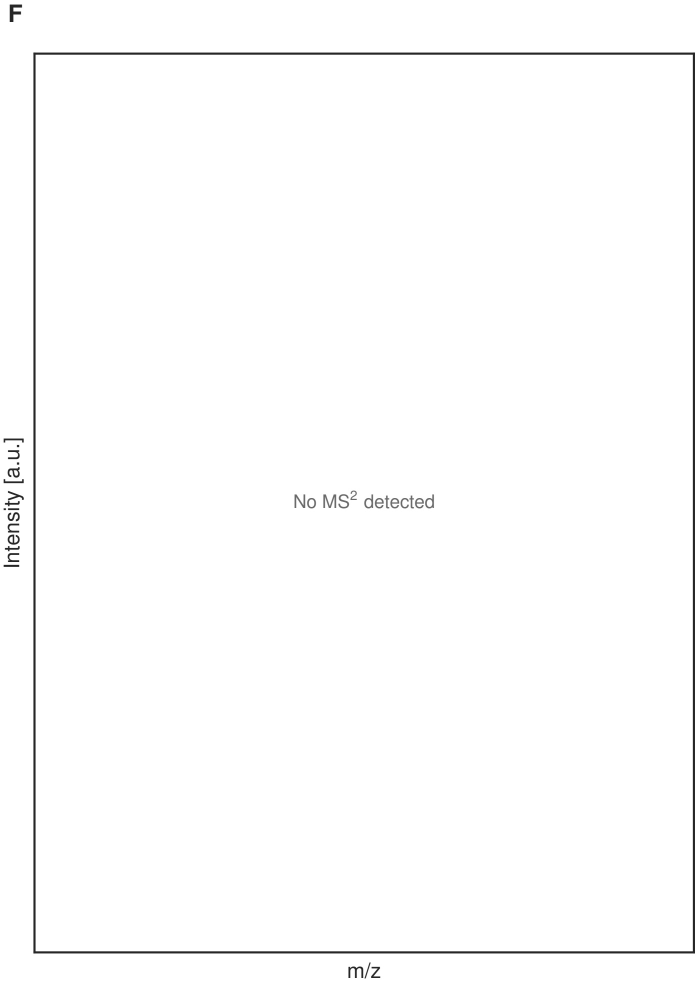


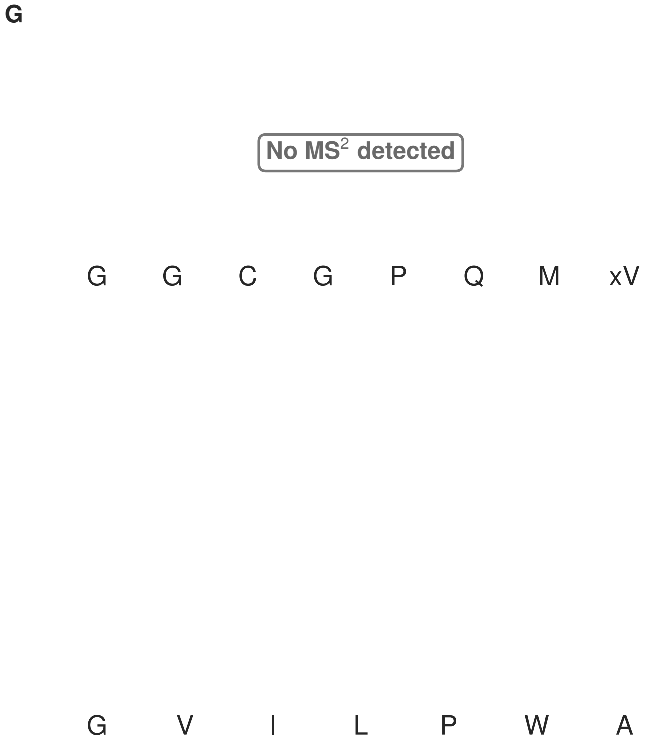


**Figure S91: Modification of the mutant_19 precursor peptide by HypXY.** LC-MS analysis of IMAC-purified precursor peptides from *E. coli* BL21(DE3)Tuner containing splicease-encoding plasmid TJ-C-689 (HypXY) and precursor peptide-encoding plasmid mutant_19 (RSTN1-HypAc-TEV-mutant-19) after expression at 24 °C for 22 h in 1.5 mL cultures after digestion with TEV. Data from n=4 independent biological replicates. Spliced amino acids have an x as a prefix. Spliced refers to a single tyramine excision. A representative experimental replicate is shown in panels **A, B.**

**A** Close-up plot of the representative extracted ion chromatogram (EIC) of the various peptide states; the blue trace corresponds to the Unmodified peptide; the orange trace corresponds to the Spliced peptide. **B** Close-up plot of the m/z values of the peptides detected by the mass spectrometer, with the shaded colored areas corresponding to where the expected m/z values for the peptide m/z traces should occur, with the observed spectrum plotted in black. The Unmodified, Spliced peptide(s) were found in the mass spectrum. **C** Bar plot showing the relative abundance of the various peptide states at charge z=2; the bars represent the average relative abundances normalized to total peak areas, with each individual experimental replicate value plotted as a black dot. **D** Mirrored $MS^2$ plot for the Unmodified peptide. In this mirror plot, the full fragmentation spectrum is plotted in black pointing upward, and the matched fragment b ions (blue) and y ions (red) are plotted pointing downward, mirrored against it, so that each matched fragment can be compared directly with the measured spectrum. **E** $MS^2$ spectrum of the fragments corresponding to the Unmodified peptide sequence, with matched b (blue, above) and y ions (red, below) plotted. **F** Mirrored $MS^2$ plot for the Spliced peptide. In this mirror plot, the full fragmentation spectrum is plotted in black pointing upward, and the matched fragment b ions (blue) and y ions (red) are plotted pointing downward, mirrored against it, so that each matched fragment can be compared directly with the measured spectrum. **G** $MS^2$ spectrum of the fragments corresponding to the Spliced peptide sequence, with matched b (blue, above) and y ions (red, below) plotted.

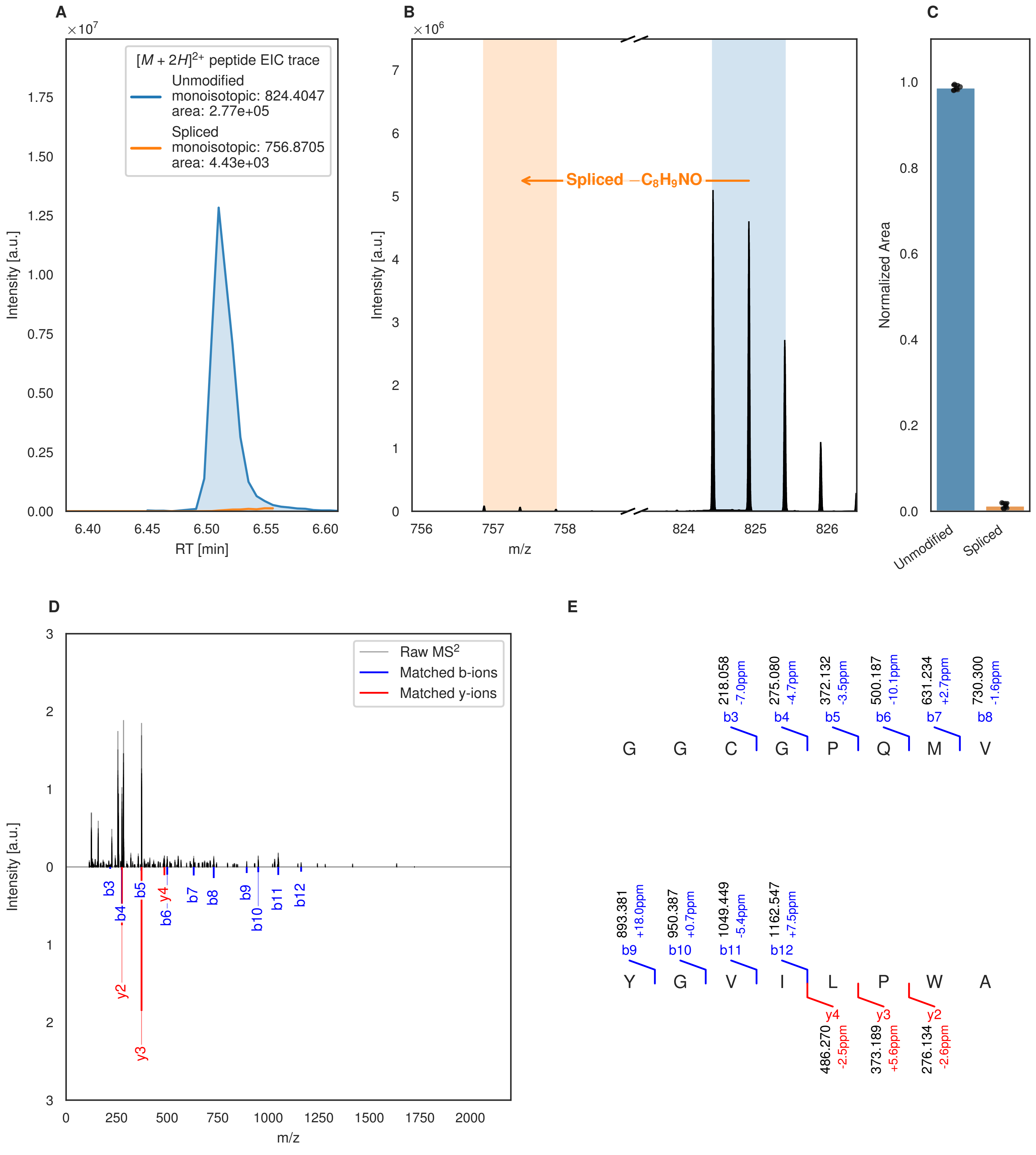
A
B
C
D
E
$[M+2H]^{2+}$ peptide EIC trace
Unmodified
monoisotopic: 824.4047
area: 2.77e+05
Spliced
monoisotopic: 756.8705
area: 4.43e+03
Intensity [a.u.]
RT [min]
Spliced $-C_8H_9NO$
m/z
Normalized Area
Unmodified
Spliced
Raw $MS^2$
Matched b-ions
Matched y-ions
218.058 -7.0ppm b3
275.080 -4.7ppm b4
372.132 -3.5ppm b5
500.187 -10.1ppm b6
631.234 +2.7ppm b7
730.300 -1.6ppm b8
G G C G P Q M V
893.381 +18.0ppm b9
950.387 +0.7ppm b10
1049.449 -5.4ppm b11
1162.547 +7.5ppm b12
Y G V I L P W A
y4 486.270 -2.5ppm
y3 373.189 +5.6ppm
y2 276.134 -2.6ppm

Figure S92 (figure continued from the previous page).

F

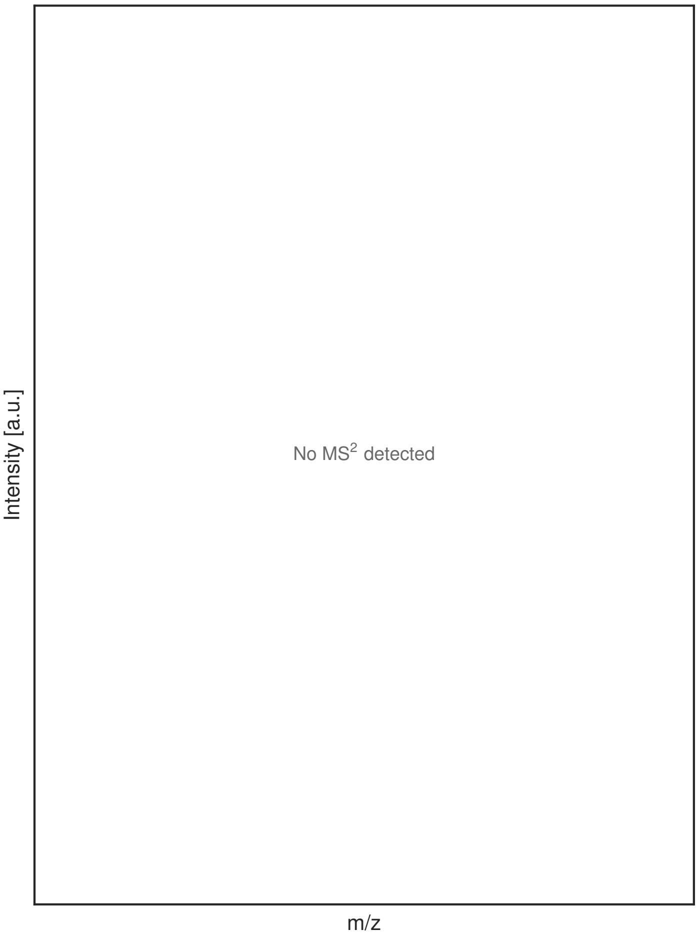


G

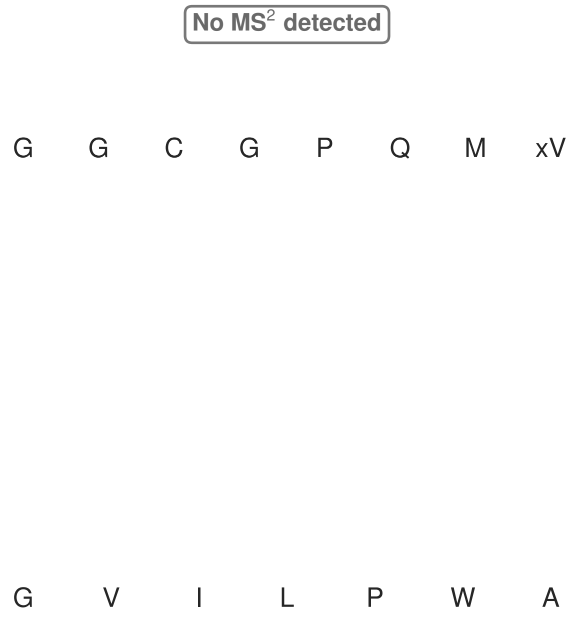


**Figure S92: Modification of the mutant_19 precursor peptide by PlpXY.** LC-MS analysis of IMAC-purified precursor peptides from *E. coli* BL21(DE3)Tuner containing splicease-encoding plasmid TJ-C-559 (PlpXY) and precursor peptide-encoding plasmid mutant_19 (RSTN1-PlpA3-TEV-mutant-19) after expression at 24 °C for 22 h in 1.5 mL cultures after digestion with TEV. Data from n=4 independent biological replicates. Spliced amino acids have an x as a prefix. Spliced refers to a single tyramine excision. A representative experimental replicate is shown in panels **A, B.**

**A** Close-up plot of the representative extracted ion chromatogram (EIC) of the various peptide states; the blue trace corresponds to the Unmodified peptide; the orange trace corresponds to the Spliced peptide. **B** Close-up plot of the m/z values of the peptides detected by the mass spectrometer, with the shaded colored areas corresponding to where the expected m/z values for the peptide m/z traces should occur, with the observed spectrum plotted in black. The Unmodified, Spliced peptide(s) were found in the mass spectrum. **C** Bar plot showing the relative abundance of the various peptide states at charge z=2; the bars represent the average relative abundances normalized to total peak areas, with each individual experimental replicate value plotted as a black dot. **D** Mirrored MS$^2$ plot for the Unmodified peptide. In this mirror plot, the full fragmentation spectrum is plotted in black pointing upward, and the matched fragment b ions (blue) and y ions (red) are plotted pointing downward, mirrored against it, so that each matched fragment can be compared directly with the measured spectrum. **E** MS$^2$ spectrum of the fragments corresponding to the Unmodified peptide sequence, with matched b (blue, above) and y ions (red, below) plotted. **F** Mirrored MS$^2$ plot for the Spliced peptide. In this mirror plot, the full fragmentation spectrum is plotted in black pointing upward, and the matched fragment b ions (blue) and y ions (red) are plotted pointing downward, mirrored against it, so that each matched fragment can be compared directly with the measured spectrum. **G** MS$^2$ spectrum of the fragments corresponding to the Spliced peptide sequence, with matched b (blue, above) and y ions (red, below) plotted.

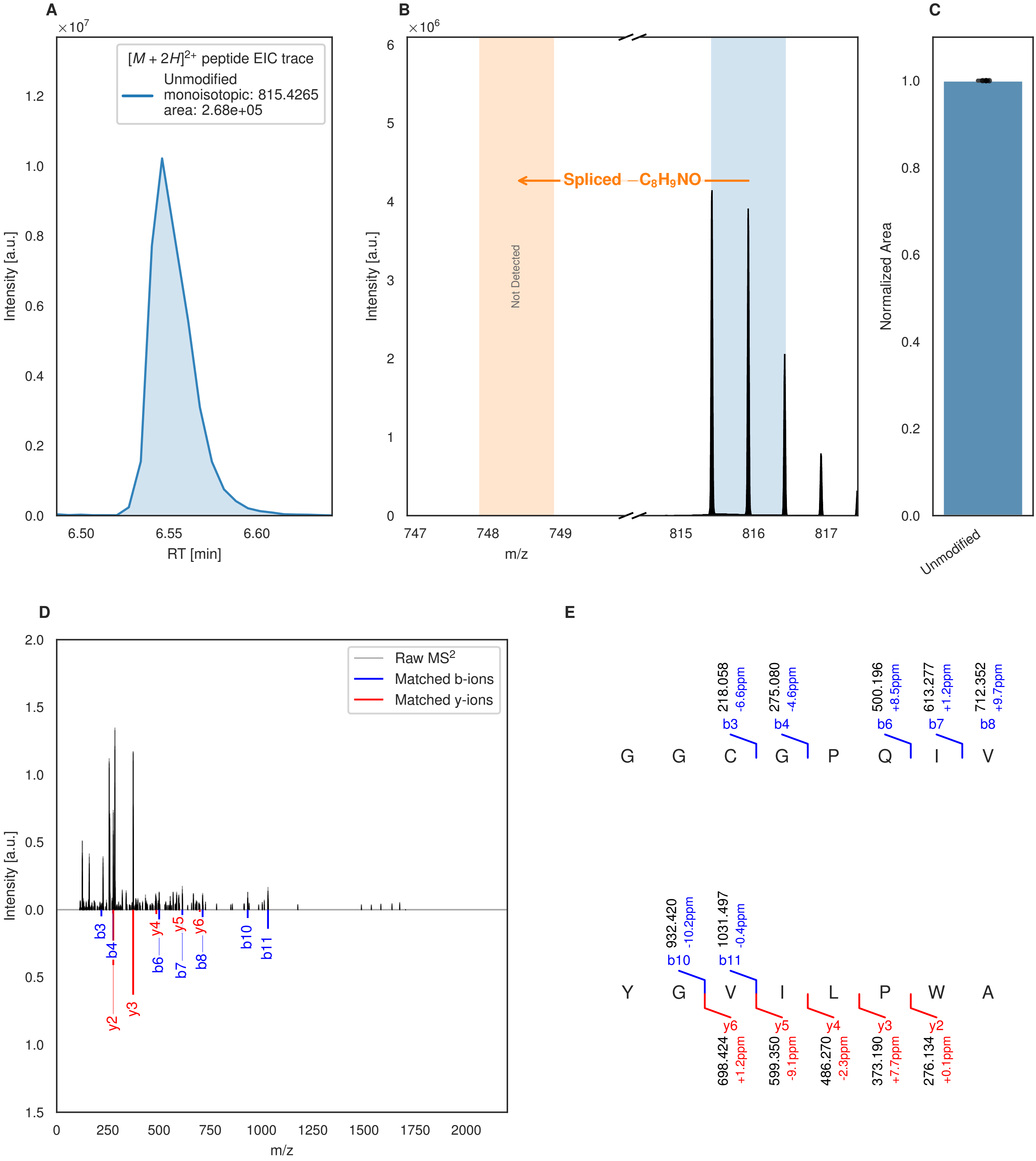


**Figure S93: Modification of the mutant_20 precursor peptide by HypXY.** LC-MS analysis of IMAC-purified precursor peptides from *E. coli* BL21(DE3)Tuner containing splicease-encoding plasmid TJ-C-689 (HypXY) and precursor peptide-encoding plasmid mutant_20 (RSTN1-HypAc-TEV-mutant-20) after expression at 24 °C for 22 h in 1.5 mL cultures after digestion with TEV. Data from n=4 independent biological replicates. Spliced amino acids have an x as a prefix. Spliced refers to a single tyramine excision. A representative experimental replicate is shown in panels **A, B.**

**A** Close-up plot of the representative extracted ion chromatogram (EIC) of the various peptide states; the blue trace corresponds to the Unmodified peptide. **B** Close-up plot of the m/z values of the peptides detected by the mass spectrometer, with the shaded colored areas corresponding to where the expected m/z values for the peptide m/z

(Caption continued on the next page.)

traces should occur, with the observed spectrum plotted in black. The Unmodified peptide(s) were found in the mass spectrum and the Spliced peptide(s) were not detected. **C** Bar plot showing the relative abundance of the various peptide states at charge z=2; the bars represent the average relative abundances normalized to total peak areas, with each individual experimental replicate value plotted as a black dot. **D** Mirrored $MS^2$ plot for the Unmodified peptide. In this mirror plot, the full fragmentation spectrum is plotted in black pointing upward, and the matched fragment b ions (blue) and y ions (red) are plotted pointing downward, mirrored against it, so that each matched fragment can be compared directly with the measured spectrum. **E** $MS^2$ spectrum of the fragments corresponding to the Unmodified peptide sequence, with matched b (blue, above) and y ions (red, below) plotted.

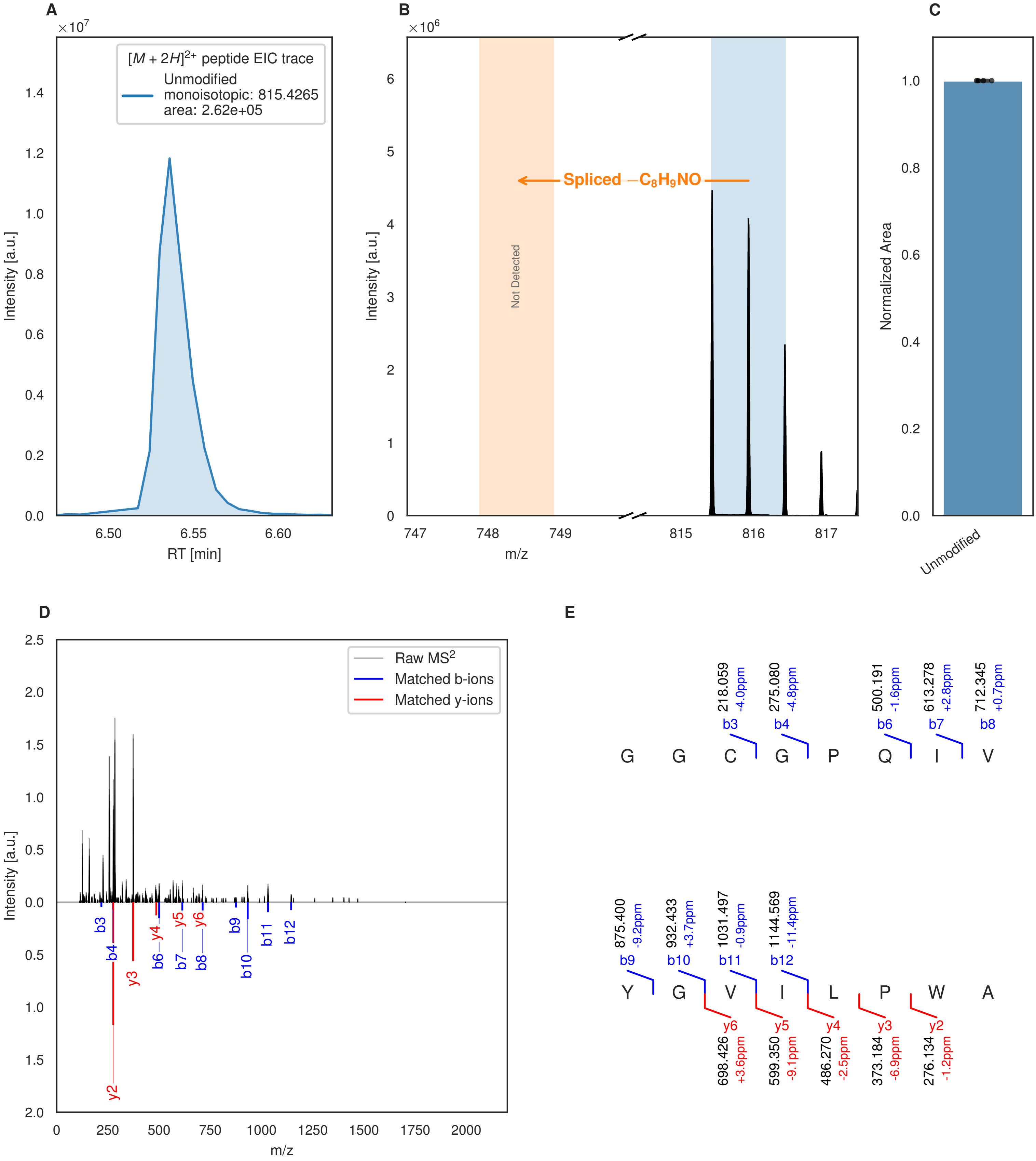


**Figure S94: Modification of the mutant_20 precursor peptide by PlpXY.** LC-MS analysis of IMAC-purified precursor peptides from *E. coli* BL21(DE3)Tuner containing splicease-encoding plasmid TJ-C-559 (PlpXY) and precursor peptide-encoding plasmid mutant_20 (RSTN1-PlpA3-TEV-mutant-20) after expression at 24 °C for 22 h in 1.5 mL cultures after digestion with TEV. Data from n=4 independent biological replicates. Spliced amino acids have an x as a prefix. Spliced refers to a single tyramine excision. A representative experimental replicate is shown in panels **A, B.**

**A** Close-up plot of the representative extracted ion chromatogram (EIC) of the various peptide states; the blue trace corresponds to the Unmodified peptide. **B** Close-up plot of the m/z values of the peptides detected by the mass spectrometer, with the shaded colored areas corresponding to where the expected m/z values for the peptide m/z

(Caption continued on the next page.)

traces should occur, with the observed spectrum plotted in black. The Unmodified peptide(s) were found in the mass spectrum and the Spliced peptide(s) were not detected. **C** Bar plot showing the relative abundance of the various peptide states at charge z=2; the bars represent the average relative abundances normalized to total peak areas, with each individual experimental replicate value plotted as a black dot. **D** Mirrored $MS^2$ plot for the Unmodified peptide. In this mirror plot, the full fragmentation spectrum is plotted in black pointing upward, and the matched fragment b ions (blue) and y ions (red) are plotted pointing downward, mirrored against it, so that each matched fragment can be compared directly with the measured spectrum. **E** $MS^2$ spectrum of the fragments corresponding to the Unmodified peptide sequence, with matched b (blue, above) and y ions (red, below) plotted.

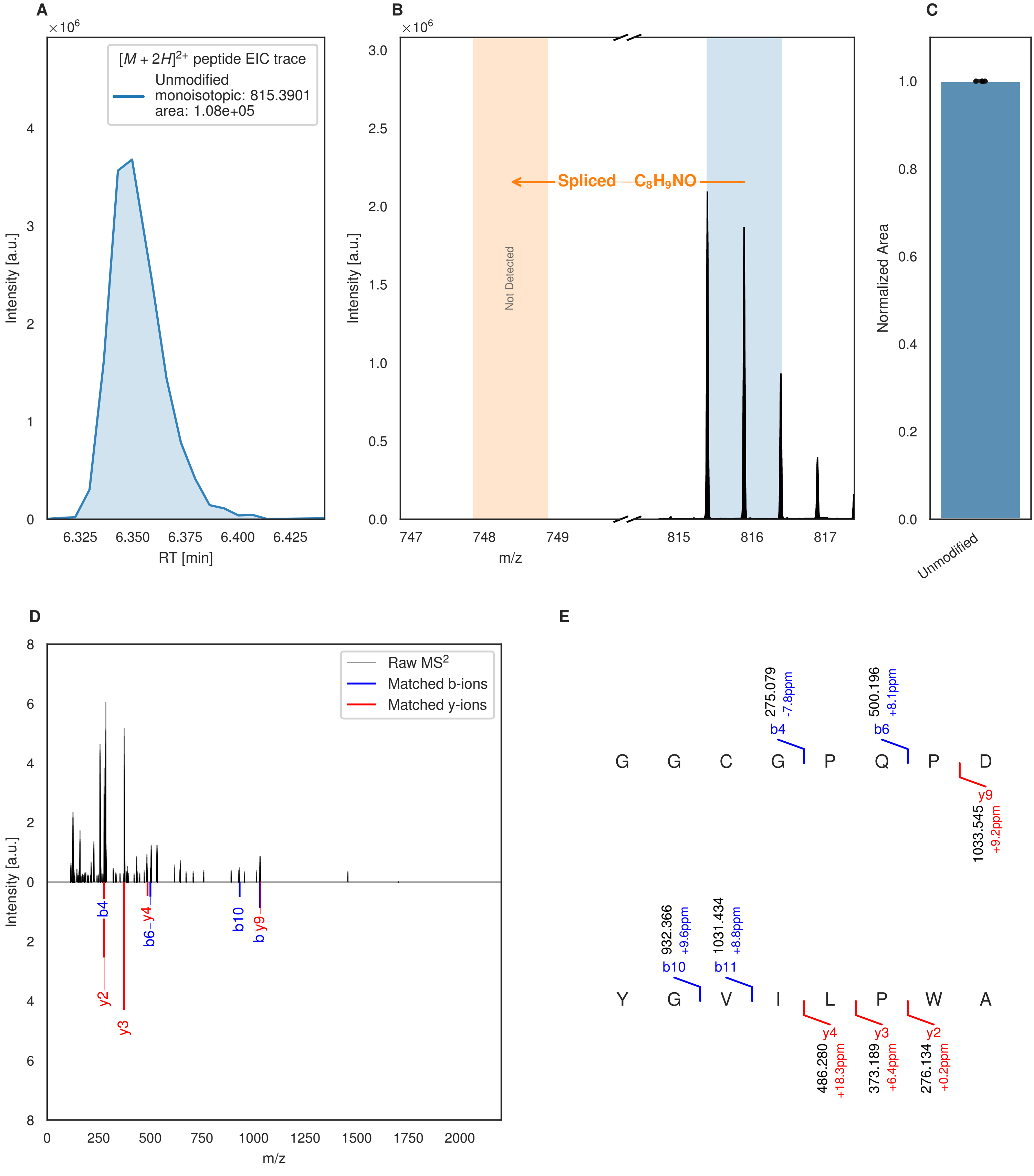


**Figure S95: Modification of the mutant_21 precursor peptide by HypXY.** LC-MS analysis of IMAC-purified precursor peptides from *E. coli* BL21(DE3)Tuner containing splicease-encoding plasmid TJ-C-689 (HypXY) and precursor peptide-encoding plasmid mutant_21 (RSTN1-HypAc-TEV-mutant-21) after expression at 24 °C for 22 h in 1.5 mL cultures after digestion with TEV. Data from n=4 independent biological replicates. Spliced amino acids have an x as a prefix. Spliced refers to a single tyramine excision. A representative experimental replicate is shown in panels **A, B.**

**A** Close-up plot of the representative extracted ion chromatogram (EIC) of the various peptide states; the blue trace corresponds to the Unmodified peptide. **B** Close-up plot of the m/z values of the peptides detected by the mass spectrometer, with the shaded colored areas corresponding to where the expected m/z values for the peptide m/z

(Caption continued on the next page.)

traces should occur, with the observed spectrum plotted in black. The Unmodified peptide(s) were found in the mass spectrum and the Spliced peptide(s) were not detected. **C** Bar plot showing the relative abundance of the various peptide states at charge z=2; the bars represent the average relative abundances normalized to total peak areas, with each individual experimental replicate value plotted as a black dot. **D** Mirrored MS$^2$ plot for the Unmodified peptide. In this mirror plot, the full fragmentation spectrum is plotted in black pointing upward, and the matched fragment b ions (blue) and y ions (red) are plotted pointing downward, mirrored against it, so that each matched fragment can be compared directly with the measured spectrum. **E** MS$^2$ spectrum of the fragments corresponding to the Unmodified peptide sequence, with matched b (blue, above) and y ions (red, below) plotted.

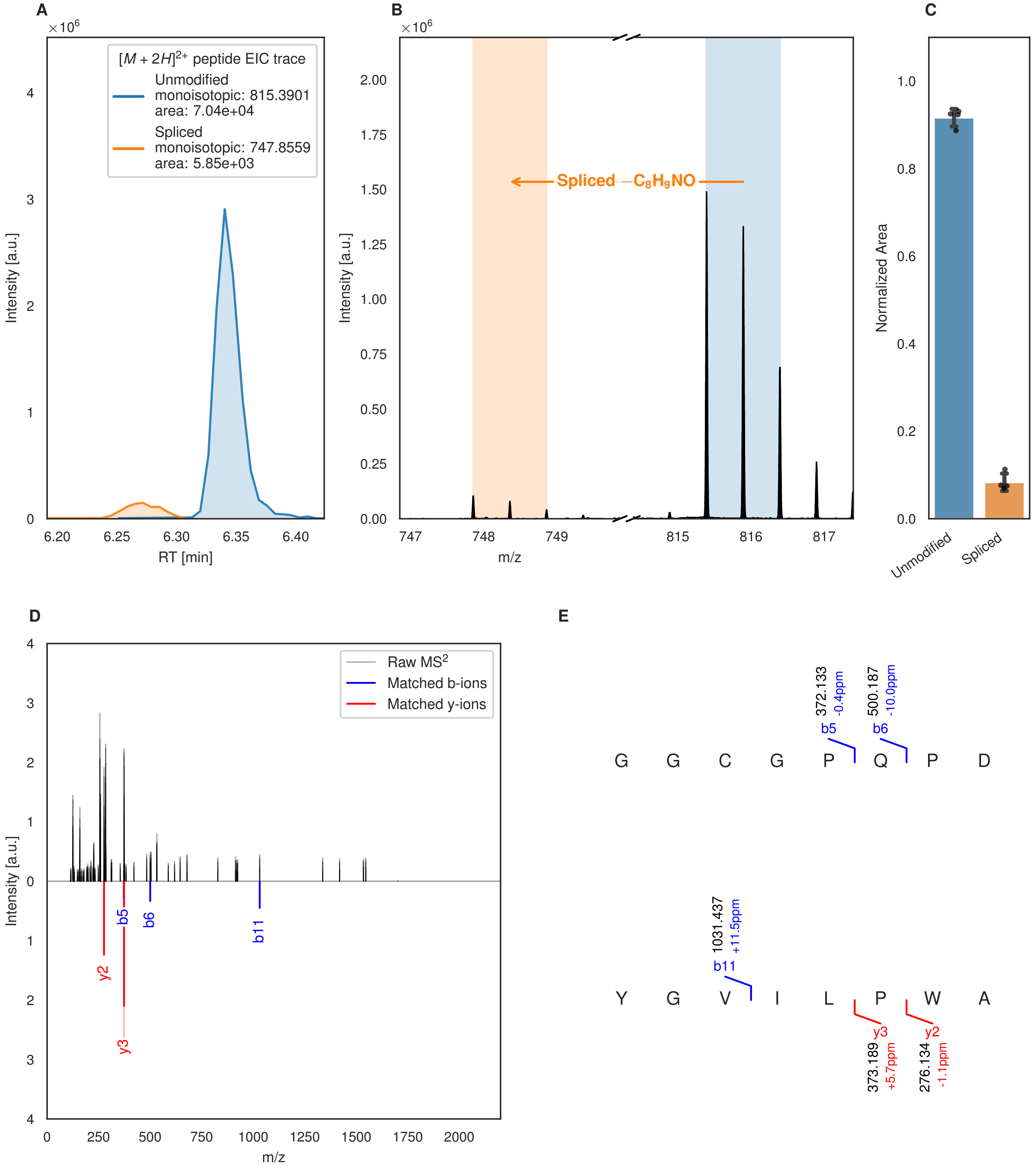
A
×10^6
[M + 2H]2+ peptide EIC trace
Unmodified
monoisotopic: 815.3901
area: 7.04e+04
Spliced
monoisotopic: 747.8559
area: 5.85e+03
Intensity [a.u.]
RT [min]
B
×10^6
Spliced −C8H9NO
Intensity [a.u.]
m/z
C
Normalized Area
Unmodified
Spliced
D
Raw MS2
Matched b-ions
Matched y-ions
Intensity [a.u.]
m/z
y2
b5
b6
b11
y3
E
372.133
-0.4ppm
500.187
-10.0ppm
b5
b6
G G C G P Q P D
1031.437
+11.5ppm
b11
Y G V I L P W A
y3
y2
373.189
+5.7ppm
276.134
-1.1ppm

Figure S96 (figure continued from the previous page).

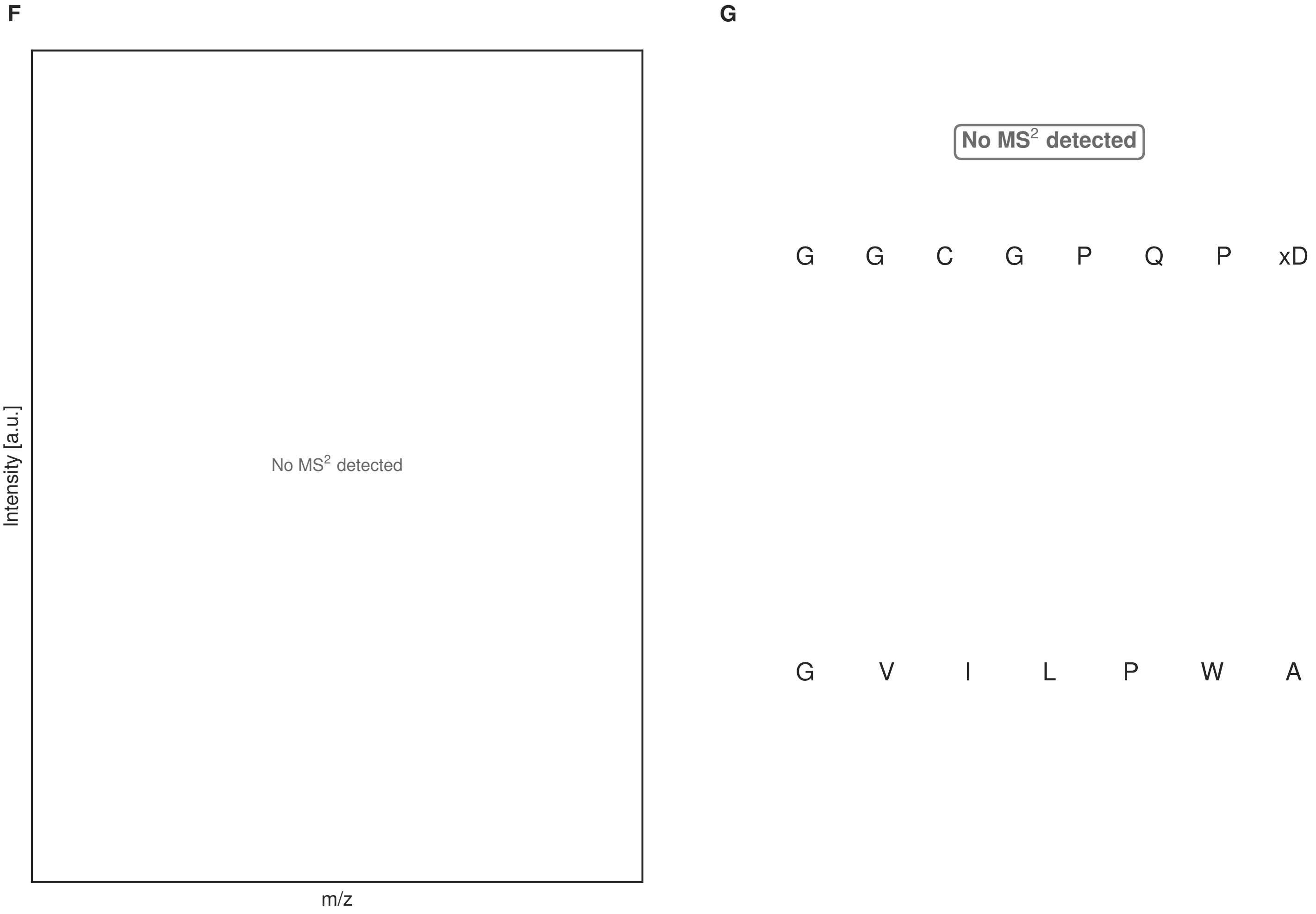


**Figure S96: Modification of the mutant_21 precursor peptide by PlpXY.** LC-MS analysis of IMAC-purified precursor peptides from *E. coli* BL21(DE3)Tuner containing splicease-encoding plasmid TJ-C-559 (PlpXY) and precursor peptide-encoding plasmid mutant_21 (RSTN1-PlpA3-TEV-mutant-21) after expression at 24 °C for 22 h in 1.5 mL cultures after digestion with TEV. Data from n=4 independent biological replicates. Spliced amino acids have an x as a prefix. Spliced refers to a single tyramine excision. A representative experimental replicate is shown in panels **A, B.**

**A** Close-up plot of the representative extracted ion chromatogram (EIC) of the various peptide states; the blue trace corresponds to the Unmodified peptide; the orange trace corresponds to the Spliced peptide. **B** Close-up plot of the m/z values of the peptides detected by the mass spectrometer, with the shaded colored areas corresponding to where the expected m/z values for the peptide m/z traces should occur, with the observed spectrum plotted in black. The Unmodified, Spliced peptide(s) were found in the mass spectrum. **C** Bar plot showing the relative abundance of the various peptide states at charge z=2; the bars represent the average relative abundances normalized to total peak areas, with each individual experimental replicate value plotted as a black dot. **D** Mirrored MS$^2$ plot for the Unmodified peptide. In this mirror plot, the full fragmentation spectrum is plotted in black pointing upward, and the matched fragment b ions (blue) and y ions (red) are plotted pointing downward, mirrored against it, so that each matched fragment can be compared directly with the measured spectrum. **E** MS$^2$ spectrum of the fragments corresponding to the Unmodified peptide sequence, with matched b (blue, above) and y ions (red, below) plotted. **F** Mirrored MS$^2$ plot for the Spliced peptide. In this mirror plot, the full fragmentation spectrum is plotted in black pointing upward, and the matched fragment b ions (blue) and y ions (red) are plotted pointing downward, mirrored against it, so that each matched fragment can be compared directly with the measured spectrum. **G** MS$^2$ spectrum of the fragments corresponding to the Spliced peptide sequence, with matched b (blue, above) and y ions (red, below) plotted.

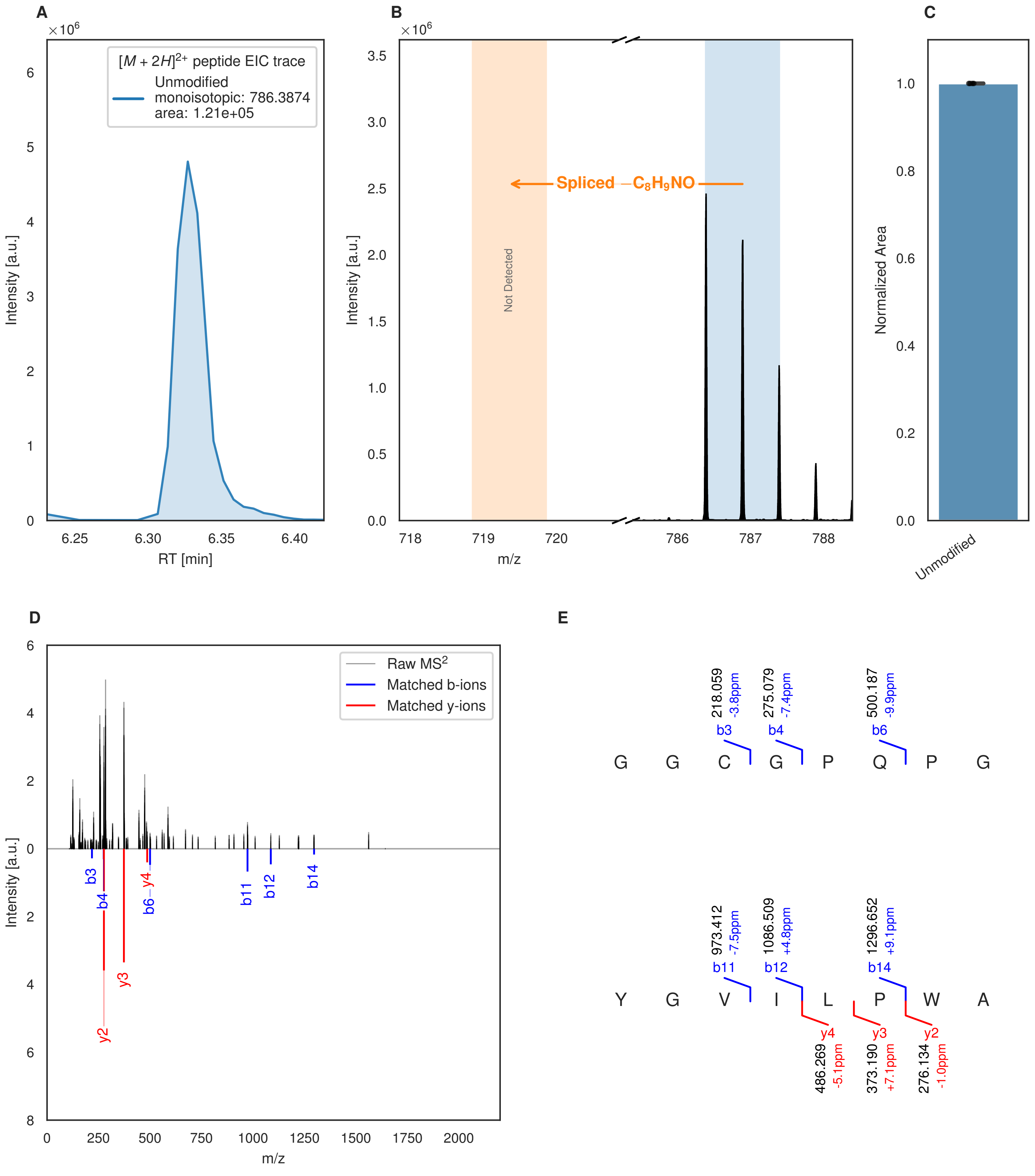


**Figure S97: Modification of the mutant_22 precursor peptide by HypXY.** LC-MS analysis of IMAC-purified precursor peptides from *E. coli* BL21(DE3)Tuner containing splicease-encoding plasmid TJ-C-689 (HypXY) and precursor peptide-encoding plasmid mutant_22 (RSTN1-HypAc-TEV-mutant-22) after expression at 24 °C for 22 h in 1.5 mL cultures after digestion with TEV. Data from n=4 independent biological replicates. Spliced amino acids have an x as a prefix. Spliced refers to a single tyramine excision. A representative experimental replicate is shown in panels **A, B.**

**A** Close-up plot of the representative extracted ion chromatogram (EIC) of the various peptide states; the blue trace corresponds to the Unmodified peptide. **B** Close-up plot of the m/z values of the peptides detected by the mass spectrometer, with the shaded colored areas corresponding to where the expected m/z values for the peptide m/z

(Caption continued on the next page.)

traces should occur, with the observed spectrum plotted in black. The Unmodified peptide(s) were found in the mass spectrum and the Spliced peptide(s) were not detected. **C** Bar plot showing the relative abundance of the various peptide states at charge z=2; the bars represent the average relative abundances normalized to total peak areas, with each individual experimental replicate value plotted as a black dot. **D** Mirrored $MS^2$ plot for the Unmodified peptide. In this mirror plot, the full fragmentation spectrum is plotted in black pointing upward, and the matched fragment b ions (blue) and y ions (red) are plotted pointing downward, mirrored against it, so that each matched fragment can be compared directly with the measured spectrum. **E** $MS^2$ spectrum of the fragments corresponding to the Unmodified peptide sequence, with matched b (blue, above) and y ions (red, below) plotted.

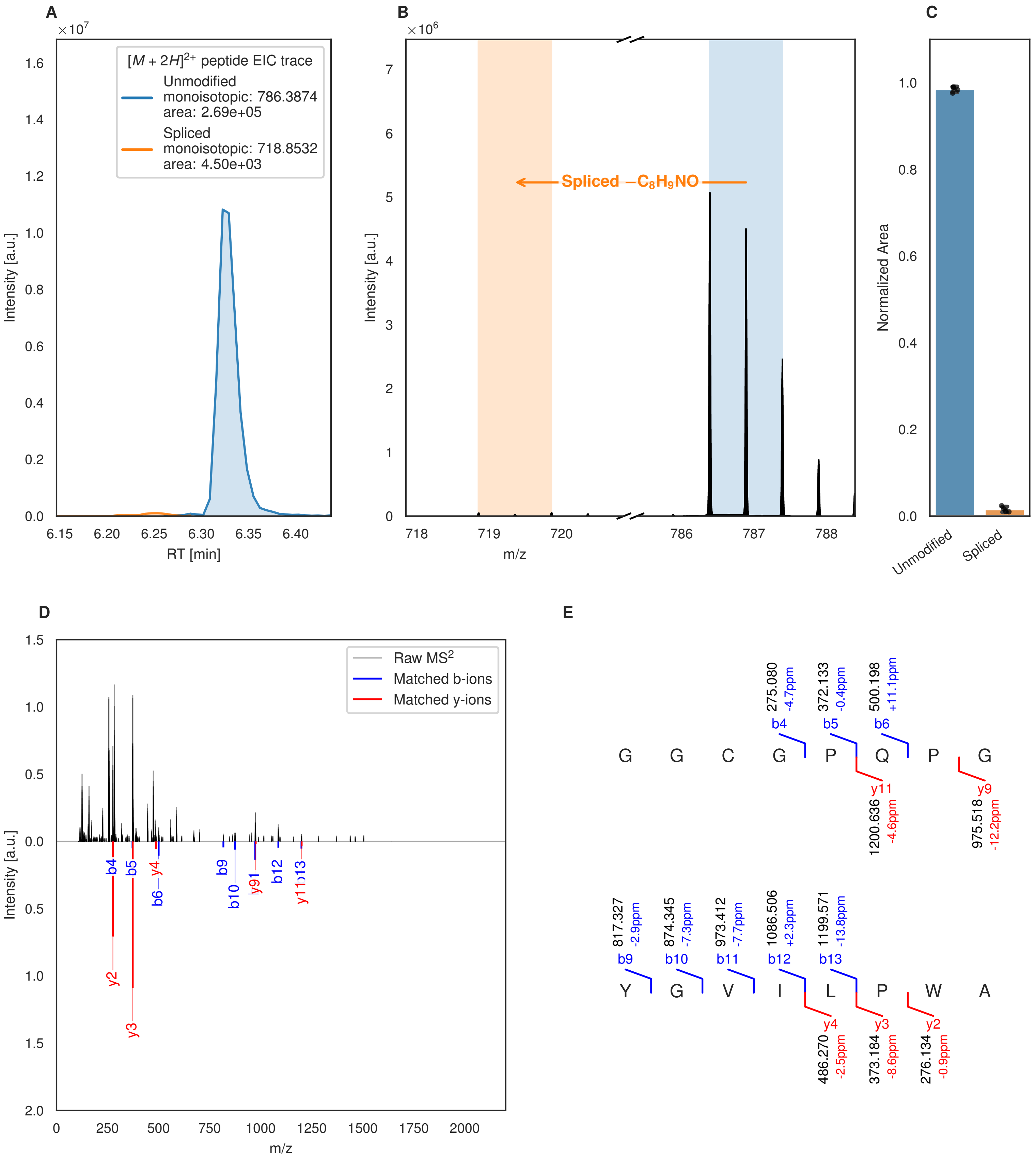
A
×10^7
[M + 2H]2+ peptide EIC trace
Unmodified
monoisotopic: 786.3874
area: 2.69e+05
Spliced
monoisotopic: 718.8532
area: 4.50e+03
Intensity [a.u.]
RT [min]
B
×10^6
Spliced −C8H9NO
Intensity [a.u.]
m/z
C
Normalized Area
Unmodified
Spliced
D
Raw MS2
Matched b-ions
Matched y-ions
Intensity [a.u.]
m/z
E
G G C G P Q P G
b4 275.080 -4.7ppm
b5 372.133 -0.4ppm
b6 500.198 +11.1ppm
y11 1200.636 -4.6ppm
y9 975.518 -12.2ppm
Y G V I L P W A
b9 817.327 -2.9ppm
b10 874.345 -7.3ppm
b11 973.412 -7.7ppm
b12 1086.506 +2.3ppm
b13 1199.571 -13.8ppm
y4 486.270 -2.5ppm
y3 373.184 -8.6ppm
y2 276.134 -0.9ppm

Figure S98 (figure continued from the previous page).

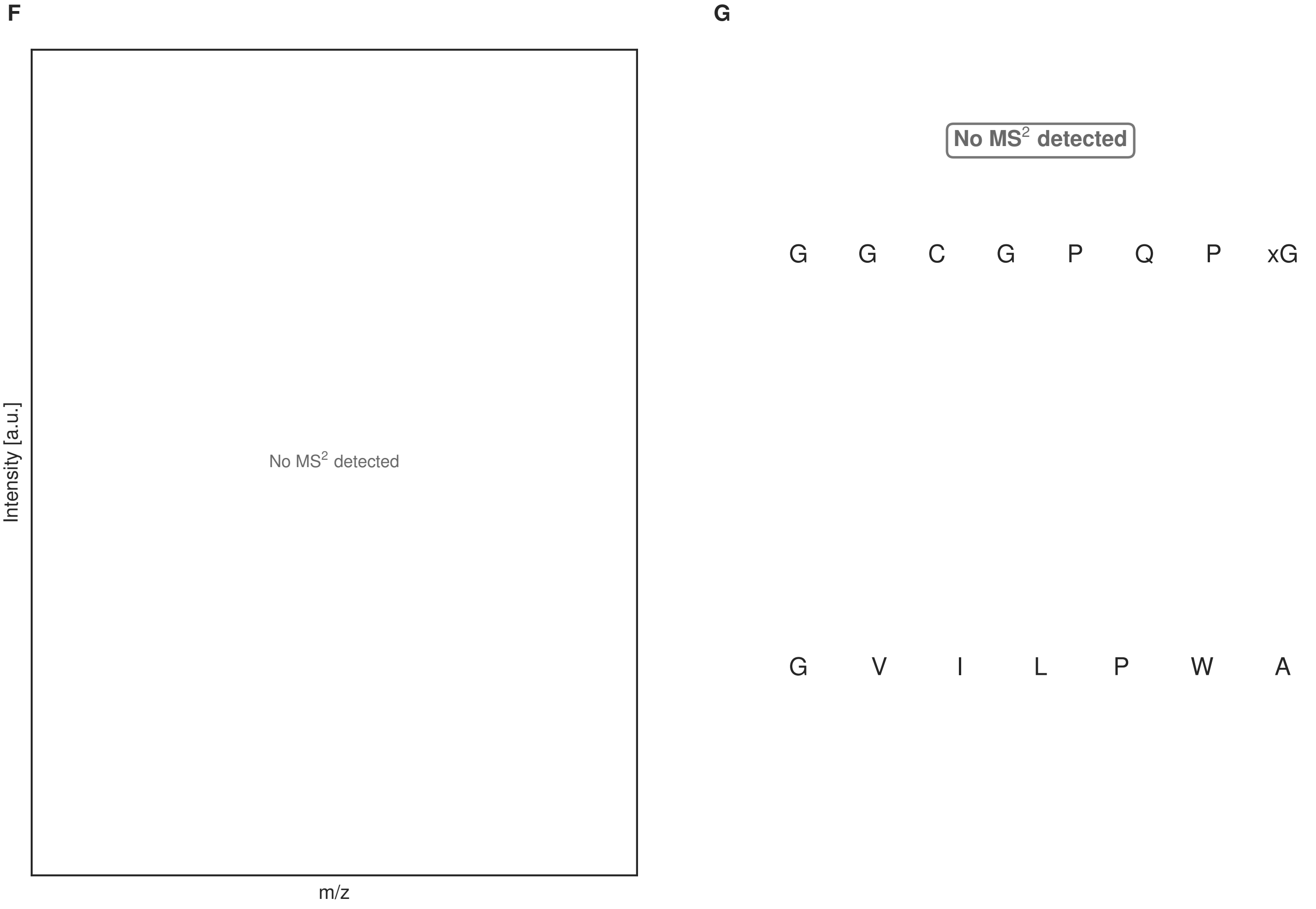


**Figure S98: Modification of the mutant_22 precursor peptide by PlpXY.** LC-MS analysis of IMAC-purified precursor peptides from *E. coli* BL21(DE3)Tuner containing splicease-encoding plasmid TJ-C-559 (PlpXY) and precursor peptide-encoding plasmid mutant_22 (RSTN1-PlpA3-TEV-mutant-22) after expression at 24 °C for 22 h in 1.5 mL cultures after digestion with TEV. Data from n=4 independent biological replicates. Spliced amino acids have an x as a prefix. Spliced refers to a single tyramine excision. A representative experimental replicate is shown in panels **A, B.**

**A** Close-up plot of the representative extracted ion chromatogram (EIC) of the various peptide states; the blue trace corresponds to the Unmodified peptide; the orange trace corresponds to the Spliced peptide. **B** Close-up plot of the m/z values of the peptides detected by the mass spectrometer, with the shaded colored areas corresponding to where the expected m/z values for the peptide m/z traces should occur, with the observed spectrum plotted in black. The Unmodified, Spliced peptide(s) were found in the mass spectrum. **C** Bar plot showing the relative abundance of the various peptide states at charge z=2; the bars represent the average relative abundances normalized to total peak areas, with each individual experimental replicate value plotted as a black dot. **D** Mirrored MS$^2$ plot for the Unmodified peptide. In this mirror plot, the full fragmentation spectrum is plotted in black pointing upward, and the matched fragment b ions (blue) and y ions (red) are plotted pointing downward, mirrored against it, so that each matched fragment can be compared directly with the measured spectrum. **E** MS$^2$ spectrum of the fragments corresponding to the Unmodified peptide sequence, with matched b (blue, above) and y ions (red, below) plotted. **F** Mirrored MS$^2$ plot for the Spliced peptide. In this mirror plot, the full fragmentation spectrum is plotted in black pointing upward, and the matched fragment b ions (blue) and y ions (red) are plotted pointing downward, mirrored against it, so that each matched fragment can be compared directly with the measured spectrum. **G** MS$^2$ spectrum of the fragments corresponding to the Spliced peptide sequence, with matched b (blue, above) and y ions (red, below) plotted.

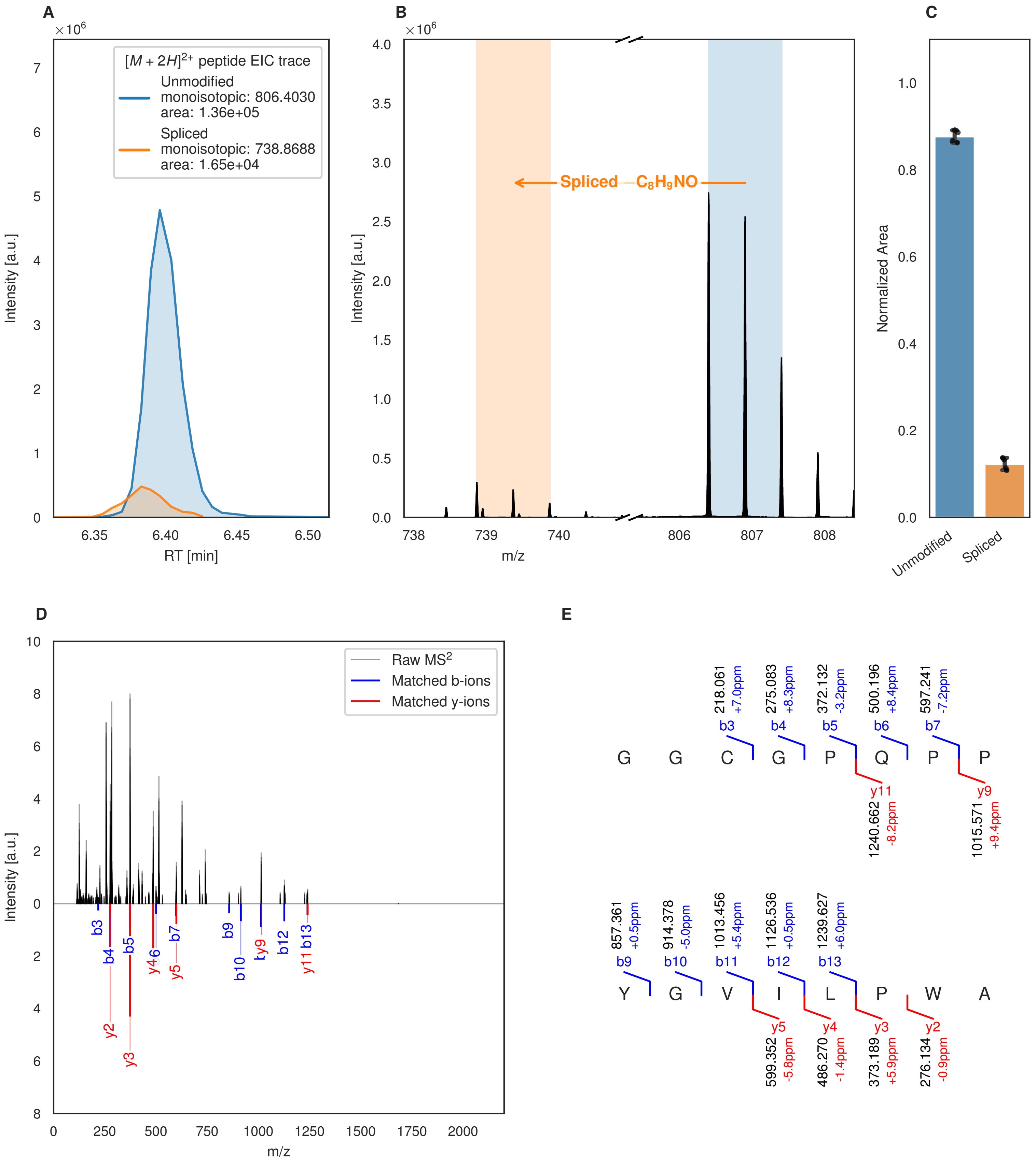
A
B
C
D
E
[M + 2H]2+ peptide EIC trace
Unmodified
monoisotopic: 806.4030
area: 1.36e+05
Spliced
monoisotopic: 738.8688
area: 1.65e+04
Intensity [a.u.]
RT [min]
Spliced −C8H9NO
m/z
Normalized Area
Unmodified
Spliced
Raw MS2
Matched b-ions
Matched y-ions
G G C G P Q P P
Y G V I L P W A
218.061 +7.0ppm b3
275.083 +8.3ppm b4
372.132 -3.2ppm b5
500.196 +8.4ppm b6
597.241 -7.2ppm b7
y11 1240.662 -8.2ppm
y9 1015.571 +9.4ppm
857.361 +0.5ppm b9
914.378 -5.0ppm b10
1013.456 +5.4ppm b11
1126.536 +0.5ppm b12
1239.627 +6.0ppm b13
y5 599.352 -5.8ppm
y4 486.270 -1.4ppm
y3 373.189 +5.9ppm
y2 276.134 -0.9ppm

Figure S99 (figure continued from the previous page).

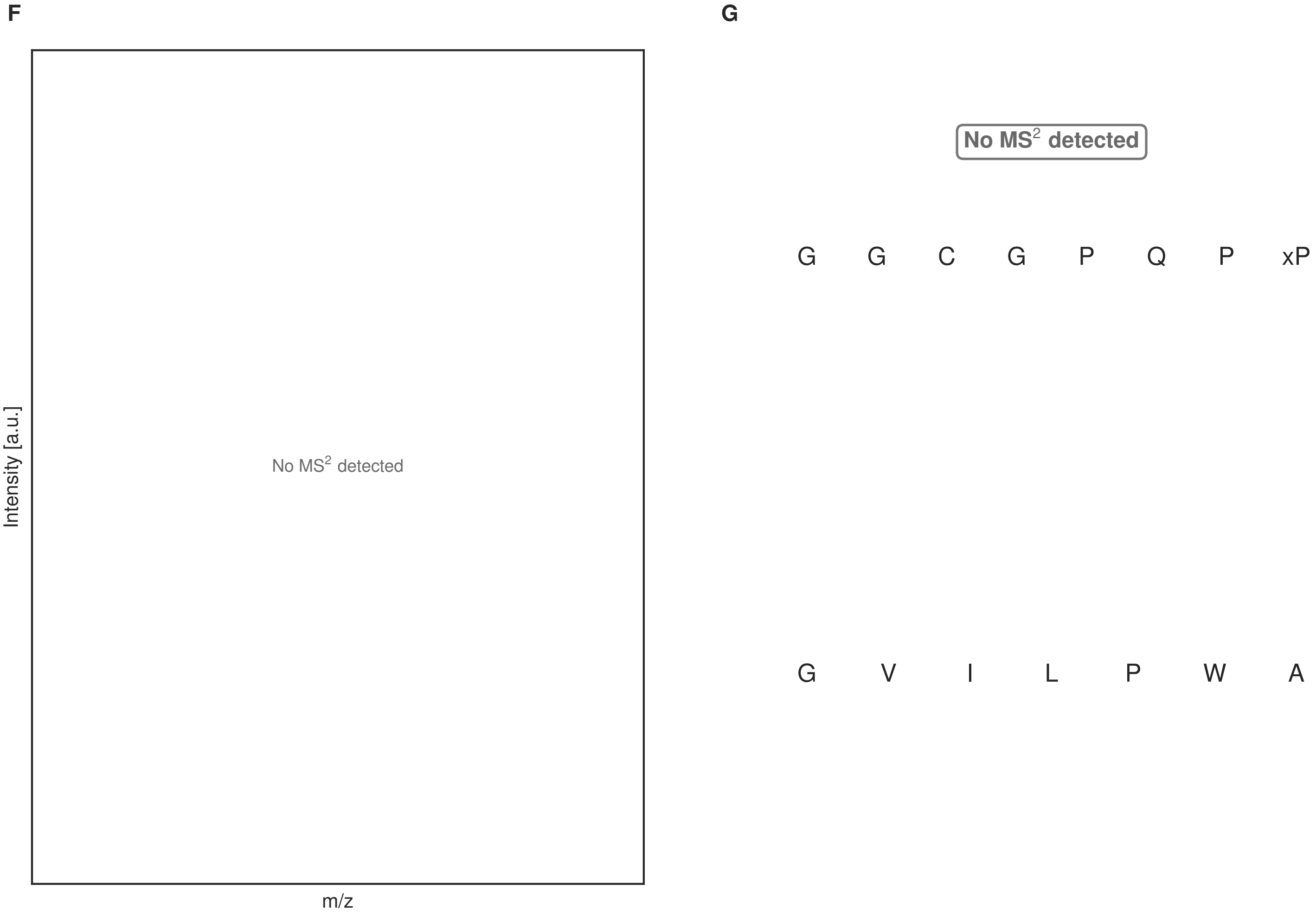


**Figure S99: Modification of the mutant_23 precursor peptide by HypXY.** LC-MS analysis of IMAC-purified precursor peptides from *E. coli* BL21(DE3)Tuner containing splicease-encoding plasmid TJ-C-689 (HypXY) and precursor peptide-encoding plasmid mutant_23 (RSTN1-HypAc-TEV-mutant-23) after expression at 24 °C for 22 h in 1.5 mL cultures after digestion with TEV. Data from n=4 independent biological replicates. Spliced amino acids have an x as a prefix. Spliced refers to a single tyramine excision. A representative experimental replicate is shown in panels **A, B.**

**A** Close-up plot of the representative extracted ion chromatogram (EIC) of the various peptide states; the blue trace corresponds to the Unmodified peptide; the orange trace corresponds to the Spliced peptide. **B** Close-up plot of the m/z values of the peptides detected by the mass spectrometer, with the shaded colored areas corresponding to where the expected m/z values for the peptide m/z traces should occur, with the observed spectrum plotted in black. The Unmodified, Spliced peptide(s) were found in the mass spectrum. **C** Bar plot showing the relative abundance of the various peptide states at charge z=2; the bars represent the average relative abundances normalized to total peak areas, with each individual experimental replicate value plotted as a black dot. **D** Mirrored $MS^2$ plot for the Unmodified peptide. In this mirror plot, the full fragmentation spectrum is plotted in black pointing upward, and the matched fragment b ions (blue) and y ions (red) are plotted pointing downward, mirrored against it, so that each matched fragment can be compared directly with the measured spectrum. **E** $MS^2$ spectrum of the fragments corresponding to the Unmodified peptide sequence, with matched b (blue, above) and y ions (red, below) plotted. **F** Mirrored $MS^2$ plot for the Spliced peptide. In this mirror plot, the full fragmentation spectrum is plotted in black pointing upward, and the matched fragment b ions (blue) and y ions (red) are plotted pointing downward, mirrored against it, so that each matched fragment can be compared directly with the measured spectrum. **G** $MS^2$ spectrum of the fragments corresponding to the Spliced peptide sequence, with matched b (blue, above) and y ions (red, below) plotted.

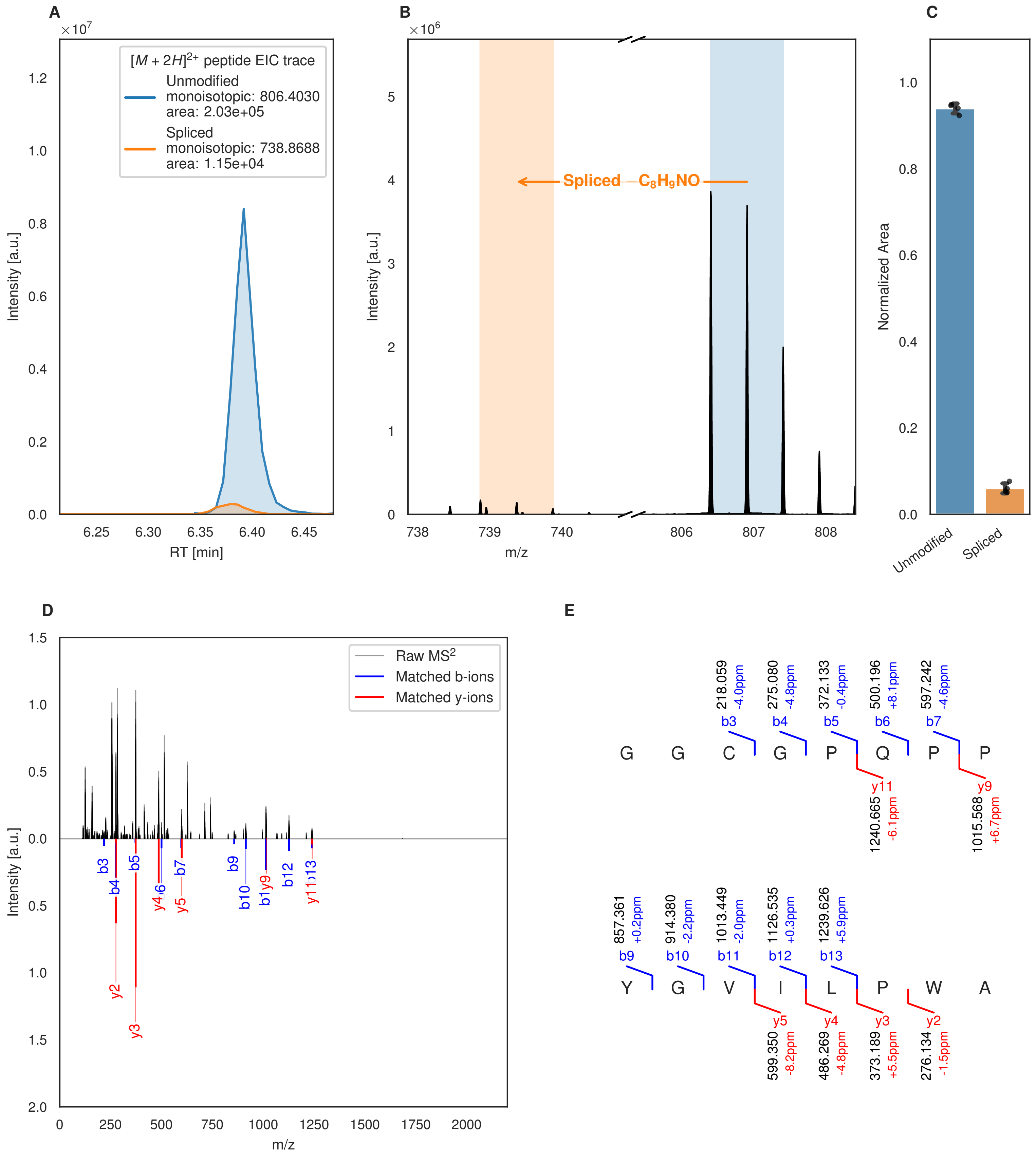

A
$[M+2H]^{2+}$ peptide EIC trace
Unmodified
monoisotopic: 806.4030
area: 2.03e+05
Spliced
monoisotopic: 738.8688
area: 1.15e+04
Intensity [a.u.]
RT [min]
B
Spliced $-C_8H_9NO$
Intensity [a.u.]
m/z
C
Normalized Area
Unmodified
Spliced
D
Raw MS2
Matched b-ions
Matched y-ions
Intensity [a.u.]
m/z
E
G G C G P Q P P
b3 218.059 -4.0ppm
b4 275.080 -4.8ppm
b5 372.133 -0.4ppm
b6 500.196 +8.1ppm
b7 597.242 -4.6ppm
y11 1240.665 -6.1ppm
y9 1015.568 +6.7ppm
Y G V I L P W A
b9 857.361 +0.2ppm
b10 914.380 -2.2ppm
b11 1013.449 -2.0ppm
b12 1126.535 +0.3ppm
b13 1239.626 +5.9ppm
y5 599.350 -8.2ppm
y4 486.269 -4.8ppm
y3 373.189 +5.5ppm
y2 276.134 -1.5ppm

Figure S100 (figure continued from the previous page).

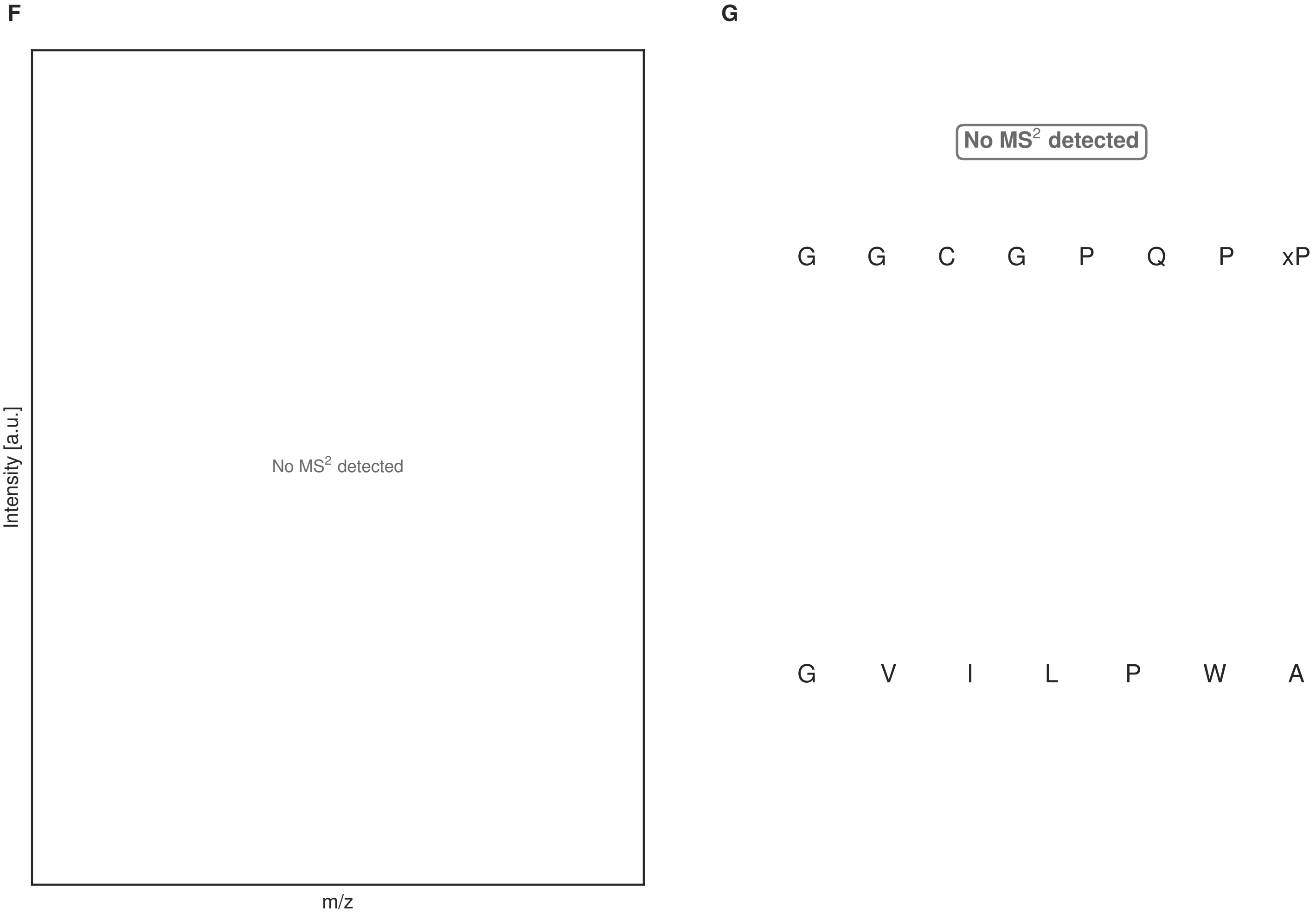


**Figure S100: Modification of the mutant_23 precursor peptide by PlpXY.** LC-MS analysis of IMAC-purified precursor peptides from *E. coli* BL21(DE3)Tuner containing splicease-encoding plasmid TJ-C-559 (PlpXY) and precursor peptide-encoding plasmid mutant_23 (RSTN1-PlpA3-TEV-mutant-23) after expression at 24 °C for 22 h in 1.5 mL cultures after digestion with TEV. Data from n=4 independent biological replicates. Spliced amino acids have an x as a prefix. Spliced refers to a single tyramine excision. A representative experimental replicate is shown in panels **A, B.**

**A** Close-up plot of the representative extracted ion chromatogram (EIC) of the various peptide states; the blue trace corresponds to the Unmodified peptide; the orange trace corresponds to the Spliced peptide. **B** Close-up plot of the m/z values of the peptides detected by the mass spectrometer, with the shaded colored areas corresponding to where the expected m/z values for the peptide m/z traces should occur, with the observed spectrum plotted in black. The Unmodified, Spliced peptide(s) were found in the mass spectrum. **C** Bar plot showing the relative abundance of the various peptide states at charge z=2; the bars represent the average relative abundances normalized to total peak areas, with each individual experimental replicate value plotted as a black dot. **D** Mirrored MS$^2$ plot for the Unmodified peptide. In this mirror plot, the full fragmentation spectrum is plotted in black pointing upward, and the matched fragment b ions (blue) and y ions (red) are plotted pointing downward, mirrored against it, so that each matched fragment can be compared directly with the measured spectrum. **E** MS$^2$ spectrum of the fragments corresponding to the Unmodified peptide sequence, with matched b (blue, above) and y ions (red, below) plotted. **F** Mirrored MS$^2$ plot for the Spliced peptide. In this mirror plot, the full fragmentation spectrum is plotted in black pointing upward, and the matched fragment b ions (blue) and y ions (red) are plotted pointing downward, mirrored against it, so that each matched fragment can be compared directly with the measured spectrum. **G** MS$^2$ spectrum of the fragments corresponding to the Spliced peptide sequence, with matched b (blue, above) and y ions (red, below) plotted.

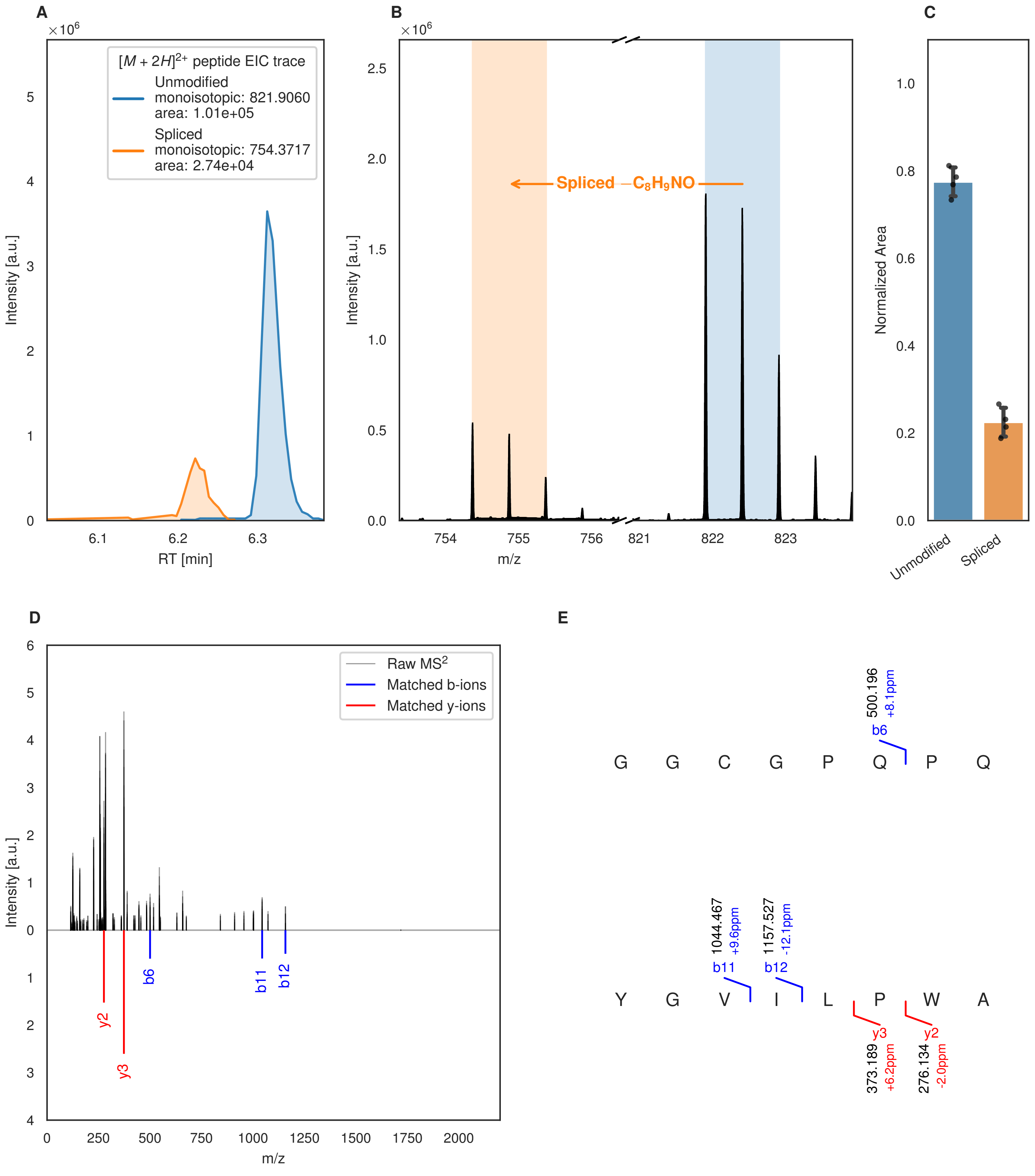
A
B
C
D
E
$[M+2H]^{2+}$ peptide EIC trace
Unmodified
monoisotopic: 821.9060
area: 1.01e+05
Spliced
monoisotopic: 754.3717
area: 2.74e+04
Intensity [a.u.]
RT [min]
Spliced $-C_8H_9NO$
m/z
Normalized Area
Unmodified
Spliced
Raw MS$^2$
Matched b-ions
Matched y-ions
G G C G P Q P Q
Y G V I L P W A
b6
500.196
+8.1ppm
b11
1044.467
+9.6ppm
b12
1157.527
-12.1ppm
y3
373.189
+6.2ppm
y2
276.134
-2.0ppm

Figure S101 (figure continued from the previous page).

F

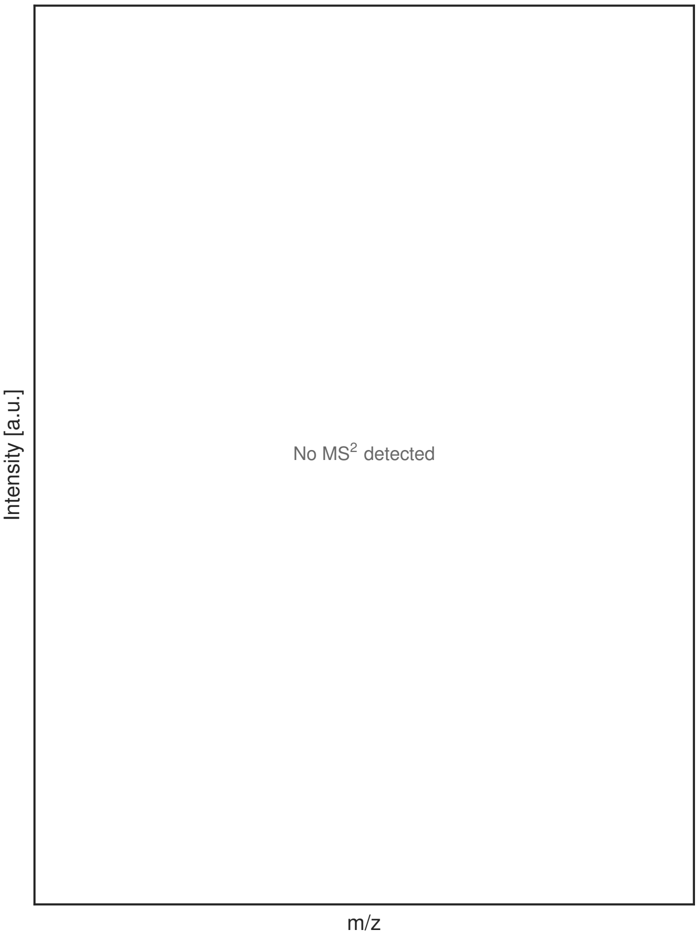


G

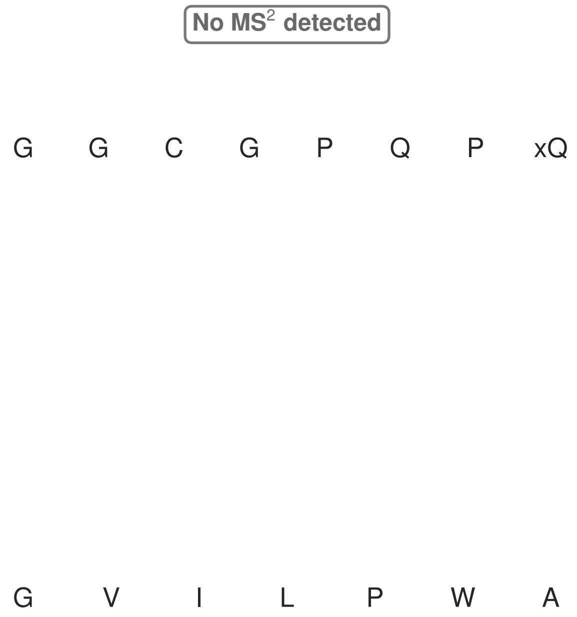


**Figure S101: Modification of the mutant_24 precursor peptide by HypXY.** LC-MS analysis of IMAC-purified precursor peptides from *E. coli* BL21(DE3)Tuner containing splicease-encoding plasmid TJ-C-689 (HypXY) and precursor peptide-encoding plasmid mutant_24 (RSTN1-HypAc-TEV-mutant-24) after expression at 24 °C for 22 h in 1.5 mL cultures after digestion with TEV. Data from n=4 independent biological replicates. Spliced amino acids have an x as a prefix. Spliced refers to a single tyramine excision. A representative experimental replicate is shown in panels **A, B.**

**A** Close-up plot of the representative extracted ion chromatogram (EIC) of the various peptide states; the blue trace corresponds to the Unmodified peptide; the orange trace corresponds to the Spliced peptide. **B** Close-up plot of the m/z values of the peptides detected by the mass spectrometer, with the shaded colored areas corresponding to where the expected m/z values for the peptide m/z traces should occur, with the observed spectrum plotted in black. The Unmodified, Spliced peptide(s) were found in the mass spectrum. **C** Bar plot showing the relative abundance of the various peptide states at charge z=2; the bars represent the average relative abundances normalized to total peak areas, with each individual experimental replicate value plotted as a black dot. **D** Mirrored $MS^2$ plot for the Unmodified peptide. In this mirror plot, the full fragmentation spectrum is plotted in black pointing upward, and the matched fragment b ions (blue) and y ions (red) are plotted pointing downward, mirrored against it, so that each matched fragment can be compared directly with the measured spectrum. **E** $MS^2$ spectrum of the fragments corresponding to the Unmodified peptide sequence, with matched b (blue, above) and y ions (red, below) plotted. **F** Mirrored $MS^2$ plot for the Spliced peptide. In this mirror plot, the full fragmentation spectrum is plotted in black pointing upward, and the matched fragment b ions (blue) and y ions (red) are plotted pointing downward, mirrored against it, so that each matched fragment can be compared directly with the measured spectrum. **G** $MS^2$ spectrum of the fragments corresponding to the Spliced peptide sequence, with matched b (blue, above) and y ions (red, below) plotted.

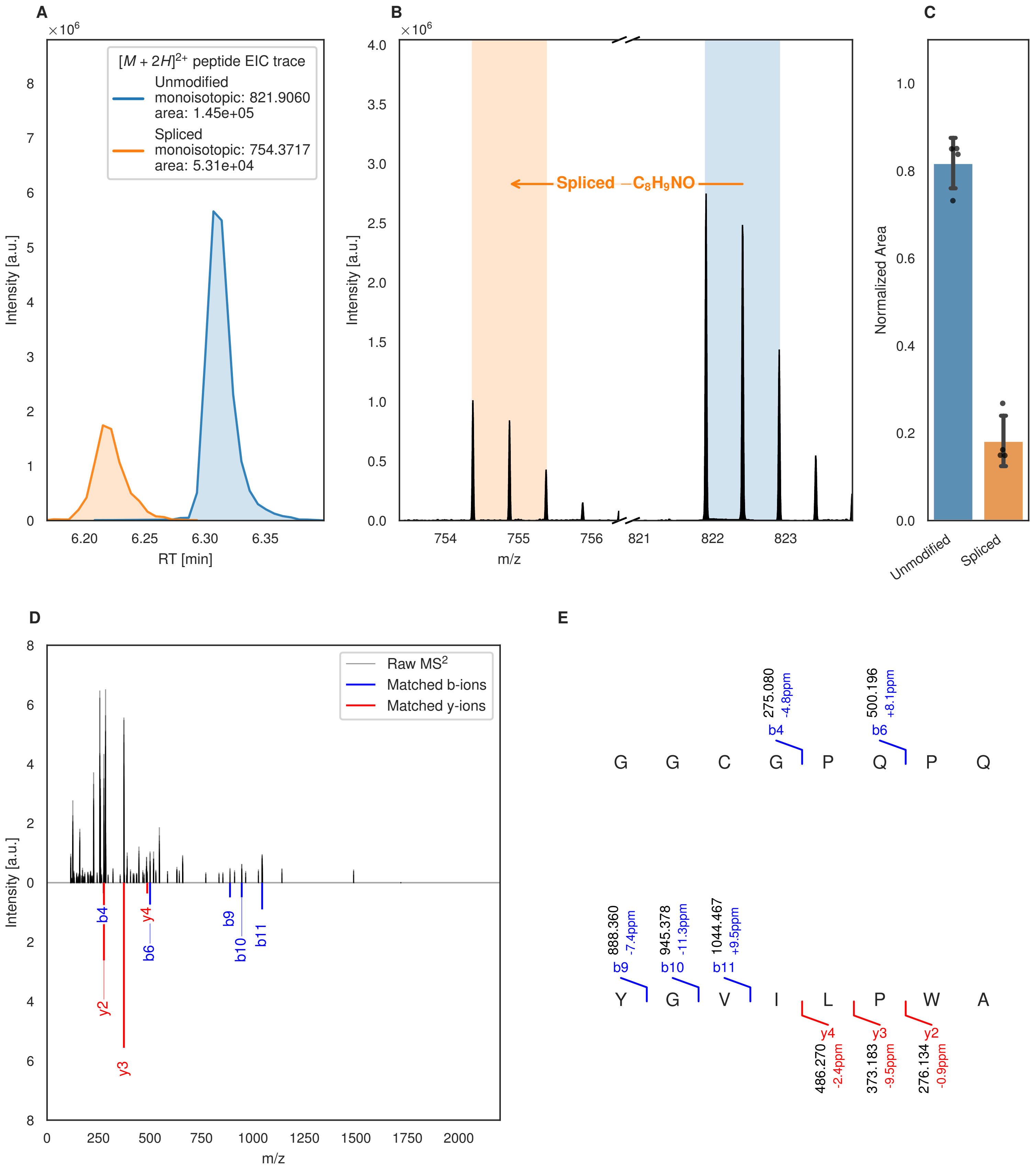
A
B
C
D
E
$[M+2H]^{2+}$ peptide EIC trace
Unmodified
monoisotopic: 821.9060
area: 1.45e+05
Spliced
monoisotopic: 754.3717
area: 5.31e+04
Intensity [a.u.]
RT [min]
Spliced $-C_8H_9NO$
m/z
Normalized Area
Unmodified
Spliced
Raw MS$^2$
Matched b-ions
Matched y-ions
b4
y4
b6
b9
b10
b11
y2
y3
G G C G P Q P Q
275.080 -4.8ppm
500.196 +8.1ppm
Y G V I L P W A
888.360 -7.4ppm
945.378 -11.3ppm
1044.467 +9.5ppm
486.270 -2.4ppm
373.183 -9.5ppm
276.134 -0.9ppm

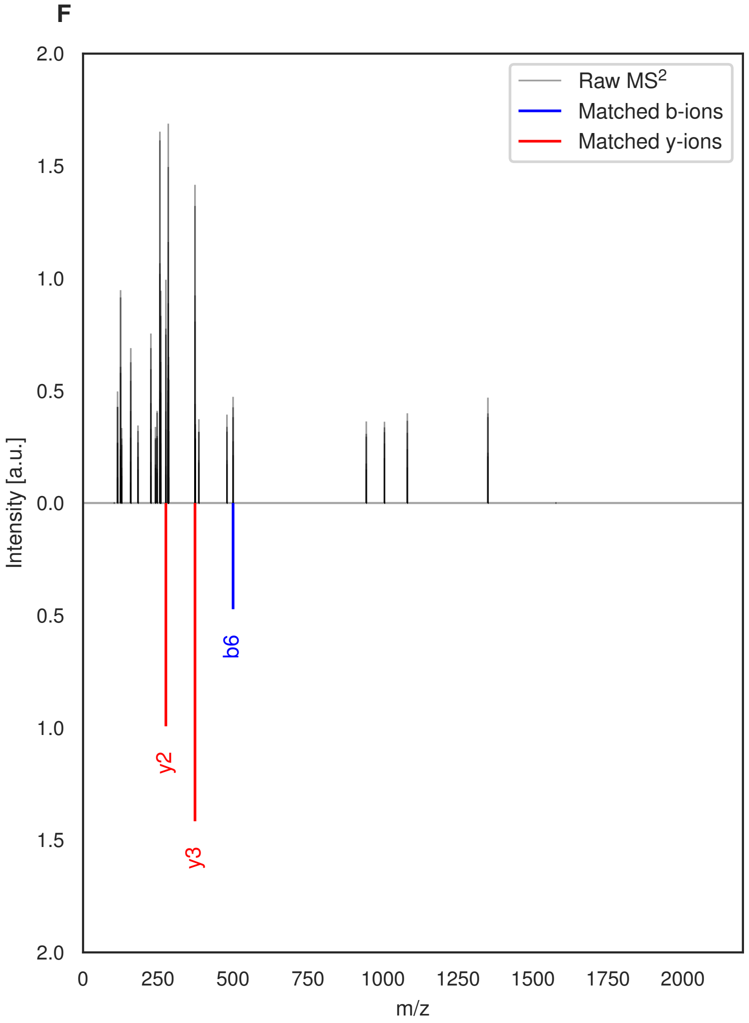


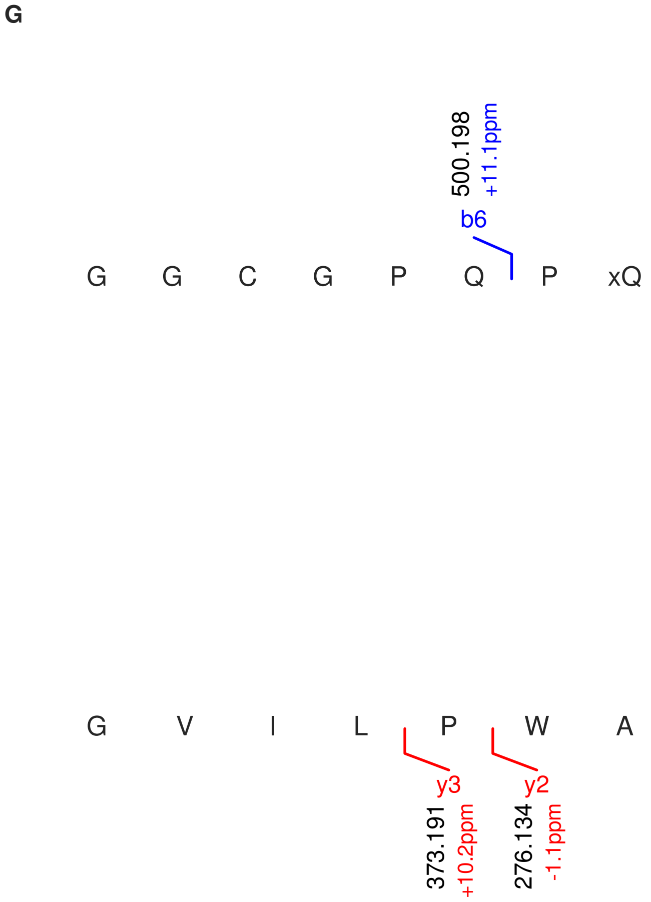


**Figure S102: Modification of the mutant_24 precursor peptide by PlpXY.** LC-MS analysis of IMAC-purified precursor peptides from *E. coli* BL21(DE3)Tuner containing splicease-encoding plasmid TJ-C-559 (PlpXY) and precursor peptide-encoding plasmid mutant_24 (RSTN1-PlpA3-TEV-mutant-24) after expression at 24 °C for 22 h in 1.5 mL cultures after digestion with TEV. Data from n=4 independent biological replicates. Spliced amino acids have an x as a prefix. Spliced refers to a single tyramine excision. A representative experimental replicate is shown in panels **A, B.**

**A** Close-up plot of the representative extracted ion chromatogram (EIC) of the various peptide states; the blue trace corresponds to the Unmodified peptide; the orange trace corresponds to the Spliced peptide. **B** Close-up plot of the m/z values of the peptides detected by the mass spectrometer, with the shaded colored areas corresponding to where the expected m/z values for the peptide m/z traces should occur, with the observed spectrum plotted in black. The Unmodified, Spliced peptide(s) were found in the mass spectrum. **C** Bar plot showing the relative abundance of the various peptide states at charge z=2; the bars represent the average relative abundances normalized to total peak areas, with each individual experimental replicate value plotted as a black dot. **D** Mirrored MS$^2$ plot for the Unmodified peptide. In this mirror plot, the full fragmentation spectrum is plotted in black pointing upward, and the matched fragment b ions (blue) and y ions (red) are plotted pointing downward, mirrored against it, so that each matched fragment can be compared directly with the measured spectrum. **E** MS$^2$ spectrum of the fragments corresponding to the Unmodified peptide sequence, with matched b (blue, above) and y ions (red, below) plotted. **F** Mirrored MS$^2$ plot for the Spliced peptide. In this mirror plot, the full fragmentation spectrum is plotted in black pointing upward, and the matched fragment b ions (blue) and y ions (red) are plotted pointing downward, mirrored against it, so that each matched fragment can be compared directly with the measured spectrum. **G** MS$^2$ spectrum of the fragments corresponding to the Spliced peptide sequence, with matched b (blue, above) and y ions (red, below) plotted.

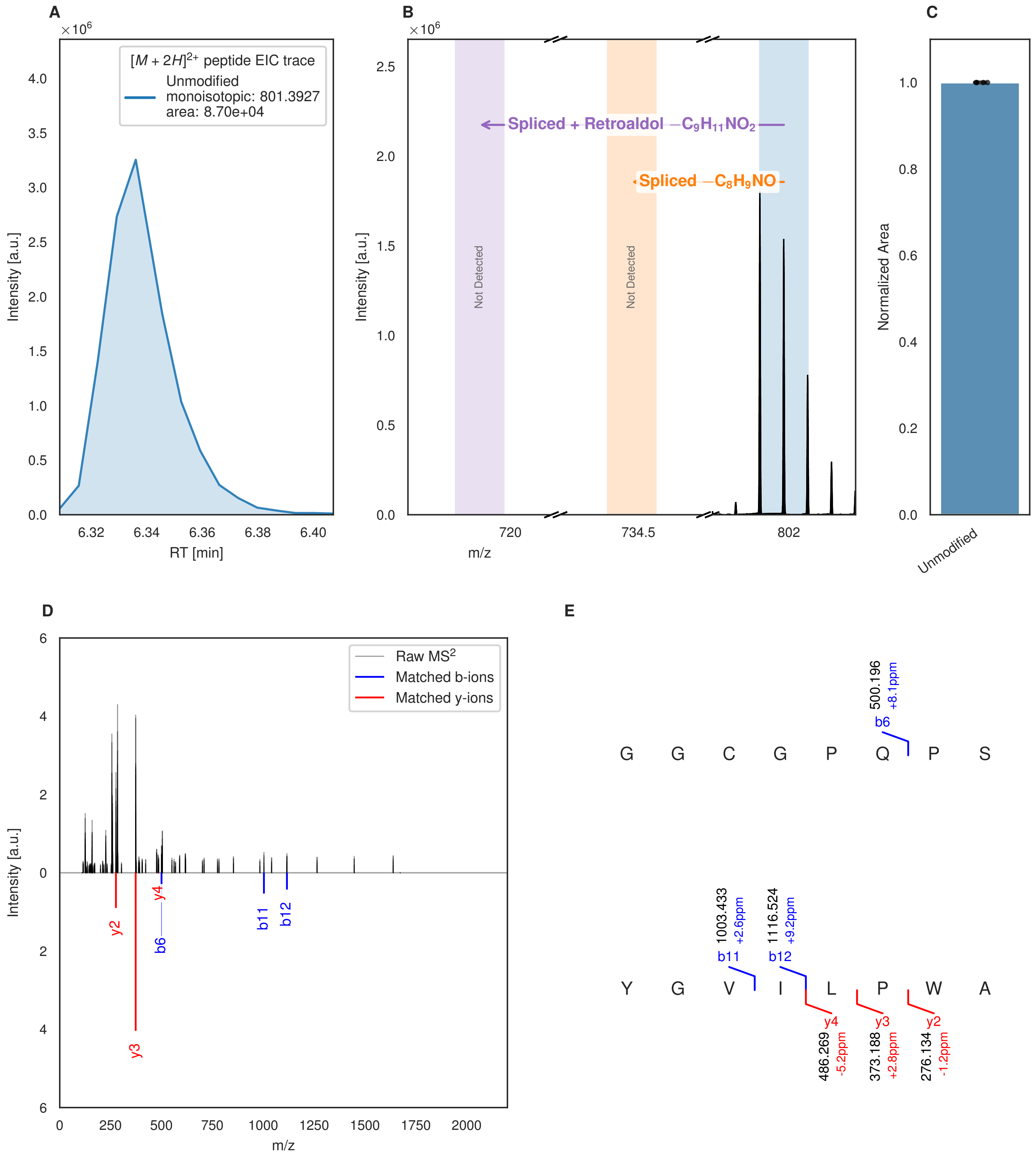


**Figure S103: Modification of the mutant_25 precursor peptide by HypXY.** LC-MS analysis of IMAC-purified precursor peptides from *E. coli* BL21(DE3)Tuner containing splicease-encoding plasmid TJ-C-689 (HypXY) and precursor peptide-encoding plasmid mutant_25 (RSTN1-HypAc-TEV-mutant-25) after expression at 24 °C for 22 h in 1.5 mL cultures after digestion with TEV. Data from n=4 independent biological replicates. Spliced amino acids have an x as a prefix. Spliced refers to a single tyramine excision; Spliced + Retroaldol refers to a glycine-derived ketoamide (xG). A representative experimental replicate is shown in panels **A, B.**
**A** Close-up plot of the representative extracted ion chromatogram (EIC) of the various peptide states; the blue trace corresponds to the Unmodified peptide. **B** Close-up plot of the m/z values of the peptides detected by the mass spectrometer, with the shaded colored areas corresponding to where the expected m/z values for the peptide m/z

(Caption continued on the next page.)

traces should occur, with the observed spectrum plotted in black. The Unmodified peptide(s) were found in the mass spectrum and the Spliced, Spliced + Retroaldol peptide(s) were not detected. **C** Bar plot showing the relative abundance of the various peptide states at charge z=2; the bars represent the average relative abundances normalized to total peak areas, with each individual experimental replicate value plotted as a black dot. **D** Mirrored $MS^2$ plot for the Unmodified peptide. In this mirror plot, the full fragmentation spectrum is plotted in black pointing upward, and the matched fragment b ions (blue) and y ions (red) are plotted pointing downward, mirrored against it, so that each matched fragment can be compared directly with the measured spectrum. **E** $MS^2$ spectrum of the fragments corresponding to the Unmodified peptide sequence, with matched b (blue, above) and y ions (red, below) plotted.

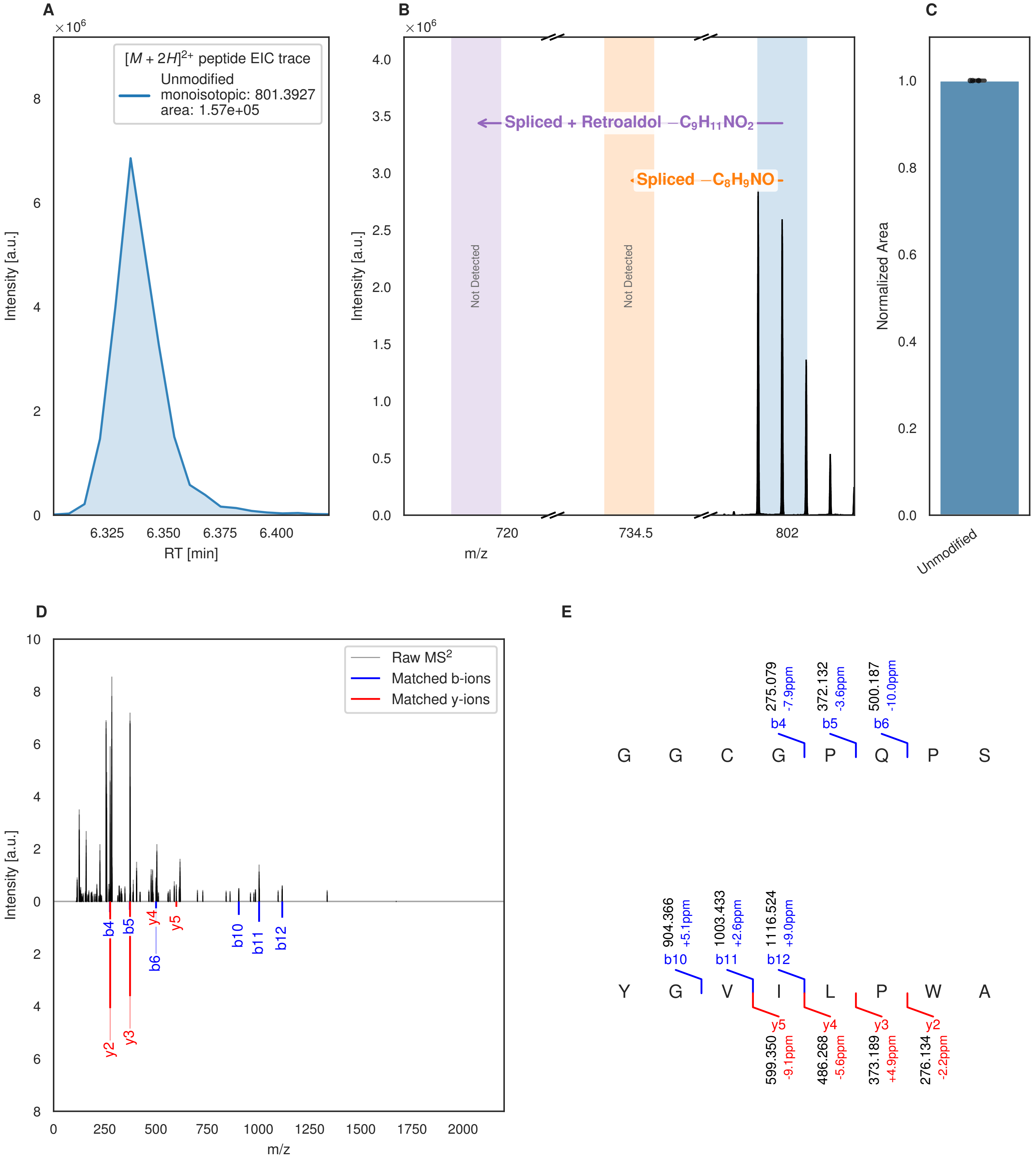


**Figure S104: Modification of the mutant_25 precursor peptide by PlpXY.** LC-MS analysis of IMAC-purified precursor peptides from *E. coli* BL21(DE3)Tuner containing splicease-encoding plasmid TJ-C-559 (PlpXY) and precursor peptide-encoding plasmid mutant_25 (RSTN1-PlpA3-TEV-mutant-25) after expression at 24 °C for 22 h in 1.5 mL cultures after digestion with TEV. Data from n=3 independent biological replicates. Spliced amino acids have an x as a prefix. Spliced refers to a single tyramine excision; Spliced + Retroaldol refers to a glycine-derived ketoamide (xG). A representative experimental replicate is shown in panels **A, B.**

**A** Close-up plot of the representative extracted ion chromatogram (EIC) of the various peptide states; the blue trace corresponds to the Unmodified peptide. **B** Close-up plot of the m/z values of the peptides detected by the mass spectrometer, with the shaded colored areas corresponding to where the expected m/z values for the peptide m/z

(Caption continued on the next page.)

traces should occur, with the observed spectrum plotted in black. The Unmodified peptide(s) were found in the mass spectrum and the Spliced, Spliced + Retroaldol peptide(s) were not detected. **C** Bar plot showing the relative abundance of the various peptide states at charge z=2; the bars represent the average relative abundances normalized to total peak areas, with each individual experimental replicate value plotted as a black dot. **D** Mirrored MS$^2$ plot for the Unmodified peptide. In this mirror plot, the full fragmentation spectrum is plotted in black pointing upward, and the matched fragment b ions (blue) and y ions (red) are plotted pointing downward, mirrored against it, so that each matched fragment can be compared directly with the measured spectrum. **E** MS$^2$ spectrum of the fragments corresponding to the Unmodified peptide sequence, with matched b (blue, above) and y ions (red, below) plotted.

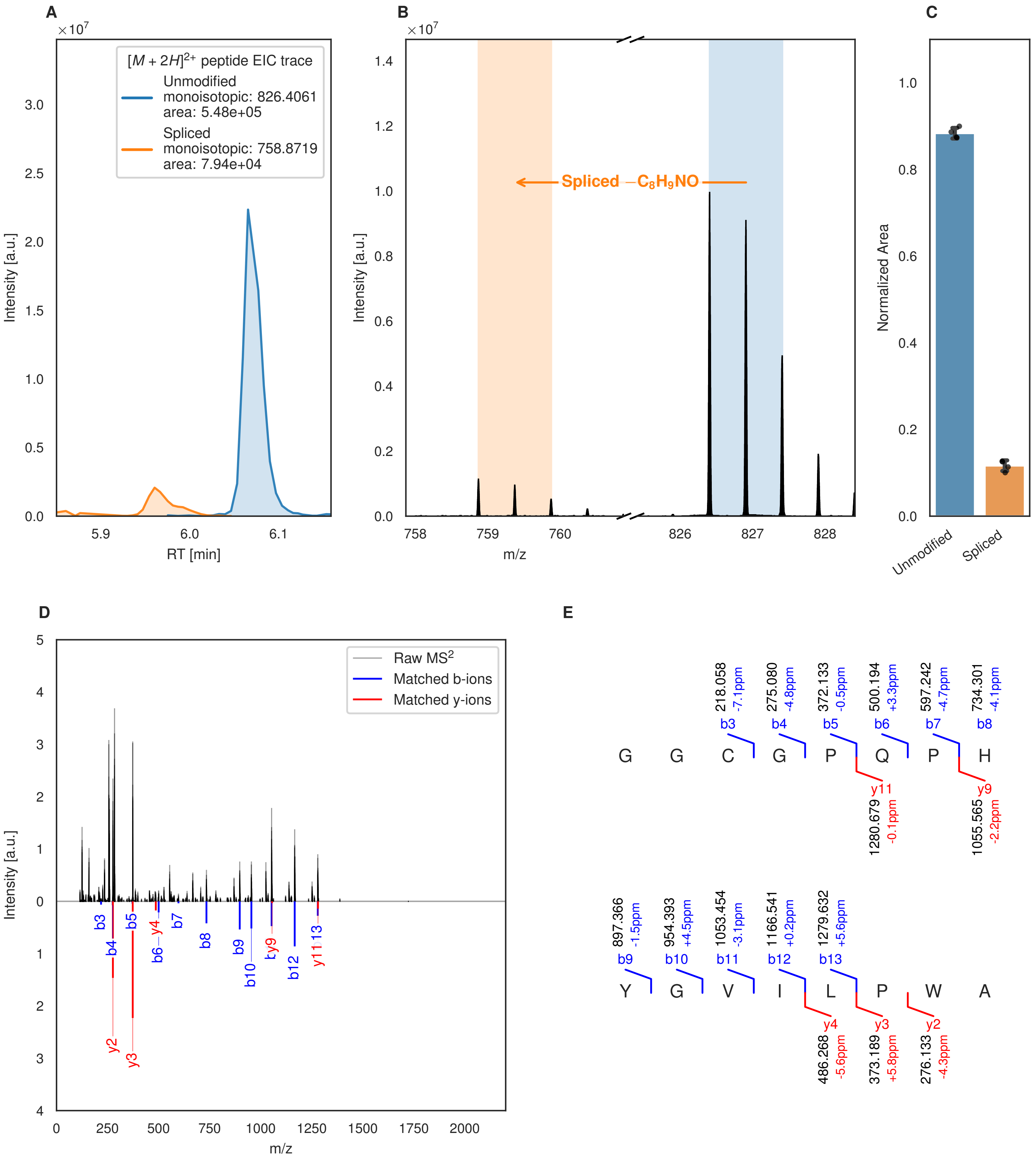
A
[M + 2H]2+ peptide EIC trace
Unmodified
monoisotopic: 826.4061
area: 5.48e+05
Spliced
monoisotopic: 758.8719
area: 7.94e+04
Intensity [a.u.]
RT [min]
B
Spliced −C8H9NO
Intensity [a.u.]
m/z
C
Normalized Area
Unmodified
Spliced
D
Raw MS2
Matched b-ions
Matched y-ions
Intensity [a.u.]
m/z
E
218.058 -7.1ppm b3
275.080 -4.8ppm b4
372.133 -0.5ppm b5
500.194 +3.3ppm b6
597.242 -4.7ppm b7
734.301 -4.1ppm b8
G G C G P Q P H
y11 1280.679 -0.1ppm
y9 1055.565 -2.2ppm
897.366 -1.5ppm b9
954.393 +4.5ppm b10
1053.454 -3.1ppm b11
1166.541 +0.2ppm b12
1279.632 +5.6ppm b13
Y G V I L P W A
y4 486.268 -5.6ppm
y3 373.189 +5.8ppm
y2 276.133 -4.3ppm

Figure S105 (figure continued from the previous page).

F

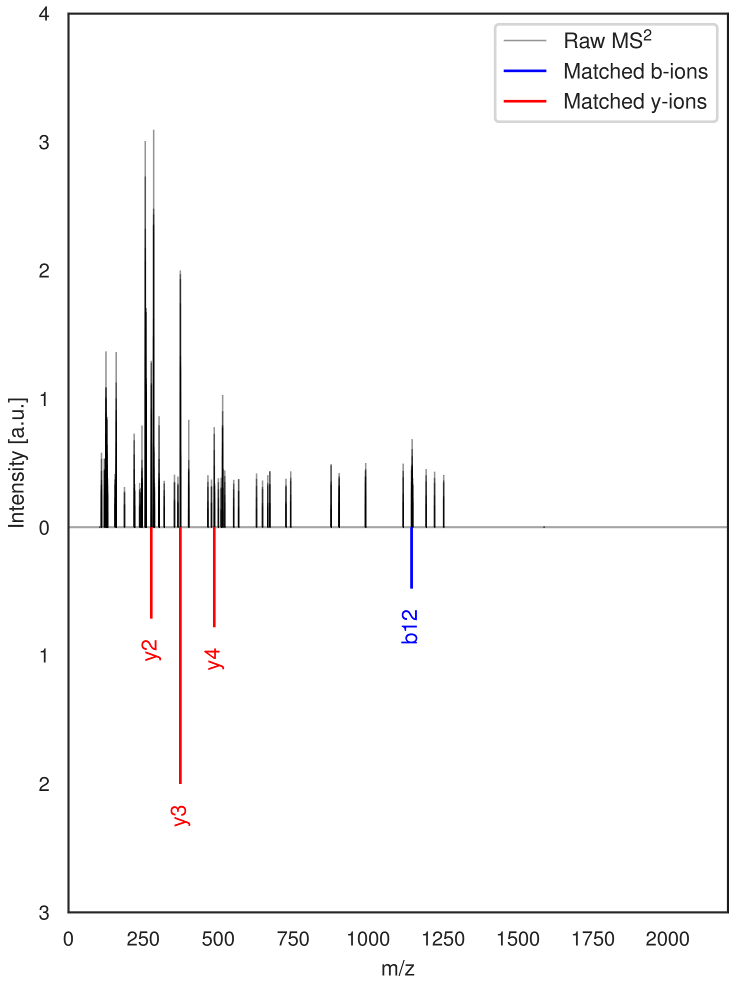


G

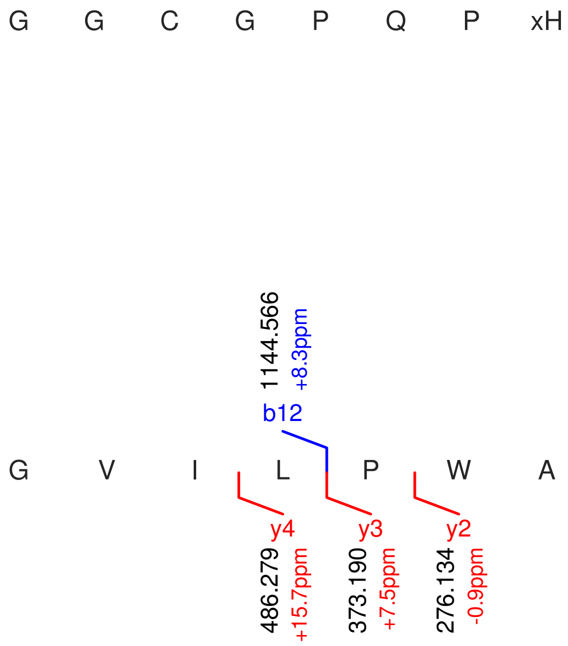


**Figure S105: Modification of the mutant_26 precursor peptide by HypXY.** LC-MS analysis of IMAC-purified precursor peptides from *E. coli* BL21(DE3)Tuner containing splicease-encoding plasmid TJ-C-689 (HypXY) and precursor peptide-encoding plasmid mutant_26 (RSTN1-HypAc-TEV-mutant-26) after expression at 24 °C for 22 h in 1.5 mL cultures after digestion with TEV. Data from n=4 independent biological replicates. Spliced amino acids have an x as a prefix. Spliced refers to a single tyramine excision. A representative experimental replicate is shown in panels **A, B.**
**A** Close-up plot of the representative extracted ion chromatogram (EIC) of the various peptide states; the blue trace corresponds to the Unmodified peptide; the orange trace corresponds to the Spliced peptide. **B** Close-up plot of the m/z values of the peptides detected by the mass spectrometer, with the shaded colored areas corresponding to where the expected m/z values for the peptide m/z traces should occur, with the observed spectrum plotted in black. The Unmodified, Spliced peptide(s) were found in the mass spectrum. **C** Bar plot showing the relative abundance of the various peptide states at charge z=2; the bars represent the average relative abundances normalized to total peak areas, with each individual experimental replicate value plotted as a black dot. **D** Mirrored MS$^2$ plot for the Unmodified peptide. In this mirror plot, the full fragmentation spectrum is plotted in black pointing upward, and the matched fragment b ions (blue) and y ions (red) are plotted pointing downward, mirrored against it, so that each matched fragment can be compared directly with the measured spectrum. **E** MS$^2$ spectrum of the fragments corresponding to the Unmodified peptide sequence, with matched b (blue, above) and y ions (red, below) plotted. **F** Mirrored MS$^2$ plot for the Spliced peptide. In this mirror plot, the full fragmentation spectrum is plotted in black pointing upward, and the matched fragment b ions (blue) and y ions (red) are plotted pointing downward, mirrored against it, so that each matched fragment can be compared directly with the measured spectrum. **G** MS$^2$ spectrum of the fragments corresponding to the Spliced peptide sequence, with matched b (blue, above) and y ions (red, below) plotted.

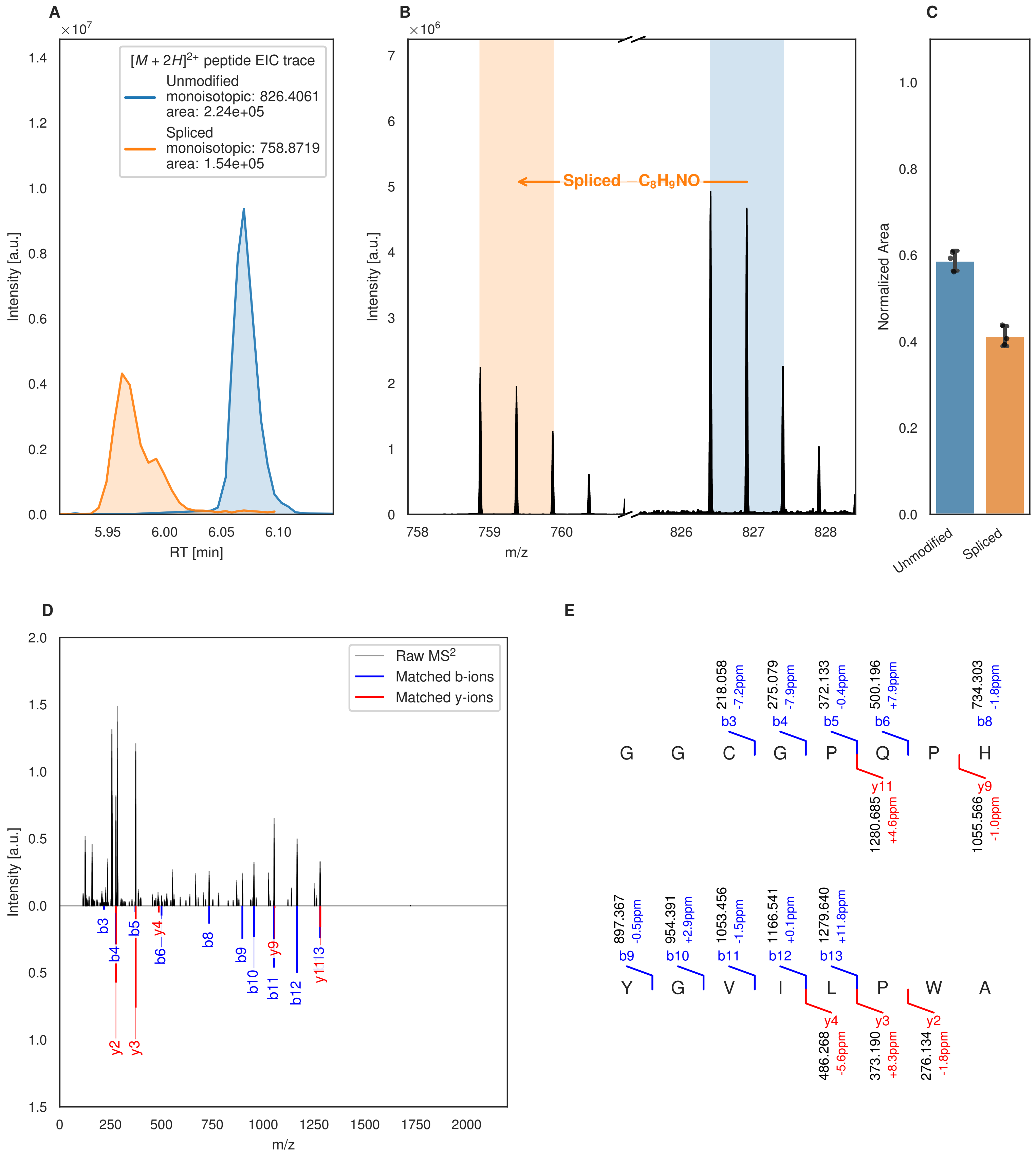
A
×10^7
[M + 2H]^2+ peptide EIC trace
Unmodified
monoisotopic: 826.4061
area: 2.24e+05
Spliced
monoisotopic: 758.8719
area: 1.54e+05
Intensity [a.u.]
RT [min]
B
×10^6
Spliced −C8H9NO
Intensity [a.u.]
m/z
C
Normalized Area
Unmodified
Spliced
D
Raw MS2
Matched b-ions
Matched y-ions
Intensity [a.u.]
m/z
b3
b4
b5
y4
b6
b8
b9
b10
y9
b11
b12
y11
y2
y3
E
218.058 -7.2ppm b3
275.079 -7.9ppm b4
372.133 -0.4ppm b5
500.196 +7.9ppm b6
734.303 -1.8ppm b8
G G C G P Q P H
y11 1280.685 +4.6ppm
y9 1055.566 -1.0ppm
897.367 -0.5ppm b9
954.391 +2.9ppm b10
1053.456 -1.5ppm b11
1166.541 +0.1ppm b12
1279.640 +11.8ppm b13
Y G V I L P W A
y4 486.268 -5.6ppm
y3 373.190 +8.3ppm
y2 276.134 -1.8ppm

F

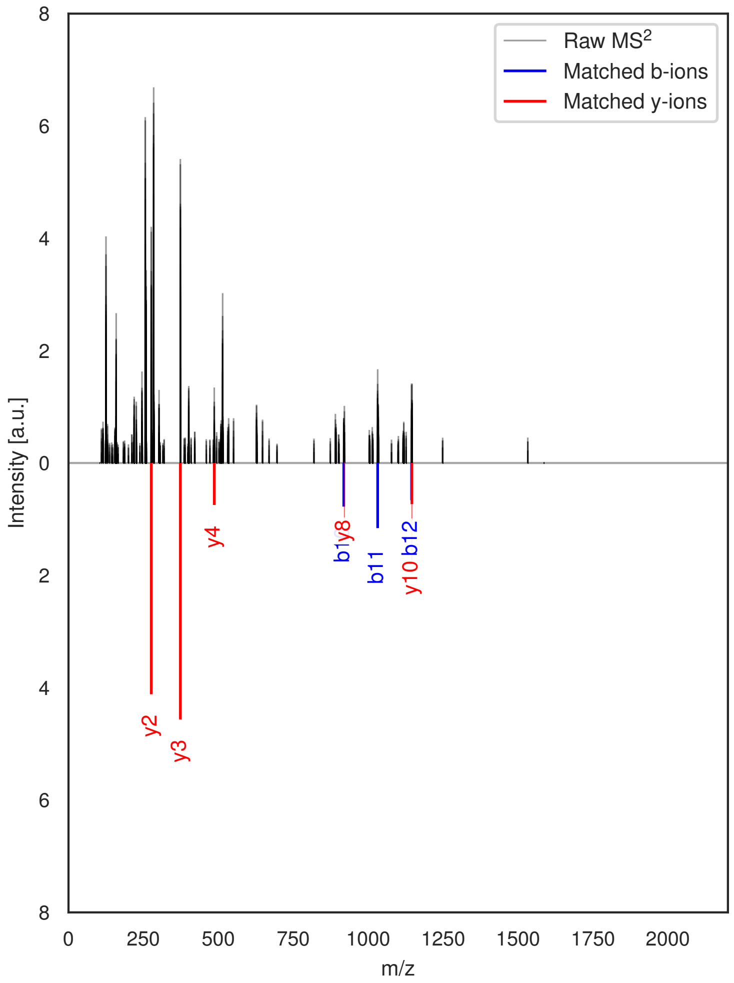


G

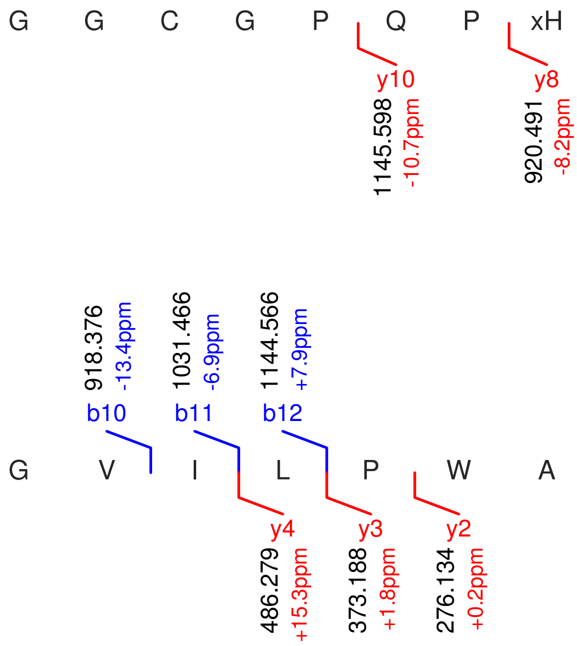


**Figure S106: Modification of the mutant_26 precursor peptide by PlpXY.** LC-MS analysis of IMAC-purified precursor peptides from *E. coli* BL21(DE3)Tuner containing splicease-encoding plasmid TJ-C-559 (PlpXY) and precursor peptide-encoding plasmid mutant_26 (RSTN1-PlpA3-TEV-mutant-26) after expression at 24 °C for 22 h in 1.5 mL cultures after digestion with TEV. Data from n=3 independent biological replicates. Spliced amino acids have an x as a prefix. Spliced refers to a single tyramine excision. A representative experimental replicate is shown in panels **A, B.**

**A** Close-up plot of the representative extracted ion chromatogram (EIC) of the various peptide states; the blue trace corresponds to the Unmodified peptide; the orange trace corresponds to the Spliced peptide. **B** Close-up plot of the m/z values of the peptides detected by the mass spectrometer, with the shaded colored areas corresponding to where the expected m/z values for the peptide m/z traces should occur, with the observed spectrum plotted in black. The Unmodified, Spliced peptide(s) were found in the mass spectrum. **C** Bar plot showing the relative abundance of the various peptide states at charge z=2; the bars represent the average relative abundances normalized to total peak areas, with each individual experimental replicate value plotted as a black dot. **D** Mirrored MS$^2$ plot for the Unmodified peptide. In this mirror plot, the full fragmentation spectrum is plotted in black pointing upward, and the matched fragment b ions (blue) and y ions (red) are plotted pointing downward, mirrored against it, so that each matched fragment can be compared directly with the measured spectrum. **E** MS$^2$ spectrum of the fragments corresponding to the Unmodified peptide sequence, with matched b (blue, above) and y ions (red, below) plotted. **F** Mirrored MS$^2$ plot for the Spliced peptide. In this mirror plot, the full fragmentation spectrum is plotted in black pointing upward, and the matched fragment b ions (blue) and y ions (red) are plotted pointing downward, mirrored against it, so that each matched fragment can be compared directly with the measured spectrum. **G** MS$^2$ spectrum of the fragments corresponding to the Spliced peptide sequence, with matched b (blue, above) and y ions (red, below) plotted.

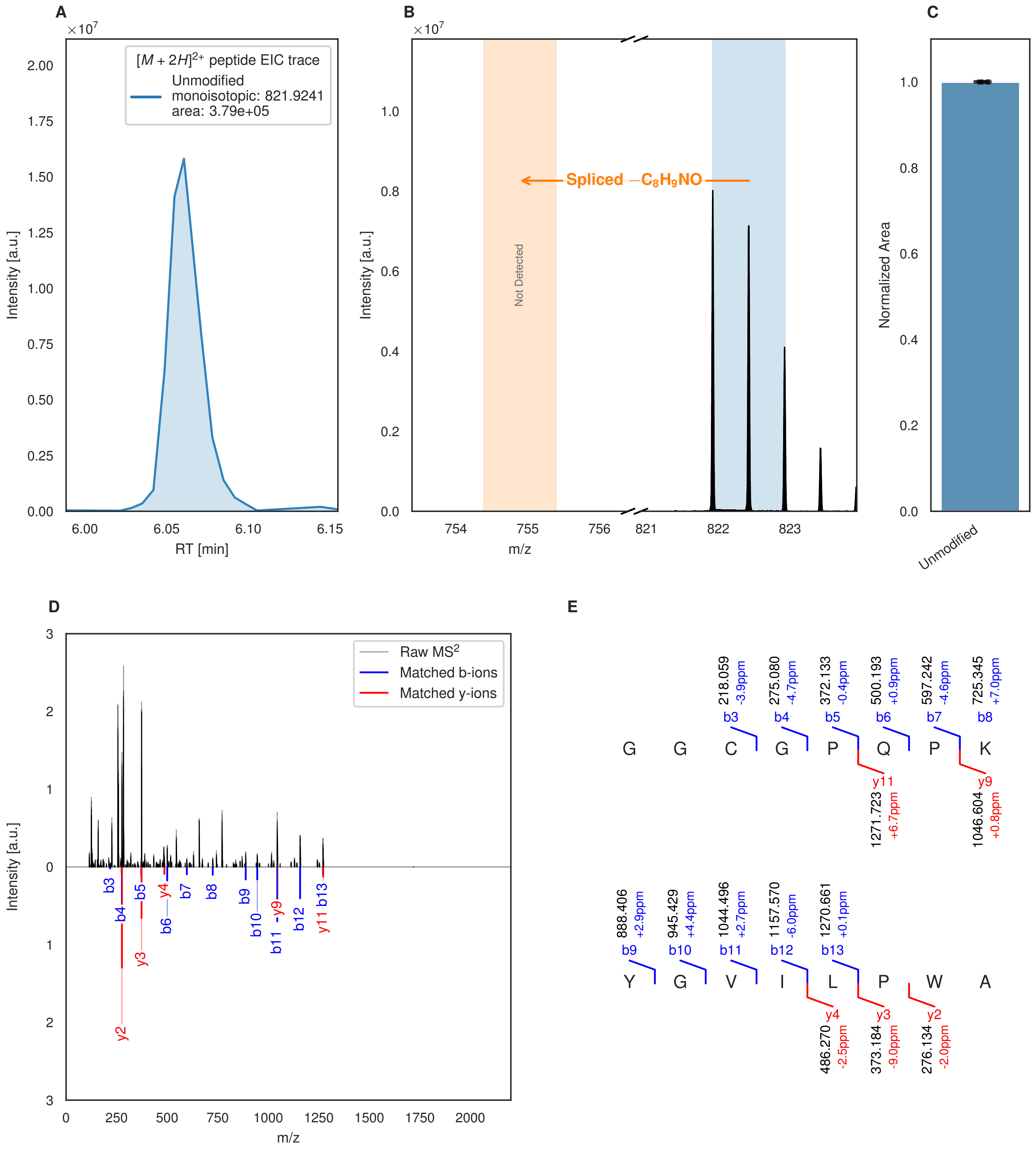


**Figure S107: Modification of the mutant_27 precursor peptide by HypXY.** LC-MS analysis of IMAC-purified precursor peptides from *E. coli* BL21(DE3)Tuner containing splicease-encoding plasmid TJ-C-689 (HypXY) and precursor peptide-encoding plasmid mutant_27 (RSTN1-HypAc-TEV-mutant-27) after expression at 24 °C for 22 h in 1.5 mL cultures after digestion with TEV. Data from n=4 independent biological replicates. Spliced amino acids have an x as a prefix. Spliced refers to a single tyramine excision. A representative experimental replicate is shown in panels **A, B.**

**A** Close-up plot of the representative extracted ion chromatogram (EIC) of the various peptide states; the blue trace corresponds to the Unmodified peptide. **B** Close-up plot of the m/z values of the peptides detected by the mass spectrometer, with the shaded colored areas corresponding to where the expected m/z values for the peptide m/z

(Caption continued on the next page.)

traces should occur, with the observed spectrum plotted in black. The Unmodified peptide(s) were found in the mass spectrum and the Spliced peptide(s) were not detected. **C** Bar plot showing the relative abundance of the various peptide states at charge z=2; the bars represent the average relative abundances normalized to total peak areas, with each individual experimental replicate value plotted as a black dot. **D** Mirrored $MS^2$ plot for the Unmodified peptide. In this mirror plot, the full fragmentation spectrum is plotted in black pointing upward, and the matched fragment b ions (blue) and y ions (red) are plotted pointing downward, mirrored against it, so that each matched fragment can be compared directly with the measured spectrum. **E** $MS^2$ spectrum of the fragments corresponding to the Unmodified peptide sequence, with matched b (blue, above) and y ions (red, below) plotted.

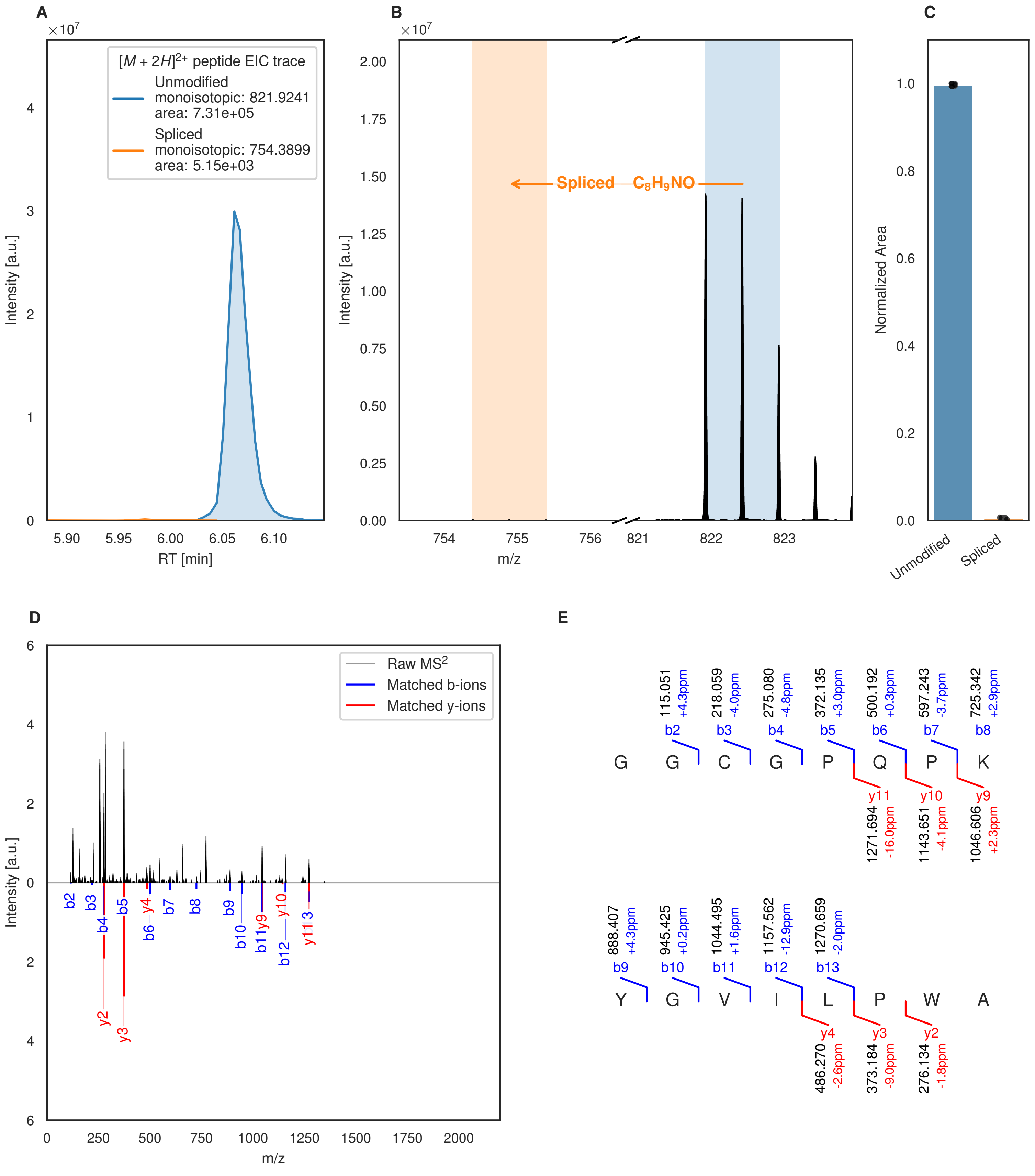
A
×10^7
[M + 2H]^2+ peptide EIC trace
Unmodified
monoisotopic: 821.9241
area: 7.31e+05
Spliced
monoisotopic: 754.3899
area: 5.15e+03
Intensity [a.u.]
RT [min]
B
×10^7
Spliced −C8H9NO
Intensity [a.u.]
m/z
C
Normalized Area
Unmodified
Spliced
D
Raw MS2
Matched b-ions
Matched y-ions
Intensity [a.u.]
m/z
E
G G C G P Q P K
115.051 +4.3ppm b2
218.059 -4.0ppm b3
275.080 -4.8ppm b4
372.135 +3.0ppm b5
500.192 +0.3ppm b6
597.243 -3.7ppm b7
725.342 +2.9ppm b8
y11 1271.694 -16.0ppm
y10 1143.651 -4.1ppm
y9 1046.606 +2.3ppm
Y G V I L P W A
888.407 +4.3ppm b9
945.425 +0.2ppm b10
1044.495 +1.6ppm b11
1157.562 -12.9ppm b12
1270.659 -2.0ppm b13
y4 486.270 -2.6ppm
y3 373.184 -9.0ppm
y2 276.134 -1.8ppm

Figure S108 (figure continued from the previous page).

F

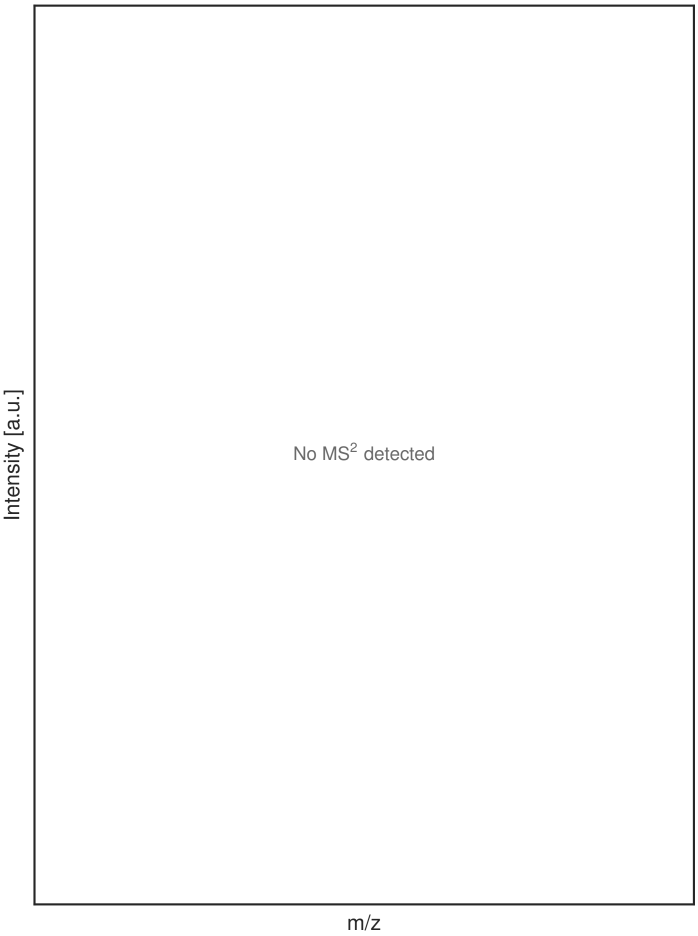


G

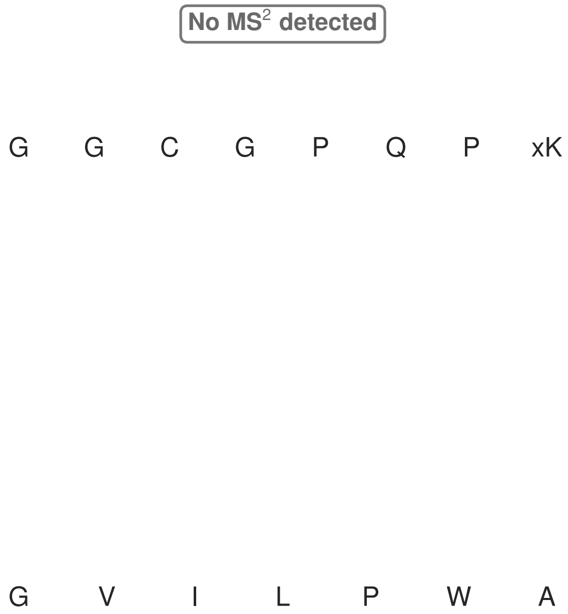


**Figure S108: Modification of the mutant_27 precursor peptide by PlpXY.** LC-MS analysis of IMAC-purified precursor peptides from *E. coli* BL21(DE3)Tuner containing splicease-encoding plasmid TJ-C-559 (PlpXY) and precursor peptide-encoding plasmid mutant_27 (RSTN1-PlpA3-TEV-mutant-27) after expression at 24 °C for 22 h in 1.5 mL cultures after digestion with TEV. Data from n=3 independent biological replicates. Spliced amino acids have an x as a prefix. Spliced refers to a single tyramine excision. A representative experimental replicate is shown in panels **A, B.**

**A** Close-up plot of the representative extracted ion chromatogram (EIC) of the various peptide states; the blue trace corresponds to the Unmodified peptide; the orange trace corresponds to the Spliced peptide. **B** Close-up plot of the m/z values of the peptides detected by the mass spectrometer, with the shaded colored areas corresponding to where the expected m/z values for the peptide m/z traces should occur, with the observed spectrum plotted in black. The Unmodified, Spliced peptide(s) were found in the mass spectrum. **C** Bar plot showing the relative abundance of the various peptide states at charge z=2; the bars represent the average relative abundances normalized to total peak areas, with each individual experimental replicate value plotted as a black dot. **D** Mirrored MS$^2$ plot for the Unmodified peptide. In this mirror plot, the full fragmentation spectrum is plotted in black pointing upward, and the matched fragment b ions (blue) and y ions (red) are plotted pointing downward, mirrored against it, so that each matched fragment can be compared directly with the measured spectrum. **E** MS$^2$ spectrum of the fragments corresponding to the Unmodified peptide sequence, with matched b (blue, above) and y ions (red, below) plotted. **F** Mirrored MS$^2$ plot for the Spliced peptide. In this mirror plot, the full fragmentation spectrum is plotted in black pointing upward, and the matched fragment b ions (blue) and y ions (red) are plotted pointing downward, mirrored against it, so that each matched fragment can be compared directly with the measured spectrum. **G** MS$^2$ spectrum of the fragments corresponding to the Spliced peptide sequence, with matched b (blue, above) and y ions (red, below) plotted.

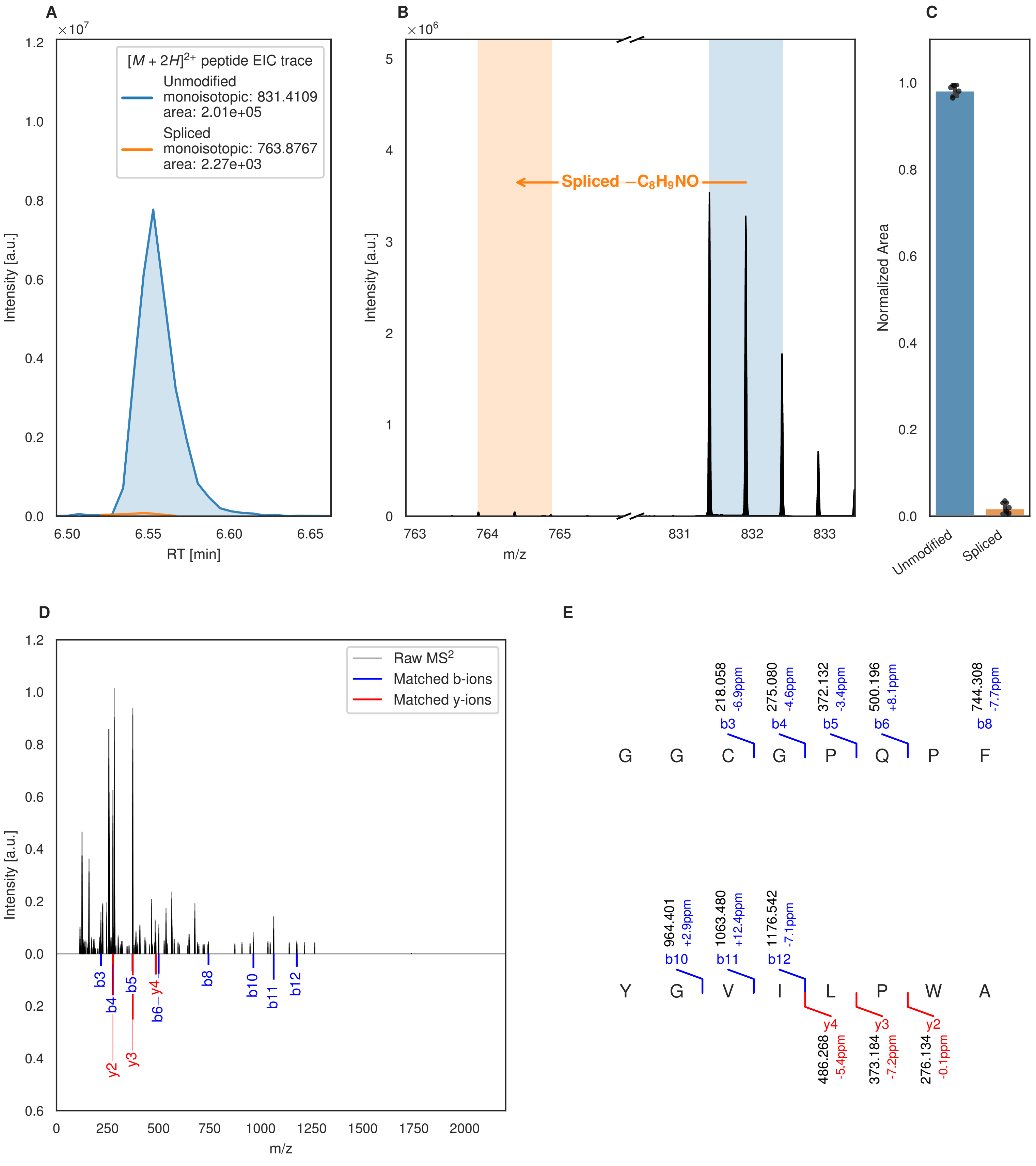
A
$\times10^7$
$[M+2H]^{2+}$ peptide EIC trace
Unmodified
monoisotopic: 831.4109
area: 2.01e+05
Spliced
monoisotopic: 763.8767
area: 2.27e+03
Intensity [a.u.]
RT [min]
B
$\times10^6$
Spliced $-C_8H_9NO$
Intensity [a.u.]
m/z
C
Normalized Area
Unmodified
Spliced
D
Raw $MS^2$
Matched b-ions
Matched y-ions
Intensity [a.u.]
m/z
E
218.058 -6.9ppm b3
275.080 -4.6ppm b4
372.132 -3.4ppm b5
500.196 +8.1ppm b6
744.308 -7.7ppm b8
G G C G P Q P F
964.401 +2.9ppm b10
1063.480 +12.4ppm b11
1176.542 -7.1ppm b12
Y G V I L P W A
y4 486.268 -5.4ppm
y3 373.184 -7.2ppm
y2 276.134 -0.1ppm

Figure S109 (figure continued from the previous page).

F G

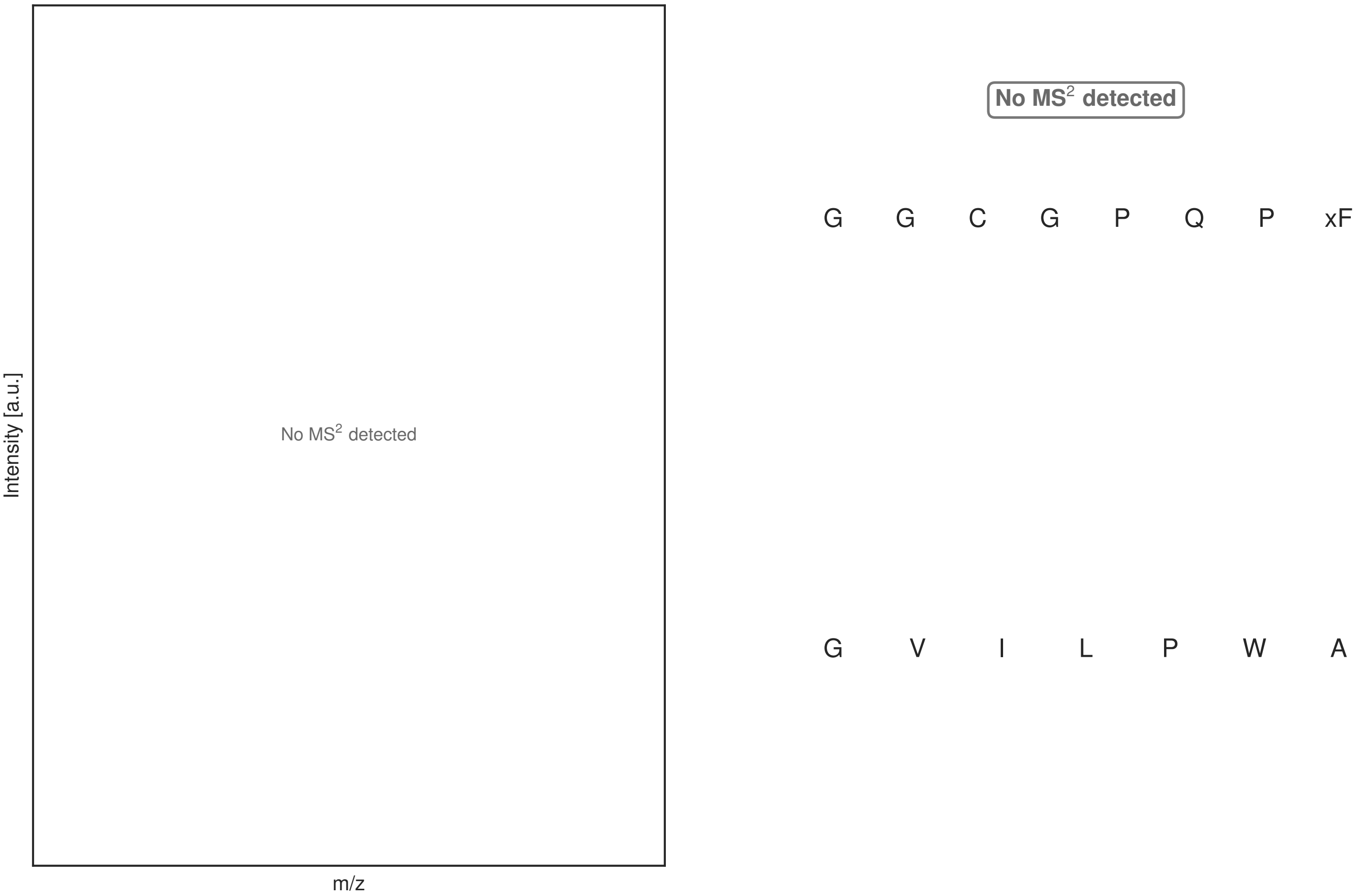


**Figure S109: Modification of the mutant_28 precursor peptide by HypXY.** LC-MS analysis of IMAC-purified precursor peptides from *E. coli* BL21(DE3)Tuner containing splicease-encoding plasmid TJ-C-689 (HypXY) and precursor peptide-encoding plasmid mutant_28 (RSTN1-HypAc-TEV-mutant-28) after expression at 24 °C for 22 h in 1.5 mL cultures after digestion with TEV. Data from n=4 independent biological replicates. Spliced amino acids have an x as a prefix. Spliced refers to a single tyramine excision. A representative experimental replicate is shown in panels **A, B.**

**A** Close-up plot of the representative extracted ion chromatogram (EIC) of the various peptide states; the blue trace corresponds to the Unmodified peptide; the orange trace corresponds to the Spliced peptide. **B** Close-up plot of the m/z values of the peptides detected by the mass spectrometer, with the shaded colored areas corresponding to where the expected m/z values for the peptide m/z traces should occur, with the observed spectrum plotted in black. The Unmodified, Spliced peptide(s) were found in the mass spectrum. **C** Bar plot showing the relative abundance of the various peptide states at charge z=2; the bars represent the average relative abundances normalized to total peak areas, with each individual experimental replicate value plotted as a black dot. **D** Mirrored MS$^2$ plot for the Unmodified peptide. In this mirror plot, the full fragmentation spectrum is plotted in black pointing upward, and the matched fragment b ions (blue) and y ions (red) are plotted pointing downward, mirrored against it, so that each matched fragment can be compared directly with the measured spectrum. **E** MS$^2$ spectrum of the fragments corresponding to the Unmodified peptide sequence, with matched b (blue, above) and y ions (red, below) plotted. **F** Mirrored MS$^2$ plot for the Spliced peptide. In this mirror plot, the full fragmentation spectrum is plotted in black pointing upward, and the matched fragment b ions (blue) and y ions (red) are plotted pointing downward, mirrored against it, so that each matched fragment can be compared directly with the measured spectrum. **G** MS$^2$ spectrum of the fragments corresponding to the Spliced peptide sequence, with matched b (blue, above) and y ions (red, below) plotted.

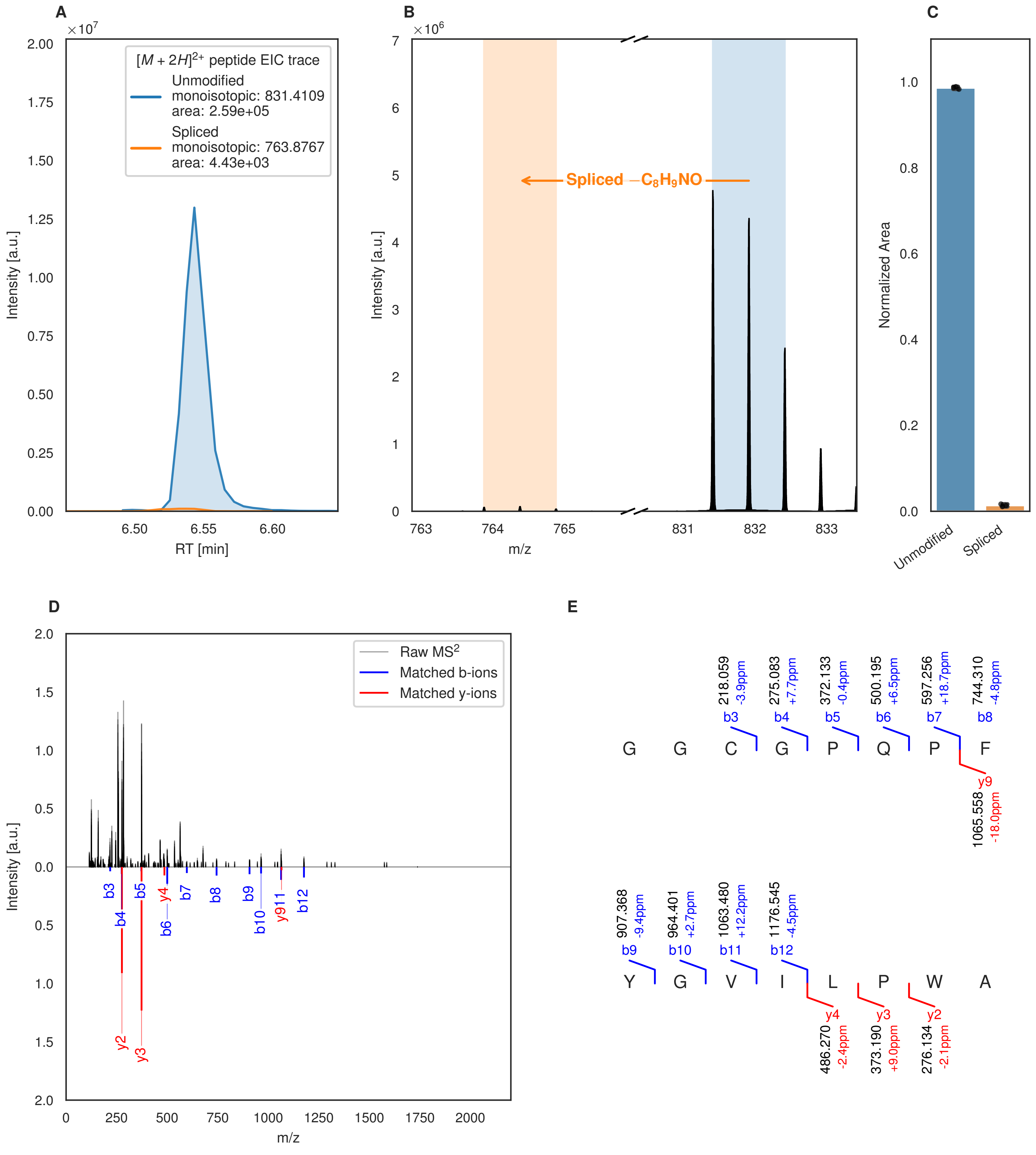
A
B
C
D
E
×10^7
[M + 2H]2+ peptide EIC trace
Unmodified
monoisotopic: 831.4109
area: 2.59e+05
Spliced
monoisotopic: 763.8767
area: 4.43e+03
Intensity [a.u.]
RT [min]
×10^6
Spliced −C8H9NO
m/z
Normalized Area
Unmodified
Spliced
Raw MS2
Matched b-ions
Matched y-ions
b3 b4 b5 y4 b6 b7 b8 b9 b10 y911 b12 y2 y3
G G C G P Q P F
218.059 -3.9ppm b3
275.083 +7.7ppm b4
372.133 -0.4ppm b5
500.195 +6.5ppm b6
597.256 +18.7ppm b7
744.310 -4.8ppm b8
y9 1065.558 -18.0ppm
Y G V I L P W A
907.368 -9.4ppm b9
964.401 +2.7ppm b10
1063.480 +12.2ppm b11
1176.545 -4.5ppm b12
y4 486.270 -2.4ppm
y3 373.190 +9.0ppm
y2 276.134 -2.1ppm

Figure S110 (figure continued from the previous page).

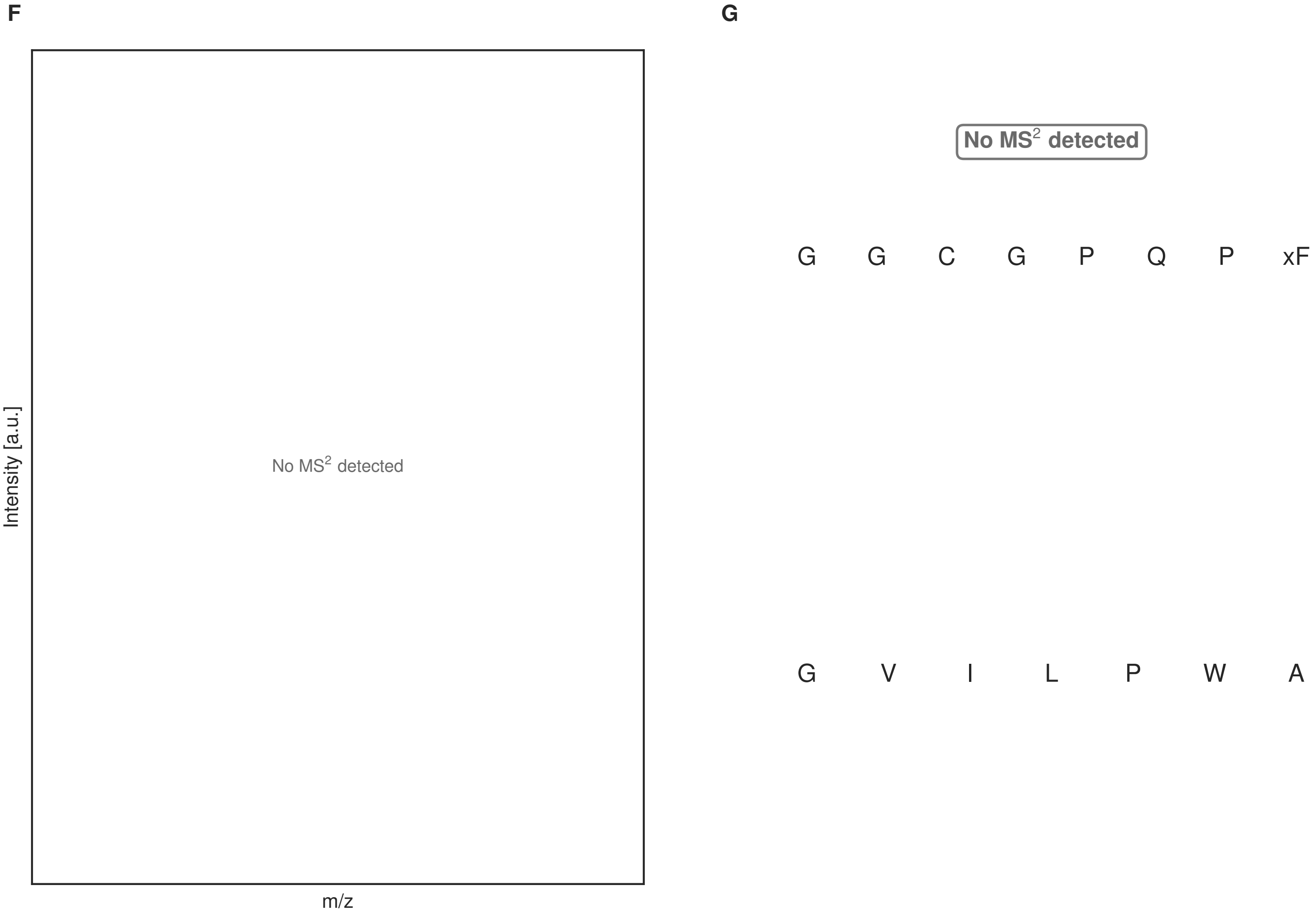


**Figure S110: Modification of the mutant_28 precursor peptide by PlpXY.** LC-MS analysis of IMAC-purified precursor peptides from *E. coli* BL21(DE3)Tuner containing splicease-encoding plasmid TJ-C-559 (PlpXY) and precursor peptide-encoding plasmid mutant_28 (RSTN1-PlpA3-TEV-mutant-28) after expression at 24 °C for 22 h in 1.5 mL cultures after digestion with TEV. Data from n=3 independent biological replicates. Spliced amino acids have an x as a prefix. Spliced refers to a single tyramine excision. A representative experimental replicate is shown in panels **A, B.**

**A** Close-up plot of the representative extracted ion chromatogram (EIC) of the various peptide states; the blue trace corresponds to the Unmodified peptide; the orange trace corresponds to the Spliced peptide. **B** Close-up plot of the m/z values of the peptides detected by the mass spectrometer, with the shaded colored areas corresponding to where the expected m/z values for the peptide m/z traces should occur, with the observed spectrum plotted in black. The Unmodified, Spliced peptide(s) were found in the mass spectrum. **C** Bar plot showing the relative abundance of the various peptide states at charge z=2; the bars represent the average relative abundances normalized to total peak areas, with each individual experimental replicate value plotted as a black dot. **D** Mirrored MS$^2$ plot for the Unmodified peptide. In this mirror plot, the full fragmentation spectrum is plotted in black pointing upward, and the matched fragment b ions (blue) and y ions (red) are plotted pointing downward, mirrored against it, so that each matched fragment can be compared directly with the measured spectrum. **E** MS$^2$ spectrum of the fragments corresponding to the Unmodified peptide sequence, with matched b (blue, above) and y ions (red, below) plotted. **F** Mirrored MS$^2$ plot for the Spliced peptide. In this mirror plot, the full fragmentation spectrum is plotted in black pointing upward, and the matched fragment b ions (blue) and y ions (red) are plotted pointing downward, mirrored against it, so that each matched fragment can be compared directly with the measured spectrum. **G** MS$^2$ spectrum of the fragments corresponding to the Spliced peptide sequence, with matched b (blue, above) and y ions (red, below) plotted.

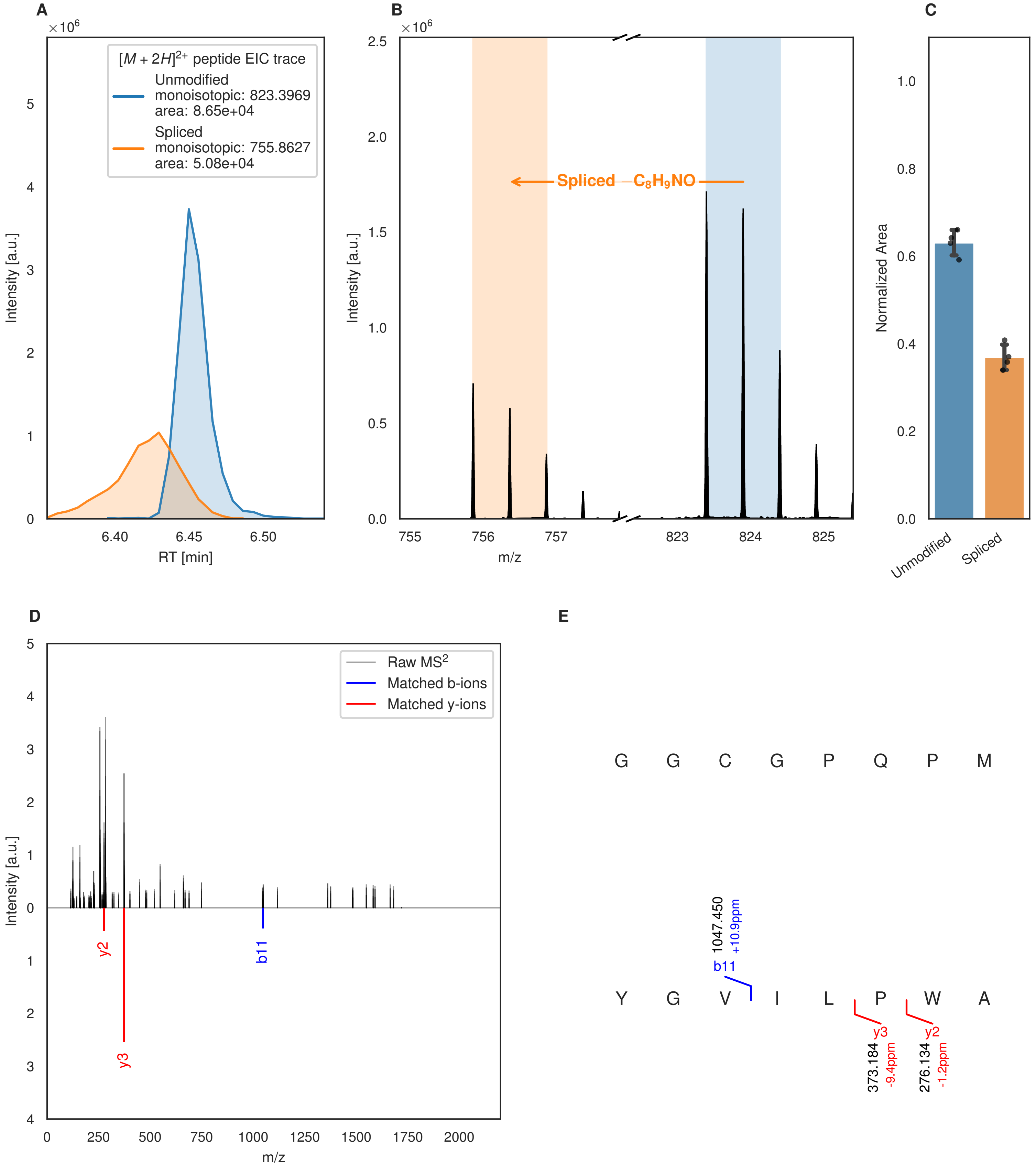
A
B
C
D
E
×10^6
[M + 2H]2+ peptide EIC trace
Unmodified
monoisotopic: 823.3969
area: 8.65e+04
Spliced
monoisotopic: 755.8627
area: 5.08e+04
Intensity [a.u.]
RT [min]
6.40
6.45
6.50
Spliced −C8H9NO
m/z
755
756
757
823
824
825
Normalized Area
Unmodified
Spliced
Raw MS2
Matched b-ions
Matched y-ions
y2
y3
b11
G G C G P Q P M
Y G V I L P W A
1047.450
+10.9ppm
373.184
-9.4ppm
276.134
-1.2ppm

F

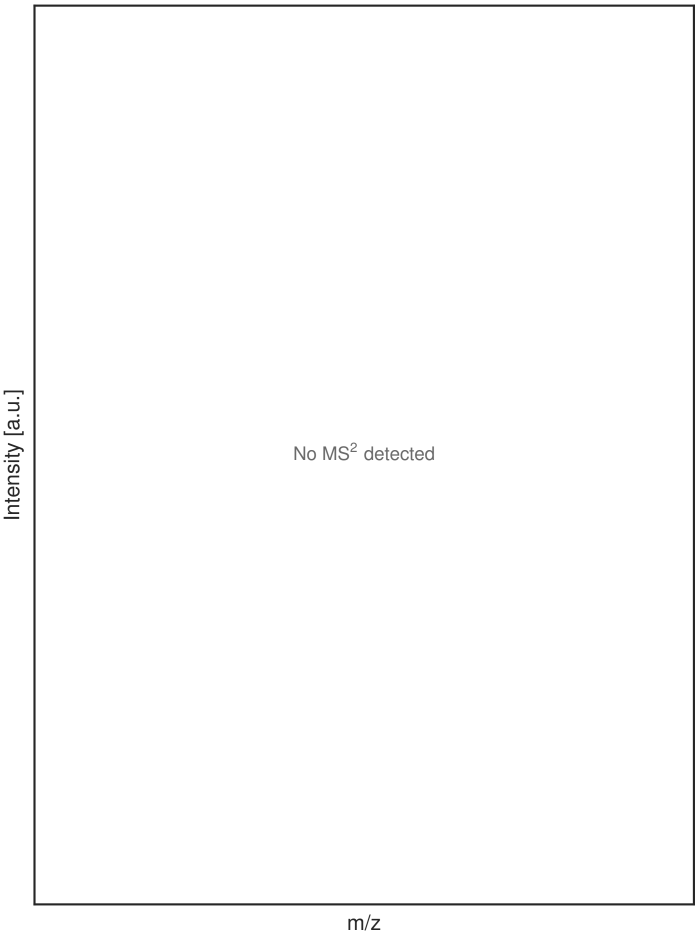


G

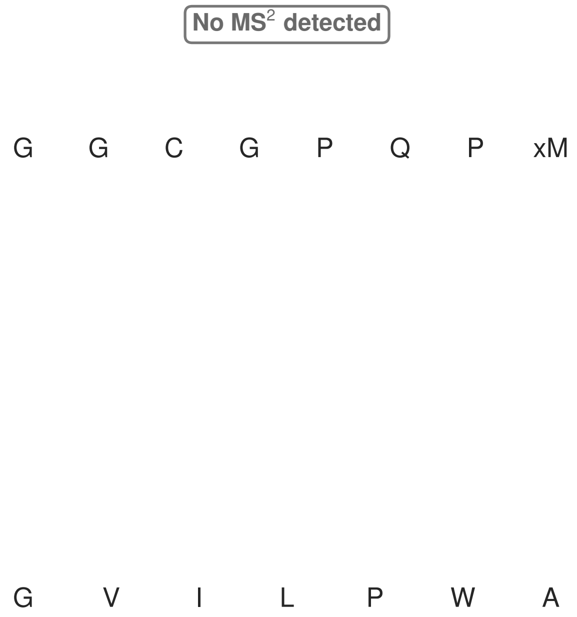


**Figure S111: Modification of the mutant_29 precursor peptide by HypXY.** LC-MS analysis of IMAC-purified precursor peptides from *E. coli* BL21(DE3)Tuner containing splicease-encoding plasmid TJ-C-689 (HypXY) and precursor peptide-encoding plasmid mutant_29 (RSTN1-HypAc-TEV-mutant-29) after expression at 24 °C for 22 h in 1.5 mL cultures after digestion with TEV. Data from n=4 independent biological replicates. Spliced amino acids have an x as a prefix. Spliced refers to a single tyramine excision. A representative experimental replicate is shown in panels **A, B.**

**A** Close-up plot of the representative extracted ion chromatogram (EIC) of the various peptide states; the blue trace corresponds to the Unmodified peptide; the orange trace corresponds to the Spliced peptide. **B** Close-up plot of the m/z values of the peptides detected by the mass spectrometer, with the shaded colored areas corresponding to where the expected m/z values for the peptide m/z traces should occur, with the observed spectrum plotted in black. The Unmodified, Spliced peptide(s) were found in the mass spectrum. **C** Bar plot showing the relative abundance of the various peptide states at charge z=2; the bars represent the average relative abundances normalized to total peak areas, with each individual experimental replicate value plotted as a black dot. **D** Mirrored MS$^2$ plot for the Unmodified peptide. In this mirror plot, the full fragmentation spectrum is plotted in black pointing upward, and the matched fragment b ions (blue) and y ions (red) are plotted pointing downward, mirrored against it, so that each matched fragment can be compared directly with the measured spectrum. **E** MS$^2$ spectrum of the fragments corresponding to the Unmodified peptide sequence, with matched b (blue, above) and y ions (red, below) plotted. **F** Mirrored MS$^2$ plot for the Spliced peptide. In this mirror plot, the full fragmentation spectrum is plotted in black pointing upward, and the matched fragment b ions (blue) and y ions (red) are plotted pointing downward, mirrored against it, so that each matched fragment can be compared directly with the measured spectrum. **G** MS$^2$ spectrum of the fragments corresponding to the Spliced peptide sequence, with matched b (blue, above) and y ions (red, below) plotted.

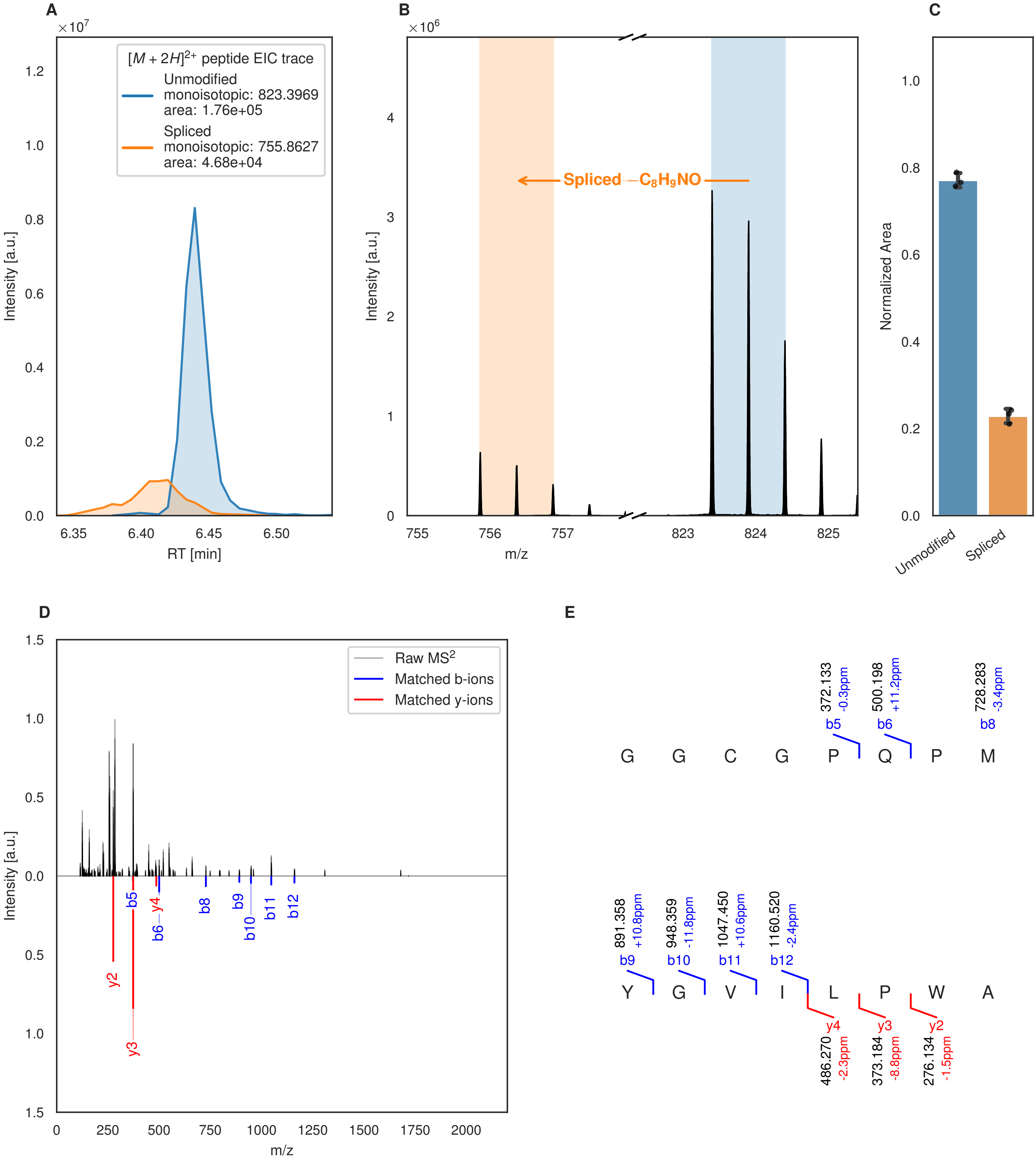
A
[M + 2H]2+ peptide EIC trace
Unmodified
monoisotopic: 823.3969
area: 1.76e+05
Spliced
monoisotopic: 755.8627
area: 4.68e+04
Intensity [a.u.]
RT [min]
B
Spliced −C8H9NO
Intensity [a.u.]
m/z
C
Normalized Area
Unmodified
Spliced
D
Raw MS2
Matched b-ions
Matched y-ions
Intensity [a.u.]
m/z
E
G G C G P Q P M
Y G V I L P W A
372.133 -0.3ppm b5
500.198 +11.2ppm b6
728.283 -3.4ppm b8
891.358 +10.8ppm b9
948.359 -11.8ppm b10
1047.450 +10.6ppm b11
1160.520 -2.4ppm b12
y4 486.270 -2.3ppm
y3 373.184 -8.8ppm
y2 276.134 -1.5ppm

**F**

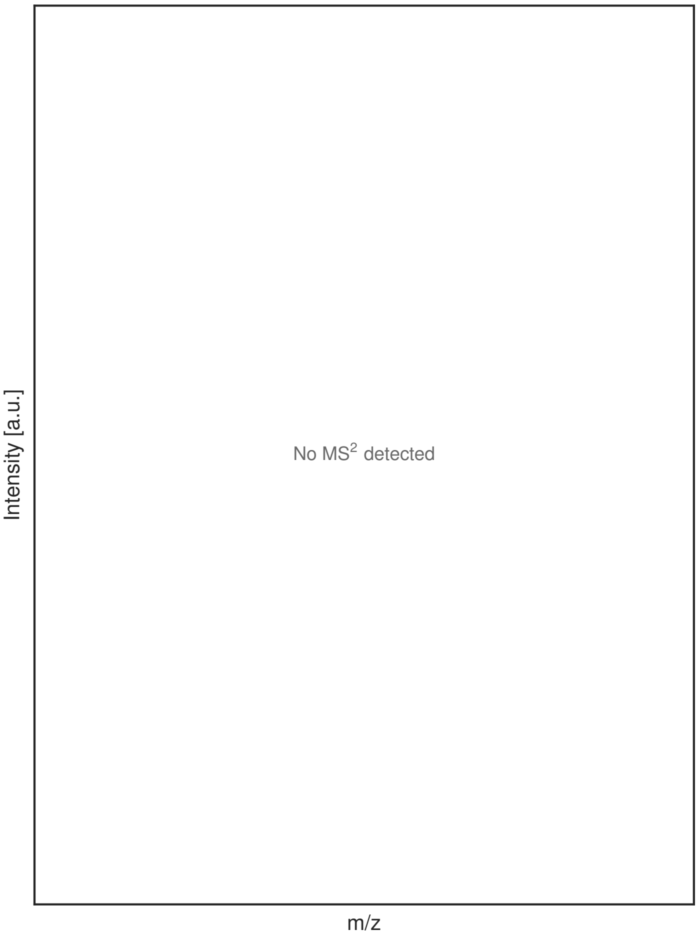


**G**

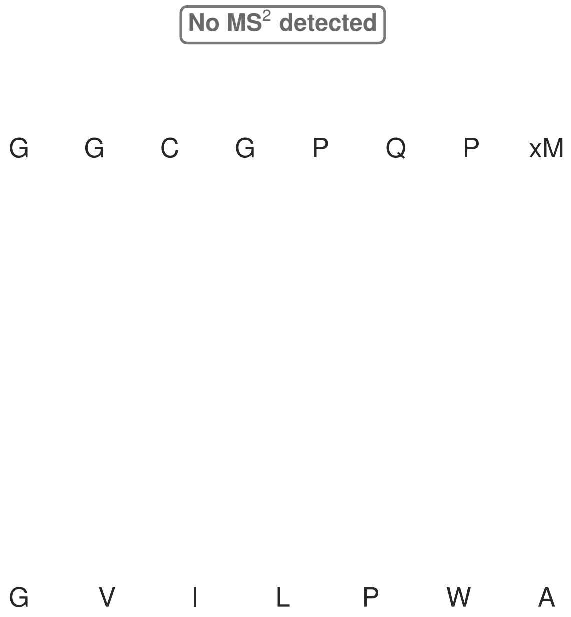


**Figure S112: Modification of the mutant_29 precursor peptide by PlpXY.** LC-MS analysis of IMAC-purified precursor peptides from *E. coli* BL21(DE3)Tuner containing splicease-encoding plasmid TJ-C-559 (PlpXY) and precursor peptide-encoding plasmid mutant_29 (RSTN1-PlpA3-TEV-mutant-29) after expression at 24 °C for 22 h in 1.5 mL cultures after digestion with TEV. Data from n=3 independent biological replicates. Spliced amino acids have an x as a prefix. Spliced refers to a single tyramine excision. A representative experimental replicate is shown in panels **A, B.**

**A** Close-up plot of the representative extracted ion chromatogram (EIC) of the various peptide states; the blue trace corresponds to the Unmodified peptide; the orange trace corresponds to the Spliced peptide. **B** Close-up plot of the m/z values of the peptides detected by the mass spectrometer, with the shaded colored areas corresponding to where the expected m/z values for the peptide m/z traces should occur, with the observed spectrum plotted in black. The Unmodified, Spliced peptide(s) were found in the mass spectrum. **C** Bar plot showing the relative abundance of the various peptide states at charge z=2; the bars represent the average relative abundances normalized to total peak areas, with each individual experimental replicate value plotted as a black dot. **D** Mirrored MS$^2$ plot for the Unmodified peptide. In this mirror plot, the full fragmentation spectrum is plotted in black pointing upward, and the matched fragment b ions (blue) and y ions (red) are plotted pointing downward, mirrored against it, so that each matched fragment can be compared directly with the measured spectrum. **E** MS$^2$ spectrum of the fragments corresponding to the Unmodified peptide sequence, with matched b (blue, above) and y ions (red, below) plotted. **F** Mirrored MS$^2$ plot for the Spliced peptide. In this mirror plot, the full fragmentation spectrum is plotted in black pointing upward, and the matched fragment b ions (blue) and y ions (red) are plotted pointing downward, mirrored against it, so that each matched fragment can be compared directly with the measured spectrum. **G** MS$^2$ spectrum of the fragments corresponding to the Spliced peptide sequence, with matched b (blue, above) and y ions (red, below) plotted.

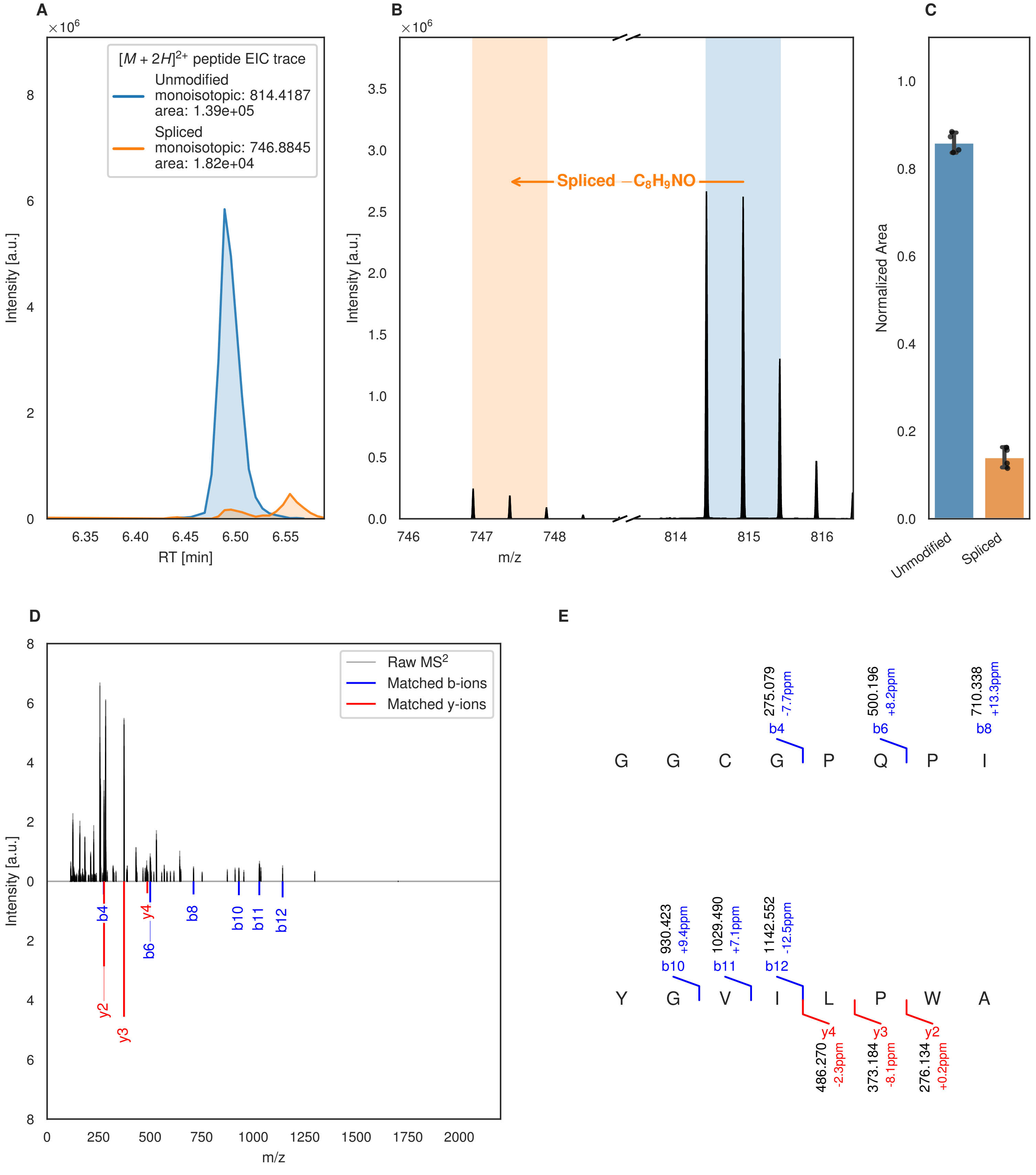
A
×10^6
[M + 2H]2+ peptide EIC trace
Unmodified
monoisotopic: 814.4187
area: 1.39e+05
Spliced
monoisotopic: 746.8845
area: 1.82e+04
Intensity [a.u.]
RT [min]
B
×10^6
Spliced −C8H9NO
Intensity [a.u.]
m/z
C
Normalized Area
Unmodified
Spliced
D
Raw MS2
Matched b-ions
Matched y-ions
Intensity [a.u.]
m/z
b4
y4
b6
b8
b10
b11
b12
y2
y3
E
275.079
-7.7ppm
b4
500.196
+8.2ppm
b6
710.338
+13.3ppm
b8
G G C G P Q P I
930.423
+9.4ppm
b10
1029.490
+7.1ppm
b11
1142.552
-12.5ppm
b12
Y G V I L P W A
y4
486.270
-2.3ppm
y3
373.184
-8.1ppm
y2
276.134
+0.2ppm

Figure S113 (figure continued from the previous page).

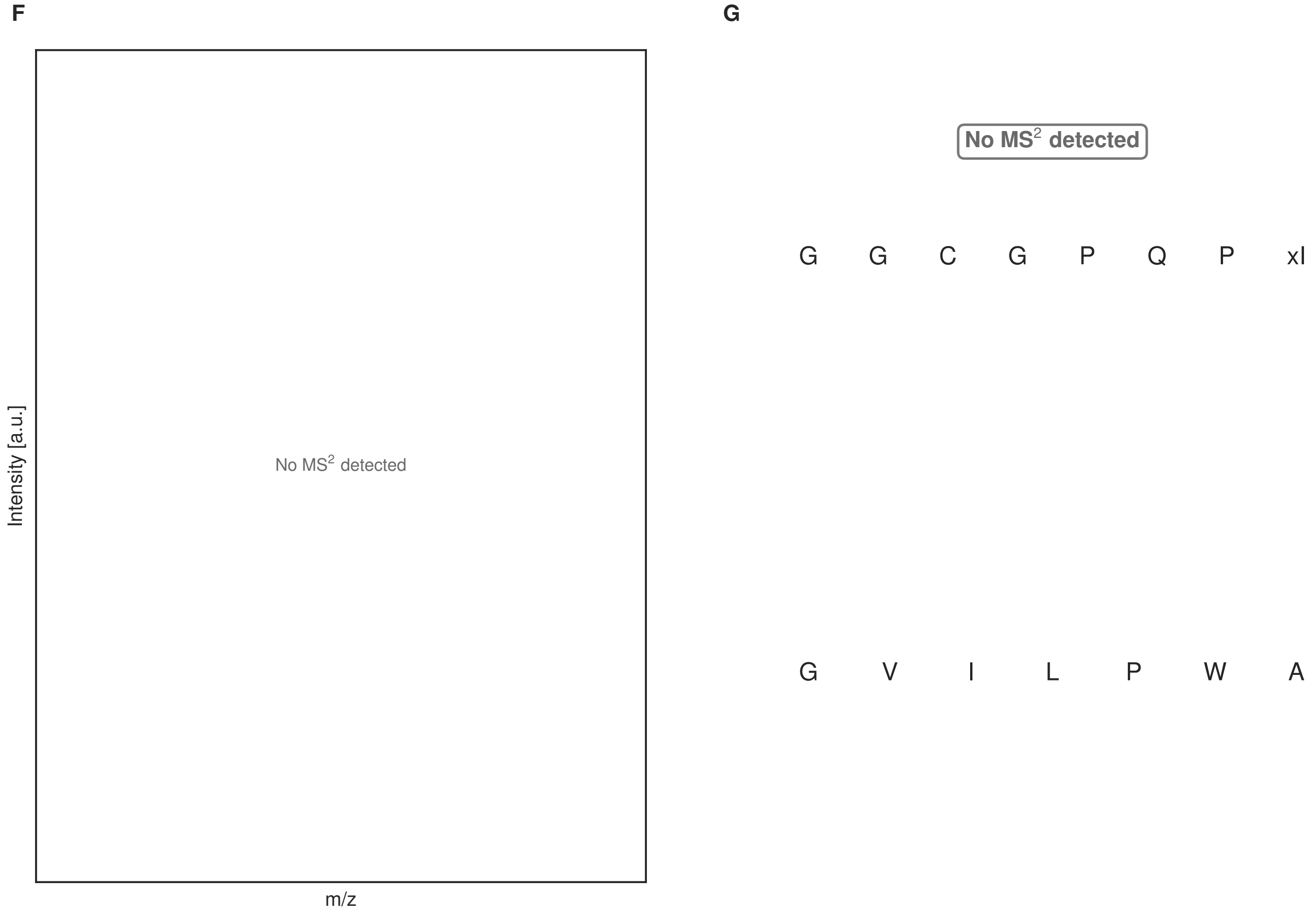


**Figure S113: Modification of the mutant_30 precursor peptide by HypXY.** LC-MS analysis of IMAC-purified precursor peptides from *E. coli* BL21(DE3)Tuner containing splicease-encoding plasmid TJ-C-689 (HypXY) and precursor peptide-encoding plasmid mutant_30 (RSTN1-HypAc-TEV-mutant-30) after expression at 24 °C for 22 h in 1.5 mL cultures after digestion with TEV. Data from n=4 independent biological replicates. Spliced amino acids have an x as a prefix. Spliced refers to a single tyramine excision. A representative experimental replicate is shown in panels **A, B.**
**A** Close-up plot of the representative extracted ion chromatogram (EIC) of the various peptide states; the blue trace corresponds to the Unmodified peptide; the orange trace corresponds to the Spliced peptide. **B** Close-up plot of the m/z values of the peptides detected by the mass spectrometer, with the shaded colored areas corresponding to where the expected m/z values for the peptide m/z traces should occur, with the observed spectrum plotted in black. The Unmodified, Spliced peptide(s) were found in the mass spectrum. **C** Bar plot showing the relative abundance of the various peptide states at charge z=2; the bars represent the average relative abundances normalized to total peak areas, with each individual experimental replicate value plotted as a black dot. **D** Mirrored $MS^2$ plot for the Unmodified peptide. In this mirror plot, the full fragmentation spectrum is plotted in black pointing upward, and the matched fragment b ions (blue) and y ions (red) are plotted pointing downward, mirrored against it, so that each matched fragment can be compared directly with the measured spectrum. **E** $MS^2$ spectrum of the fragments corresponding to the Unmodified peptide sequence, with matched b (blue, above) and y ions (red, below) plotted. **F** Mirrored $MS^2$ plot for the Spliced peptide. In this mirror plot, the full fragmentation spectrum is plotted in black pointing upward, and the matched fragment b ions (blue) and y ions (red) are plotted pointing downward, mirrored against it, so that each matched fragment can be compared directly with the measured spectrum. **G** $MS^2$ spectrum of the fragments corresponding to the Spliced peptide sequence, with matched b (blue, above) and y ions (red, below) plotted.

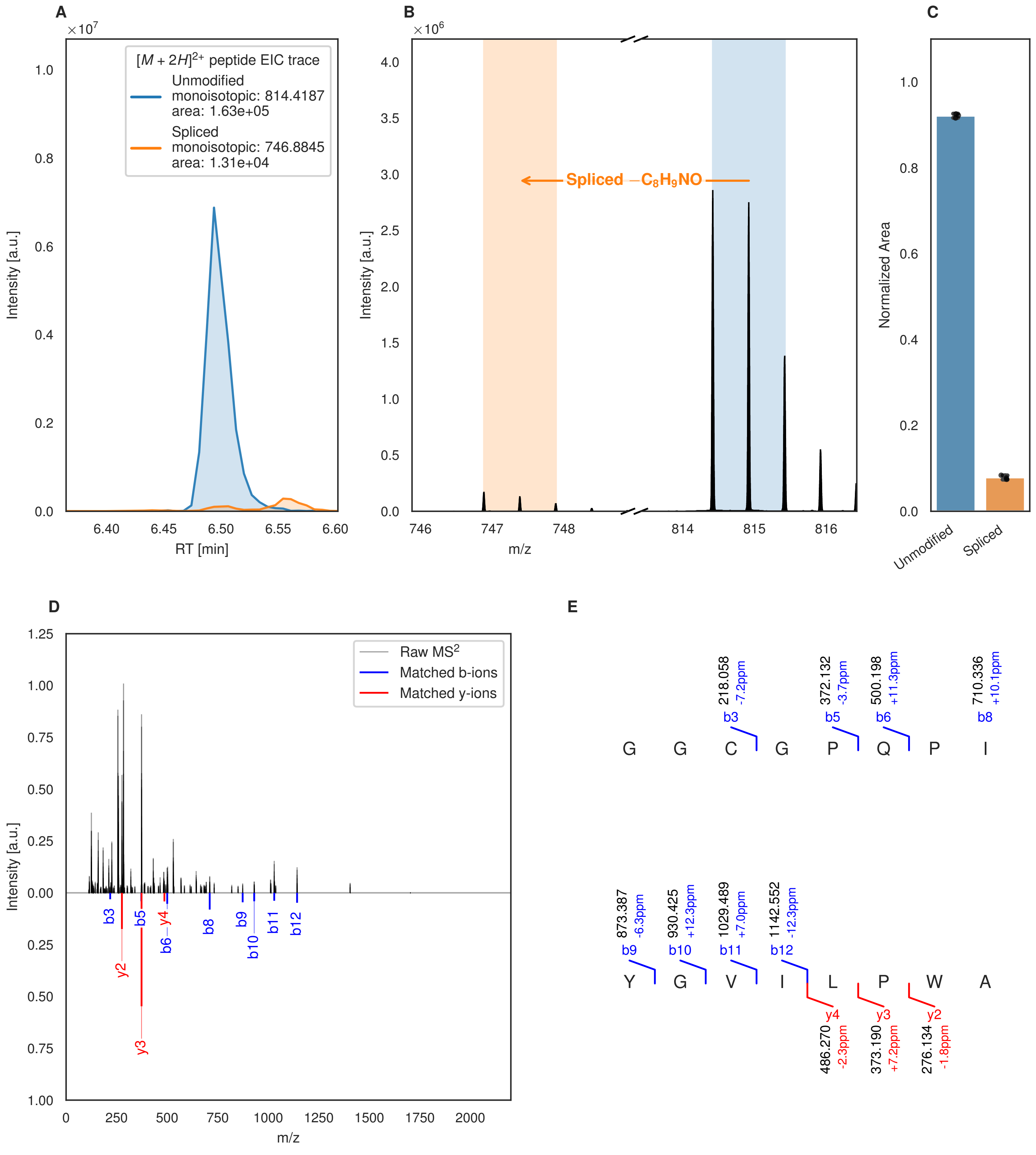
A
$[M+2H]^{2+}$ peptide EIC trace
Unmodified
monoisotopic: 814.4187
area: 1.63e+05
Spliced
monoisotopic: 746.8845
area: 1.31e+04
Intensity [a.u.]
RT [min]
B
Spliced $-C_8H_9NO$
Intensity [a.u.]
m/z
C
Normalized Area
Unmodified
Spliced
D
Raw MS2
Matched b-ions
Matched y-ions
Intensity [a.u.]
m/z
E
218.058 -7.2ppm b3
372.132 -3.7ppm b5
500.198 +11.3ppm b6
710.336 +10.1ppm b8
G G C G P Q P I
873.387 -6.3ppm b9
930.425 +12.3ppm b10
1029.489 +7.0ppm b11
1142.552 -12.3ppm b12
Y G V I L P W A
y4 486.270 -2.3ppm
y3 373.190 +7.2ppm
y2 276.134 -1.8ppm

Figure S114 (figure continued from the previous page).

F

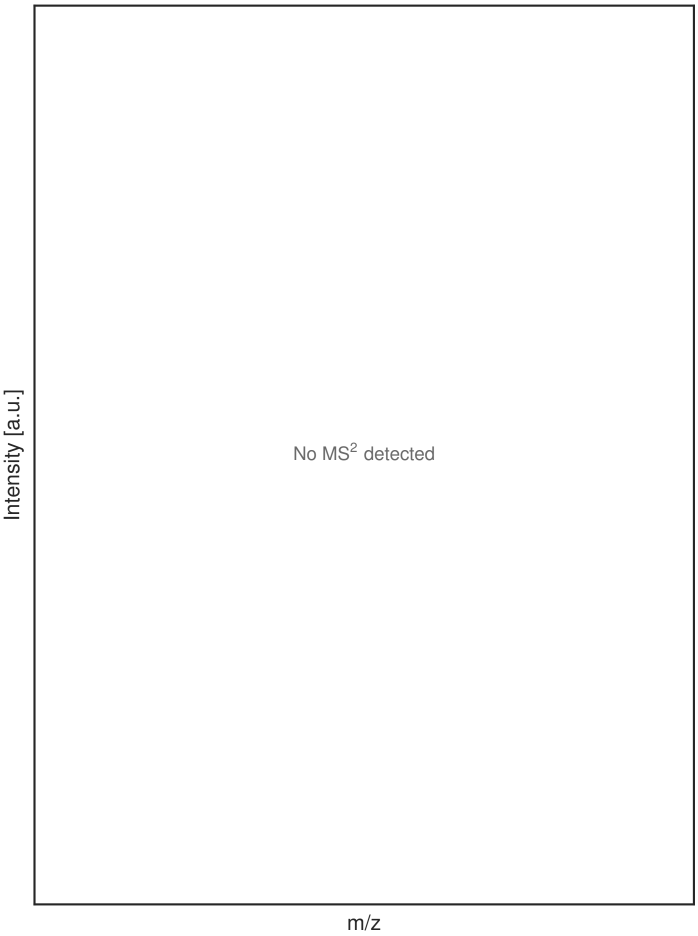


G

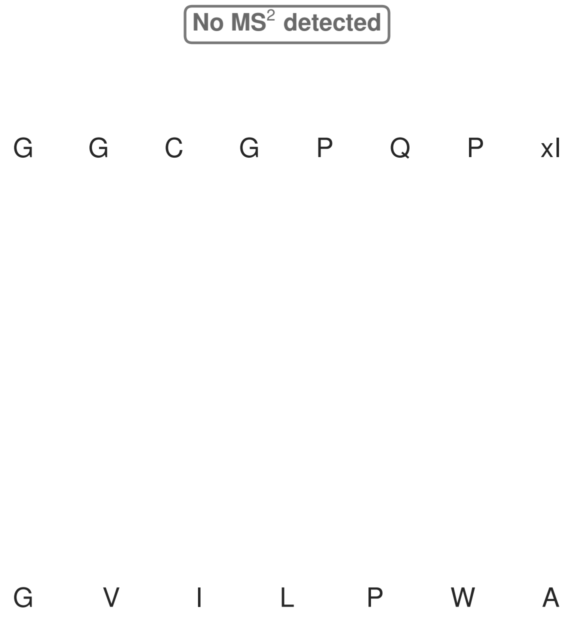


**Figure S114: Modification of the mutant_30 precursor peptide by PlpXY.** LC-MS analysis of IMAC-purified precursor peptides from *E. coli* BL21(DE3)Tuner containing splicease-encoding plasmid TJ-C-559 (PlpXY) and precursor peptide-encoding plasmid mutant_30 (RSTN1-PlpA3-TEV-mutant-30) after expression at 24 °C for 22 h in 1.5 mL cultures after digestion with TEV. Data from n=3 independent biological replicates. Spliced amino acids have an x as a prefix. Spliced refers to a single tyramine excision. A representative experimental replicate is shown in panels **A, B.**

**A** Close-up plot of the representative extracted ion chromatogram (EIC) of the various peptide states; the blue trace corresponds to the Unmodified peptide; the orange trace corresponds to the Spliced peptide. **B** Close-up plot of the m/z values of the peptides detected by the mass spectrometer, with the shaded colored areas corresponding to where the expected m/z values for the peptide m/z traces should occur, with the observed spectrum plotted in black. The Unmodified, Spliced peptide(s) were found in the mass spectrum. **C** Bar plot showing the relative abundance of the various peptide states at charge z=2; the bars represent the average relative abundances normalized to total peak areas, with each individual experimental replicate value plotted as a black dot. **D** Mirrored MS$^2$ plot for the Unmodified peptide. In this mirror plot, the full fragmentation spectrum is plotted in black pointing upward, and the matched fragment b ions (blue) and y ions (red) are plotted pointing downward, mirrored against it, so that each matched fragment can be compared directly with the measured spectrum. **E** MS$^2$ spectrum of the fragments corresponding to the Unmodified peptide sequence, with matched b (blue, above) and y ions (red, below) plotted. **F** Mirrored MS$^2$ plot for the Spliced peptide. In this mirror plot, the full fragmentation spectrum is plotted in black pointing upward, and the matched fragment b ions (blue) and y ions (red) are plotted pointing downward, mirrored against it, so that each matched fragment can be compared directly with the measured spectrum. **G** MS$^2$ spectrum of the fragments corresponding to the Spliced peptide sequence, with matched b (blue, above) and y ions (red, below) plotted.

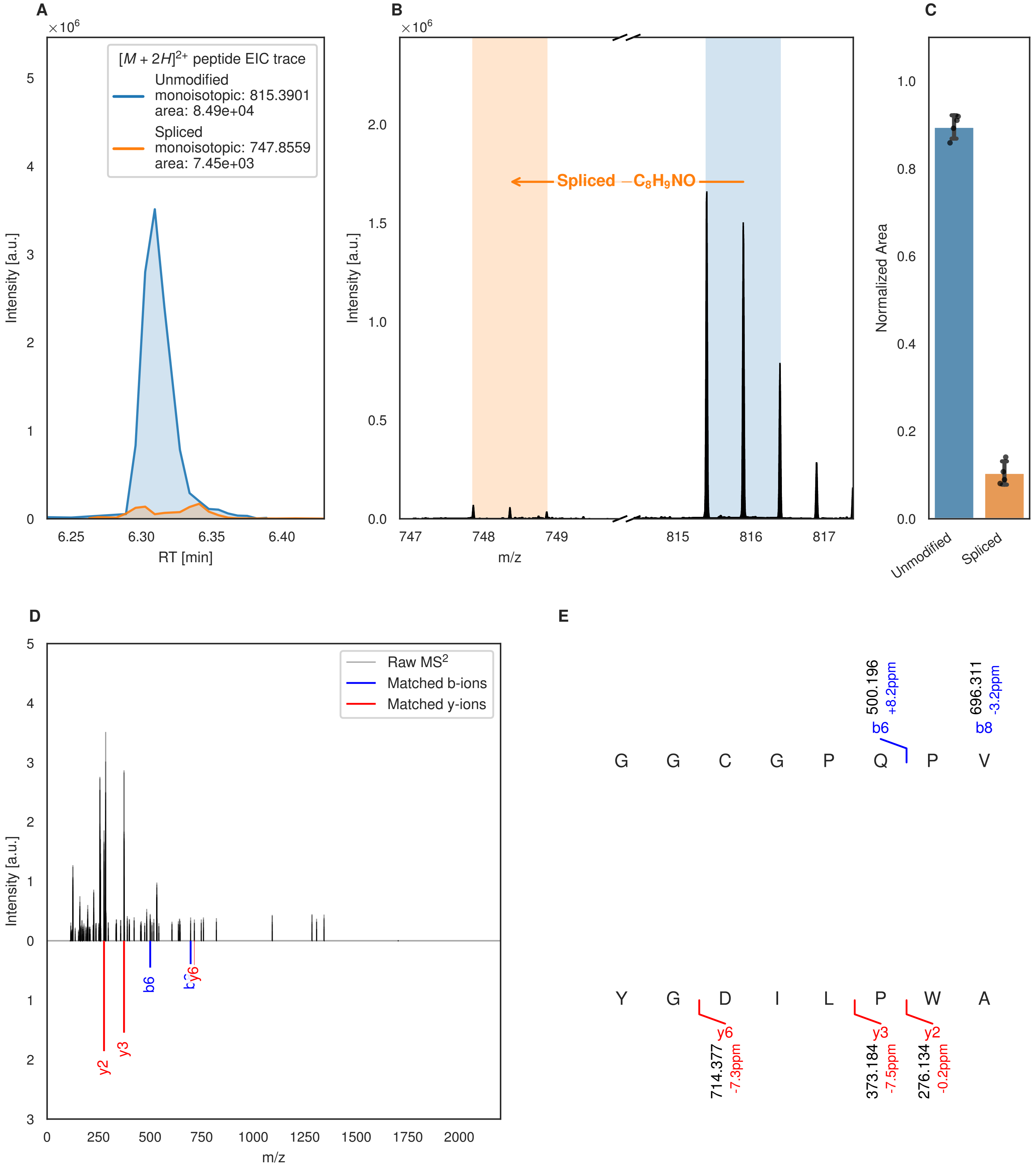
A
B
C
D
E
×10^6
$[M+2H]^{2+}$ peptide EIC trace
Unmodified
monoisotopic: 815.3901
area: 8.49e+04
Spliced
monoisotopic: 747.8559
area: 7.45e+03
Intensity [a.u.]
RT [min]
6.25
6.30
6.35
6.40
Spliced $-C_8H_9NO$
m/z
747
748
749
815
816
817
Normalized Area
Unmodified
Spliced
Raw MS$^2$
Matched b-ions
Matched y-ions
y2
y3
b6
y6
0
250
500
750
1000
1250
1500
1750
2000
G G C G P Q P V
b6
500.196
+8.2ppm
b8
696.311
-3.2ppm
Y G D I L P W A
y6
714.377
-7.3ppm
y3
373.184
-7.5ppm
y2
276.134
-0.2ppm

Figure S115 (figure continued from the previous page).

F

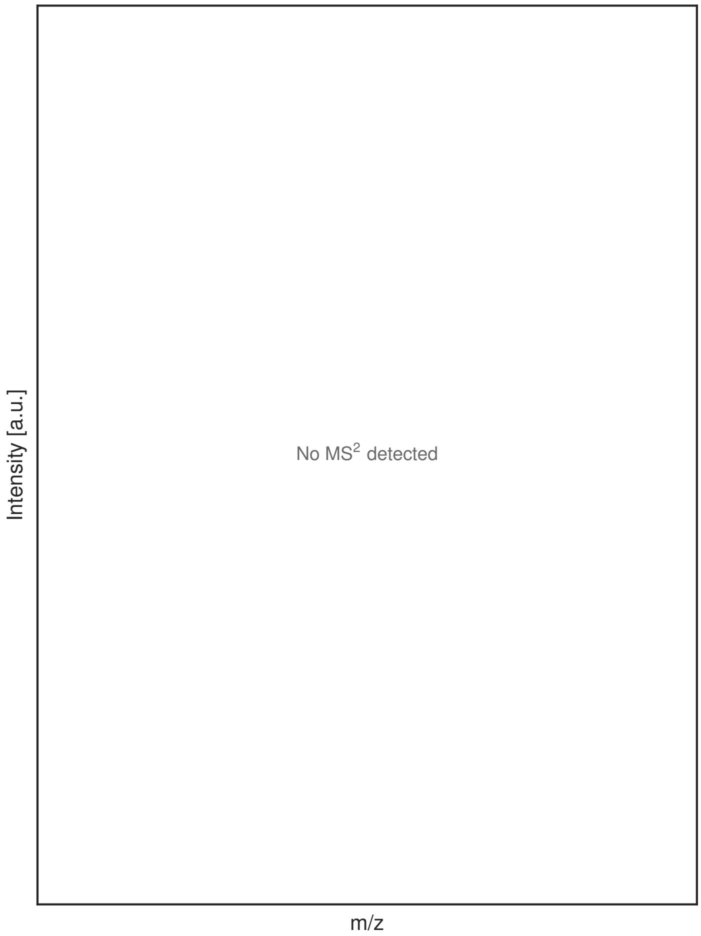


G

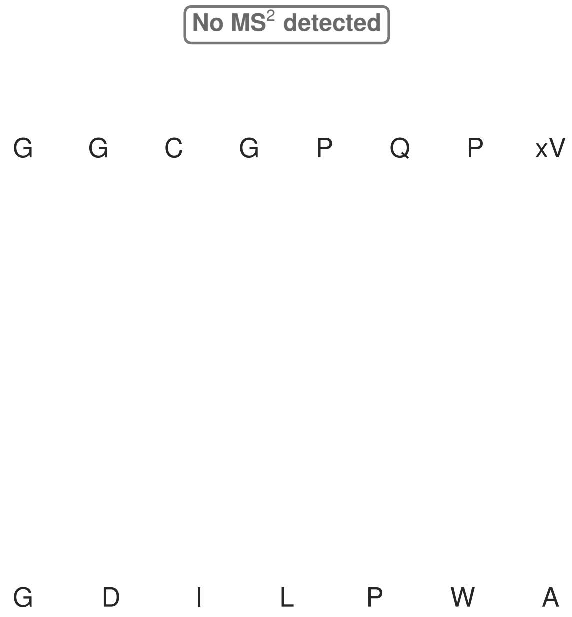


**Figure S115: Modification of the mutant_31 precursor peptide by HypXY.** LC-MS analysis of IMAC-purified precursor peptides from *E. coli* BL21(DE3)Tuner containing splicease-encoding plasmid TJ-C-689 (HypXY) and precursor peptide-encoding plasmid mutant_31 (RSTN1-HypAc-TEV-mutant-31) after expression at 24 °C for 22 h in 1.5 mL cultures after digestion with TEV. Data from n=4 independent biological replicates. Spliced amino acids have an x as a prefix. Spliced refers to a single tyramine excision. A representative experimental replicate is shown in panels **A, B.**

**A** Close-up plot of the representative extracted ion chromatogram (EIC) of the various peptide states; the blue trace corresponds to the Unmodified peptide; the orange trace corresponds to the Spliced peptide. **B** Close-up plot of the m/z values of the peptides detected by the mass spectrometer, with the shaded colored areas corresponding to where the expected m/z values for the peptide m/z traces should occur, with the observed spectrum plotted in black. The Unmodified, Spliced peptide(s) were found in the mass spectrum. **C** Bar plot showing the relative abundance of the various peptide states at charge z=2; the bars represent the average relative abundances normalized to total peak areas, with each individual experimental replicate value plotted as a black dot. **D** Mirrored MS$^2$ plot for the Unmodified peptide. In this mirror plot, the full fragmentation spectrum is plotted in black pointing upward, and the matched fragment b ions (blue) and y ions (red) are plotted pointing downward, mirrored against it, so that each matched fragment can be compared directly with the measured spectrum. **E** MS$^2$ spectrum of the fragments corresponding to the Unmodified peptide sequence, with matched b (blue, above) and y ions (red, below) plotted. **F** Mirrored MS$^2$ plot for the Spliced peptide. In this mirror plot, the full fragmentation spectrum is plotted in black pointing upward, and the matched fragment b ions (blue) and y ions (red) are plotted pointing downward, mirrored against it, so that each matched fragment can be compared directly with the measured spectrum. **G** MS$^2$ spectrum of the fragments corresponding to the Spliced peptide sequence, with matched b (blue, above) and y ions (red, below) plotted.

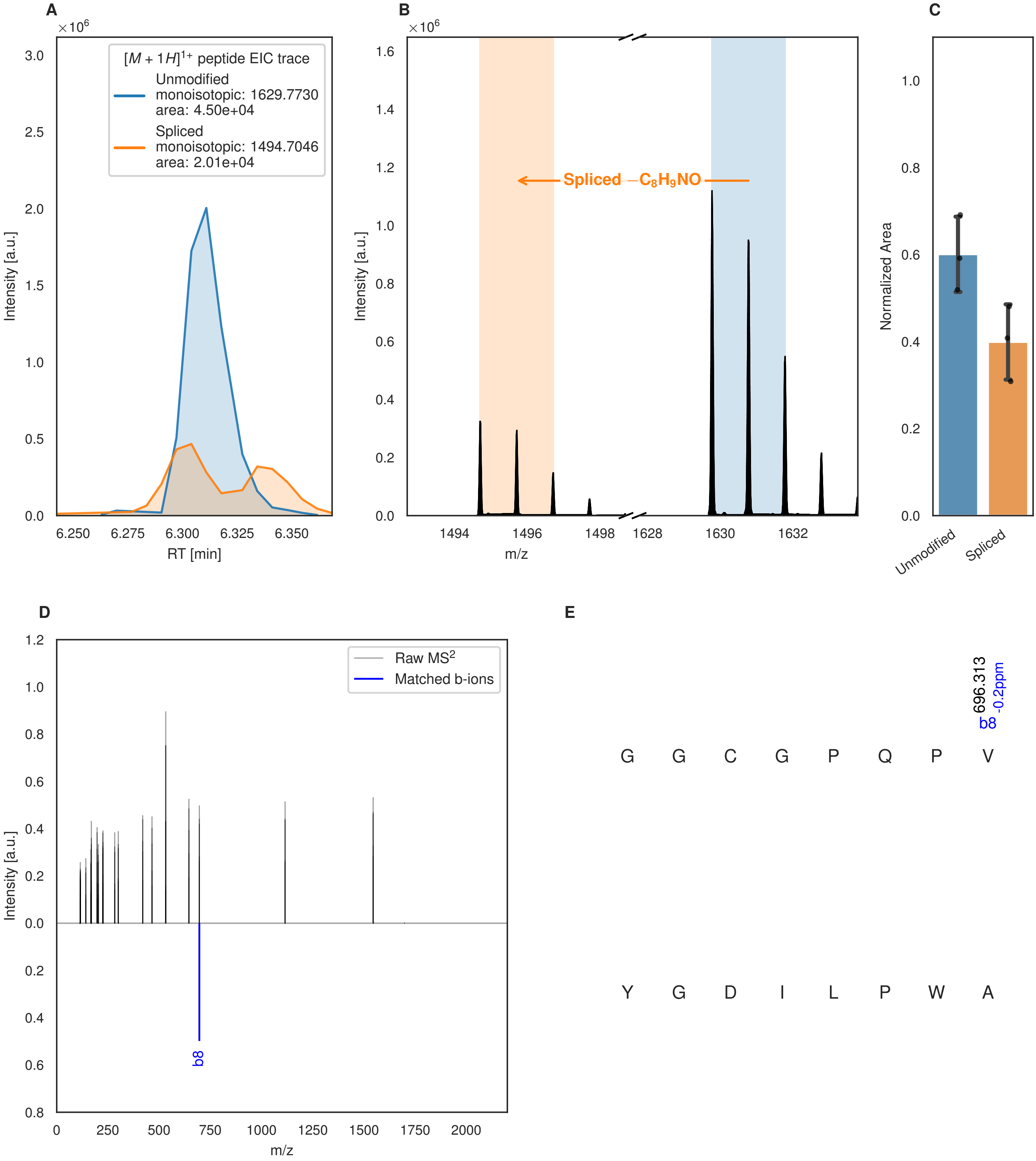
A
×10^6
[M + 1H]1+ peptide EIC trace
Unmodified
monoisotopic: 1629.7730
area: 4.50e+04
Spliced
monoisotopic: 1494.7046
area: 2.01e+04
Intensity [a.u.]
RT [min]
B
×10^6
Spliced −C8H9NO
Intensity [a.u.]
m/z
C
Normalized Area
Unmodified
Spliced
D
Raw MS2
Matched b-ions
Intensity [a.u.]
b8
m/z
E
696.313
-0.2ppm
b8
G G C G P Q P V
Y G D I L P W A

Figure S116 (figure continued from the previous page).

F

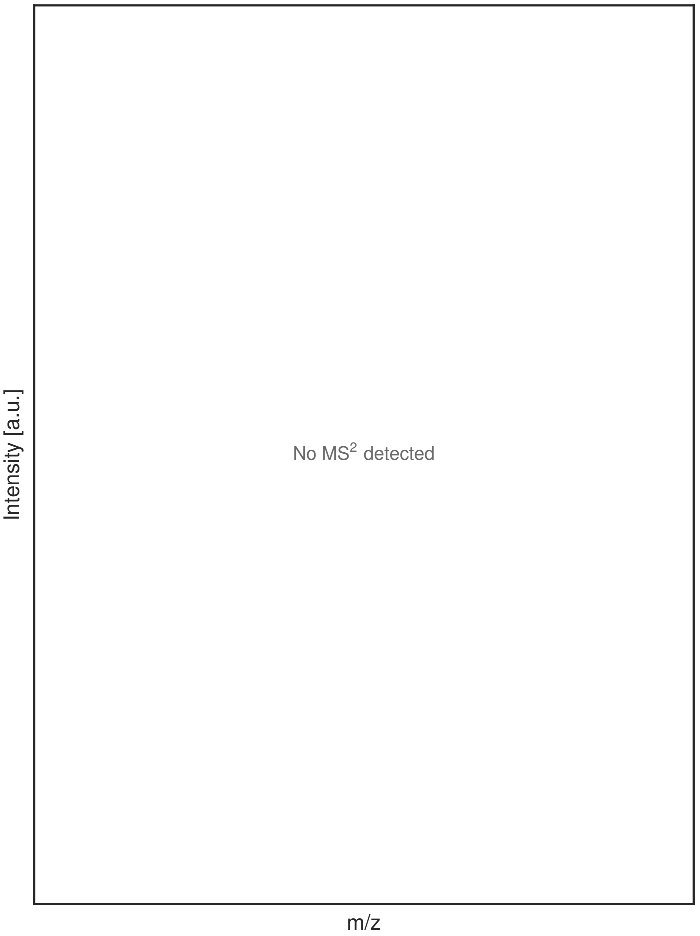


G

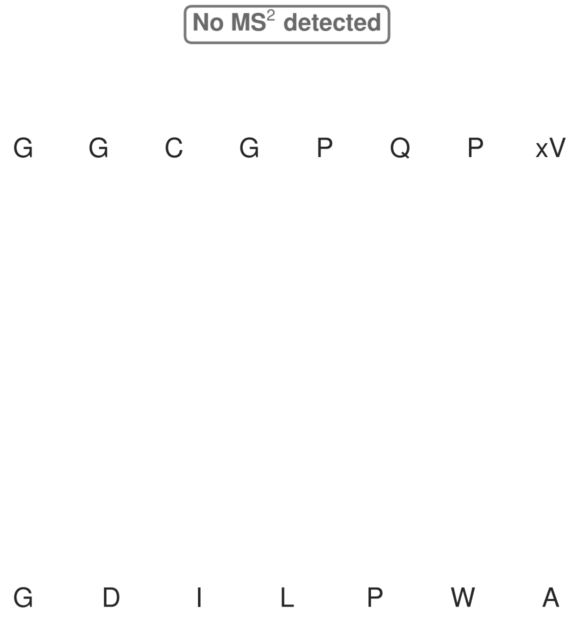


**Figure S116: Modification of the mutant_31 precursor peptide by PlpXY.** LC-MS analysis of IMAC-purified precursor peptides from *E. coli* BL21(DE3)Tuner containing splicease-encoding plasmid TJ-C-559 (PlpXY) and precursor peptide-encoding plasmid mutant_31 (RSTN1-PlpA3-TEV-mutant-31) after expression at 24 °C for 22 h in 1.5 mL cultures after digestion with TEV. Data from n=3 independent biological replicates. Spliced amino acids have an x as a prefix. Spliced refers to a single tyramine excision. A representative experimental replicate is shown in panels **A, B.**

**A** Close-up plot of the representative extracted ion chromatogram (EIC) of the various peptide states; the blue trace corresponds to the Unmodified peptide; the orange trace corresponds to the Spliced peptide. **B** Close-up plot of the m/z values of the peptides detected by the mass spectrometer, with the shaded colored areas corresponding to where the expected m/z values for the peptide m/z traces should occur, with the observed spectrum plotted in black. The Unmodified, Spliced peptide(s) were found in the mass spectrum. **C** Bar plot showing the relative abundance of the various peptide states at charge z=1; the bars represent the average relative abundances normalized to total peak areas, with each individual experimental replicate value plotted as a black dot. **D** Mirrored MS$^2$ plot for the Unmodified peptide. In this mirror plot, the full fragmentation spectrum is plotted in black pointing upward, and the matched fragment b ions (blue) and y ions (red) are plotted pointing downward, mirrored against it, so that each matched fragment can be compared directly with the measured spectrum. **E** MS$^2$ spectrum of the fragments corresponding to the Unmodified peptide sequence, with matched b (blue, above) and y ions (red, below) plotted. **F** Mirrored MS$^2$ plot for the Spliced peptide. In this mirror plot, the full fragmentation spectrum is plotted in black pointing upward, and the matched fragment b ions (blue) and y ions (red) are plotted pointing downward, mirrored against it, so that each matched fragment can be compared directly with the measured spectrum. **G** MS$^2$ spectrum of the fragments corresponding to the Spliced peptide sequence, with matched b (blue, above) and y ions (red, below) plotted.

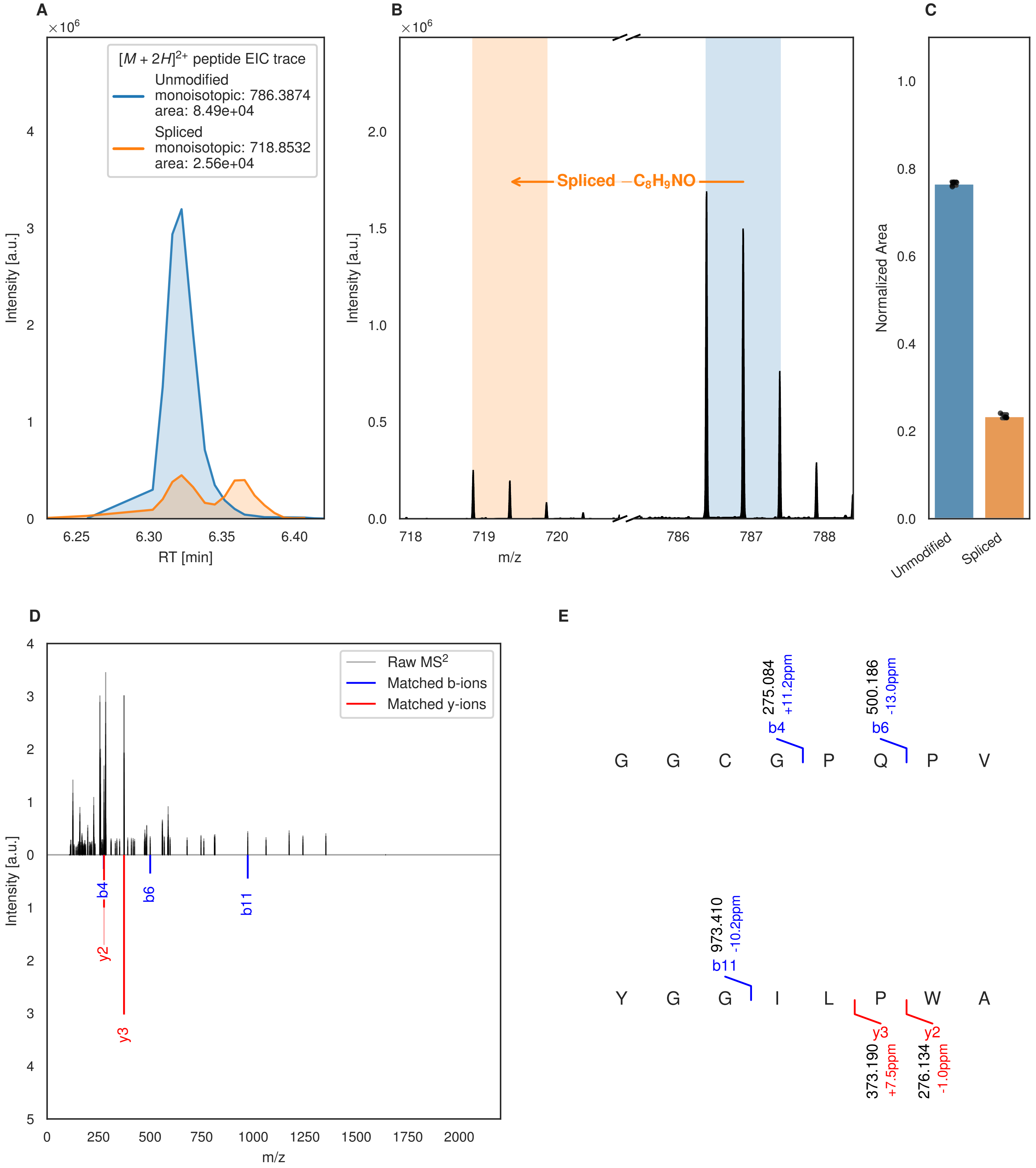
A
×10^6
[M + 2H]2+ peptide EIC trace
Unmodified
monoisotopic: 786.3874
area: 8.49e+04
Spliced
monoisotopic: 718.8532
area: 2.56e+04
Intensity [a.u.]
RT [min]
B
×10^6
Spliced −C8H9NO
Intensity [a.u.]
m/z
C
Normalized Area
Unmodified
Spliced
D
Raw MS2
Matched b-ions
Matched y-ions
Intensity [a.u.]
m/z
b4
b6
b11
y2
y3
E
275.084
+11.2ppm
b4
500.186
-13.0ppm
b6
G G C G P Q P V
973.410
-10.2ppm
b11
Y G G I L P W A
y3
373.190
+7.5ppm
y2
276.134
-1.0ppm

Figure S117 (figure continued from the previous page).

F

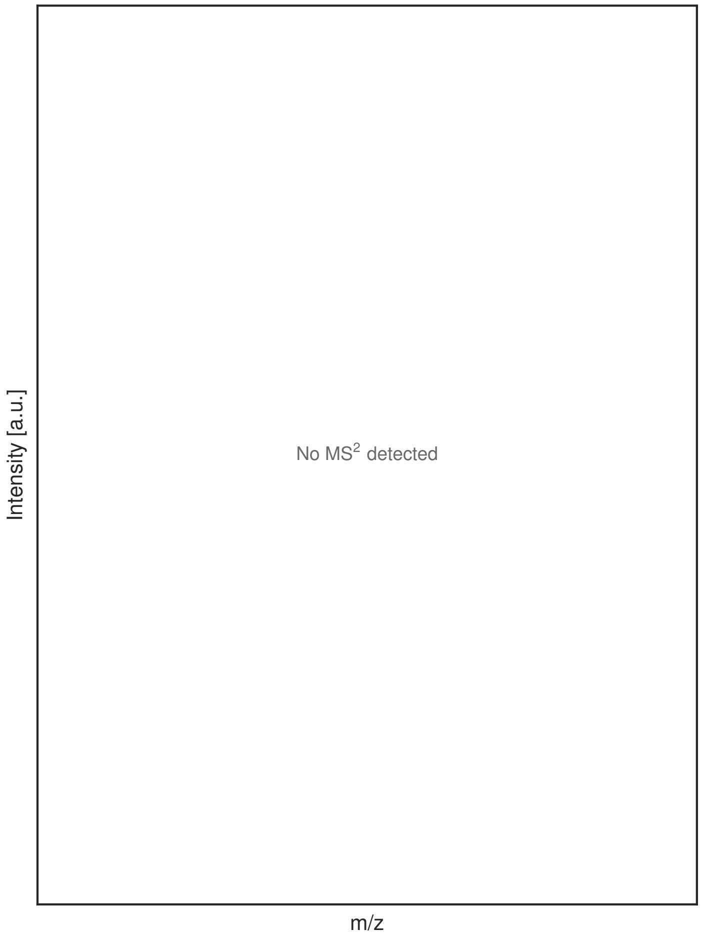


G

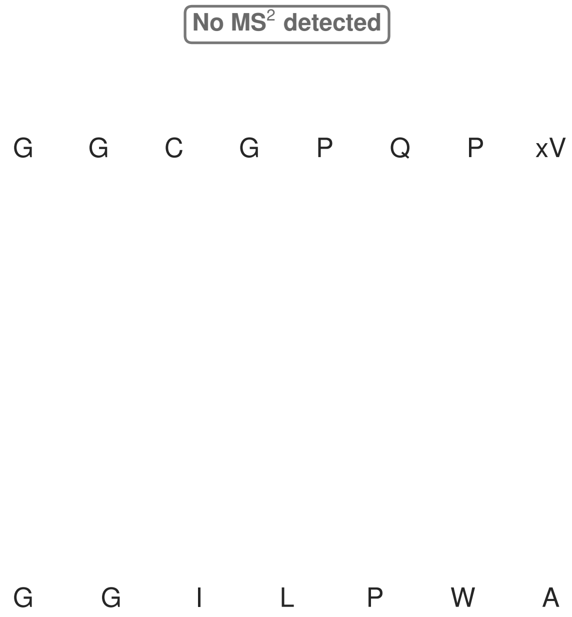


**Figure S117: Modification of the mutant_32 precursor peptide by HypXY.** LC-MS analysis of IMAC-purified precursor peptides from *E. coli* BL21(DE3)Tuner containing splicease-encoding plasmid TJ-C-689 (HypXY) and precursor peptide-encoding plasmid mutant_32 (RSTN1-HypAc-TEV-mutant-32) after expression at 24 °C for 22 h in 1.5 mL cultures after digestion with TEV. Data from n=4 independent biological replicates. Spliced amino acids have an x as a prefix. Spliced refers to a single tyramine excision. A representative experimental replicate is shown in panels **A, B.**

**A** Close-up plot of the representative extracted ion chromatogram (EIC) of the various peptide states; the blue trace corresponds to the Unmodified peptide; the orange trace corresponds to the Spliced peptide. **B** Close-up plot of the m/z values of the peptides detected by the mass spectrometer, with the shaded colored areas corresponding to where the expected m/z values for the peptide m/z traces should occur, with the observed spectrum plotted in black. The Unmodified, Spliced peptide(s) were found in the mass spectrum. **C** Bar plot showing the relative abundance of the various peptide states at charge z=2; the bars represent the average relative abundances normalized to total peak areas, with each individual experimental replicate value plotted as a black dot. **D** Mirrored $MS^2$ plot for the Unmodified peptide. In this mirror plot, the full fragmentation spectrum is plotted in black pointing upward, and the matched fragment b ions (blue) and y ions (red) are plotted pointing downward, mirrored against it, so that each matched fragment can be compared directly with the measured spectrum. **E** $MS^2$ spectrum of the fragments corresponding to the Unmodified peptide sequence, with matched b (blue, above) and y ions (red, below) plotted. **F** Mirrored $MS^2$ plot for the Spliced peptide. In this mirror plot, the full fragmentation spectrum is plotted in black pointing upward, and the matched fragment b ions (blue) and y ions (red) are plotted pointing downward, mirrored against it, so that each matched fragment can be compared directly with the measured spectrum. **G** $MS^2$ spectrum of the fragments corresponding to the Spliced peptide sequence, with matched b (blue, above) and y ions (red, below) plotted.

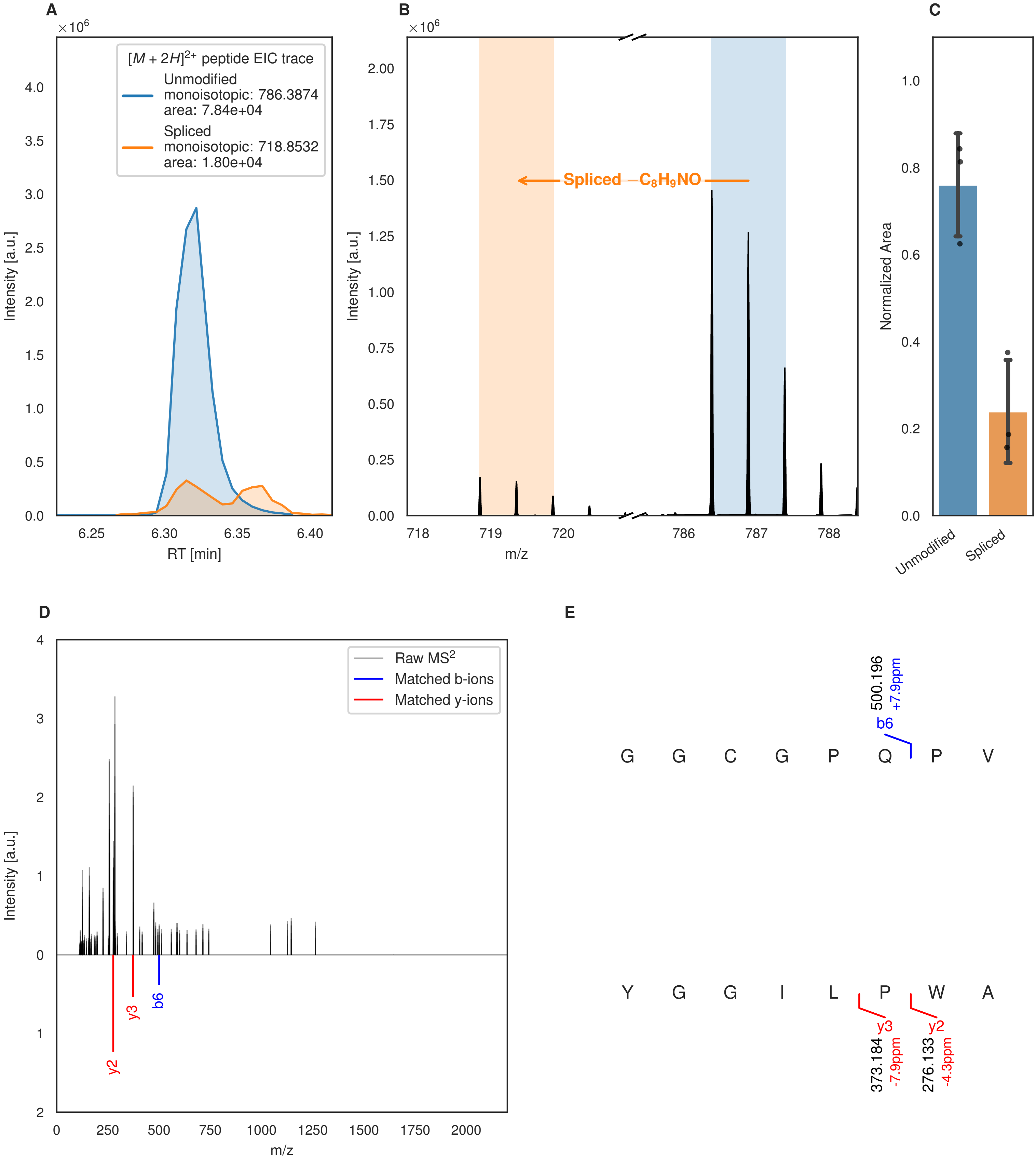
A
×10^6
[M + 2H]^2+ peptide EIC trace
Unmodified
monoisotopic: 786.3874
area: 7.84e+04
Spliced
monoisotopic: 718.8532
area: 1.80e+04
Intensity [a.u.]
RT [min]
B
Spliced −C8H9NO
m/z
C
Normalized Area
Unmodified
Spliced
D
Raw MS2
Matched b-ions
Matched y-ions
y2
y3
b6
E
500.196
+7.9ppm
b6
G G C G P Q P V
Y G G I L P W A
y3
y2
373.184
-7.9ppm
276.133
-4.3ppm

Figure S118 (figure continued from the previous page).

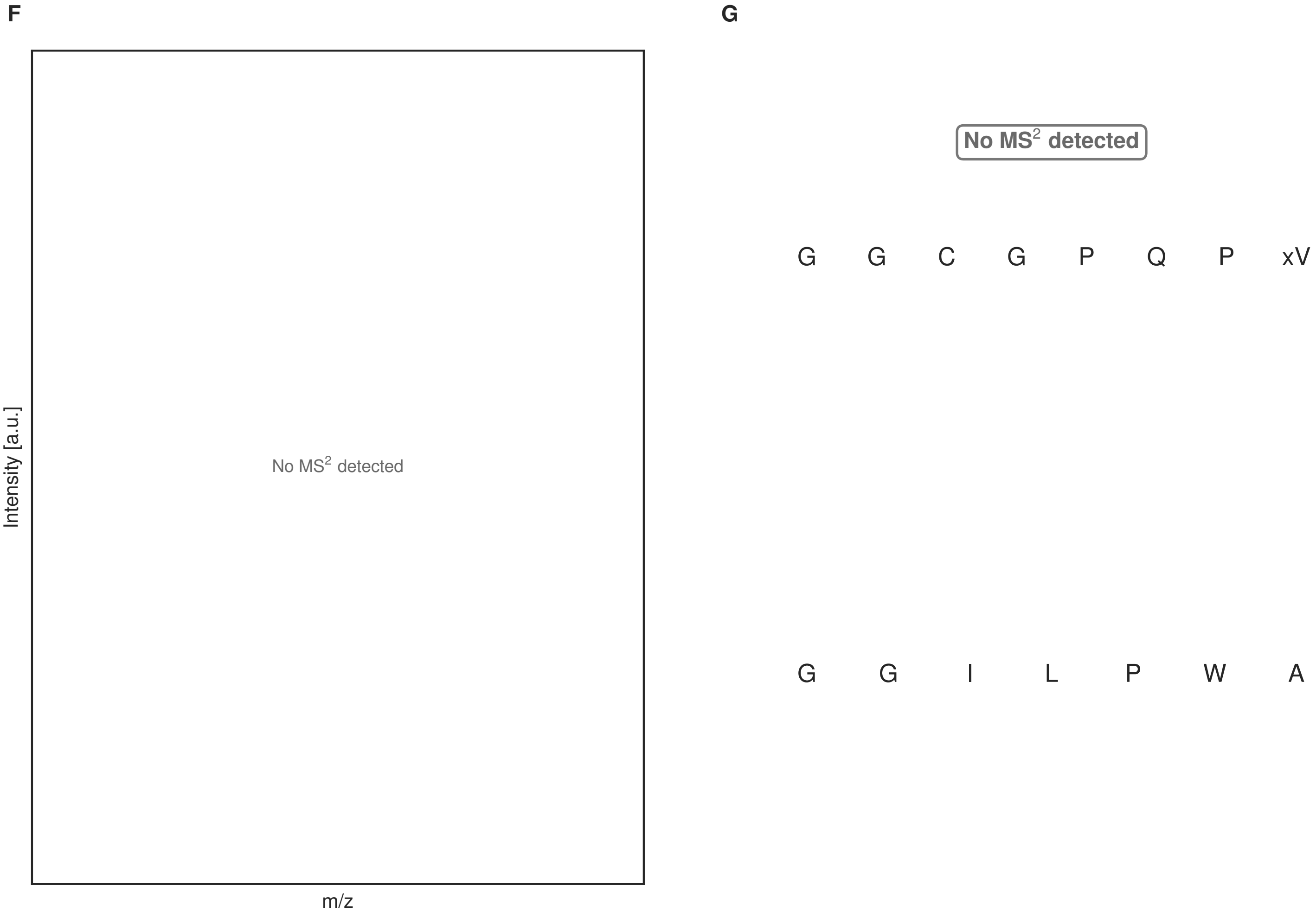


**Figure S118: Modification of the mutant_32 precursor peptide by PlpXY.** LC-MS analysis of IMAC-purified precursor peptides from *E. coli* BL21(DE3)Tuner containing splicease-encoding plasmid TJ-C-559 (PlpXY) and precursor peptide-encoding plasmid mutant_32 (RSTN1-PlpA3-TEV-mutant-32) after expression at 24 °C for 22 h in 1.5 mL cultures after digestion with TEV. Data from n=3 independent biological replicates. Spliced amino acids have an x as a prefix. Spliced refers to a single tyramine excision. A representative experimental replicate is shown in panels **A, B.**

**A** Close-up plot of the representative extracted ion chromatogram (EIC) of the various peptide states; the blue trace corresponds to the Unmodified peptide; the orange trace corresponds to the Spliced peptide. **B** Close-up plot of the m/z values of the peptides detected by the mass spectrometer, with the shaded colored areas corresponding to where the expected m/z values for the peptide m/z traces should occur, with the observed spectrum plotted in black. The Unmodified, Spliced peptide(s) were found in the mass spectrum. **C** Bar plot showing the relative abundance of the various peptide states at charge z=2; the bars represent the average relative abundances normalized to total peak areas, with each individual experimental replicate value plotted as a black dot. **D** Mirrored $MS^2$ plot for the Unmodified peptide. In this mirror plot, the full fragmentation spectrum is plotted in black pointing upward, and the matched fragment b ions (blue) and y ions (red) are plotted pointing downward, mirrored against it, so that each matched fragment can be compared directly with the measured spectrum. **E** $MS^2$ spectrum of the fragments corresponding to the Unmodified peptide sequence, with matched b (blue, above) and y ions (red, below) plotted. **F** Mirrored $MS^2$ plot for the Spliced peptide. In this mirror plot, the full fragmentation spectrum is plotted in black pointing upward, and the matched fragment b ions (blue) and y ions (red) are plotted pointing downward, mirrored against it, so that each matched fragment can be compared directly with the measured spectrum. **G** $MS^2$ spectrum of the fragments corresponding to the Spliced peptide sequence, with matched b (blue, above) and y ions (red, below) plotted.

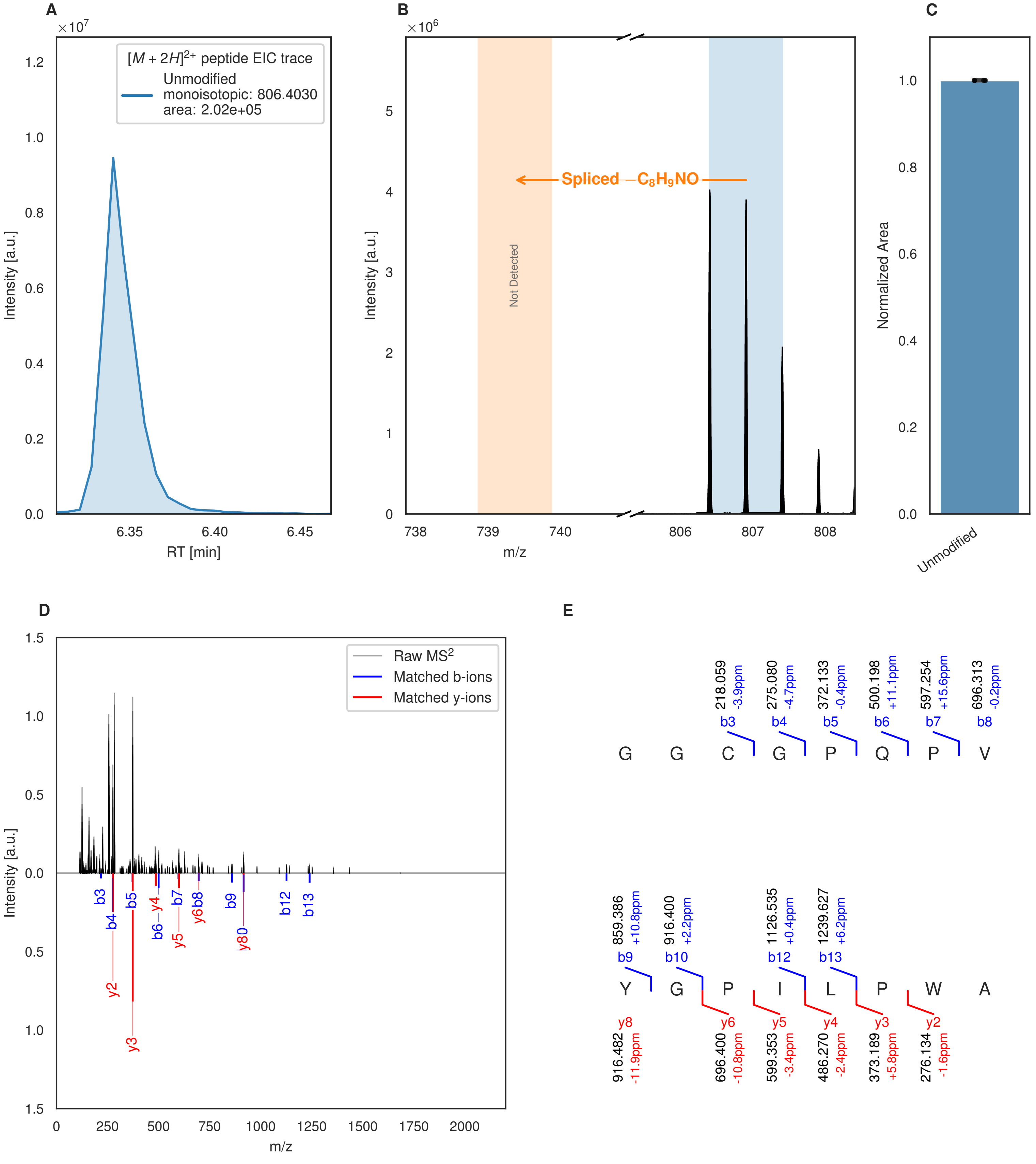


**Figure S119: Modification of the mutant_33 precursor peptide by HypXY.** LC-MS analysis of IMAC-purified precursor peptides from *E. coli* BL21(DE3)Tuner containing splicease-encoding plasmid TJ-C-689 (HypXY) and precursor peptide-encoding plasmid mutant_33 (RSTN1-HypAc-TEV-mutant-33) after expression at 24 °C for 22 h in 1.5 mL cultures after digestion with TEV. Data from n=4 independent biological replicates. Spliced amino acids have an x as a prefix. Spliced refers to a single tyramine excision. A representative experimental replicate is shown in panels **A, B.**

**A** Close-up plot of the representative extracted ion chromatogram (EIC) of the various peptide states; the blue trace corresponds to the Unmodified peptide. **B** Close-up plot of the m/z values of the peptides detected by the mass spectrometer, with the shaded colored areas corresponding to where the expected m/z values for the peptide m/z

(Caption continued on the next page.)

traces should occur, with the observed spectrum plotted in black. The Unmodified peptide(s) were found in the mass spectrum and the Spliced peptide(s) were not detected. **C** Bar plot showing the relative abundance of the various peptide states at charge z=2; the bars represent the average relative abundances normalized to total peak areas, with each individual experimental replicate value plotted as a black dot. **D** Mirrored $MS^2$ plot for the Unmodified peptide. In this mirror plot, the full fragmentation spectrum is plotted in black pointing upward, and the matched fragment b ions (blue) and y ions (red) are plotted pointing downward, mirrored against it, so that each matched fragment can be compared directly with the measured spectrum. **E** $MS^2$ spectrum of the fragments corresponding to the Unmodified peptide sequence, with matched b (blue, above) and y ions (red, below) plotted.

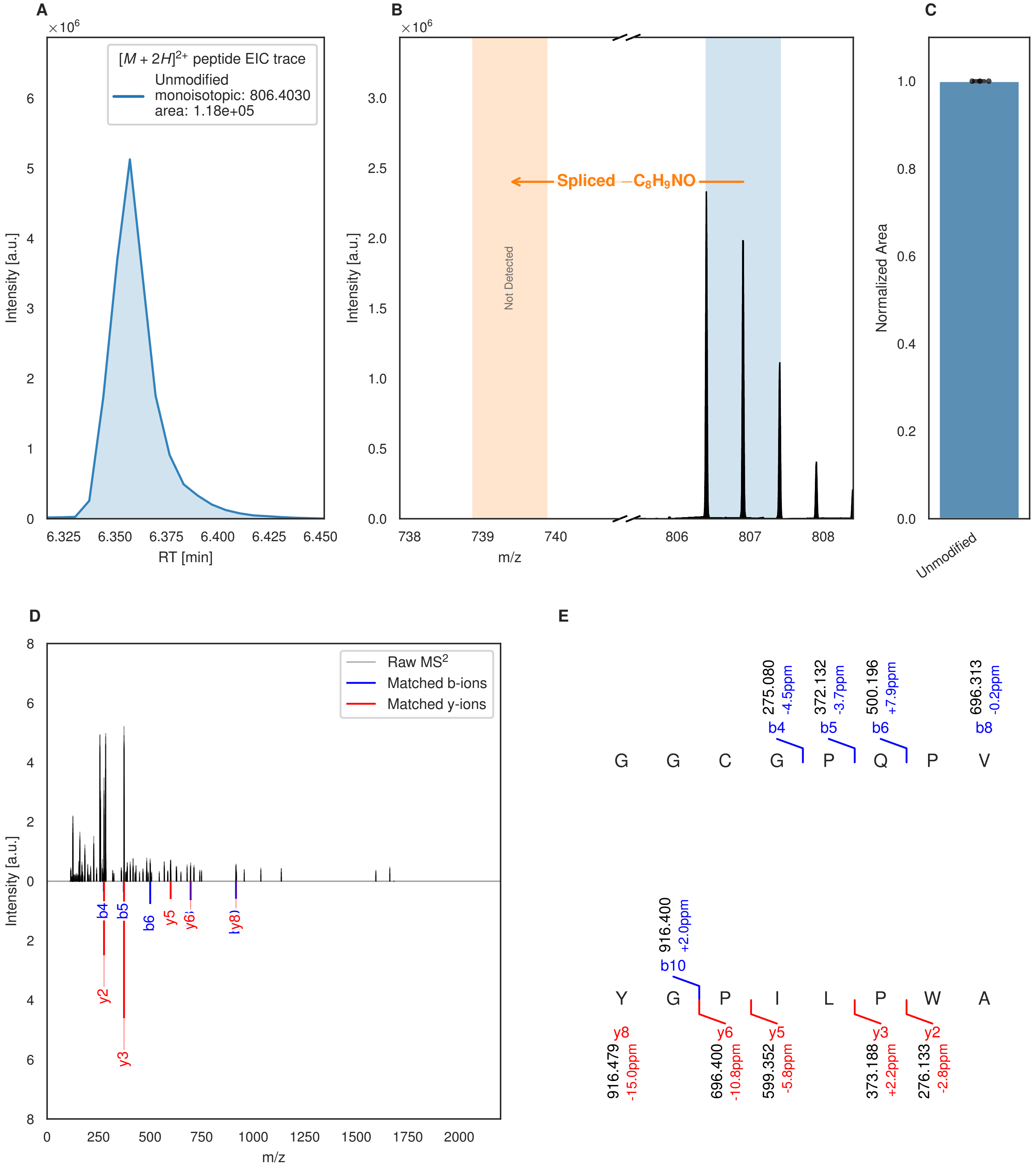


**Figure S120: Modification of the mutant_33 precursor peptide by PlpXY.** LC-MS analysis of IMAC-purified precursor peptides from *E. coli* BL21(DE3)Tuner containing splicease-encoding plasmid TJ-C-559 (PlpXY) and precursor peptide-encoding plasmid mutant_33 (RSTN1-PlpA3-TEV-mutant-33) after expression at 24 °C for 22 h in 1.5 mL cultures after digestion with TEV. Data from n=3 independent biological replicates. Spliced amino acids have an x as a prefix. Spliced refers to a single tyramine excision. A representative experimental replicate is shown in panels **A, B.**

**A** Close-up plot of the representative extracted ion chromatogram (EIC) of the various peptide states; the blue trace corresponds to the Unmodified peptide. **B** Close-up plot of the m/z values of the peptides detected by the mass spectrometer, with the shaded colored areas corresponding to where the expected m/z values for the peptide m/z

(Caption continued on the next page.)

traces should occur, with the observed spectrum plotted in black. The Unmodified peptide(s) were found in the mass spectrum and the Spliced peptide(s) were not detected. **C** Bar plot showing the relative abundance of the various peptide states at charge z=2; the bars represent the average relative abundances normalized to total peak areas, with each individual experimental replicate value plotted as a black dot. **D** Mirrored $MS^2$ plot for the Unmodified peptide. In this mirror plot, the full fragmentation spectrum is plotted in black pointing upward, and the matched fragment b ions (blue) and y ions (red) are plotted pointing downward, mirrored against it, so that each matched fragment can be compared directly with the measured spectrum. **E** $MS^2$ spectrum of the fragments corresponding to the Unmodified peptide sequence, with matched b (blue, above) and y ions (red, below) plotted.

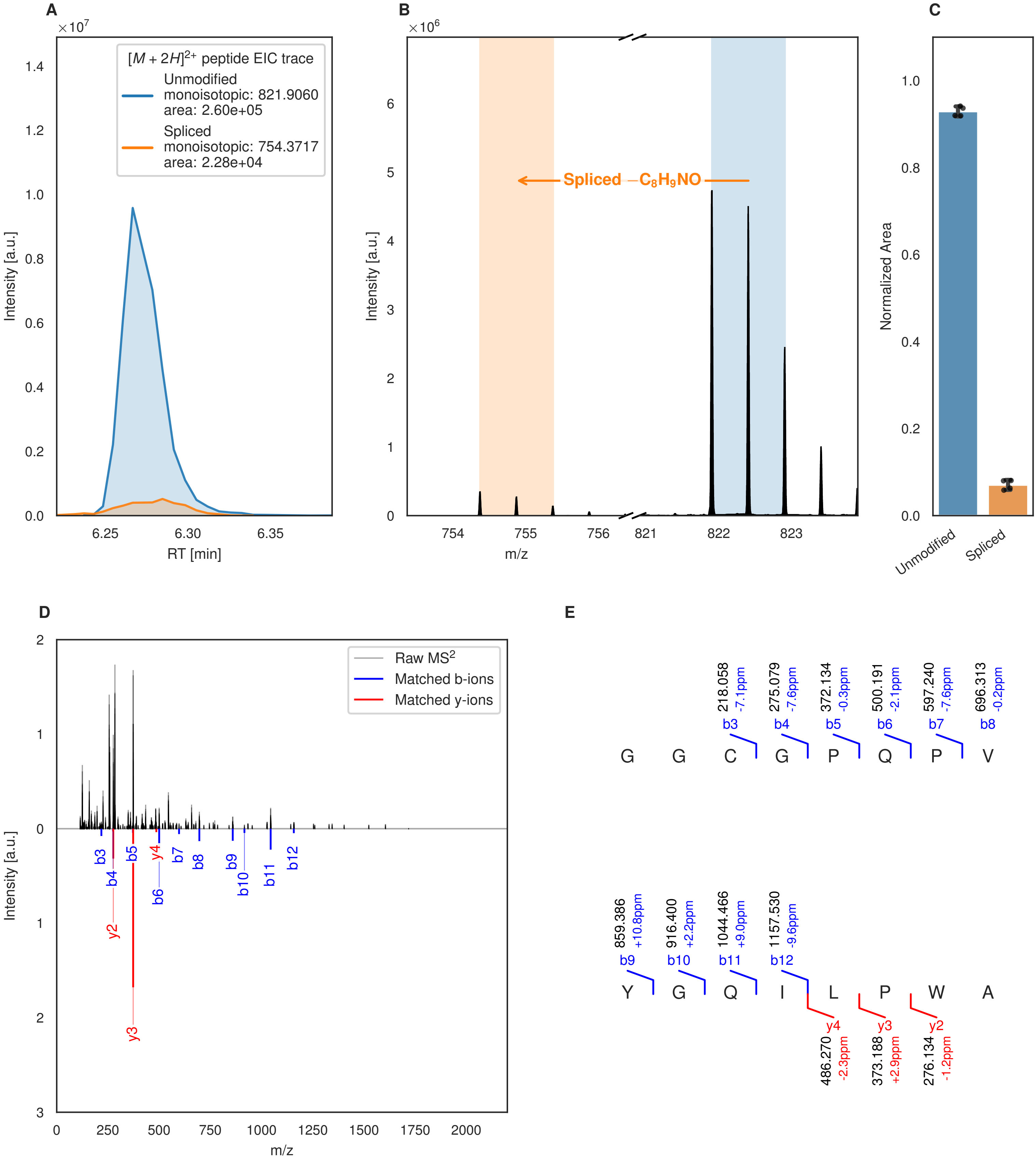
A
×10^7
[M + 2H]2+ peptide EIC trace
Unmodified
monoisotopic: 821.9060
area: 2.60e+05
Spliced
monoisotopic: 754.3717
area: 2.28e+04
Intensity [a.u.]
RT [min]
B
×10^6
Spliced −C8H9NO
Intensity [a.u.]
m/z
C
Normalized Area
Unmodified
Spliced
D
Raw MS2
Matched b-ions
Matched y-ions
Intensity [a.u.]
m/z
b3
b4
b5
y4
b6
b7
b8
b9
b10
b11
b12
y2
y3
E
218.058 -7.1ppm b3
275.079 -7.6ppm b4
372.134 -0.3ppm b5
500.191 -2.1ppm b6
597.240 -7.6ppm b7
696.313 -0.2ppm b8
G G C G P Q P V
859.386 +10.8ppm b9
916.400 +2.2ppm b10
1044.466 +9.0ppm b11
1157.530 -9.6ppm b12
Y G Q I L P W A
y4 486.270 -2.3ppm
y3 373.188 +2.9ppm
y2 276.134 -1.2ppm

Figure S121 (figure continued from the previous page).

F G

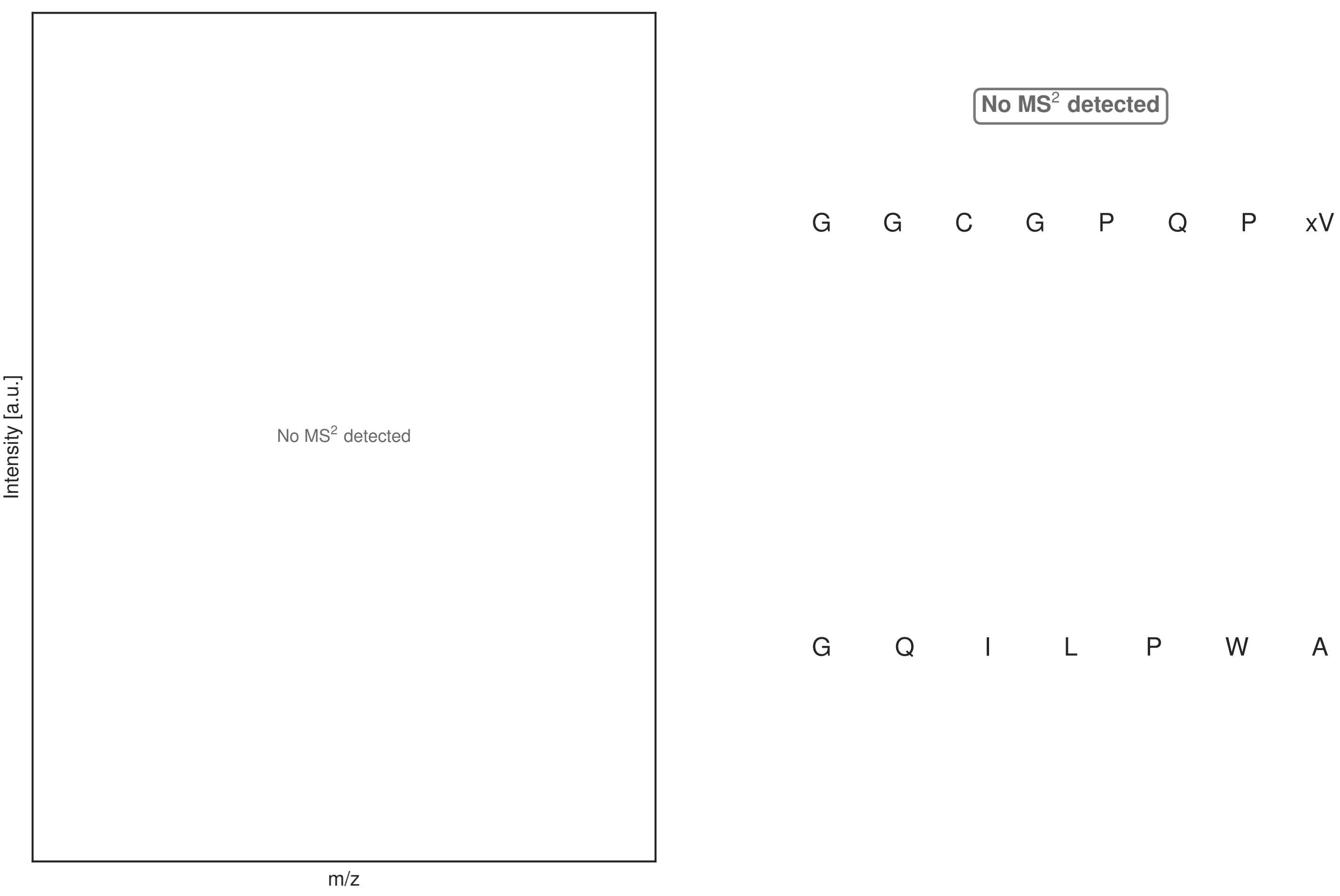


**Figure S121: Modification of the mutant_34 precursor peptide by HypXY.** LC-MS analysis of IMAC-purified precursor peptides from *E. coli* BL21(DE3)Tuner containing splicease-encoding plasmid TJ-C-689 (HypXY) and precursor peptide-encoding plasmid mutant_34 (RSTN1-HypAc-TEV-mutant-34) after expression at 24 °C for 22 h in 1.5 mL cultures after digestion with TEV. Data from n=4 independent biological replicates. Spliced amino acids have an x as a prefix. Spliced refers to a single tyramine excision. A representative experimental replicate is shown in panels **A, B.**

**A** Close-up plot of the representative extracted ion chromatogram (EIC) of the various peptide states; the blue trace corresponds to the Unmodified peptide; the orange trace corresponds to the Spliced peptide. **B** Close-up plot of the m/z values of the peptides detected by the mass spectrometer, with the shaded colored areas corresponding to where the expected m/z values for the peptide m/z traces should occur, with the observed spectrum plotted in black. The Unmodified, Spliced peptide(s) were found in the mass spectrum. **C** Bar plot showing the relative abundance of the various peptide states at charge z=2; the bars represent the average relative abundances normalized to total peak areas, with each individual experimental replicate value plotted as a black dot. **D** Mirrored MS$^2$ plot for the Unmodified peptide. In this mirror plot, the full fragmentation spectrum is plotted in black pointing upward, and the matched fragment b ions (blue) and y ions (red) are plotted pointing downward, mirrored against it, so that each matched fragment can be compared directly with the measured spectrum. **E** MS$^2$ spectrum of the fragments corresponding to the Unmodified peptide sequence, with matched b (blue, above) and y ions (red, below) plotted. **F** Mirrored MS$^2$ plot for the Spliced peptide. In this mirror plot, the full fragmentation spectrum is plotted in black pointing upward, and the matched fragment b ions (blue) and y ions (red) are plotted pointing downward, mirrored against it, so that each matched fragment can be compared directly with the measured spectrum. **G** MS$^2$ spectrum of the fragments corresponding to the Spliced peptide sequence, with matched b (blue, above) and y ions (red, below) plotted.

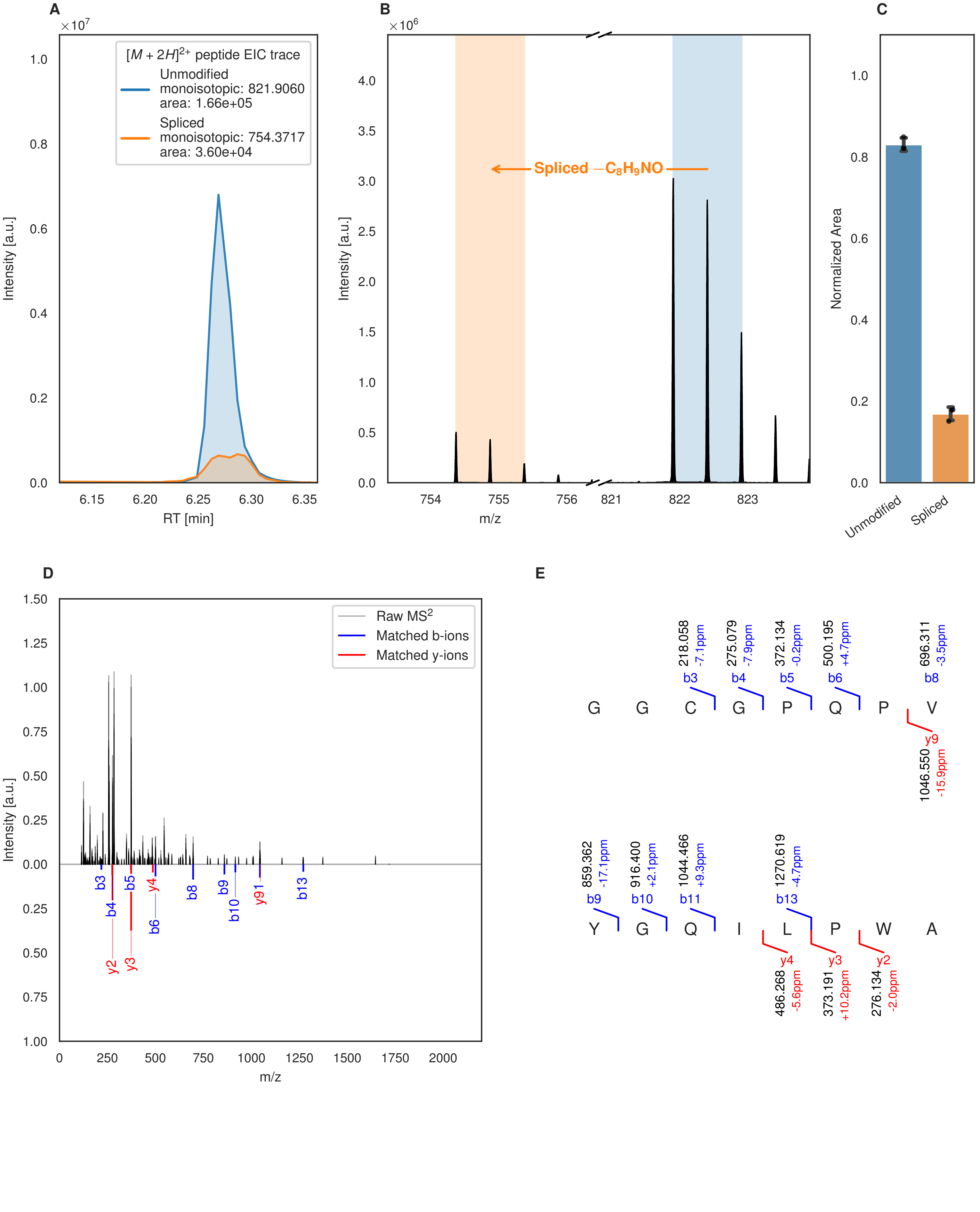
A
[M + 2H]2+ peptide EIC trace
Unmodified
monoisotopic: 821.9060
area: 1.66e+05
Spliced
monoisotopic: 754.3717
area: 3.60e+04
Intensity [a.u.]
RT [min]
B
Spliced −C8H9NO
Intensity [a.u.]
m/z
C
Normalized Area
Unmodified
Spliced
D
Raw MS2
Matched b-ions
Matched y-ions
Intensity [a.u.]
m/z
E
218.058 -7.1ppm b3
275.079 -7.9ppm b4
372.134 -0.2ppm b5
500.195 +4.7ppm b6
696.311 -3.5ppm b8
G G C G P Q P V
y9 1046.550 -15.9ppm
859.362 -17.1ppm b9
916.400 +2.1ppm b10
1044.466 +9.3ppm b11
1270.619 -4.7ppm b13
Y G Q I L P W A
y4 486.268 -5.6ppm
y3 373.191 +10.2ppm
y2 276.134 -2.0ppm

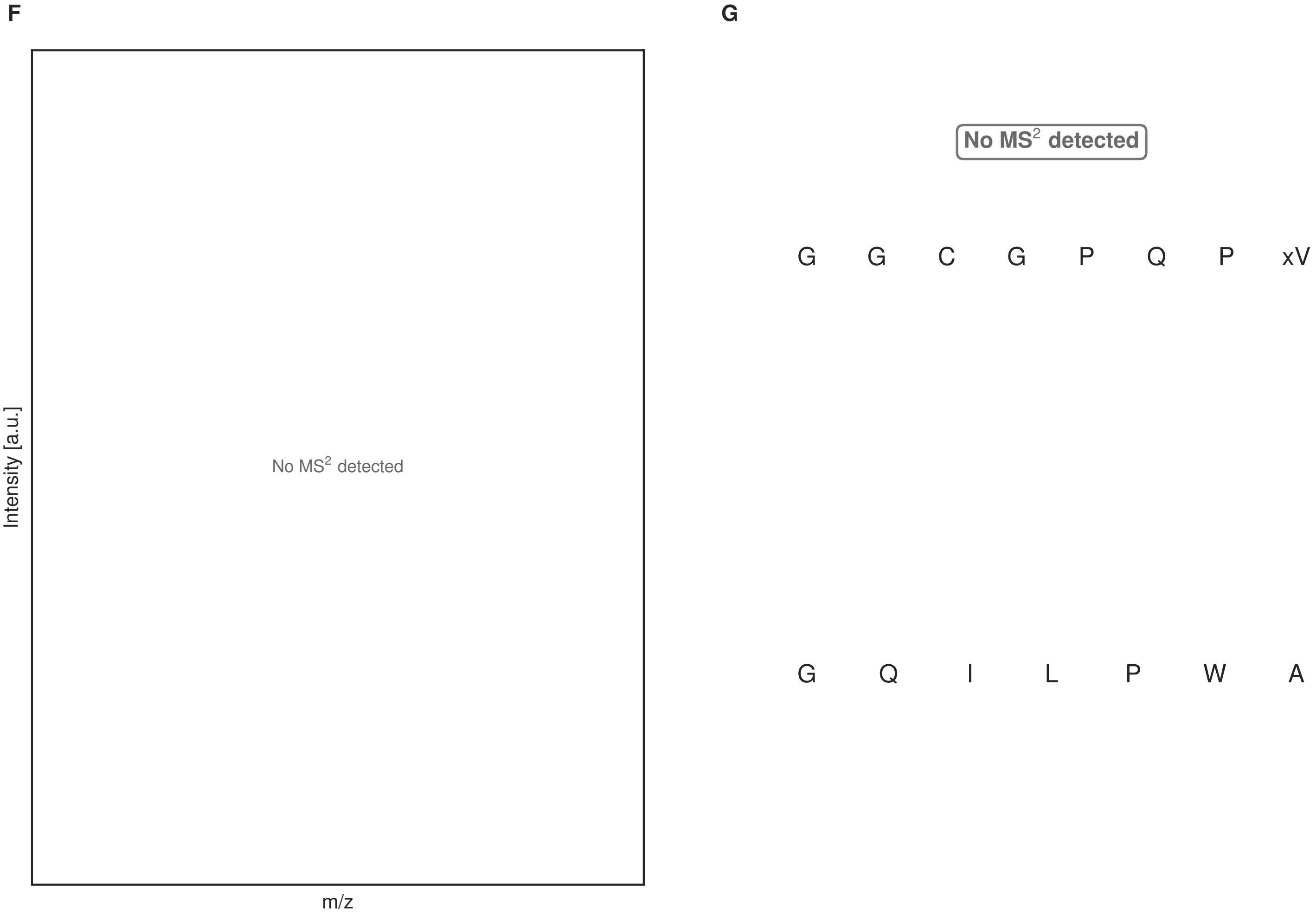


**Figure S122: Modification of the mutant_34 precursor peptide by PlpXY.** LC-MS analysis of IMAC-purified precursor peptides from *E. coli* BL21(DE3)Tuner containing splicease-encoding plasmid TJ-C-559 (PlpXY) and precursor peptide-encoding plasmid mutant_34 (RSTN1-PlpA3-TEV-mutant-34) after expression at 24 °C for 22 h in 1.5 mL cultures after digestion with TEV. Data from n=3 independent biological replicates. Spliced amino acids have an x as a prefix. Spliced refers to a single tyramine excision. A representative experimental replicate is shown in panels **A, B.**

**A** Close-up plot of the representative extracted ion chromatogram (EIC) of the various peptide states; the blue trace corresponds to the Unmodified peptide; the orange trace corresponds to the Spliced peptide. **B** Close-up plot of the m/z values of the peptides detected by the mass spectrometer, with the shaded colored areas corresponding to where the expected m/z values for the peptide m/z traces should occur, with the observed spectrum plotted in black. The Unmodified, Spliced peptide(s) were found in the mass spectrum. **C** Bar plot showing the relative abundance of the various peptide states at charge z=2; the bars represent the average relative abundances normalized to total peak areas, with each individual experimental replicate value plotted as a black dot. **D** Mirrored MS$^2$ plot for the Unmodified peptide. In this mirror plot, the full fragmentation spectrum is plotted in black pointing upward, and the matched fragment b ions (blue) and y ions (red) are plotted pointing downward, mirrored against it, so that each matched fragment can be compared directly with the measured spectrum. **E** MS$^2$ spectrum of the fragments corresponding to the Unmodified peptide sequence, with matched b (blue, above) and y ions (red, below) plotted. **F** Mirrored MS$^2$ plot for the Spliced peptide. In this mirror plot, the full fragmentation spectrum is plotted in black pointing upward, and the matched fragment b ions (blue) and y ions (red) are plotted pointing downward, mirrored against it, so that each matched fragment can be compared directly with the measured spectrum. **G** MS$^2$ spectrum of the fragments corresponding to the Spliced peptide sequence, with matched b (blue, above) and y ions (red, below) plotted.

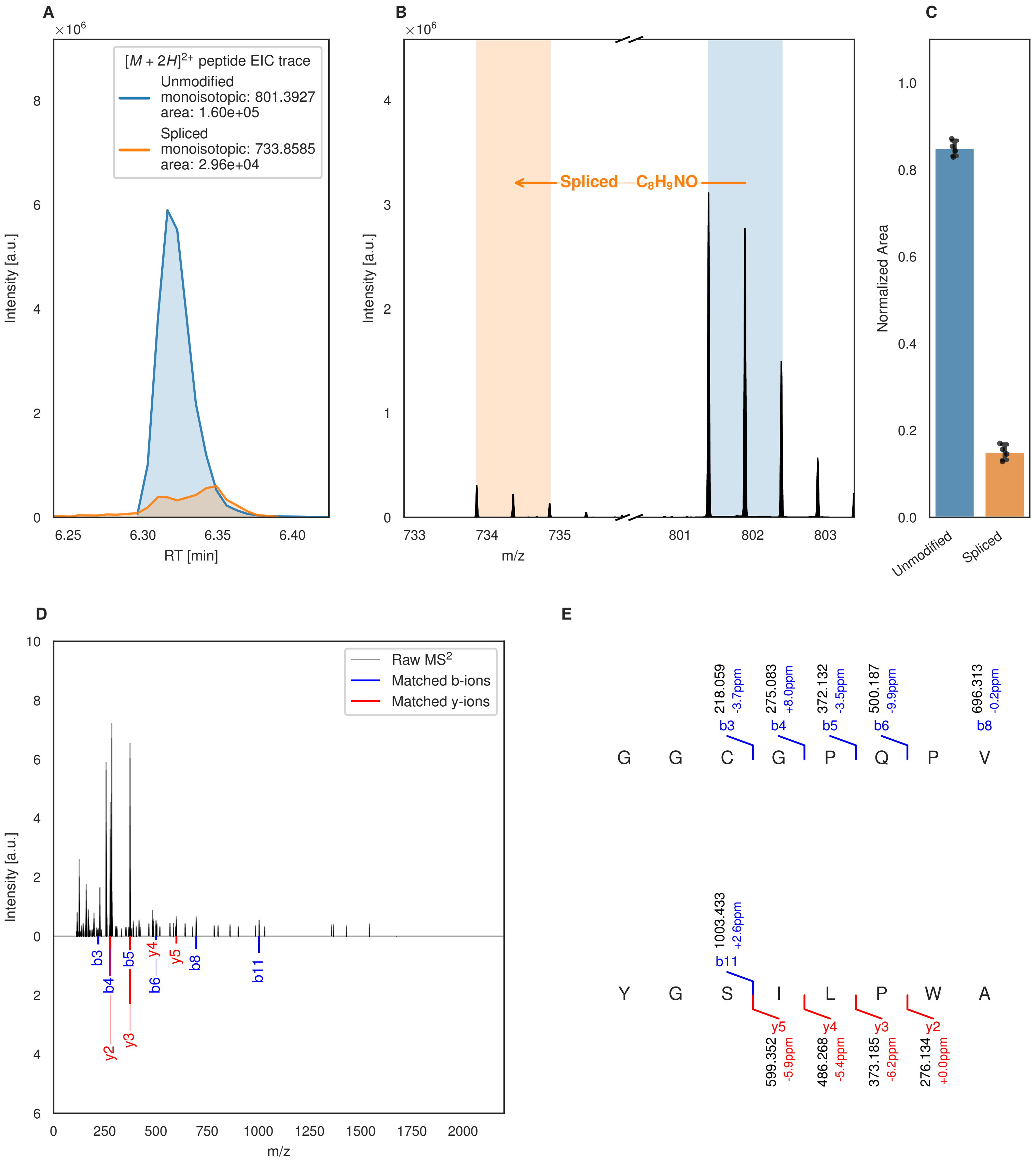
A
B
C
D
E
[M + 2H]2+ peptide EIC trace
Unmodified
monoisotopic: 801.3927
area: 1.60e+05
Spliced
monoisotopic: 733.8585
area: 2.96e+04
Intensity [a.u.]
RT [min]
Spliced −C8H9NO
m/z
Normalized Area
Unmodified
Spliced
Raw MS2
Matched b-ions
Matched y-ions
G G C G P Q P V
Y G S I L P W A
218.059 -3.7ppm b3
275.083 +8.0ppm b4
372.132 -3.5ppm b5
500.187 -9.9ppm b6
696.313 -0.2ppm b8
1003.433 +2.6ppm b11
599.352 -5.9ppm y5
486.268 -5.4ppm y4
373.185 -6.2ppm y3
276.134 +0.0ppm y2

Figure S123 (figure continued from the previous page).

F

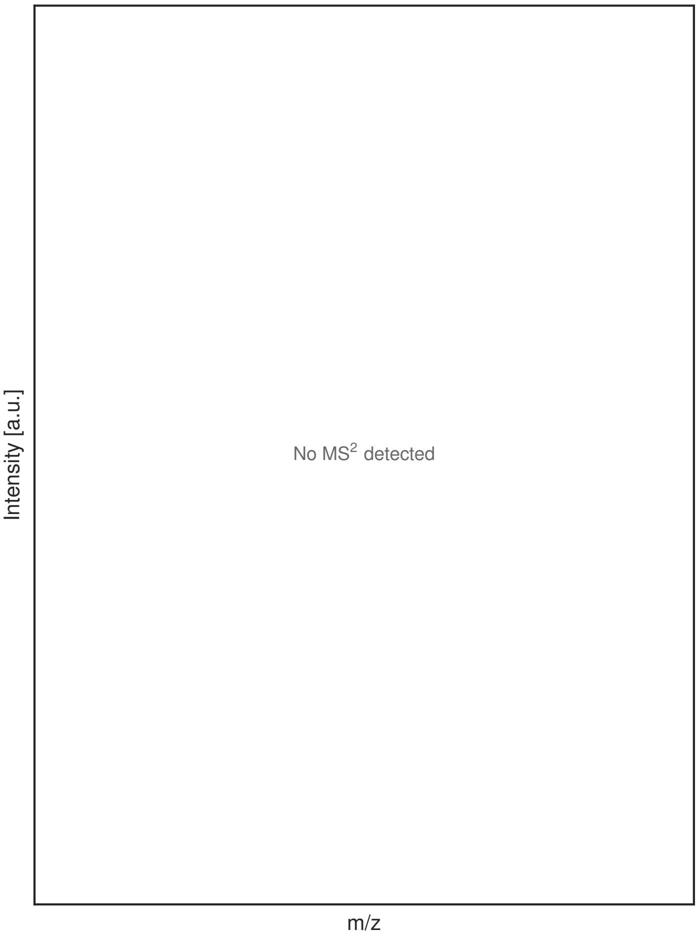


G

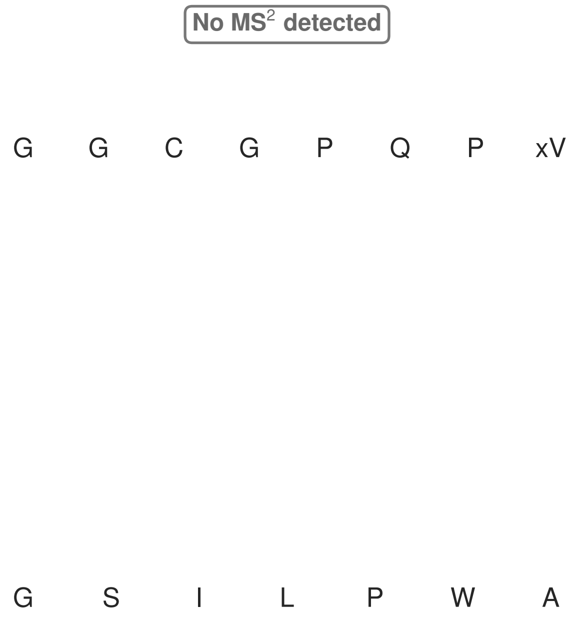


**Figure S123: Modification of the mutant_35 precursor peptide by HypXY.** LC-MS analysis of IMAC-purified precursor peptides from *E. coli* BL21(DE3)Tuner containing splicease-encoding plasmid TJ-C-689 (HypXY) and precursor peptide-encoding plasmid mutant_35 (RSTN1-HypAc-TEV-mutant-35) after expression at 24 °C for 22 h in 1.5 mL cultures after digestion with TEV. Data from n=4 independent biological replicates. Spliced amino acids have an x as a prefix. Spliced refers to a single tyramine excision. A representative experimental replicate is shown in panels **A, B.**

**A** Close-up plot of the representative extracted ion chromatogram (EIC) of the various peptide states; the blue trace corresponds to the Unmodified peptide; the orange trace corresponds to the Spliced peptide. **B** Close-up plot of the m/z values of the peptides detected by the mass spectrometer, with the shaded colored areas corresponding to where the expected m/z values for the peptide m/z traces should occur, with the observed spectrum plotted in black. The Unmodified, Spliced peptide(s) were found in the mass spectrum. **C** Bar plot showing the relative abundance of the various peptide states at charge z=2; the bars represent the average relative abundances normalized to total peak areas, with each individual experimental replicate value plotted as a black dot. **D** Mirrored $MS^2$ plot for the Unmodified peptide. In this mirror plot, the full fragmentation spectrum is plotted in black pointing upward, and the matched fragment b ions (blue) and y ions (red) are plotted pointing downward, mirrored against it, so that each matched fragment can be compared directly with the measured spectrum. **E** $MS^2$ spectrum of the fragments corresponding to the Unmodified peptide sequence, with matched b (blue, above) and y ions (red, below) plotted. **F** Mirrored $MS^2$ plot for the Spliced peptide. In this mirror plot, the full fragmentation spectrum is plotted in black pointing upward, and the matched fragment b ions (blue) and y ions (red) are plotted pointing downward, mirrored against it, so that each matched fragment can be compared directly with the measured spectrum. **G** $MS^2$ spectrum of the fragments corresponding to the Spliced peptide sequence, with matched b (blue, above) and y ions (red, below) plotted.

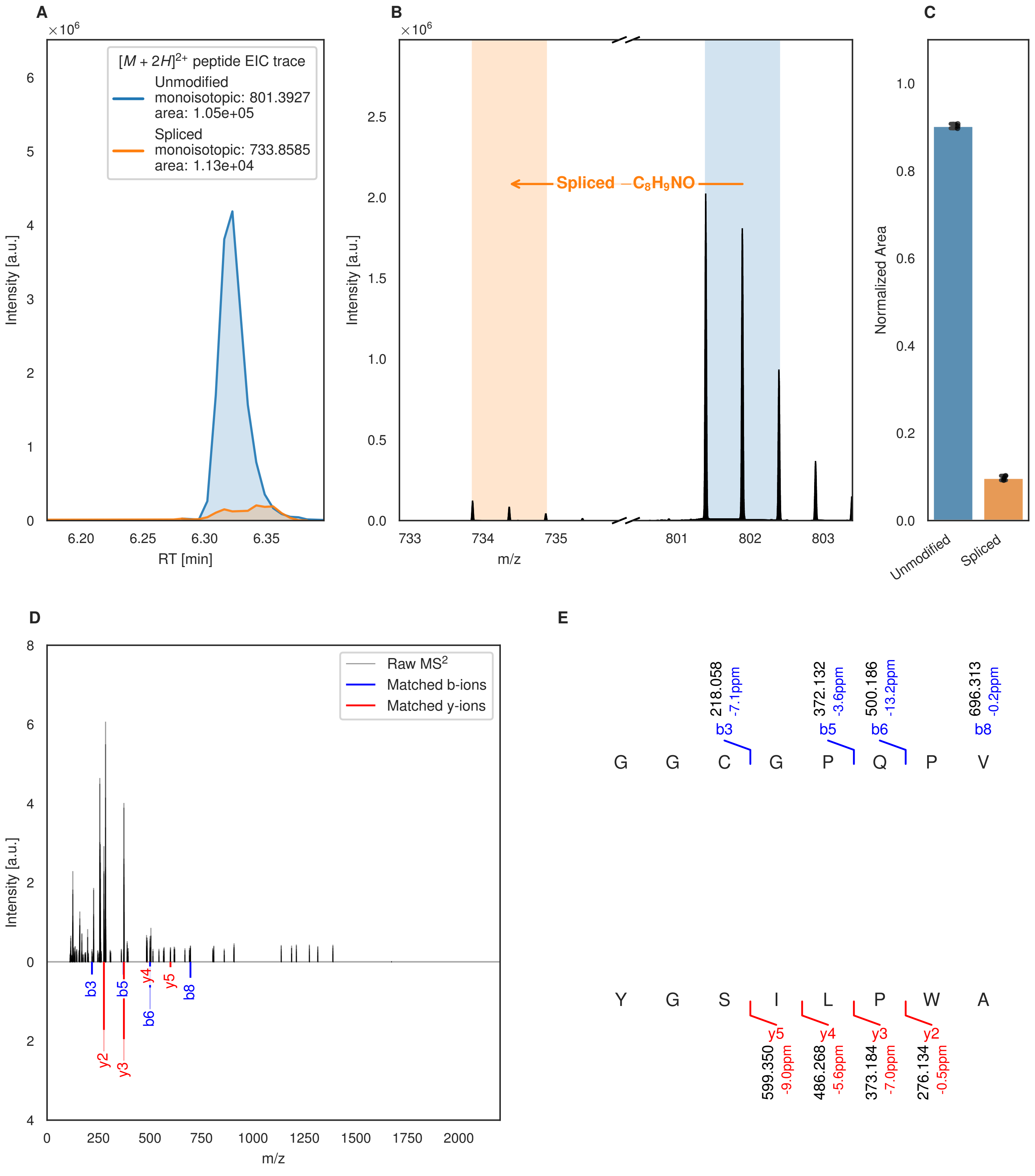
A
$\times 10^6$
$[M+2H]^{2+}$ peptide EIC trace
Unmodified
monoisotopic: 801.3927
area: 1.05e+05
Spliced
monoisotopic: 733.8585
area: 1.13e+04
Intensity [a.u.]
RT [min]
B
Spliced $-C_8H_9NO$
m/z
C
Normalized Area
Unmodified
Spliced
D
Raw $MS^2$
Matched b-ions
Matched y-ions
b3
b5
b6
b8
y2
y3
y4
y5
E
218.058 -7.1ppm b3
372.132 -3.6ppm b5
500.186 -13.2ppm b6
696.313 -0.2ppm b8
G G C G P Q P V
Y G S I L P W A
y5 599.350 -9.0ppm
y4 486.268 -5.6ppm
y3 373.184 -7.0ppm
y2 276.134 -0.5ppm

Figure S124 (figure continued from the previous page).

F

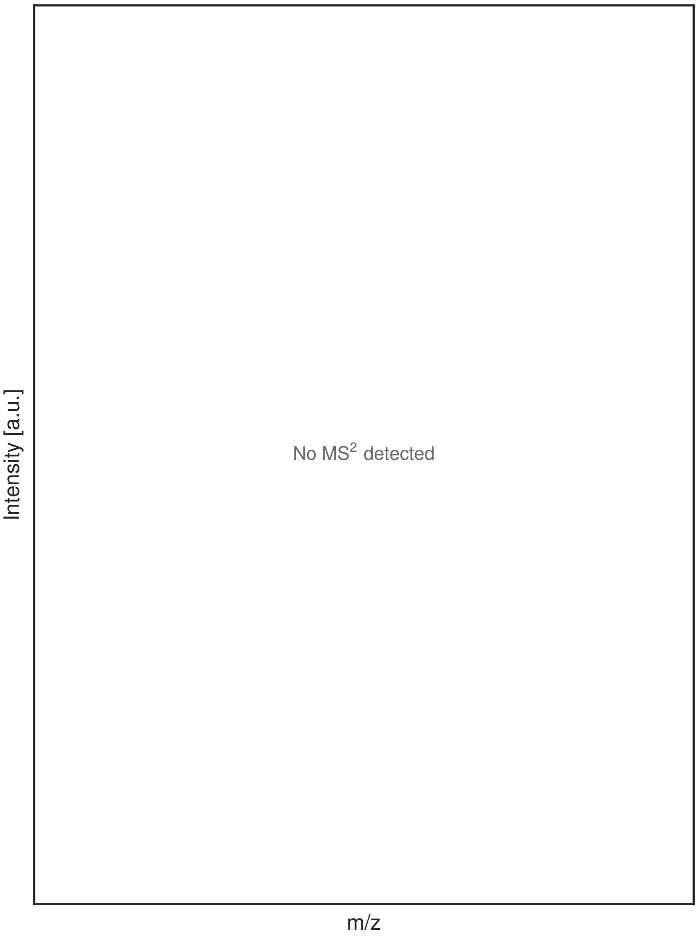


G

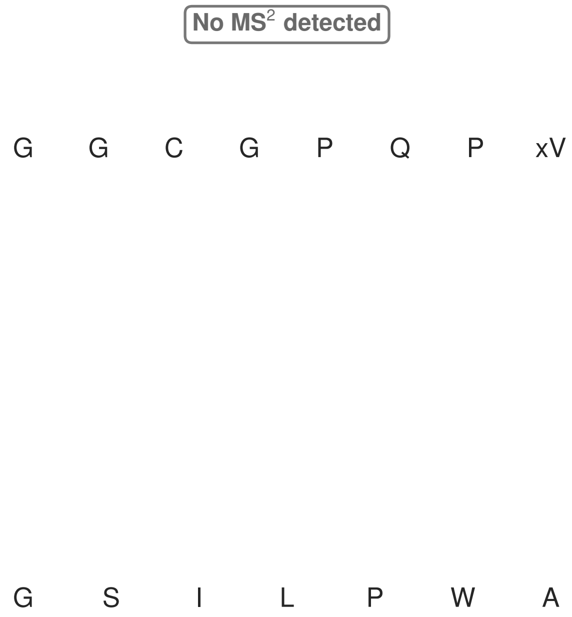


**Figure S124: Modification of the mutant_35 precursor peptide by PlpXY.** LC-MS analysis of IMAC-purified precursor peptides from *E. coli* BL21(DE3)Tuner containing splicease-encoding plasmid TJ-C-559 (PlpXY) and precursor peptide-encoding plasmid mutant_35 (RSTN1-PlpA3-TEV-mutant-35) after expression at 24 °C for 22 h in 1.5 mL cultures after digestion with TEV. Data from n=3 independent biological replicates. Spliced amino acids have an x as a prefix. Spliced refers to a single tyramine excision. A representative experimental replicate is shown in panels **A, B.**

**A** Close-up plot of the representative extracted ion chromatogram (EIC) of the various peptide states; the blue trace corresponds to the Unmodified peptide; the orange trace corresponds to the Spliced peptide. **B** Close-up plot of the m/z values of the peptides detected by the mass spectrometer, with the shaded colored areas corresponding to where the expected m/z values for the peptide m/z traces should occur, with the observed spectrum plotted in black. The Unmodified, Spliced peptide(s) were found in the mass spectrum. **C** Bar plot showing the relative abundance of the various peptide states at charge z=2; the bars represent the average relative abundances normalized to total peak areas, with each individual experimental replicate value plotted as a black dot. **D** Mirrored MS$^2$ plot for the Unmodified peptide. In this mirror plot, the full fragmentation spectrum is plotted in black pointing upward, and the matched fragment b ions (blue) and y ions (red) are plotted pointing downward, mirrored against it, so that each matched fragment can be compared directly with the measured spectrum. **E** MS$^2$ spectrum of the fragments corresponding to the Unmodified peptide sequence, with matched b (blue, above) and y ions (red, below) plotted. **F** Mirrored MS$^2$ plot for the Spliced peptide. In this mirror plot, the full fragmentation spectrum is plotted in black pointing upward, and the matched fragment b ions (blue) and y ions (red) are plotted pointing downward, mirrored against it, so that each matched fragment can be compared directly with the measured spectrum. **G** MS$^2$ spectrum of the fragments corresponding to the Spliced peptide sequence, with matched b (blue, above) and y ions (red, below) plotted.

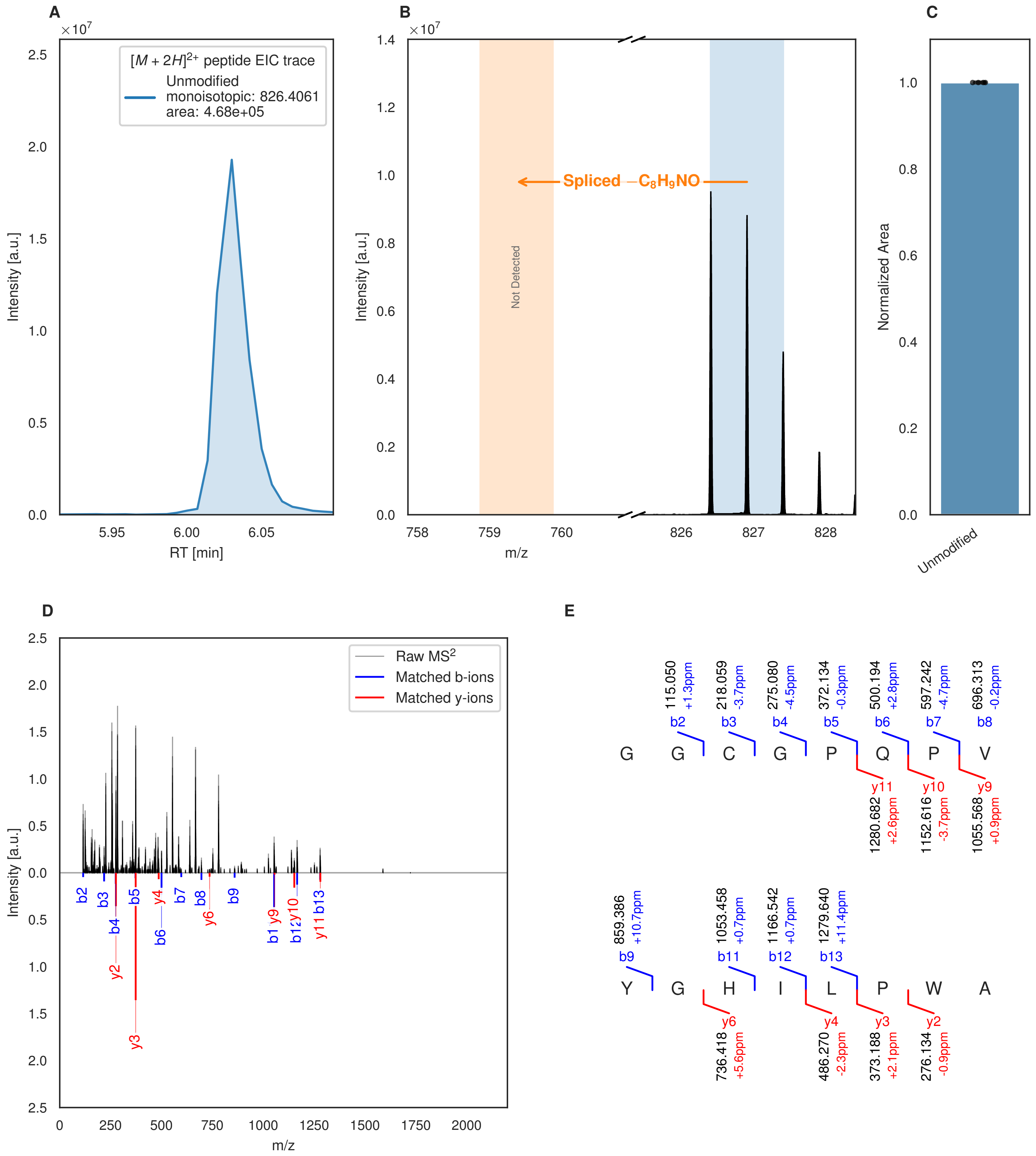


**Figure S125: Modification of the mutant_36 precursor peptide by HypXY.** LC-MS analysis of IMAC-purified precursor peptides from *E. coli* BL21(DE3)Tuner containing splicease-encoding plasmid TJ-C-689 (HypXY) and precursor peptide-encoding plasmid mutant_36 (RSTN1-HypAc-TEV-mutant-36) after expression at 24 °C for 22 h in 1.5 mL cultures after digestion with TEV. Data from n=4 independent biological replicates. Spliced amino acids have an x as a prefix. Spliced refers to a single tyramine excision. A representative experimental replicate is shown in panels **A, B.**

**A** Close-up plot of the representative extracted ion chromatogram (EIC) of the various peptide states; the blue trace corresponds to the Unmodified peptide. **B** Close-up plot of the m/z values of the peptides detected by the mass spectrometer, with the shaded colored areas corresponding to where the expected m/z values for the peptide m/z

(Caption continued on the next page.)

traces should occur, with the observed spectrum plotted in black. The Unmodified peptide(s) were found in the mass spectrum and the Spliced peptide(s) were not detected. **C** Bar plot showing the relative abundance of the various peptide states at charge z=2; the bars represent the average relative abundances normalized to total peak areas, with each individual experimental replicate value plotted as a black dot. **D** Mirrored $MS^2$ plot for the Unmodified peptide. In this mirror plot, the full fragmentation spectrum is plotted in black pointing upward, and the matched fragment b ions (blue) and y ions (red) are plotted pointing downward, mirrored against it, so that each matched fragment can be compared directly with the measured spectrum. **E** $MS^2$ spectrum of the fragments corresponding to the Unmodified peptide sequence, with matched b (blue, above) and y ions (red, below) plotted.

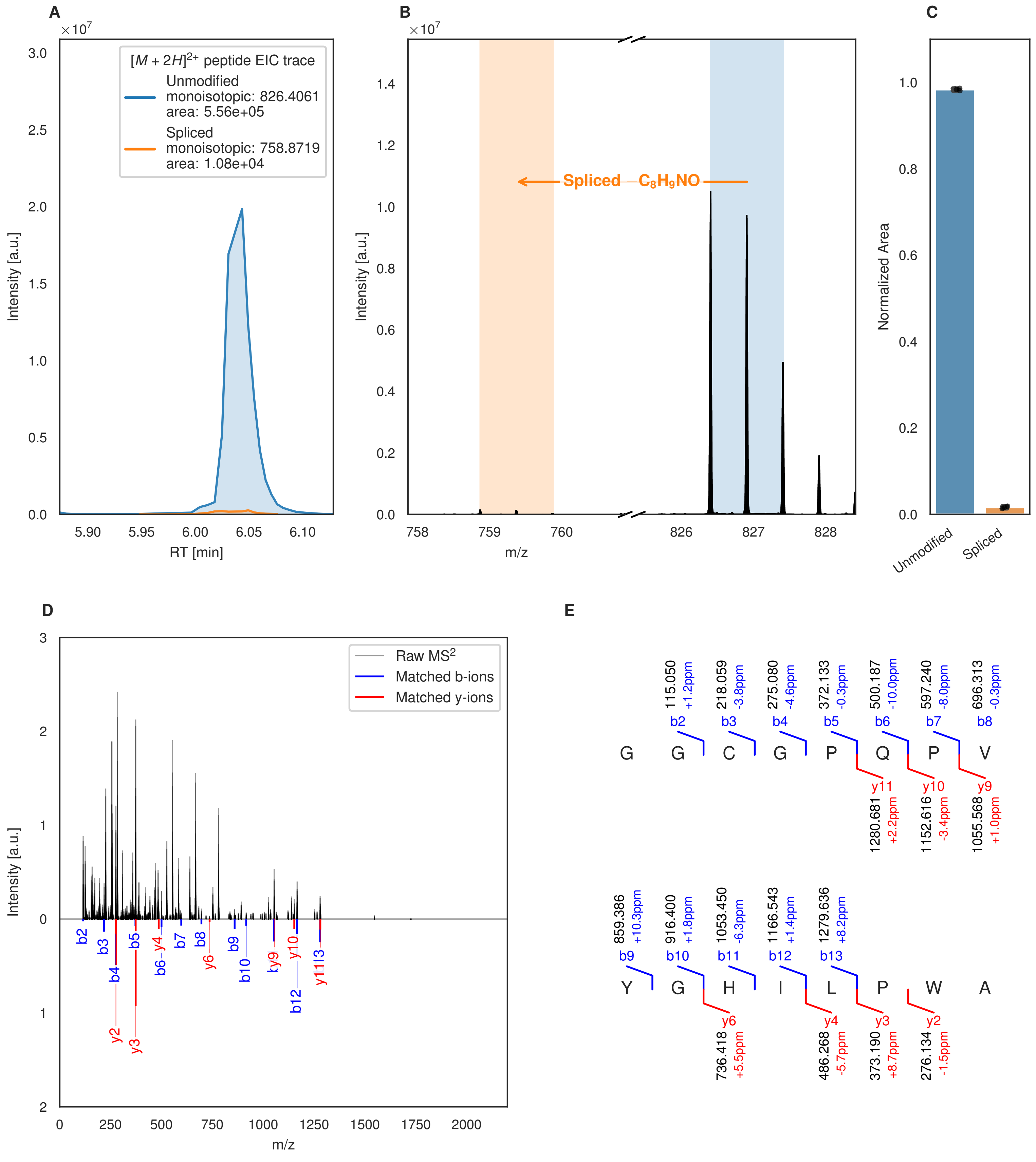
A
[M + 2H]2+ peptide EIC trace
Unmodified
monoisotopic: 826.4061
area: 5.56e+05
Spliced
monoisotopic: 758.8719
area: 1.08e+04
Intensity [a.u.]
RT [min]
B
Spliced −C8H9NO
Intensity [a.u.]
m/z
C
Normalized Area
Unmodified
Spliced
D
Raw MS2
Matched b-ions
Matched y-ions
Intensity [a.u.]
m/z
E
G G C G P Q P V
Y G H I L P W A

Figure S126 (figure continued from the previous page).

F

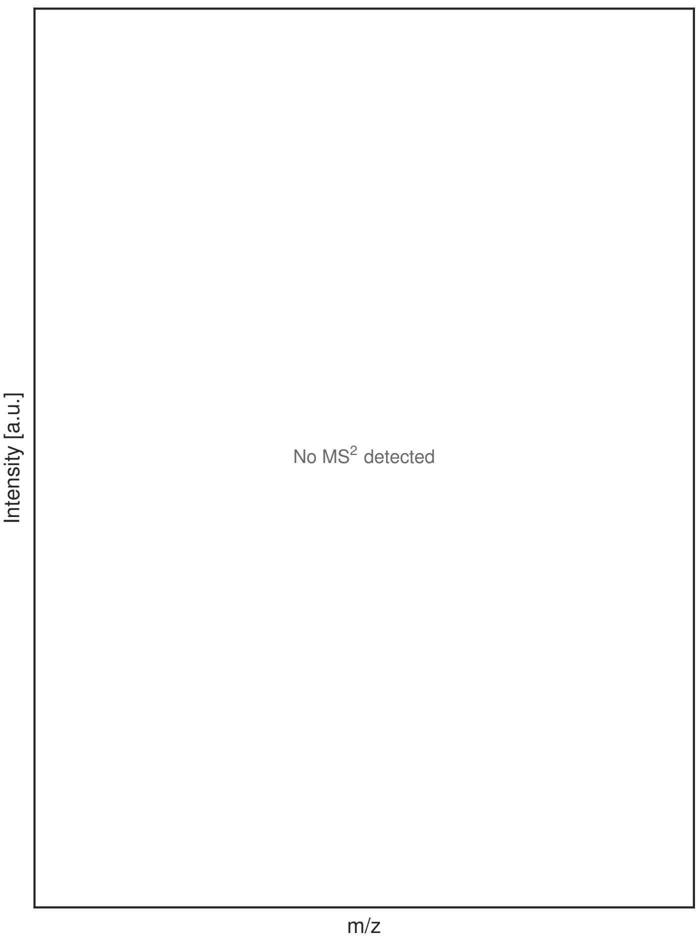


G

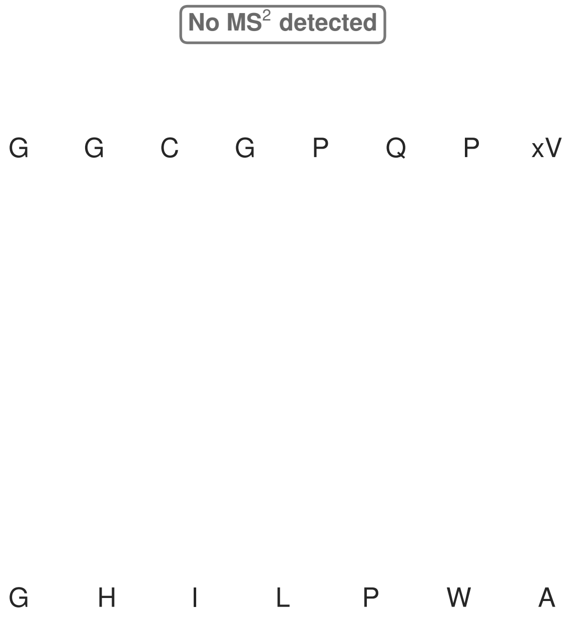


**Figure S126: Modification of the mutant_36 precursor peptide by PlpXY.** LC-MS analysis of IMAC-purified precursor peptides from *E. coli* BL21(DE3)Tuner containing splicease-encoding plasmid TJ-C-559 (PlpXY) and precursor peptide-encoding plasmid mutant_36 (RSTN1-PlpA3-TEV-mutant-36) after expression at 24 °C for 22 h in 1.5 mL cultures after digestion with TEV. Data from n=3 independent biological replicates. Spliced amino acids have an x as a prefix. Spliced refers to a single tyramine excision. A representative experimental replicate is shown in panels **A, B.**

**A** Close-up plot of the representative extracted ion chromatogram (EIC) of the various peptide states; the blue trace corresponds to the Unmodified peptide; the orange trace corresponds to the Spliced peptide. **B** Close-up plot of the m/z values of the peptides detected by the mass spectrometer, with the shaded colored areas corresponding to where the expected m/z values for the peptide m/z traces should occur, with the observed spectrum plotted in black. The Unmodified, Spliced peptide(s) were found in the mass spectrum. **C** Bar plot showing the relative abundance of the various peptide states at charge z=2; the bars represent the average relative abundances normalized to total peak areas, with each individual experimental replicate value plotted as a black dot. **D** Mirrored MS$^2$ plot for the Unmodified peptide. In this mirror plot, the full fragmentation spectrum is plotted in black pointing upward, and the matched fragment b ions (blue) and y ions (red) are plotted pointing downward, mirrored against it, so that each matched fragment can be compared directly with the measured spectrum. **E** MS$^2$ spectrum of the fragments corresponding to the Unmodified peptide sequence, with matched b (blue, above) and y ions (red, below) plotted. **F** Mirrored MS$^2$ plot for the Spliced peptide. In this mirror plot, the full fragmentation spectrum is plotted in black pointing upward, and the matched fragment b ions (blue) and y ions (red) are plotted pointing downward, mirrored against it, so that each matched fragment can be compared directly with the measured spectrum. **G** MS$^2$ spectrum of the fragments corresponding to the Spliced peptide sequence, with matched b (blue, above) and y ions (red, below) plotted.

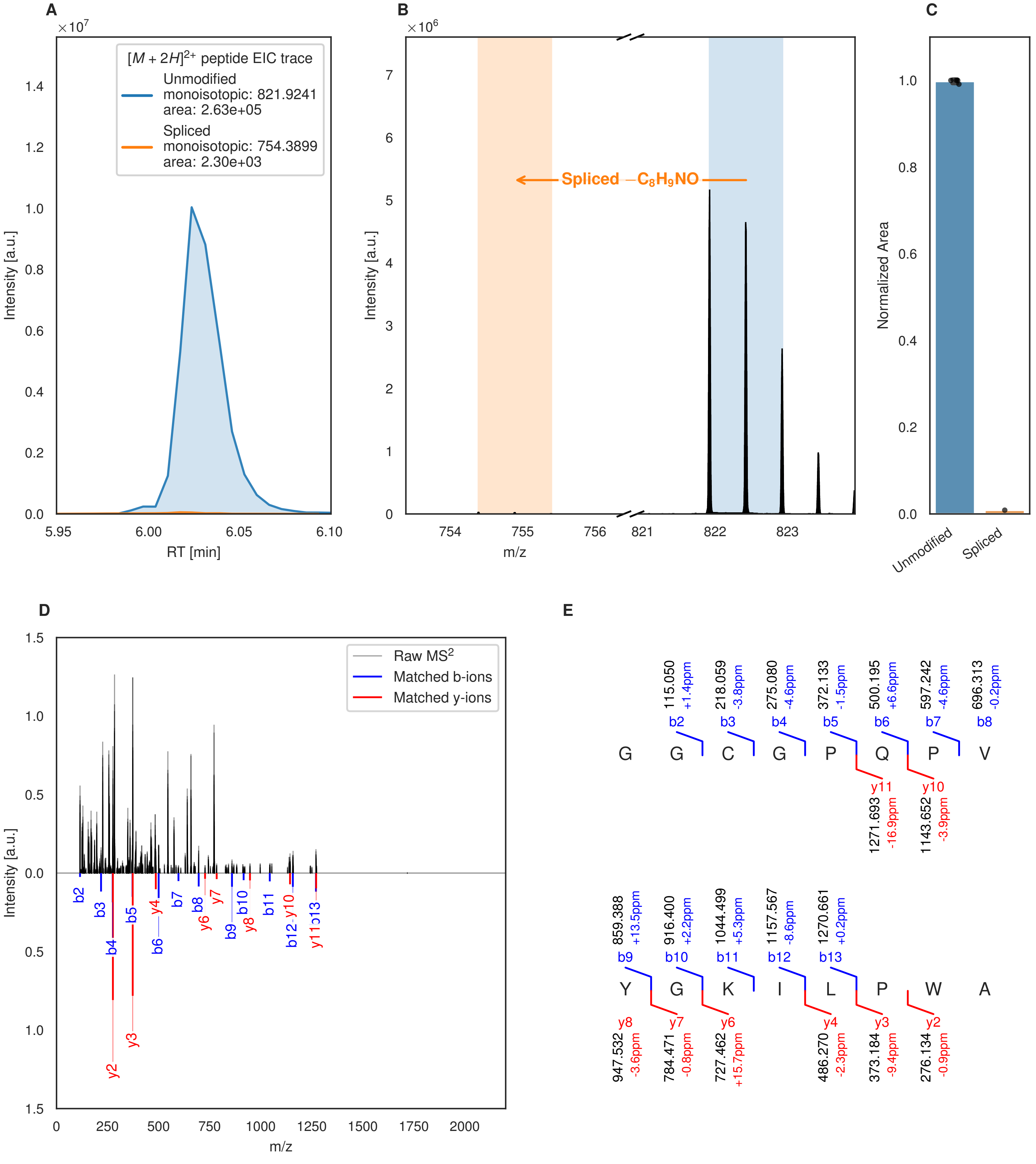
A
×10^7
[M + 2H]^2+ peptide EIC trace
Unmodified
monoisotopic: 821.9241
area: 2.63e+05
Spliced
monoisotopic: 754.3899
area: 2.30e+03
Intensity [a.u.]
RT [min]
B
×10^6
Spliced −C8H9NO
Intensity [a.u.]
m/z
C
Normalized Area
Unmodified
Spliced
D
Raw MS2
Matched b-ions
Matched y-ions
Intensity [a.u.]
m/z
E
G G C G P Q P V
b2 115.050 +1.4ppm
b3 218.059 -3.8ppm
b4 275.080 -4.6ppm
b5 372.133 -1.5ppm
b6 500.195 +6.6ppm
b7 597.242 -4.6ppm
b8 696.313 -0.2ppm
y11 1271.693 -16.9ppm
y10 1143.652 -3.9ppm
Y G K I L P W A
b9 859.388 +13.5ppm
b10 916.400 +2.2ppm
b11 1044.499 +5.3ppm
b12 1157.567 -8.6ppm
b13 1270.661 +0.2ppm
y8 947.532 -3.6ppm
y7 784.471 -0.8ppm
y6 727.462 +15.7ppm
y4 486.270 -2.3ppm
y3 373.184 -9.4ppm
y2 276.134 -0.9ppm

Figure S127 (figure continued from the previous page).

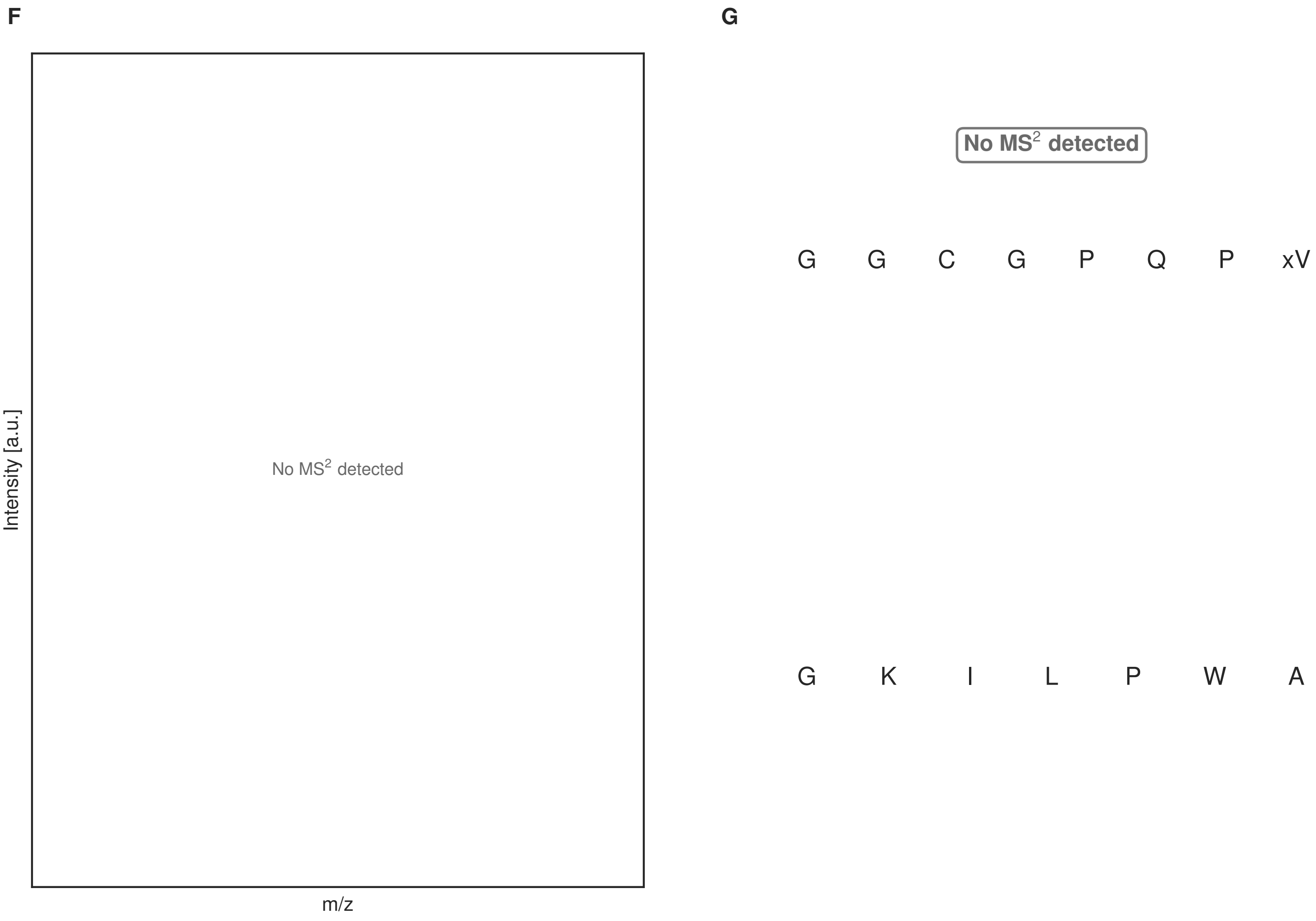


**Figure S127: Modification of the mutant_37 precursor peptide by HypXY.** LC-MS analysis of IMAC-purified precursor peptides from *E. coli* BL21(DE3)Tuner containing splicease-encoding plasmid TJ-C-689 (HypXY) and precursor peptide-encoding plasmid mutant_37 (RSTN1-HypAc-TEV-mutant-37) after expression at 24 °C for 22 h in 1.5 mL cultures after digestion with TEV. Data from n=4 independent biological replicates. Spliced amino acids have an x as a prefix. Spliced refers to a single tyramine excision. A representative experimental replicate is shown in panels **A, B.**

**A** Close-up plot of the representative extracted ion chromatogram (EIC) of the various peptide states; the blue trace corresponds to the Unmodified peptide; the orange trace corresponds to the Spliced peptide. **B** Close-up plot of the m/z values of the peptides detected by the mass spectrometer, with the shaded colored areas corresponding to where the expected m/z values for the peptide m/z traces should occur, with the observed spectrum plotted in black. The Unmodified, Spliced peptide(s) were found in the mass spectrum. **C** Bar plot showing the relative abundance of the various peptide states at charge z=2; the bars represent the average relative abundances normalized to total peak areas, with each individual experimental replicate value plotted as a black dot. **D** Mirrored MS$^2$ plot for the Unmodified peptide. In this mirror plot, the full fragmentation spectrum is plotted in black pointing upward, and the matched fragment b ions (blue) and y ions (red) are plotted pointing downward, mirrored against it, so that each matched fragment can be compared directly with the measured spectrum. **E** MS$^2$ spectrum of the fragments corresponding to the Unmodified peptide sequence, with matched b (blue, above) and y ions (red, below) plotted. **F** Mirrored MS$^2$ plot for the Spliced peptide. In this mirror plot, the full fragmentation spectrum is plotted in black pointing upward, and the matched fragment b ions (blue) and y ions (red) are plotted pointing downward, mirrored against it, so that each matched fragment can be compared directly with the measured spectrum. **G** MS$^2$ spectrum of the fragments corresponding to the Spliced peptide sequence, with matched b (blue, above) and y ions (red, below) plotted.

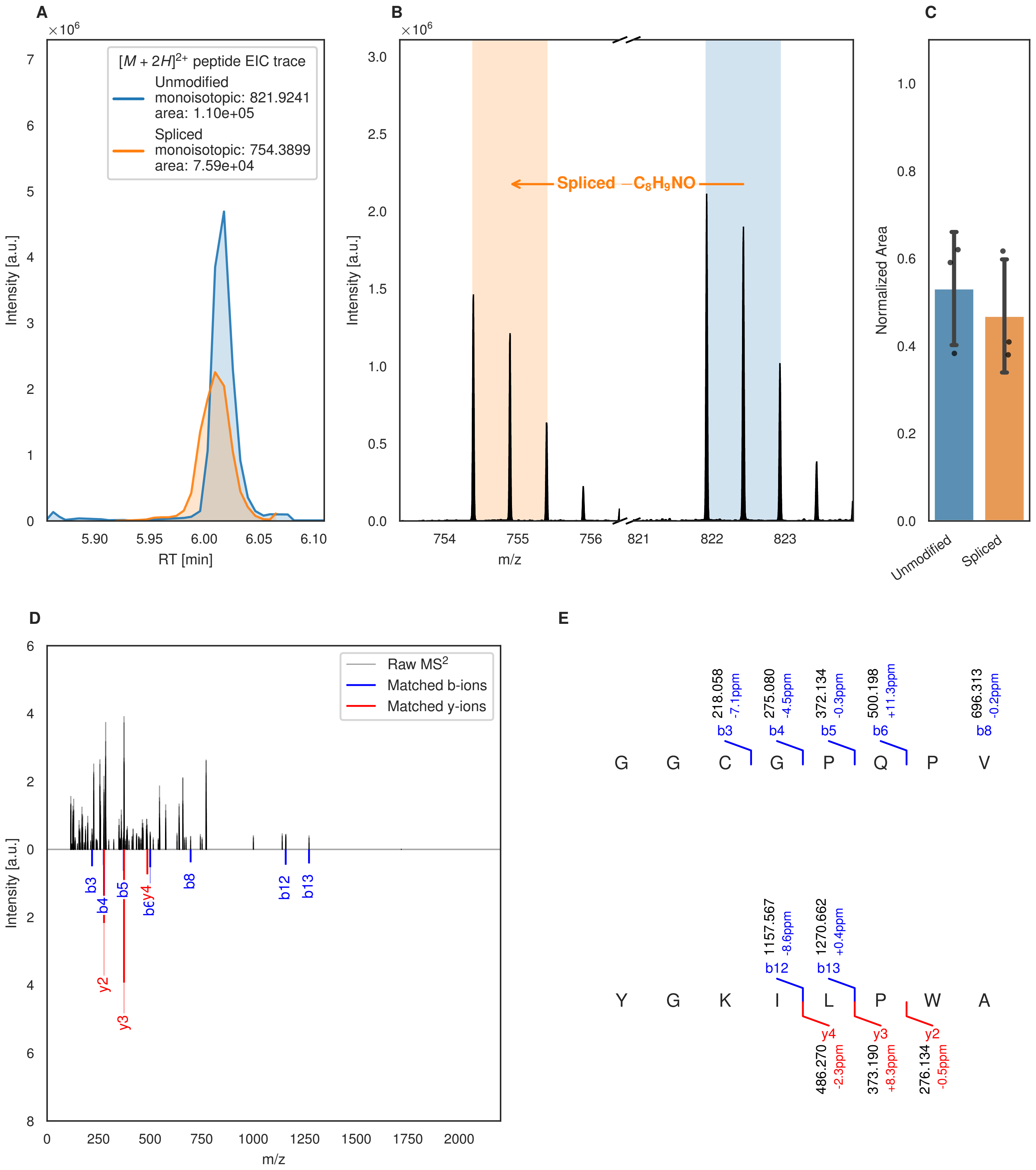
A
×10^6
[M + 2H]2+ peptide EIC trace
Unmodified
monoisotopic: 821.9241
area: 1.10e+05
Spliced
monoisotopic: 754.3899
area: 7.59e+04
Intensity [a.u.]
RT [min]
B
×10^6
Spliced −C8H9NO
Intensity [a.u.]
m/z
C
Normalized Area
Unmodified
Spliced
D
Raw MS2
Matched b-ions
Matched y-ions
Intensity [a.u.]
m/z
E
G G C G P Q P V
Y G K I L P W A
218.058 -7.1ppm b3
275.080 -4.5ppm b4
372.134 -0.3ppm b5
500.198 +11.3ppm b6
696.313 -0.2ppm b8
1157.567 -8.6ppm b12
1270.662 +0.4ppm b13
y4 486.270 -2.3ppm
y3 373.190 +8.3ppm
y2 276.134 -0.5ppm

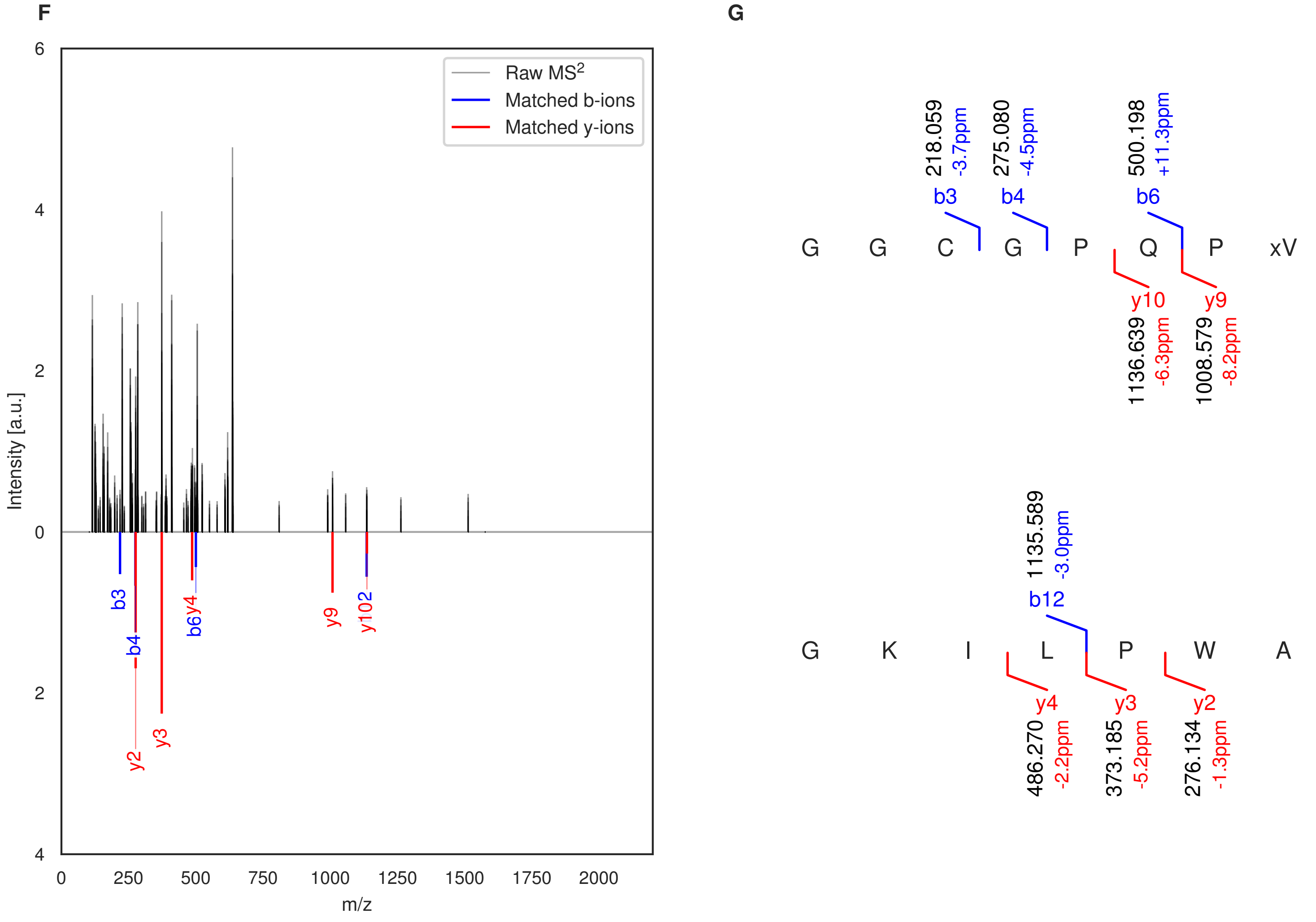


**Figure S128: Modification of the mutant_37 precursor peptide by PlpXY.** LC-MS analysis of IMAC-purified precursor peptides from *E. coli* BL21(DE3)Tuner containing splicease-encoding plasmid TJ-C-559 (PlpXY) and precursor peptide-encoding plasmid mutant_37 (RSTN1-PlpA3-TEV-mutant-37) after expression at 24 °C for 22 h in 1.5 mL cultures after digestion with TEV. Data from n=3 independent biological replicates. Spliced amino acids have an x as a prefix. Spliced refers to a single tyramine excision. A representative experimental replicate is shown in panels **A, B.**

**A** Close-up plot of the representative extracted ion chromatogram (EIC) of the various peptide states; the blue trace corresponds to the Unmodified peptide; the orange trace corresponds to the Spliced peptide. **B** Close-up plot of the m/z values of the peptides detected by the mass spectrometer, with the shaded colored areas corresponding to where the expected m/z values for the peptide m/z traces should occur, with the observed spectrum plotted in black. The Unmodified, Spliced peptide(s) were found in the mass spectrum. **C** Bar plot showing the relative abundance of the various peptide states at charge z=2; the bars represent the average relative abundances normalized to total peak areas, with each individual experimental replicate value plotted as a black dot. **D** Mirrored MS$^2$ plot for the Unmodified peptide. In this mirror plot, the full fragmentation spectrum is plotted in black pointing upward, and the matched fragment b ions (blue) and y ions (red) are plotted pointing downward, mirrored against it, so that each matched fragment can be compared directly with the measured spectrum. **E** MS$^2$ spectrum of the fragments corresponding to the Unmodified peptide sequence, with matched b (blue, above) and y ions (red, below) plotted. **F** Mirrored MS$^2$ plot for the Spliced peptide. In this mirror plot, the full fragmentation spectrum is plotted in black pointing upward, and the matched fragment b ions (blue) and y ions (red) are plotted pointing downward, mirrored against it, so that each matched fragment can be compared directly with the measured spectrum. **G** MS$^2$ spectrum of the fragments corresponding to the Spliced peptide sequence, with matched b (blue, above) and y ions (red, below) plotted.

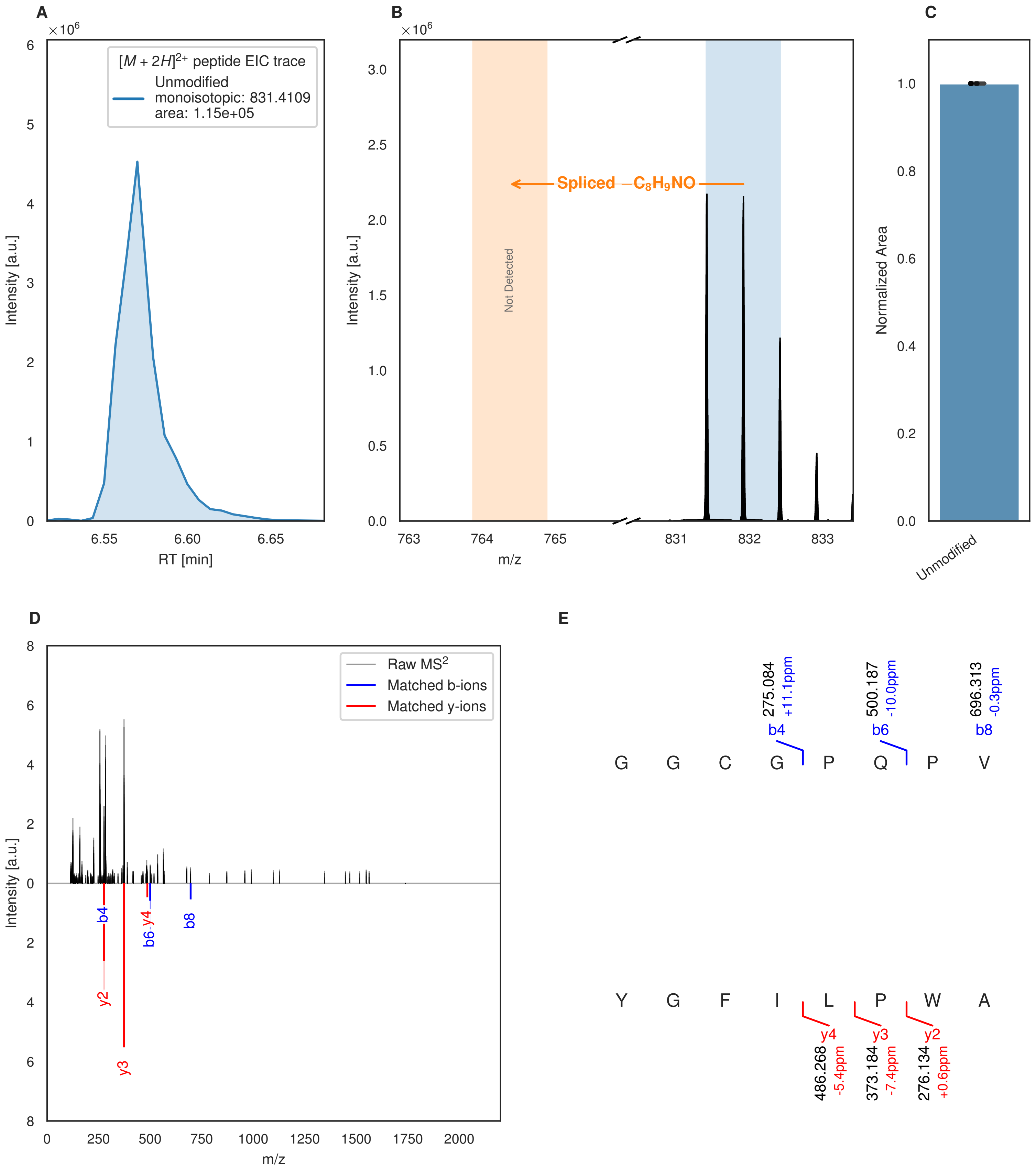


**Figure S129: Modification of the mutant_38 precursor peptide by HypXY.** LC-MS analysis of IMAC-purified precursor peptides from *E. coli* BL21(DE3)Tuner containing splicease-encoding plasmid TJ-C-689 (HypXY) and precursor peptide-encoding plasmid mutant_38 (RSTN1-HypAc-TEV-mutant-38) after expression at 24 °C for 22 h in 1.5 mL cultures after digestion with TEV. Data from n=4 independent biological replicates. Spliced amino acids have an x as a prefix. Spliced refers to a single tyramine excision. A representative experimental replicate is shown in panels **A, B.**

**A** Close-up plot of the representative extracted ion chromatogram (EIC) of the various peptide states; the blue trace corresponds to the Unmodified peptide. **B** Close-up plot of the m/z values of the peptides detected by the mass spectrometer, with the shaded colored areas corresponding to where the expected m/z values for the peptide m/z

(Caption continued on the next page.)

traces should occur, with the observed spectrum plotted in black. The Unmodified peptide(s) were found in the mass spectrum and the Spliced peptide(s) were not detected. **C** Bar plot showing the relative abundance of the various peptide states at charge z=2; the bars represent the average relative abundances normalized to total peak areas, with each individual experimental replicate value plotted as a black dot. **D** Mirrored $MS^2$ plot for the Unmodified peptide. In this mirror plot, the full fragmentation spectrum is plotted in black pointing upward, and the matched fragment b ions (blue) and y ions (red) are plotted pointing downward, mirrored against it, so that each matched fragment can be compared directly with the measured spectrum. **E** $MS^2$ spectrum of the fragments corresponding to the Unmodified peptide sequence, with matched b (blue, above) and y ions (red, below) plotted.

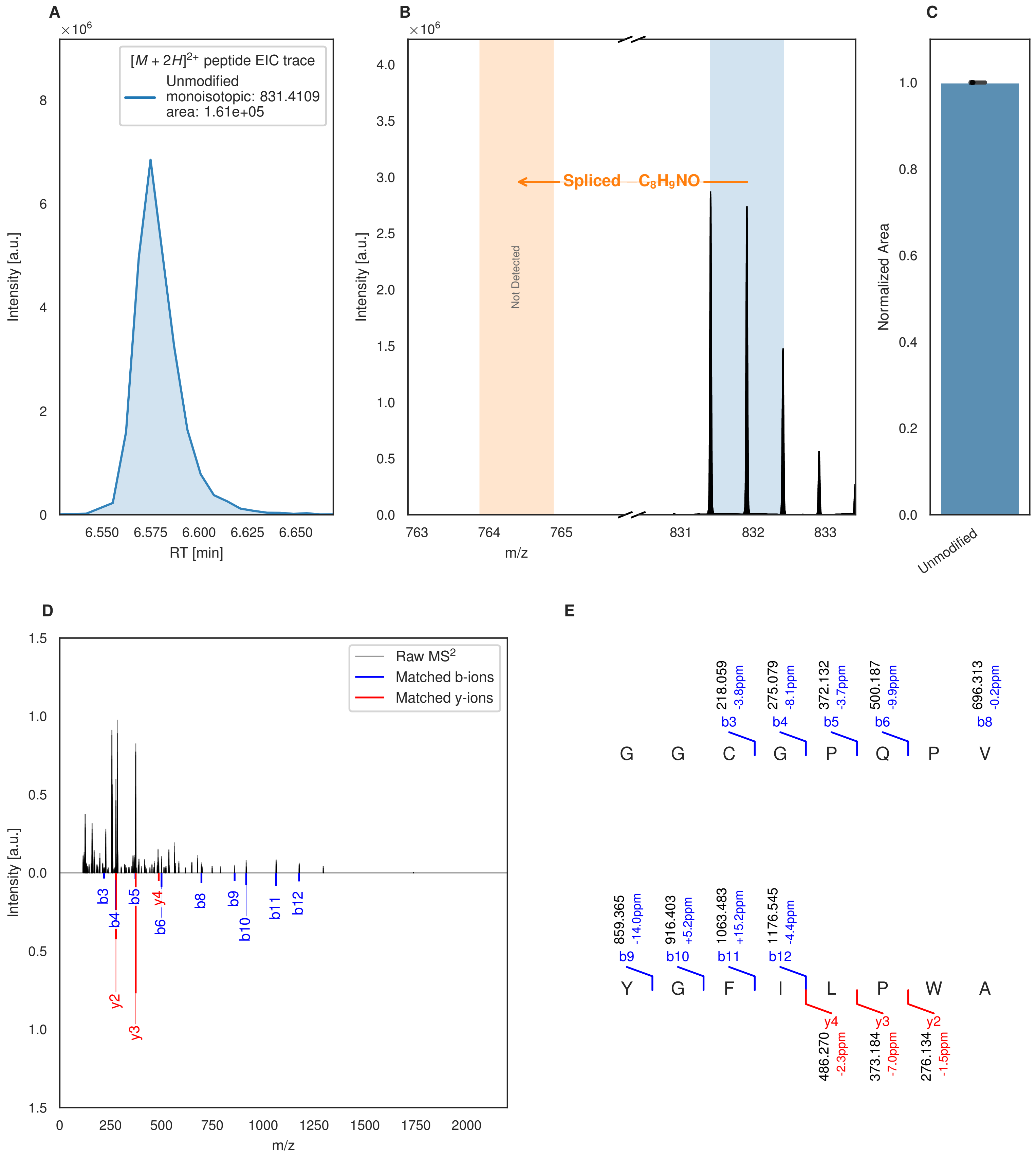


**Figure S130: Modification of the mutant_38 precursor peptide by PlpXY.** LC-MS analysis of IMAC-purified precursor peptides from *E. coli* BL21(DE3)Tuner containing splicease-encoding plasmid TJ-C-559 (PlpXY) and precursor peptide-encoding plasmid mutant_38 (RSTN1-PlpA3-TEV-mutant-38) after expression at 24 °C for 22 h in 1.5 mL cultures after digestion with TEV. Data from n=3 independent biological replicates. Spliced amino acids have an x as a prefix. Spliced refers to a single tyramine excision. A representative experimental replicate is shown in panels **A, B.**

**A** Close-up plot of the representative extracted ion chromatogram (EIC) of the various peptide states; the blue trace corresponds to the Unmodified peptide. **B** Close-up plot of the m/z values of the peptides detected by the mass spectrometer, with the shaded colored areas corresponding to where the expected m/z values for the peptide m/z

(Caption continued on the next page.)

traces should occur, with the observed spectrum plotted in black. The Unmodified peptide(s) were found in the mass spectrum and the Spliced peptide(s) were not detected. **C** Bar plot showing the relative abundance of the various peptide states at charge z=2; the bars represent the average relative abundances normalized to total peak areas, with each individual experimental replicate value plotted as a black dot. **D** Mirrored $MS^2$ plot for the Unmodified peptide. In this mirror plot, the full fragmentation spectrum is plotted in black pointing upward, and the matched fragment b ions (blue) and y ions (red) are plotted pointing downward, mirrored against it, so that each matched fragment can be compared directly with the measured spectrum. **E** $MS^2$ spectrum of the fragments corresponding to the Unmodified peptide sequence, with matched b (blue, above) and y ions (red, below) plotted.

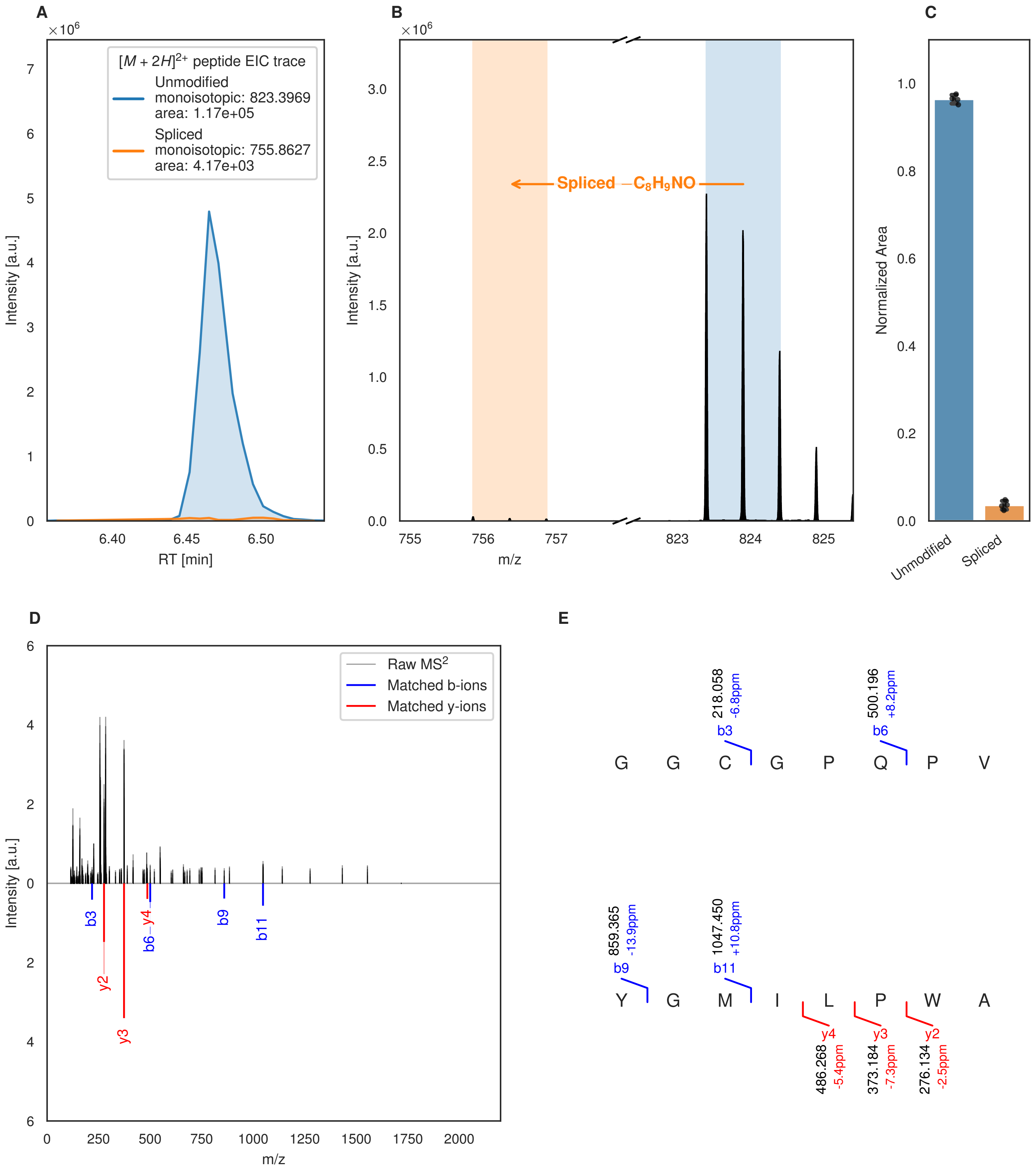

A
B
C
D
E
[M + 2H]2+ peptide EIC trace
Unmodified
monoisotopic: 823.3969
area: 1.17e+05
Spliced
monoisotopic: 755.8627
area: 4.17e+03
Intensity [a.u.]
RT [min]
Spliced −C8H9NO
m/z
Normalized Area
Unmodified
Spliced
Raw MS2
Matched b-ions
Matched y-ions
b3
b6
b9
b11
y2
y3
y4
G G C G P Q P V
Y G M I L P W A
218.058 -6.8ppm
500.196 +8.2ppm
859.365 -13.9ppm
1047.450 +10.8ppm
486.268 -5.4ppm
373.184 -7.3ppm
276.134 -2.5ppm

F

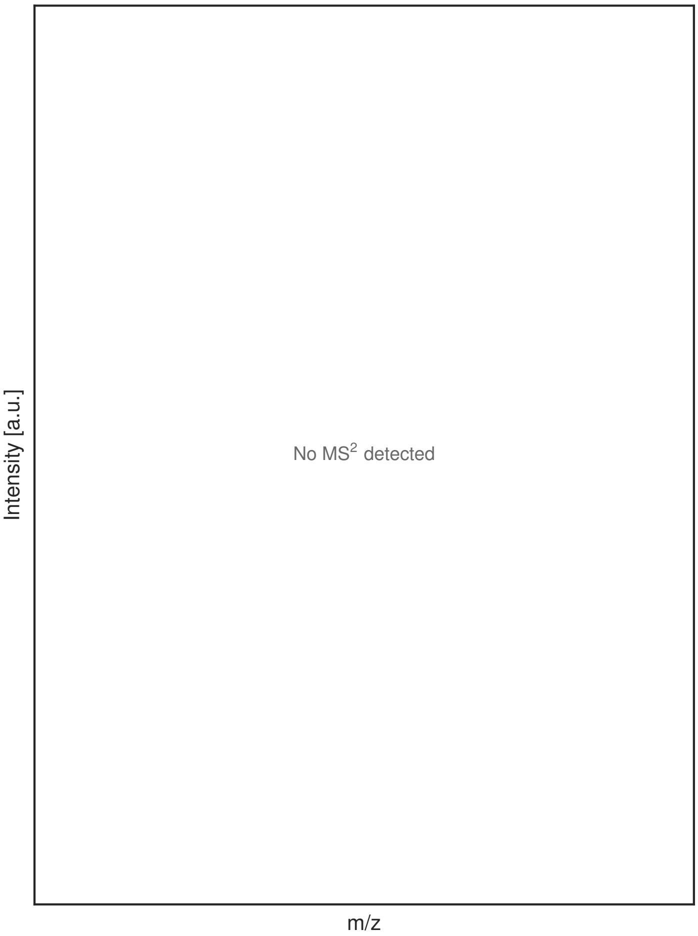


G

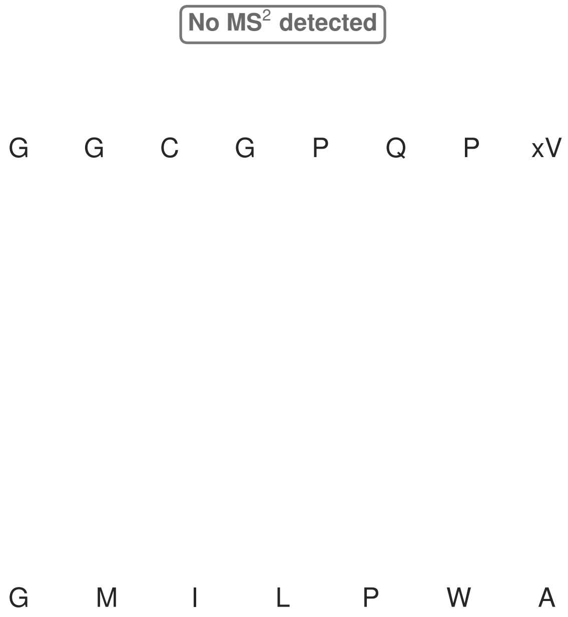


**Figure S131: Modification of the mutant_39 precursor peptide by HypXY.** LC-MS analysis of IMAC-purified precursor peptides from *E. coli* BL21(DE3)Tuner containing splicease-encoding plasmid TJ-C-689 (HypXY) and precursor peptide-encoding plasmid mutant_39 (RSTN1-HypAc-TEV-mutant-39) after expression at 24 °C for 22 h in 1.5 mL cultures after digestion with TEV. Data from n=4 independent biological replicates. Spliced amino acids have an x as a prefix. Spliced refers to a single tyramine excision. A representative experimental replicate is shown in panels **A, B.**

**A** Close-up plot of the representative extracted ion chromatogram (EIC) of the various peptide states; the blue trace corresponds to the Unmodified peptide; the orange trace corresponds to the Spliced peptide. **B** Close-up plot of the m/z values of the peptides detected by the mass spectrometer, with the shaded colored areas corresponding to where the expected m/z values for the peptide m/z traces should occur, with the observed spectrum plotted in black. The Unmodified, Spliced peptide(s) were found in the mass spectrum. **C** Bar plot showing the relative abundance of the various peptide states at charge z=2; the bars represent the average relative abundances normalized to total peak areas, with each individual experimental replicate value plotted as a black dot. **D** Mirrored $MS^2$ plot for the Unmodified peptide. In this mirror plot, the full fragmentation spectrum is plotted in black pointing upward, and the matched fragment b ions (blue) and y ions (red) are plotted pointing downward, mirrored against it, so that each matched fragment can be compared directly with the measured spectrum. **E** $MS^2$ spectrum of the fragments corresponding to the Unmodified peptide sequence, with matched b (blue, above) and y ions (red, below) plotted. **F** Mirrored $MS^2$ plot for the Spliced peptide. In this mirror plot, the full fragmentation spectrum is plotted in black pointing upward, and the matched fragment b ions (blue) and y ions (red) are plotted pointing downward, mirrored against it, so that each matched fragment can be compared directly with the measured spectrum. **G** $MS^2$ spectrum of the fragments corresponding to the Spliced peptide sequence, with matched b (blue, above) and y ions (red, below) plotted.

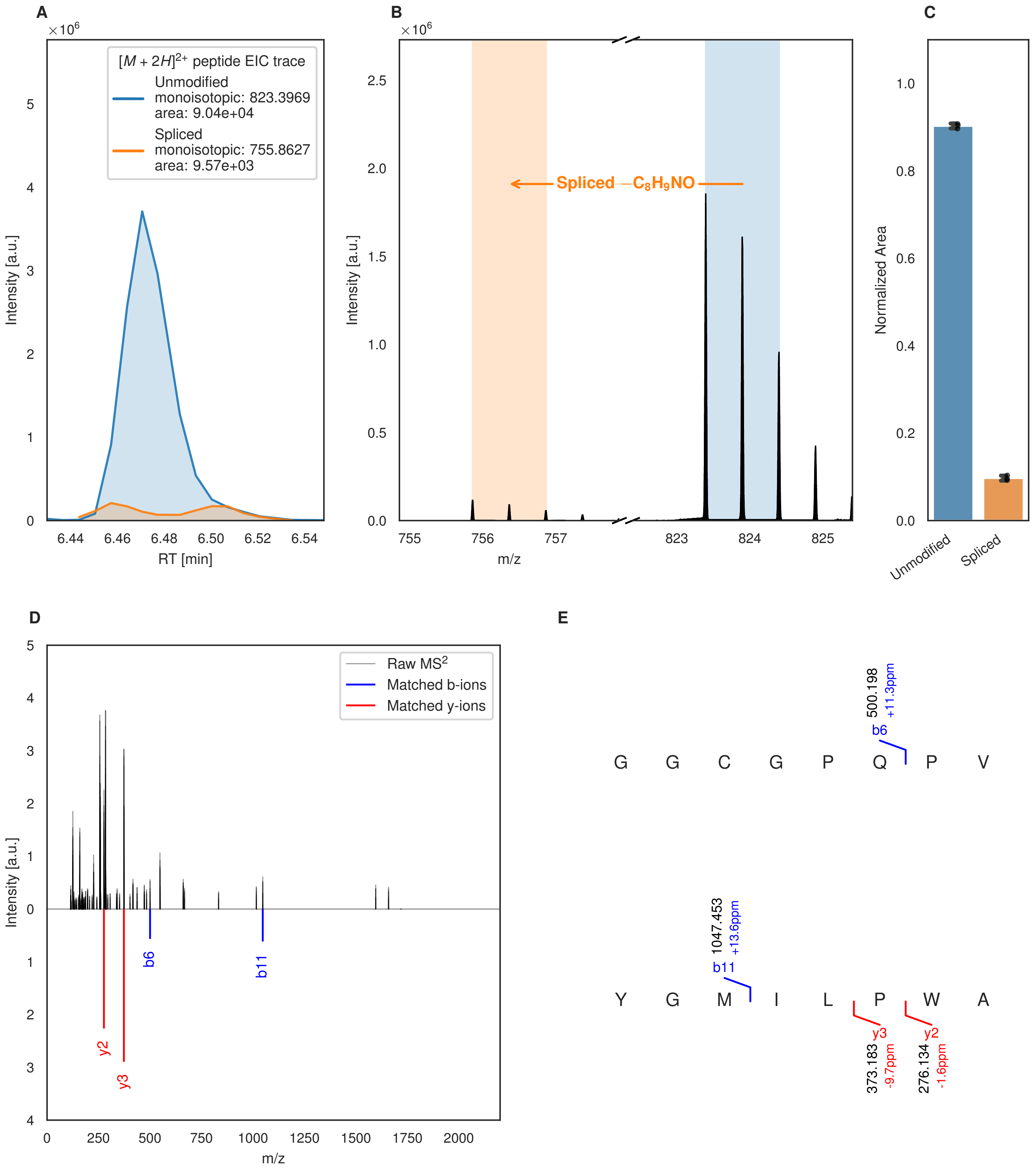
A
B
C
D
E
$[M+2H]^{2+}$ peptide EIC trace
Unmodified
monoisotopic: 823.3969
area: 9.04e+04
Spliced
monoisotopic: 755.8627
area: 9.57e+03
Intensity [a.u.]
RT [min]
Spliced $-C_8H_9NO$
m/z
Normalized Area
Unmodified
Spliced
Raw MS$^2$
Matched b-ions
Matched y-ions
b6
b11
y2
y3
G G C G P Q P V
500.198
+11.3ppm
Y G M I L P W A
1047.453
+13.6ppm
373.183
-9.7ppm
276.134
-1.6ppm

Figure S132 (figure continued from the previous page).

F

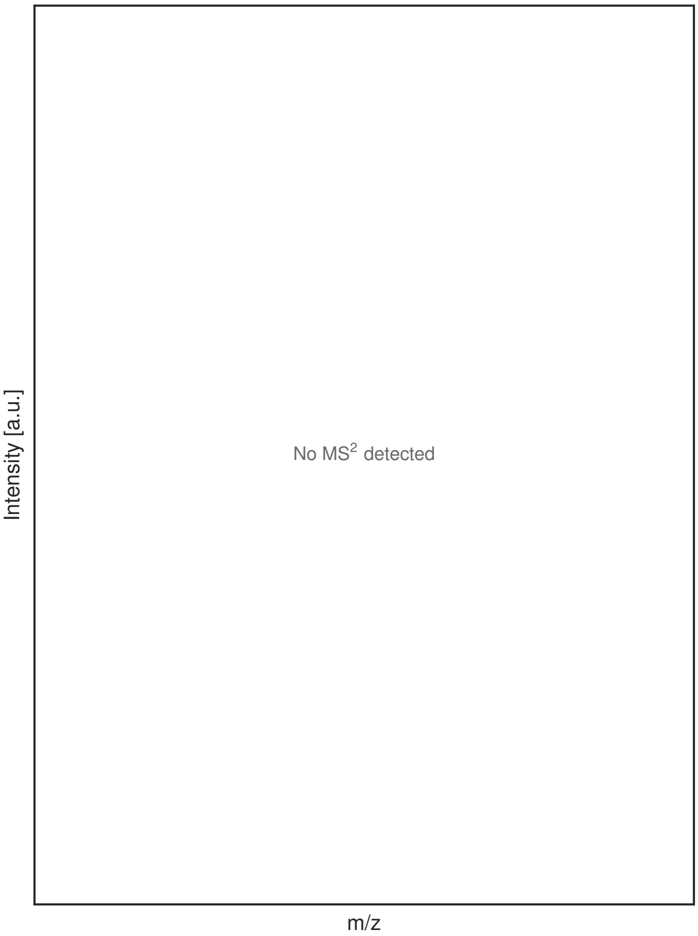


G

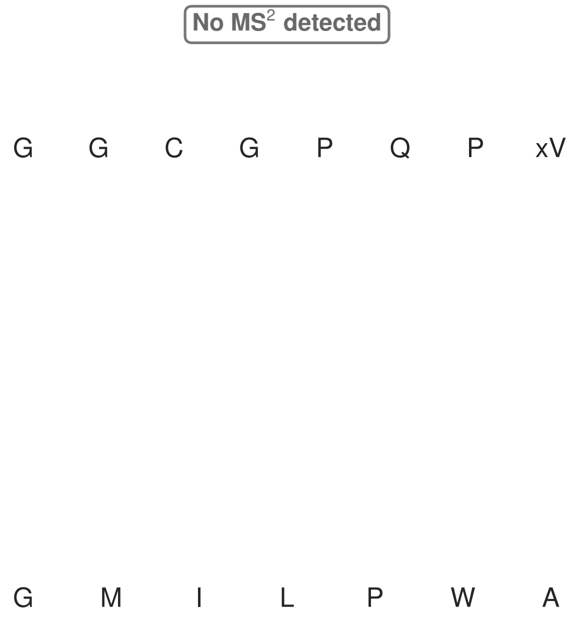


**Figure S132: Modification of the mutant_39 precursor peptide by PlpXY.** LC-MS analysis of IMAC-purified precursor peptides from *E. coli* BL21(DE3)Tuner containing splicease-encoding plasmid TJ-C-559 (PlpXY) and precursor peptide-encoding plasmid mutant_39 (RSTN1-PlpA3-TEV-mutant-39) after expression at 24 °C for 22 h in 1.5 mL cultures after digestion with TEV. Data from n=3 independent biological replicates. Spliced amino acids have an x as a prefix. Spliced refers to a single tyramine excision. A representative experimental replicate is shown in panels **A, B.**

**A** Close-up plot of the representative extracted ion chromatogram (EIC) of the various peptide states; the blue trace corresponds to the Unmodified peptide; the orange trace corresponds to the Spliced peptide. **B** Close-up plot of the m/z values of the peptides detected by the mass spectrometer, with the shaded colored areas corresponding to where the expected m/z values for the peptide m/z traces should occur, with the observed spectrum plotted in black. The Unmodified, Spliced peptide(s) were found in the mass spectrum. **C** Bar plot showing the relative abundance of the various peptide states at charge z=2; the bars represent the average relative abundances normalized to total peak areas, with each individual experimental replicate value plotted as a black dot. **D** Mirrored $MS^2$ plot for the Unmodified peptide. In this mirror plot, the full fragmentation spectrum is plotted in black pointing upward, and the matched fragment b ions (blue) and y ions (red) are plotted pointing downward, mirrored against it, so that each matched fragment can be compared directly with the measured spectrum. **E** $MS^2$ spectrum of the fragments corresponding to the Unmodified peptide sequence, with matched b (blue, above) and y ions (red, below) plotted. **F** Mirrored $MS^2$ plot for the Spliced peptide. In this mirror plot, the full fragmentation spectrum is plotted in black pointing upward, and the matched fragment b ions (blue) and y ions (red) are plotted pointing downward, mirrored against it, so that each matched fragment can be compared directly with the measured spectrum. **G** $MS^2$ spectrum of the fragments corresponding to the Spliced peptide sequence, with matched b (blue, above) and y ions (red, below) plotted.

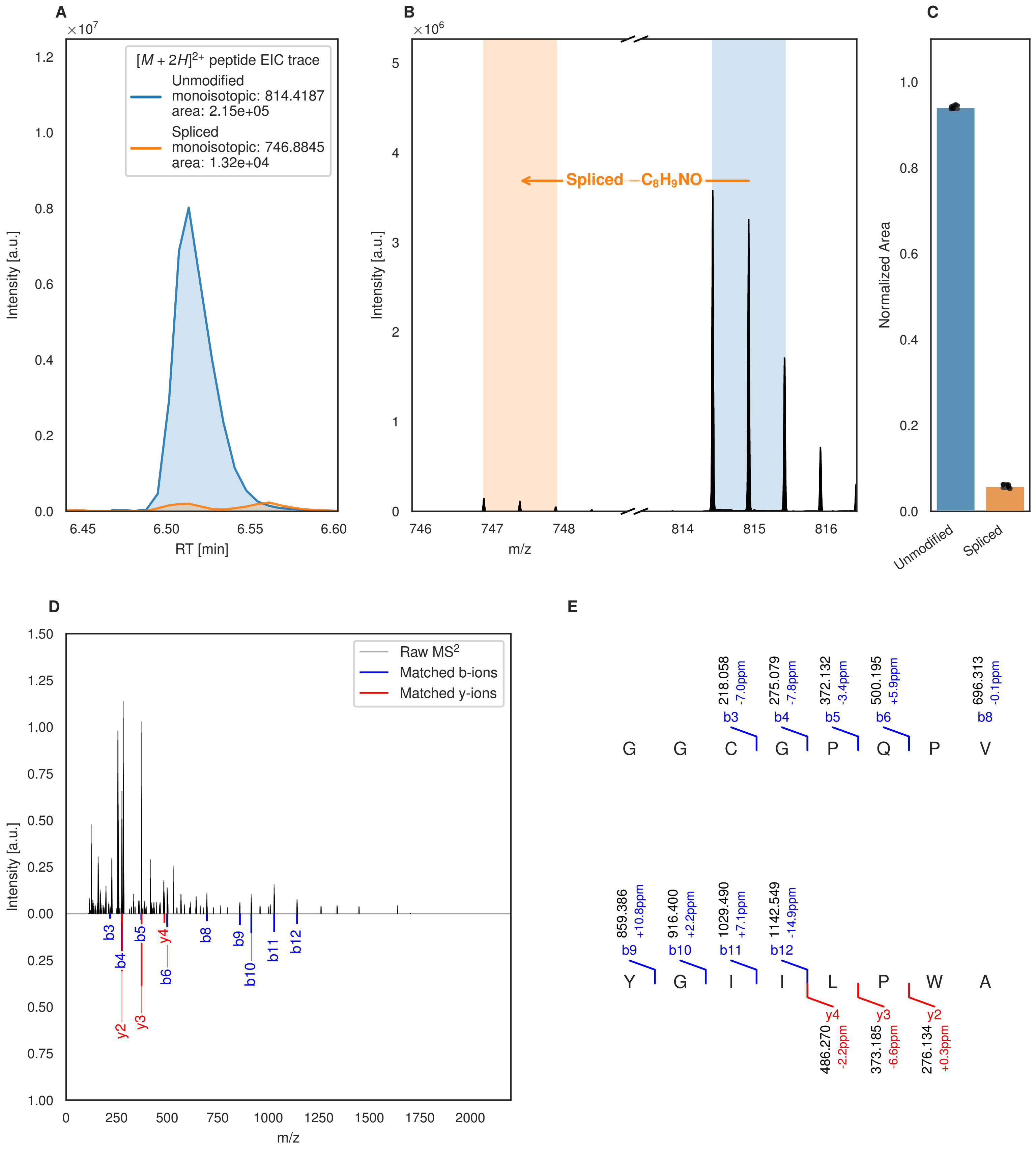

A
B
C
D
E
[M + 2H]2+ peptide EIC trace
Unmodified
monoisotopic: 814.4187
area: 2.15e+05
Spliced
monoisotopic: 746.8845
area: 1.32e+04
Intensity [a.u.]
RT [min]
Spliced −C8H9NO
m/z
Normalized Area
Unmodified
Spliced
Raw MS2
Matched b-ions
Matched y-ions
G G C G P Q P V
Y G I I L P W A
218.058 -7.0ppm b3
275.079 -7.8ppm b4
372.132 -3.4ppm b5
500.195 +5.9ppm b6
696.313 -0.1ppm b8
859.386 +10.8ppm b9
916.400 +2.2ppm b10
1029.490 +7.1ppm b11
1142.549 -14.9ppm b12
y4 486.270 -2.2ppm
y3 373.185 -6.6ppm
y2 276.134 +0.3ppm

Figure S133 (figure continued from the previous page).

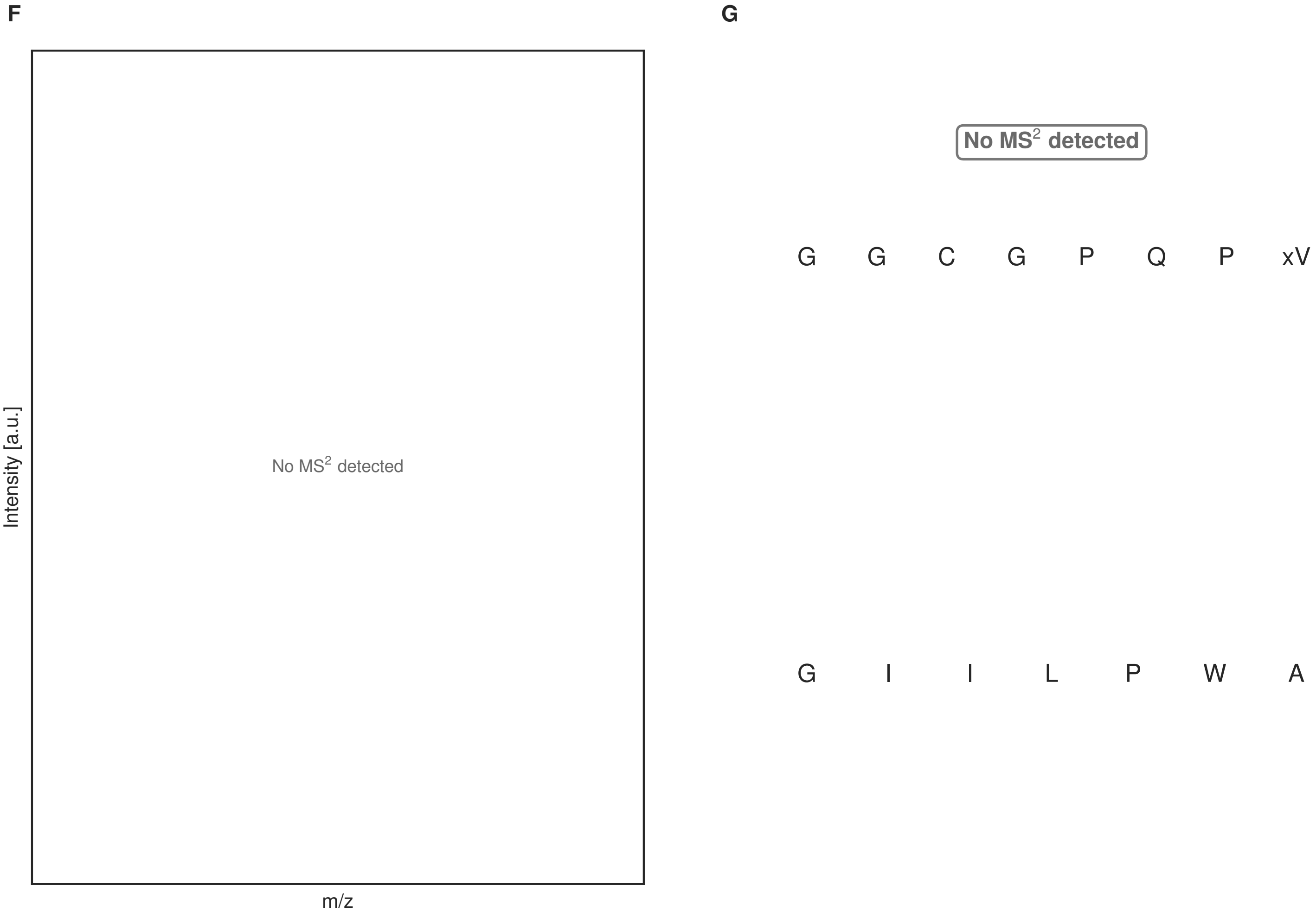


**Figure S133: Modification of the mutant_40 precursor peptide by HypXY.** LC-MS analysis of IMAC-purified precursor peptides from *E. coli* BL21(DE3)Tuner containing splicease-encoding plasmid TJ-C-689 (HypXY) and precursor peptide-encoding plasmid mutant_40 (RSTN1-HypAc-TEV-mutant-40) after expression at 24 °C for 22 h in 1.5 mL cultures after digestion with TEV. Data from n=4 independent biological replicates. Spliced amino acids have an x as a prefix. Spliced refers to a single tyramine excision. A representative experimental replicate is shown in panels **A, B.**

**A** Close-up plot of the representative extracted ion chromatogram (EIC) of the various peptide states; the blue trace corresponds to the Unmodified peptide; the orange trace corresponds to the Spliced peptide. **B** Close-up plot of the m/z values of the peptides detected by the mass spectrometer, with the shaded colored areas corresponding to where the expected m/z values for the peptide m/z traces should occur, with the observed spectrum plotted in black. The Unmodified, Spliced peptide(s) were found in the mass spectrum. **C** Bar plot showing the relative abundance of the various peptide states at charge z=2; the bars represent the average relative abundances normalized to total peak areas, with each individual experimental replicate value plotted as a black dot. **D** Mirrored MS$^2$ plot for the Unmodified peptide. In this mirror plot, the full fragmentation spectrum is plotted in black pointing upward, and the matched fragment b ions (blue) and y ions (red) are plotted pointing downward, mirrored against it, so that each matched fragment can be compared directly with the measured spectrum. **E** MS$^2$ spectrum of the fragments corresponding to the Unmodified peptide sequence, with matched b (blue, above) and y ions (red, below) plotted. **F** Mirrored MS$^2$ plot for the Spliced peptide. In this mirror plot, the full fragmentation spectrum is plotted in black pointing upward, and the matched fragment b ions (blue) and y ions (red) are plotted pointing downward, mirrored against it, so that each matched fragment can be compared directly with the measured spectrum. **G** MS$^2$ spectrum of the fragments corresponding to the Spliced peptide sequence, with matched b (blue, above) and y ions (red, below) plotted.

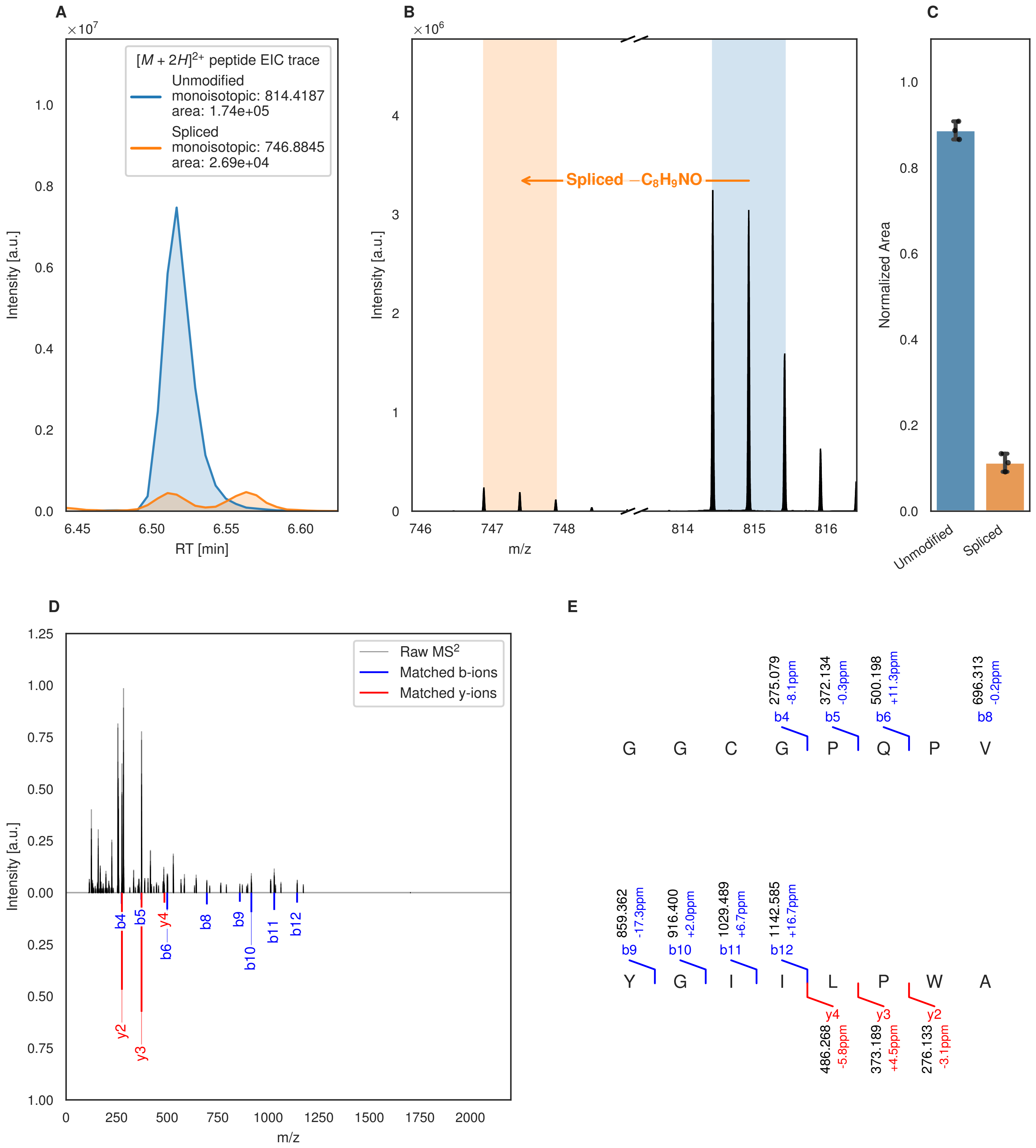
A
B
C
D
E
$[M+2H]^{2+}$ peptide EIC trace
Unmodified
monoisotopic: 814.4187
area: 1.74e+05
Spliced
monoisotopic: 746.8845
area: 2.69e+04
Intensity [a.u.]
RT [min]
Spliced $-C_8H_9NO$
m/z
Normalized Area
Unmodified
Spliced
Raw $MS^2$
Matched b-ions
Matched y-ions
G G C G P Q P V
Y G I I L P W A
275.079 -8.1ppm b4
372.134 -0.3ppm b5
500.198 +11.3ppm b6
696.313 -0.2ppm b8
859.362 -17.3ppm b9
916.400 +2.0ppm b10
1029.489 +6.7ppm b11
1142.585 +16.7ppm b12
y4 486.268 -5.8ppm
y3 373.189 +4.5ppm
y2 276.133 -3.1ppm

Figure S134 (figure continued from the previous page).

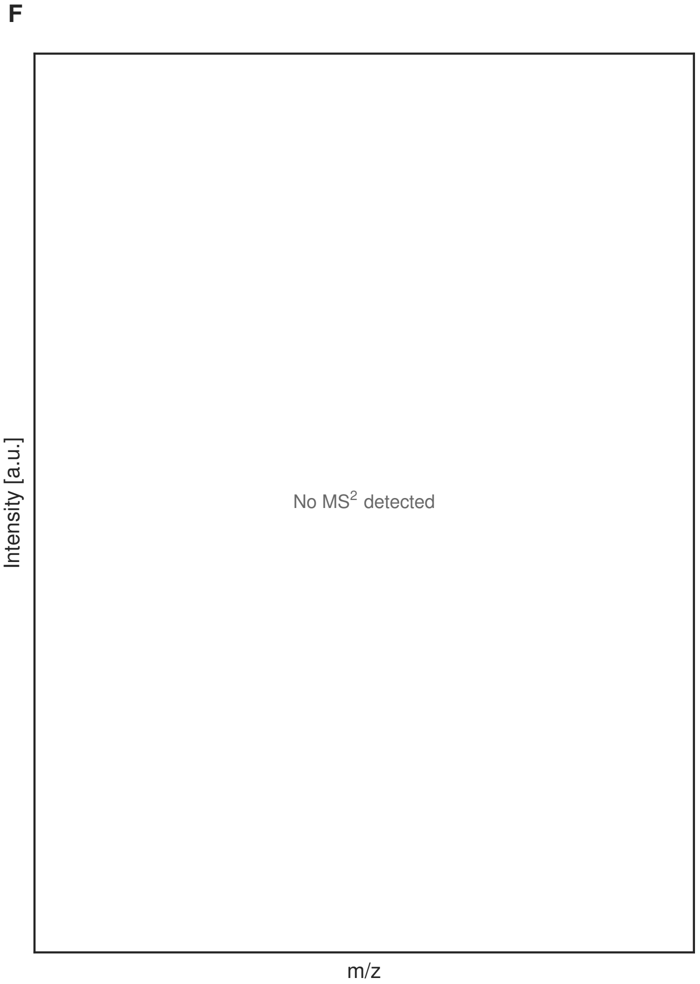


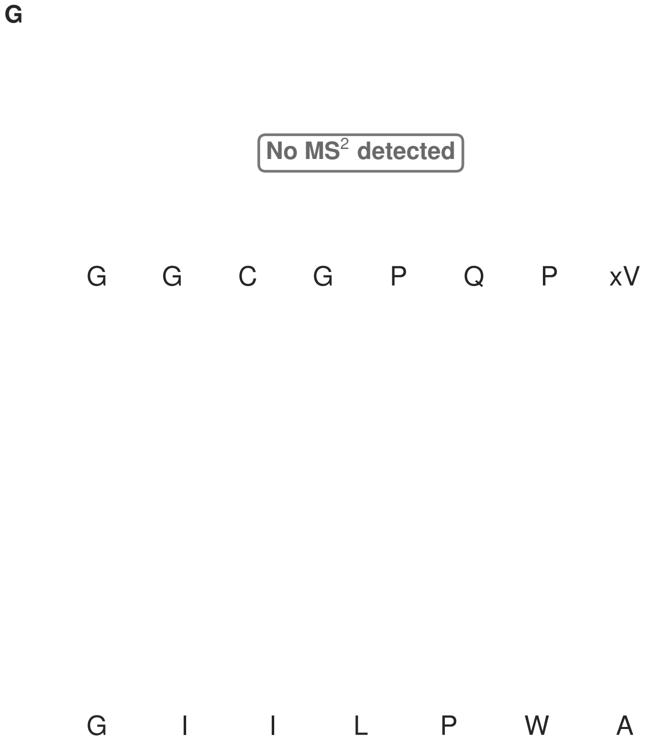


**Figure S134: Modification of the mutant_40 precursor peptide by PlpXY.** LC-MS analysis of IMAC-purified precursor peptides from *E. coli* BL21(DE3)Tuner containing splicease-encoding plasmid TJ-C-559 (PlpXY) and precursor peptide-encoding plasmid mutant_40 (RSTN1-PlpA3-TEV-mutant-40) after expression at 24 °C for 22 h in 1.5 mL cultures after digestion with TEV. Data from n=3 independent biological replicates. Spliced amino acids have an x as a prefix. Spliced refers to a single tyramine excision. A representative experimental replicate is shown in panels **A, B.**

**A** Close-up plot of the representative extracted ion chromatogram (EIC) of the various peptide states; the blue trace corresponds to the Unmodified peptide; the orange trace corresponds to the Spliced peptide. **B** Close-up plot of the m/z values of the peptides detected by the mass spectrometer, with the shaded colored areas corresponding to where the expected m/z values for the peptide m/z traces should occur, with the observed spectrum plotted in black. The Unmodified, Spliced peptide(s) were found in the mass spectrum. **C** Bar plot showing the relative abundance of the various peptide states at charge z=2; the bars represent the average relative abundances normalized to total peak areas, with each individual experimental replicate value plotted as a black dot. **D** Mirrored $MS^2$ plot for the Unmodified peptide. In this mirror plot, the full fragmentation spectrum is plotted in black pointing upward, and the matched fragment b ions (blue) and y ions (red) are plotted pointing downward, mirrored against it, so that each matched fragment can be compared directly with the measured spectrum. **E** $MS^2$ spectrum of the fragments corresponding to the Unmodified peptide sequence, with matched b (blue, above) and y ions (red, below) plotted. **F** Mirrored $MS^2$ plot for the Spliced peptide. In this mirror plot, the full fragmentation spectrum is plotted in black pointing upward, and the matched fragment b ions (blue) and y ions (red) are plotted pointing downward, mirrored against it, so that each matched fragment can be compared directly with the measured spectrum. **G** $MS^2$ spectrum of the fragments corresponding to the Spliced peptide sequence, with matched b (blue, above) and y ions (red, below) plotted.

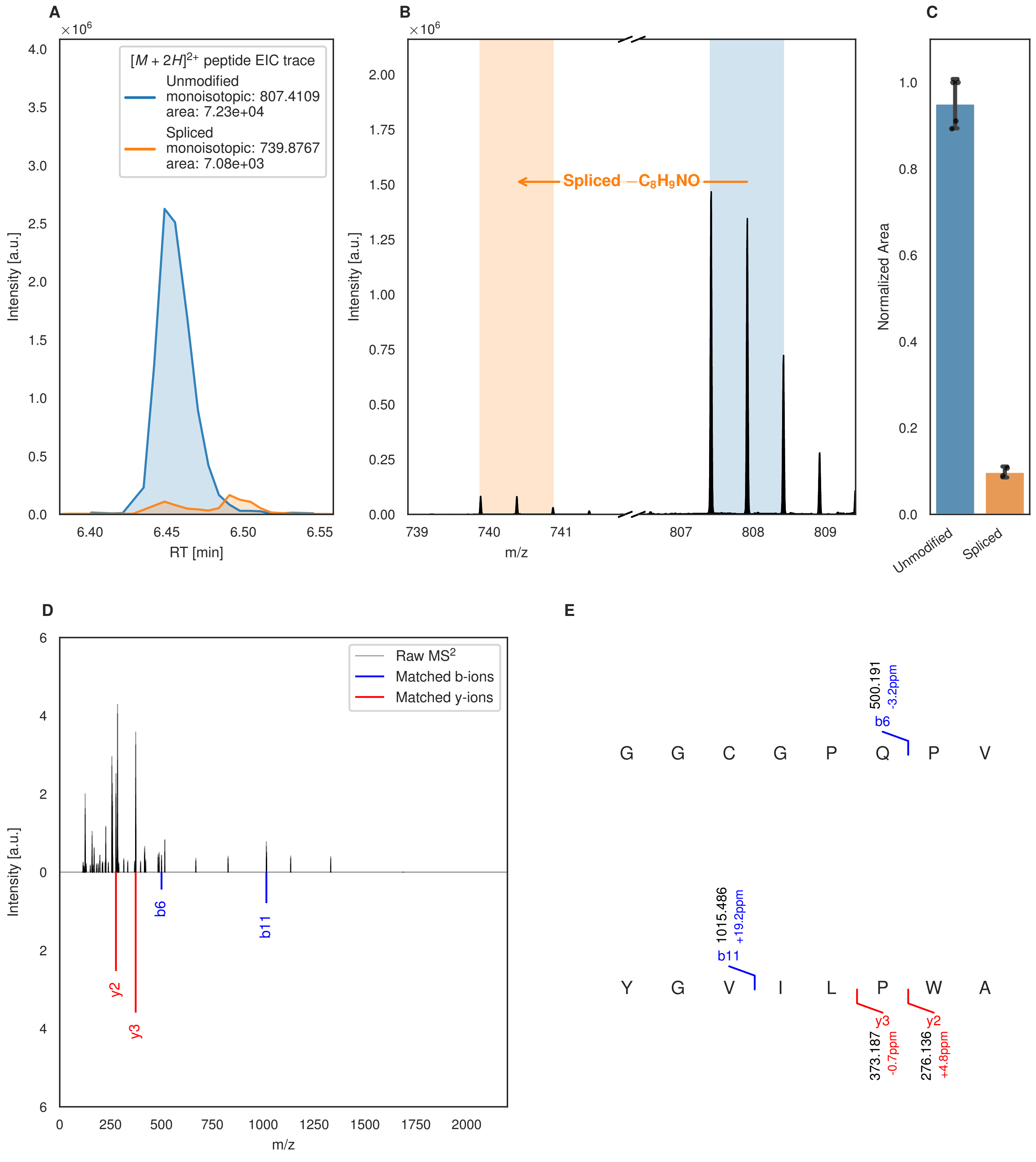
A
[M + 2H]2+ peptide EIC trace
Unmodified
monoisotopic: 807.4109
area: 7.23e+04
Spliced
monoisotopic: 739.8767
area: 7.08e+03
Intensity [a.u.]
RT [min]
B
Spliced −C8H9NO
m/z
C
Normalized Area
Unmodified
Spliced
D
Raw MS2
Matched b-ions
Matched y-ions
b6
b11
y2
y3
E
G G C G P Q P V
500.191
-3.2ppm
b6
Y G V I L P W A
1015.486
+19.2ppm
b11
y3
373.187
-0.7ppm
y2
276.136
+4.8ppm

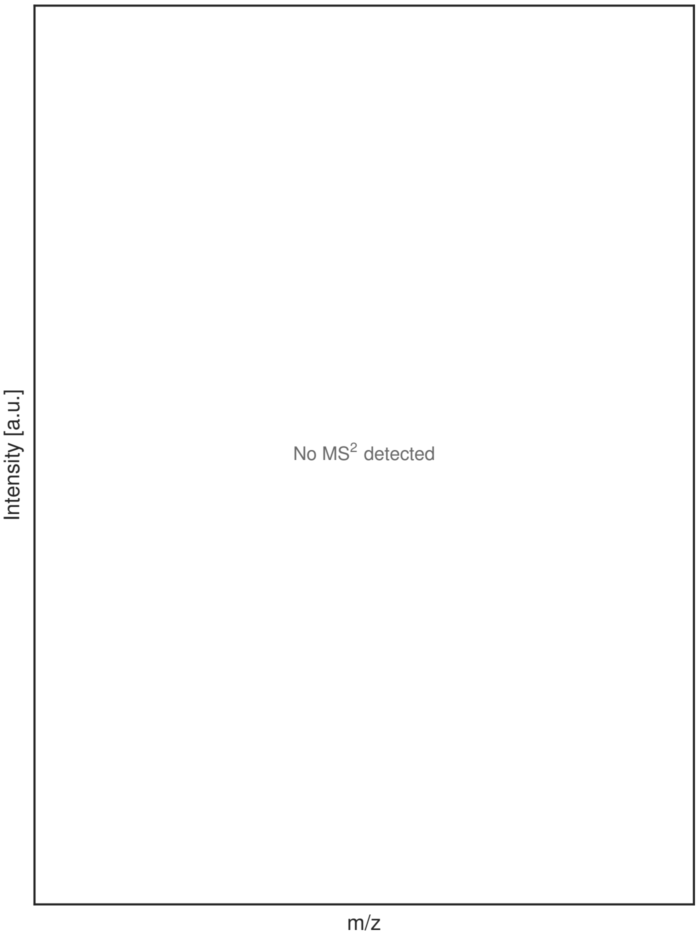


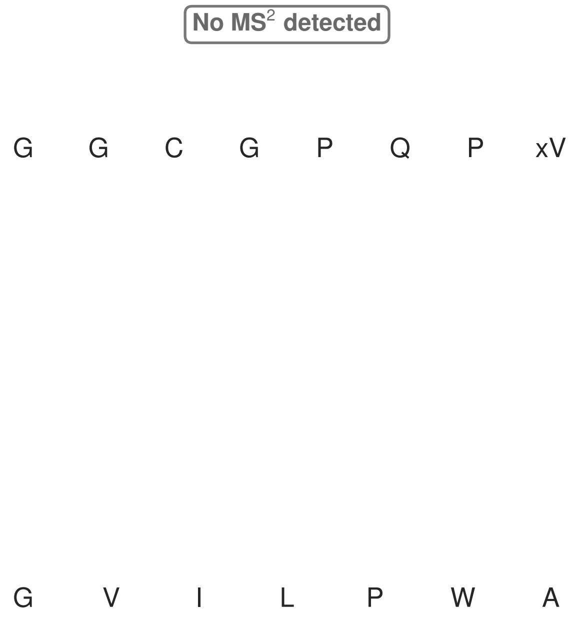


**Figure S135: Modification of the TJ-C-710 precursor peptide by PlpXY.** LC-MS analysis of IMAC-purified precursor peptides from *E. coli* BL21(DE3)Tuner containing splicease-encoding plasmid TJ-C-599 (PlpXY) and precursor peptide-encoding plasmid TJ-C-710 (RSTN-PlpA3-TEV-Artificial-2) after expression at 24 °C for 22 h in 1.5 mL cultures after digestion with TEV. Data from n=4 independent biological replicates. Spliced amino acids have an x as a prefix. Spliced refers to a single tyramine excision. A representative experimental replicate is shown in panels **A, B.**

**A** Close-up plot of the representative extracted ion chromatogram (EIC) of the various peptide states; the blue trace corresponds to the Unmodified peptide; the orange trace corresponds to the Spliced peptide. **B** Close-up plot of the m/z values of the peptides detected by the mass spectrometer, with the shaded colored areas corresponding to where the expected m/z values for the peptide m/z traces should occur, with the observed spectrum plotted in black. The Unmodified, Spliced peptide(s) were found in the mass spectrum. **C** Bar plot showing the relative abundance of the various peptide states at charge z=2; the bars represent the average relative abundances normalized to total peak areas, with each individual experimental replicate value plotted as a black dot. **D** Mirrored MS$^2$ plot for the Unmodified peptide. In this mirror plot, the full fragmentation spectrum is plotted in black pointing upward, and the matched fragment b ions (blue) and y ions (red) are plotted pointing downward, mirrored against it, so that each matched fragment can be compared directly with the measured spectrum. **E** MS$^2$ spectrum of the fragments corresponding to the Unmodified peptide sequence, with matched b (blue, above) and y ions (red, below) plotted. **F** Mirrored MS$^2$ plot for the Spliced peptide. In this mirror plot, the full fragmentation spectrum is plotted in black pointing upward, and the matched fragment b ions (blue) and y ions (red) are plotted pointing downward, mirrored against it, so that each matched fragment can be compared directly with the measured spectrum. **G** MS$^2$ spectrum of the fragments corresponding to the Spliced peptide sequence, with matched b (blue, above) and y ions (red, below) plotted.

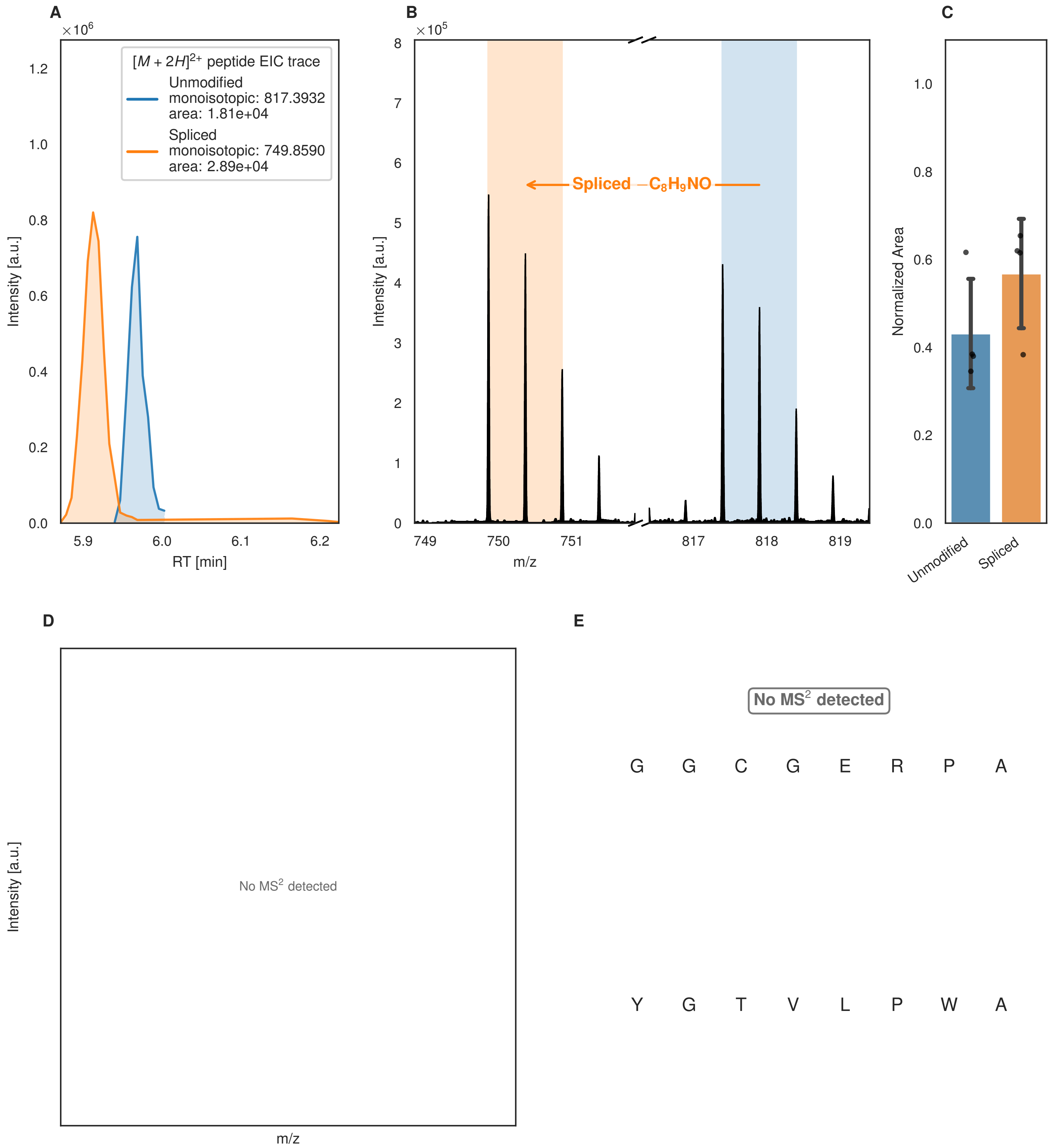
A
×10^6
[M + 2H]2+ peptide EIC trace
Unmodified
monoisotopic: 817.3932
area: 1.81e+04
Spliced
monoisotopic: 749.8590
area: 2.89e+04
Intensity [a.u.]
RT [min]
B
×10^5
Spliced −C8H9NO
Intensity [a.u.]
m/z
C
Normalized Area
Unmodified
Spliced
D
No MS2 detected
Intensity [a.u.]
m/z
E
No MS2 detected
G G C G E R P A
Y G T V L P W A

F

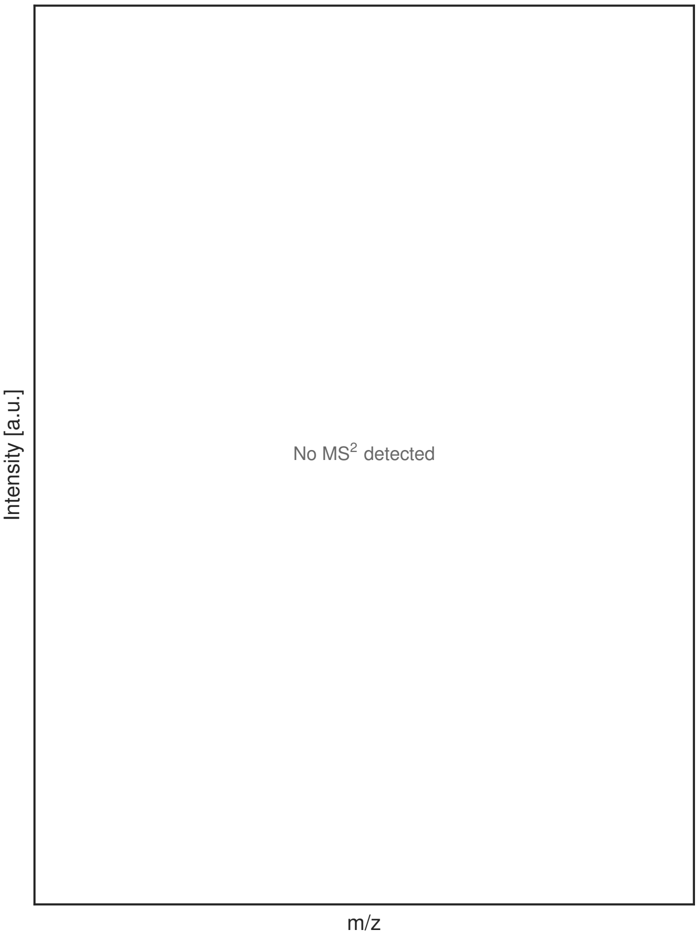


G

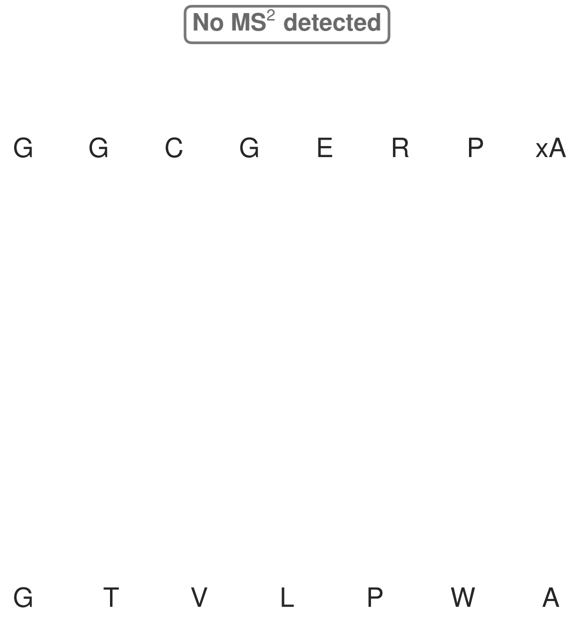


**Figure S136: Modification of the design_1 precursor peptide by PlpXY.** LC-MS analysis of IMAC-purified precursor peptides from *E. coli* BL21(DE3)Tuner containing splicease-encoding plasmid TJ-C-599 (PlpXY) and precursor peptide-encoding plasmid design_1 (RSTN-PlpA3-TEV-designed-1) after expression at 24 °C for 22 h in 1.5 mL cultures after digestion with TEV. Data from n=4 independent biological replicates. Spliced amino acids have an x as a prefix. Spliced refers to a single tyramine excision. A representative experimental replicate is shown in panels **A, B.**

**A** Close-up plot of the representative extracted ion chromatogram (EIC) of the various peptide states; the blue trace corresponds to the Unmodified peptide; the orange trace corresponds to the Spliced peptide. **B** Close-up plot of the m/z values of the peptides detected by the mass spectrometer, with the shaded colored areas corresponding to where the expected m/z values for the peptide m/z traces should occur, with the observed spectrum plotted in black. The Unmodified, Spliced peptide(s) were found in the mass spectrum. **C** Bar plot showing the relative abundance of the various peptide states at charge z=2; the bars represent the average relative abundances normalized to total peak areas, with each individual experimental replicate value plotted as a black dot. **D** Mirrored MS$^2$ plot for the Unmodified peptide. In this mirror plot, the full fragmentation spectrum is plotted in black pointing upward, and the matched fragment b ions (blue) and y ions (red) are plotted pointing downward, mirrored against it, so that each matched fragment can be compared directly with the measured spectrum. **E** MS$^2$ spectrum of the fragments corresponding to the Unmodified peptide sequence, with matched b (blue, above) and y ions (red, below) plotted. **F** Mirrored MS$^2$ plot for the Spliced peptide. In this mirror plot, the full fragmentation spectrum is plotted in black pointing upward, and the matched fragment b ions (blue) and y ions (red) are plotted pointing downward, mirrored against it, so that each matched fragment can be compared directly with the measured spectrum. **G** MS$^2$ spectrum of the fragments corresponding to the Spliced peptide sequence, with matched b (blue, above) and y ions (red, below) plotted.

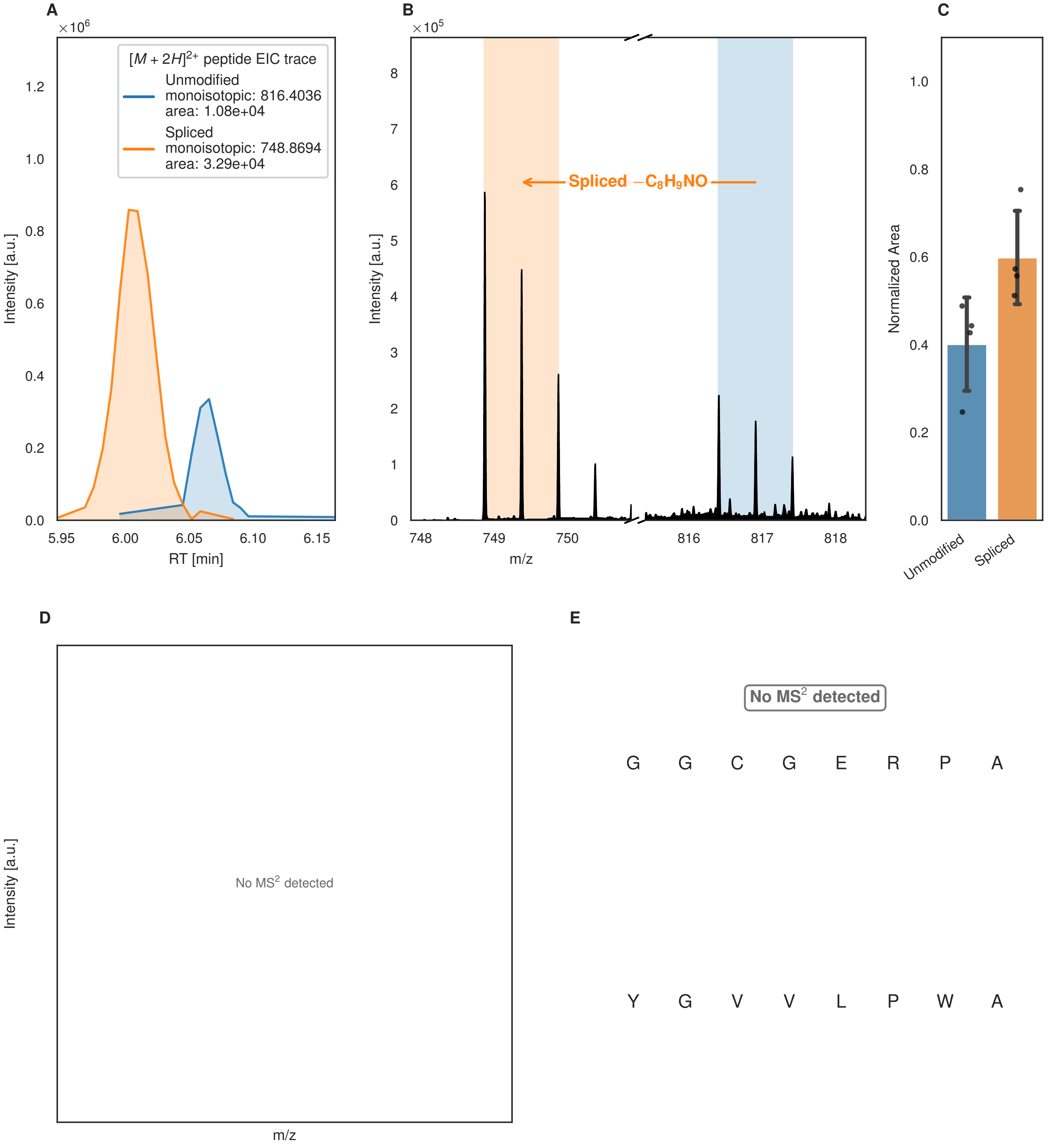

A
×10^6
[M + 2H]2+ peptide EIC trace
Unmodified
monoisotopic: 816.4036
area: 1.08e+04
Spliced
monoisotopic: 748.8694
area: 3.29e+04
Intensity [a.u.]
RT [min]
B
×10^5
Spliced −C8H9NO
Intensity [a.u.]
m/z
C
Normalized Area
Unmodified
Spliced
D
Intensity [a.u.]
No MS2 detected
m/z
E
No MS2 detected
G G C G E R P A
Y G V V L P W A

Figure S137 (figure continued from the previous page).

F

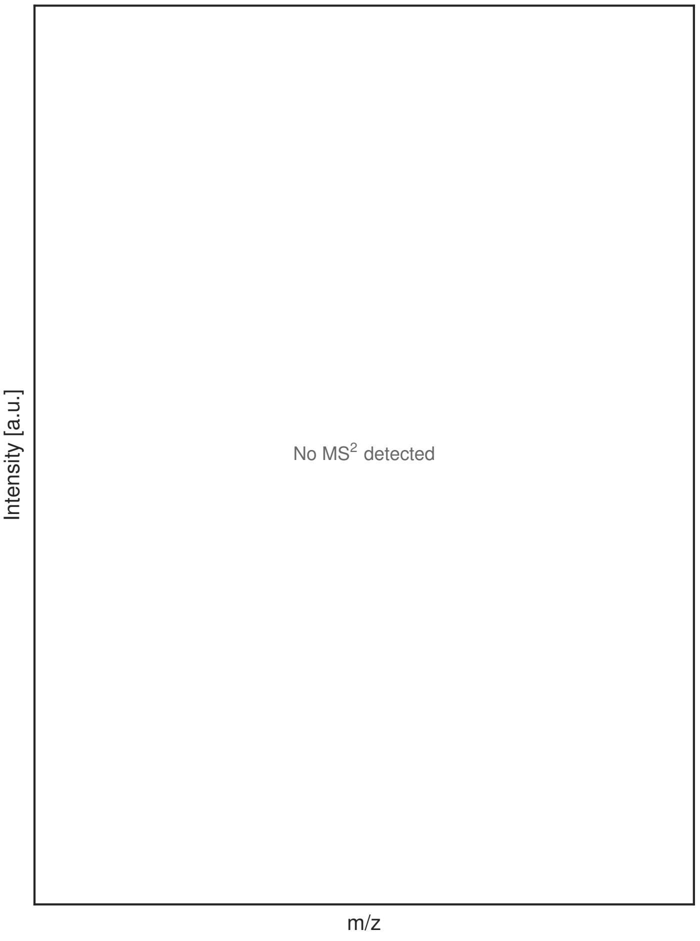


G

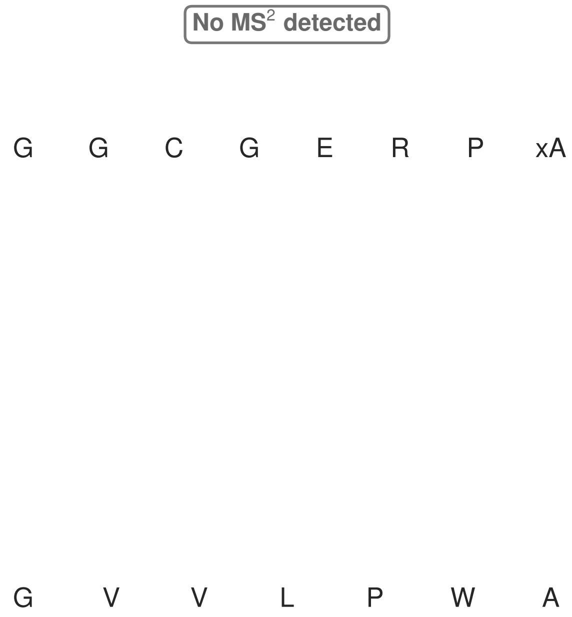


**Figure S137: Modification of the design_2 precursor peptide by PlpXY.** LC-MS analysis of IMAC-purified precursor peptides from *E. coli* BL21(DE3)Tuner containing splicease-encoding plasmid TJ-C-599 (PlpXY) and precursor peptide-encoding plasmid design_2 (RSTN-PlpA3-TEV-designed-2) after expression at 24 °C for 22 h in 1.5 mL cultures after digestion with TEV. Data from n=4 independent biological replicates. Spliced amino acids have an x as a prefix. Spliced refers to a single tyramine excision. A representative experimental replicate is shown in panels **A, B.**

**A** Close-up plot of the representative extracted ion chromatogram (EIC) of the various peptide states; the blue trace corresponds to the Unmodified peptide; the orange trace corresponds to the Spliced peptide. **B** Close-up plot of the m/z values of the peptides detected by the mass spectrometer, with the shaded colored areas corresponding to where the expected m/z values for the peptide m/z traces should occur, with the observed spectrum plotted in black. The Unmodified, Spliced peptide(s) were found in the mass spectrum. **C** Bar plot showing the relative abundance of the various peptide states at charge z=2; the bars represent the average relative abundances normalized to total peak areas, with each individual experimental replicate value plotted as a black dot. **D** Mirrored MS$^2$ plot for the Unmodified peptide. In this mirror plot, the full fragmentation spectrum is plotted in black pointing upward, and the matched fragment b ions (blue) and y ions (red) are plotted pointing downward, mirrored against it, so that each matched fragment can be compared directly with the measured spectrum. **E** MS$^2$ spectrum of the fragments corresponding to the Unmodified peptide sequence, with matched b (blue, above) and y ions (red, below) plotted. **F** Mirrored MS$^2$ plot for the Spliced peptide. In this mirror plot, the full fragmentation spectrum is plotted in black pointing upward, and the matched fragment b ions (blue) and y ions (red) are plotted pointing downward, mirrored against it, so that each matched fragment can be compared directly with the measured spectrum. **G** MS$^2$ spectrum of the fragments corresponding to the Spliced peptide sequence, with matched b (blue, above) and y ions (red, below) plotted.

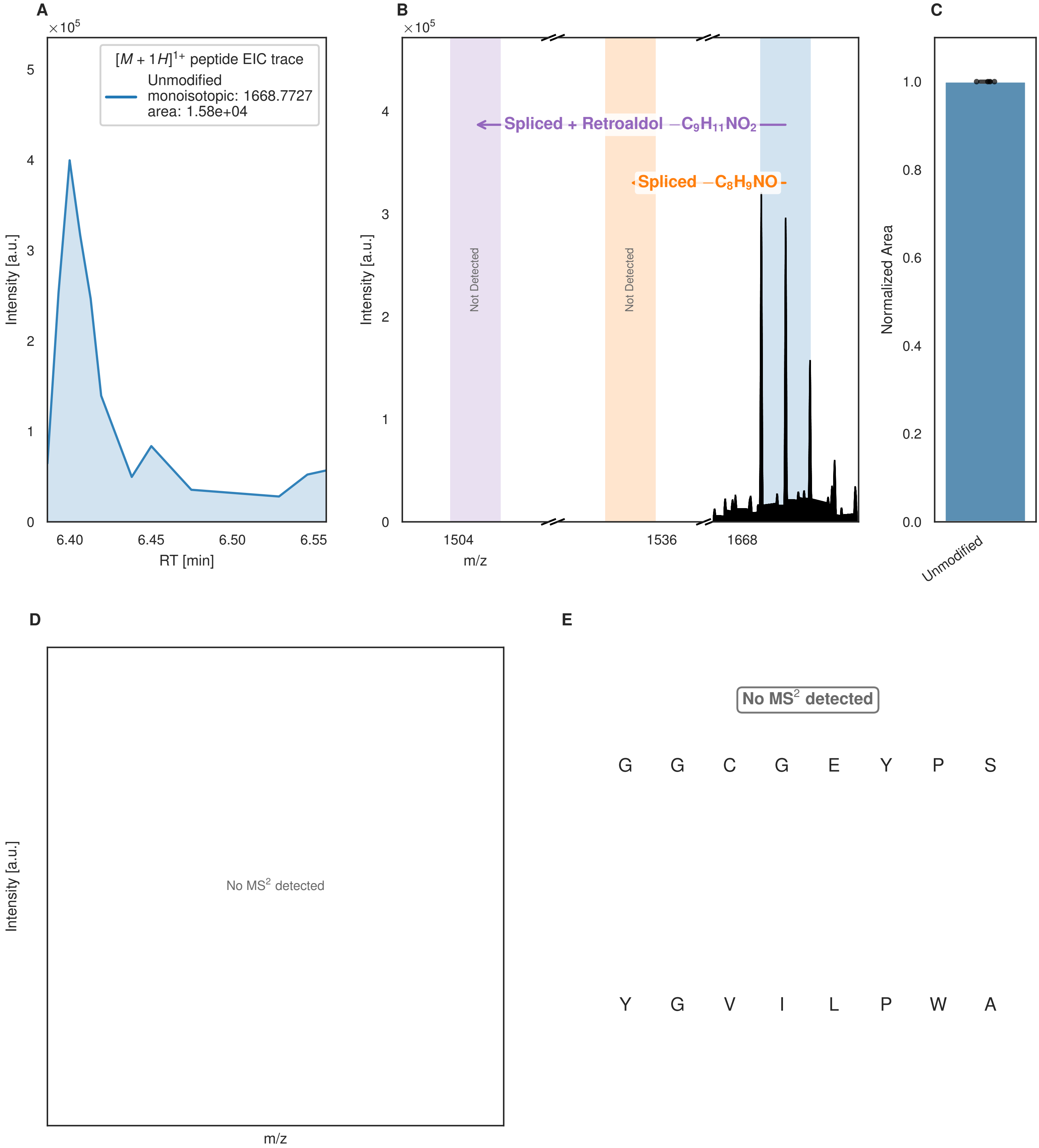


**Figure S138: Modification of the design_3 precursor peptide by PlpXY.** LC-MS analysis of IMAC-purified precursor peptides from *E. coli* BL21(DE3)Tuner containing splicease-encoding plasmid TJ-C-599 (PlpXY) and precursor peptide-encoding plasmid design_3 (RSTN-PlpA3-TEV-designed-3) after expression at 24 °C for 22 h in 1.5 mL cultures after digestion with TEV. Data from n=4 independent biological replicates. Spliced amino acids have an x as a prefix. Spliced refers to a single tyramine excision; Spliced + Retroaldol refers to a glycine-derived ketoamide (xG). A representative experimental replicate is shown in panels **A, B.**
**A** Close-up plot of the representative extracted ion chromatogram (EIC) of the various peptide states; the blue trace corresponds to the Unmodified peptide. **B** Close-up plot of the m/z values of the peptides detected by the mass spectrometer, with the shaded colored areas corresponding to where the expected m/z values for the peptide m/z

(Caption continued on the next page.)

traces should occur, with the observed spectrum plotted in black. The Unmodified peptide(s) were found in the mass spectrum and the Spliced, Spliced + Retroaldol peptide(s) were not detected. **C** Bar plot showing the relative abundance of the various peptide states at charge z=1; the bars represent the average relative abundances normalized to total peak areas, with each individual experimental replicate value plotted as a black dot. **D** Mirrored $MS^2$ plot for the Unmodified peptide. In this mirror plot, the full fragmentation spectrum is plotted in black pointing upward, and the matched fragment b ions (blue) and y ions (red) are plotted pointing downward, mirrored against it, so that each matched fragment can be compared directly with the measured spectrum. **E** $MS^2$ spectrum of the fragments corresponding to the Unmodified peptide sequence, with matched b (blue, above) and y ions (red, below) plotted.

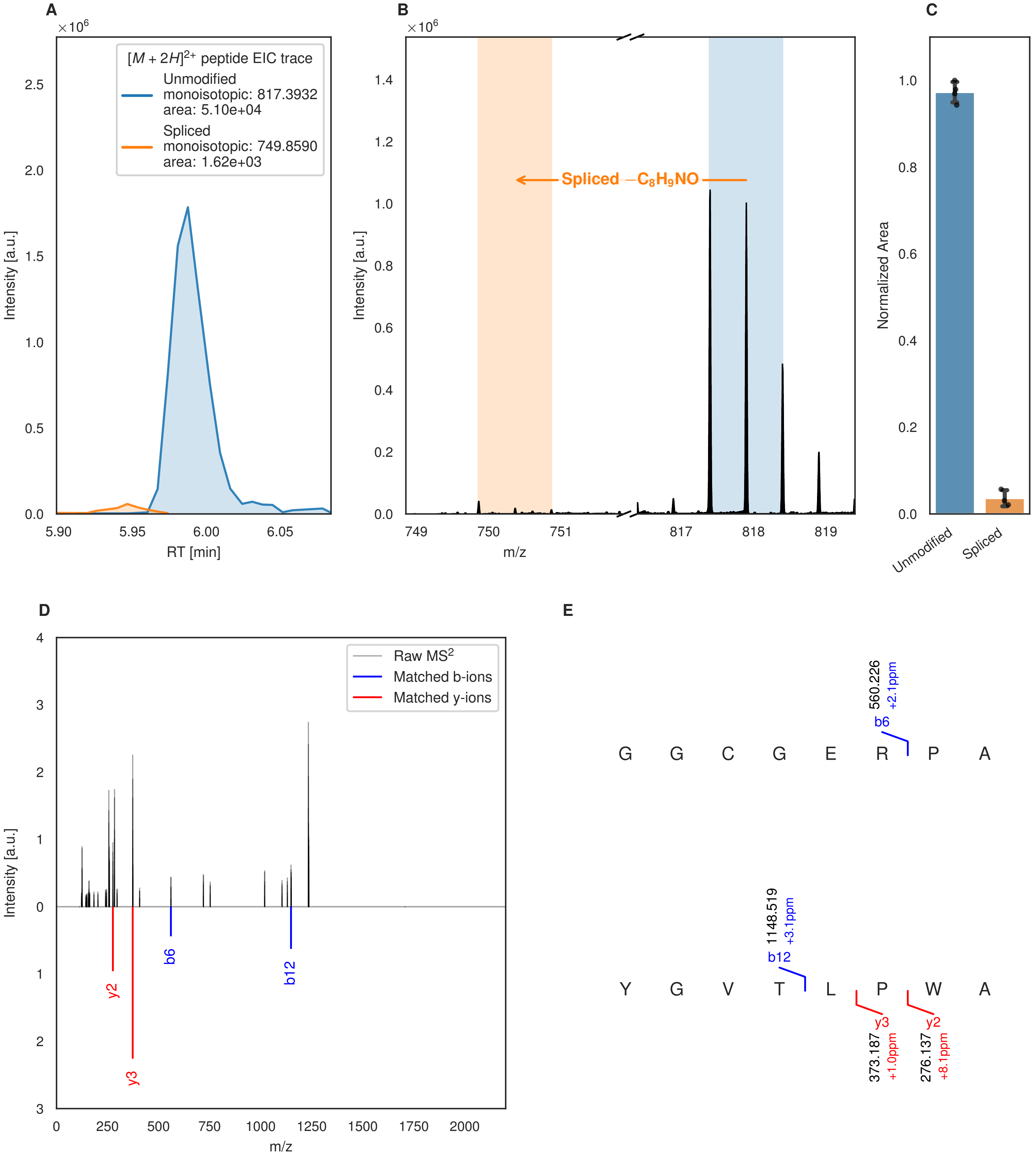

A
[M + 2H]2+ peptide EIC trace
Unmodified
monoisotopic: 817.3932
area: 5.10e+04
Spliced
monoisotopic: 749.8590
area: 1.62e+03
Intensity [a.u.]
RT [min]
B
Spliced −C8H9NO
Intensity [a.u.]
m/z
C
Normalized Area
Unmodified
Spliced
D
Raw MS2
Matched b-ions
Matched y-ions
Intensity [a.u.]
m/z
y2
y3
b6
b12
E
G G C G E R P A
560.226
+2.1ppm
b6
Y G V T L P W A
1148.519
+3.1ppm
b12
y3
373.187
+1.0ppm
y2
276.137
+8.1ppm

Figure S139 (figure continued from the previous page).

F

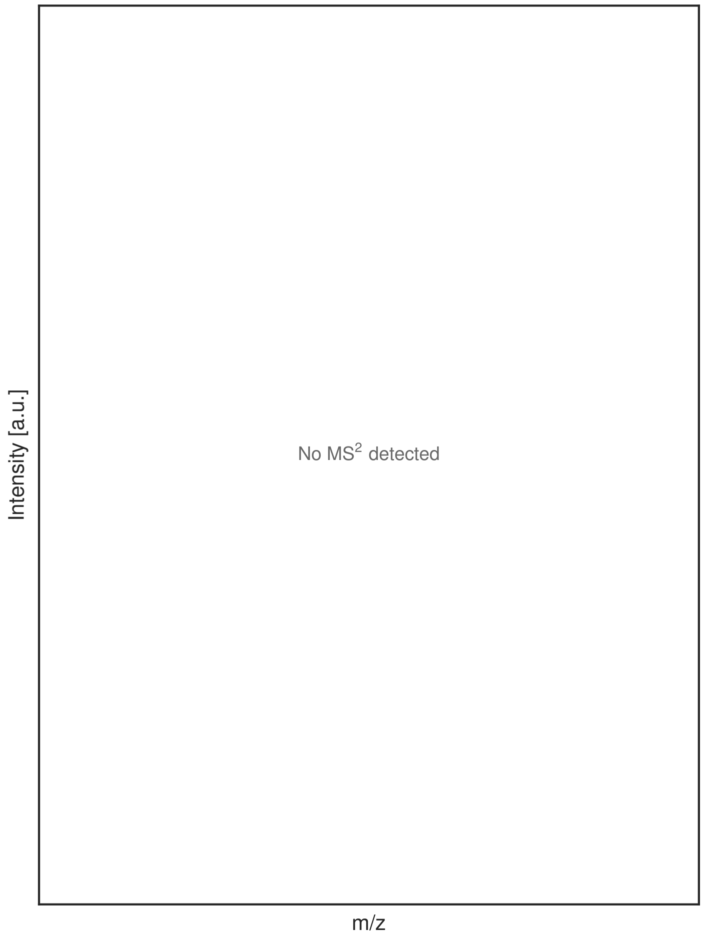


G

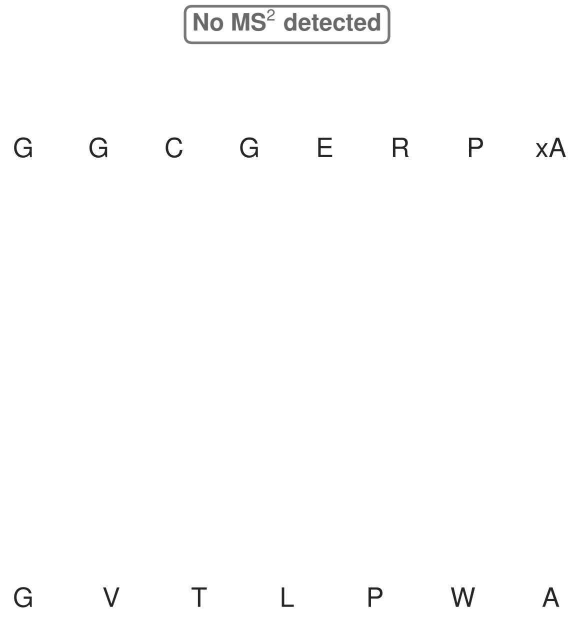


**Figure S139: Modification of the design_4 precursor peptide by PlpXY.** LC-MS analysis of IMAC-purified precursor peptides from *E. coli* BL21(DE3)Tuner containing splicease-encoding plasmid TJ-C-599 (PlpXY) and precursor peptide-encoding plasmid design_4 (RSTN-PlpA3-TEV-designed-4) after expression at 24 °C for 22 h in 1.5 mL cultures after digestion with TEV. Data from n=4 independent biological replicates. Spliced amino acids have an x as a prefix. Spliced refers to a single tyramine excision. A representative experimental replicate is shown in panels **A, B.**
**A** Close-up plot of the representative extracted ion chromatogram (EIC) of the various peptide states; the blue trace corresponds to the Unmodified peptide; the orange trace corresponds to the Spliced peptide. **B** Close-up plot of the m/z values of the peptides detected by the mass spectrometer, with the shaded colored areas corresponding to where the expected m/z values for the peptide m/z traces should occur, with the observed spectrum plotted in black. The Unmodified, Spliced peptide(s) were found in the mass spectrum. **C** Bar plot showing the relative abundance of the various peptide states at charge z=2; the bars represent the average relative abundances normalized to total peak areas, with each individual experimental replicate value plotted as a black dot. **D** Mirrored $MS^2$ plot for the Unmodified peptide. In this mirror plot, the full fragmentation spectrum is plotted in black pointing upward, and the matched fragment b ions (blue) and y ions (red) are plotted pointing downward, mirrored against it, so that each matched fragment can be compared directly with the measured spectrum. **E** $MS^2$ spectrum of the fragments corresponding to the Unmodified peptide sequence, with matched b (blue, above) and y ions (red, below) plotted. **F** Mirrored $MS^2$ plot for the Spliced peptide. In this mirror plot, the full fragmentation spectrum is plotted in black pointing upward, and the matched fragment b ions (blue) and y ions (red) are plotted pointing downward, mirrored against it, so that each matched fragment can be compared directly with the measured spectrum. **G** $MS^2$ spectrum of the fragments corresponding to the Spliced peptide sequence, with matched b (blue, above) and y ions (red, below) plotted.

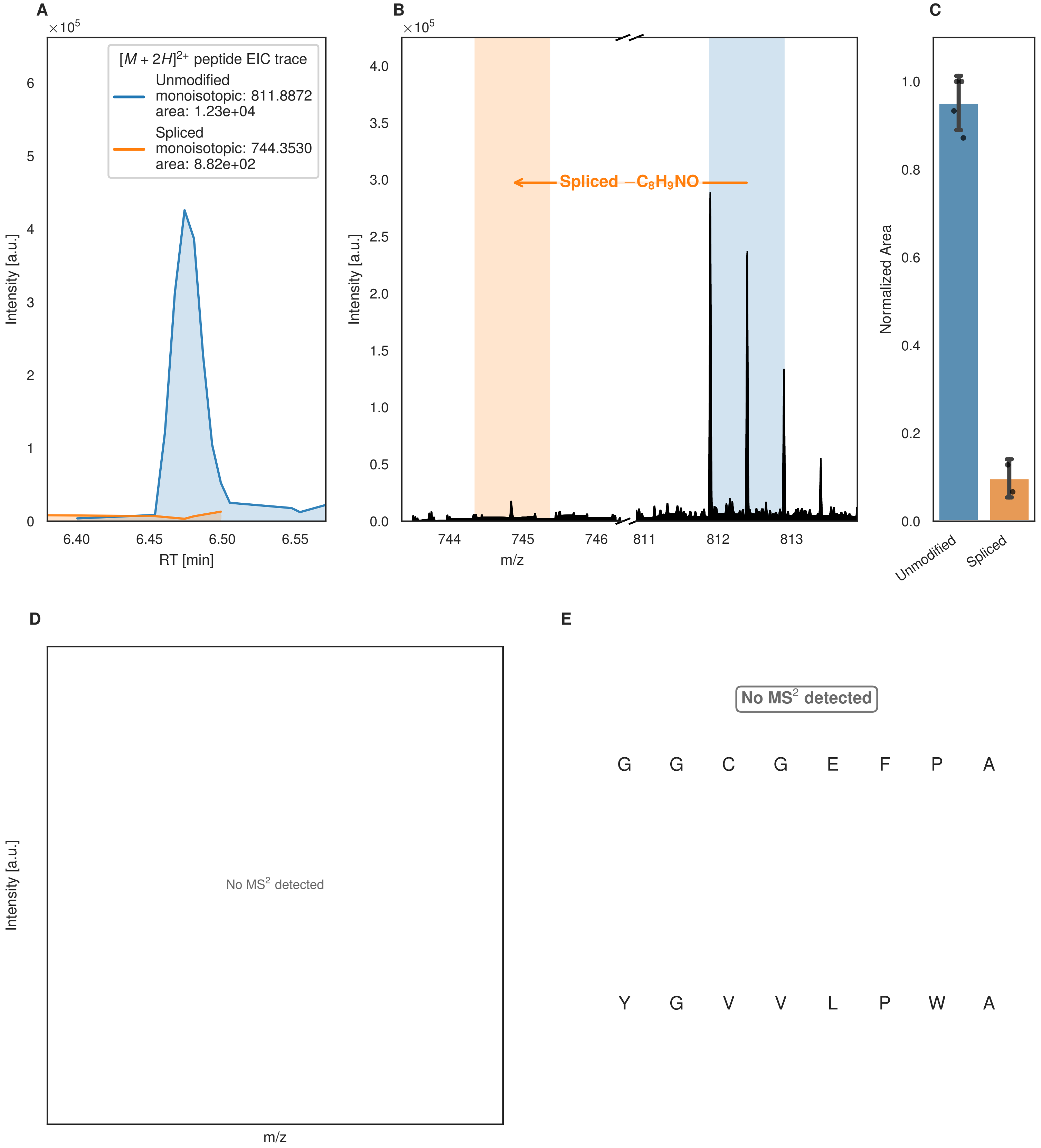
A
B
C
D
E
×10^5
[M + 2H]2+ peptide EIC trace
Unmodified
monoisotopic: 811.8872
area: 1.23e+04
Spliced
monoisotopic: 744.3530
area: 8.82e+02
Intensity [a.u.]
RT [min]
Spliced −C8H9NO
m/z
Normalized Area
Unmodified
Spliced
No MS2 detected
No MS2 detected
G G C G E F P A
Y G V V L P W A

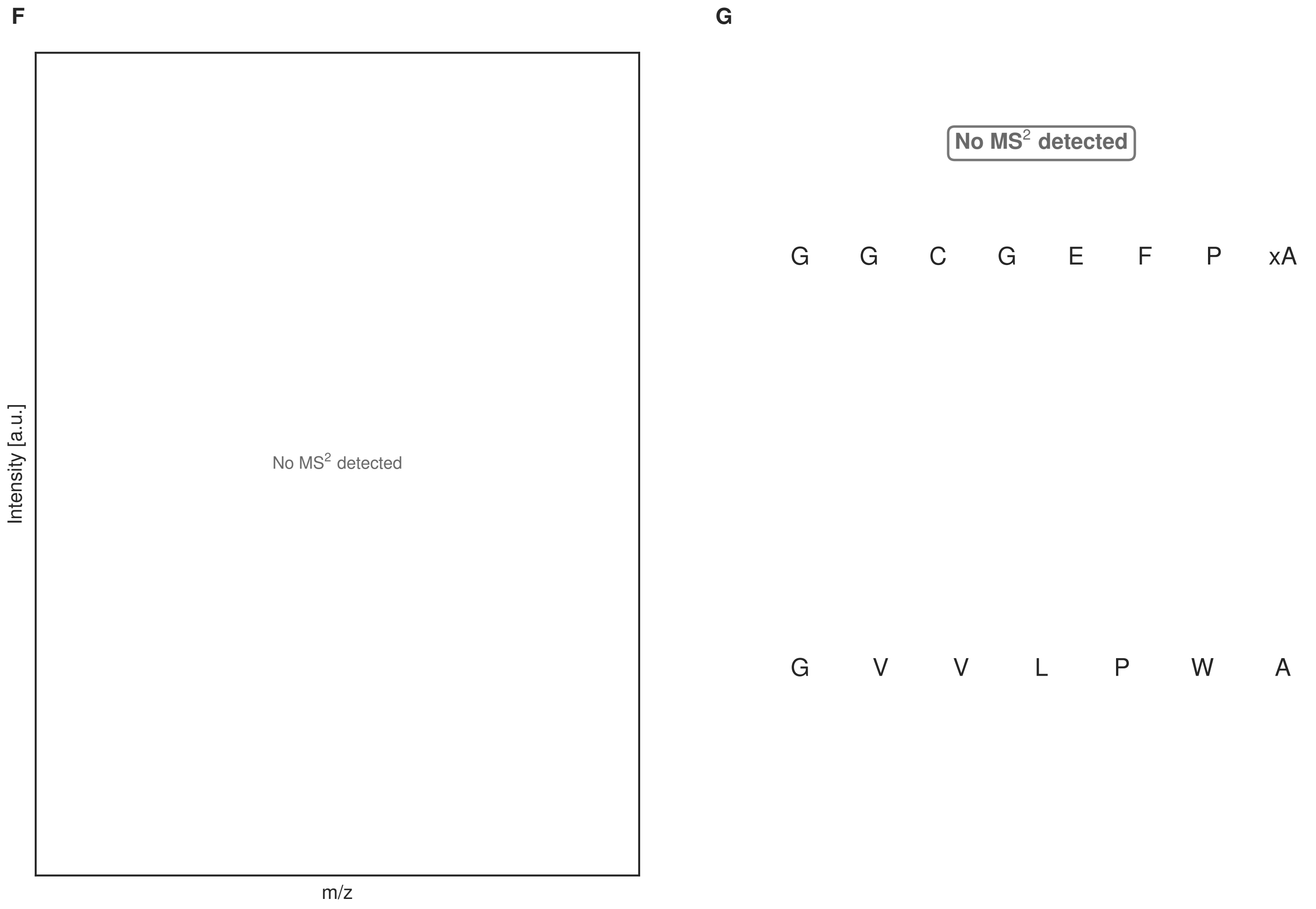


**Figure S140: Modification of the design_5 precursor peptide by PlpXY.** LC-MS analysis of IMAC-purified precursor peptides from *E. coli* BL21(DE3)Tuner containing splicease-encoding plasmid TJ-C-599 (PlpXY) and precursor peptide-encoding plasmid design_5 (RSTN-PlpA3-TEV-designed-5) after expression at 24 °C for 22 h in 1.5 mL cultures after digestion with TEV. Data from n=4 independent biological replicates. Spliced amino acids have an x as a prefix. Spliced refers to a single tyramine excision. A representative experimental replicate is shown in panels **A, B.**

**A** Close-up plot of the representative extracted ion chromatogram (EIC) of the various peptide states; the blue trace corresponds to the Unmodified peptide; the orange trace corresponds to the Spliced peptide. **B** Close-up plot of the m/z values of the peptides detected by the mass spectrometer, with the shaded colored areas corresponding to where the expected m/z values for the peptide m/z traces should occur, with the observed spectrum plotted in black. The Unmodified, Spliced peptide(s) were found in the mass spectrum. **C** Bar plot showing the relative abundance of the various peptide states at charge z=2; the bars represent the average relative abundances normalized to total peak areas, with each individual experimental replicate value plotted as a black dot. **D** Mirrored $MS^2$ plot for the Unmodified peptide. In this mirror plot, the full fragmentation spectrum is plotted in black pointing upward, and the matched fragment b ions (blue) and y ions (red) are plotted pointing downward, mirrored against it, so that each matched fragment can be compared directly with the measured spectrum. **E** $MS^2$ spectrum of the fragments corresponding to the Unmodified peptide sequence, with matched b (blue, above) and y ions (red, below) plotted. **F** Mirrored $MS^2$ plot for the Spliced peptide. In this mirror plot, the full fragmentation spectrum is plotted in black pointing upward, and the matched fragment b ions (blue) and y ions (red) are plotted pointing downward, mirrored against it, so that each matched fragment can be compared directly with the measured spectrum. **G** $MS^2$ spectrum of the fragments corresponding to the Spliced peptide sequence, with matched b (blue, above) and y ions (red, below) plotted.

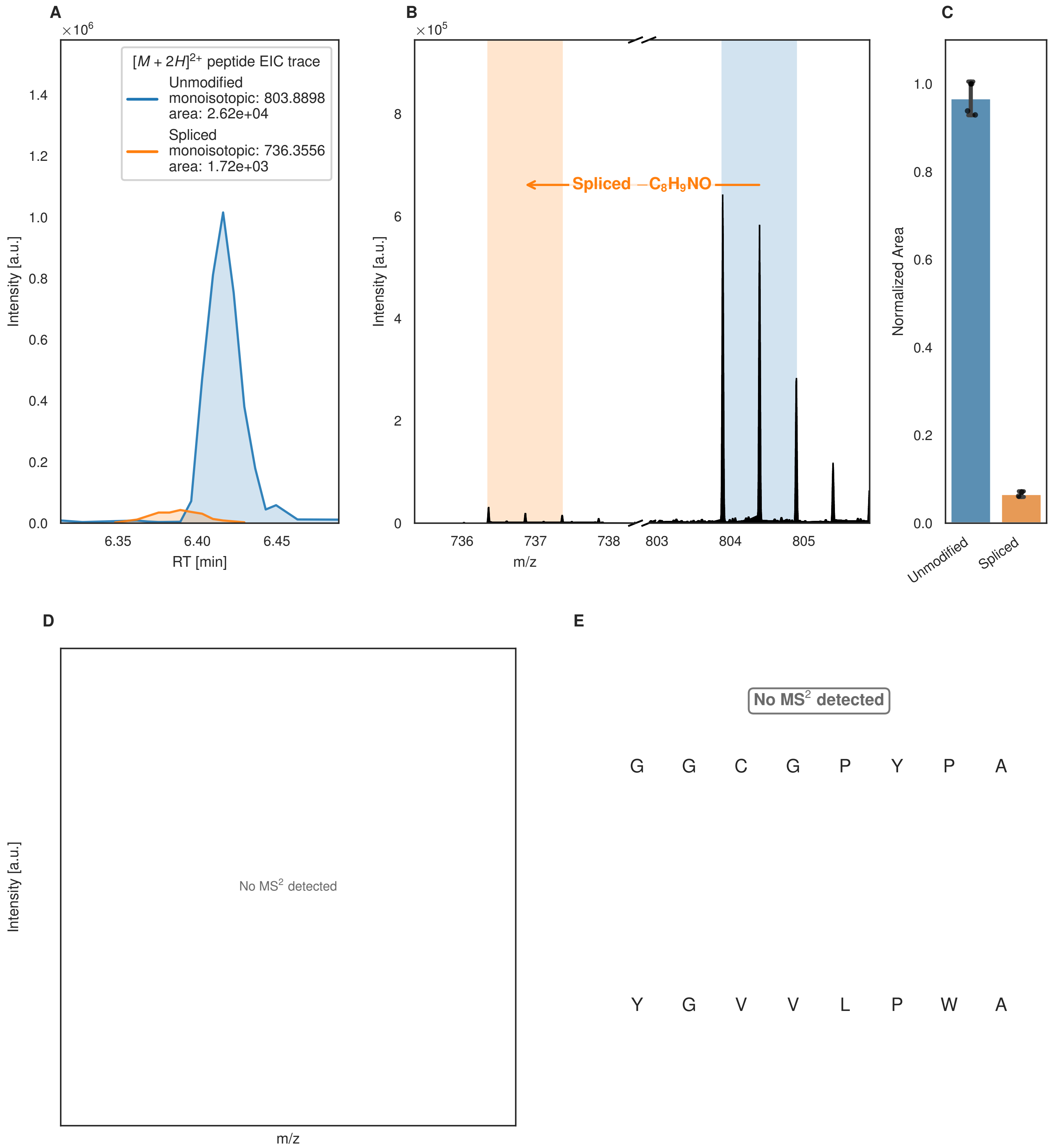
A
B
C
D
E
$\times10^6$
$[M+2H]^{2+}$ peptide EIC trace
Unmodified
monoisotopic: 803.8898
area: 2.62e+04
Spliced
monoisotopic: 736.3556
area: 1.72e+03
Intensity [a.u.]
RT [min]
$\times10^5$
Spliced $-C_8H_9NO$
m/z
Normalized Area
Unmodified
Spliced
No $MS^2$ detected
G G C G P Y P A
Y G V V L P W A

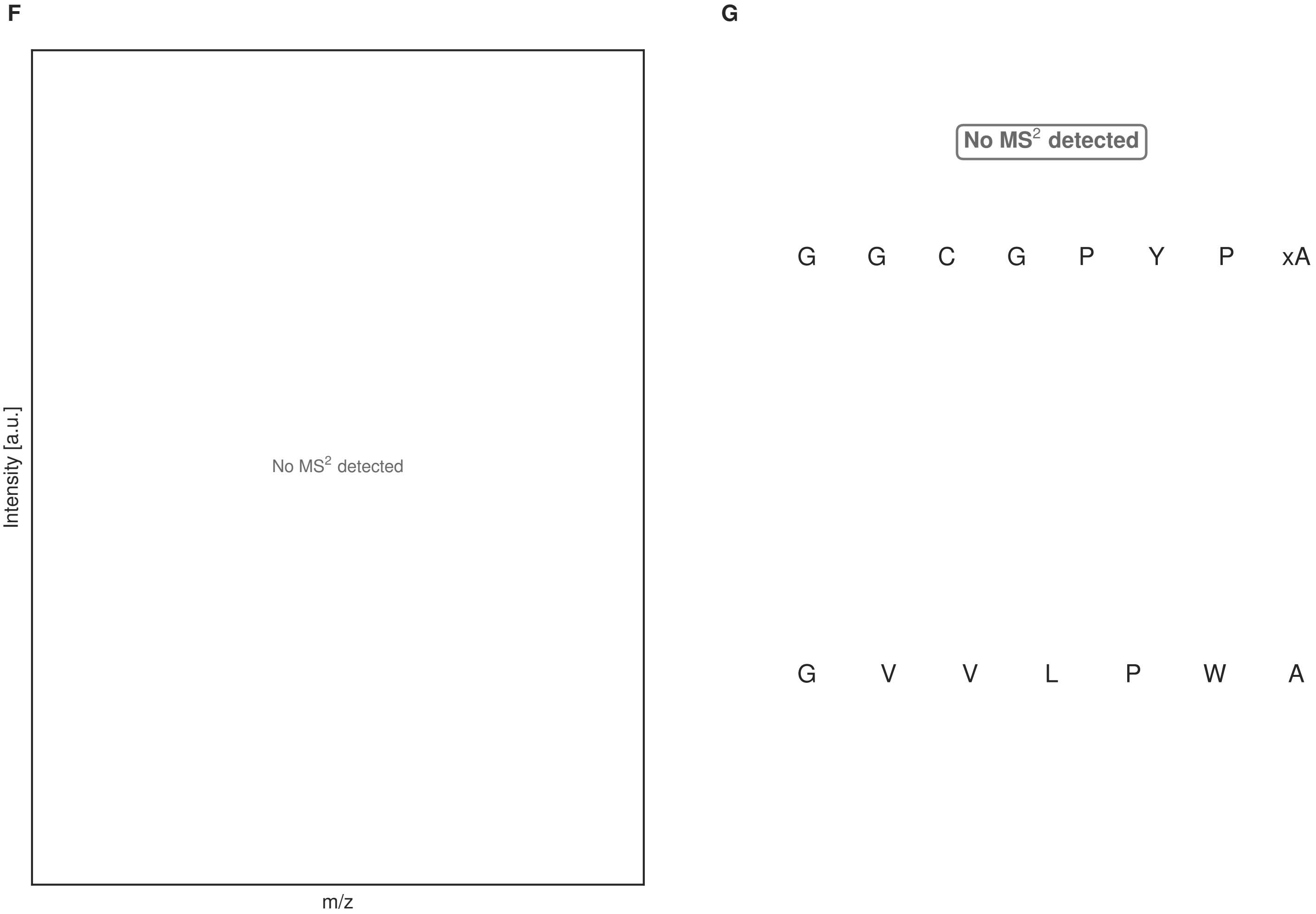


**Figure S141: Modification of the design_6 precursor peptide by PlpXY.** LC-MS analysis of IMAC-purified precursor peptides from *E. coli* BL21(DE3)Tuner containing splicease-encoding plasmid TJ-C-599 (PlpXY) and precursor peptide-encoding plasmid design_6 (RSTN-PlpA3-TEV-designed-6) after expression at 24 °C for 22 h in 1.5 mL cultures after digestion with TEV. Data from n=4 independent biological replicates. Spliced amino acids have an x as a prefix. Spliced refers to a single tyramine excision. A representative experimental replicate is shown in panels **A, B.**

**A** Close-up plot of the representative extracted ion chromatogram (EIC) of the various peptide states; the blue trace corresponds to the Unmodified peptide; the orange trace corresponds to the Spliced peptide. **B** Close-up plot of the m/z values of the peptides detected by the mass spectrometer, with the shaded colored areas corresponding to where the expected m/z values for the peptide m/z traces should occur, with the observed spectrum plotted in black. The Unmodified, Spliced peptide(s) were found in the mass spectrum. **C** Bar plot showing the relative abundance of the various peptide states at charge z=2; the bars represent the average relative abundances normalized to total peak areas, with each individual experimental replicate value plotted as a black dot. **D** Mirrored MS$^2$ plot for the Unmodified peptide. In this mirror plot, the full fragmentation spectrum is plotted in black pointing upward, and the matched fragment b ions (blue) and y ions (red) are plotted pointing downward, mirrored against it, so that each matched fragment can be compared directly with the measured spectrum. **E** MS$^2$ spectrum of the fragments corresponding to the Unmodified peptide sequence, with matched b (blue, above) and y ions (red, below) plotted. **F** Mirrored MS$^2$ plot for the Spliced peptide. In this mirror plot, the full fragmentation spectrum is plotted in black pointing upward, and the matched fragment b ions (blue) and y ions (red) are plotted pointing downward, mirrored against it, so that each matched fragment can be compared directly with the measured spectrum. **G** MS$^2$ spectrum of the fragments corresponding to the Spliced peptide sequence, with matched b (blue, above) and y ions (red, below) plotted.

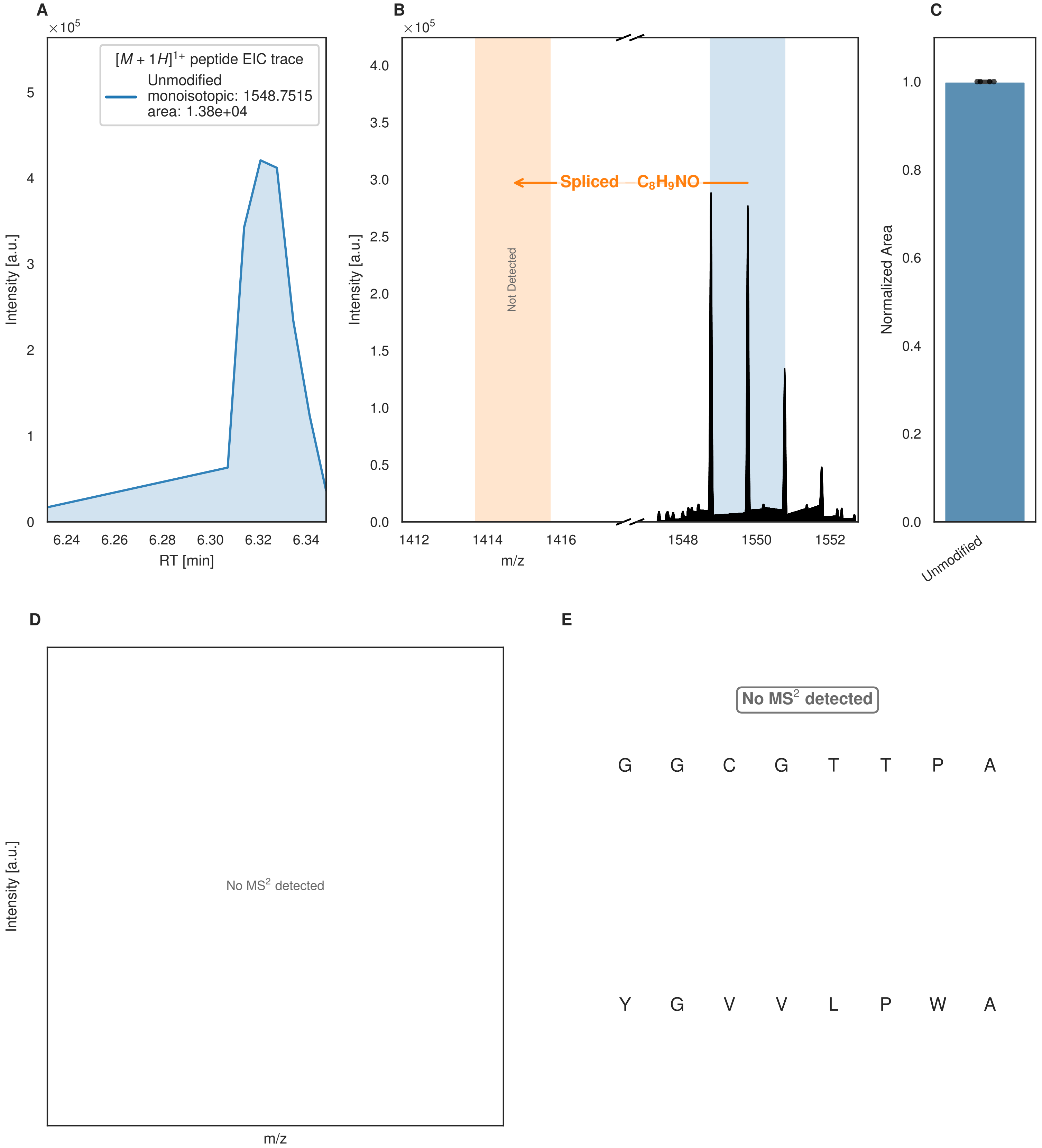


**Figure S142: Modification of the design_7 precursor peptide by PlpXY.** LC-MS analysis of IMAC-purified precursor peptides from *E. coli* BL21(DE3)Tuner containing splicease-encoding plasmid TJ-C-599 (PlpXY) and precursor peptide-encoding plasmid design_7 (RSTN-PlpA3-TEV-designed-7) after expression at 24 °C for 22 h in 1.5 mL cultures after digestion with TEV. Data from n=4 independent biological replicates. Spliced amino acids have an x as a prefix. Spliced refers to a single tyramine excision. A representative experimental replicate is shown in panels **A, B.**

**A** Close-up plot of the representative extracted ion chromatogram (EIC) of the various peptide states; the blue trace corresponds to the Unmodified peptide. **B** Close-up plot of the m/z values of the peptides detected by the mass spectrometer, with the shaded colored areas corresponding to where the expected m/z values for the peptide m/z

(Caption continued on the next page.)

traces should occur, with the observed spectrum plotted in black. The Unmodified peptide(s) were found in the mass spectrum and the Spliced peptide(s) were not detected. **C** Bar plot showing the relative abundance of the various peptide states at charge z=1; the bars represent the average relative abundances normalized to total peak areas, with each individual experimental replicate value plotted as a black dot. **D** Mirrored $MS^2$ plot for the Unmodified peptide. In this mirror plot, the full fragmentation spectrum is plotted in black pointing upward, and the matched fragment b ions (blue) and y ions (red) are plotted pointing downward, mirrored against it, so that each matched fragment can be compared directly with the measured spectrum. **E** $MS^2$ spectrum of the fragments corresponding to the Unmodified peptide sequence, with matched b (blue, above) and y ions (red, below) plotted.

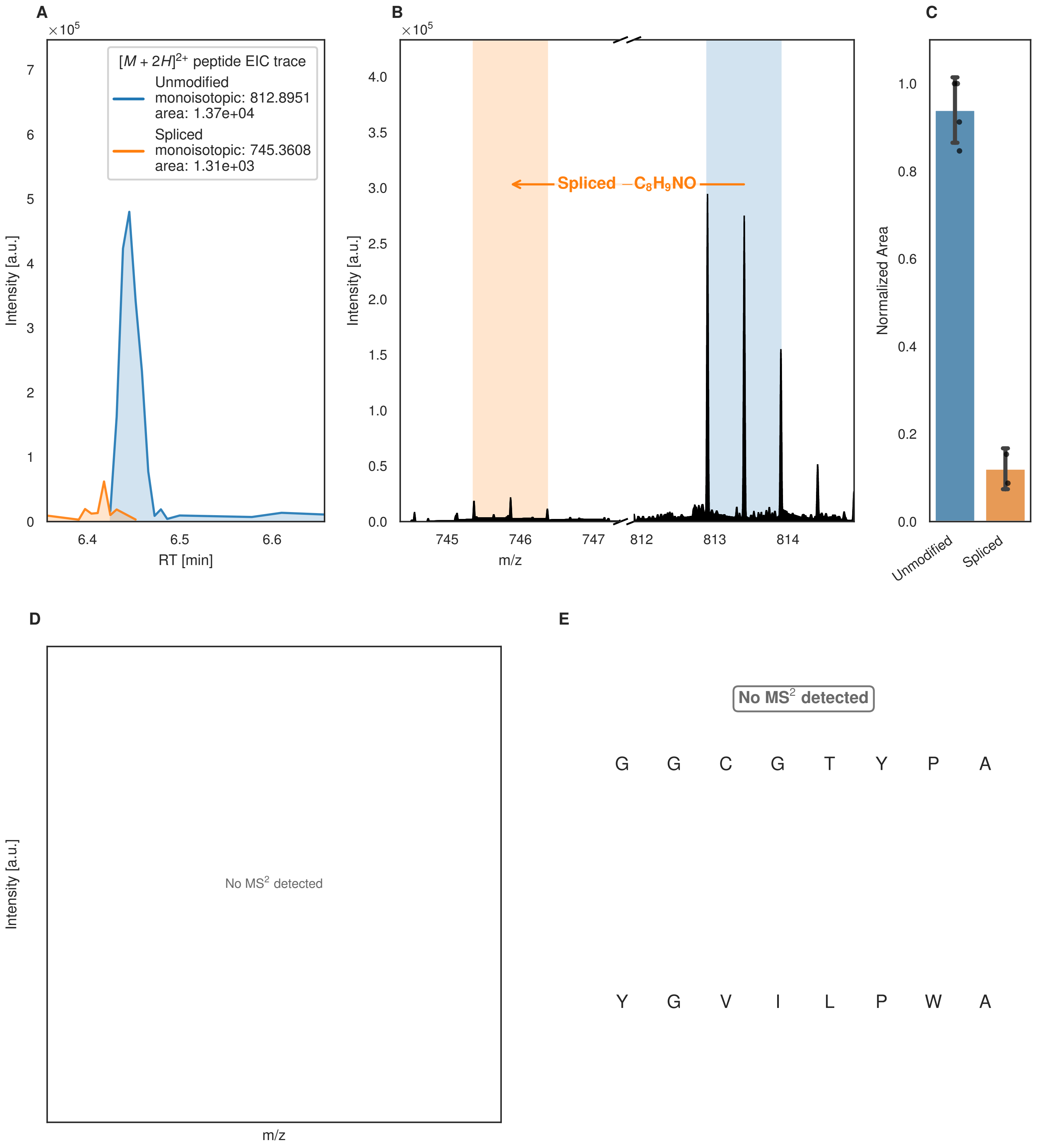
A
×10^5
[M + 2H]2+ peptide EIC trace
Unmodified
monoisotopic: 812.8951
area: 1.37e+04
Spliced
monoisotopic: 745.3608
area: 1.31e+03
Intensity [a.u.]
RT [min]
B
×10^5
Spliced −C8H9NO
Intensity [a.u.]
m/z
C
Normalized Area
Unmodified
Spliced
D
Intensity [a.u.]
No MS2 detected
m/z
E
No MS2 detected
G G C G T Y P A
Y G V I L P W A

Figure S143 (figure continued from the previous page).

F

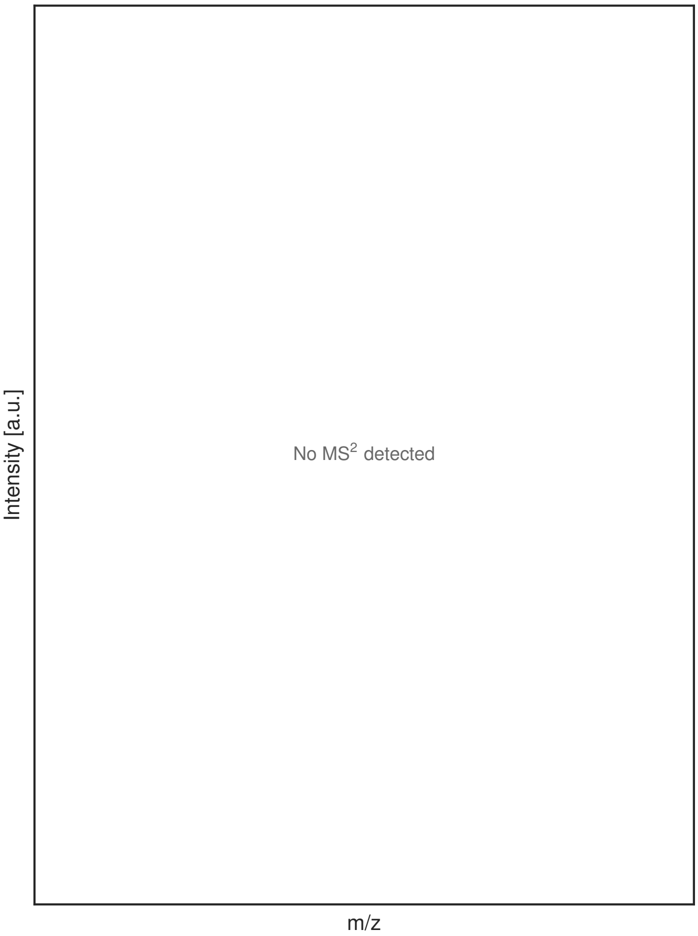


G

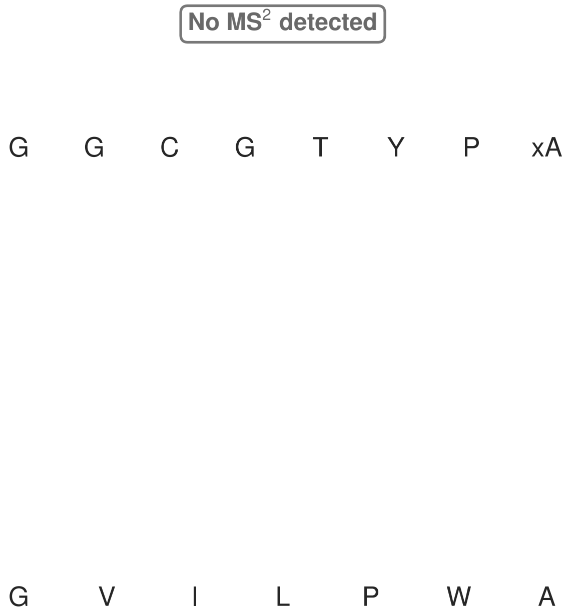


**Figure S143: Modification of the design_8 precursor peptide by PlpXY.** LC-MS analysis of IMAC-purified precursor peptides from *E. coli* BL21(DE3)Tuner containing splicease-encoding plasmid TJ-C-599 (PlpXY) and precursor peptide-encoding plasmid design_8 (RSTN-PlpA3-TEV-designed-8) after expression at 24 °C for 22 h in 1.5 mL cultures after digestion with TEV. Data from n=4 independent biological replicates. Spliced amino acids have an x as a prefix. Spliced refers to a single tyramine excision. A representative experimental replicate is shown in panels **A, B.**

**A** Close-up plot of the representative extracted ion chromatogram (EIC) of the various peptide states; the blue trace corresponds to the Unmodified peptide; the orange trace corresponds to the Spliced peptide. **B** Close-up plot of the m/z values of the peptides detected by the mass spectrometer, with the shaded colored areas corresponding to where the expected m/z values for the peptide m/z traces should occur, with the observed spectrum plotted in black. The Unmodified, Spliced peptide(s) were found in the mass spectrum. **C** Bar plot showing the relative abundance of the various peptide states at charge z=2; the bars represent the average relative abundances normalized to total peak areas, with each individual experimental replicate value plotted as a black dot. **D** Mirrored MS$^2$ plot for the Unmodified peptide. In this mirror plot, the full fragmentation spectrum is plotted in black pointing upward, and the matched fragment b ions (blue) and y ions (red) are plotted pointing downward, mirrored against it, so that each matched fragment can be compared directly with the measured spectrum. **E** MS$^2$ spectrum of the fragments corresponding to the Unmodified peptide sequence, with matched b (blue, above) and y ions (red, below) plotted. **F** Mirrored MS$^2$ plot for the Spliced peptide. In this mirror plot, the full fragmentation spectrum is plotted in black pointing upward, and the matched fragment b ions (blue) and y ions (red) are plotted pointing downward, mirrored against it, so that each matched fragment can be compared directly with the measured spectrum. **G** MS$^2$ spectrum of the fragments corresponding to the Spliced peptide sequence, with matched b (blue, above) and y ions (red, below) plotted.

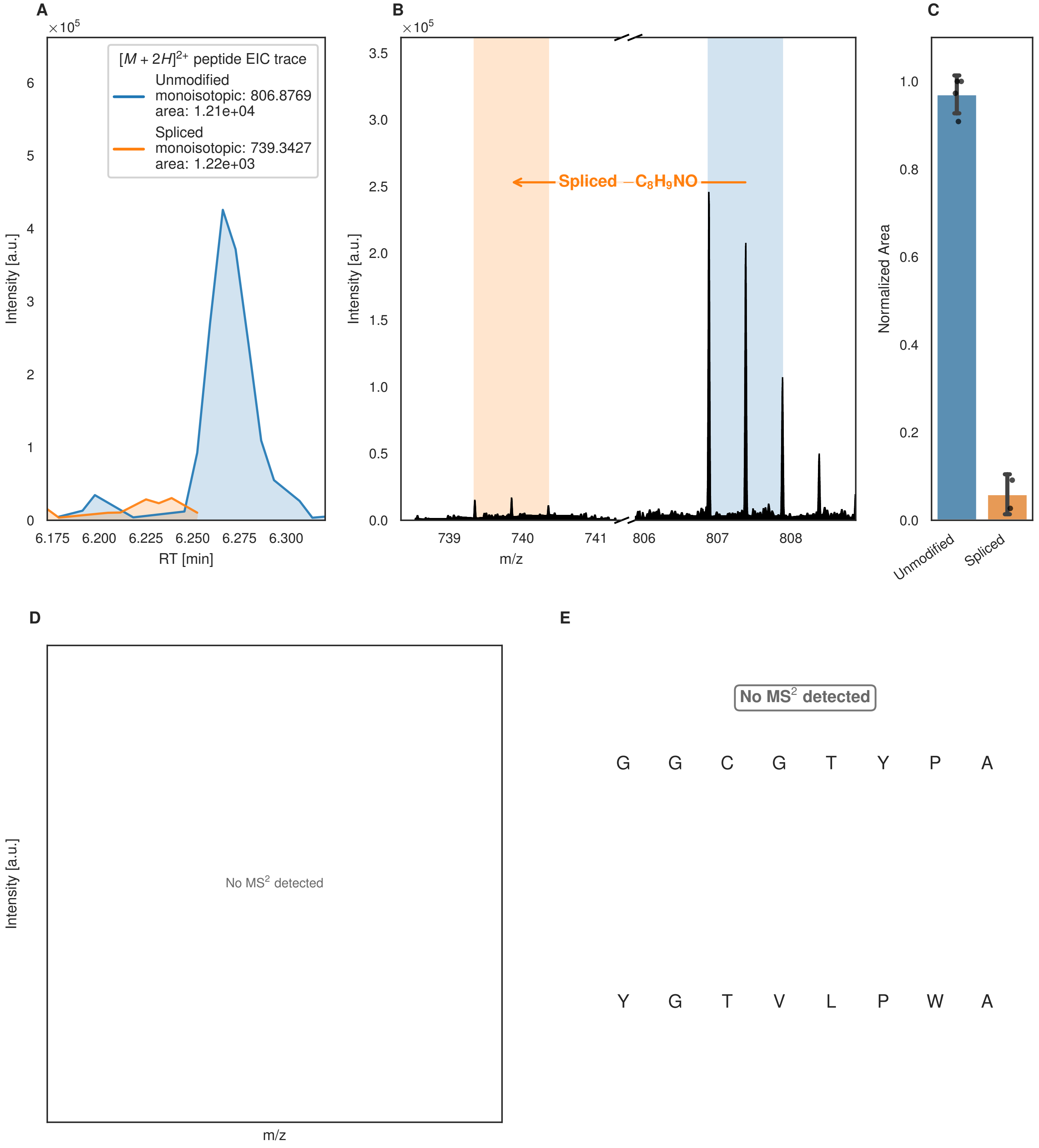

A
×10^5
[M + 2H]2+ peptide EIC trace
Unmodified
monoisotopic: 806.8769
area: 1.21e+04
Spliced
monoisotopic: 739.3427
area: 1.22e+03
Intensity [a.u.]
RT [min]
6.175
6.200
6.225
6.250
6.275
6.300
B
×10^5
Spliced −C8H9NO
Intensity [a.u.]
m/z
739
740
741
806
807
808
C
Normalized Area
Unmodified
Spliced
D
Intensity [a.u.]
No MS2 detected
m/z
E
No MS2 detected
G G C G T Y P A
Y G T V L P W A

Figure S144 (figure continued from the previous page).

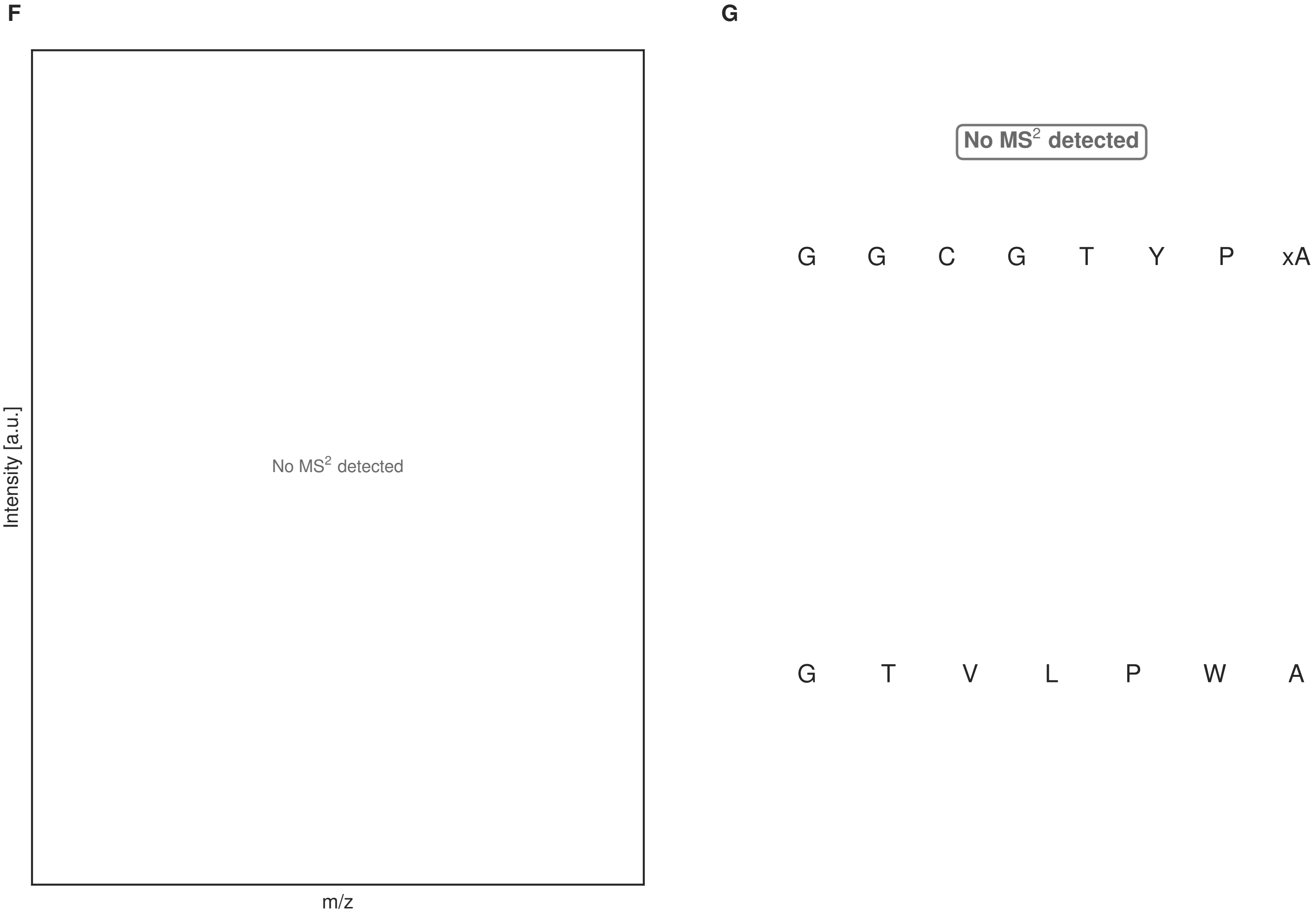


**Figure S144: Modification of the design_10 precursor peptide by PlpXY.** LC-MS analysis of IMAC-purified precursor peptides from *E. coli* BL21(DE3)Tuner containing splicease-encoding plasmid TJ-C-599 (PlpXY) and precursor peptide-encoding plasmid design_10 (RSTN-PlpA3-TEV-designed-10) after expression at 24 °C for 22 h in 1.5 mL cultures after digestion with TEV. Data from n=4 independent biological replicates. Spliced amino acids have an x as a prefix. Spliced refers to a single tyramine excision. A representative experimental replicate is shown in panels **A, B.**

**A** Close-up plot of the representative extracted ion chromatogram (EIC) of the various peptide states; the blue trace corresponds to the Unmodified peptide; the orange trace corresponds to the Spliced peptide. **B** Close-up plot of the m/z values of the peptides detected by the mass spectrometer, with the shaded colored areas corresponding to where the expected m/z values for the peptide m/z traces should occur, with the observed spectrum plotted in black. The Unmodified, Spliced peptide(s) were found in the mass spectrum. **C** Bar plot showing the relative abundance of the various peptide states at charge z=2; the bars represent the average relative abundances normalized to total peak areas, with each individual experimental replicate value plotted as a black dot. **D** Mirrored $MS^2$ plot for the Unmodified peptide. In this mirror plot, the full fragmentation spectrum is plotted in black pointing upward, and the matched fragment b ions (blue) and y ions (red) are plotted pointing downward, mirrored against it, so that each matched fragment can be compared directly with the measured spectrum. **E** $MS^2$ spectrum of the fragments corresponding to the Unmodified peptide sequence, with matched b (blue, above) and y ions (red, below) plotted. **F** Mirrored $MS^2$ plot for the Spliced peptide. In this mirror plot, the full fragmentation spectrum is plotted in black pointing upward, and the matched fragment b ions (blue) and y ions (red) are plotted pointing downward, mirrored against it, so that each matched fragment can be compared directly with the measured spectrum. **G** $MS^2$ spectrum of the fragments corresponding to the Spliced peptide sequence, with matched b (blue, above) and y ions (red, below) plotted.

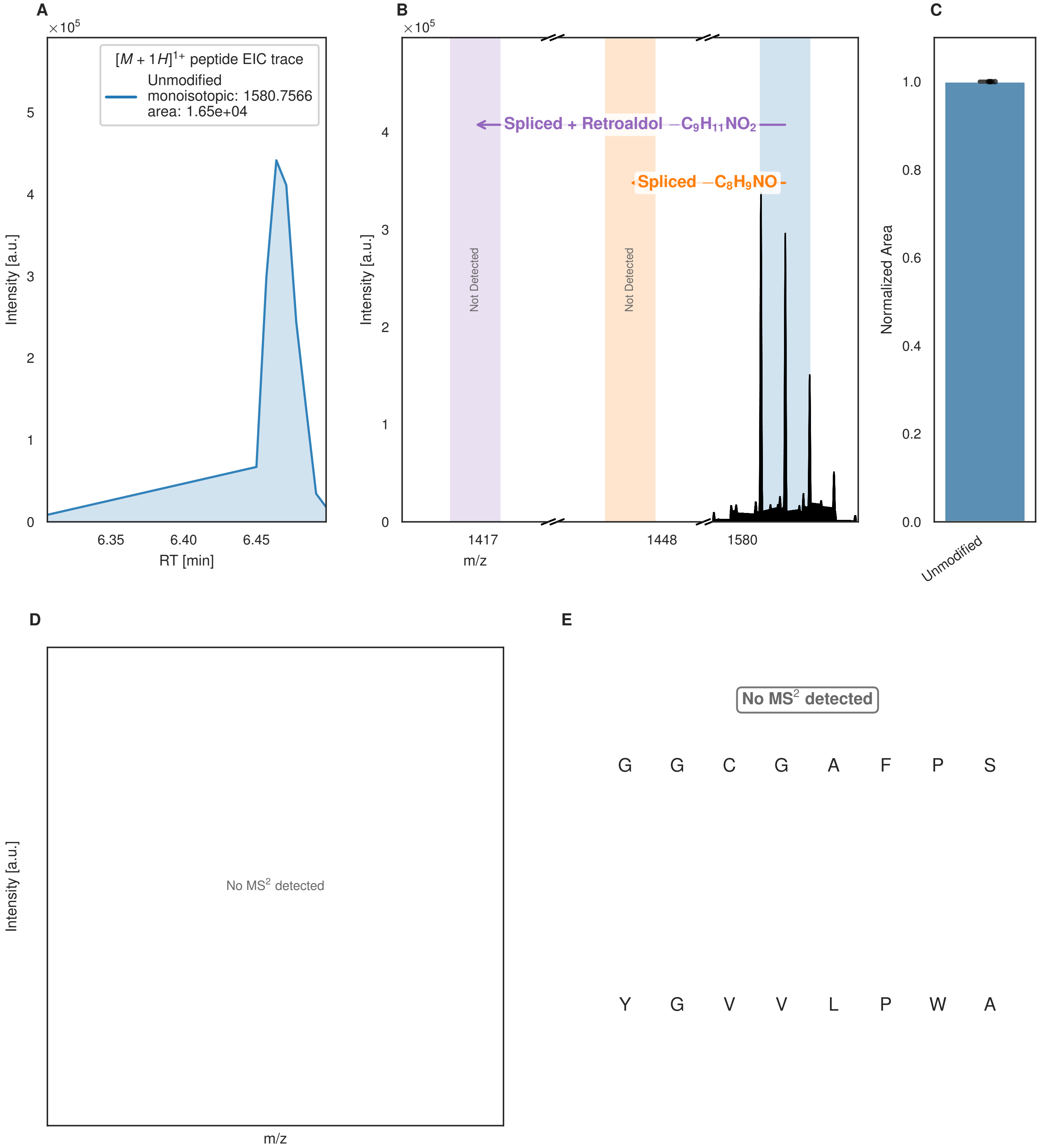


**Figure S145: Modification of the design_11 precursor peptide by PlpXY.** LC-MS analysis of IMAC-purified precursor peptides from *E. coli* BL21(DE3)Tuner containing splicease-encoding plasmid TJ-C-599 (PlpXY) and precursor peptide-encoding plasmid design_11 (RSTN-PlpA3-TEV-designed-11) after expression at 24 °C for 22 h in 1.5 mL cultures after digestion with TEV. Data from n=4 independent biological replicates. Spliced amino acids have an x as a prefix. Spliced refers to a single tyramine excision; Spliced + Retroaldol refers to a glycine-derived ketoamide (xG). A representative experimental replicate is shown in panels **A, B.**

**A** Close-up plot of the representative extracted ion chromatogram (EIC) of the various peptide states; the blue trace corresponds to the Unmodified peptide. **B** Close-up plot of the m/z values of the peptides detected by the mass spectrometer, with the shaded colored areas corresponding to where the expected m/z values for the peptide m/z

(Caption continued on the next page.)

traces should occur, with the observed spectrum plotted in black. The Unmodified peptide(s) were found in the mass spectrum and the Spliced, Spliced + Retroaldol peptide(s) were not detected. **C** Bar plot showing the relative abundance of the various peptide states at charge z=1; the bars represent the average relative abundances normalized to total peak areas, with each individual experimental replicate value plotted as a black dot. **D** Mirrored $MS^2$ plot for the Unmodified peptide. In this mirror plot, the full fragmentation spectrum is plotted in black pointing upward, and the matched fragment b ions (blue) and y ions (red) are plotted pointing downward, mirrored against it, so that each matched fragment can be compared directly with the measured spectrum. **E** $MS^2$ spectrum of the fragments corresponding to the Unmodified peptide sequence, with matched b (blue, above) and y ions (red, below) plotted.

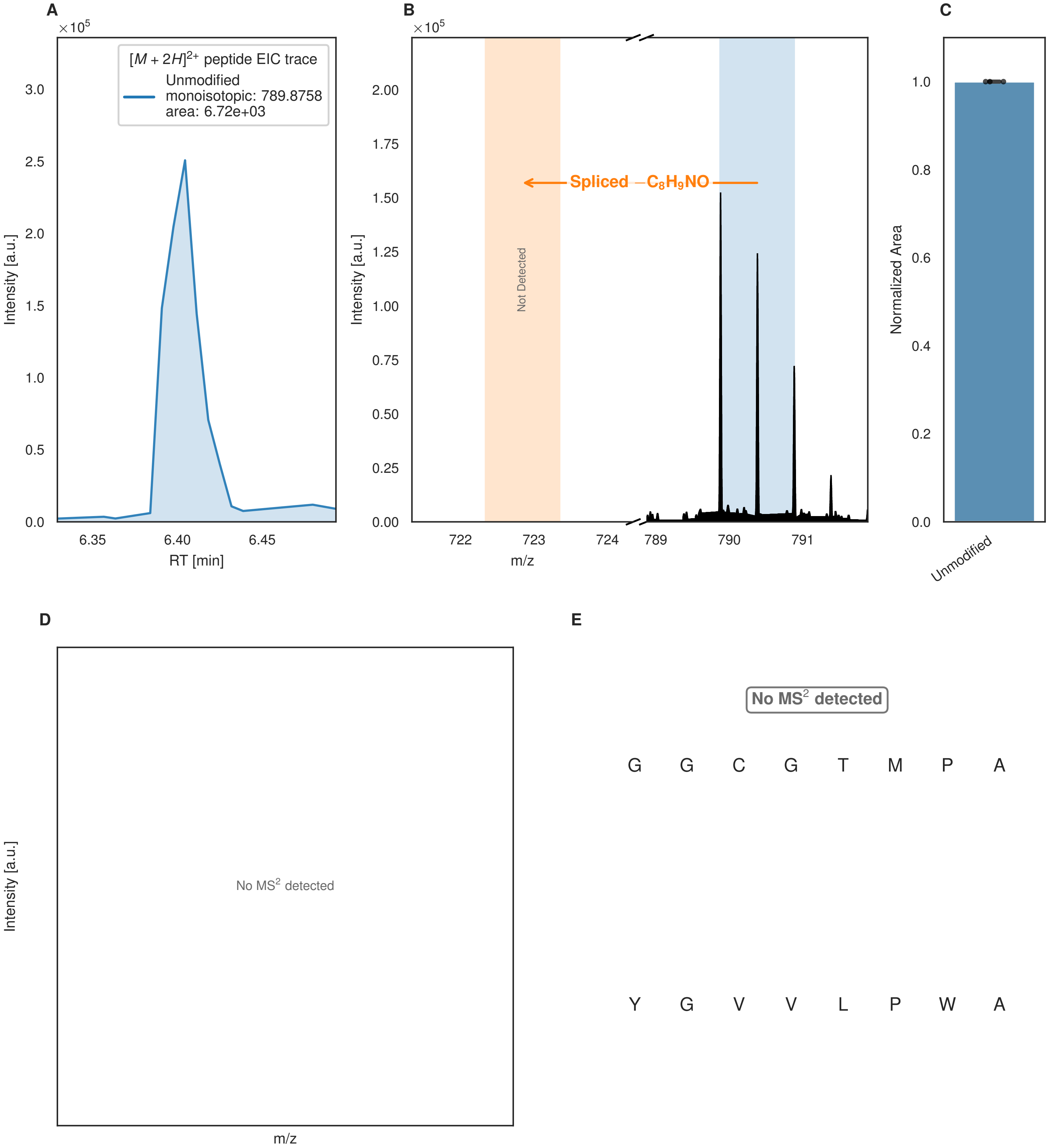


**Figure S146: Modification of the design_12 precursor peptide by PlpXY.** LC-MS analysis of IMAC-purified precursor peptides from *E. coli* BL21(DE3)Tuner containing splicease-encoding plasmid TJ-C-599 (PlpXY) and precursor peptide-encoding plasmid design_12 (RSTN-PlpA3-TEV-designed-12) after expression at 24 °C for 22 h in 1.5 mL cultures after digestion with TEV. Data from n=4 independent biological replicates. Spliced amino acids have an x as a prefix. Spliced refers to a single tyramine excision. A representative experimental replicate is shown in panels **A, B.**

**A** Close-up plot of the representative extracted ion chromatogram (EIC) of the various peptide states; the blue trace corresponds to the Unmodified peptide. **B** Close-up plot of the m/z values of the peptides detected by the mass spectrometer, with the shaded colored areas corresponding to where the expected m/z values for the peptide m/z

(Caption continued on the next page.)

traces should occur, with the observed spectrum plotted in black. The Unmodified peptide(s) were found in the mass spectrum and the Spliced peptide(s) were not detected. **C** Bar plot showing the relative abundance of the various peptide states at charge z=2; the bars represent the average relative abundances normalized to total peak areas, with each individual experimental replicate value plotted as a black dot. **D** Mirrored $MS^2$ plot for the Unmodified peptide. In this mirror plot, the full fragmentation spectrum is plotted in black pointing upward, and the matched fragment b ions (blue) and y ions (red) are plotted pointing downward, mirrored against it, so that each matched fragment can be compared directly with the measured spectrum. **E** $MS^2$ spectrum of the fragments corresponding to the Unmodified peptide sequence, with matched b (blue, above) and y ions (red, below) plotted.

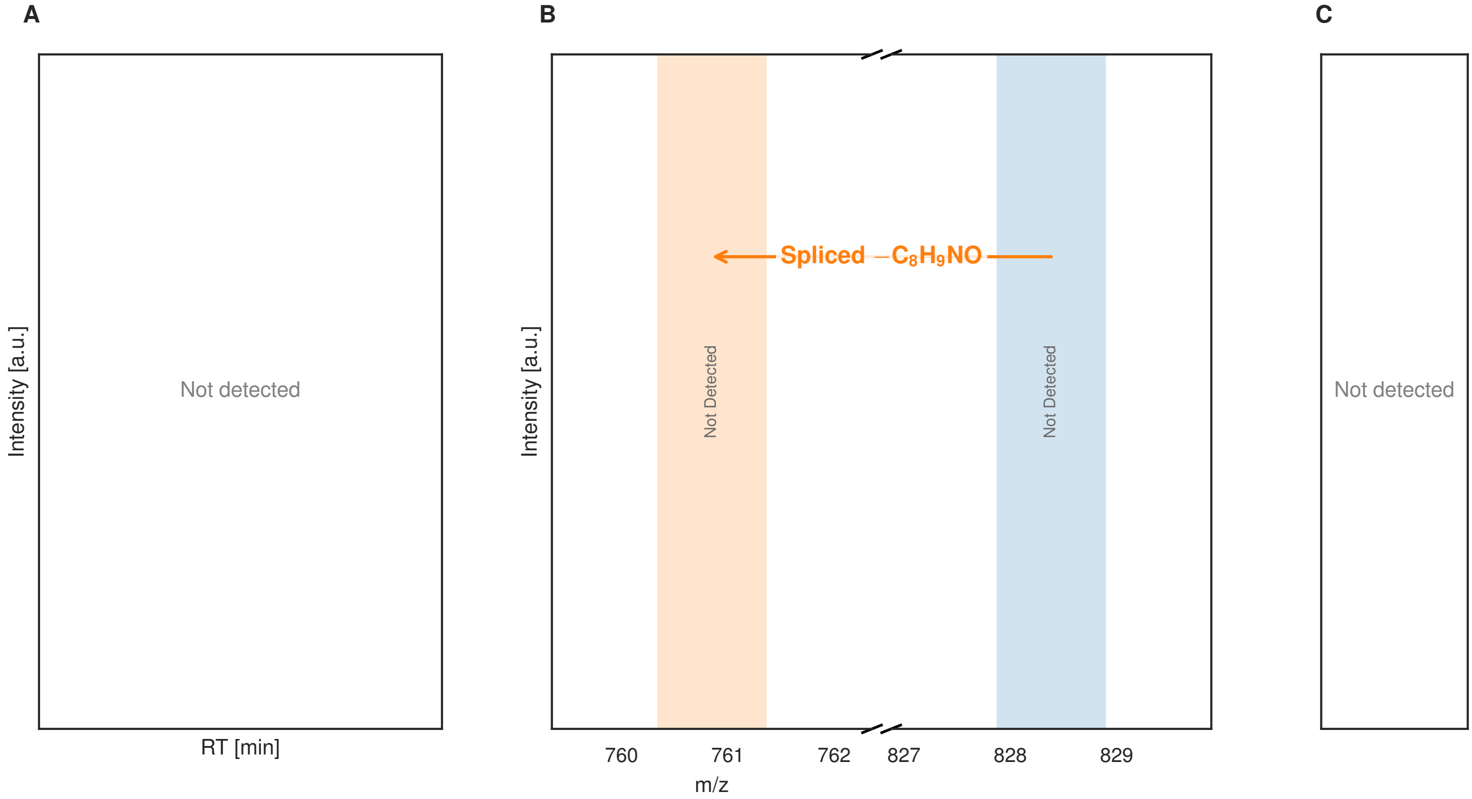


**Figure S147: Modification of the design_13 precursor peptide by PlpXY.** LC-MS analysis of IMAC-purified precursor peptides from *E. coli* BL21(DE3)Tuner containing splicease-encoding plasmid TJ-C-599 (PlpXY) and precursor peptide-encoding plasmid design_13 (RSTN-PlpA3-TEV-designed-13) after expression at 24 °C for 22 h in 1.5 mL cultures after digestion with TEV. Data from n=4 independent biological replicates. Spliced amino acids have an x as a prefix. Spliced refers to a single tyramine excision. A representative experimental replicate is shown in panels **A, B.**

**A** Close-up plot of the representative extracted ion chromatogram (EIC) of the various peptide states; no peptide state was detected. **B** Close-up plot of the m/z values of the peptides, with the shaded colored areas corresponding to where the expected m/z values for the peptide m/z traces should occur. The Unmodified, Spliced peptide(s) were not detected. **C** Bar plot of the relative abundance of the various peptide states at charge z=2; no peptide state was detected, so no abundances are shown.

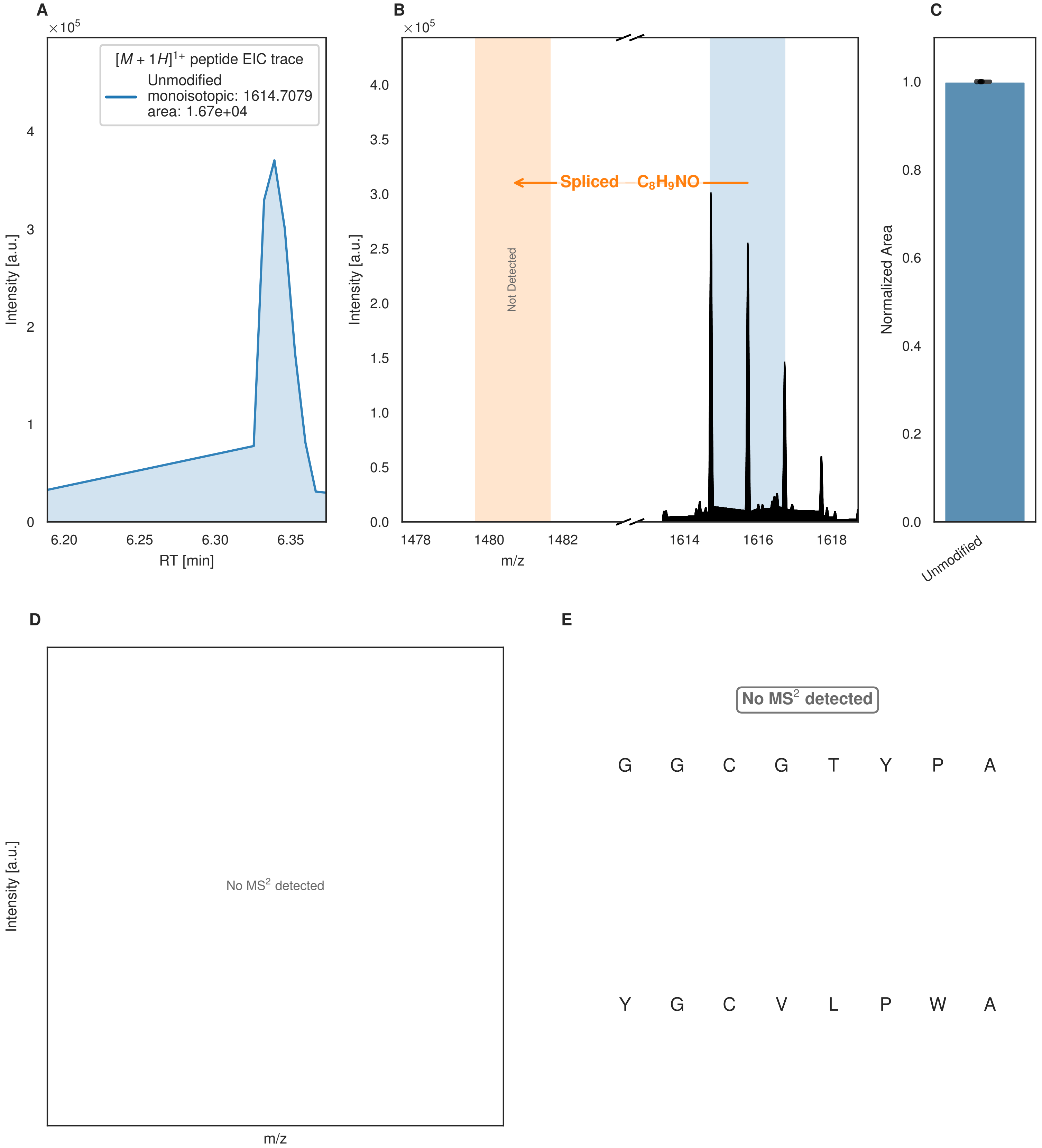


**Figure S148: Modification of the design_14 precursor peptide by PlpXY.** LC-MS analysis of IMAC-purified precursor peptides from *E. coli* BL21(DE3)Tuner containing splicease-encoding plasmid TJ-C-599 (PlpXY) and precursor peptide-encoding plasmid design_14 (RSTN-PlpA3-TEV-designed-14) after expression at 24 °C for 22 h in 1.5 mL cultures after digestion with TEV. Data from n=4 independent biological replicates. Spliced amino acids have an x as a prefix. Spliced refers to a single tyramine excision. A representative experimental replicate is shown in panels **A, B.**

**A** Close-up plot of the representative extracted ion chromatogram (EIC) of the various peptide states; the blue trace corresponds to the Unmodified peptide. **B** Close-up plot of the m/z values of the peptides detected by the mass spectrometer, with the shaded colored areas corresponding to where the expected m/z values for the peptide m/z

(Caption continued on the next page.)

traces should occur, with the observed spectrum plotted in black. The Unmodified peptide(s) were found in the mass spectrum and the Spliced peptide(s) were not detected. **C** Bar plot showing the relative abundance of the various peptide states at charge z=1; the bars represent the average relative abundances normalized to total peak areas, with each individual experimental replicate value plotted as a black dot. **D** Mirrored $MS^2$ plot for the Unmodified peptide. In this mirror plot, the full fragmentation spectrum is plotted in black pointing upward, and the matched fragment b ions (blue) and y ions (red) are plotted pointing downward, mirrored against it, so that each matched fragment can be compared directly with the measured spectrum. **E** $MS^2$ spectrum of the fragments corresponding to the Unmodified peptide sequence, with matched b (blue, above) and y ions (red, below) plotted.

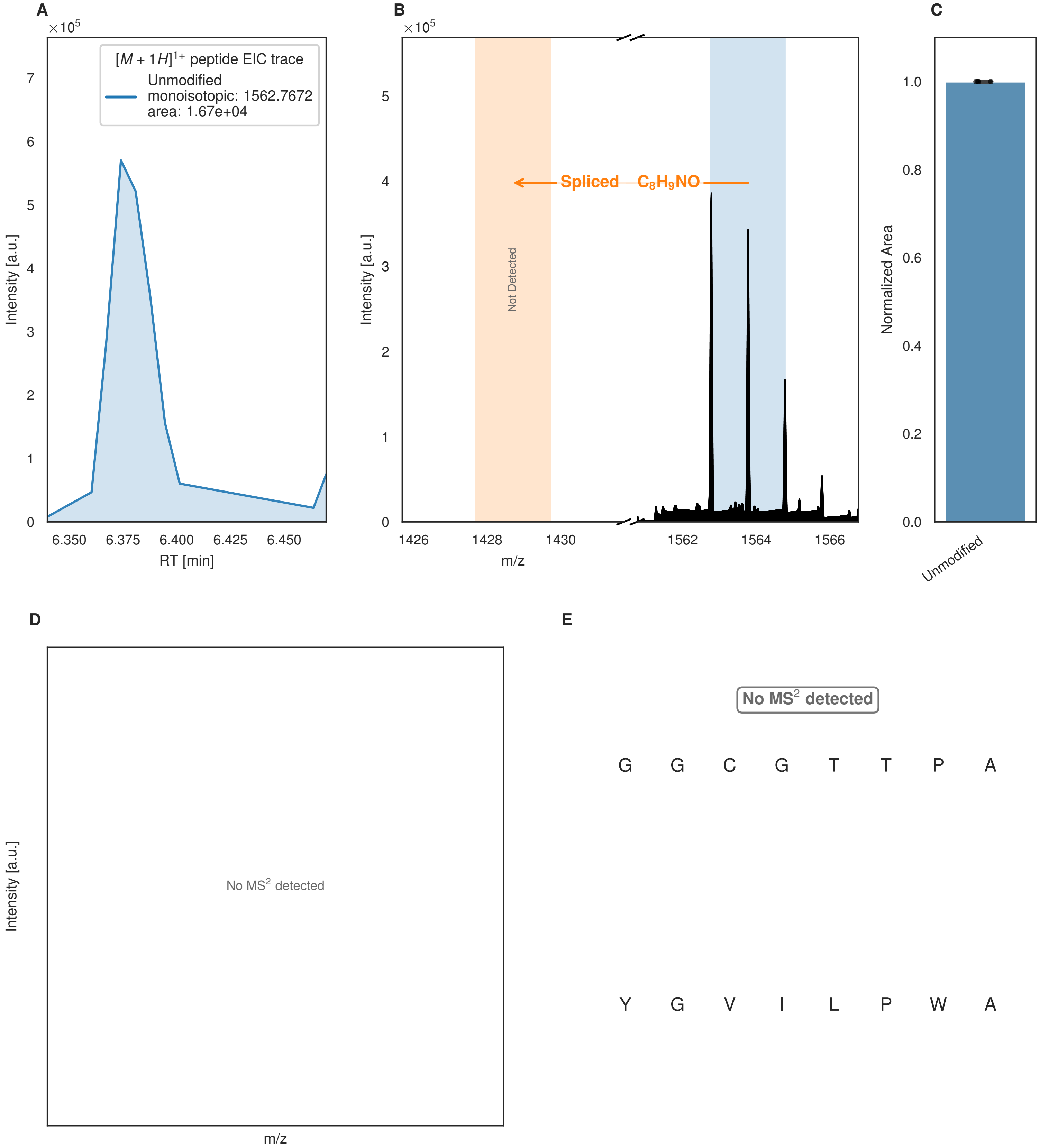


**Figure S149: Modification of the design_15 precursor peptide by PlpXY.** LC-MS analysis of IMAC-purified precursor peptides from *E. coli* BL21(DE3)Tuner containing splicease-encoding plasmid TJ-C-599 (PlpXY) and precursor peptide-encoding plasmid design_15 (RSTN-PlpA3-TEV-designed-15) after expression at 24 °C for 22 h in 1.5 mL cultures after digestion with TEV. Data from n=4 independent biological replicates. Spliced amino acids have an x as a prefix. Spliced refers to a single tyramine excision. A representative experimental replicate is shown in panels **A, B.**

**A** Close-up plot of the representative extracted ion chromatogram (EIC) of the various peptide states; the blue trace corresponds to the Unmodified peptide. **B** Close-up plot of the m/z values of the peptides detected by the mass spectrometer, with the shaded colored areas corresponding to where the expected m/z values for the peptide m/z

(Caption continued on the next page.)

traces should occur, with the observed spectrum plotted in black. The Unmodified peptide(s) were found in the mass spectrum and the Spliced peptide(s) were not detected. **C** Bar plot showing the relative abundance of the various peptide states at charge z=1; the bars represent the average relative abundances normalized to total peak areas, with each individual experimental replicate value plotted as a black dot. **D** Mirrored $MS^2$ plot for the Unmodified peptide. In this mirror plot, the full fragmentation spectrum is plotted in black pointing upward, and the matched fragment b ions (blue) and y ions (red) are plotted pointing downward, mirrored against it, so that each matched fragment can be compared directly with the measured spectrum. **E** $MS^2$ spectrum of the fragments corresponding to the Unmodified peptide sequence, with matched b (blue, above) and y ions (red, below) plotted.

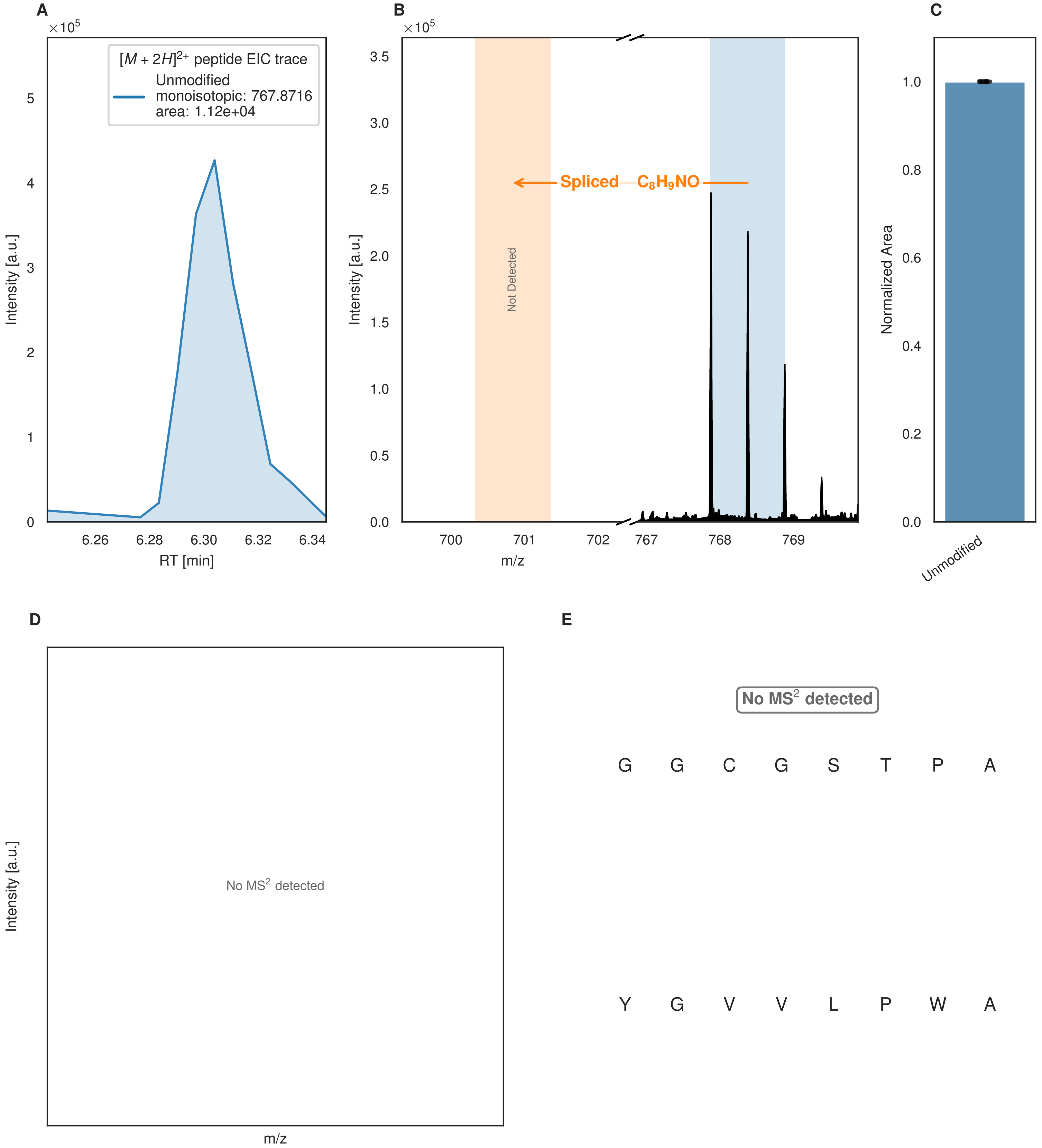


**Figure S150: Modification of the design_16 precursor peptide by PlpXY.** LC-MS analysis of IMAC-purified precursor peptides from *E. coli* BL21(DE3)Tuner containing splicease-encoding plasmid TJ-C-599 (PlpXY) and precursor peptide-encoding plasmid design_16 (RSTN-PlpA3-TEV-designed-16) after expression at 24 °C for 22 h in 1.5 mL cultures after digestion with TEV. Data from n=4 independent biological replicates. Spliced amino acids have an x as a prefix. Spliced refers to a single tyramine excision. A representative experimental replicate is shown in panels **A, B.**

**A** Close-up plot of the representative extracted ion chromatogram (EIC) of the various peptide states; the blue trace corresponds to the Unmodified peptide. **B** Close-up plot of the m/z values of the peptides detected by the mass spectrometer, with the shaded colored areas corresponding to where the expected m/z values for the peptide m/z

(Caption continued on the next page.)

traces should occur, with the observed spectrum plotted in black. The Unmodified peptide(s) were found in the mass spectrum and the Spliced peptide(s) were not detected. **C** Bar plot showing the relative abundance of the various peptide states at charge z=2; the bars represent the average relative abundances normalized to total peak areas, with each individual experimental replicate value plotted as a black dot. **D** Mirrored $MS^2$ plot for the Unmodified peptide. In this mirror plot, the full fragmentation spectrum is plotted in black pointing upward, and the matched fragment b ions (blue) and y ions (red) are plotted pointing downward, mirrored against it, so that each matched fragment can be compared directly with the measured spectrum. **E** $MS^2$ spectrum of the fragments corresponding to the Unmodified peptide sequence, with matched b (blue, above) and y ions (red, below) plotted.

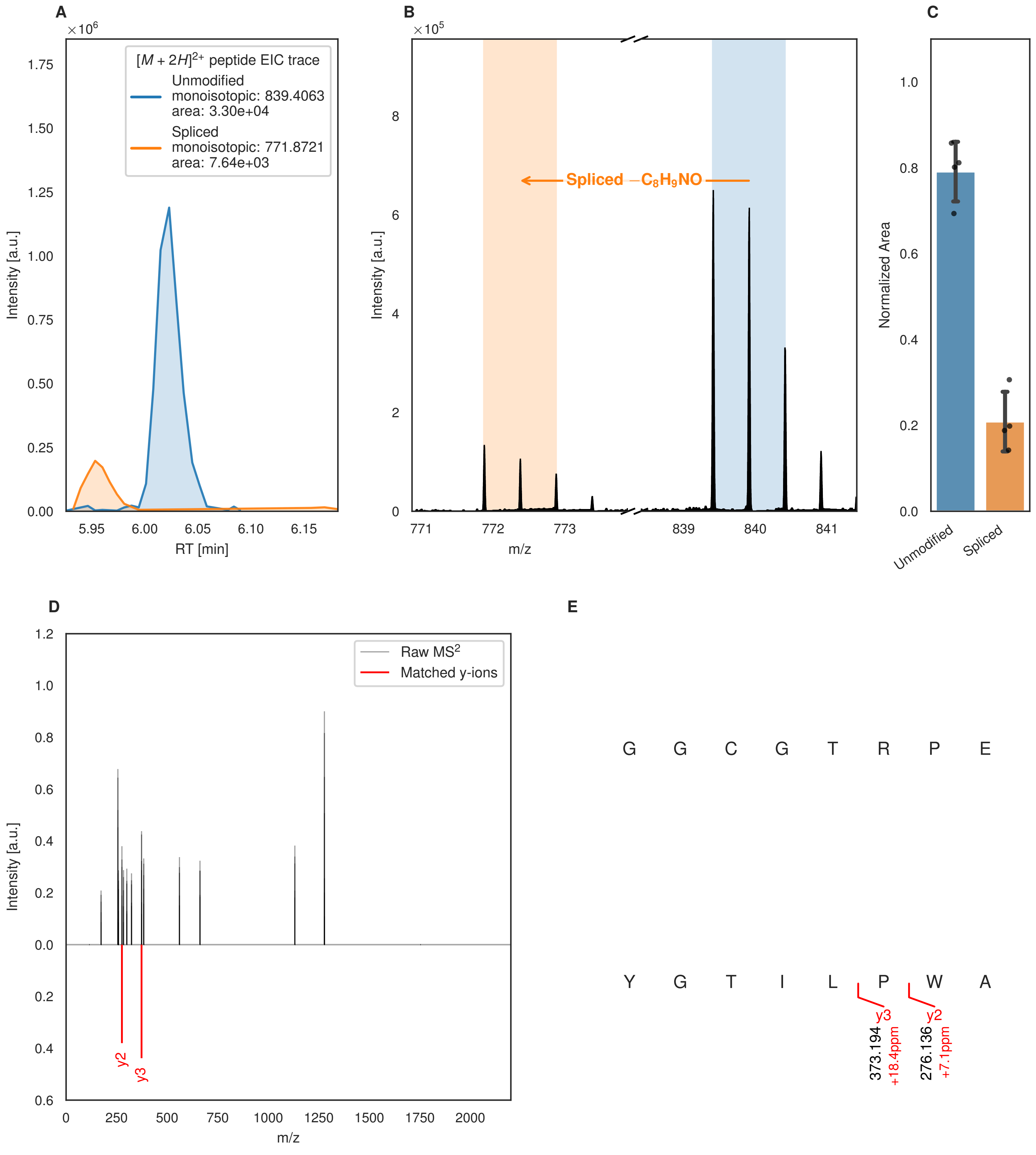
A
B
C
D
E
×10^6
[M + 2H]2+ peptide EIC trace
Unmodified
monoisotopic: 839.4063
area: 3.30e+04
Spliced
monoisotopic: 771.8721
area: 7.64e+03
Intensity [a.u.]
RT [min]
×10^5
Spliced −C8H9NO
m/z
Normalized Area
Unmodified
Spliced
Raw MS2
Matched y-ions
y2
y3
G G C G T R P E
Y G T I L P W A
373.194
+18.4ppm
276.136
+7.1ppm

Figure S151 (figure continued from the previous page).

F

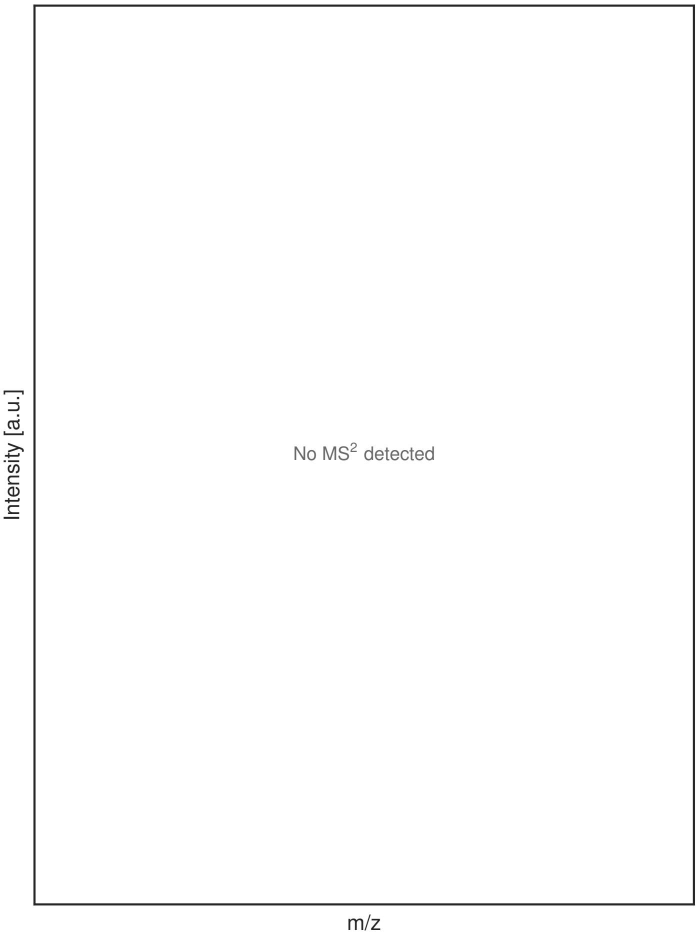


G

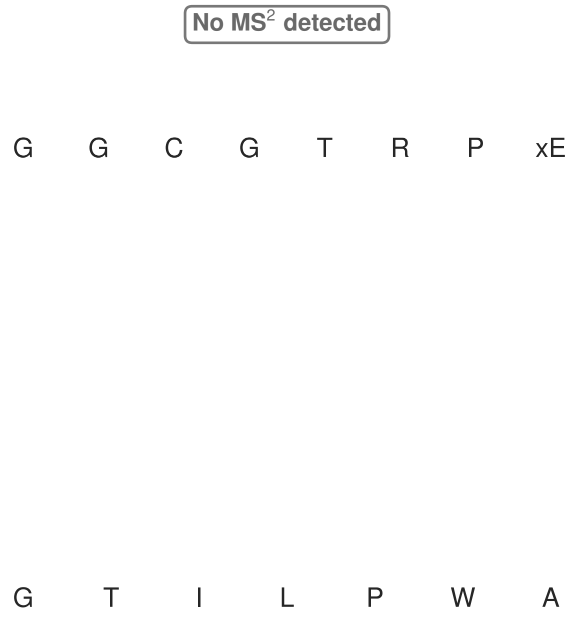


**Figure S151: Modification of the design_17 precursor peptide by PlpXY.** LC-MS analysis of IMAC-purified precursor peptides from *E. coli* BL21(DE3)Tuner containing splicease-encoding plasmid TJ-C-599 (PlpXY) and precursor peptide-encoding plasmid design_17 (RSTN-PlpA3-TEV-designed-17) after expression at 24 °C for 22 h in 1.5 mL cultures after digestion with TEV. Data from n=4 independent biological replicates. Spliced amino acids have an x as a prefix. Spliced refers to a single tyramine excision. A representative experimental replicate is shown in panels **A, B.**

**A** Close-up plot of the representative extracted ion chromatogram (EIC) of the various peptide states; the blue trace corresponds to the Unmodified peptide; the orange trace corresponds to the Spliced peptide. **B** Close-up plot of the m/z values of the peptides detected by the mass spectrometer, with the shaded colored areas corresponding to where the expected m/z values for the peptide m/z traces should occur, with the observed spectrum plotted in black. The Unmodified, Spliced peptide(s) were found in the mass spectrum. **C** Bar plot showing the relative abundance of the various peptide states at charge z=2; the bars represent the average relative abundances normalized to total peak areas, with each individual experimental replicate value plotted as a black dot. **D** Mirrored MS$^2$ plot for the Unmodified peptide. In this mirror plot, the full fragmentation spectrum is plotted in black pointing upward, and the matched fragment b ions (blue) and y ions (red) are plotted pointing downward, mirrored against it, so that each matched fragment can be compared directly with the measured spectrum. **E** MS$^2$ spectrum of the fragments corresponding to the Unmodified peptide sequence, with matched b (blue, above) and y ions (red, below) plotted. **F** Mirrored MS$^2$ plot for the Spliced peptide. In this mirror plot, the full fragmentation spectrum is plotted in black pointing upward, and the matched fragment b ions (blue) and y ions (red) are plotted pointing downward, mirrored against it, so that each matched fragment can be compared directly with the measured spectrum. **G** MS$^2$ spectrum of the fragments corresponding to the Spliced peptide sequence, with matched b (blue, above) and y ions (red, below) plotted.

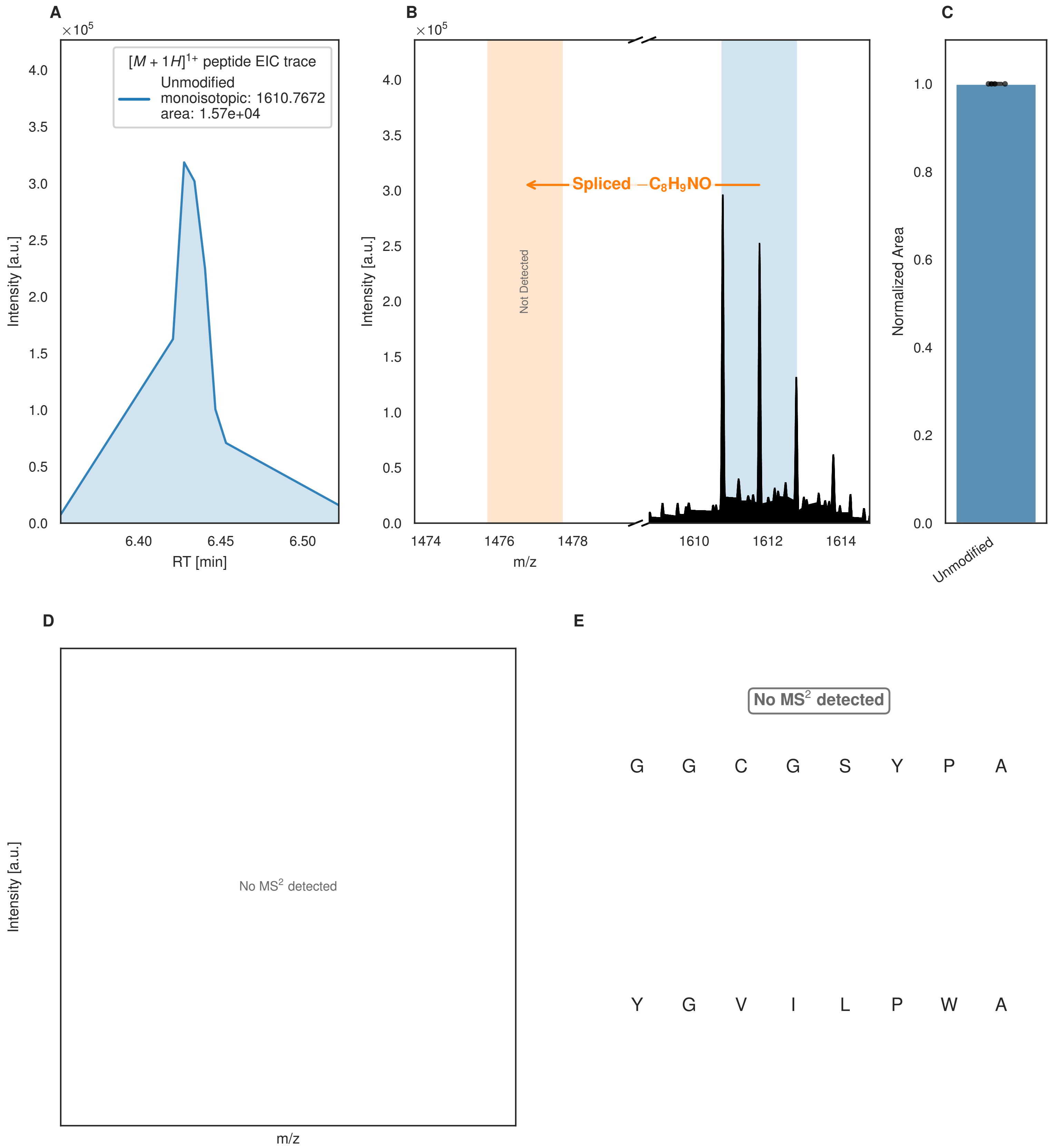


**Figure S152: Modification of the design_18 precursor peptide by PlpXY.** LC-MS analysis of IMAC-purified precursor peptides from *E. coli* BL21(DE3)Tuner containing splicease-encoding plasmid TJ-C-599 (PlpXY) and precursor peptide-encoding plasmid design_18 (RSTN-PlpA3-TEV-designed-18) after expression at 24 °C for 22 h in 1.5 mL cultures after digestion with TEV. Data from n=4 independent biological replicates. Spliced amino acids have an x as a prefix. Spliced refers to a single tyramine excision. A representative experimental replicate is shown in panels **A, B.**

**A** Close-up plot of the representative extracted ion chromatogram (EIC) of the various peptide states; the blue trace corresponds to the Unmodified peptide. **B** Close-up plot of the m/z values of the peptides detected by the mass spectrometer, with the shaded colored areas corresponding to where the expected m/z values for the peptide m/z

(Caption continued on the next page.)

traces should occur, with the observed spectrum plotted in black. The Unmodified peptide(s) were found in the mass spectrum and the Spliced peptide(s) were not detected. **C** Bar plot showing the relative abundance of the various peptide states at charge z=1; the bars represent the average relative abundances normalized to total peak areas, with each individual experimental replicate value plotted as a black dot. **D** Mirrored $MS^2$ plot for the Unmodified peptide. In this mirror plot, the full fragmentation spectrum is plotted in black pointing upward, and the matched fragment b ions (blue) and y ions (red) are plotted pointing downward, mirrored against it, so that each matched fragment can be compared directly with the measured spectrum. **E** $MS^2$ spectrum of the fragments corresponding to the Unmodified peptide sequence, with matched b (blue, above) and y ions (red, below) plotted.

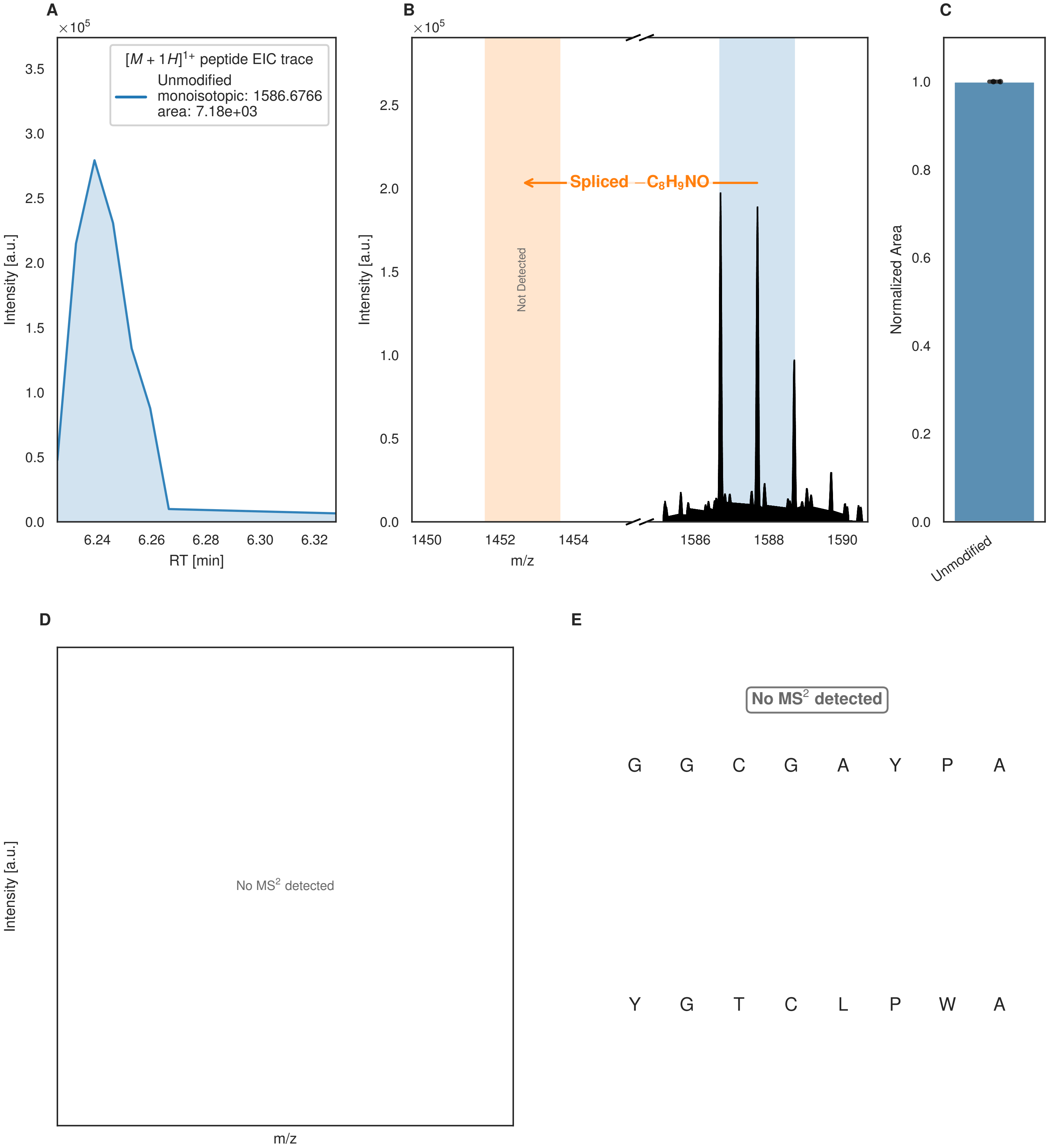


**Figure S153: Modification of the design_19 precursor peptide by PlpXY.** LC-MS analysis of IMAC-purified precursor peptides from *E. coli* BL21(DE3)Tuner containing splicease-encoding plasmid TJ-C-599 (PlpXY) and precursor peptide-encoding plasmid design_19 (RSTN-PlpA3-TEV-designed-19) after expression at 24 °C for 22 h in 1.5 mL cultures after digestion with TEV. Data from n=4 independent biological replicates. Spliced amino acids have an x as a prefix. Spliced refers to a single tyramine excision. A representative experimental replicate is shown in panels **A, B.**

**A** Close-up plot of the representative extracted ion chromatogram (EIC) of the various peptide states; the blue trace corresponds to the Unmodified peptide. **B** Close-up plot of the m/z values of the peptides detected by the mass spectrometer, with the shaded colored areas corresponding to where the expected m/z values for the peptide m/z

(Caption continued on the next page.)

traces should occur, with the observed spectrum plotted in black. The Unmodified peptide(s) were found in the mass spectrum and the Spliced peptide(s) were not detected. **C** Bar plot showing the relative abundance of the various peptide states at charge z=1; the bars represent the average relative abundances normalized to total peak areas, with each individual experimental replicate value plotted as a black dot. **D** Mirrored $MS^2$ plot for the Unmodified peptide. In this mirror plot, the full fragmentation spectrum is plotted in black pointing upward, and the matched fragment b ions (blue) and y ions (red) are plotted pointing downward, mirrored against it, so that each matched fragment can be compared directly with the measured spectrum. **E** $MS^2$ spectrum of the fragments corresponding to the Unmodified peptide sequence, with matched b (blue, above) and y ions (red, below) plotted.

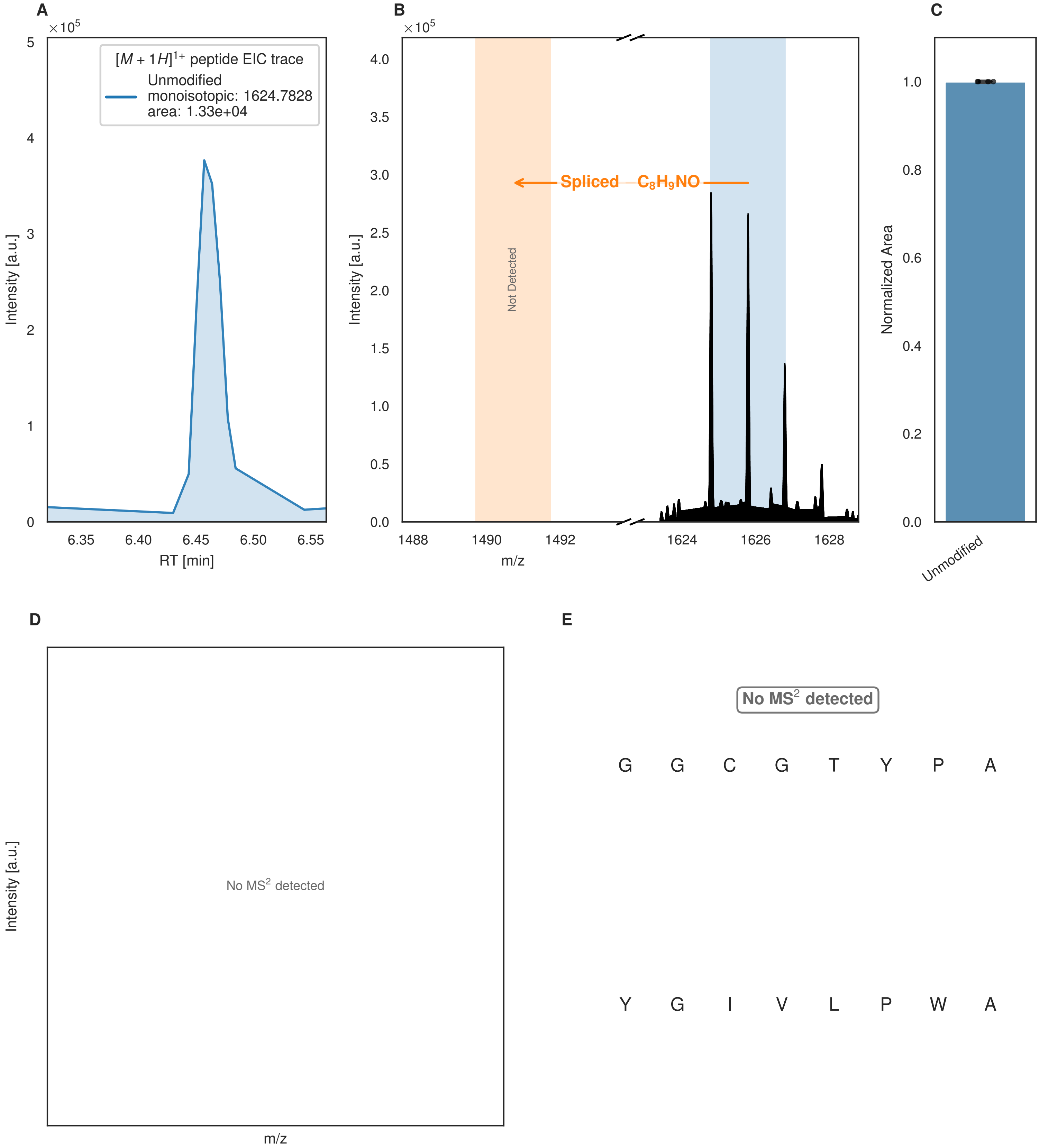


**Figure S154: Modification of the design_20 precursor peptide by PlpXY.** LC-MS analysis of IMAC-purified precursor peptides from *E. coli* BL21(DE3)Tuner containing splicease-encoding plasmid TJ-C-599 (PlpXY) and precursor peptide-encoding plasmid design_20 (RSTN-PlpA3-TEV-designed-20) after expression at 24 °C for 22 h in 1.5 mL cultures after digestion with TEV. Data from n=4 independent biological replicates. Spliced amino acids have an x as a prefix. Spliced refers to a single tyramine excision. A representative experimental replicate is shown in panels **A, B.**

**A** Close-up plot of the representative extracted ion chromatogram (EIC) of the various peptide states; the blue trace corresponds to the Unmodified peptide. **B** Close-up plot of the m/z values of the peptides detected by the mass spectrometer, with the shaded colored areas corresponding to where the expected m/z values for the peptide m/z

(Caption continued on the next page.)

traces should occur, with the observed spectrum plotted in black. The Unmodified peptide(s) were found in the mass spectrum and the Spliced peptide(s) were not detected. **C** Bar plot showing the relative abundance of the various peptide states at charge z=1; the bars represent the average relative abundances normalized to total peak areas, with each individual experimental replicate value plotted as a black dot. **D** Mirrored $MS^2$ plot for the Unmodified peptide. In this mirror plot, the full fragmentation spectrum is plotted in black pointing upward, and the matched fragment b ions (blue) and y ions (red) are plotted pointing downward, mirrored against it, so that each matched fragment can be compared directly with the measured spectrum. **E** $MS^2$ spectrum of the fragments corresponding to the Unmodified peptide sequence, with matched b (blue, above) and y ions (red, below) plotted.

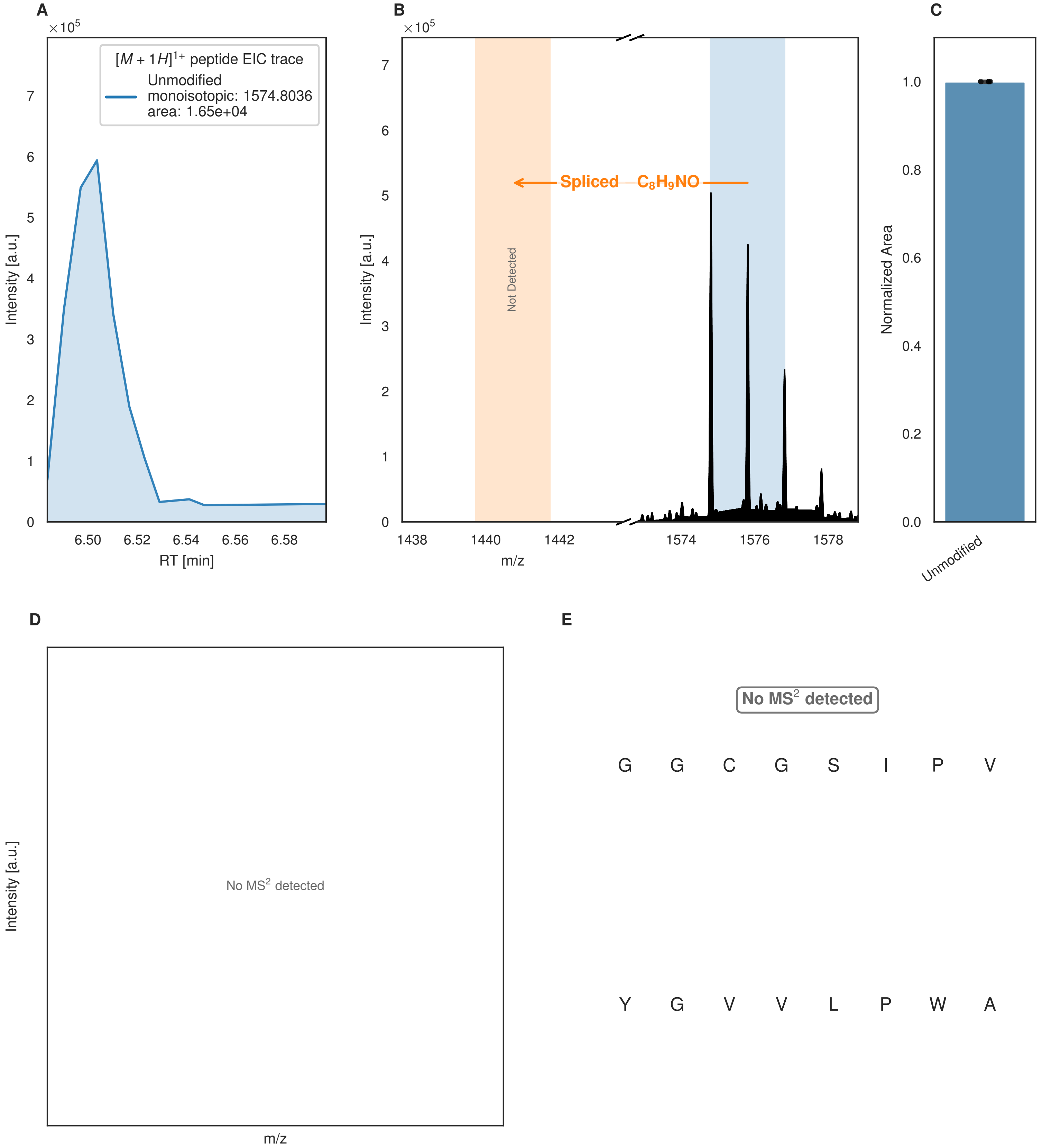


**Figure S155: Modification of the design_22 precursor peptide by PlpXY.** LC-MS analysis of IMAC-purified precursor peptides from *E. coli* BL21(DE3)Tuner containing splicease-encoding plasmid TJ-C-599 (PlpXY) and precursor peptide-encoding plasmid design_22 (RSTN-PlpA3-TEV-designed-22) after expression at 24 °C for 22 h in 1.5 mL cultures after digestion with TEV. Data from n=4 independent biological replicates. Spliced amino acids have an x as a prefix. Spliced refers to a single tyramine excision. A representative experimental replicate is shown in panels **A, B.**

**A** Close-up plot of the representative extracted ion chromatogram (EIC) of the various peptide states; the blue trace corresponds to the Unmodified peptide. **B** Close-up plot of the m/z values of the peptides detected by the mass spectrometer, with the shaded colored areas corresponding to where the expected m/z values for the peptide m/z

(Caption continued on the next page.)

traces should occur, with the observed spectrum plotted in black. The Unmodified peptide(s) were found in the mass spectrum and the Spliced peptide(s) were not detected. **C** Bar plot showing the relative abundance of the various peptide states at charge z=1; the bars represent the average relative abundances normalized to total peak areas, with each individual experimental replicate value plotted as a black dot. **D** Mirrored $MS^2$ plot for the Unmodified peptide. In this mirror plot, the full fragmentation spectrum is plotted in black pointing upward, and the matched fragment b ions (blue) and y ions (red) are plotted pointing downward, mirrored against it, so that each matched fragment can be compared directly with the measured spectrum. **E** $MS^2$ spectrum of the fragments corresponding to the Unmodified peptide sequence, with matched b (blue, above) and y ions (red, below) plotted.

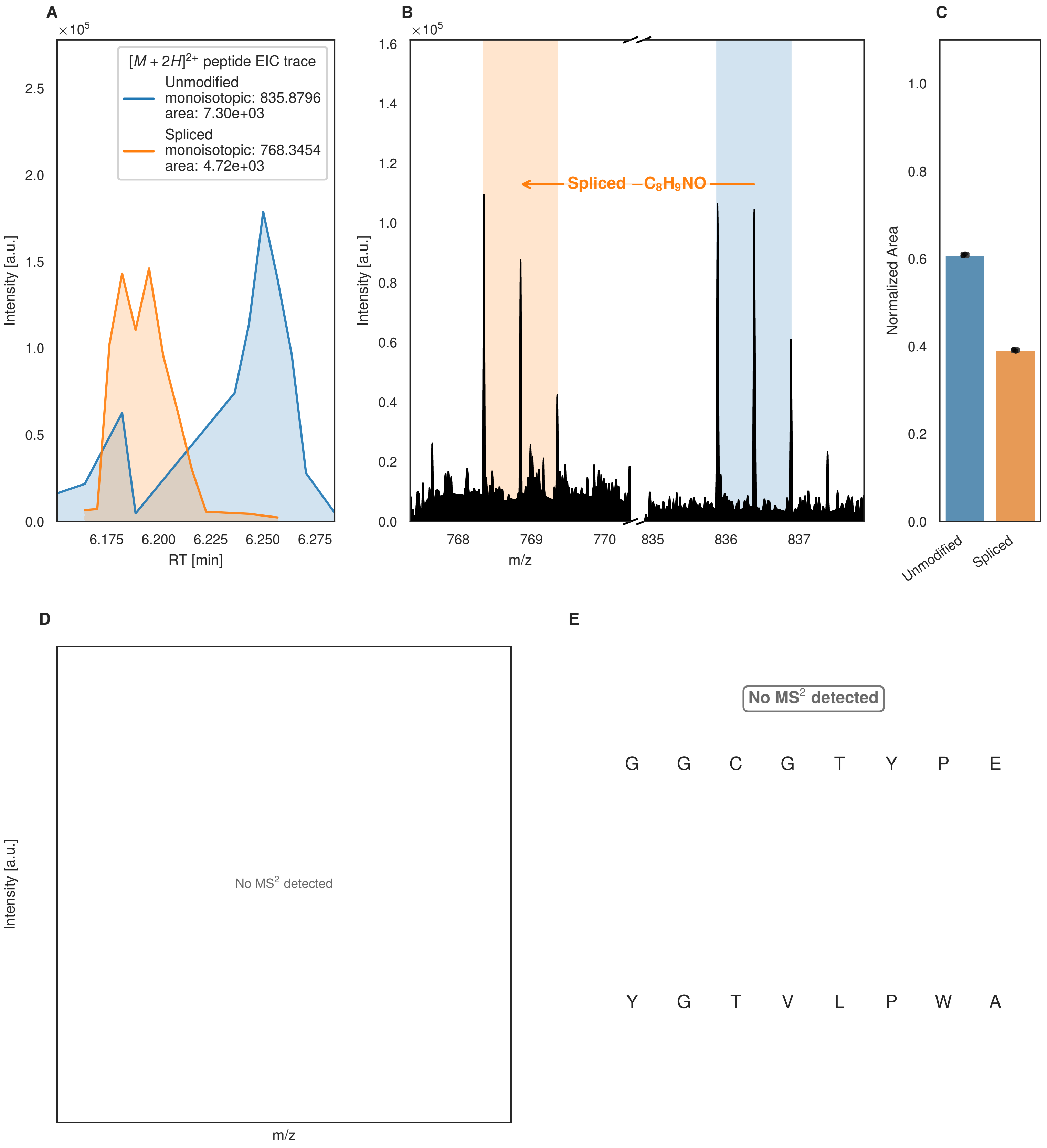
A
×10^5
[M + 2H]2+ peptide EIC trace
Unmodified
monoisotopic: 835.8796
area: 7.30e+03
Spliced
monoisotopic: 768.3454
area: 4.72e+03
Intensity [a.u.]
RT [min]
B
Spliced −C8H9NO
m/z
C
Normalized Area
Unmodified
Spliced
D
No MS2 detected
E
No MS2 detected
G G C G T Y P E
Y G T V L P W A

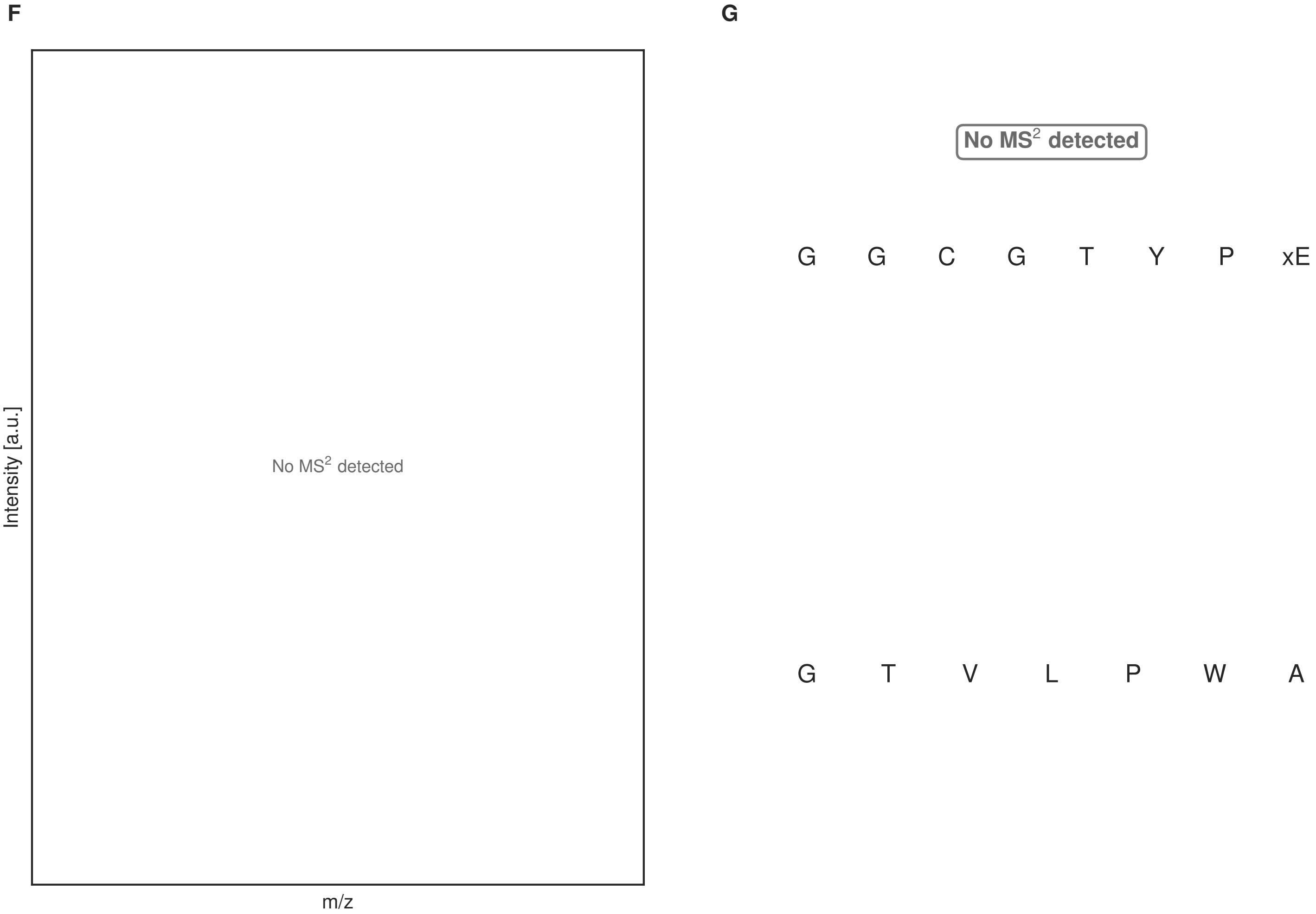


**Figure S156: Modification of the design_23 precursor peptide by PlpXY.** LC-MS analysis of IMAC-purified precursor peptides from *E. coli* BL21(DE3)Tuner containing splicease-encoding plasmid TJ-C-599 (PlpXY) and precursor peptide-encoding plasmid design_23 (RSTN-PlpA3-TEV-designed-23) after expression at 24 °C for 22 h in 1.5 mL cultures after digestion with TEV. Data from n=4 independent biological replicates. Spliced amino acids have an x as a prefix. Spliced refers to a single tyramine excision. A representative experimental replicate is shown in panels **A, B.**

**A** Close-up plot of the representative extracted ion chromatogram (EIC) of the various peptide states; the blue trace corresponds to the Unmodified peptide; the orange trace corresponds to the Spliced peptide. **B** Close-up plot of the m/z values of the peptides detected by the mass spectrometer, with the shaded colored areas corresponding to where the expected m/z values for the peptide m/z traces should occur, with the observed spectrum plotted in black. The Unmodified, Spliced peptide(s) were found in the mass spectrum. **C** Bar plot showing the relative abundance of the various peptide states at charge z=2; the bars represent the average relative abundances normalized to total peak areas, with each individual experimental replicate value plotted as a black dot. **D** Mirrored $MS^2$ plot for the Unmodified peptide. In this mirror plot, the full fragmentation spectrum is plotted in black pointing upward, and the matched fragment b ions (blue) and y ions (red) are plotted pointing downward, mirrored against it, so that each matched fragment can be compared directly with the measured spectrum. **E** $MS^2$ spectrum of the fragments corresponding to the Unmodified peptide sequence, with matched b (blue, above) and y ions (red, below) plotted. **F** Mirrored $MS^2$ plot for the Spliced peptide. In this mirror plot, the full fragmentation spectrum is plotted in black pointing upward, and the matched fragment b ions (blue) and y ions (red) are plotted pointing downward, mirrored against it, so that each matched fragment can be compared directly with the measured spectrum. **G** $MS^2$ spectrum of the fragments corresponding to the Spliced peptide sequence, with matched b (blue, above) and y ions (red, below) plotted.

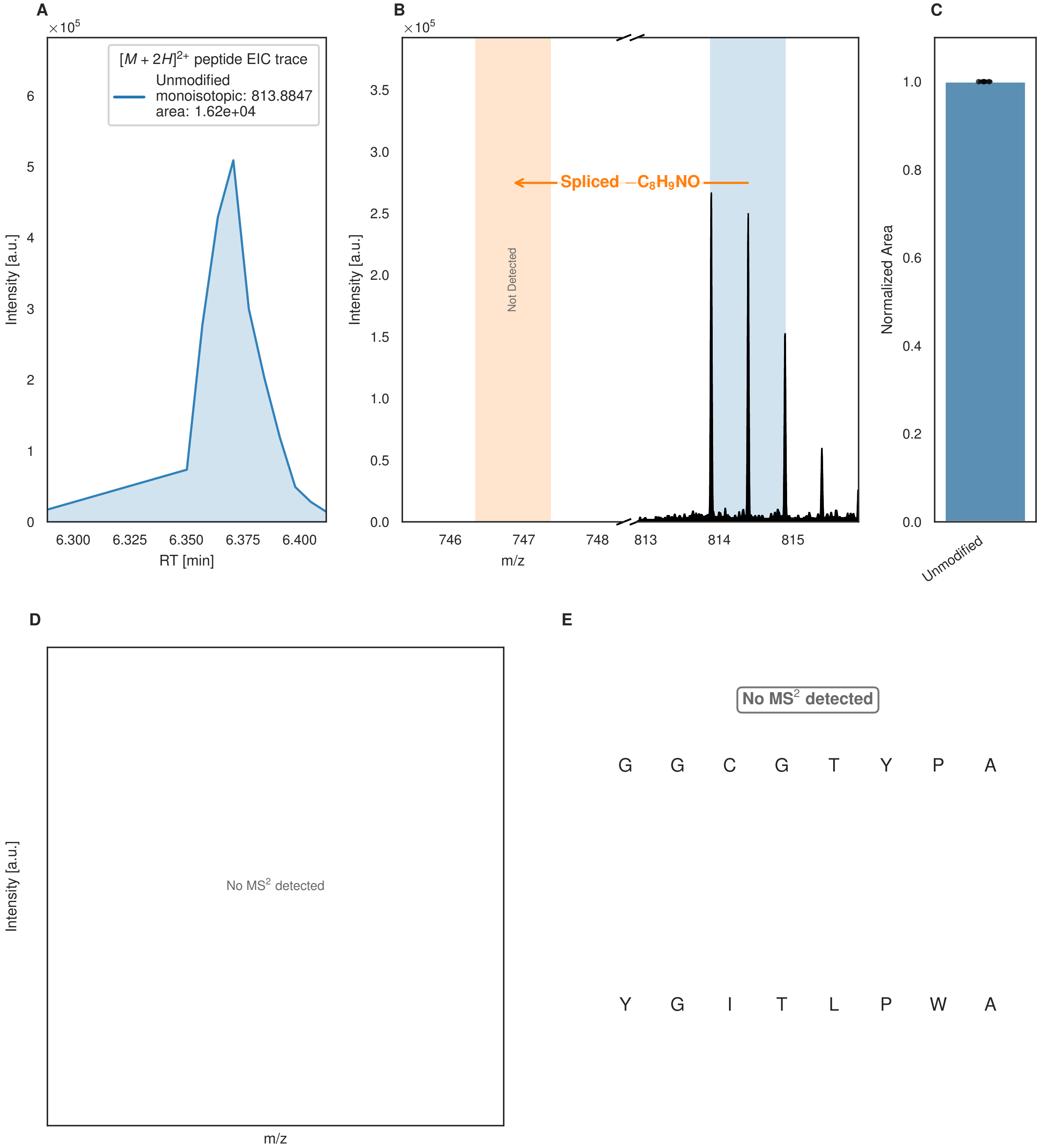


**Figure S157: Modification of the design_24 precursor peptide by PlpXY.** LC-MS analysis of IMAC-purified precursor peptides from *E. coli* BL21(DE3)Tuner containing splicease-encoding plasmid TJ-C-599 (PlpXY) and precursor peptide-encoding plasmid design_24 (RSTN-PlpA3-TEV-designed-24) after expression at 24 °C for 22 h in 1.5 mL cultures after digestion with TEV. Data from n=4 independent biological replicates. Spliced amino acids have an x as a prefix. Spliced refers to a single tyramine excision. A representative experimental replicate is shown in panels **A, B.**

**A** Close-up plot of the representative extracted ion chromatogram (EIC) of the various peptide states; the blue trace corresponds to the Unmodified peptide. **B** Close-up plot of the m/z values of the peptides detected by the mass spectrometer, with the shaded colored areas corresponding to where the expected m/z values for the peptide m/z

(Caption continued on the next page.)

traces should occur, with the observed spectrum plotted in black. The Unmodified peptide(s) were found in the mass spectrum and the Spliced peptide(s) were not detected. **C** Bar plot showing the relative abundance of the various peptide states at charge z=2; the bars represent the average relative abundances normalized to total peak areas, with each individual experimental replicate value plotted as a black dot. **D** Mirrored MS$^2$ plot for the Unmodified peptide. In this mirror plot, the full fragmentation spectrum is plotted in black pointing upward, and the matched fragment b ions (blue) and y ions (red) are plotted pointing downward, mirrored against it, so that each matched fragment can be compared directly with the measured spectrum. **E** MS$^2$ spectrum of the fragments corresponding to the Unmodified peptide sequence, with matched b (blue, above) and y ions (red, below) plotted.

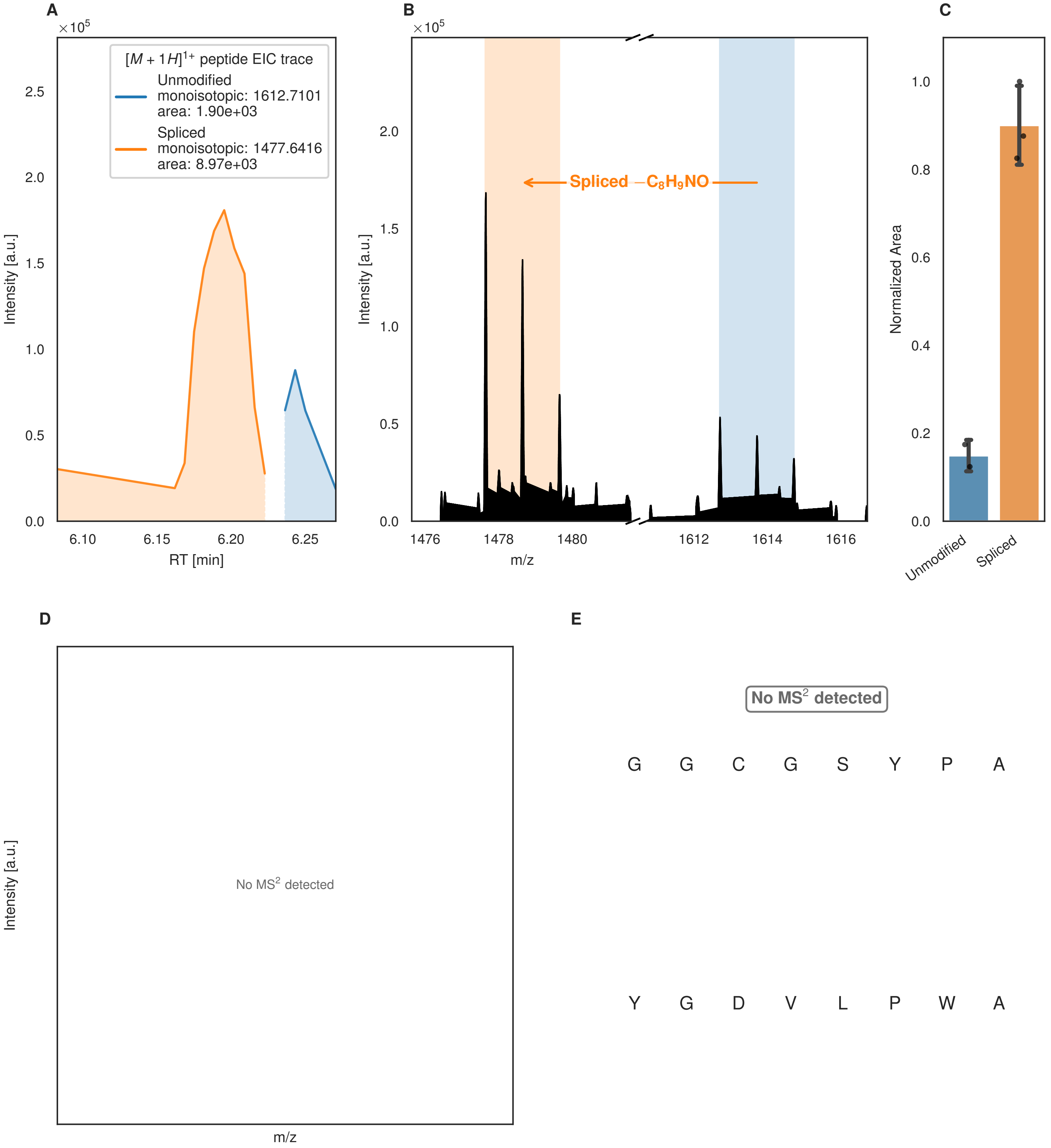
A
×10^5
[M + 1H]1+ peptide EIC trace
Unmodified
monoisotopic: 1612.7101
area: 1.90e+03
Spliced
monoisotopic: 1477.6416
area: 8.97e+03
Intensity [a.u.]
RT [min]
6.10
6.15
6.20
6.25
0.0
0.5
1.0
1.5
2.0
2.5
B
×10^5
Spliced −C8H9NO
Intensity [a.u.]
m/z
1476
1478
1480
1612
1614
1616
C
Normalized Area
Unmodified
Spliced
0.0
0.2
0.4
0.6
0.8
1.0
D
Intensity [a.u.]
No MS2 detected
m/z
E
No MS2 detected
G G C G S Y P A
Y G D V L P W A

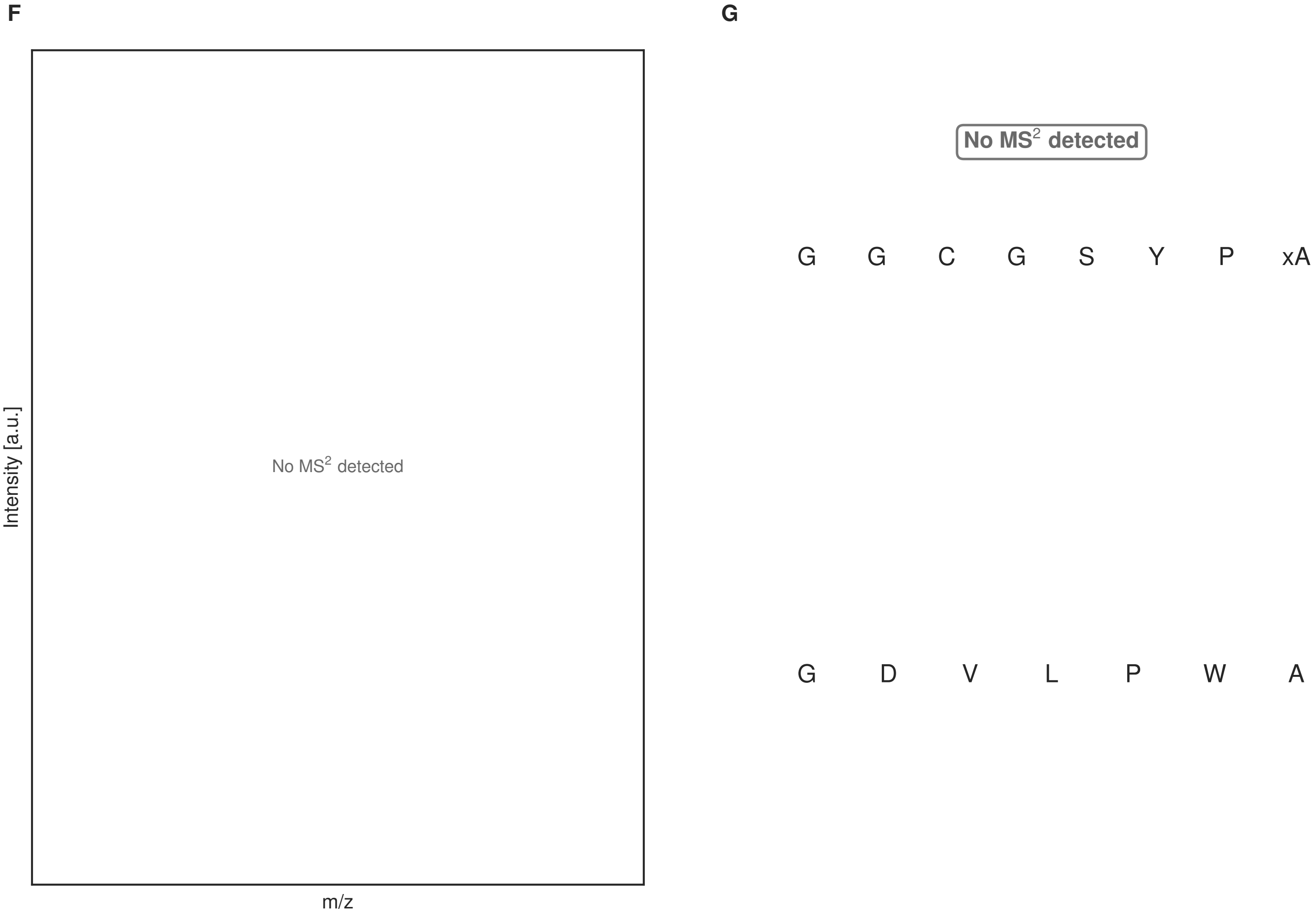


**Figure S158: Modification of the design_25 precursor peptide by PlpXY.** LC-MS analysis of IMAC-purified precursor peptides from *E. coli* BL21(DE3)Tuner containing splicease-encoding plasmid TJ-C-599 (PlpXY) and precursor peptide-encoding plasmid design_25 (RSTN-PlpA3-TEV-designed-25) after expression at 24 °C for 22 h in 1.5 mL cultures after digestion with TEV. Data from n=4 independent biological replicates. Spliced amino acids have an x as a prefix. Spliced refers to a single tyramine excision. A representative experimental replicate is shown in panels **A, B.**
**A** Close-up plot of the representative extracted ion chromatogram (EIC) of the various peptide states; the blue trace corresponds to the Unmodified peptide; the orange trace corresponds to the Spliced peptide. **B** Close-up plot of the m/z values of the peptides detected by the mass spectrometer, with the shaded colored areas corresponding to where the expected m/z values for the peptide m/z traces should occur, with the observed spectrum plotted in black. The Unmodified, Spliced peptide(s) were found in the mass spectrum. **C** Bar plot showing the relative abundance of the various peptide states at charge z=1; the bars represent the average relative abundances normalized to total peak areas, with each individual experimental replicate value plotted as a black dot. **D** Mirrored $MS^2$ plot for the Unmodified peptide. In this mirror plot, the full fragmentation spectrum is plotted in black pointing upward, and the matched fragment b ions (blue) and y ions (red) are plotted pointing downward, mirrored against it, so that each matched fragment can be compared directly with the measured spectrum. **E** $MS^2$ spectrum of the fragments corresponding to the Unmodified peptide sequence, with matched b (blue, above) and y ions (red, below) plotted. **F** Mirrored $MS^2$ plot for the Spliced peptide. In this mirror plot, the full fragmentation spectrum is plotted in black pointing upward, and the matched fragment b ions (blue) and y ions (red) are plotted pointing downward, mirrored against it, so that each matched fragment can be compared directly with the measured spectrum. **G** $MS^2$ spectrum of the fragments corresponding to the Spliced peptide sequence, with matched b (blue, above) and y ions (red, below) plotted.

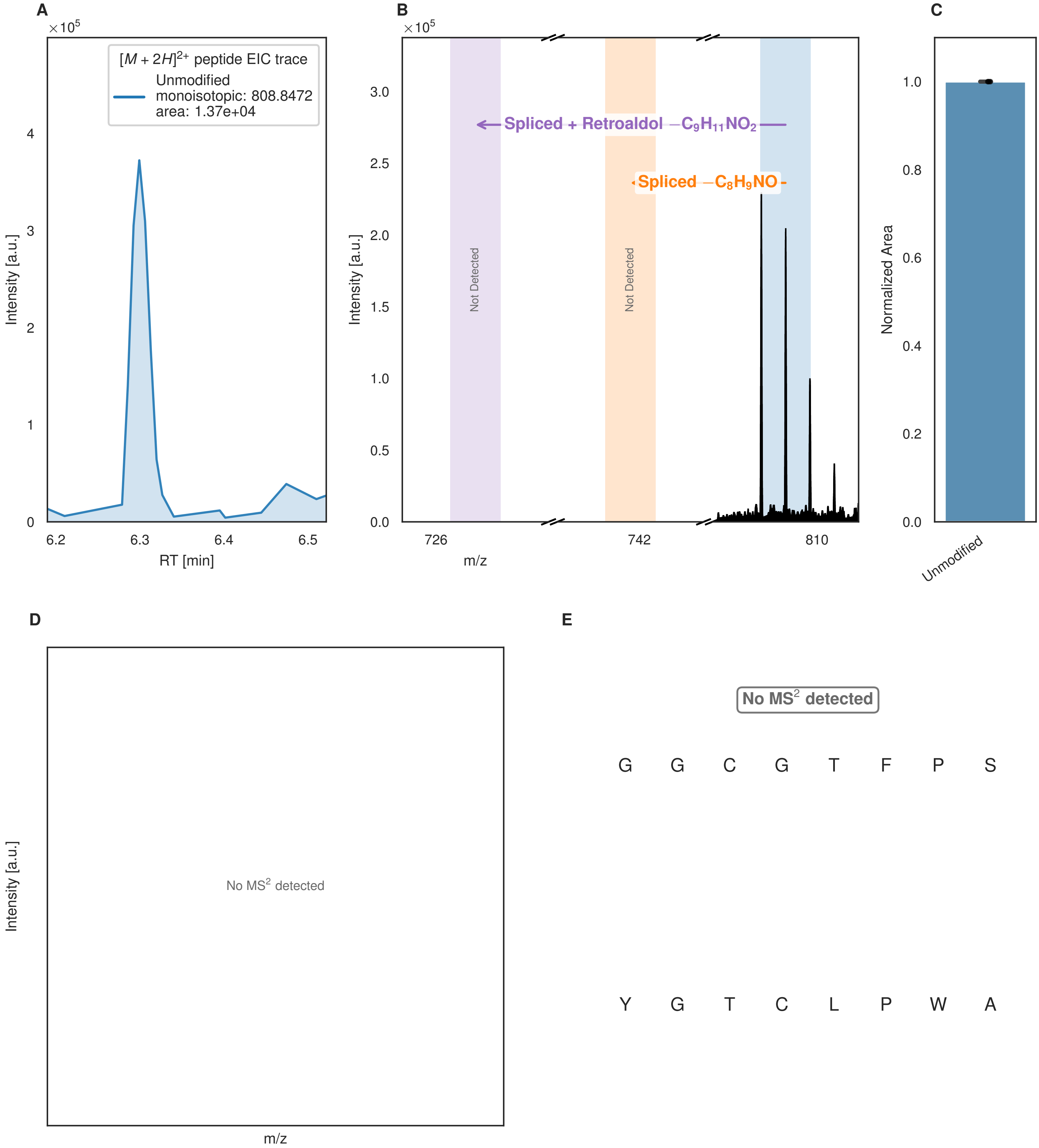


**Figure S159: Modification of the design_26 precursor peptide by PlpXY.** LC-MS analysis of IMAC-purified precursor peptides from *E. coli* BL21(DE3)Tuner containing splicease-encoding plasmid TJ-C-599 (PlpXY) and precursor peptide-encoding plasmid design_26 (RSTN-PlpA3-TEV-designed-26) after expression at 24 °C for 22 h in 1.5 mL cultures after digestion with TEV. Data from n=4 independent biological replicates. Spliced amino acids have an x as a prefix. Spliced refers to a single tyramine excision; Spliced + Retroaldol refers to a glycine-derived ketoamide (xG). A representative experimental replicate is shown in panels **A, B.**
**A** Close-up plot of the representative extracted ion chromatogram (EIC) of the various peptide states; the blue trace corresponds to the Unmodified peptide. **B** Close-up plot of the m/z values of the peptides detected by the mass spectrometer, with the shaded colored areas corresponding to where the expected m/z values for the peptide m/z

(Caption continued on the next page.)

traces should occur, with the observed spectrum plotted in black. The Unmodified peptide(s) were found in the mass spectrum and the Spliced, Spliced + Retroaldol peptide(s) were not detected. **C** Bar plot showing the relative abundance of the various peptide states at charge z=2; the bars represent the average relative abundances normalized to total peak areas, with each individual experimental replicate value plotted as a black dot. **D** Mirrored $MS^2$ plot for the Unmodified peptide. In this mirror plot, the full fragmentation spectrum is plotted in black pointing upward, and the matched fragment b ions (blue) and y ions (red) are plotted pointing downward, mirrored against it, so that each matched fragment can be compared directly with the measured spectrum. **E** $MS^2$ spectrum of the fragments corresponding to the Unmodified peptide sequence, with matched b (blue, above) and y ions (red, below) plotted.

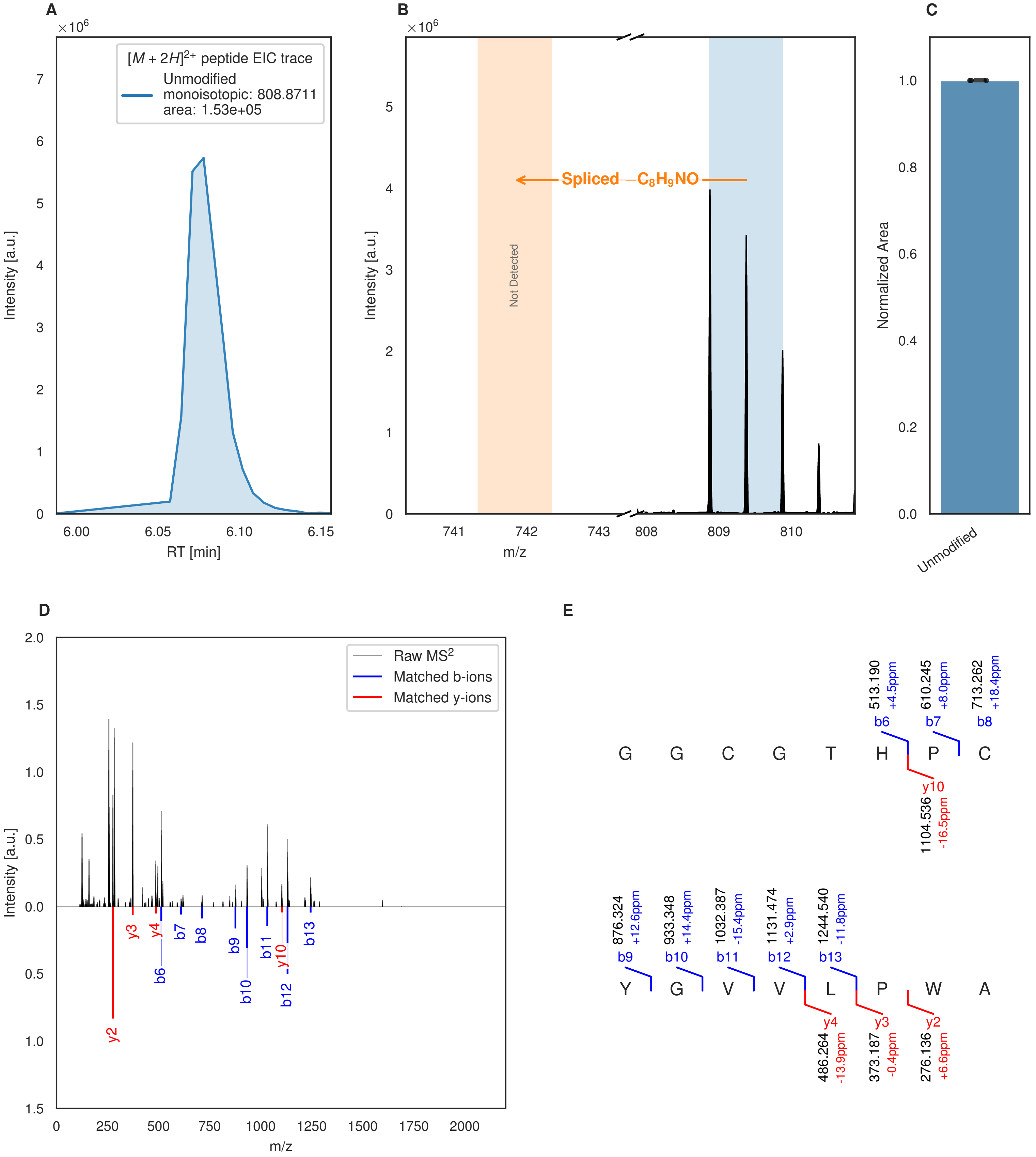


**Figure S160: Modification of the design_27 precursor peptide by PlpXY.** LC-MS analysis of IMAC-purified precursor peptides from *E. coli* BL21(DE3)Tuner containing splicease-encoding plasmid TJ-C-599 (PlpXY) and precursor peptide-encoding plasmid design_27 (RSTN-PlpA3-TEV-designed-27) after expression at 24 °C for 22 h in 1.5 mL cultures after digestion with TEV. Data from n=4 independent biological replicates. Spliced amino acids have an x as a prefix. Spliced refers to a single tyramine excision. A representative experimental replicate is shown in panels **A, B.**

**A** Close-up plot of the representative extracted ion chromatogram (EIC) of the various peptide states; the blue trace corresponds to the Unmodified peptide. **B** Close-up plot of the m/z values of the peptides detected by the mass spectrometer, with the shaded colored areas corresponding to where the expected m/z values for the peptide m/z

(Caption continued on the next page.)

traces should occur, with the observed spectrum plotted in black. The Unmodified peptide(s) were found in the mass spectrum and the Spliced peptide(s) were not detected. **C** Bar plot showing the relative abundance of the various peptide states at charge z=2; the bars represent the average relative abundances normalized to total peak areas, with each individual experimental replicate value plotted as a black dot. **D** Mirrored $MS^2$ plot for the Unmodified peptide. In this mirror plot, the full fragmentation spectrum is plotted in black pointing upward, and the matched fragment b ions (blue) and y ions (red) are plotted pointing downward, mirrored against it, so that each matched fragment can be compared directly with the measured spectrum. **E** $MS^2$ spectrum of the fragments corresponding to the Unmodified peptide sequence, with matched b (blue, above) and y ions (red, below) plotted.

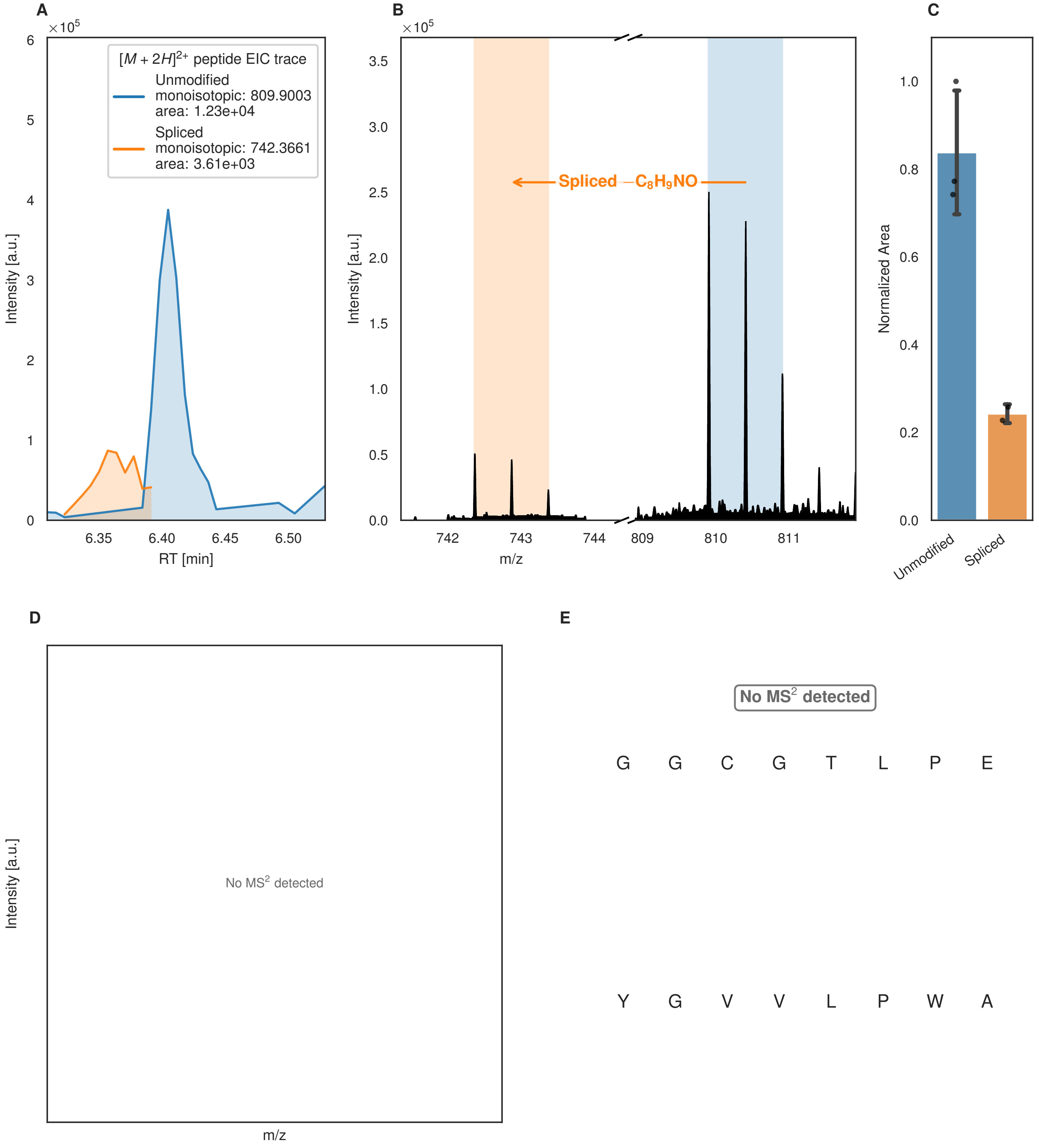

A
×10^5
[M + 2H]2+ peptide EIC trace
Unmodified
monoisotopic: 809.9003
area: 1.23e+04
Spliced
monoisotopic: 742.3661
area: 3.61e+03
Intensity [a.u.]
RT [min]
6.35
6.40
6.45
6.50
B
×10^5
Spliced −C8H9NO
Intensity [a.u.]
m/z
742
743
744
809
810
811
C
Normalized Area
Unmodified
Spliced
D
Intensity [a.u.]
No MS2 detected
m/z
E
No MS2 detected
G G C G T L P E
Y G V V L P W A

Figure S161 (figure continued from the previous page).

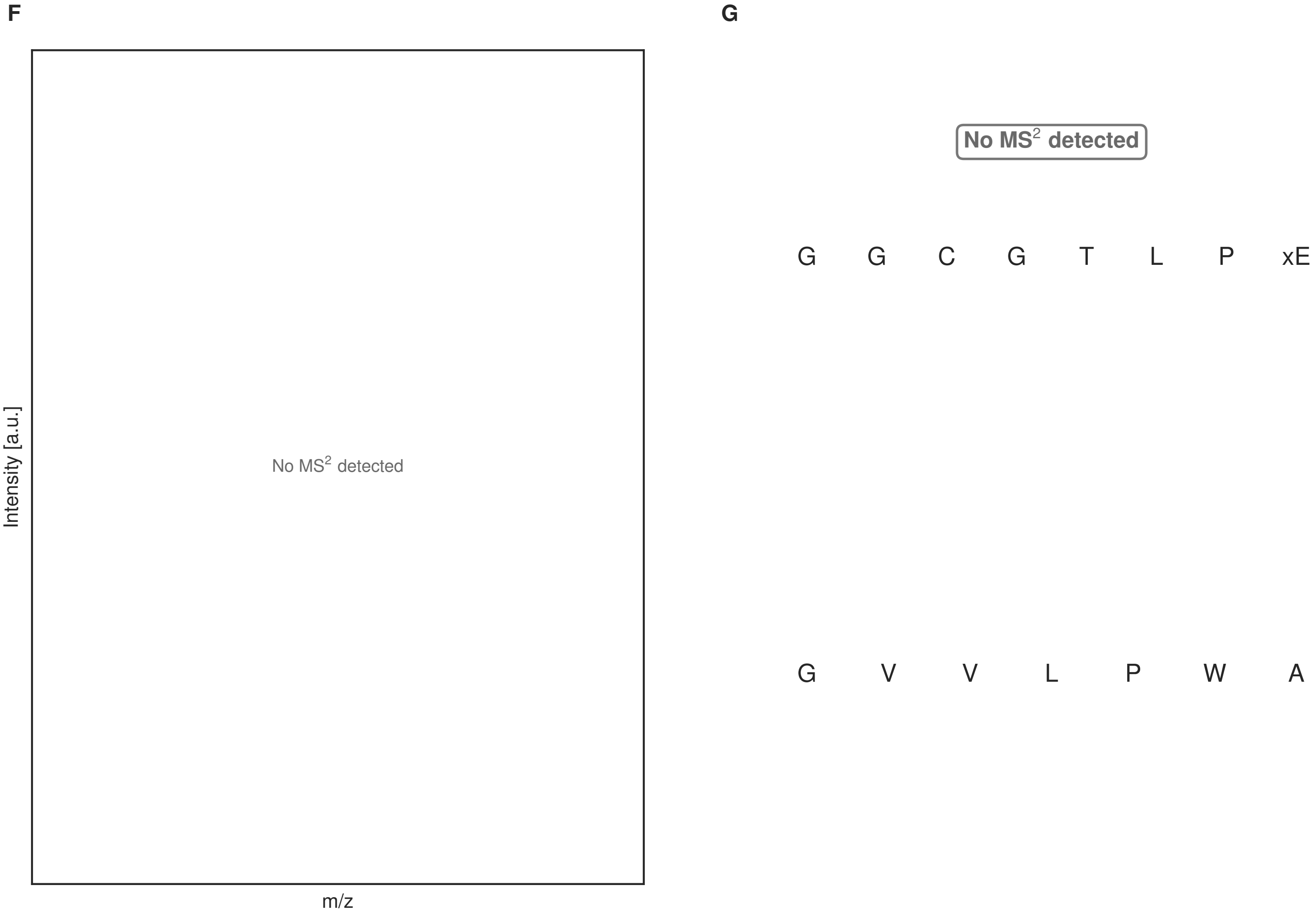


**Figure S161: Modification of the design_28 precursor peptide by PlpXY.** LC-MS analysis of IMAC-purified precursor peptides from *E. coli* BL21(DE3)Tuner containing splicease-encoding plasmid TJ-C-599 (PlpXY) and precursor peptide-encoding plasmid design_28 (RSTN-PlpA3-TEV-designed-28) after expression at 24 °C for 22 h in 1.5 mL cultures after digestion with TEV. Data from n=4 independent biological replicates. Spliced amino acids have an x as a prefix. Spliced refers to a single tyramine excision. A representative experimental replicate is shown in panels **A, B.**

**A** Close-up plot of the representative extracted ion chromatogram (EIC) of the various peptide states; the blue trace corresponds to the Unmodified peptide; the orange trace corresponds to the Spliced peptide. **B** Close-up plot of the m/z values of the peptides detected by the mass spectrometer, with the shaded colored areas corresponding to where the expected m/z values for the peptide m/z traces should occur, with the observed spectrum plotted in black. The Unmodified, Spliced peptide(s) were found in the mass spectrum. **C** Bar plot showing the relative abundance of the various peptide states at charge z=2; the bars represent the average relative abundances normalized to total peak areas, with each individual experimental replicate value plotted as a black dot. **D** Mirrored MS$^2$ plot for the Unmodified peptide. In this mirror plot, the full fragmentation spectrum is plotted in black pointing upward, and the matched fragment b ions (blue) and y ions (red) are plotted pointing downward, mirrored against it, so that each matched fragment can be compared directly with the measured spectrum. **E** MS$^2$ spectrum of the fragments corresponding to the Unmodified peptide sequence, with matched b (blue, above) and y ions (red, below) plotted. **F** Mirrored MS$^2$ plot for the Spliced peptide. In this mirror plot, the full fragmentation spectrum is plotted in black pointing upward, and the matched fragment b ions (blue) and y ions (red) are plotted pointing downward, mirrored against it, so that each matched fragment can be compared directly with the measured spectrum. **G** MS$^2$ spectrum of the fragments corresponding to the Spliced peptide sequence, with matched b (blue, above) and y ions (red, below) plotted.

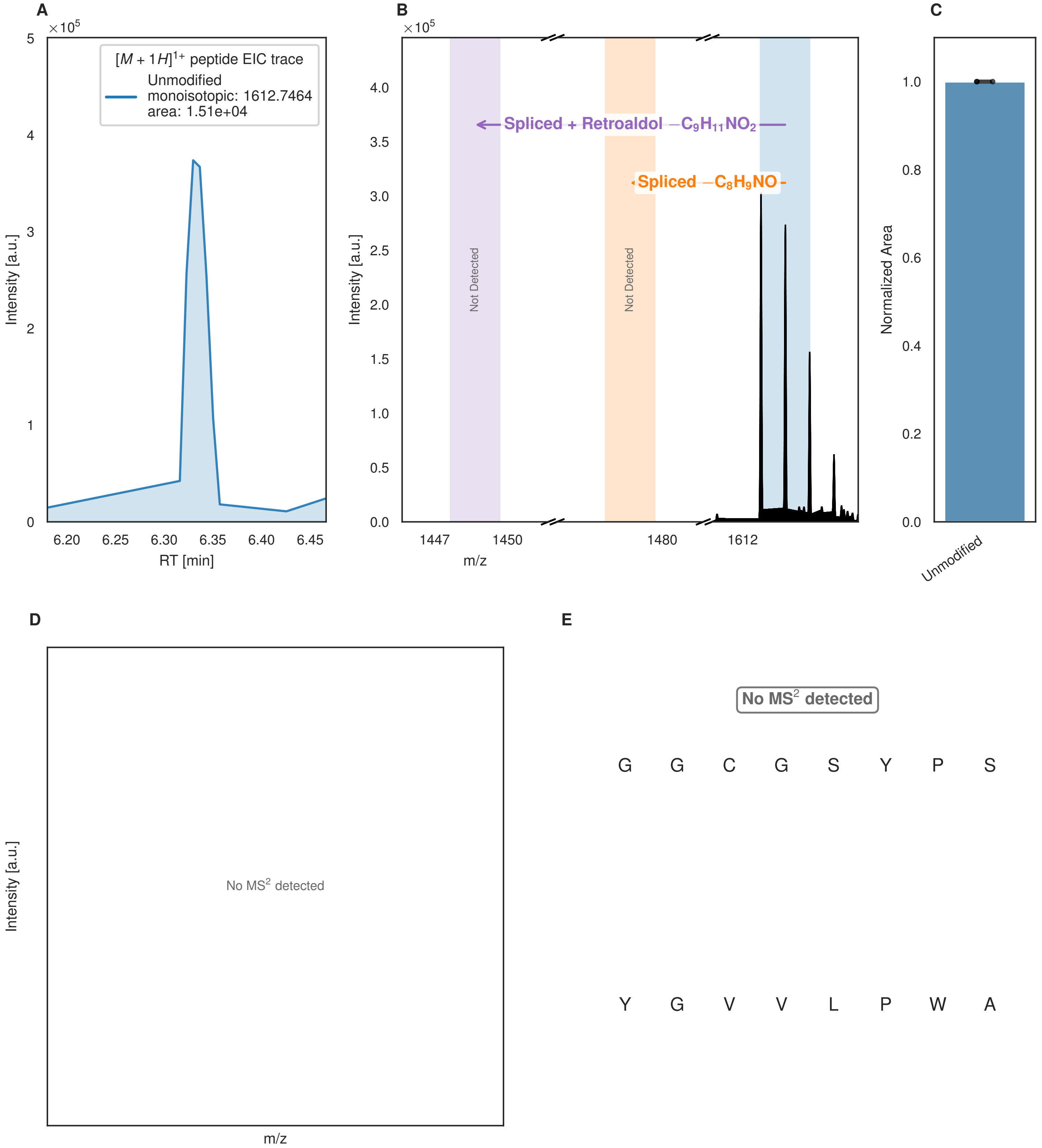


**Figure S162: Modification of the design_29 precursor peptide by PlpXY.** LC-MS analysis of IMAC-purified precursor peptides from *E. coli* BL21(DE3)Tuner containing splicease-encoding plasmid TJ-C-599 (PlpXY) and precursor peptide-encoding plasmid design_29 (RSTN-PlpA3-TEV-designed-29) after expression at 24 °C for 22 h in 1.5 mL cultures after digestion with TEV. Data from n=4 independent biological replicates. Spliced amino acids have an x as a prefix. Spliced refers to a single tyramine excision; Spliced + Retroaldol refers to a glycine-derived ketoamide (xG). A representative experimental replicate is shown in panels **A, B.**
**A** Close-up plot of the representative extracted ion chromatogram (EIC) of the various peptide states; the blue trace corresponds to the Unmodified peptide. **B** Close-up plot of the m/z values of the peptides detected by the mass spectrometer, with the shaded colored areas corresponding to where the expected m/z values for the peptide m/z

(Caption continued on the next page.)

traces should occur, with the observed spectrum plotted in black. The Unmodified peptide(s) were found in the mass spectrum and the Spliced, Spliced + Retroaldol peptide(s) were not detected. **C** Bar plot showing the relative abundance of the various peptide states at charge z=1; the bars represent the average relative abundances normalized to total peak areas, with each individual experimental replicate value plotted as a black dot. **D** Mirrored $MS^2$ plot for the Unmodified peptide. In this mirror plot, the full fragmentation spectrum is plotted in black pointing upward, and the matched fragment b ions (blue) and y ions (red) are plotted pointing downward, mirrored against it, so that each matched fragment can be compared directly with the measured spectrum. **E** $MS^2$ spectrum of the fragments corresponding to the Unmodified peptide sequence, with matched b (blue, above) and y ions (red, below) plotted.

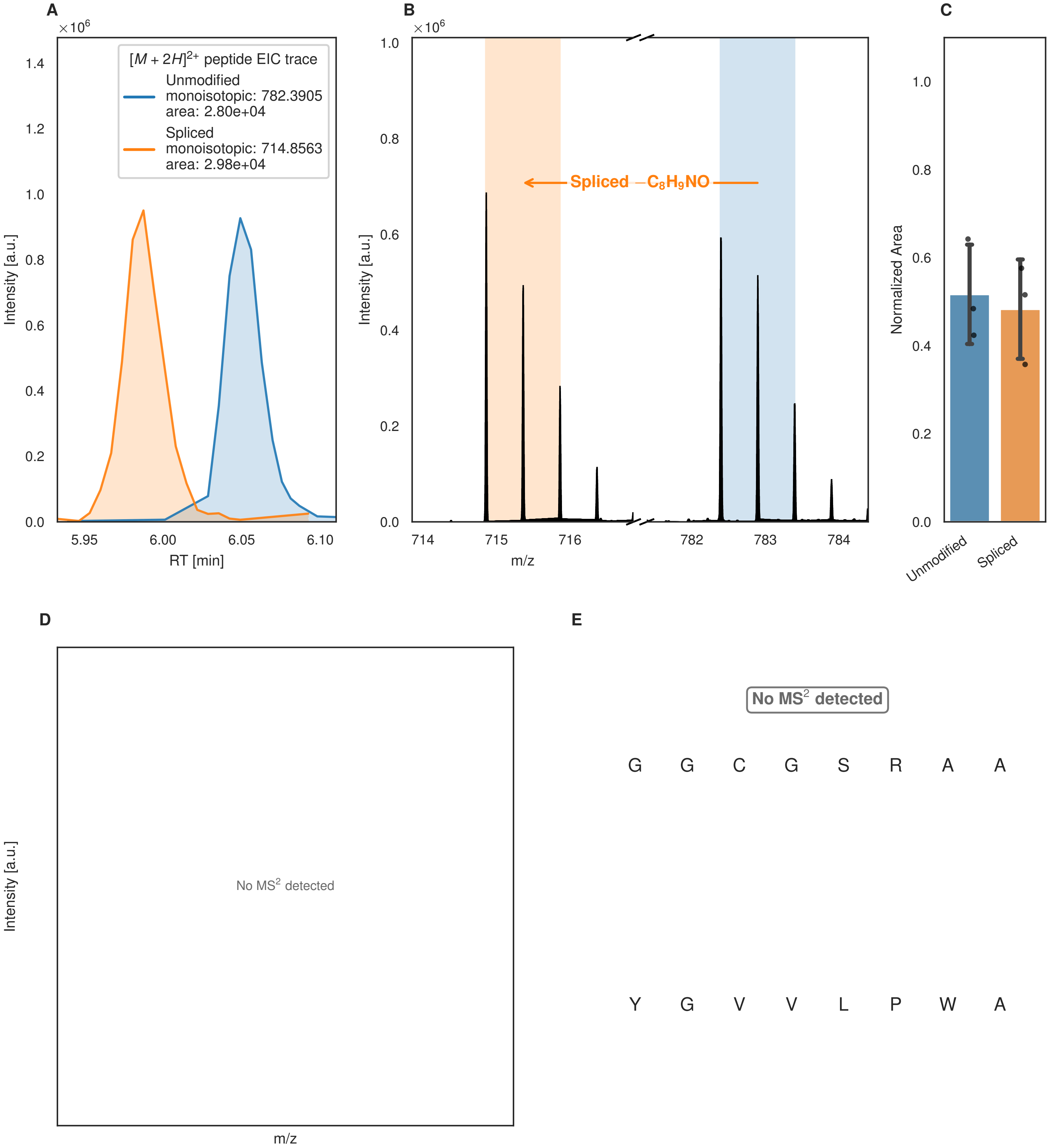

A
×10^6
[M + 2H]2+ peptide EIC trace
Unmodified
monoisotopic: 782.3905
area: 2.80e+04
Spliced
monoisotopic: 714.8563
area: 2.98e+04
Intensity [a.u.]
RT [min]
B
×10^6
Spliced −C8H9NO
Intensity [a.u.]
m/z
C
Normalized Area
Unmodified
Spliced
D
No MS2 detected
Intensity [a.u.]
m/z
E
No MS2 detected
G G C G S R A A
Y G V V L P W A

Figure S163 (figure continued from the previous page).

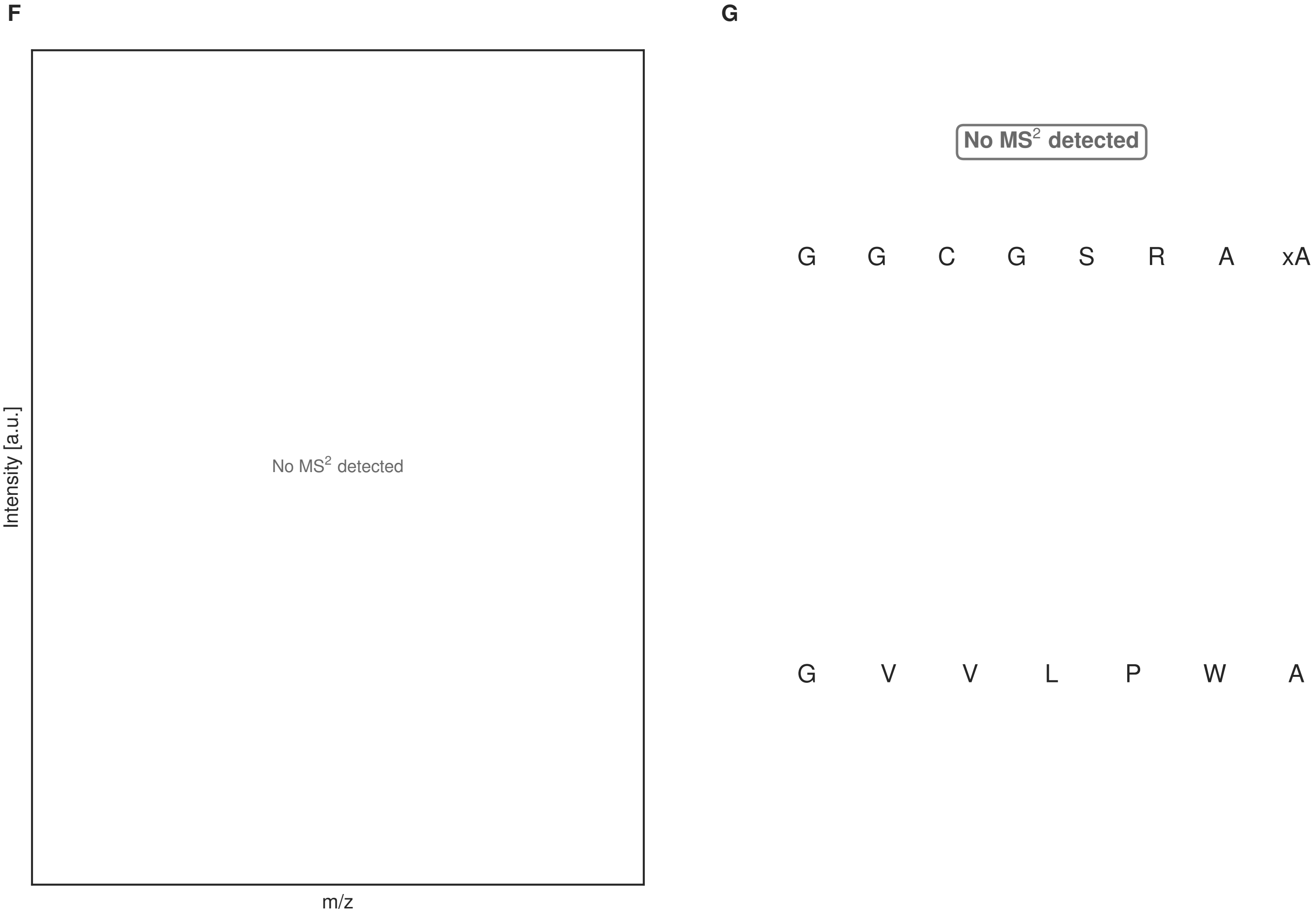


**Figure S163: Modification of the design_30 precursor peptide by PlpXY.** LC-MS analysis of IMAC-purified precursor peptides from *E. coli* BL21(DE3)Tuner containing splicease-encoding plasmid TJ-C-599 (PlpXY) and precursor peptide-encoding plasmid design_30 (RSTN-PlpA3-TEV-designed-30) after expression at 24 °C for 22 h in 1.5 mL cultures after digestion with TEV. Data from n=4 independent biological replicates. Spliced amino acids have an x as a prefix. Spliced refers to a single tyramine excision. A representative experimental replicate is shown in panels **A, B.**

**A** Close-up plot of the representative extracted ion chromatogram (EIC) of the various peptide states; the blue trace corresponds to the Unmodified peptide; the orange trace corresponds to the Spliced peptide. **B** Close-up plot of the m/z values of the peptides detected by the mass spectrometer, with the shaded colored areas corresponding to where the expected m/z values for the peptide m/z traces should occur, with the observed spectrum plotted in black. The Unmodified, Spliced peptide(s) were found in the mass spectrum. **C** Bar plot showing the relative abundance of the various peptide states at charge z=2; the bars represent the average relative abundances normalized to total peak areas, with each individual experimental replicate value plotted as a black dot. **D** Mirrored MS$^2$ plot for the Unmodified peptide. In this mirror plot, the full fragmentation spectrum is plotted in black pointing upward, and the matched fragment b ions (blue) and y ions (red) are plotted pointing downward, mirrored against it, so that each matched fragment can be compared directly with the measured spectrum. **E** MS$^2$ spectrum of the fragments corresponding to the Unmodified peptide sequence, with matched b (blue, above) and y ions (red, below) plotted. **F** Mirrored MS$^2$ plot for the Spliced peptide. In this mirror plot, the full fragmentation spectrum is plotted in black pointing upward, and the matched fragment b ions (blue) and y ions (red) are plotted pointing downward, mirrored against it, so that each matched fragment can be compared directly with the measured spectrum. **G** MS$^2$ spectrum of the fragments corresponding to the Spliced peptide sequence, with matched b (blue, above) and y ions (red, below) plotted.

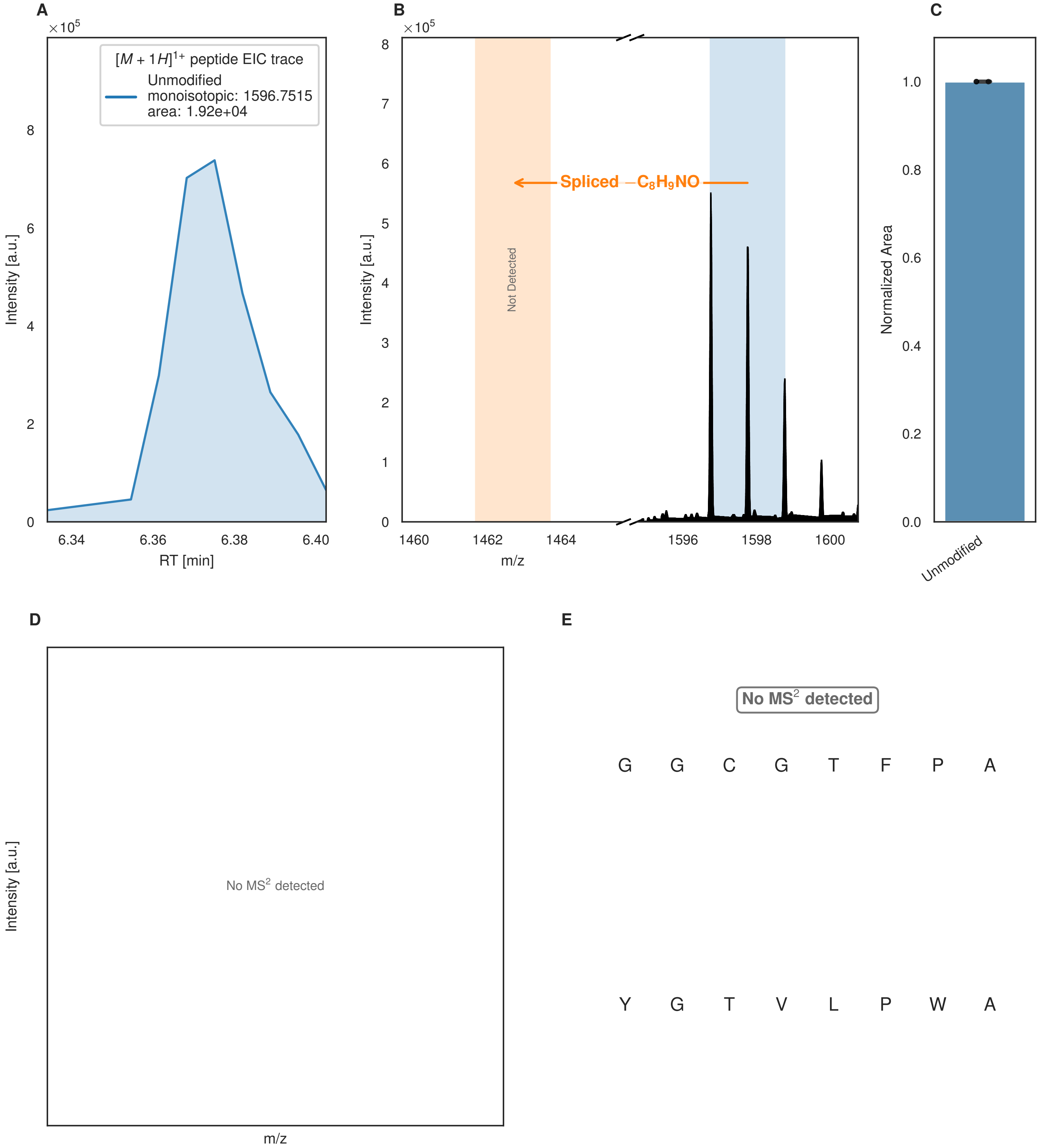


**Figure S164: Modification of the design_31 precursor peptide by PlpXY.** LC-MS analysis of IMAC-purified precursor peptides from *E. coli* BL21(DE3)Tuner containing splicease-encoding plasmid TJ-C-599 (PlpXY) and precursor peptide-encoding plasmid design_31 (RSTN-PlpA3-TEV-designed-31) after expression at 24 °C for 22 h in 1.5 mL cultures after digestion with TEV. Data from n=4 independent biological replicates. Spliced amino acids have an x as a prefix. Spliced refers to a single tyramine excision. A representative experimental replicate is shown in panels **A, B.**

**A** Close-up plot of the representative extracted ion chromatogram (EIC) of the various peptide states; the blue trace corresponds to the Unmodified peptide. **B** Close-up plot of the m/z values of the peptides detected by the mass spectrometer, with the shaded colored areas corresponding to where the expected m/z values for the peptide m/z

(Caption continued on the next page.)

traces should occur, with the observed spectrum plotted in black. The Unmodified peptide(s) were found in the mass spectrum and the Spliced peptide(s) were not detected. **C** Bar plot showing the relative abundance of the various peptide states at charge z=1; the bars represent the average relative abundances normalized to total peak areas, with each individual experimental replicate value plotted as a black dot. **D** Mirrored $MS^2$ plot for the Unmodified peptide. In this mirror plot, the full fragmentation spectrum is plotted in black pointing upward, and the matched fragment b ions (blue) and y ions (red) are plotted pointing downward, mirrored against it, so that each matched fragment can be compared directly with the measured spectrum. **E** $MS^2$ spectrum of the fragments corresponding to the Unmodified peptide sequence, with matched b (blue, above) and y ions (red, below) plotted.

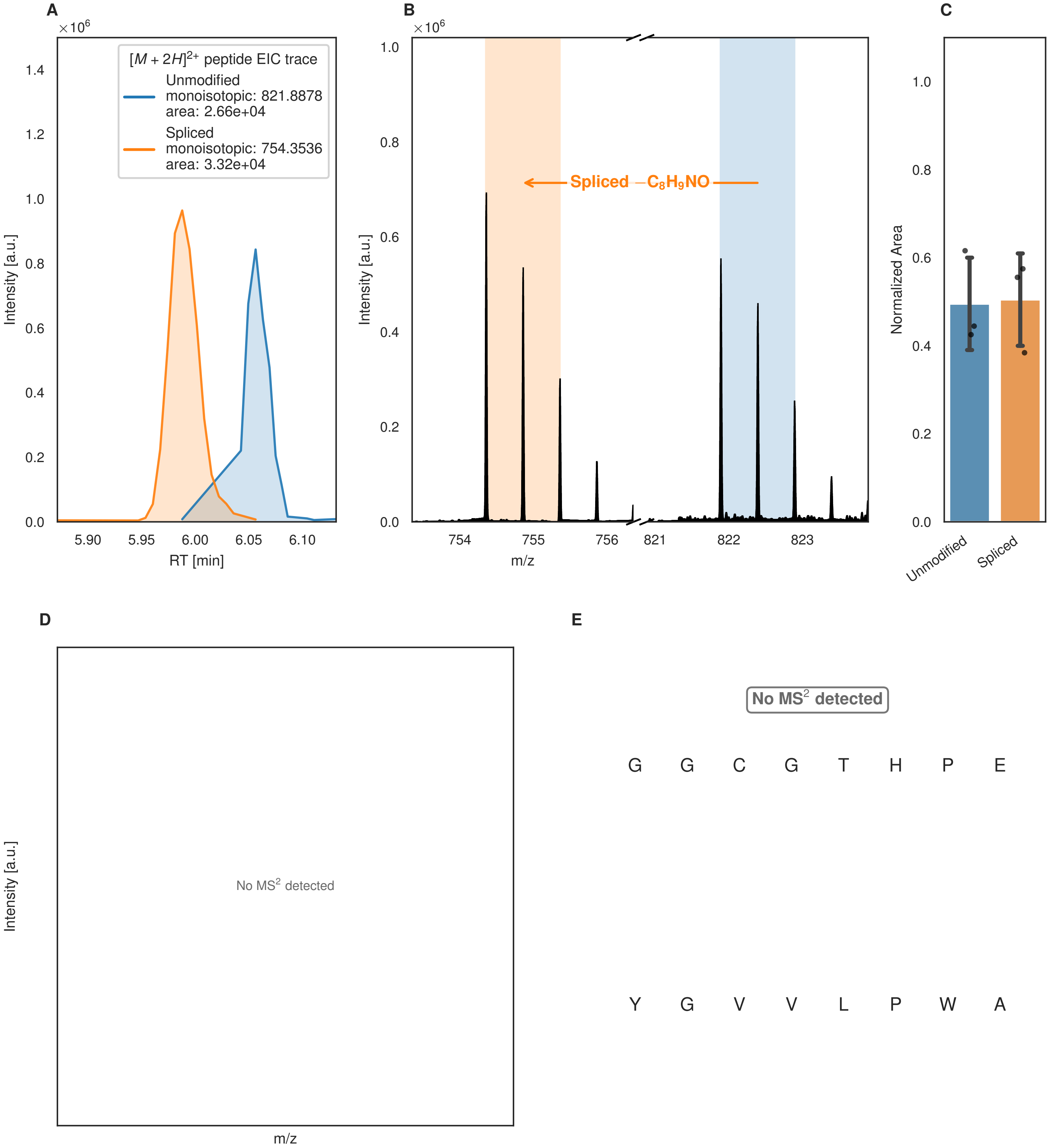
A
B
C
D
E
$\times10^6$
$[M+2H]^{2+}$ peptide EIC trace
Unmodified
monoisotopic: 821.8878
area: 2.66e+04
Spliced
monoisotopic: 754.3536
area: 3.32e+04
Intensity [a.u.]
RT [min]
Spliced $-C_8H_9NO$
m/z
Normalized Area
Unmodified
Spliced
No MS$^2$ detected
No MS$^2$ detected
G G C G T H P E
Y G V V L P W A

Figure S165 (figure continued from the previous page).

F

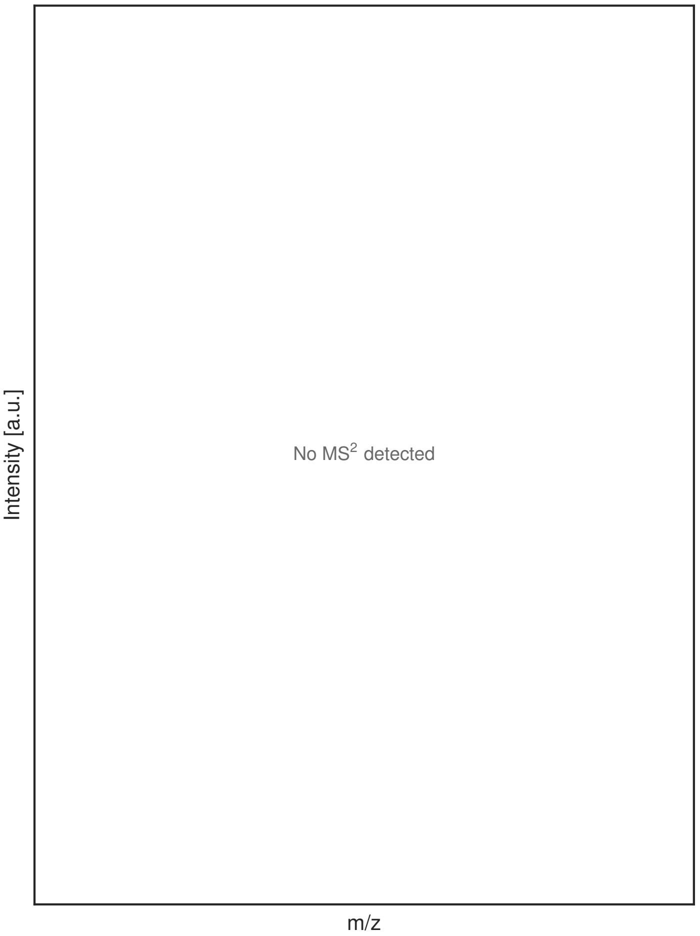


G

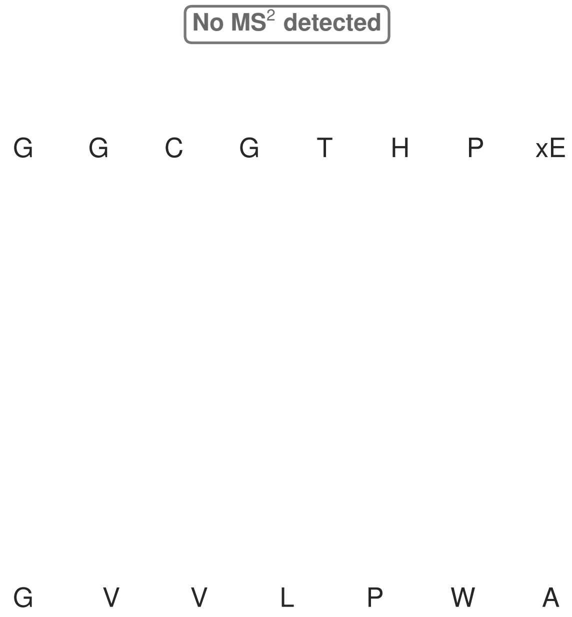


**Figure S165: Modification of the design_32 precursor peptide by PlpXY.** LC-MS analysis of IMAC-purified precursor peptides from *E. coli* BL21(DE3)Tuner containing splicease-encoding plasmid TJ-C-599 (PlpXY) and precursor peptide-encoding plasmid design_32 (RSTN-PlpA3-TEV-designed-32) after expression at 24 °C for 22 h in 1.5 mL cultures after digestion with TEV. Data from n=4 independent biological replicates. Spliced amino acids have an x as a prefix. Spliced refers to a single tyramine excision. A representative experimental replicate is shown in panels **A, B.**

**A** Close-up plot of the representative extracted ion chromatogram (EIC) of the various peptide states; the blue trace corresponds to the Unmodified peptide; the orange trace corresponds to the Spliced peptide. **B** Close-up plot of the m/z values of the peptides detected by the mass spectrometer, with the shaded colored areas corresponding to where the expected m/z values for the peptide m/z traces should occur, with the observed spectrum plotted in black. The Unmodified, Spliced peptide(s) were found in the mass spectrum. **C** Bar plot showing the relative abundance of the various peptide states at charge z=2; the bars represent the average relative abundances normalized to total peak areas, with each individual experimental replicate value plotted as a black dot. **D** Mirrored MS$^2$ plot for the Unmodified peptide. In this mirror plot, the full fragmentation spectrum is plotted in black pointing upward, and the matched fragment b ions (blue) and y ions (red) are plotted pointing downward, mirrored against it, so that each matched fragment can be compared directly with the measured spectrum. **E** MS$^2$ spectrum of the fragments corresponding to the Unmodified peptide sequence, with matched b (blue, above) and y ions (red, below) plotted. **F** Mirrored MS$^2$ plot for the Spliced peptide. In this mirror plot, the full fragmentation spectrum is plotted in black pointing upward, and the matched fragment b ions (blue) and y ions (red) are plotted pointing downward, mirrored against it, so that each matched fragment can be compared directly with the measured spectrum. **G** MS$^2$ spectrum of the fragments corresponding to the Spliced peptide sequence, with matched b (blue, above) and y ions (red, below) plotted.

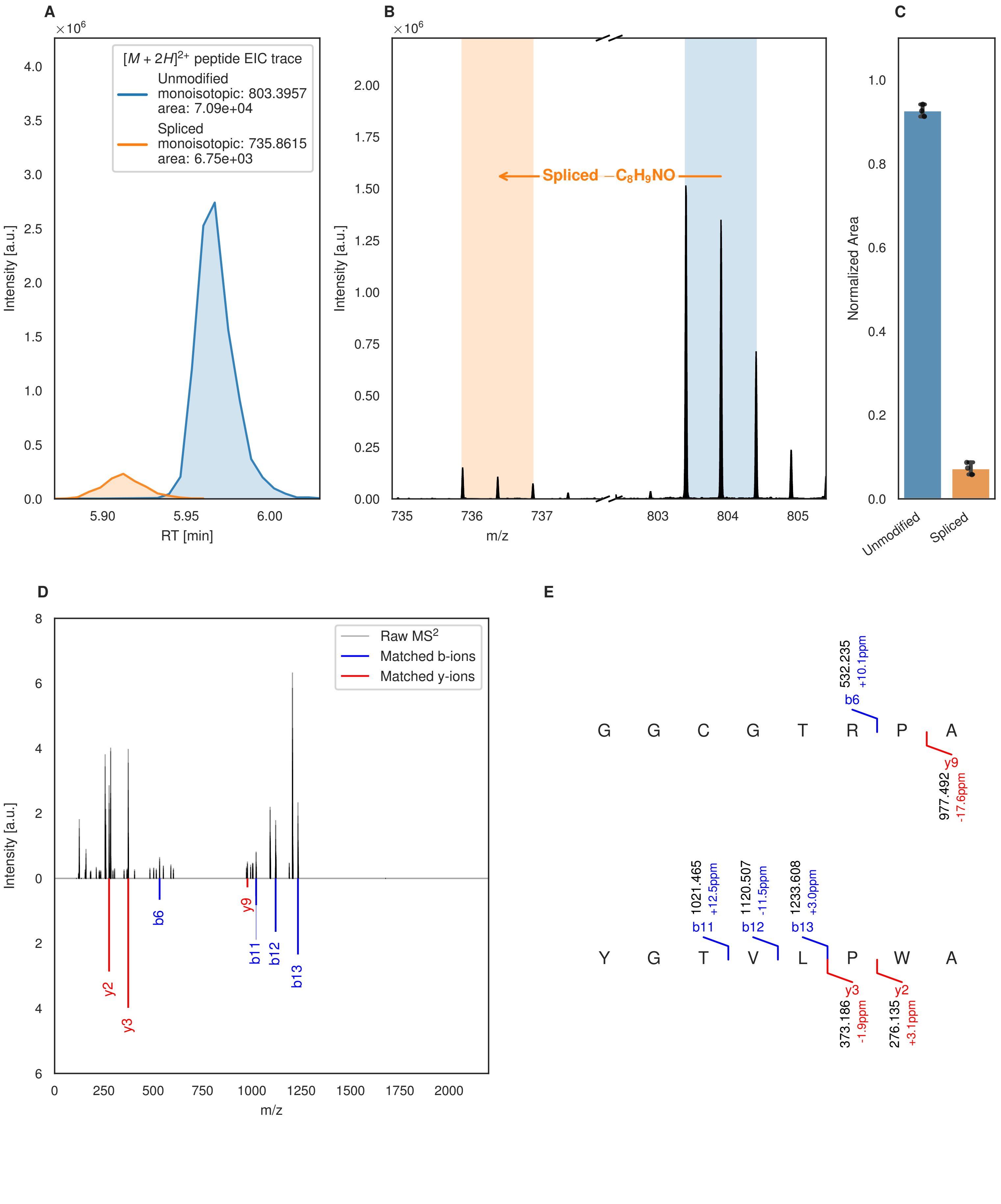

A
B
C
D
E
[M + 2H]2+ peptide EIC trace
Unmodified
monoisotopic: 803.3957
area: 7.09e+04
Spliced
monoisotopic: 735.8615
area: 6.75e+03
Intensity [a.u.]
RT [min]
Spliced −C8H9NO
m/z
Normalized Area
Unmodified
Spliced
Raw MS2
Matched b-ions
Matched y-ions
b6
b11
b12
b13
y2
y3
y9
G G C G T R P A
532.235
+10.1ppm
977.492
-17.6ppm
Y G T V L P W A
1021.465
+12.5ppm
1120.507
-11.5ppm
1233.608
+3.0ppm
373.186
-1.9ppm
276.135
+3.1ppm

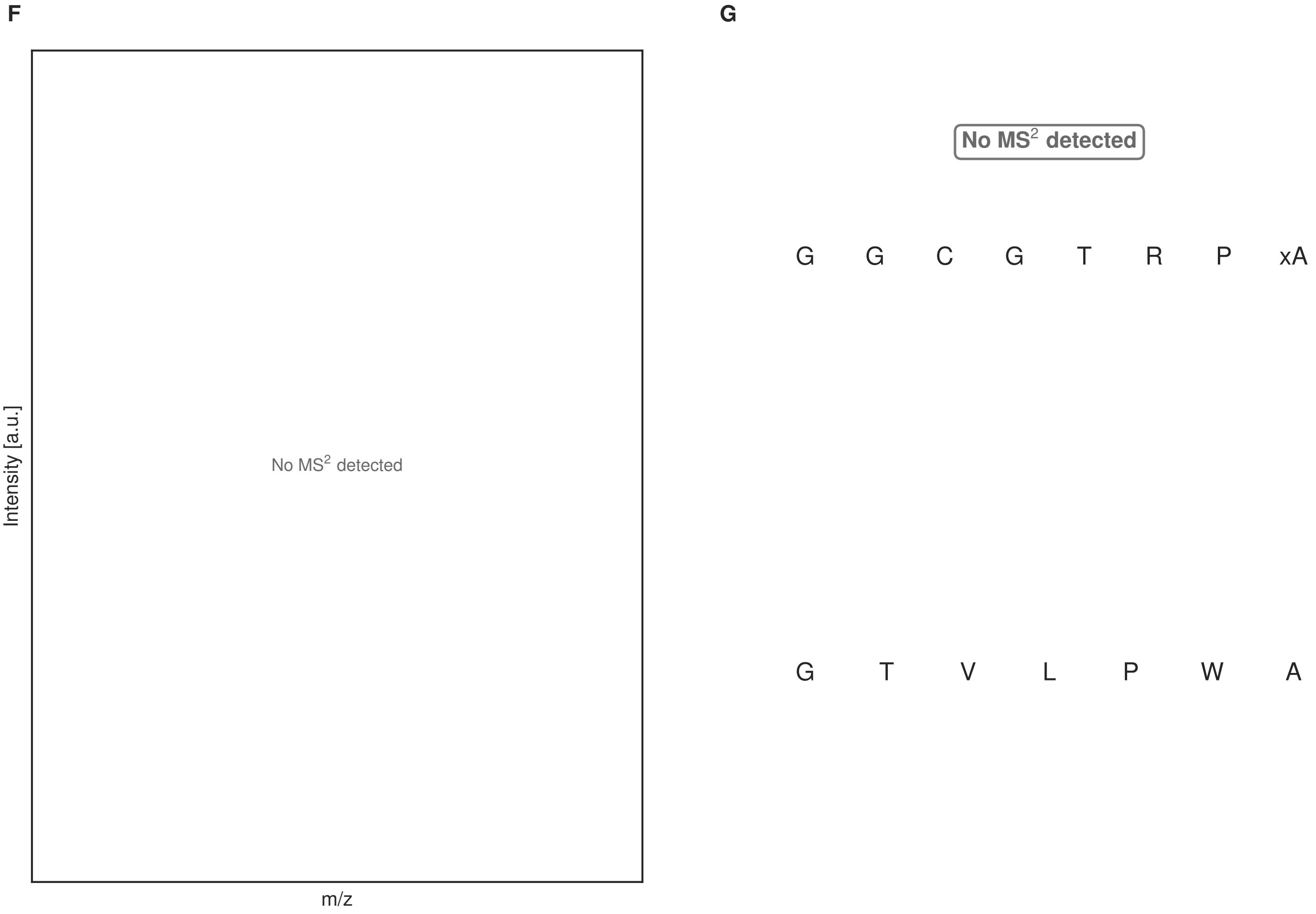


**Figure S166: Modification of the design_34 precursor peptide by PlpXY.** LC-MS analysis of IMAC-purified precursor peptides from *E. coli* BL21(DE3)Tuner containing splicease-encoding plasmid TJ-C-599 (PlpXY) and precursor peptide-encoding plasmid design_34 (RSTN-PlpA3-TEV-designed-34) after expression at 24 °C for 22 h in 1.5 mL cultures after digestion with TEV. Data from n=4 independent biological replicates. Spliced amino acids have an x as a prefix. Spliced refers to a single tyramine excision. A representative experimental replicate is shown in panels **A, B.**

**A** Close-up plot of the representative extracted ion chromatogram (EIC) of the various peptide states; the blue trace corresponds to the Unmodified peptide; the orange trace corresponds to the Spliced peptide. **B** Close-up plot of the m/z values of the peptides detected by the mass spectrometer, with the shaded colored areas corresponding to where the expected m/z values for the peptide m/z traces should occur, with the observed spectrum plotted in black. The Unmodified, Spliced peptide(s) were found in the mass spectrum. **C** Bar plot showing the relative abundance of the various peptide states at charge z=2; the bars represent the average relative abundances normalized to total peak areas, with each individual experimental replicate value plotted as a black dot. **D** Mirrored MS$^2$ plot for the Unmodified peptide. In this mirror plot, the full fragmentation spectrum is plotted in black pointing upward, and the matched fragment b ions (blue) and y ions (red) are plotted pointing downward, mirrored against it, so that each matched fragment can be compared directly with the measured spectrum. **E** MS$^2$ spectrum of the fragments corresponding to the Unmodified peptide sequence, with matched b (blue, above) and y ions (red, below) plotted. **F** Mirrored MS$^2$ plot for the Spliced peptide. In this mirror plot, the full fragmentation spectrum is plotted in black pointing upward, and the matched fragment b ions (blue) and y ions (red) are plotted pointing downward, mirrored against it, so that each matched fragment can be compared directly with the measured spectrum. **G** MS$^2$ spectrum of the fragments corresponding to the Spliced peptide sequence, with matched b (blue, above) and y ions (red, below) plotted.

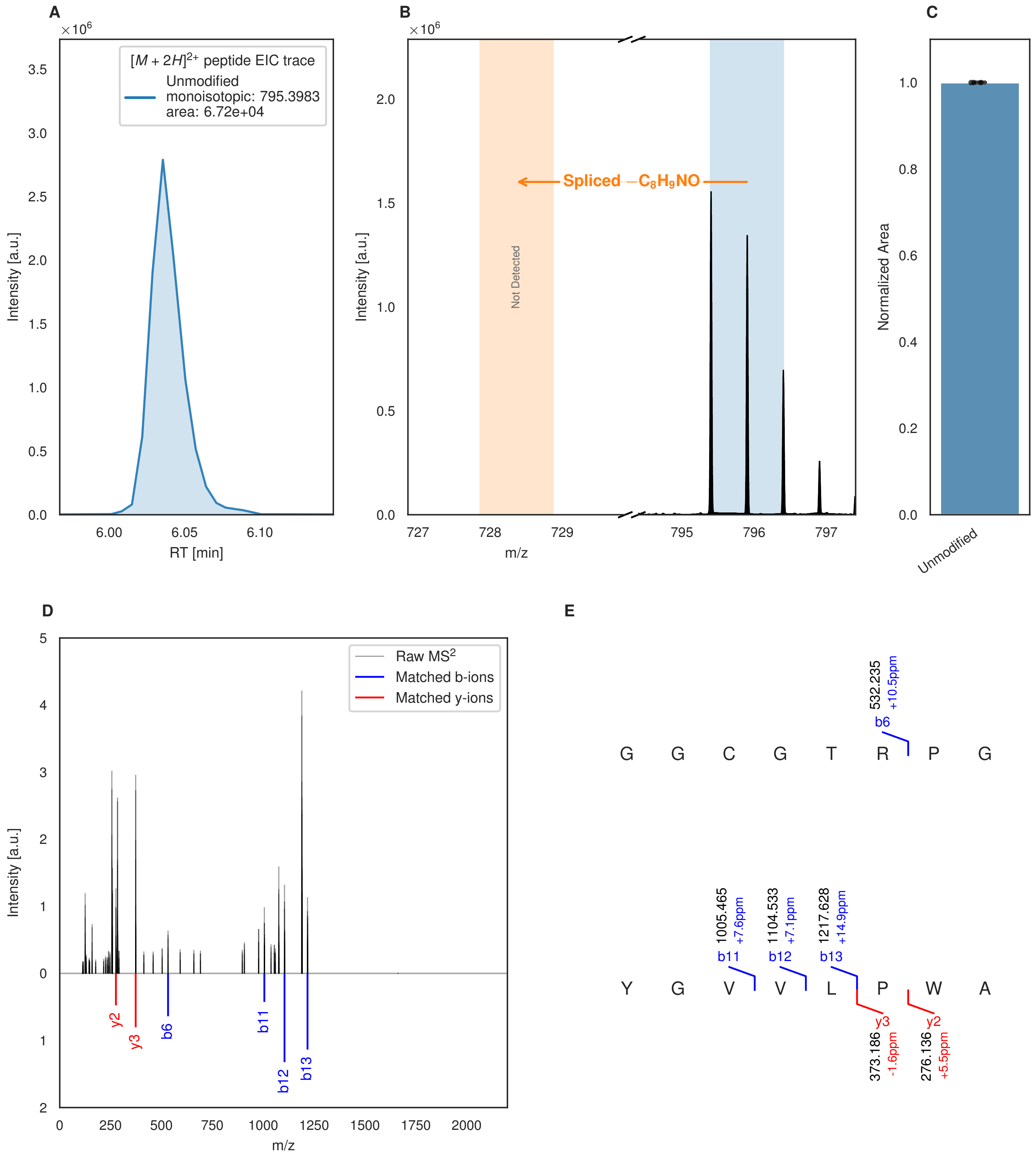


**Figure S167: Modification of the design_35 precursor peptide by PlpXY.** LC-MS analysis of IMAC-purified precursor peptides from *E. coli* BL21(DE3)Tuner containing splicease-encoding plasmid TJ-C-599 (PlpXY) and precursor peptide-encoding plasmid design_35 (RSTN-PlpA3-TEV-designed-35) after expression at 24 °C for 22 h in 1.5 mL cultures after digestion with TEV. Data from n=4 independent biological replicates. Spliced amino acids have an x as a prefix. Spliced refers to a single tyramine excision. A representative experimental replicate is shown in panels **A, B.**

**A** Close-up plot of the representative extracted ion chromatogram (EIC) of the various peptide states; the blue trace corresponds to the Unmodified peptide. **B** Close-up plot of the m/z values of the peptides detected by the mass spectrometer, with the shaded colored areas corresponding to where the expected m/z values for the peptide m/z

(Caption continued on the next page.)

traces should occur, with the observed spectrum plotted in black. The Unmodified peptide(s) were found in the mass spectrum and the Spliced peptide(s) were not detected. **C** Bar plot showing the relative abundance of the various peptide states at charge z=2; the bars represent the average relative abundances normalized to total peak areas, with each individual experimental replicate value plotted as a black dot. **D** Mirrored $MS^2$ plot for the Unmodified peptide. In this mirror plot, the full fragmentation spectrum is plotted in black pointing upward, and the matched fragment b ions (blue) and y ions (red) are plotted pointing downward, mirrored against it, so that each matched fragment can be compared directly with the measured spectrum. **E** $MS^2$ spectrum of the fragments corresponding to the Unmodified peptide sequence, with matched b (blue, above) and y ions (red, below) plotted.

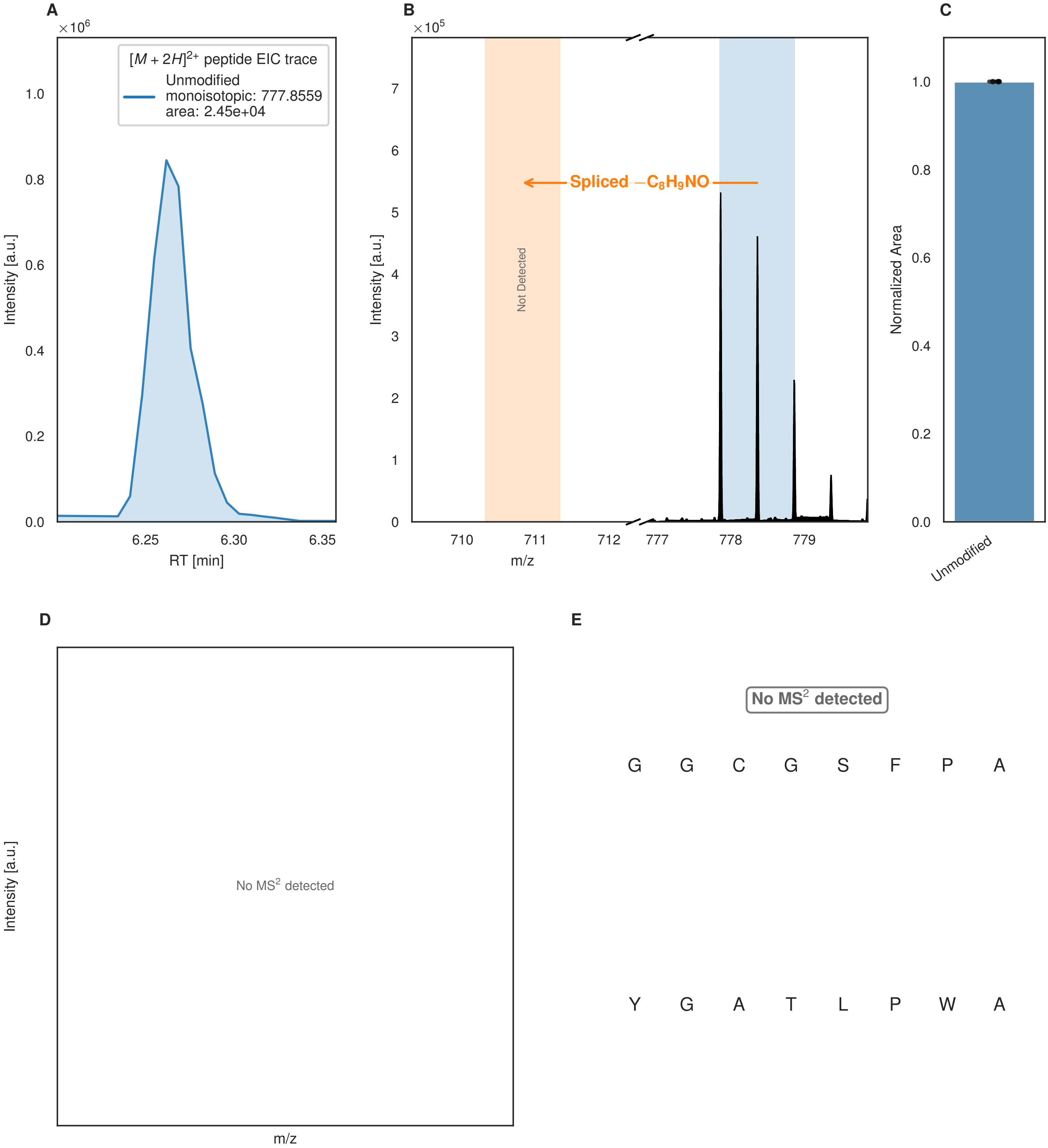


**Figure S168: Modification of the design_36 precursor peptide by PlpXY.** LC-MS analysis of IMAC-purified precursor peptides from *E. coli* BL21(DE3)Tuner containing splicease-encoding plasmid TJ-C-599 (PlpXY) and precursor peptide-encoding plasmid design_36 (RSTN-PlpA3-TEV-designed-36) after expression at 24 °C for 22 h in 1.5 mL cultures after digestion with TEV. Data from n=4 independent biological replicates. Spliced amino acids have an x as a prefix. Spliced refers to a single tyramine excision. A representative experimental replicate is shown in panels **A, B.**

**A** Close-up plot of the representative extracted ion chromatogram (EIC) of the various peptide states; the blue trace corresponds to the Unmodified peptide. **B** Close-up plot of the m/z values of the peptides detected by the mass spectrometer, with the shaded colored areas corresponding to where the expected m/z values for the peptide m/z

(Caption continued on the next page.)

traces should occur, with the observed spectrum plotted in black. The Unmodified peptide(s) were found in the mass spectrum and the Spliced peptide(s) were not detected. **C** Bar plot showing the relative abundance of the various peptide states at charge z=2; the bars represent the average relative abundances normalized to total peak areas, with each individual experimental replicate value plotted as a black dot. **D** Mirrored $MS^2$ plot for the Unmodified peptide. In this mirror plot, the full fragmentation spectrum is plotted in black pointing upward, and the matched fragment b ions (blue) and y ions (red) are plotted pointing downward, mirrored against it, so that each matched fragment can be compared directly with the measured spectrum. **E** $MS^2$ spectrum of the fragments corresponding to the Unmodified peptide sequence, with matched b (blue, above) and y ions (red, below) plotted.

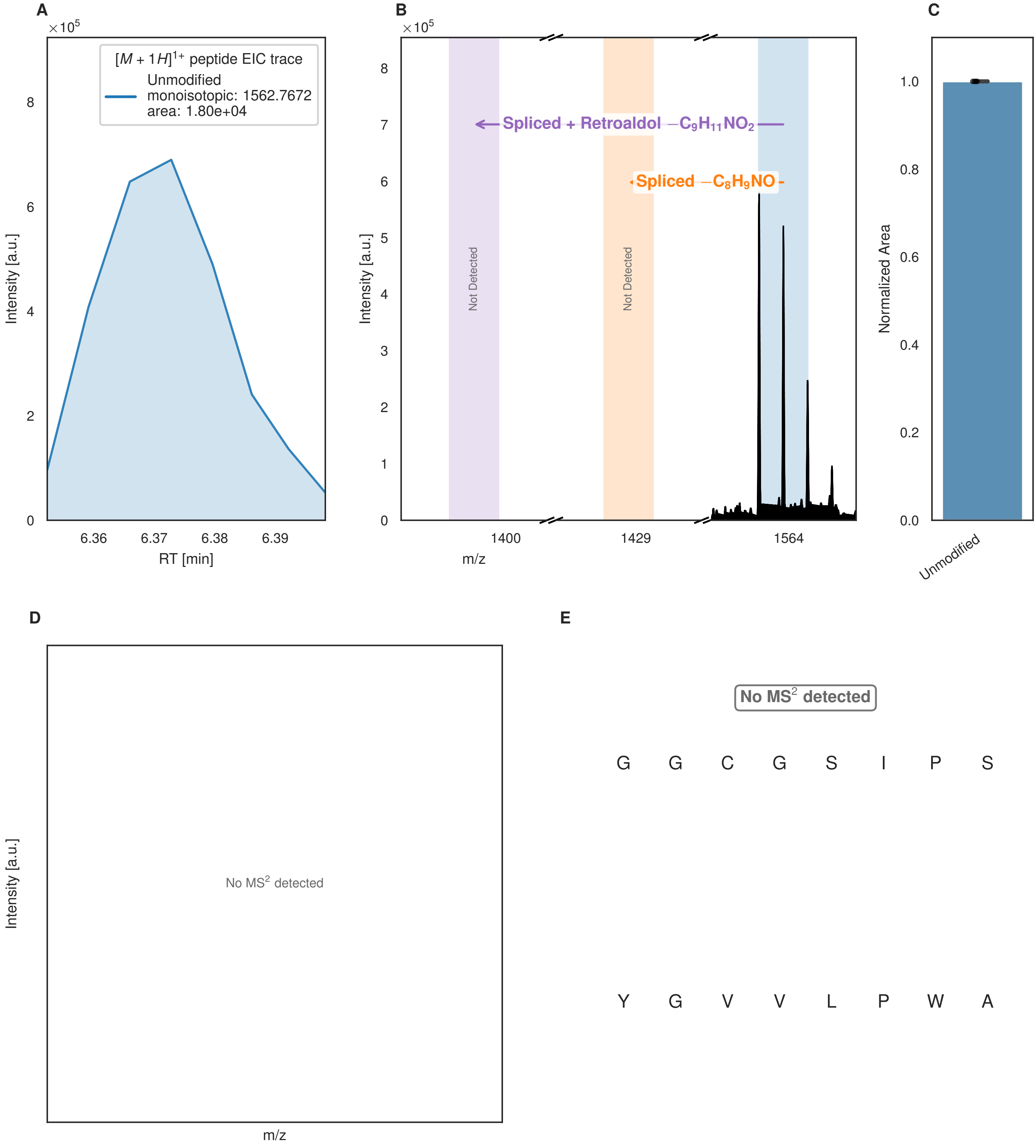


**Figure S169: Modification of the design_37 precursor peptide by PlpXY.** LC-MS analysis of IMAC-purified precursor peptides from *E. coli* BL21(DE3)Tuner containing splicease-encoding plasmid TJ-C-599 (PlpXY) and precursor peptide-encoding plasmid design_37 (RSTN-PlpA3-TEV-designed-37) after expression at 24 °C for 22 h in 1.5 mL cultures after digestion with TEV. Data from n=4 independent biological replicates. Spliced amino acids have an x as a prefix. Spliced refers to a single tyramine excision; Spliced + Retroaldol refers to a glycine-derived ketoamide (xG). A representative experimental replicate is shown in panels **A, B.**
**A** Close-up plot of the representative extracted ion chromatogram (EIC) of the various peptide states; the blue trace corresponds to the Unmodified peptide. **B** Close-up plot of the m/z values of the peptides detected by the mass spectrometer, with the shaded colored areas corresponding to where the expected m/z values for the peptide m/z

(Caption continued on the next page.)

traces should occur, with the observed spectrum plotted in black. The Unmodified peptide(s) were found in the mass spectrum and the Spliced, Spliced + Retroaldol peptide(s) were not detected. **C** Bar plot showing the relative abundance of the various peptide states at charge z=1; the bars represent the average relative abundances normalized to total peak areas, with each individual experimental replicate value plotted as a black dot. **D** Mirrored $MS^2$ plot for the Unmodified peptide. In this mirror plot, the full fragmentation spectrum is plotted in black pointing upward, and the matched fragment b ions (blue) and y ions (red) are plotted pointing downward, mirrored against it, so that each matched fragment can be compared directly with the measured spectrum. **E** $MS^2$ spectrum of the fragments corresponding to the Unmodified peptide sequence, with matched b (blue, above) and y ions (red, below) plotted.

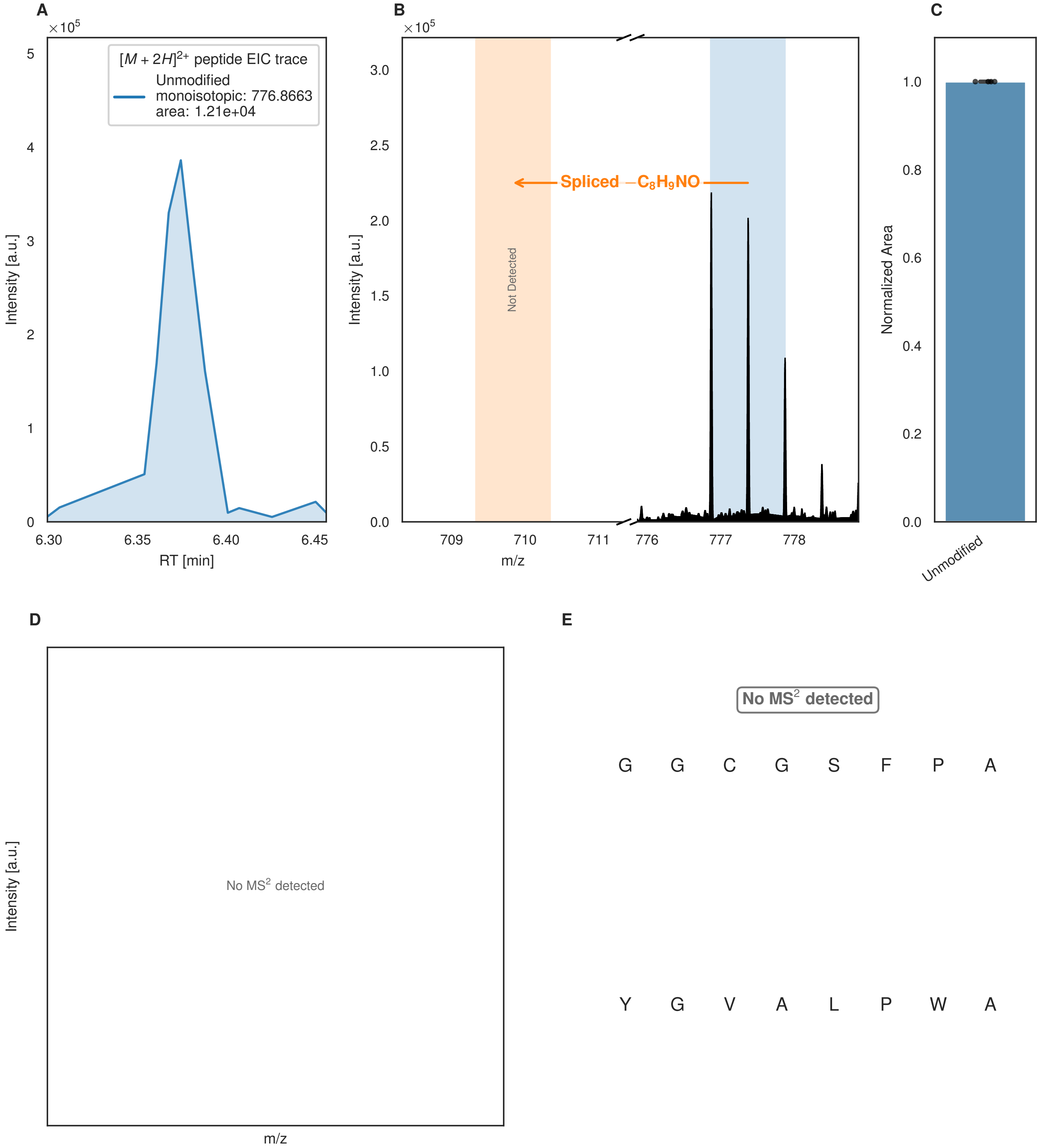


**Figure S170: Modification of the design_38 precursor peptide by PlpXY.** LC-MS analysis of IMAC-purified precursor peptides from *E. coli* BL21(DE3)Tuner containing splicease-encoding plasmid TJ-C-599 (PlpXY) and precursor peptide-encoding plasmid design_38 (RSTN-PlpA3-TEV-designed-38) after expression at 24 °C for 22 h in 1.5 mL cultures after digestion with TEV. Data from n=4 independent biological replicates. Spliced amino acids have an x as a prefix. Spliced refers to a single tyramine excision. A representative experimental replicate is shown in panels **A, B.**

**A** Close-up plot of the representative extracted ion chromatogram (EIC) of the various peptide states; the blue trace corresponds to the Unmodified peptide. **B** Close-up plot of the m/z values of the peptides detected by the mass spectrometer, with the shaded colored areas corresponding to where the expected m/z values for the peptide m/z

(Caption continued on the next page.)

traces should occur, with the observed spectrum plotted in black. The Unmodified peptide(s) were found in the mass spectrum and the Spliced peptide(s) were not detected. **C** Bar plot showing the relative abundance of the various peptide states at charge z=2; the bars represent the average relative abundances normalized to total peak areas, with each individual experimental replicate value plotted as a black dot. **D** Mirrored $MS^2$ plot for the Unmodified peptide. In this mirror plot, the full fragmentation spectrum is plotted in black pointing upward, and the matched fragment b ions (blue) and y ions (red) are plotted pointing downward, mirrored against it, so that each matched fragment can be compared directly with the measured spectrum. **E** $MS^2$ spectrum of the fragments corresponding to the Unmodified peptide sequence, with matched b (blue, above) and y ions (red, below) plotted.

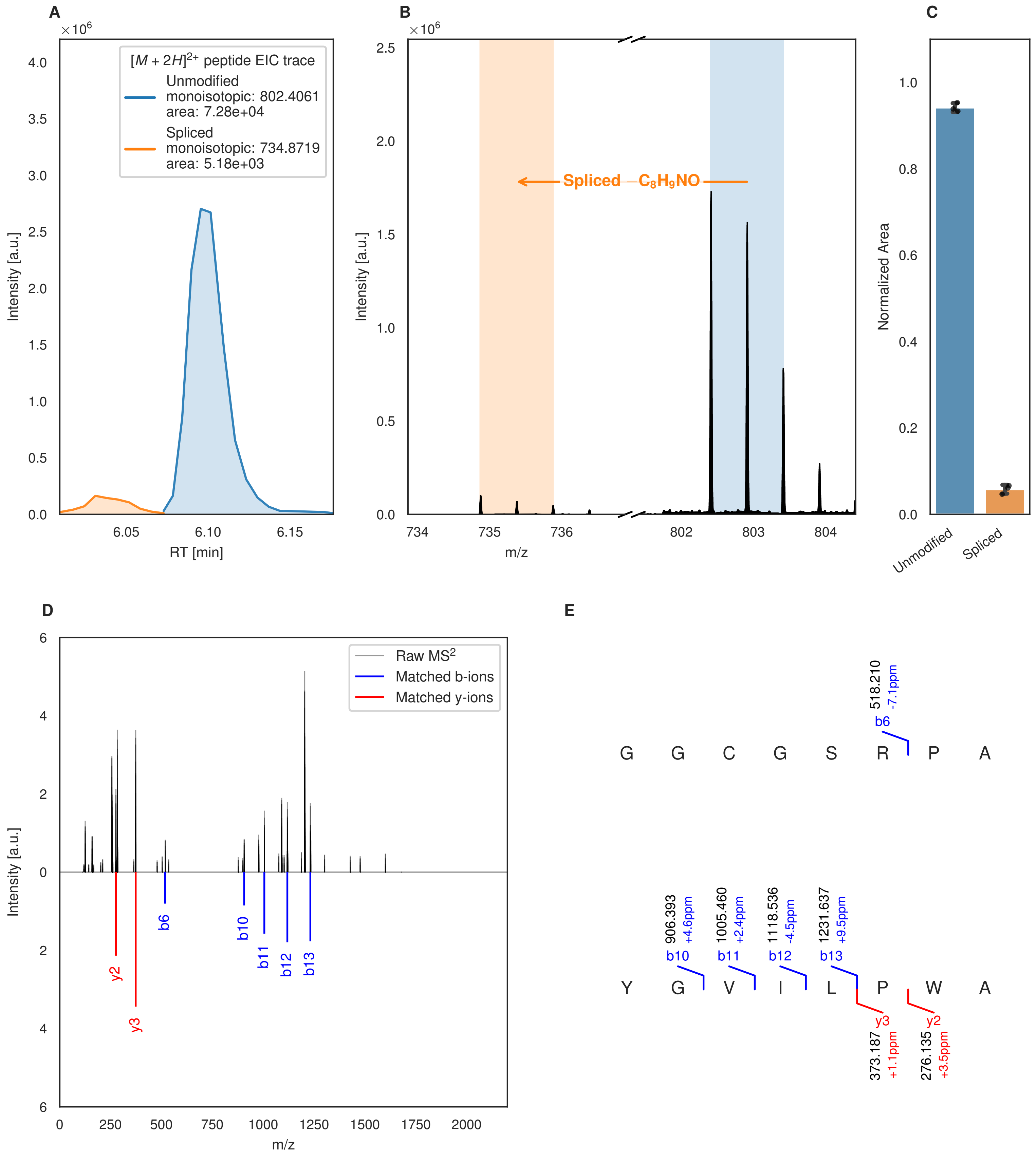

A
B
C
$[M+2H]^{2+}$ peptide EIC trace
Unmodified
monoisotopic: 802.4061
area: 7.28e+04
Spliced
monoisotopic: 734.8719
area: 5.18e+03
Intensity [a.u.]
RT [min]
Spliced −C8H9NO
m/z
Normalized Area
Unmodified
Spliced
D
E
Raw MS2
Matched b-ions
Matched y-ions
b6
b10
b11
b12
b13
y2
y3
G G C G S R P A
518.210
-7.1ppm
Y G V I L P W A
906.393
+4.6ppm
1005.460
+2.4ppm
1118.536
-4.5ppm
1231.637
+9.5ppm
373.187
+1.1ppm
276.135
+3.5ppm

Figure S171 (figure continued from the previous page).

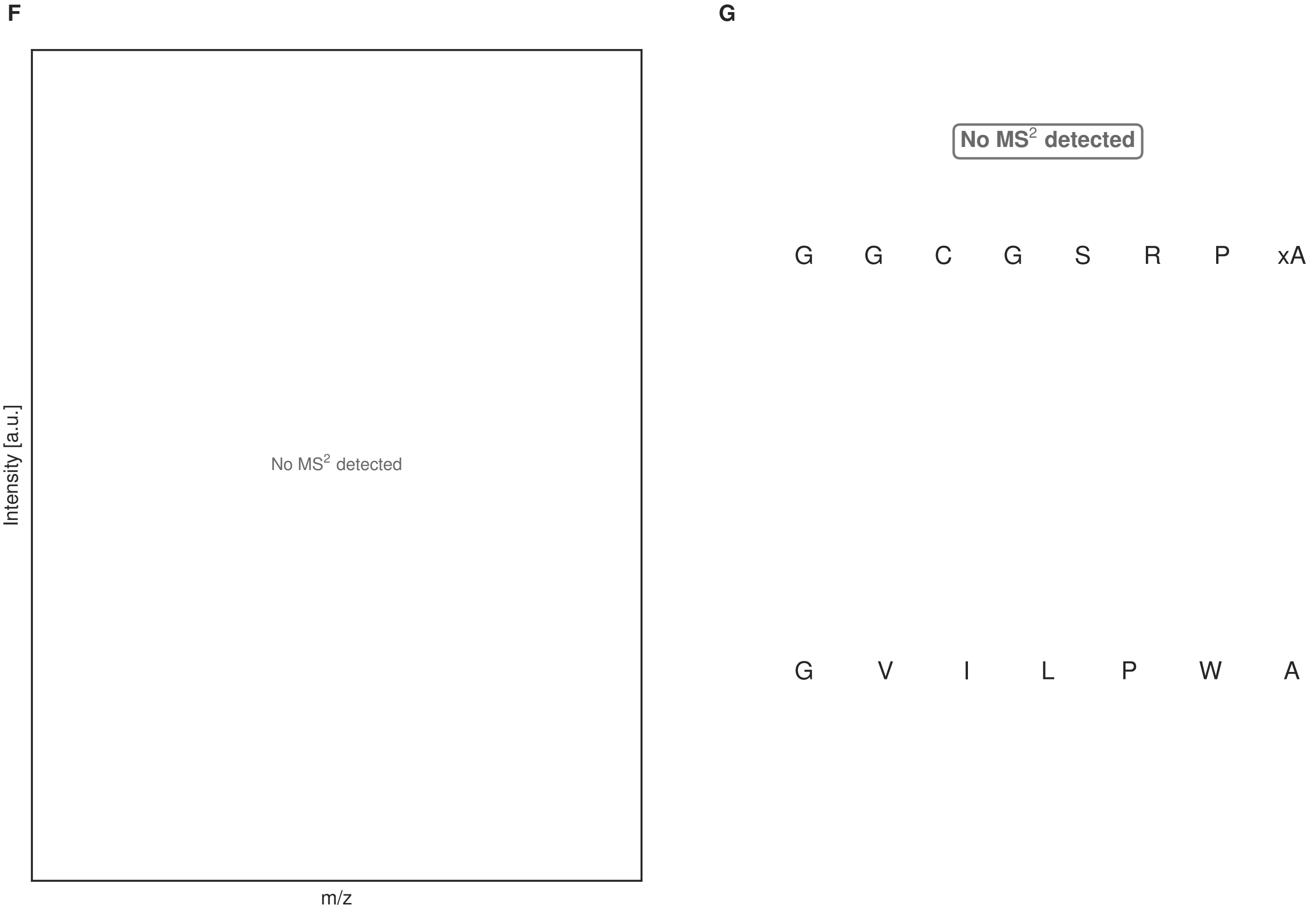


**Figure S171: Modification of the design_39 precursor peptide by PlpXY.** LC-MS analysis of IMAC-purified precursor peptides from *E. coli* BL21(DE3)Tuner containing splicease-encoding plasmid TJ-C-599 (PlpXY) and precursor peptide-encoding plasmid design_39 (RSTN-PlpA3-TEV-designed-39) after expression at 24 °C for 22 h in 1.5 mL cultures after digestion with TEV. Data from n=4 independent biological replicates. Spliced amino acids have an x as a prefix. Spliced refers to a single tyramine excision. A representative experimental replicate is shown in panels **A, B.**

**A** Close-up plot of the representative extracted ion chromatogram (EIC) of the various peptide states; the blue trace corresponds to the Unmodified peptide; the orange trace corresponds to the Spliced peptide. **B** Close-up plot of the m/z values of the peptides detected by the mass spectrometer, with the shaded colored areas corresponding to where the expected m/z values for the peptide m/z traces should occur, with the observed spectrum plotted in black. The Unmodified, Spliced peptide(s) were found in the mass spectrum. **C** Bar plot showing the relative abundance of the various peptide states at charge z=2; the bars represent the average relative abundances normalized to total peak areas, with each individual experimental replicate value plotted as a black dot. **D** Mirrored MS$^2$ plot for the Unmodified peptide. In this mirror plot, the full fragmentation spectrum is plotted in black pointing upward, and the matched fragment b ions (blue) and y ions (red) are plotted pointing downward, mirrored against it, so that each matched fragment can be compared directly with the measured spectrum. **E** MS$^2$ spectrum of the fragments corresponding to the Unmodified peptide sequence, with matched b (blue, above) and y ions (red, below) plotted. **F** Mirrored MS$^2$ plot for the Spliced peptide. In this mirror plot, the full fragmentation spectrum is plotted in black pointing upward, and the matched fragment b ions (blue) and y ions (red) are plotted pointing downward, mirrored against it, so that each matched fragment can be compared directly with the measured spectrum. **G** MS$^2$ spectrum of the fragments corresponding to the Spliced peptide sequence, with matched b (blue, above) and y ions (red, below) plotted.

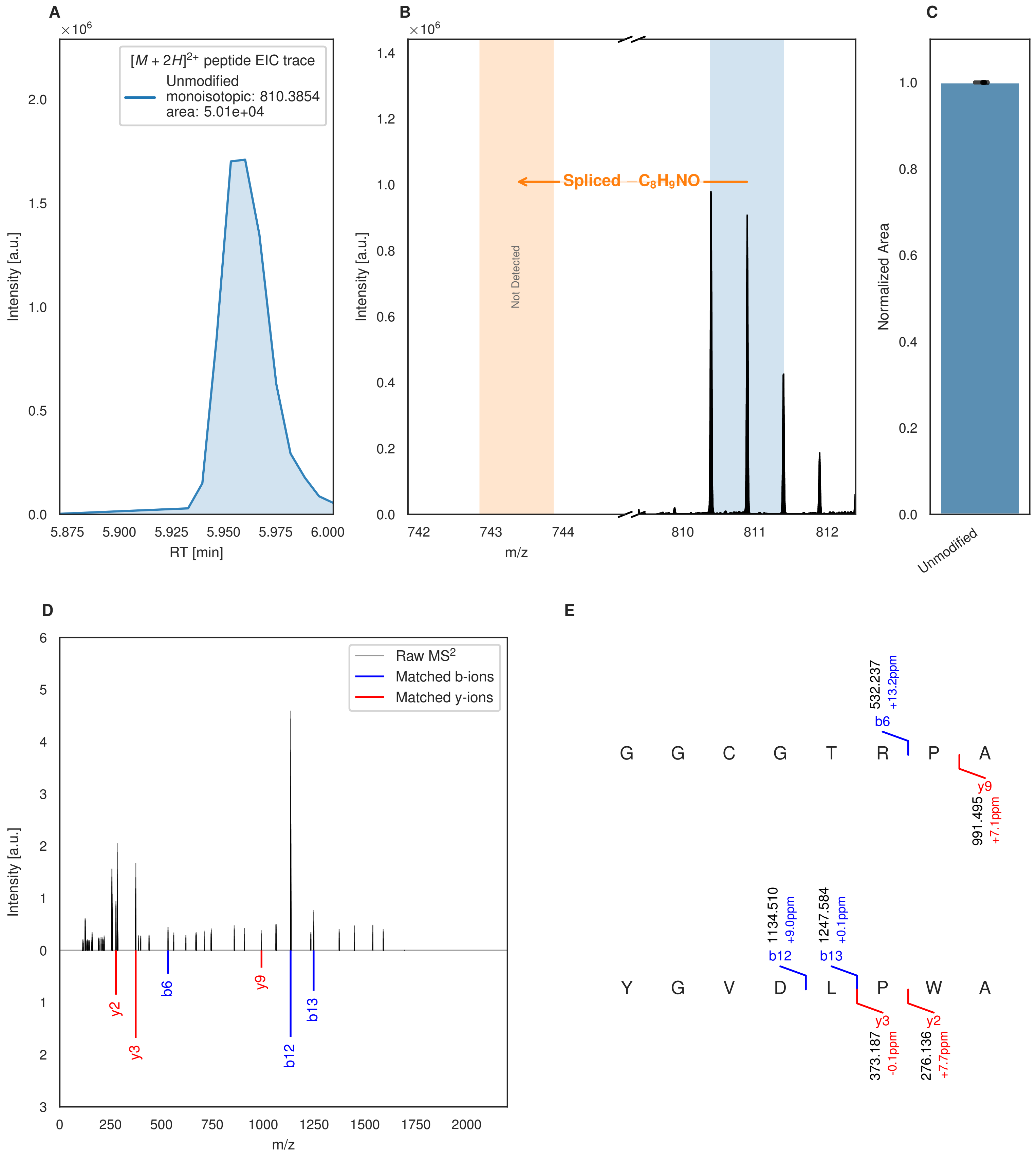


**Figure S172: Modification of the design_40 precursor peptide by PlpXY.** LC-MS analysis of IMAC-purified precursor peptides from *E. coli* BL21(DE3)Tuner containing splicease-encoding plasmid TJ-C-599 (PlpXY) and precursor peptide-encoding plasmid design_40 (RSTN-PlpA3-TEV-designed-40) after expression at 24 °C for 22 h in 1.5 mL cultures after digestion with TEV. Data from n=4 independent biological replicates. Spliced amino acids have an x as a prefix. Spliced refers to a single tyramine excision. A representative experimental replicate is shown in panels **A, B.**

**A** Close-up plot of the representative extracted ion chromatogram (EIC) of the various peptide states; the blue trace corresponds to the Unmodified peptide. **B** Close-up plot of the m/z values of the peptides detected by the mass spectrometer, with the shaded colored areas corresponding to where the expected m/z values for the peptide m/z

(Caption continued on the next page.)

traces should occur, with the observed spectrum plotted in black. The Unmodified peptide(s) were found in the mass spectrum and the Spliced peptide(s) were not detected. **C** Bar plot showing the relative abundance of the various peptide states at charge z=2; the bars represent the average relative abundances normalized to total peak areas, with each individual experimental replicate value plotted as a black dot. **D** Mirrored $MS^2$ plot for the Unmodified peptide. In this mirror plot, the full fragmentation spectrum is plotted in black pointing upward, and the matched fragment b ions (blue) and y ions (red) are plotted pointing downward, mirrored against it, so that each matched fragment can be compared directly with the measured spectrum. **E** $MS^2$ spectrum of the fragments corresponding to the Unmodified peptide sequence, with matched b (blue, above) and y ions (red, below) plotted.

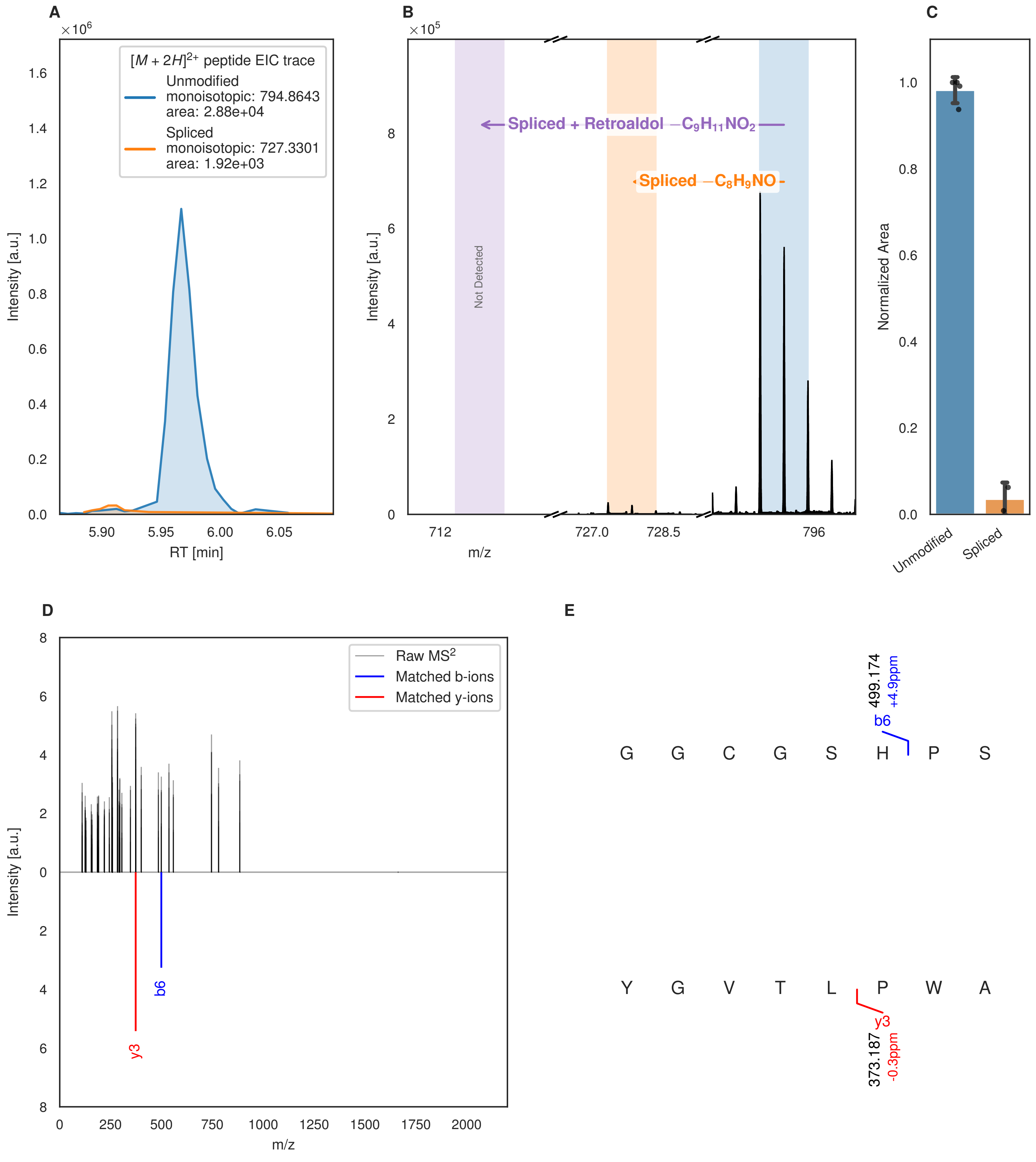

A
B
C
D
E
$[M+2H]^{2+}$ peptide EIC trace
Unmodified
monoisotopic: 794.8643
area: 2.88e+04
Spliced
monoisotopic: 727.3301
area: 1.92e+03
Intensity [a.u.]
RT [min]
Spliced + Retroaldol $-C_9H_{11}NO_2$
Spliced $-C_8H_9NO$
Not Detected
m/z
Normalized Area
Unmodified
Spliced
Raw MS$^2$
Matched b-ions
Matched y-ions
b6
y3
499.174
+4.9ppm
G G C G S H P S
Y G V T L P W A
373.187
-0.3ppm

Figure S173 (figure continued from the previous page).

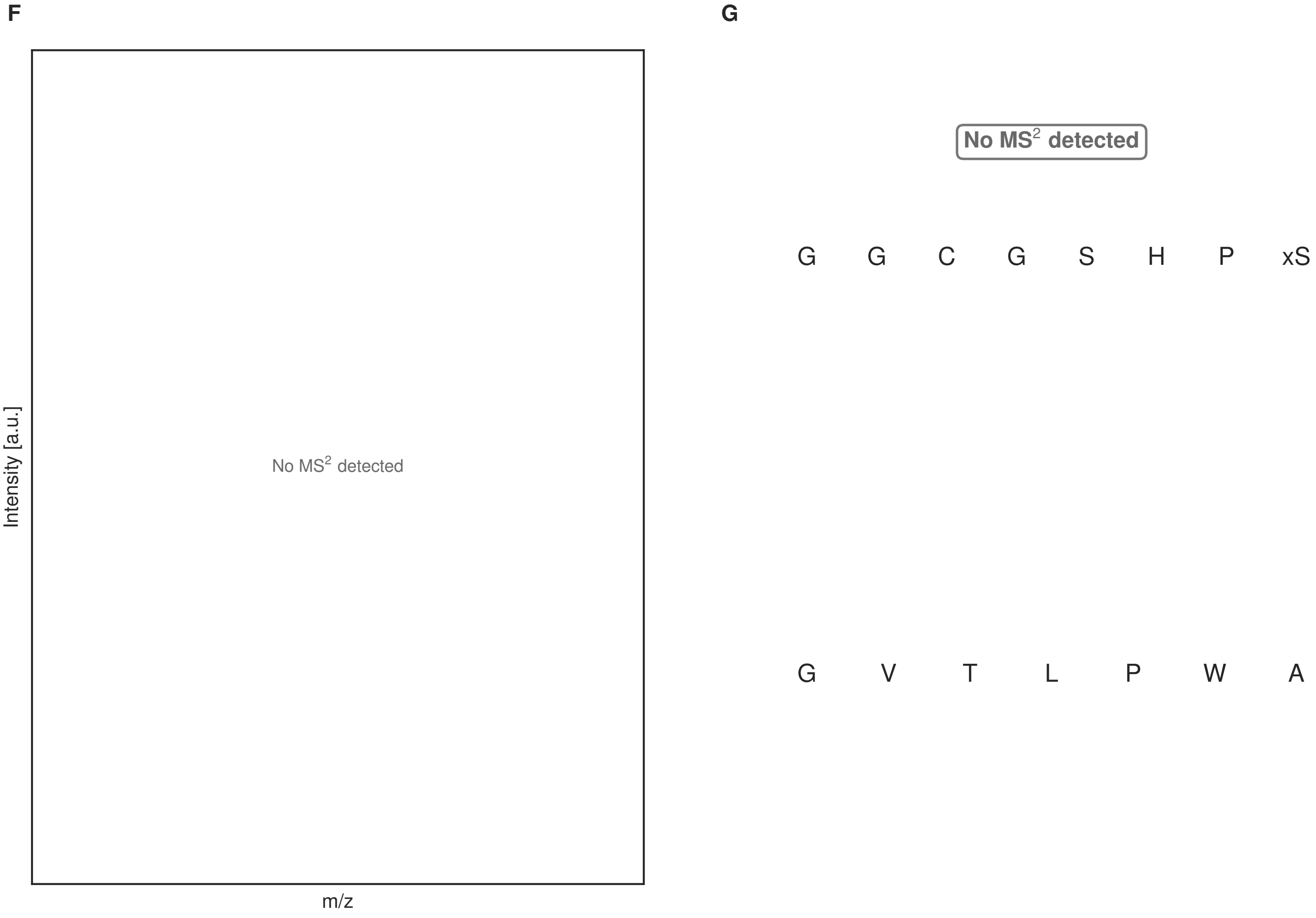


**Figure S173: Modification of the design_41 precursor peptide by PlpXY.** LC-MS analysis of IMAC-purified precursor peptides from *E. coli* BL21(DE3)Tuner containing splicease-encoding plasmid TJ-C-599 (PlpXY) and precursor peptide-encoding plasmid design_41 (RSTN-PlpA3-TEV-designed-41) after expression at 24 °C for 22 h in 1.5 mL cultures after digestion with TEV. Data from n=4 independent biological replicates. Spliced amino acids have an x as a prefix. Spliced refers to a single tyramine excision; Spliced + Retroaldol refers to a glycine-derived ketoamide (xG). A representative experimental replicate is shown in panels **A, B.**

**A** Close-up plot of the representative extracted ion chromatogram (EIC) of the various peptide states; the blue trace corresponds to the Unmodified peptide; the orange trace corresponds to the Spliced peptide. **B** Close-up plot of the m/z values of the peptides detected by the mass spectrometer, with the shaded colored areas corresponding to where the expected m/z values for the peptide m/z traces should occur, with the observed spectrum plotted in black. The Unmodified, Spliced peptide(s) were found in the mass spectrum and the Spliced + Retroaldol peptide(s) were not detected. **C** Bar plot showing the relative abundance of the various peptide states at charge z=2; the bars represent the average relative abundances normalized to total peak areas, with each individual experimental replicate value plotted as a black dot. **D** Mirrored $MS^2$ plot for the Unmodified peptide. In this mirror plot, the full fragmentation spectrum is plotted in black pointing upward, and the matched fragment b ions (blue) and y ions (red) are plotted pointing downward, mirrored against it, so that each matched fragment can be compared directly with the measured spectrum. **E** $MS^2$ spectrum of the fragments corresponding to the Unmodified peptide sequence, with matched b (blue, above) and y ions (red, below) plotted. **F** Mirrored $MS^2$ plot for the Spliced peptide. In this mirror plot, the full fragmentation spectrum is plotted in black pointing upward, and the matched fragment b ions (blue) and y ions (red) are plotted pointing downward, mirrored against it, so that each matched fragment can be compared directly with the measured spectrum. **G** $MS^2$ spectrum of the fragments corresponding to the Spliced peptide sequence, with matched b (blue, above) and y ions (red, below) plotted.

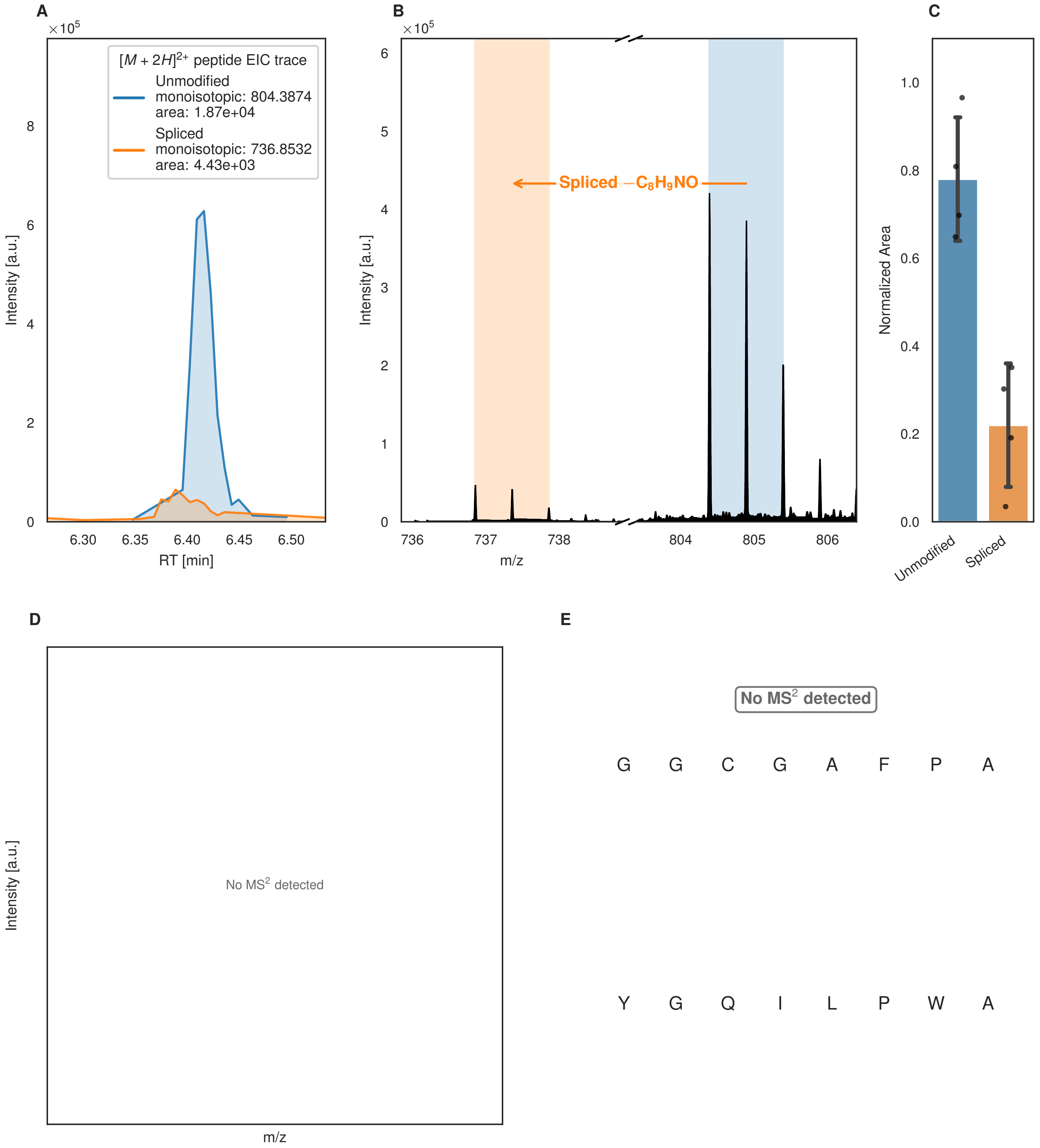
A
×10^5
$[M+2H]^{2+}$ peptide EIC trace
Unmodified
monoisotopic: 804.3874
area: 1.87e+04
Spliced
monoisotopic: 736.8532
area: 4.43e+03
Intensity [a.u.]
0
2
4
6
8
6.30
6.35
6.40
6.45
6.50
RT [min]
B
×10^5
Spliced $-C_8H_9NO$
Intensity [a.u.]
0
1
2
3
4
5
6
736
737
738
804
805
806
m/z
C
Normalized Area
0.0
0.2
0.4
0.6
0.8
1.0
Unmodified
Spliced
D
Intensity [a.u.]
No MS$^2$ detected
m/z
E
No MS$^2$ detected
G G C G A F P A
Y G Q I L P W A

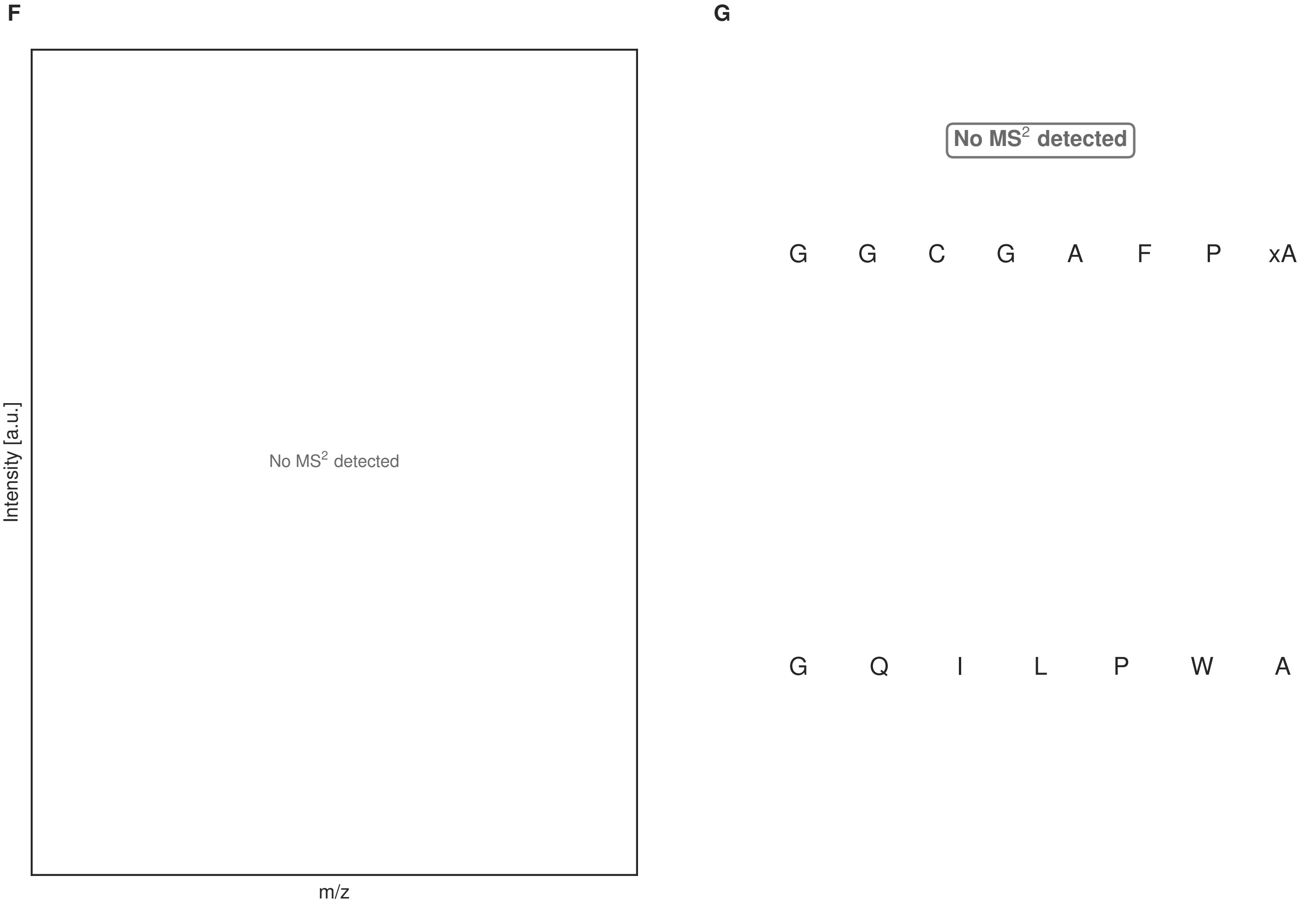


**Figure S174: Modification of the design_42 precursor peptide by PlpXY.** LC-MS analysis of IMAC-purified precursor peptides from *E. coli* BL21(DE3)Tuner containing splicease-encoding plasmid TJ-C-599 (PlpXY) and precursor peptide-encoding plasmid design_42 (RSTN-PlpA3-TEV-designed-42) after expression at 24 °C for 22 h in 1.5 mL cultures after digestion with TEV. Data from n=4 independent biological replicates. Spliced amino acids have an x as a prefix. Spliced refers to a single tyramine excision. A representative experimental replicate is shown in panels **A, B.**

**A** Close-up plot of the representative extracted ion chromatogram (EIC) of the various peptide states; the blue trace corresponds to the Unmodified peptide; the orange trace corresponds to the Spliced peptide. **B** Close-up plot of the m/z values of the peptides detected by the mass spectrometer, with the shaded colored areas corresponding to where the expected m/z values for the peptide m/z traces should occur, with the observed spectrum plotted in black. The Unmodified, Spliced peptide(s) were found in the mass spectrum. **C** Bar plot showing the relative abundance of the various peptide states at charge z=2; the bars represent the average relative abundances normalized to total peak areas, with each individual experimental replicate value plotted as a black dot. **D** Mirrored $MS^2$ plot for the Unmodified peptide. In this mirror plot, the full fragmentation spectrum is plotted in black pointing upward, and the matched fragment b ions (blue) and y ions (red) are plotted pointing downward, mirrored against it, so that each matched fragment can be compared directly with the measured spectrum. **E** $MS^2$ spectrum of the fragments corresponding to the Unmodified peptide sequence, with matched b (blue, above) and y ions (red, below) plotted. **F** Mirrored $MS^2$ plot for the Spliced peptide. In this mirror plot, the full fragmentation spectrum is plotted in black pointing upward, and the matched fragment b ions (blue) and y ions (red) are plotted pointing downward, mirrored against it, so that each matched fragment can be compared directly with the measured spectrum. **G** $MS^2$ spectrum of the fragments corresponding to the Spliced peptide sequence, with matched b (blue, above) and y ions (red, below) plotted.

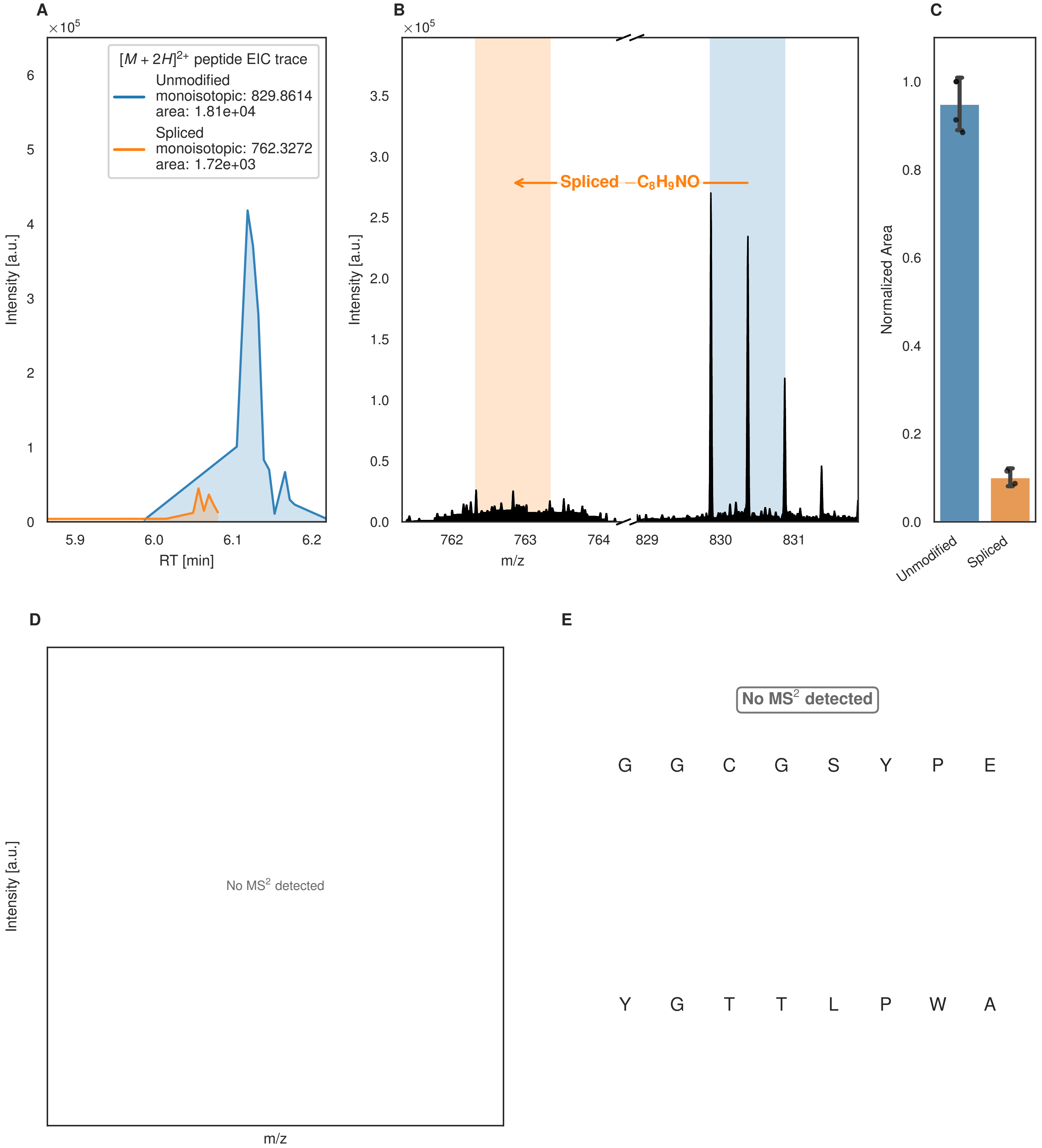
A
B
C
D
E
×10^5
$[M+2H]^{2+}$ peptide EIC trace
Unmodified
monoisotopic: 829.8614
area: 1.81e+04
Spliced
monoisotopic: 762.3272
area: 1.72e+03
Intensity [a.u.]
RT [min]
Spliced −$C_8H_9NO$
m/z
Normalized Area
Unmodified
Spliced
No MS² detected
G G C G S Y P E
Y G T T L P W A

F

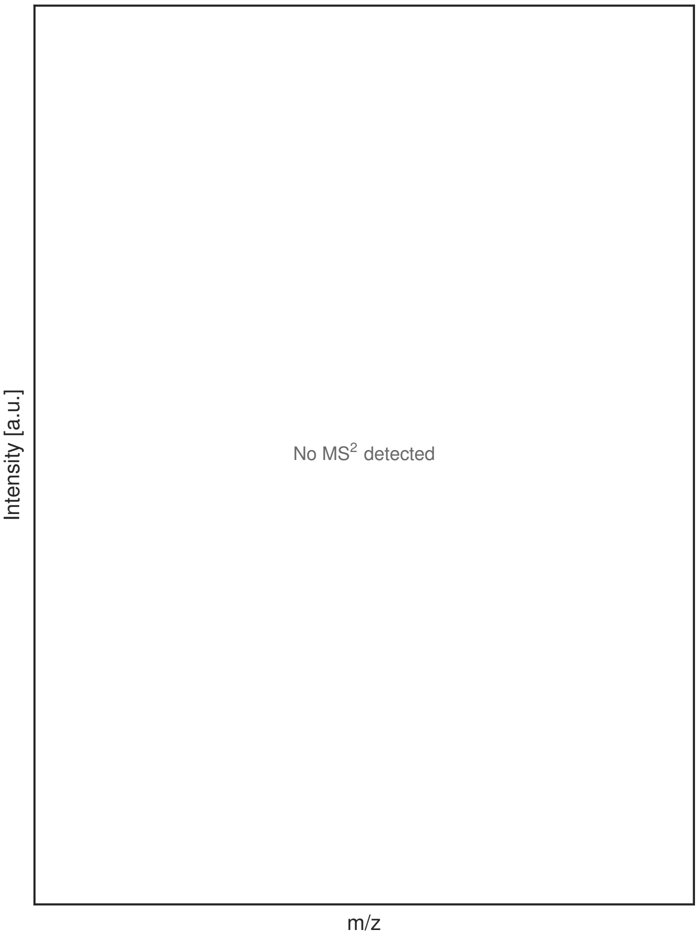


G

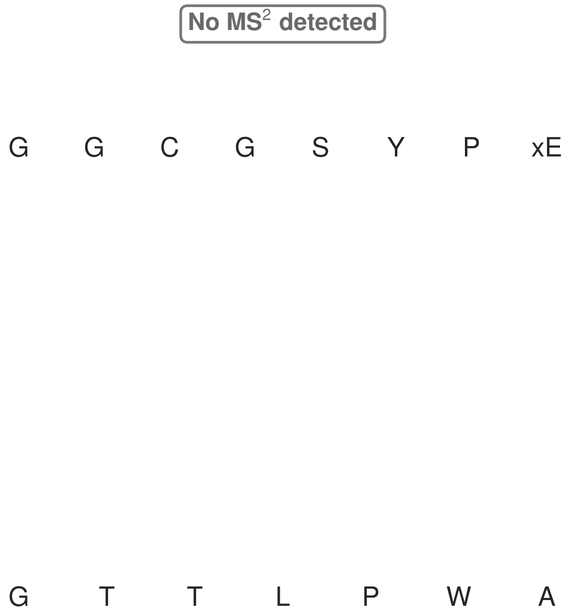


**Figure S175: Modification of the design_43 precursor peptide by PlpXY.** LC-MS analysis of IMAC-purified precursor peptides from *E. coli* BL21(DE3)Tuner containing splicease-encoding plasmid TJ-C-599 (PlpXY) and precursor peptide-encoding plasmid design_43 (RSTN-PlpA3-TEV-designed-43) after expression at 24 °C for 22 h in 1.5 mL cultures after digestion with TEV. Data from n=4 independent biological replicates. Spliced amino acids have an x as a prefix. Spliced refers to a single tyramine excision. A representative experimental replicate is shown in panels **A, B.**

**A** Close-up plot of the representative extracted ion chromatogram (EIC) of the various peptide states; the blue trace corresponds to the Unmodified peptide; the orange trace corresponds to the Spliced peptide. **B** Close-up plot of the m/z values of the peptides detected by the mass spectrometer, with the shaded colored areas corresponding to where the expected m/z values for the peptide m/z traces should occur, with the observed spectrum plotted in black. The Unmodified, Spliced peptide(s) were found in the mass spectrum. **C** Bar plot showing the relative abundance of the various peptide states at charge z=2; the bars represent the average relative abundances normalized to total peak areas, with each individual experimental replicate value plotted as a black dot. **D** Mirrored $MS^2$ plot for the Unmodified peptide. In this mirror plot, the full fragmentation spectrum is plotted in black pointing upward, and the matched fragment b ions (blue) and y ions (red) are plotted pointing downward, mirrored against it, so that each matched fragment can be compared directly with the measured spectrum. **E** $MS^2$ spectrum of the fragments corresponding to the Unmodified peptide sequence, with matched b (blue, above) and y ions (red, below) plotted. **F** Mirrored $MS^2$ plot for the Spliced peptide. In this mirror plot, the full fragmentation spectrum is plotted in black pointing upward, and the matched fragment b ions (blue) and y ions (red) are plotted pointing downward, mirrored against it, so that each matched fragment can be compared directly with the measured spectrum. **G** $MS^2$ spectrum of the fragments corresponding to the Spliced peptide sequence, with matched b (blue, above) and y ions (red, below) plotted.

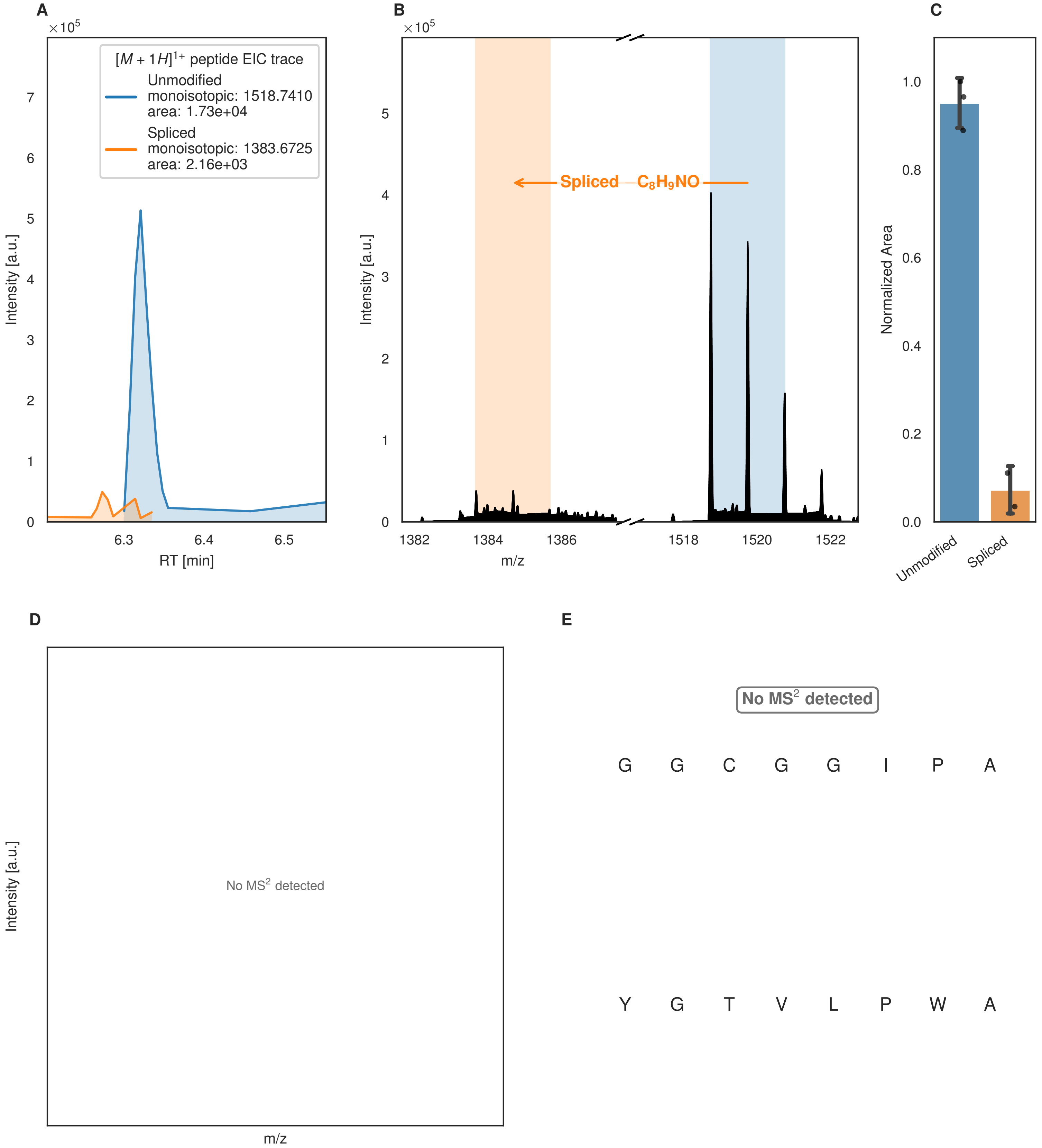

A
×10^5
[M + 1H]^1+ peptide EIC trace
Unmodified
monoisotopic: 1518.7410
area: 1.73e+04
Spliced
monoisotopic: 1383.6725
area: 2.16e+03
Intensity [a.u.]
RT [min]
B
×10^5
Spliced −C8H9NO
Intensity [a.u.]
m/z
C
Normalized Area
Unmodified
Spliced
D
Intensity [a.u.]
No MS2 detected
m/z
E
No MS2 detected
G G C G G I P A
Y G T V L P W A

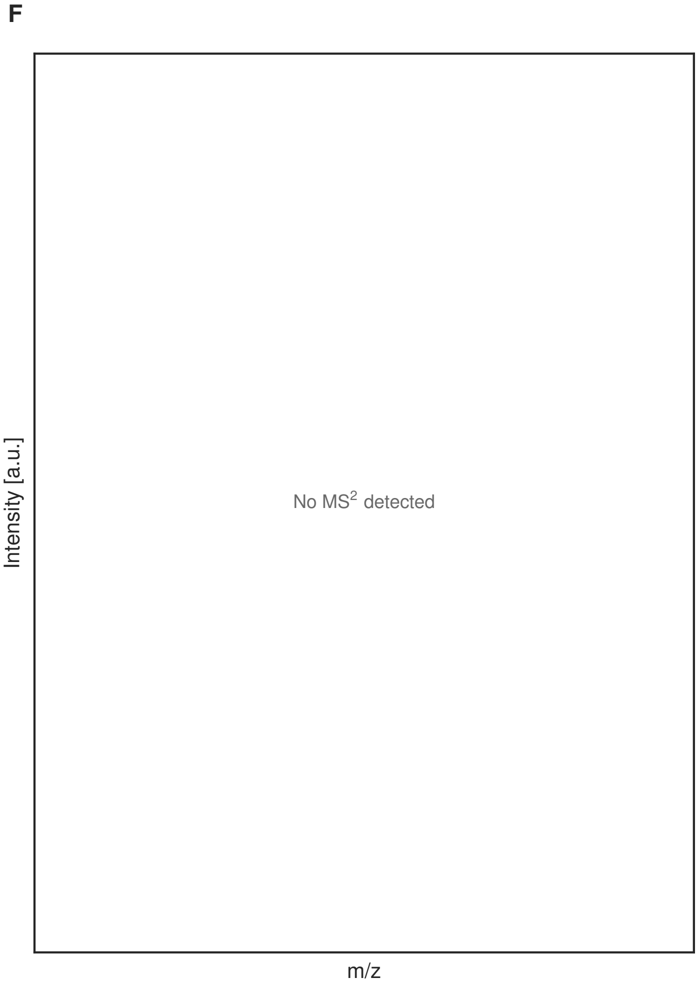


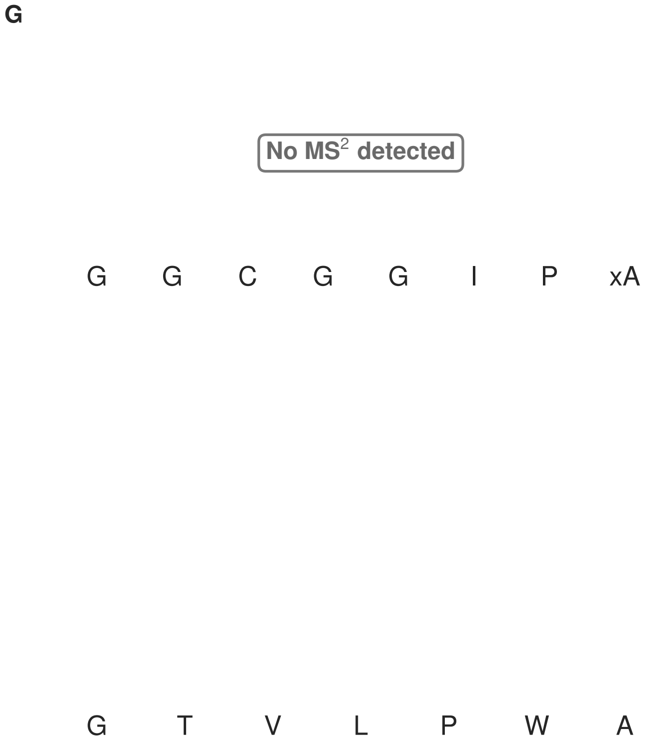


**Figure S176: Modification of the design_44 precursor peptide by PlpXY.** LC-MS analysis of IMAC-purified precursor peptides from *E. coli* BL21(DE3)Tuner containing splicease-encoding plasmid TJ-C-599 (PlpXY) and precursor peptide-encoding plasmid design_44 (RSTN-PlpA3-TEV-designed-44) after expression at 24 °C for 22 h in 1.5 mL cultures after digestion with TEV. Data from n=4 independent biological replicates. Spliced amino acids have an x as a prefix. Spliced refers to a single tyramine excision. A representative experimental replicate is shown in panels **A, B.**

**A** Close-up plot of the representative extracted ion chromatogram (EIC) of the various peptide states; the blue trace corresponds to the Unmodified peptide; the orange trace corresponds to the Spliced peptide. **B** Close-up plot of the m/z values of the peptides detected by the mass spectrometer, with the shaded colored areas corresponding to where the expected m/z values for the peptide m/z traces should occur, with the observed spectrum plotted in black. The Unmodified, Spliced peptide(s) were found in the mass spectrum. **C** Bar plot showing the relative abundance of the various peptide states at charge z=1; the bars represent the average relative abundances normalized to total peak areas, with each individual experimental replicate value plotted as a black dot. **D** Mirrored MS$^2$ plot for the Unmodified peptide. In this mirror plot, the full fragmentation spectrum is plotted in black pointing upward, and the matched fragment b ions (blue) and y ions (red) are plotted pointing downward, mirrored against it, so that each matched fragment can be compared directly with the measured spectrum. **E** MS$^2$ spectrum of the fragments corresponding to the Unmodified peptide sequence, with matched b (blue, above) and y ions (red, below) plotted. **F** Mirrored MS$^2$ plot for the Spliced peptide. In this mirror plot, the full fragmentation spectrum is plotted in black pointing upward, and the matched fragment b ions (blue) and y ions (red) are plotted pointing downward, mirrored against it, so that each matched fragment can be compared directly with the measured spectrum. **G** MS$^2$ spectrum of the fragments corresponding to the Spliced peptide sequence, with matched b (blue, above) and y ions (red, below) plotted.

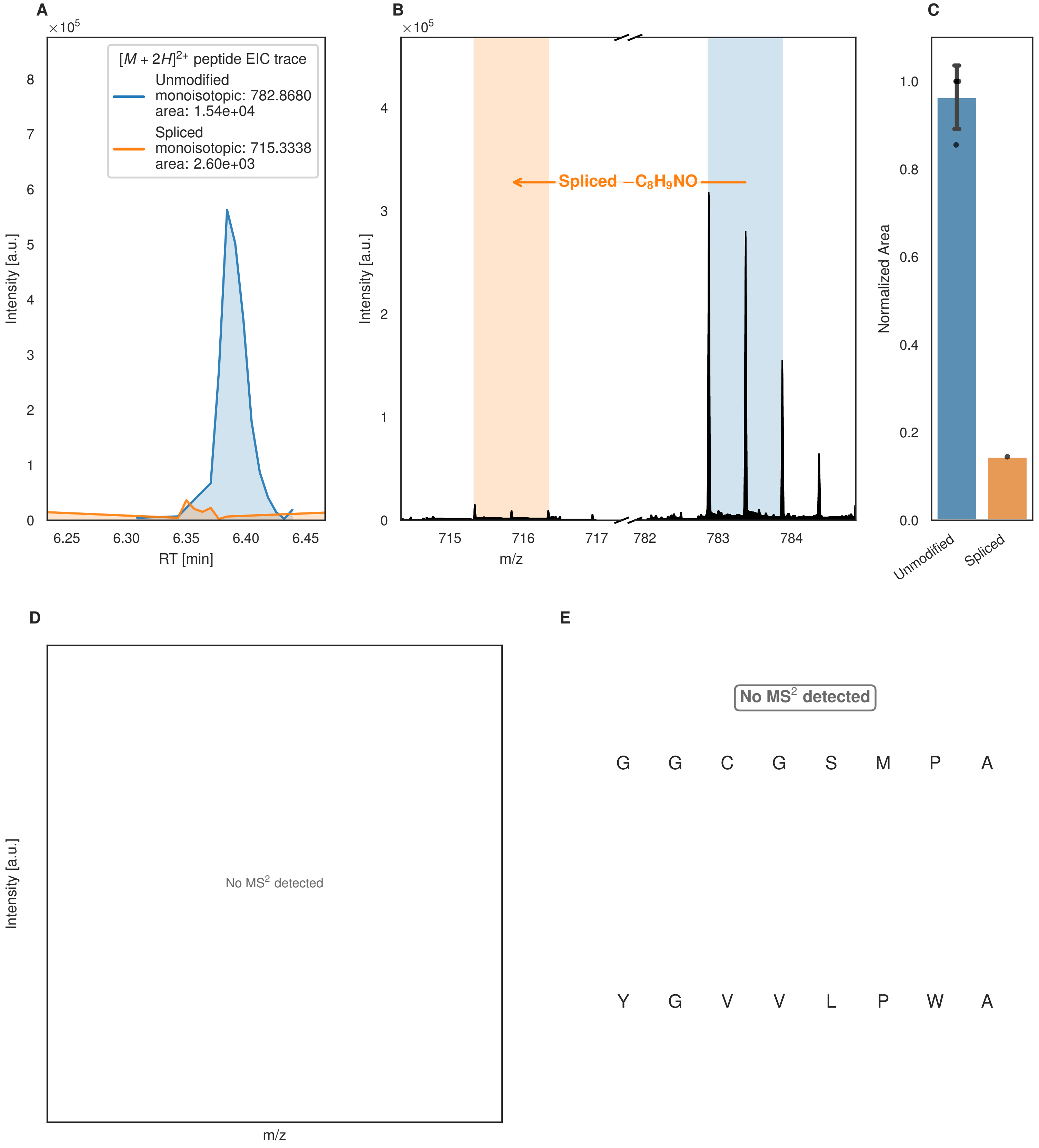
A
B
C
D
E
×10^5
[M + 2H]2+ peptide EIC trace
Unmodified
monoisotopic: 782.8680
area: 1.54e+04
Spliced
monoisotopic: 715.3338
area: 2.60e+03
Intensity [a.u.]
RT [min]
6.25
6.30
6.35
6.40
6.45
Spliced −C8H9NO
m/z
715
716
717
782
783
784
Normalized Area
Unmodified
Spliced
No MS2 detected
G G C G S M P A
Y G V V L P W A

Figure S177 (figure continued from the previous page).

F

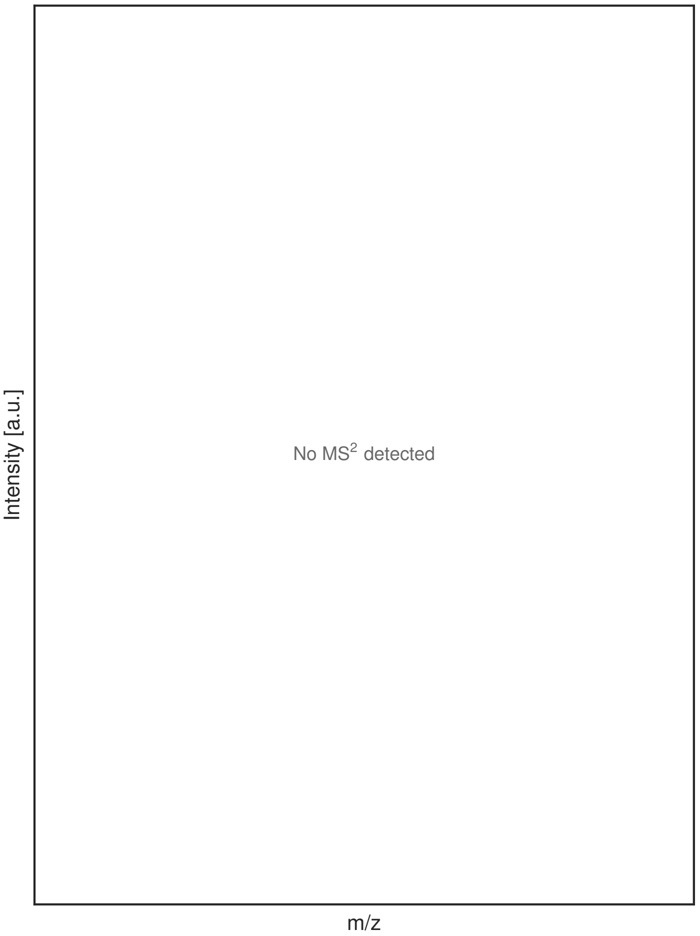


G

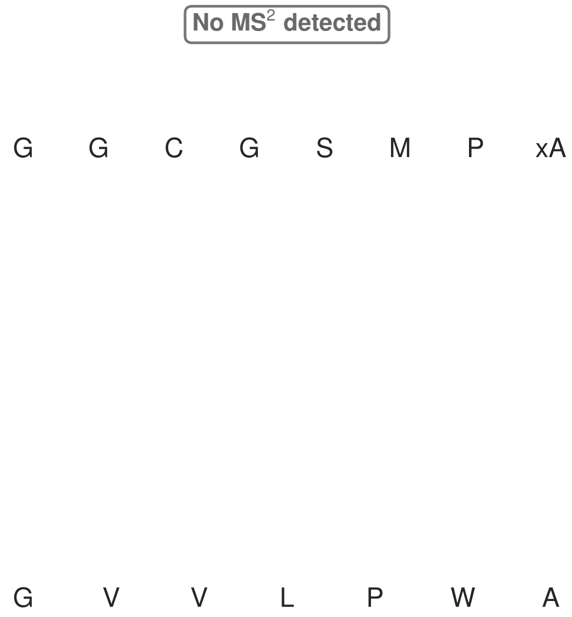


**Figure S177: Modification of the design_46 precursor peptide by PlpXY.** LC-MS analysis of IMAC-purified precursor peptides from *E. coli* BL21(DE3)Tuner containing splicease-encoding plasmid TJ-C-599 (PlpXY) and precursor peptide-encoding plasmid design_46 (RSTN-PlpA3-TEV-designed-46) after expression at 24 °C for 22 h in 1.5 mL cultures after digestion with TEV. Data from n=4 independent biological replicates. Spliced amino acids have an x as a prefix. Spliced refers to a single tyramine excision. A representative experimental replicate is shown in panels **A, B.**

**A** Close-up plot of the representative extracted ion chromatogram (EIC) of the various peptide states; the blue trace corresponds to the Unmodified peptide; the orange trace corresponds to the Spliced peptide. **B** Close-up plot of the m/z values of the peptides detected by the mass spectrometer, with the shaded colored areas corresponding to where the expected m/z values for the peptide m/z traces should occur, with the observed spectrum plotted in black. The Unmodified, Spliced peptide(s) were found in the mass spectrum. **C** Bar plot showing the relative abundance of the various peptide states at charge z=2; the bars represent the average relative abundances normalized to total peak areas, with each individual experimental replicate value plotted as a black dot. **D** Mirrored MS$^2$ plot for the Unmodified peptide. In this mirror plot, the full fragmentation spectrum is plotted in black pointing upward, and the matched fragment b ions (blue) and y ions (red) are plotted pointing downward, mirrored against it, so that each matched fragment can be compared directly with the measured spectrum. **E** MS$^2$ spectrum of the fragments corresponding to the Unmodified peptide sequence, with matched b (blue, above) and y ions (red, below) plotted. **F** Mirrored MS$^2$ plot for the Spliced peptide. In this mirror plot, the full fragmentation spectrum is plotted in black pointing upward, and the matched fragment b ions (blue) and y ions (red) are plotted pointing downward, mirrored against it, so that each matched fragment can be compared directly with the measured spectrum. **G** MS$^2$ spectrum of the fragments corresponding to the Spliced peptide sequence, with matched b (blue, above) and y ions (red, below) plotted.

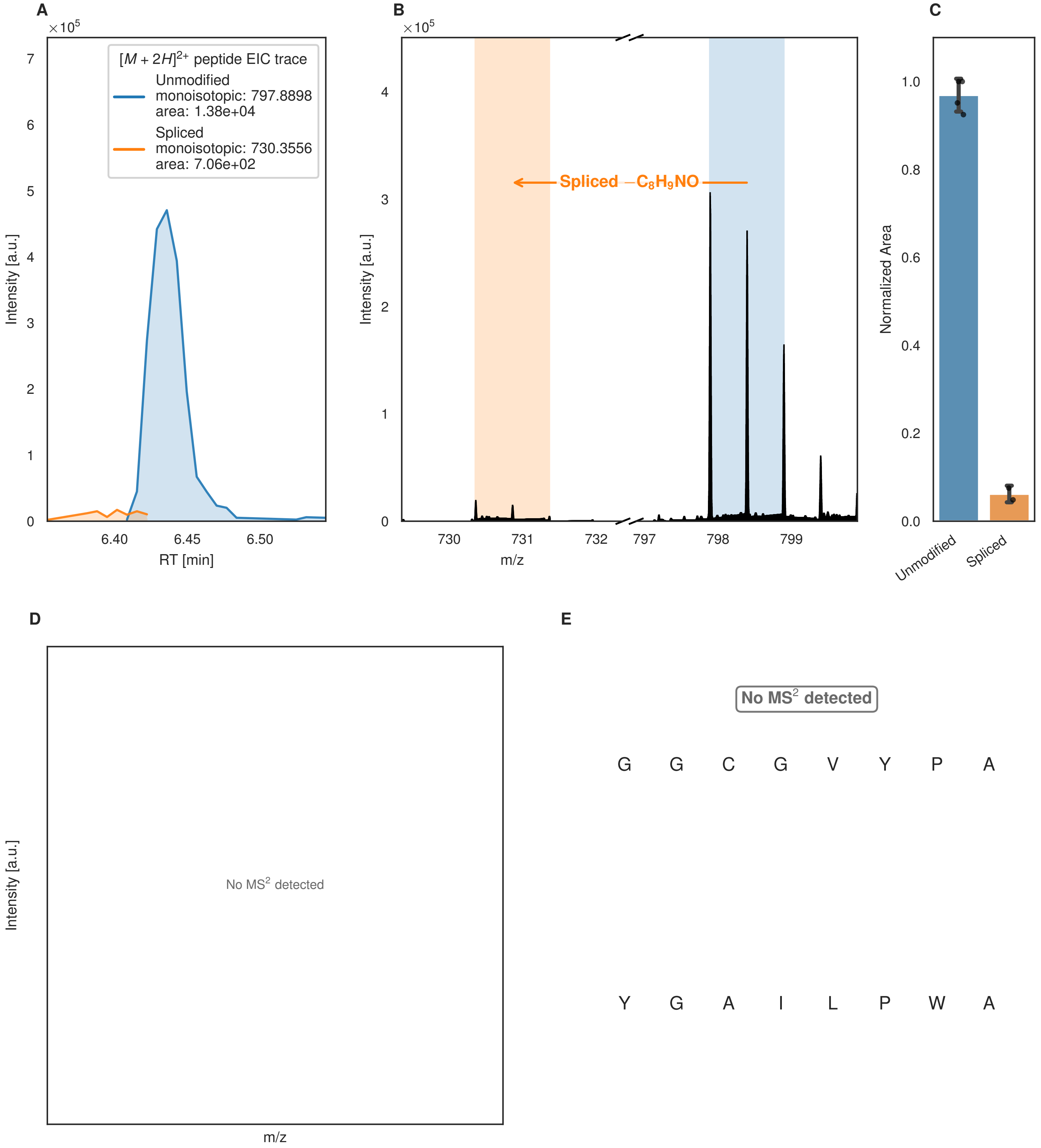
A
×10^5
[M + 2H]2+ peptide EIC trace
Unmodified
monoisotopic: 797.8898
area: 1.38e+04
Spliced
monoisotopic: 730.3556
area: 7.06e+02
Intensity [a.u.]
RT [min]
6.40
6.45
6.50
B
×10^5
Spliced −C8H9NO
Intensity [a.u.]
m/z
730
731
732
797
798
799
C
Normalized Area
Unmodified
Spliced
D
No MS2 detected
Intensity [a.u.]
m/z
E
No MS2 detected
G G C G V Y P A
Y G A I L P W A

F

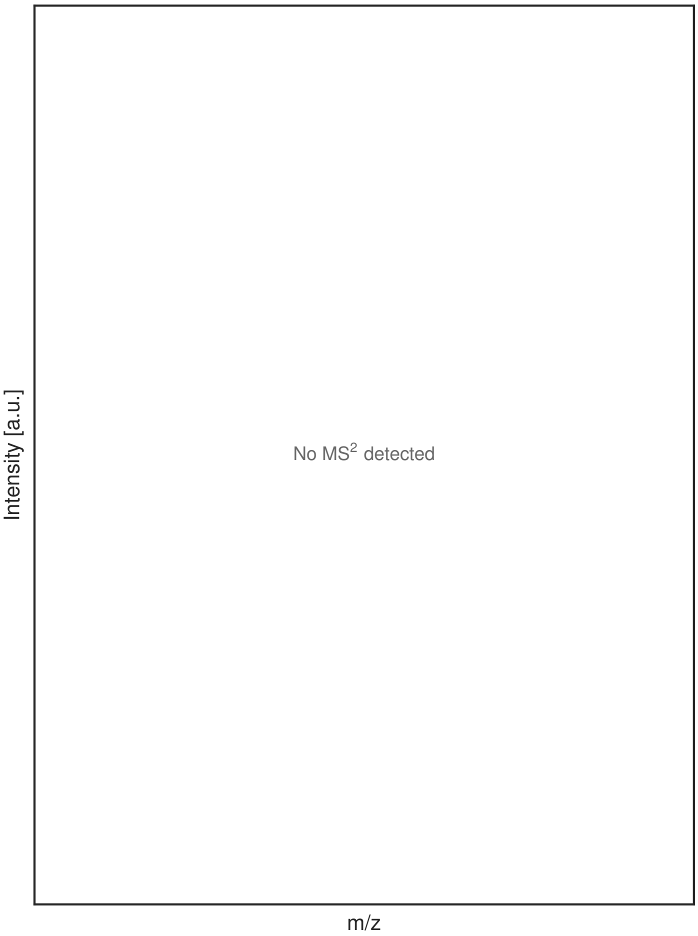


G

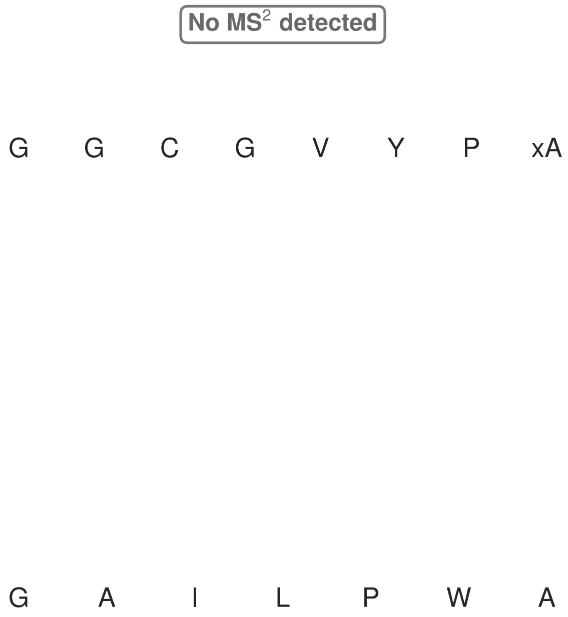


**Figure S178: Modification of the design_47 precursor peptide by PlpXY.** LC-MS analysis of IMAC-purified precursor peptides from *E. coli* BL21(DE3)Tuner containing splicease-encoding plasmid TJ-C-599 (PlpXY) and precursor peptide-encoding plasmid design_47 (RSTN-PlpA3-TEV-designed-47) after expression at 24 °C for 22 h in 1.5 mL cultures after digestion with TEV. Data from n=4 independent biological replicates. Spliced amino acids have an x as a prefix. Spliced refers to a single tyramine excision. A representative experimental replicate is shown in panels **A, B.**

**A** Close-up plot of the representative extracted ion chromatogram (EIC) of the various peptide states; the blue trace corresponds to the Unmodified peptide; the orange trace corresponds to the Spliced peptide. **B** Close-up plot of the m/z values of the peptides detected by the mass spectrometer, with the shaded colored areas corresponding to where the expected m/z values for the peptide m/z traces should occur, with the observed spectrum plotted in black. The Unmodified, Spliced peptide(s) were found in the mass spectrum. **C** Bar plot showing the relative abundance of the various peptide states at charge z=2; the bars represent the average relative abundances normalized to total peak areas, with each individual experimental replicate value plotted as a black dot. **D** Mirrored $MS^2$ plot for the Unmodified peptide. In this mirror plot, the full fragmentation spectrum is plotted in black pointing upward, and the matched fragment b ions (blue) and y ions (red) are plotted pointing downward, mirrored against it, so that each matched fragment can be compared directly with the measured spectrum. **E** $MS^2$ spectrum of the fragments corresponding to the Unmodified peptide sequence, with matched b (blue, above) and y ions (red, below) plotted. **F** Mirrored $MS^2$ plot for the Spliced peptide. In this mirror plot, the full fragmentation spectrum is plotted in black pointing upward, and the matched fragment b ions (blue) and y ions (red) are plotted pointing downward, mirrored against it, so that each matched fragment can be compared directly with the measured spectrum. **G** $MS^2$ spectrum of the fragments corresponding to the Spliced peptide sequence, with matched b (blue, above) and y ions (red, below) plotted.
