## Supplementary material for "Mapping and Probing the Substrate Scope of a Ketoamide-Forming Splicease by Computational Design": Main Supplementary Infromation File

* To whom correspondence should be addressed

### 1. Overview of Supplementary Materials

Two supplementary files accompany the manuscript in total (including this document):

- Main Supplementary Information File (this file)
- SI_report_plates.pdf

Figures found in this document.

| Figures | Construct group | Digestion | Notes |
|---|---|---|---|
| S1-7 | LC-MS traces of the wild-type *Hyella* precursors. | Trypsin | |
| S8 | Overview figure of the wild-type *Hyella* spliceotides and their modification. | Trypsin or GluC | |
| S9-10 | LC-MS traces of wild-type *Hyella* precursors. | GluC | |
| S11 | Overview figure of the *Hyella* spliceotides expressed as a fusion to the HypAc consensus leader sequence, and their modification. | Trypsin or GluC | |
| S12-16 | LC-MS traces of precursors expressed as fusions to the consensus leader sequence HypAc. | Trypsin or GluC | |
| S17 | In-depth overview of the HypA4 core and its modification states. | GluC | |
| S18 | In-depth overview of the HypA4 core and its modification states according to the $MS^2$ fragments. | GluC | |
| S19-26 | Artificial-1–4 peptides × HypXY. | TEV | Establishes Artificial-2 as the shared reference construct; underlies Fig. 2B. |
| S29-32 | Artificial-1–4 peptides × PlpX-PlpY | TEV | Establishes Artificial-2 as the shared reference construct; underlies Fig. 2B. |
| S31 | Overview figure of the effect of solubility tags and expression | TEV | |

| | | | |
|---|---|---|---|
| | temperature on the modification efficiency of HypX and PlpX. | | |
| S32–52 | PlpA3-boc-trunc, PlpA3-Trunc × Fh8 / Fh8-chaos / RSTN1 solubility tags × 24°C/30°C × HypX-HypY / PlpX-PlpY | TEV | Existing temperature and solubility-tag comparison data; corresponds to Figure S5 |

Figures found in the document SI_report_plates.pdf

| **Figures** | **Construct group** | **Digestion** | **Notes** |
|---|---|---|---|
| S53-133 | LC-MS traces of the point mutants of the artificial core. | TEV | Underlines Figure 3. |
| S134-S177 | LC-MS traces of the Rosetta-designed precursors | TEV | Underlines Figure 4C. |

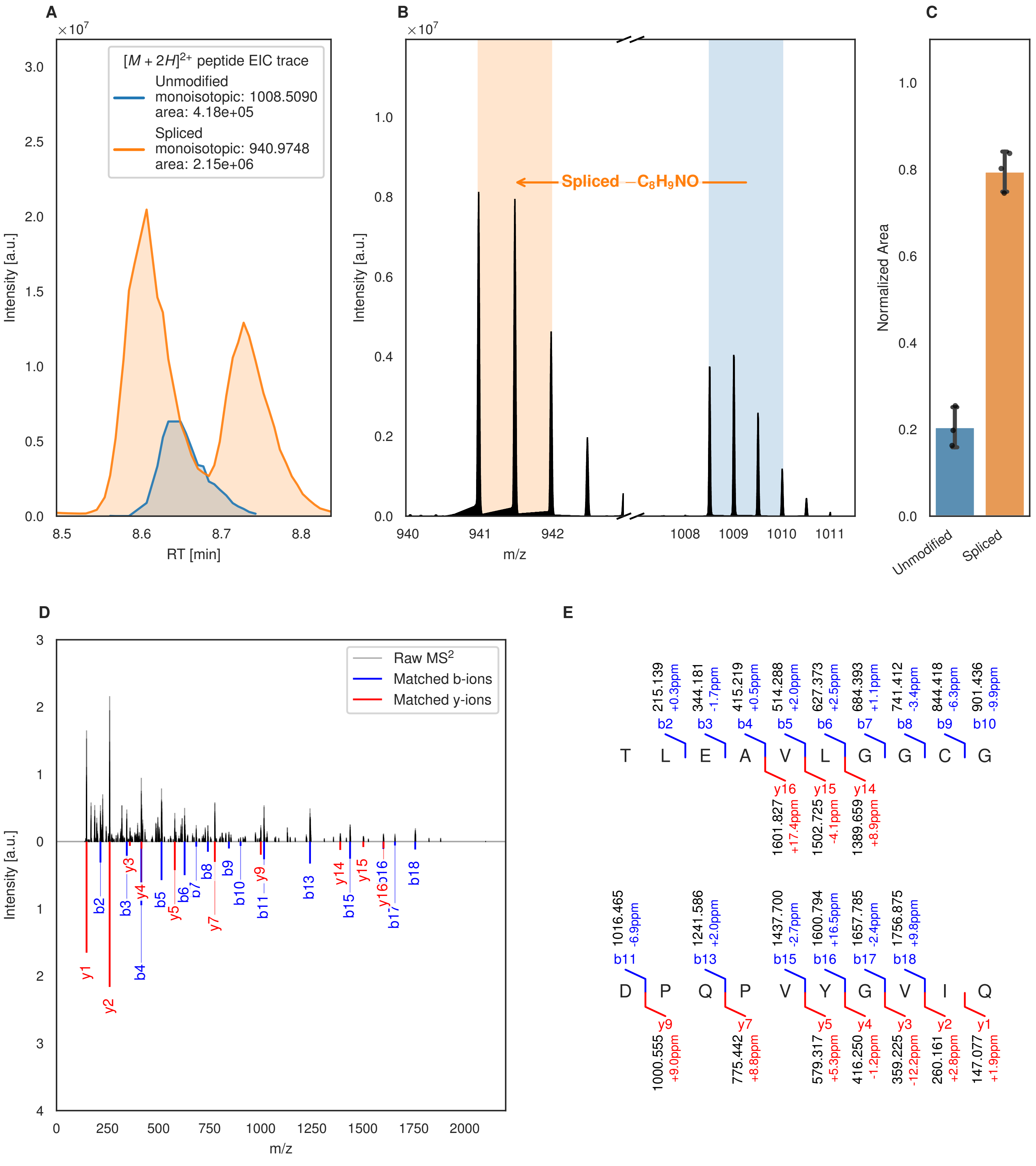
A
B
C
D
E
×10^7
[M + 2H]^2+ peptide EIC trace
Unmodified
monoisotopic: 1008.5090
area: 4.18e+05
Spliced
monoisotopic: 940.9748
area: 2.15e+06
Intensity [a.u.]
RT [min]
Spliced −C8H9NO
m/z
Normalized Area
Unmodified
Spliced
Raw MS2
Matched b-ions
Matched y-ions
T L E A V L G G C G
D P Q P V Y G V I Q
b2 215.139 +0.3ppm
b3 344.181 -1.7ppm
b4 415.219 +0.5ppm
b5 514.288 +2.0ppm
b6 627.373 +2.5ppm
b7 684.393 +1.1ppm
b8 741.412 -3.4ppm
b9 844.418 -6.3ppm
b10 901.436 -9.9ppm
y16 1601.827 +17.4ppm
y15 1502.725 -4.1ppm
y14 1389.659 +8.9ppm
b11 1016.465 -6.9ppm
b13 1241.586 +2.0ppm
b15 1437.700 -2.7ppm
b16 1600.794 +16.5ppm
b17 1657.785 -2.4ppm
b18 1756.875 +9.8ppm
y9 1000.555 +9.0ppm
y7 775.442 +8.8ppm
y5 579.317 +5.3ppm
y4 416.250 -1.2ppm
y3 359.225 -12.2ppm
y2 260.161 +2.8ppm
y1 147.077 +1.9ppm

Figure S1 (figure continued from the previous page).

F G

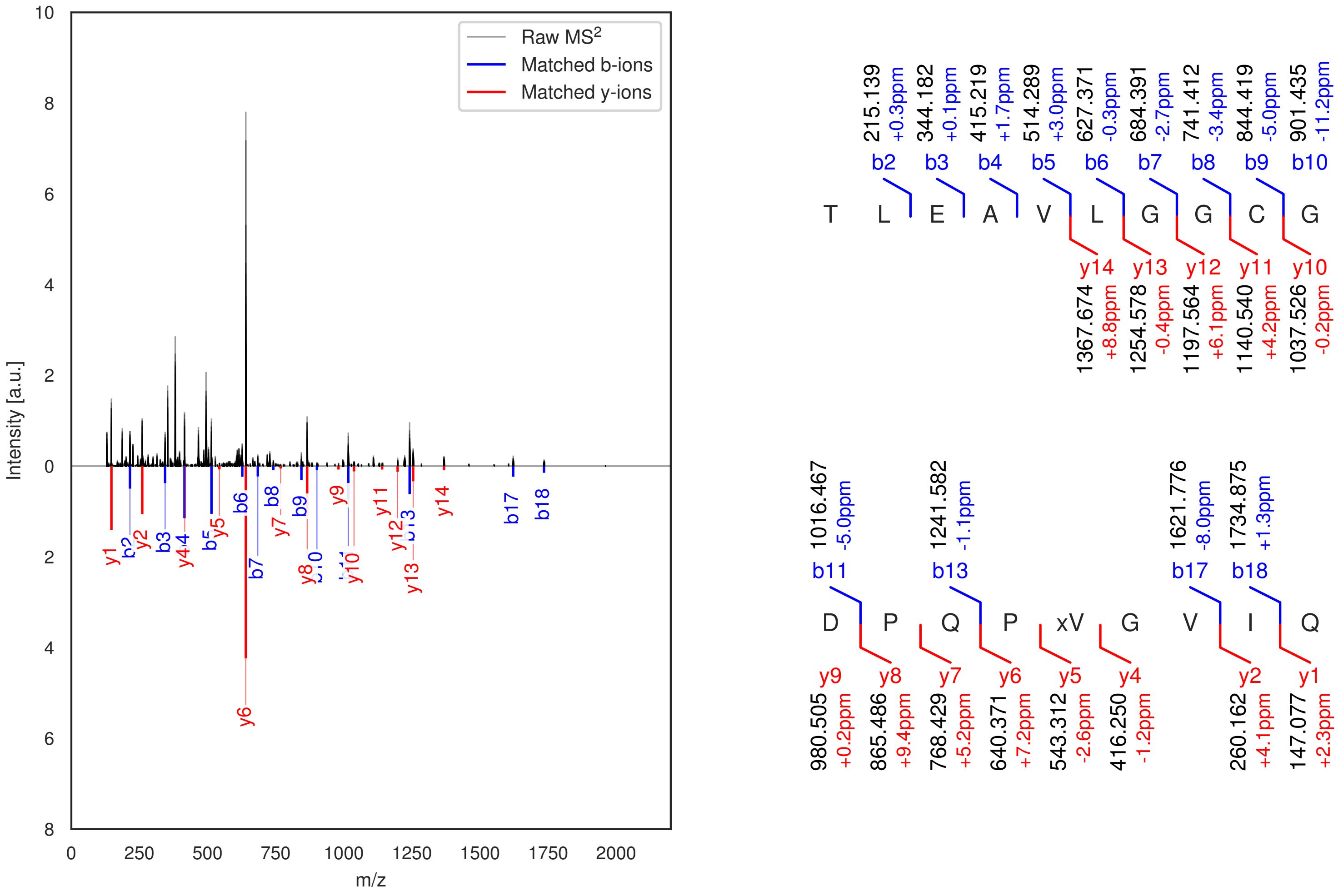

**Figure S1: Modification of the TJ-C-593 precursor peptide by HypXY.** LC-MS analysis of IMAC-purified precursor peptides from *E. coli* BL21(DE3)Tuner containing splicease-encoding plasmid TJ-C-560 (HypXY) and precursor peptide-encoding plasmid TJ-C-593 (Fh8-chaos-HypA1-HypA1) after expression at 24 °C for 22 h in 30 mL cultures after digestion with trypsin. Data from n=3 independent biological replicates. Spliced amino acids have an x as a prefix. Spliced refers to a single tyramine excision. A representative experimental replicate is shown in panels **A, B.**

**A** Close-up plot of the representative extracted ion chromatogram (EIC) of the various peptide states; the blue trace corresponds to the Unmodified peptide; the orange trace corresponds to the Spliced peptide. **B** Close-up plot of the m/z values of the peptides detected by the mass spectrometer, with the shaded colored areas corresponding to where the expected m/z values for the peptide m/z traces should occur, with the observed spectrum plotted in black. The Unmodified, Spliced peptide(s) were found in the mass spectrum. **C** Bar plot showing the relative abundance of the various peptide states at charge z=2; the bars represent the average relative abundances normalized to total peak areas, with each individual experimental replicate value plotted as a black dot. **D** Mirrored MS$^2$ plot for the Unmodified peptide. In this mirror plot, the full fragmentation spectrum is plotted in black pointing upward, and the matched fragment b ions (blue) and y ions (red) are plotted pointing downward, mirrored against it, so that each matched fragment can be compared directly with the measured spectrum. **E** MS$^2$ spectrum of the fragments corresponding to the Unmodified peptide sequence, with matched b (blue, above) and y ions (red, below) plotted. **F** Mirrored MS$^2$ plot for the Spliced peptide. In this mirror plot, the full fragmentation spectrum is plotted in black pointing upward, and the matched fragment b ions (blue) and y ions (red) are plotted pointing downward, mirrored against it, so that each matched fragment can be compared directly with the measured spectrum. **G** MS$^2$ spectrum of the fragments corresponding to the Spliced peptide sequence, with matched b (blue, above) and y ions (red, below) plotted.

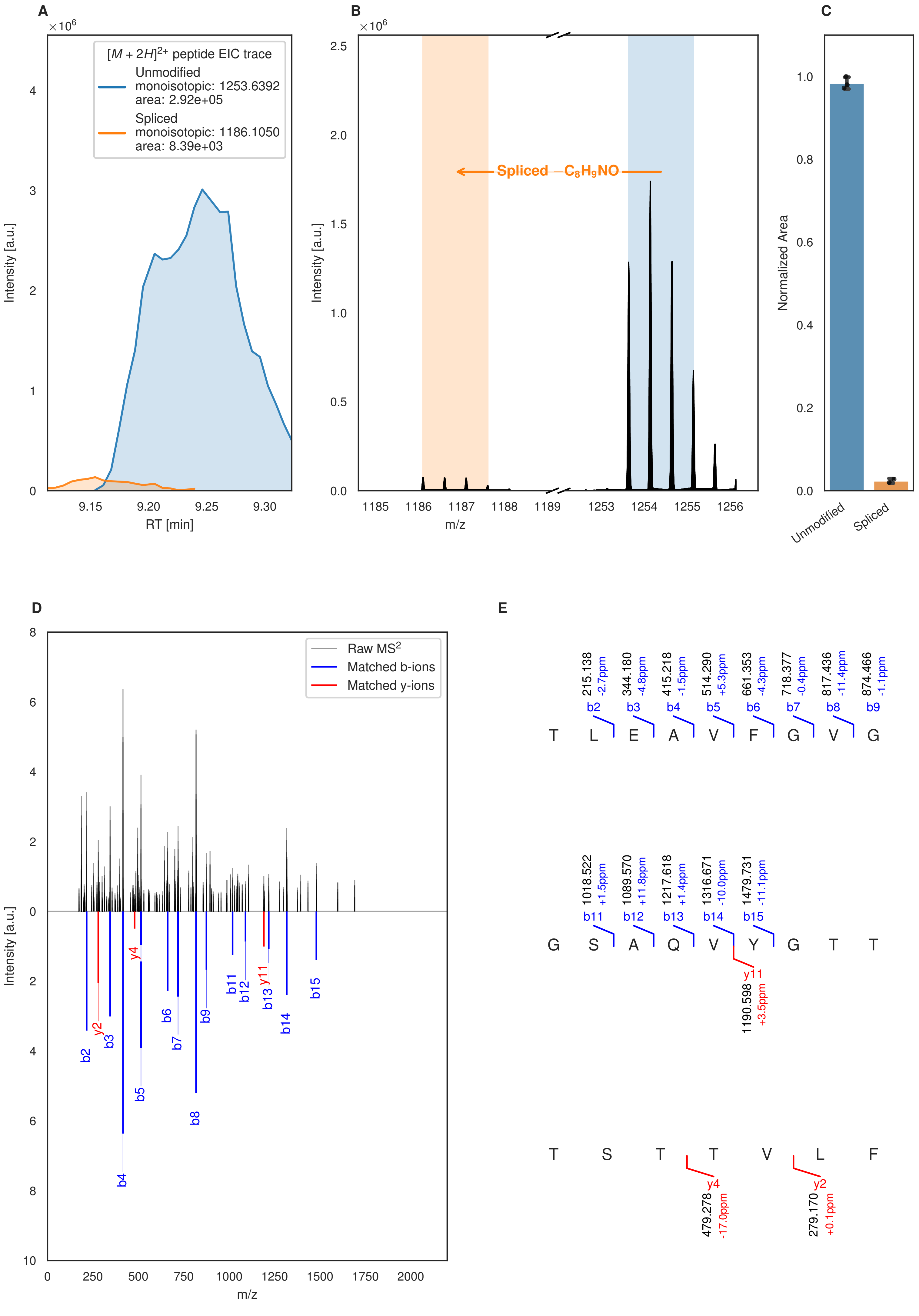
A
×10^6
[M + 2H]2+ peptide EIC trace
Unmodified
monoisotopic: 1253.6392
area: 2.92e+05
Spliced
monoisotopic: 1186.1050
area: 8.39e+03
Intensity [a.u.]
RT [min]
B
×10^6
Spliced −C8H9NO
Intensity [a.u.]
m/z
C
Normalized Area
Unmodified
Spliced
D
Raw MS2
Matched b-ions
Matched y-ions
Intensity [a.u.]
m/z
E
215.138 -2.7ppm b2
344.180 -4.8ppm b3
415.218 -1.5ppm b4
514.290 +5.3ppm b5
661.353 -4.3ppm b6
718.377 -0.4ppm b7
817.436 -11.4ppm b8
874.466 -1.1ppm b9
T L E A V F G V G
1018.522 +1.5ppm b11
1089.570 +11.8ppm b12
1217.618 +1.4ppm b13
1316.671 -10.0ppm b14
1479.731 -11.1ppm b15
G S A Q V Y G T T
y11 1190.598 +3.5ppm
T S T T V L F
y4 479.278 -17.0ppm
y2 279.170 +0.1ppm

Figure S2 (figure continued from the previous page).

F G

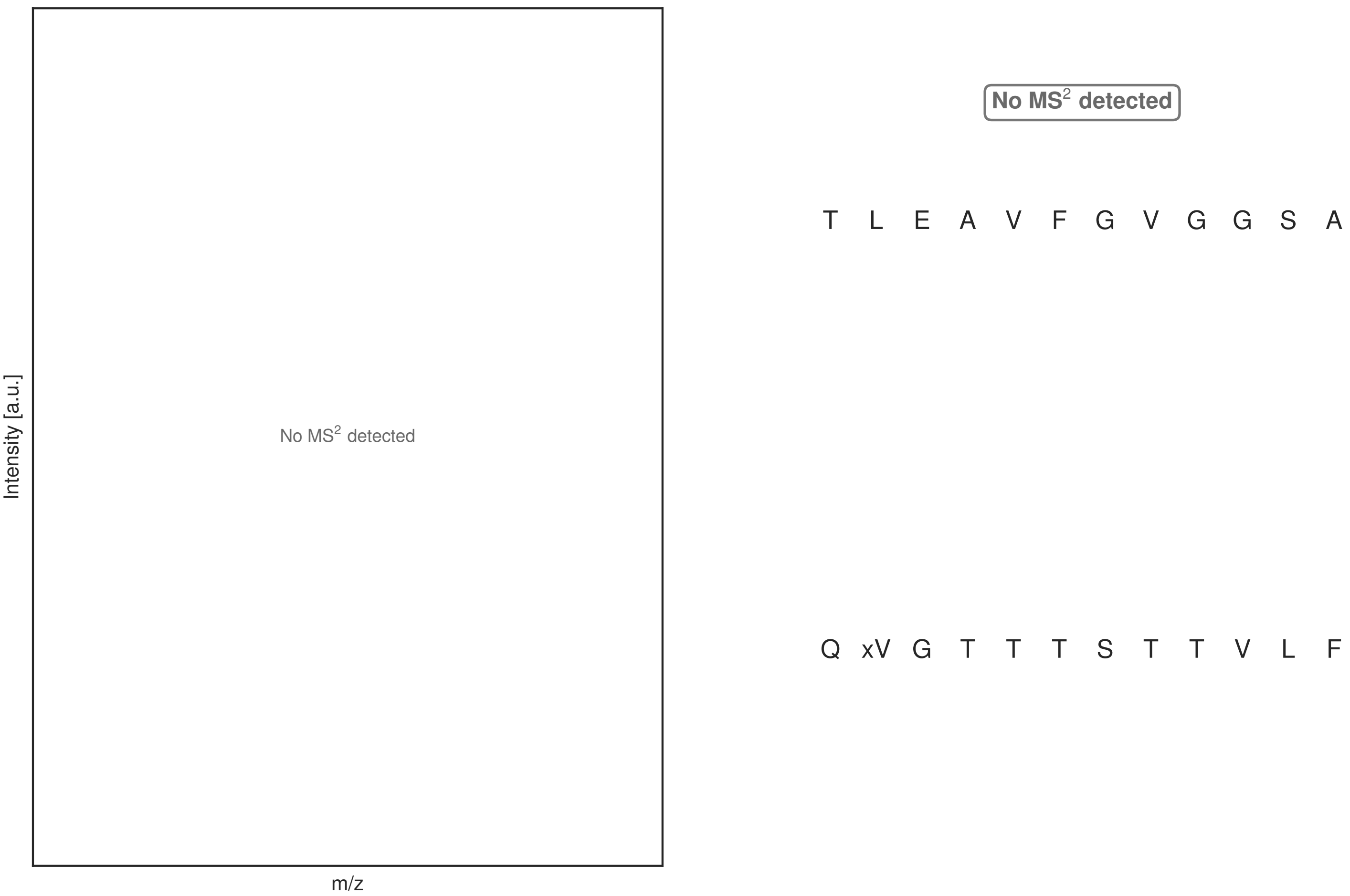

**Figure S2: Modification of the TJ-C-594 precursor peptide by HypXY.** LC-MS analysis of IMAC-purified precursor peptides from *E. coli* BL21(DE3)Tuner containing splicease-encoding plasmid TJ-C-560 (HypXY) and precursor peptide-encoding plasmid TJ-C-594 (Fh8-chaos-HypA2-HypA2) after expression at 24 °C for 22 h in 30 mL cultures after digestion with trypsin. Data from n=3 independent biological replicates. Spliced amino acids have an x as a prefix. Spliced refers to a single tyramine excision. A representative experimental replicate is shown in panels **A, B.**

**A** Close-up plot of the representative extracted ion chromatogram (EIC) of the various peptide states; the blue trace corresponds to the Unmodified peptide; the orange trace corresponds to the Spliced peptide. **B** Close-up plot of the m/z values of the peptides detected by the mass spectrometer, with the shaded colored areas corresponding to where the expected m/z values for the peptide m/z traces should occur, with the observed spectrum plotted in black. The Unmodified, Spliced peptide(s) were found in the mass spectrum. **C** Bar plot showing the relative abundance of the various peptide states at charge z=2; the bars represent the average relative abundances normalized to total peak areas, with each individual experimental replicate value plotted as a black dot. **D** Mirrored MS$^2$ plot for the Unmodified peptide. In this mirror plot, the full fragmentation spectrum is plotted in black pointing upward, and the matched fragment b ions (blue) and y ions (red) are plotted pointing downward, mirrored against it, so that each matched fragment can be compared directly with the measured spectrum. **E** MS$^2$ spectrum of the fragments corresponding to the Unmodified peptide sequence, with matched b (blue, above) and y ions (red, below) plotted. **F** Mirrored MS$^2$ plot for the Spliced peptide. In this mirror plot, the full fragmentation spectrum is plotted in black pointing upward, and the matched fragment b ions (blue) and y ions (red) are plotted pointing downward, mirrored against it, so that each matched fragment can be compared directly with the measured spectrum. **G** MS$^2$ spectrum of the fragments corresponding to the Spliced peptide sequence, with matched b (blue, above) and y ions (red, below) plotted.

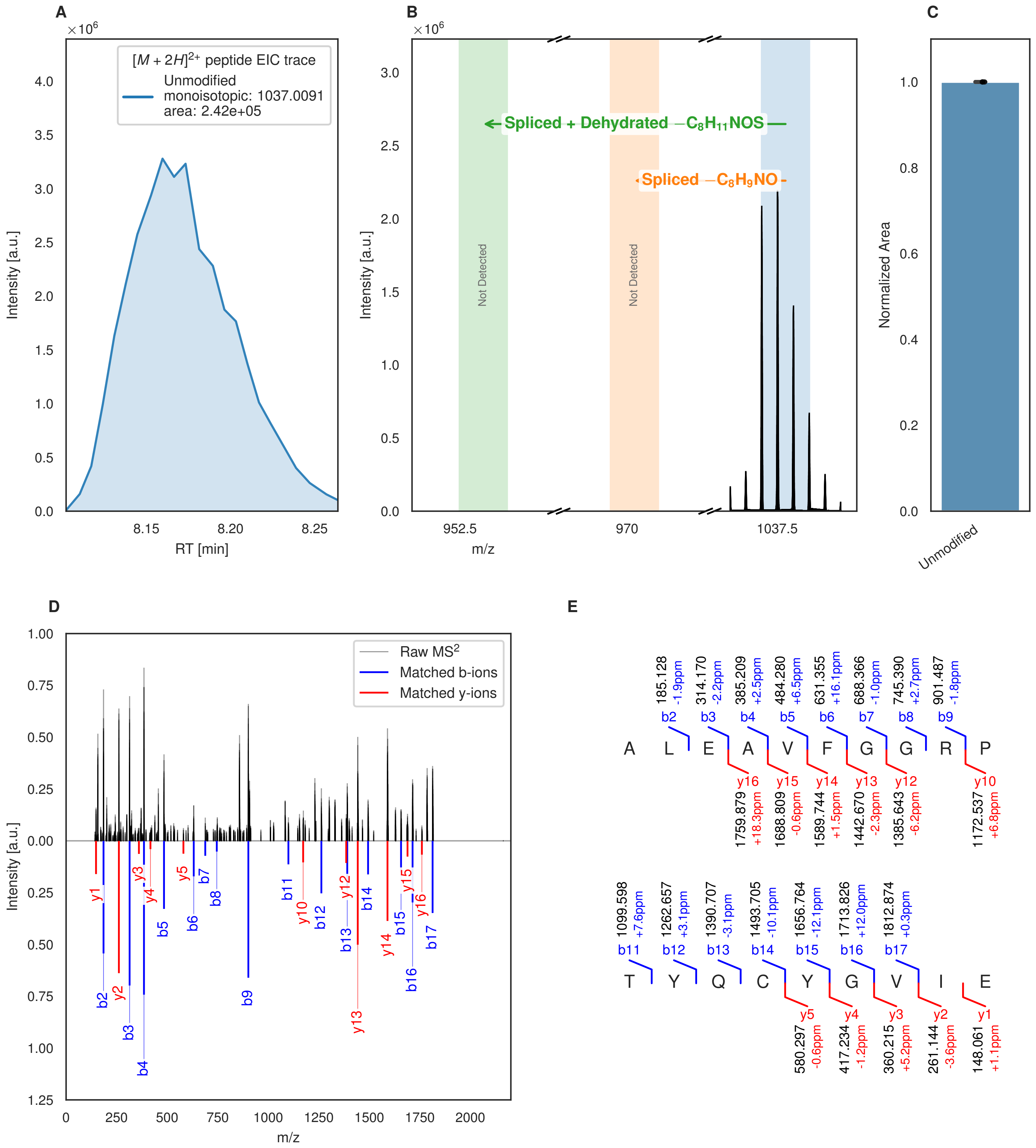

**Figure S3: Modification of the TJ-C-595 precursor peptide by HypXY.** LC-MS analysis of IMAC-purified precursor peptides from *E. coli* BL21(DE3)Tuner containing splicease-encoding plasmid TJ-C-560 (HypXY) and precursor peptide-encoding plasmid TJ-C-595 (Fh8-chaos-HypA3-HypA3) after expression at 24 °C for 22 h in 30 mL cultures after digestion with trypsin. Data from n=3 independent biological replicates. Spliced amino acids have an x as a prefix. Spliced refers to a single tyramine excision; Spliced + Dehydrated refers to a dehydroalanine-derived ketoamide residue (xdhA). A representative experimental replicate is shown in panels **A, B.** **A** Close-up plot of the representative extracted ion chromatogram (EIC) of the various peptide states; the blue trace corresponds to the Unmodified peptide. **B** Close-up plot of the m/z values of the peptides detected by the mass spectrometer, with the shaded colored areas corresponding to where the expected m/z values for the peptide m/z

(Caption continued on the next page.)

traces should occur, with the observed spectrum plotted in black. The Unmodified peptide(s) were found in the mass spectrum and the Spliced, Spliced + Dehydrated peptide(s) were not detected. **C** Bar plot showing the relative abundance of the various peptide states at charge z=2; the bars represent the average relative abundances normalized to total peak areas, with each individual experimental replicate value plotted as a black dot. **D** Mirrored $MS^2$ plot for the Unmodified peptide. In this mirror plot, the full fragmentation spectrum is plotted in black pointing upward, and the matched fragment b ions (blue) and y ions (red) are plotted pointing downward, mirrored against it, so that each matched fragment can be compared directly with the measured spectrum. **E** $MS^2$ spectrum of the fragments corresponding to the Unmodified peptide sequence, with matched b (blue, above) and y ions (red, below) plotted.

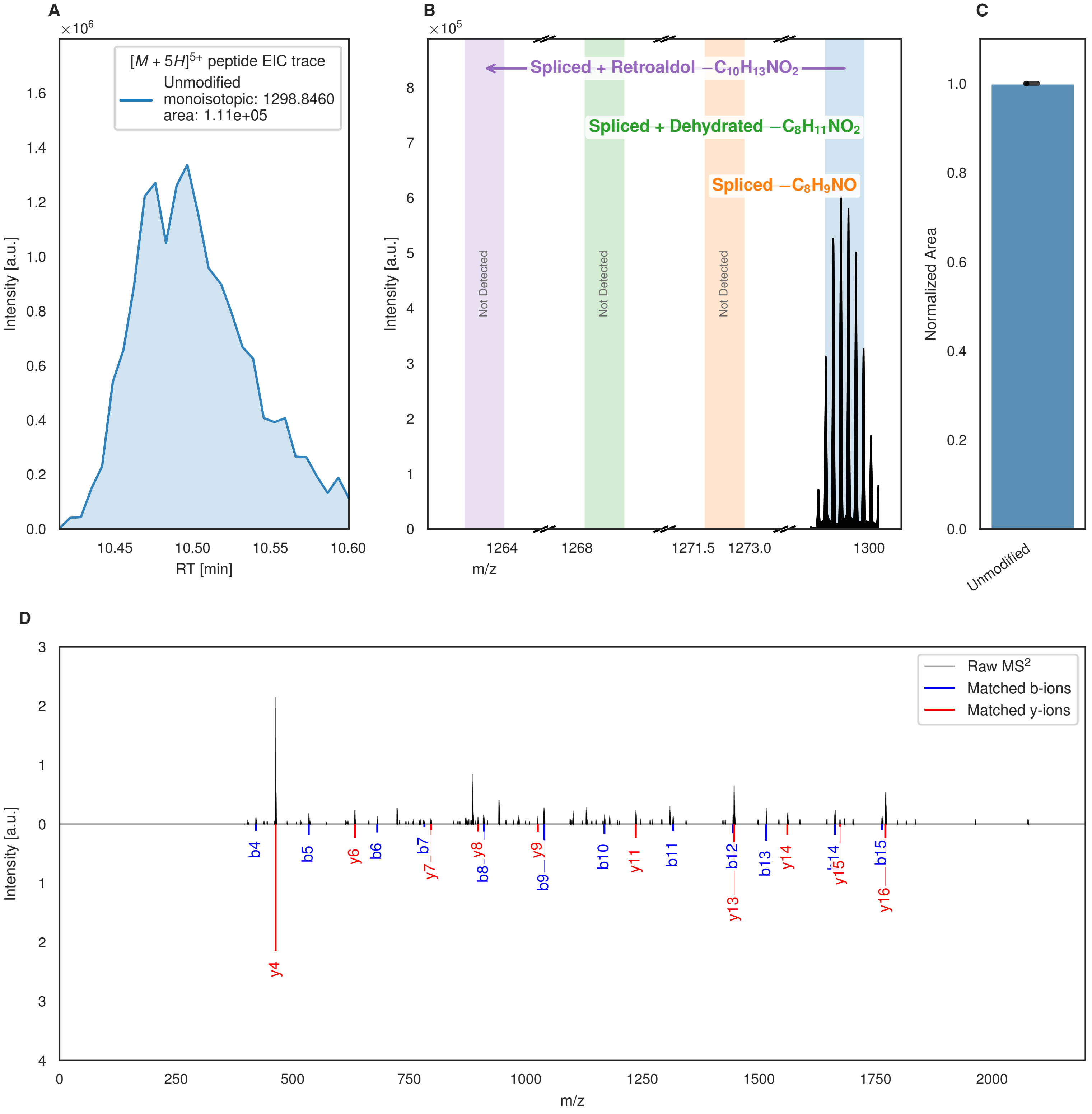

A
$\times10^6$
$[M+5H]^{5+}$ peptide EIC trace
Unmodified
monoisotopic: 1298.8460
area: 1.11e+05
Intensity [a.u.]
RT [min]
B
$\times10^5$
Spliced + Retroaldol $-C_{10}H_{13}NO_2$
Spliced + Dehydrated $-C_8H_{11}NO_2$
Spliced $-C_8H_9NO$
Not Detected
Not Detected
Not Detected
Intensity [a.u.]
m/z
C
Normalized Area
Unmodified
D
Raw $MS^2$
Matched b-ions
Matched y-ions
Intensity [a.u.]
m/z
b4 b5 y6 b6 b7 y7 y8 b8 y9 b9 b10 y11 b11 b12 y13 b13 y14 b14 y15 b15 y16 y4

Figure S4 (figure continued from the previous page).

**E**

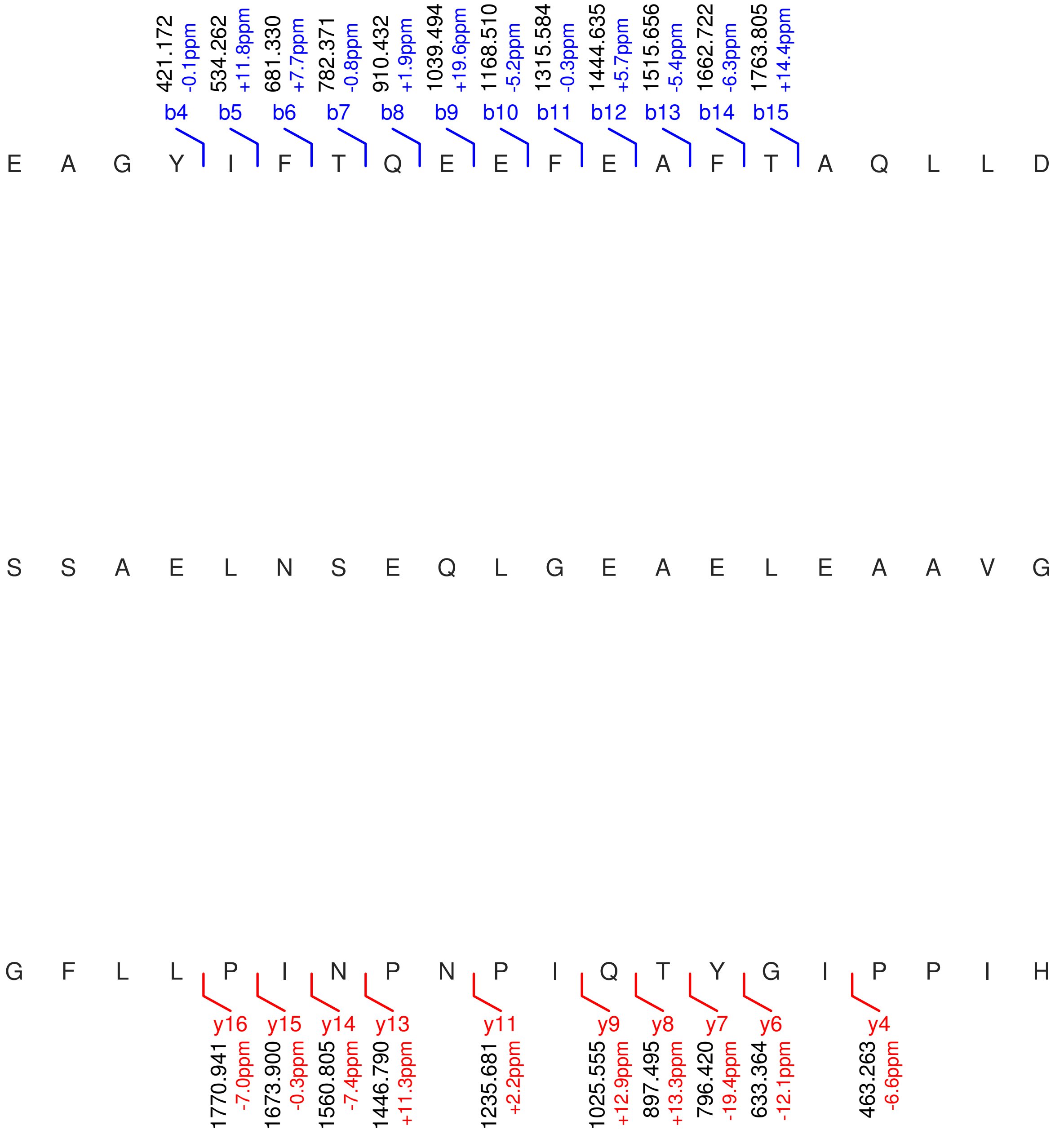

**Figure S4: Modification of the TJ-C-596 precursor peptide by HypXY.** LC-MS analysis of IMAC-purified precursor peptides from *E. coli* BL21(DE3)Tuner containing splicease-encoding plasmid TJ-C-560 (HypXY) and precursor peptide-encoding plasmid TJ-C-596 (Fh8-chaos-HypA4-HypA4) after expression at 24 °C for 22 h in 30 mL cultures after digestion with trypsin. Data from n=2 independent biological replicates. Spliced amino acids have an x as a prefix. Spliced refers to a single tyramine excision; Spliced + Dehydrated refers to a dehydrobutyrine-derived ketoamide residue (xdhB); Spliced + Retroaldol refers to a glycine-derived ketoamide (xG). A representative experimental replicate is shown in panels **A, B.**

**A** Close-up plot of the representative extracted ion chromatogram (EIC) of the various peptide states; the blue trace corresponds to the Unmodified peptide. **B** Close-up plot of the m/z values of the peptides detected by the mass spectrometer, with the shaded colored areas corresponding to where the expected m/z values for the peptide m/z traces should occur, with the observed spectrum plotted in black. The Unmodified peptide(s) were found in the mass spectrum and the Spliced, Spliced + Dehydrated, Spliced + Retroaldol peptide(s) were not detected. **C** Bar

(Caption continued on the next page.)

plot showing the relative abundance of the various peptide states at charge z=5; the bars represent the average relative abundances normalized to total peak areas, with each individual experimental replicate value plotted as a black dot. **D** Mirrored $MS^2$ plot for the Unmodified peptide. In this mirror plot, the full fragmentation spectrum is plotted in black pointing upward, and the matched fragment b ions (blue) and y ions (red) are plotted pointing downward, mirrored against it, so that each matched fragment can be compared directly with the measured spectrum. **E** $MS^2$ spectrum of the fragments corresponding to the Unmodified peptide sequence, with matched b (blue, above) and y ions (red, below) plotted.

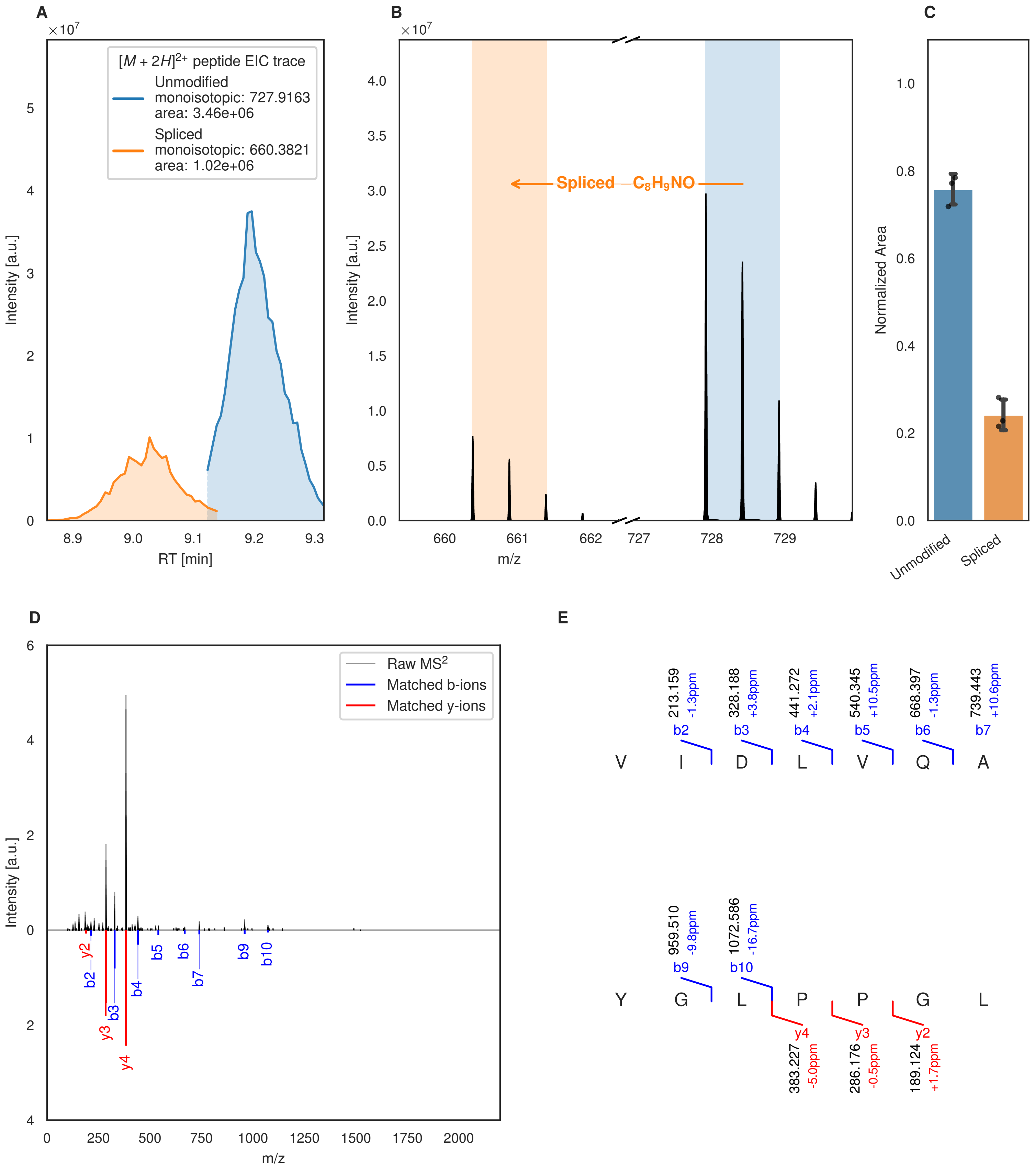
A
B
C
D
E
$[M+2H]^{2+}$ peptide EIC trace
Unmodified
monoisotopic: 727.9163
area: 3.46e+06
Spliced
monoisotopic: 660.3821
area: 1.02e+06
Intensity [a.u.]
RT [min]
Spliced $-C_8H_9NO$
m/z
Normalized Area
Unmodified
Spliced
Raw $MS^2$
Matched b-ions
Matched y-ions
V I D L V Q A
213.159 -1.3ppm b2
328.188 +3.8ppm b3
441.272 +2.1ppm b4
540.345 +10.5ppm b5
668.397 -1.3ppm b6
739.443 +10.6ppm b7
Y G L P P G L
959.510 -9.8ppm b9
1072.586 -16.7ppm b10
y4 383.227 -5.0ppm
y3 286.176 -0.5ppm
y2 189.124 +1.7ppm

Figure S5 (figure continued from the previous page).

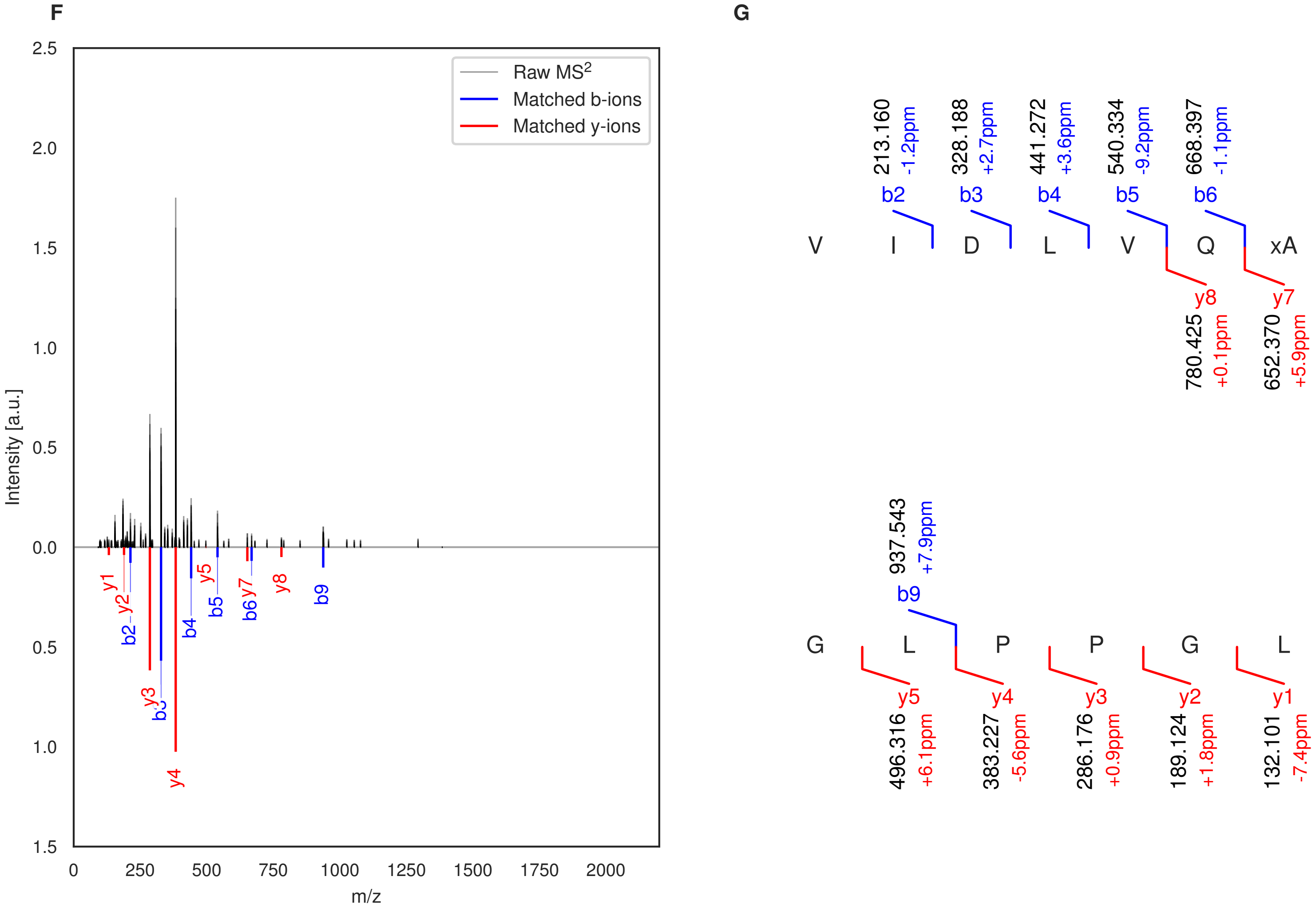

**Figure S5: Modification of the TJ-C-597 precursor peptide by HypXY.** LC-MS analysis of IMAC-purified precursor peptides from *E. coli* BL21(DE3)Tuner containing splicease-encoding plasmid TJ-C-560 (HypXY) and precursor peptide-encoding plasmid TJ-C-597 (Fh8-chaos-HypA5-HypA5) after expression at 24 °C for 22 h in 30 mL cultures after digestion with trypsin. Data from n=3 independent biological replicates. Spliced amino acids have an x as a prefix. Spliced refers to a single tyramine excision. A representative experimental replicate is shown in panels **A, B.**

**A** Close-up plot of the representative extracted ion chromatogram (EIC) of the various peptide states; the blue trace corresponds to the Unmodified peptide; the orange trace corresponds to the Spliced peptide. **B** Close-up plot of the m/z values of the peptides detected by the mass spectrometer, with the shaded colored areas corresponding to where the expected m/z values for the peptide m/z traces should occur, with the observed spectrum plotted in black. The Unmodified, Spliced peptide(s) were found in the mass spectrum. **C** Bar plot showing the relative abundance of the various peptide states at charge z=2; the bars represent the average relative abundances normalized to total peak areas, with each individual experimental replicate value plotted as a black dot. **D** Mirrored MS$^2$ plot for the Unmodified peptide. In this mirror plot, the full fragmentation spectrum is plotted in black pointing upward, and the matched fragment b ions (blue) and y ions (red) are plotted pointing downward, mirrored against it, so that each matched fragment can be compared directly with the measured spectrum. **E** MS$^2$ spectrum of the fragments corresponding to the Unmodified peptide sequence, with matched b (blue, above) and y ions (red, below) plotted. **F** Mirrored MS$^2$ plot for the Spliced peptide. In this mirror plot, the full fragmentation spectrum is plotted in black pointing upward, and the matched fragment b ions (blue) and y ions (red) are plotted pointing downward, mirrored against it, so that each matched fragment can be compared directly with the measured spectrum. **G** MS$^2$ spectrum of the fragments corresponding to the Spliced peptide sequence, with matched b (blue, above) and y ions (red, below) plotted.

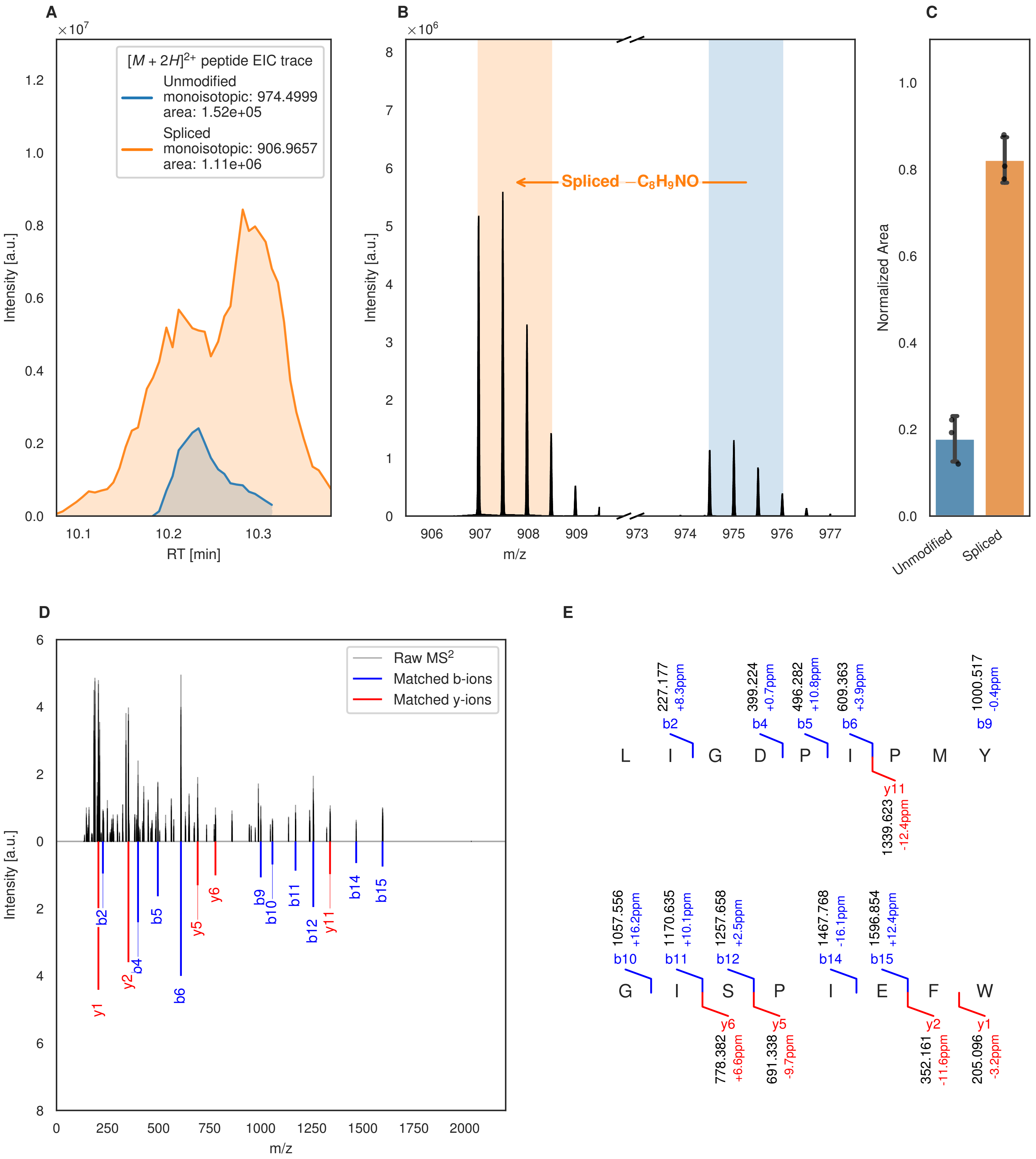
A
×10^7
[M + 2H]^2+ peptide EIC trace
Unmodified
monoisotopic: 974.4999
area: 1.52e+05
Spliced
monoisotopic: 906.9657
area: 1.11e+06
Intensity [a.u.]
RT [min]
10.1
10.2
10.3
B
×10^6
Spliced −C8H9NO
Intensity [a.u.]
m/z
906
907
908
909
973
974
975
976
977
C
Normalized Area
Unmodified
Spliced
D
Raw MS2
Matched b-ions
Matched y-ions
Intensity [a.u.]
m/z
b2
b4
b5
b6
b9
b10
b11
b12
b14
b15
y1
y2
y5
y6
y11
E
L I G D P I P M Y
227.177 +8.3ppm b2
399.224 +0.7ppm b4
496.282 +10.8ppm b5
609.363 +3.9ppm b6
1000.517 -0.4ppm b9
y11 1339.623 -12.4ppm
G I S P I E F W
1057.556 +16.2ppm b10
1170.635 +10.1ppm b11
1257.658 +2.5ppm b12
1467.768 -16.1ppm b14
1596.854 +12.4ppm b15
y6 778.382 +6.6ppm
y5 691.338 -9.7ppm
y2 352.161 -11.6ppm
y1 205.096 -3.2ppm

Figure S6 (figure continued from the previous page).

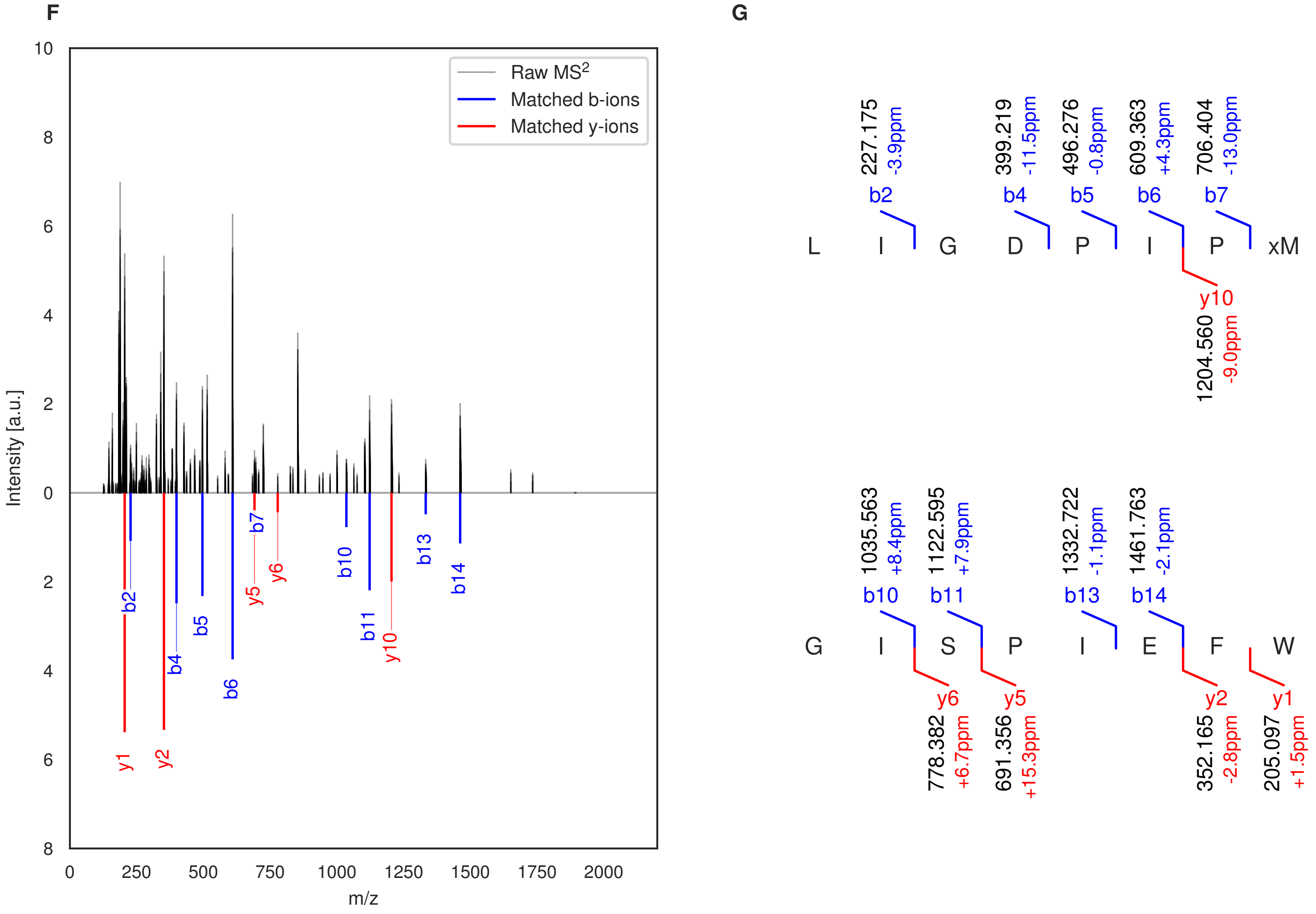

**Figure S6: Modification of the TJ-C-598 precursor peptide by HypXY.** LC-MS analysis of IMAC-purified precursor peptides from *E. coli* BL21(DE3)Tuner containing splicease-encoding plasmid TJ-C-560 (HypXY) and precursor peptide-encoding plasmid TJ-C-598 (Fh8-chaos-HypA6-HypA6) after expression at 24 °C for 22 h in 30 mL cultures after digestion with trypsin. Data from n=3 independent biological replicates. Spliced amino acids have an x as a prefix. Spliced refers to a single tyramine excision. A representative experimental replicate is shown in panels **A, B.**

**A** Close-up plot of the representative extracted ion chromatogram (EIC) of the various peptide states; the blue trace corresponds to the Unmodified peptide; the orange trace corresponds to the Spliced peptide. **B** Close-up plot of the m/z values of the peptides detected by the mass spectrometer, with the shaded colored areas corresponding to where the expected m/z values for the peptide m/z traces should occur, with the observed spectrum plotted in black. The Unmodified, Spliced peptide(s) were found in the mass spectrum. **C** Bar plot showing the relative abundance of the various peptide states at charge z=2; the bars represent the average relative abundances normalized to total peak areas, with each individual experimental replicate value plotted as a black dot. **D** Mirrored MS$^2$ plot for the Unmodified peptide. In this mirror plot, the full fragmentation spectrum is plotted in black pointing upward, and the matched fragment b ions (blue) and y ions (red) are plotted pointing downward, mirrored against it, so that each matched fragment can be compared directly with the measured spectrum. **E** MS$^2$ spectrum of the fragments corresponding to the Unmodified peptide sequence, with matched b (blue, above) and y ions (red, below) plotted. **F** Mirrored MS$^2$ plot for the Spliced peptide. In this mirror plot, the full fragmentation spectrum is plotted in black pointing upward, and the matched fragment b ions (blue) and y ions (red) are plotted pointing downward, mirrored against it, so that each matched fragment can be compared directly with the measured spectrum. **G** MS$^2$ spectrum of the fragments corresponding to the Spliced peptide sequence, with matched b (blue, above) and y ions (red, below) plotted.

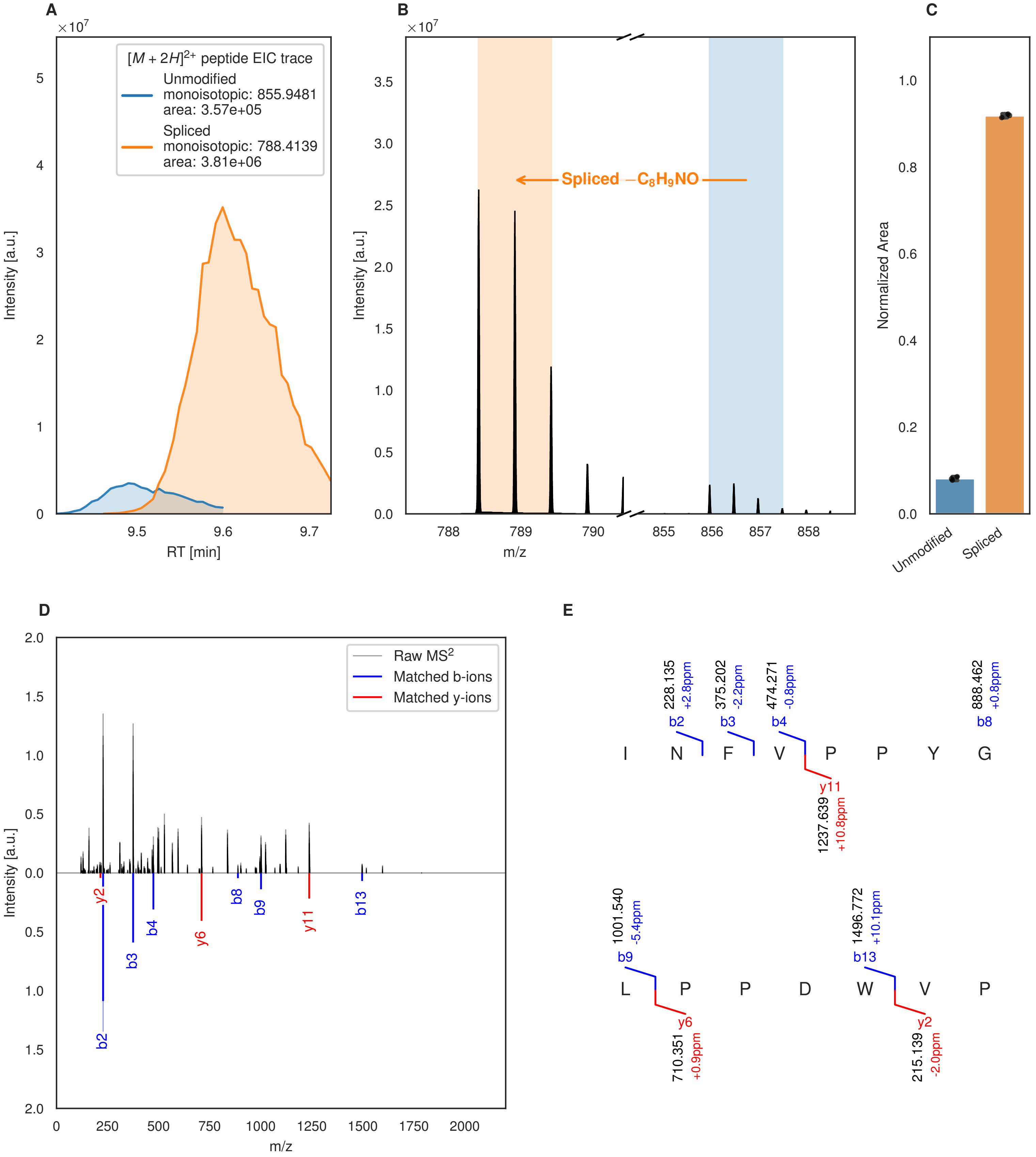
A
B
C
D
E
[M + 2H]2+ peptide EIC trace
Unmodified
monoisotopic: 855.9481
area: 3.57e+05
Spliced
monoisotopic: 788.4139
area: 3.81e+06
Intensity [a.u.]
RT [min]
Spliced −C8H9NO
m/z
Normalized Area
Unmodified
Spliced
Raw MS2
Matched b-ions
Matched y-ions
I N F V P P Y G
L P P D W V P
b2 228.135 +2.8ppm
b3 375.202 -2.2ppm
b4 474.271 -0.8ppm
b8 888.462 +0.8ppm
y11 1237.639 +10.8ppm
b9 1001.540 -5.4ppm
b13 1496.772 +10.1ppm
y6 710.351 +0.9ppm
y2 215.139 -2.0ppm

Figure S7 (figure continued from the previous page).

**F**

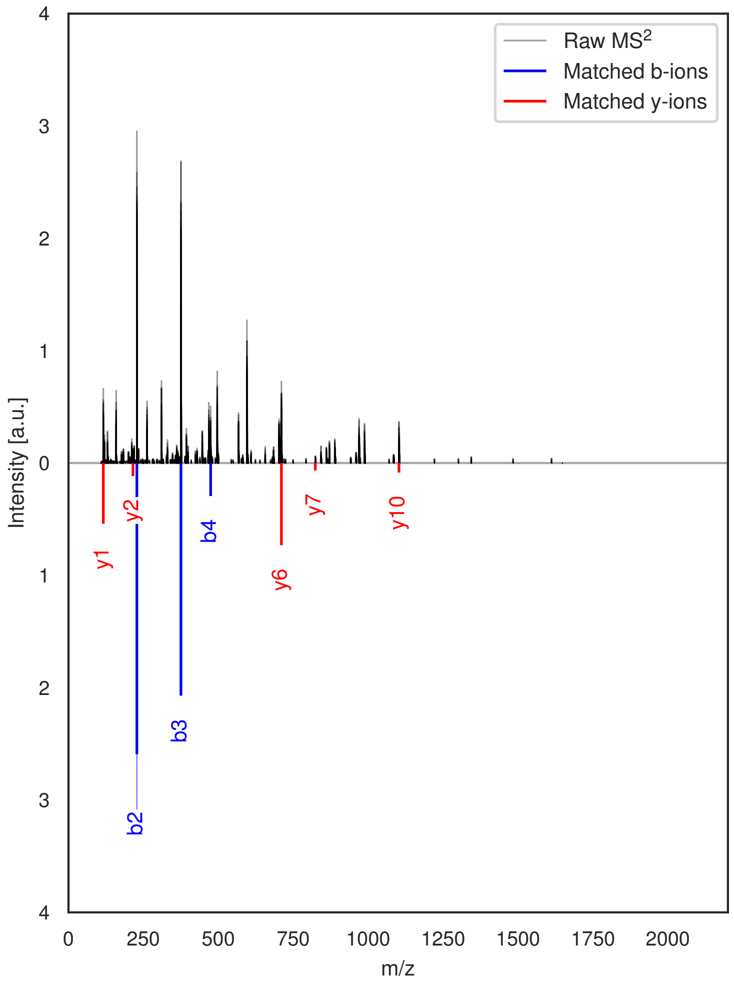

**G**

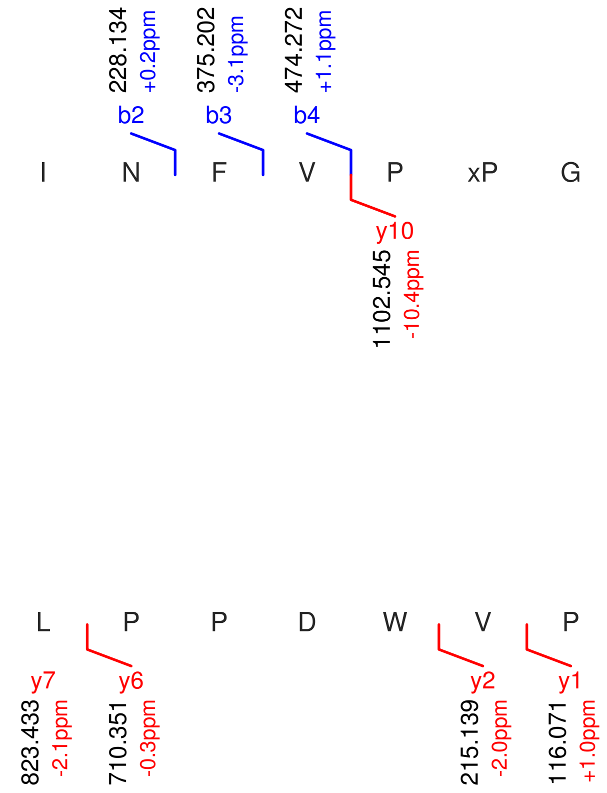

**Figure S7: Modification of the TJ-C-599 precursor peptide by HypXY.** LC-MS analysis of IMAC-purified precursor peptides from *E. coli* BL21(DE3)Tuner containing splicease-encoding plasmid TJ-C-560 (HypXY) and precursor peptide-encoding plasmid TJ-C-599 (Fh8-chaos-HypA7-HypA7) after expression at 24 °C for 22 h in 30 mL cultures after digestion with trypsin. Data from n=3 independent biological replicates. Spliced amino acids have an x as a prefix. Spliced refers to a single tyramine excision. A representative experimental replicate is shown in panels **A, B.**

**A** Close-up plot of the representative extracted ion chromatogram (EIC) of the various peptide states; the blue trace corresponds to the Unmodified peptide; the orange trace corresponds to the Spliced peptide. **B** Close-up plot of the m/z values of the peptides detected by the mass spectrometer, with the shaded colored areas corresponding to where the expected m/z values for the peptide m/z traces should occur, with the observed spectrum plotted in black. The Unmodified, Spliced peptide(s) were found in the mass spectrum. **C** Bar plot showing the relative abundance of the various peptide states at charge z=2; the bars represent the average relative abundances normalized to total peak areas, with each individual experimental replicate value plotted as a black dot. **D** Mirrored $MS^2$ plot for the Unmodified peptide. In this mirror plot, the full fragmentation spectrum is plotted in black pointing upward, and the matched fragment b ions (blue) and y ions (red) are plotted pointing downward, mirrored against it, so that each matched fragment can be compared directly with the measured spectrum. **E** $MS^2$ spectrum of the fragments corresponding to the Unmodified peptide sequence, with matched b (blue, above) and y ions (red, below) plotted. **F** Mirrored $MS^2$ plot for the Spliced peptide. In this mirror plot, the full fragmentation spectrum is plotted in black pointing upward, and the matched fragment b ions (blue) and y ions (red) are plotted pointing downward, mirrored against it, so that each matched fragment can be compared directly with the measured spectrum. **G** $MS^2$ spectrum of the fragments corresponding to the Spliced peptide sequence, with matched b (blue, above) and y ions (red, below) plotted.

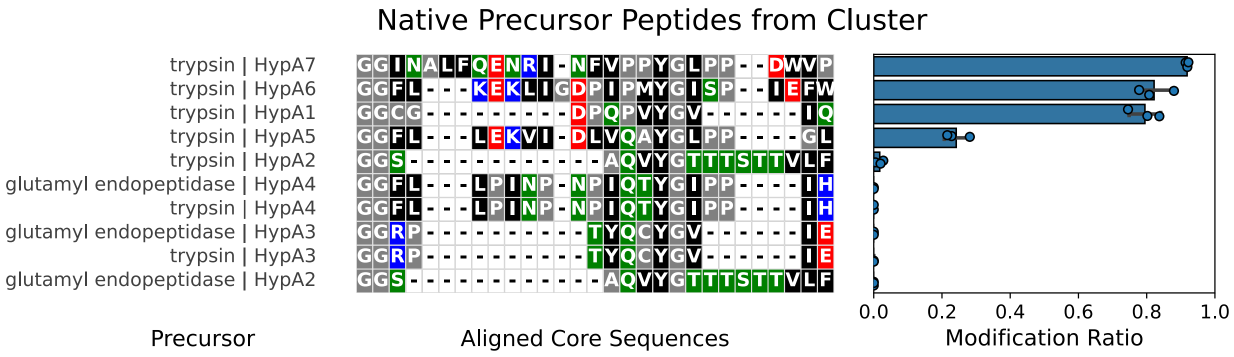

**Figure S8: Modification efficiency of various peptides by the *Hyella patelloides* splicease HypXY.** The precursors were derived from the native *hyp* gene cluster and co-produced alongside the *Hyella* splicease, the precursor peptides were then purified and digested with the indicated protease, and liquid chromatography coupled to mass spectrometry (LC-MS) was used to identify the unmodified and modified precursor fragments. Bars show the mean ± SD of n = 2-3 biological replicates; points are individual biological replicates.

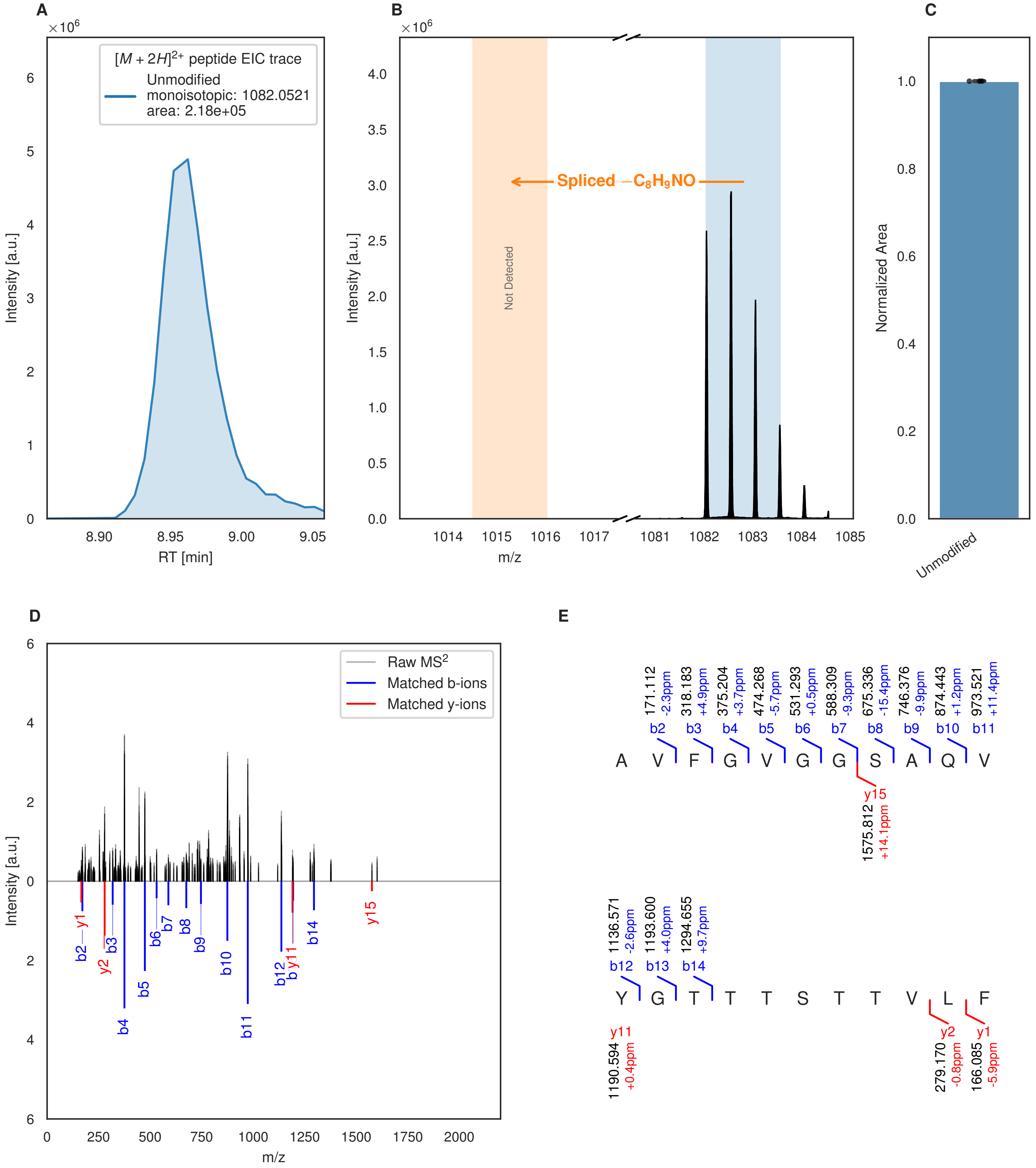

**Figure S9: Modification of the TJ-C-594 precursor peptide by HypXY.** LC-MS analysis of IMAC-purified precursor peptides from *E. coli* BL21(DE3)Tuner containing splicease-encoding plasmid TJ-C-560 (HypXY) and precursor peptide-encoding plasmid TJ-C-594 (Fh8-chaos-HypA2-HypA2) after expression at 24 °C for 22 h in 30 mL cultures after digestion with glutamyl endopeptidase. Data from n=3 independent biological replicates. Spliced amino acids have an x as a prefix. Spliced refers to a single tyramine excision. A representative experimental replicate is shown in panels **A, B.**

**A** Close-up plot of the representative extracted ion chromatogram (EIC) of the various peptide states; the blue trace corresponds to the Unmodified peptide. **B** Close-up plot of the m/z values of the peptides detected by the mass spectrometer, with the shaded colored areas corresponding to where the expected m/z values for the peptide m/z

(Caption continued on the next page.)

traces should occur, with the observed spectrum plotted in black. The Unmodified peptide(s) were found in the mass spectrum and the Spliced peptide(s) were not detected. **C** Bar plot showing the relative abundance of the various peptide states at charge z=2; the bars represent the average relative abundances normalized to total peak areas, with each individual experimental replicate value plotted as a black dot. **D** Mirrored $MS^2$ plot for the Unmodified peptide. In this mirror plot, the full fragmentation spectrum is plotted in black pointing upward, and the matched fragment b ions (blue) and y ions (red) are plotted pointing downward, mirrored against it, so that each matched fragment can be compared directly with the measured spectrum. **E** $MS^2$ spectrum of the fragments corresponding to the Unmodified peptide sequence, with matched b (blue, above) and y ions (red, below) plotted.

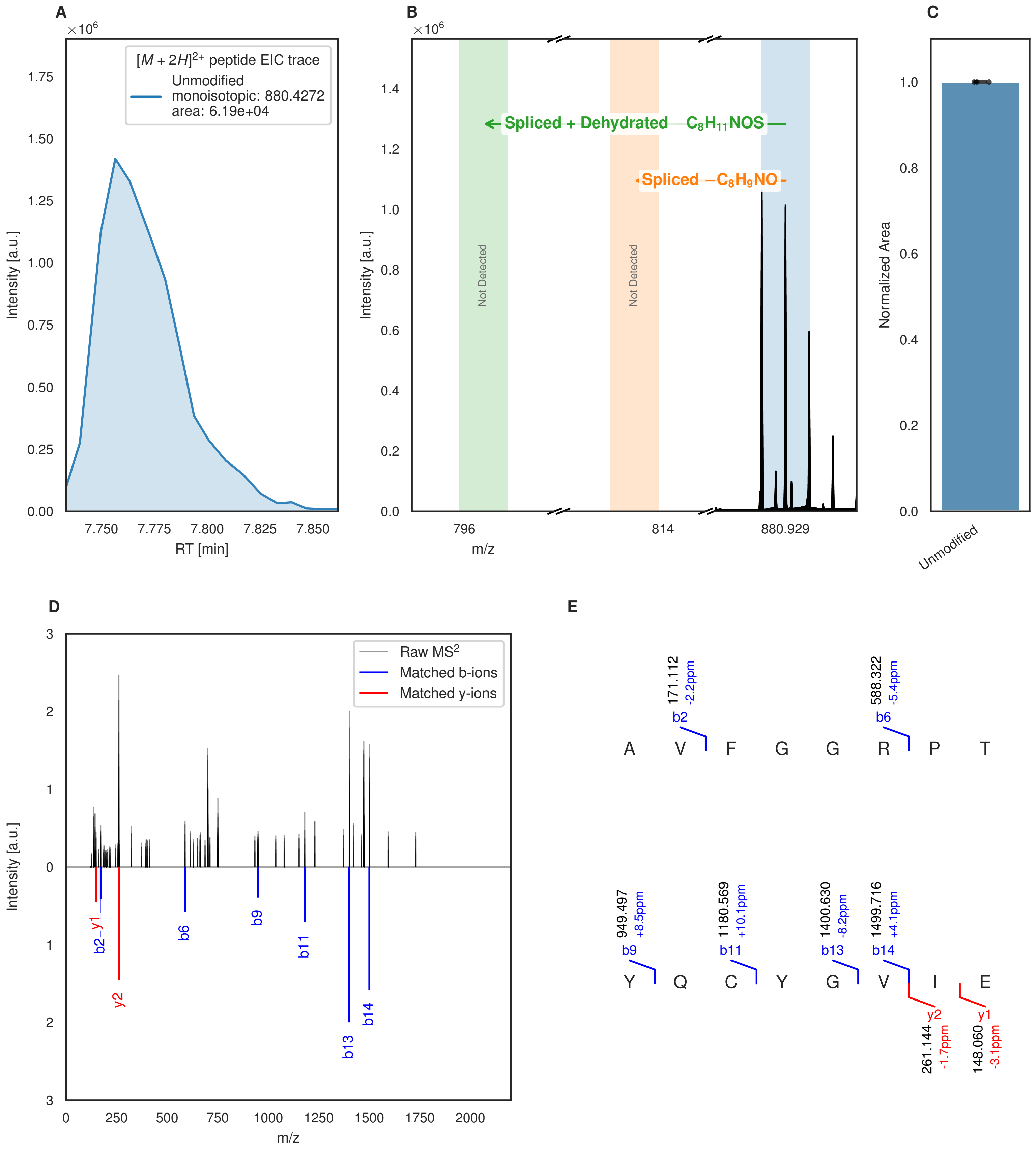

**Figure S10: Modification of the TJ-C-595 precursor peptide by HypXY.** LC-MS analysis of IMAC-purified precursor peptides from *E. coli* BL21(DE3)Tuner containing splicease-encoding plasmid TJ-C-560 (HypXY) and precursor peptide-encoding plasmid TJ-C-595 (Fh8-chaos-HypA3-HypA3) after expression at 24 °C for 22 h in 30 mL cultures after digestion with glutamyl endopeptidase. Data from n=3 independent biological replicates. Spliced amino acids have an x as a prefix. Spliced refers to a single tyramine excision; Spliced + Dehydrated refers to a dehydroalanine-derived ketoamide residue (xdhA). A representative experimental replicate is shown in panels **A, B.** **A** Close-up plot of the representative extracted ion chromatogram (EIC) of the various peptide states; the blue trace corresponds to the Unmodified peptide. **B** Close-up plot of the m/z values of the peptides detected by the mass spectrometer, with the shaded colored areas corresponding to where the expected m/z values for the peptide m/z

(Caption continued on the next page.)

traces should occur, with the observed spectrum plotted in black. The Unmodified peptide(s) were found in the mass spectrum and the Spliced, Spliced + Dehydrated peptide(s) were not detected. **C** Bar plot showing the relative abundance of the various peptide states at charge z=2; the bars represent the average relative abundances normalized to total peak areas, with each individual experimental replicate value plotted as a black dot. **D** Mirrored $MS^2$ plot for the Unmodified peptide. In this mirror plot, the full fragmentation spectrum is plotted in black pointing upward, and the matched fragment b ions (blue) and y ions (red) are plotted pointing downward, mirrored against it, so that each matched fragment can be compared directly with the measured spectrum. **E** $MS^2$ spectrum of the fragments corresponding to the Unmodified peptide sequence, with matched b (blue, above) and y ions (red, below) plotted.

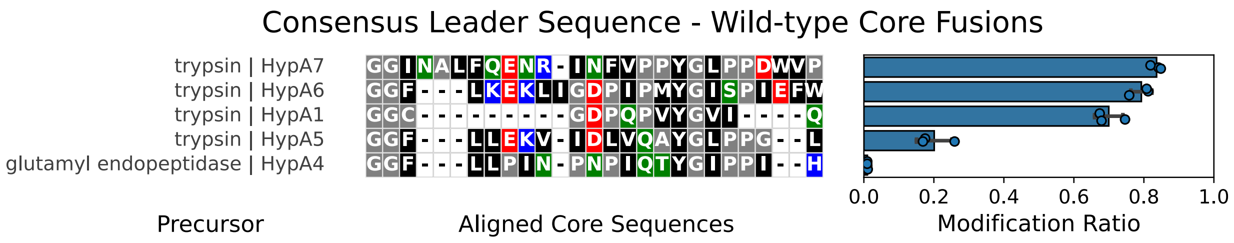

**Figure S11**: **Modification efficiency of various consensus leader-fused peptides by HypXY.** The consensus *Hyella* leader peptide sequence (Table S4) was fused to the wild-type core sequences, and the plasmid encoding the precursors was used alongside a plasmid encoding the splicease to produce the peptides. Precursor peptides were then purified, digested with the indicated protease, and LC-MS was used to identify both unmodified and modified precursor fragments. The overall modification efficiency is comparable to that of the experiment performed with the wild-type precursor peptides (Figure S1). Bars show the mean ± SD of n = 2-3 biological replicates; points are individual biological replicates.

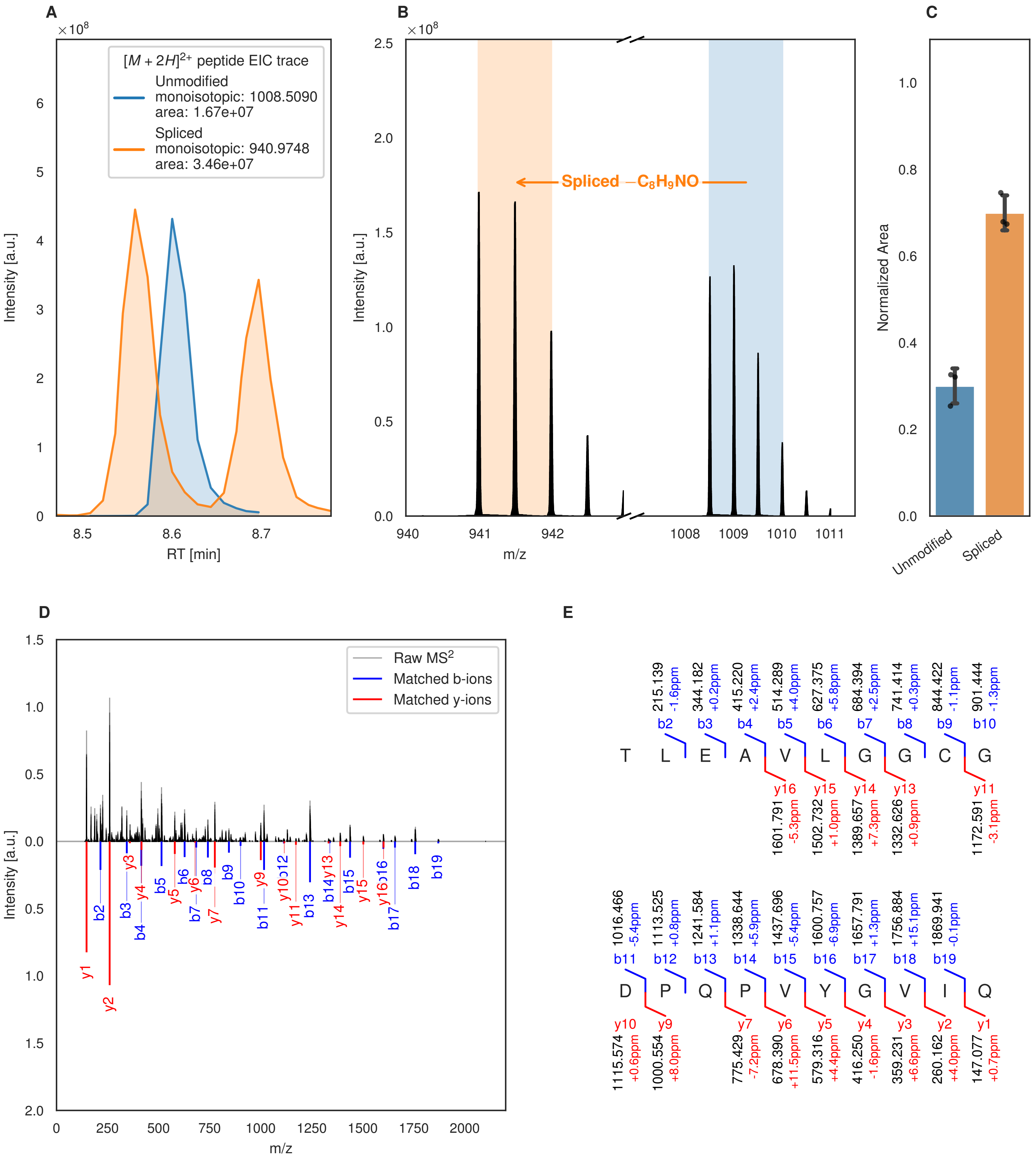
A
×10^8
[M + 2H]^2+ peptide EIC trace
Unmodified
monoisotopic: 1008.5090
area: 1.67e+07
Spliced
monoisotopic: 940.9748
area: 3.46e+07
Intensity [a.u.]
RT [min]
B
×10^8
Spliced −C8H9NO
Intensity [a.u.]
m/z
C
Normalized Area
Unmodified
Spliced
D
Raw MS2
Matched b-ions
Matched y-ions
Intensity [a.u.]
m/z
E
T L E A V L G G C G
b2 215.139 -1.6ppm
b3 344.182 +0.2ppm
b4 415.220 +2.4ppm
b5 514.289 +4.0ppm
b6 627.375 +5.8ppm
b7 684.394 +2.5ppm
b8 741.414 +0.3ppm
b9 844.422 -1.1ppm
b10 901.444 -1.3ppm
y16 1601.791 -5.3ppm
y15 1502.732 +1.0ppm
y14 1389.657 +7.3ppm
y13 1332.626 +0.9ppm
y11 1172.591 -3.1ppm
D P Q P V Y G V I Q
b11 1016.466 -5.4ppm
b12 1113.525 +0.8ppm
b13 1241.584 +1.1ppm
b14 1338.644 +5.9ppm
b15 1437.696 -5.4ppm
b16 1600.757 -6.9ppm
b17 1657.791 +1.3ppm
b18 1756.884 +15.1ppm
b19 1869.941 -0.1ppm
y10 1115.574 +0.6ppm
y9 1000.554 +8.0ppm
y7 775.429 -7.2ppm
y6 678.390 +11.5ppm
y5 579.316 +4.4ppm
y4 416.250 -1.6ppm
y3 359.231 +6.6ppm
y2 260.162 +4.0ppm
y1 147.077 +0.7ppm

Figure S12 (figure continued from the previous page).

F

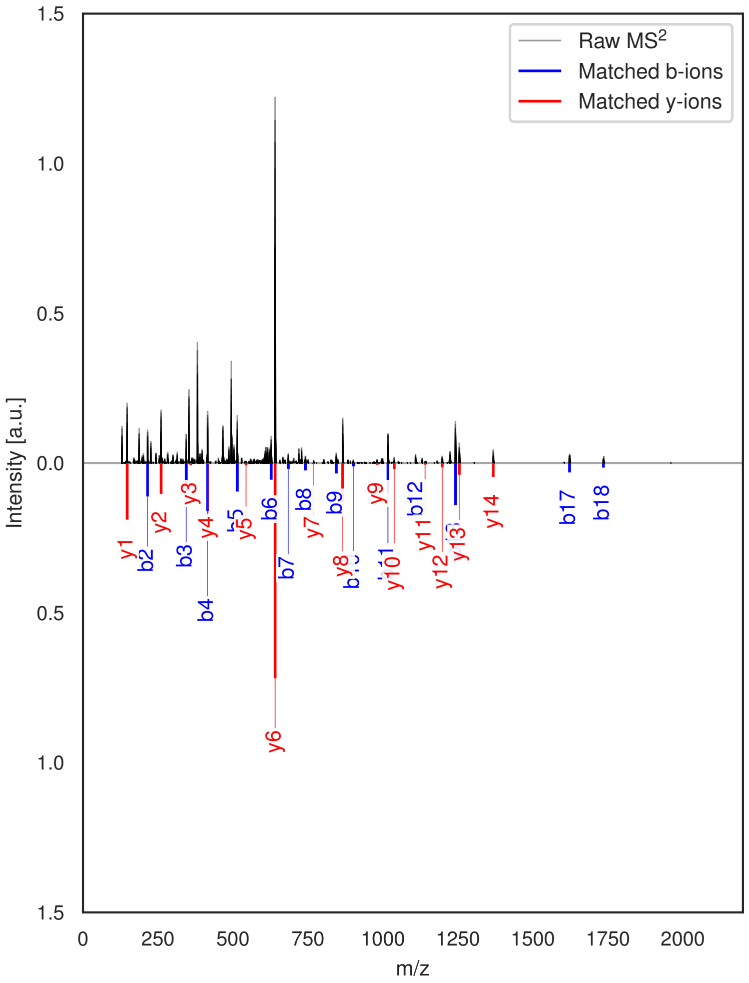

G

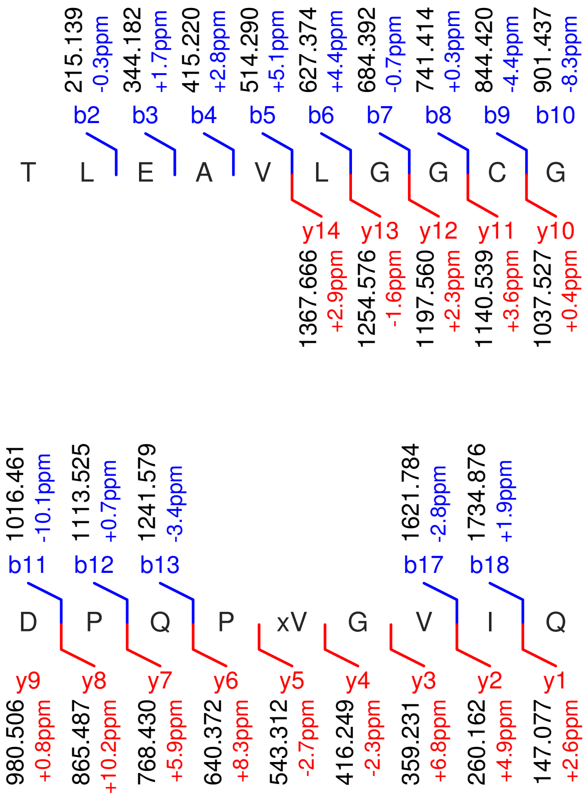

**Figure S12: Modification of the TJ-C-614 precursor peptide by HypXY.** LC-MS analysis of IMAC-purified precursor peptides from *E. coli* BL21(DE3)Tuner containing splicease-encoding plasmid TJ-C-689 (HypXY) and precursor peptide-encoding plasmid TJ-C-614 (Fh8-HypAc-HypA1) after expression at 24 °C for 22 h in 30 mL cultures after digestion with trypsin. Data from n=3 independent biological replicates. Spliced amino acids have an x as a prefix. Spliced refers to a single tyramine excision. A representative experimental replicate is shown in panels **A, B.**

**A** Close-up plot of the representative extracted ion chromatogram (EIC) of the various peptide states; the blue trace corresponds to the Unmodified peptide; the orange trace corresponds to the Spliced peptide. **B** Close-up plot of the m/z values of the peptides detected by the mass spectrometer, with the shaded colored areas corresponding to where the expected m/z values for the peptide m/z traces should occur, with the observed spectrum plotted in black. The Unmodified, Spliced peptide(s) were found in the mass spectrum. **C** Bar plot showing the relative abundance of the various peptide states at charge z=2; the bars represent the average relative abundances normalized to total peak areas, with each individual experimental replicate value plotted as a black dot. **D** Mirrored MS$^2$ plot for the Unmodified peptide. In this mirror plot, the full fragmentation spectrum is plotted in black pointing upward, and the matched fragment b ions (blue) and y ions (red) are plotted pointing downward, mirrored against it, so that each matched fragment can be compared directly with the measured spectrum. **E** MS$^2$ spectrum of the fragments corresponding to the Unmodified peptide sequence, with matched b (blue, above) and y ions (red, below) plotted. **F** Mirrored MS$^2$ plot for the Spliced peptide. In this mirror plot, the full fragmentation spectrum is plotted in black pointing upward, and the matched fragment b ions (blue) and y ions (red) are plotted pointing downward, mirrored against it, so that each matched fragment can be compared directly with the measured spectrum. **G** MS$^2$ spectrum of the fragments corresponding to the Spliced peptide sequence, with matched b (blue, above) and y ions (red, below) plotted.

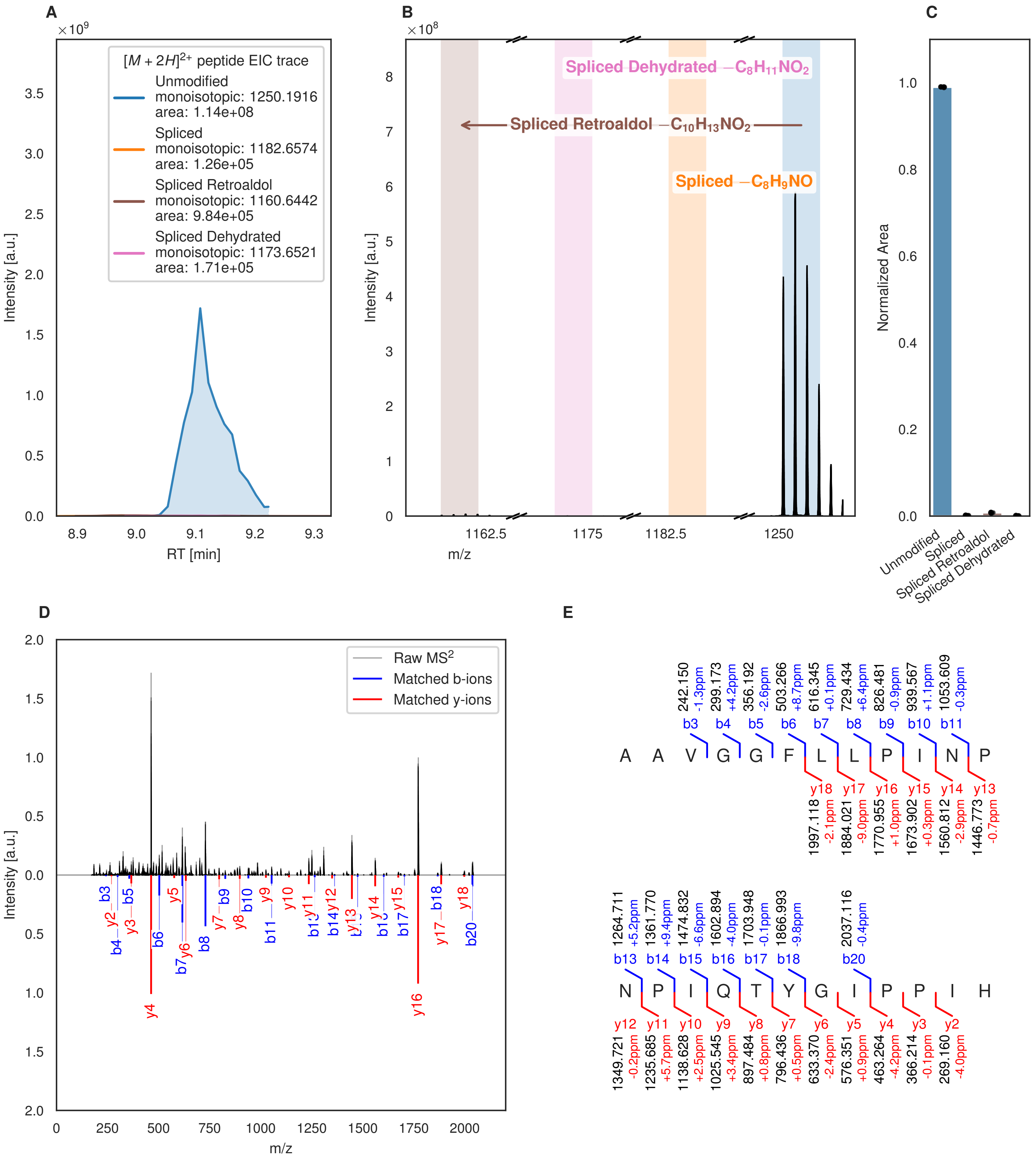
A
B
C
D
E
[M + 2H]2+ peptide EIC trace
Unmodified
monoisotopic: 1250.1916
area: 1.14e+08
Spliced
monoisotopic: 1182.6574
area: 1.26e+05
Spliced Retroaldol
monoisotopic: 1160.6442
area: 9.84e+05
Spliced Dehydrated
monoisotopic: 1173.6521
area: 1.71e+05
Intensity [a.u.]
RT [min]
Spliced Dehydrated −C8H11NO2
Spliced Retroaldol −C10H13NO2
Spliced −C8H9NO
m/z
Normalized Area
Unmodified
Spliced
Spliced Retroaldol
Spliced Dehydrated
Raw MS2
Matched b-ions
Matched y-ions
A A V G G F L L P I N P
N P I Q T Y G I P P I H

Figure S13 (figure continued from the previous page).

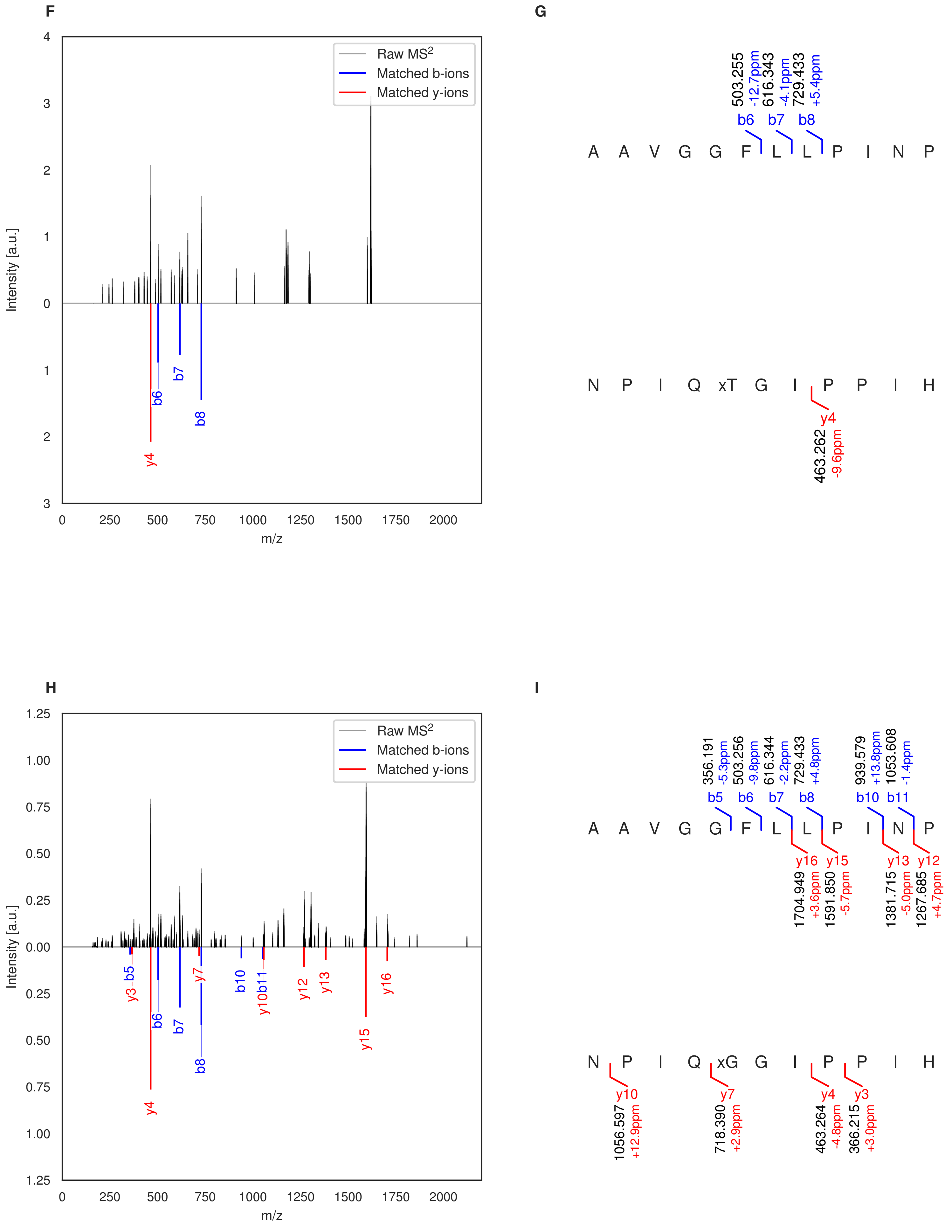

Figure S13 (figure continued from the previous page).

**J**

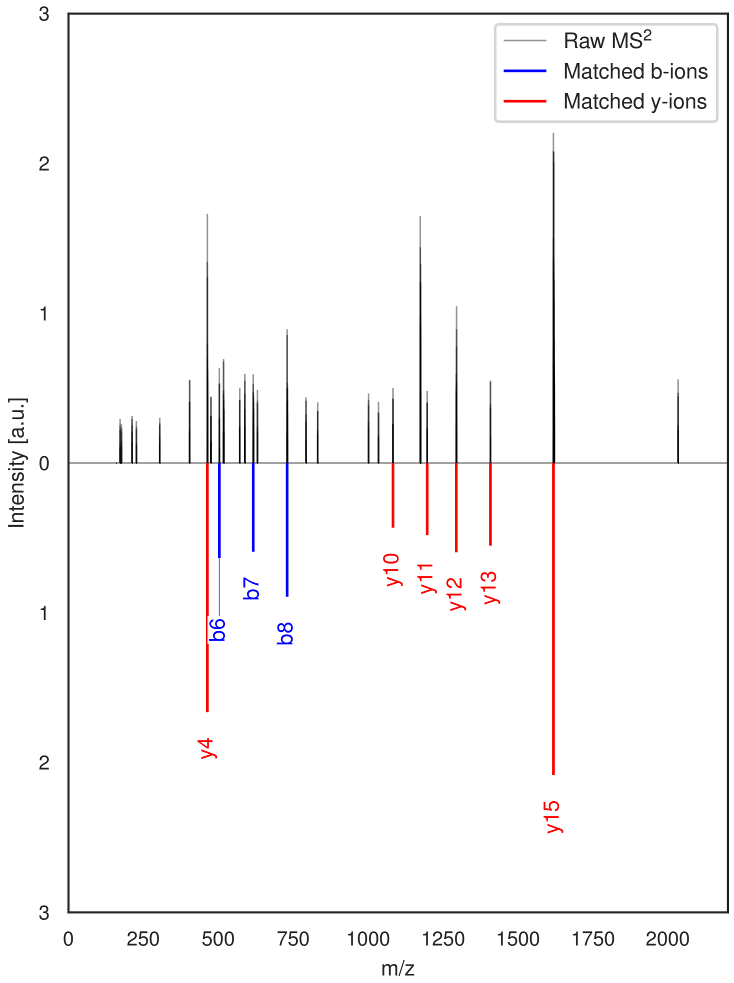

**K**

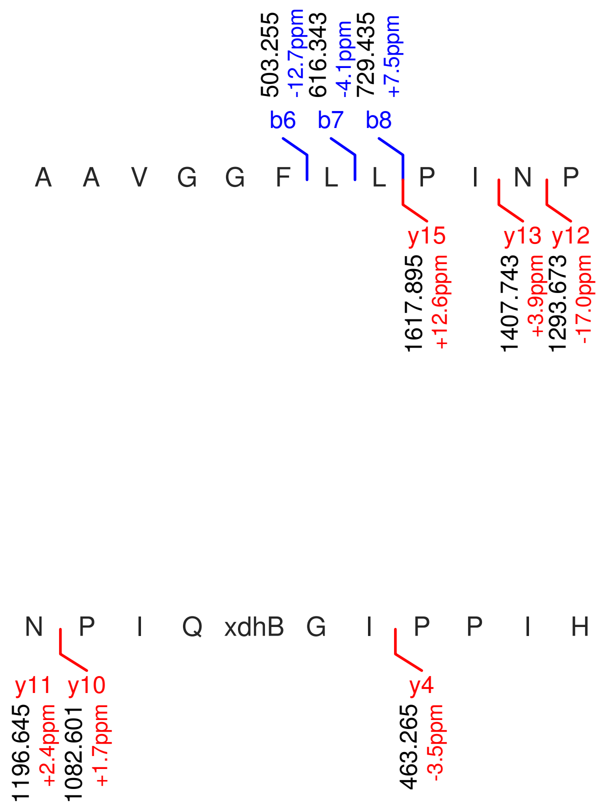

**Figure S13: Modification of the TJ-C-617 precursor peptide by HypXY.** LC-MS analysis of IMAC-purified precursor peptides from *E. coli* BL21(DE3)Tuner containing splicease-encoding plasmid TJ-C-689 (HypXY) and precursor peptide-encoding plasmid TJ-C-617 (Fh8-HypAc-HypA4) after expression at 24 °C for 22 h in 30 mL cultures after digestion with glutamyl endopeptidase. Data from n=5 independent biological replicates. Spliced amino acids have an x as a prefix. Spliced refers to a single tyramine excision; Spliced Retroaldol refers to a glycine-derived ketoamide (xG); Spliced Dehydrated refers to a dehydrobutyrine-derived ketoamide residue (xdhB). A representative experimental replicate is shown in panels **A, B.**

**A** Close-up plot of the representative extracted ion chromatogram (EIC) of the various peptide states; the blue trace corresponds to the Unmodified peptide; the orange trace corresponds to the Spliced peptide; the brown trace corresponds to the Spliced Retroaldol peptide; the pink trace corresponds to the Spliced Dehydrated peptide. **B** Close-up plot of the m/z values of the peptides detected by the mass spectrometer, with the shaded colored areas corresponding to where the expected m/z values for the peptide m/z traces should occur, with the observed spectrum plotted in black. The Unmodified, Spliced, Spliced Retroaldol, Spliced Dehydrated peptide(s) were found in the mass spectrum. **C** Bar plot showing the relative abundance of the various peptide states at charge z=2; the bars represent the average relative abundances normalized to total peak areas, with each individual experimental replicate value plotted as a black dot. **D** Mirrored MS$^2$ plot for the Unmodified peptide. In this mirror plot, the full fragmentation spectrum is plotted in black pointing upward, and the matched fragment b ions (blue) and y ions (red) are plotted pointing downward, mirrored against it, so that each matched fragment can be compared directly with the measured spectrum. **E** MS$^2$ spectrum of the fragments corresponding to the Unmodified peptide sequence, with matched b (blue, above) and y ions (red, below) plotted. **F** Mirrored MS$^2$ plot for the Spliced peptide. In this mirror plot, the full fragmentation spectrum is plotted in black pointing upward, and the matched fragment b ions (blue) and y ions (red) are plotted pointing downward, mirrored against it, so that each matched fragment can be compared directly with the measured spectrum. **G** MS$^2$ spectrum of the fragments corresponding to the Spliced peptide sequence, with matched b (blue, above) and y ions (red, below) plotted. **H** Mirrored MS$^2$ plot for the Spliced Retroaldol peptide. In this mirror plot, the full fragmentation spectrum is plotted in black pointing upward, and the

(Caption continued on the next page.)

matched fragment b ions (blue) and y ions (red) are plotted pointing downward, mirrored against it, so that each matched fragment can be compared directly with the measured spectrum. **I** $MS^2$ spectrum of the fragments corresponding to the Spliced Retroaldol peptide sequence, with matched b (blue, above) and y ions (red, below) plotted. **J** Mirrored $MS^2$ plot for the Spliced Dehydrated peptide. In this mirror plot, the full fragmentation spectrum is plotted in black pointing upward, and the matched fragment b ions (blue) and y ions (red) are plotted pointing downward, mirrored against it, so that each matched fragment can be compared directly with the measured spectrum. **K** $MS^2$ spectrum of the fragments corresponding to the Spliced Dehydrated peptide sequence, with matched b (blue, above) and y ions (red, below) plotted.

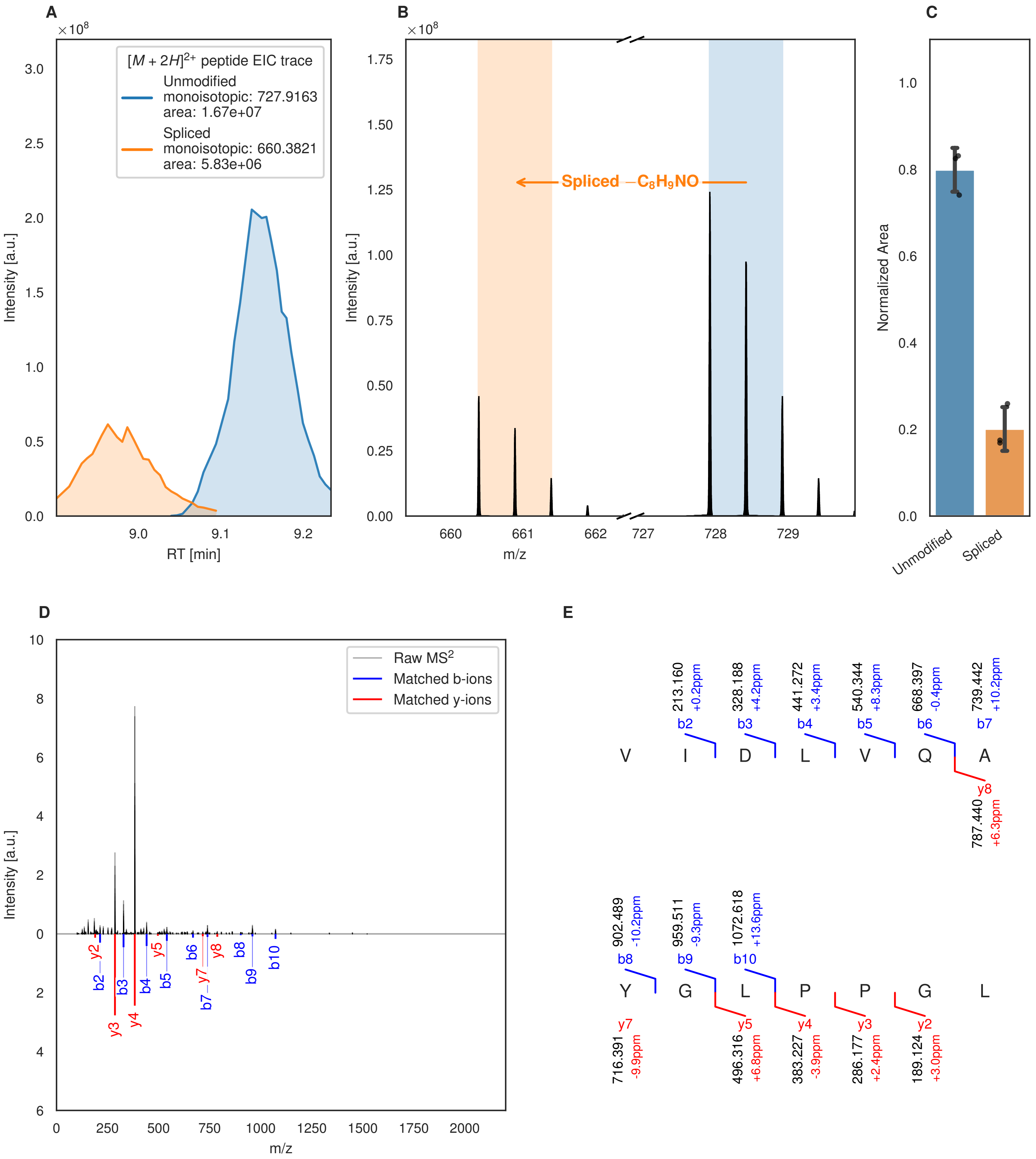

A
×10^8
[M + 2H]2+ peptide EIC trace
Unmodified
monoisotopic: 727.9163
area: 1.67e+07
Spliced
monoisotopic: 660.3821
area: 5.83e+06
Intensity [a.u.]
RT [min]
B
×10^8
Spliced −C8H9NO
Intensity [a.u.]
m/z
C
Normalized Area
Unmodified
Spliced
D
Raw MS2
Matched b-ions
Matched y-ions
Intensity [a.u.]
m/z
E
213.160 +0.2ppm b2
328.188 +4.2ppm b3
441.272 +3.4ppm b4
540.344 +8.3ppm b5
668.397 -0.4ppm b6
739.442 +10.2ppm b7
V I D L V Q A
y8 787.440 +6.3ppm
902.489 -10.2ppm b8
959.511 -9.3ppm b9
1072.618 +13.6ppm b10
Y G L P P G L
y7 716.391 -9.9ppm
y5 496.316 +6.8ppm
y4 383.227 -3.9ppm
y3 286.177 +2.4ppm
y2 189.124 +3.0ppm

Figure S14 (figure continued from the previous page).

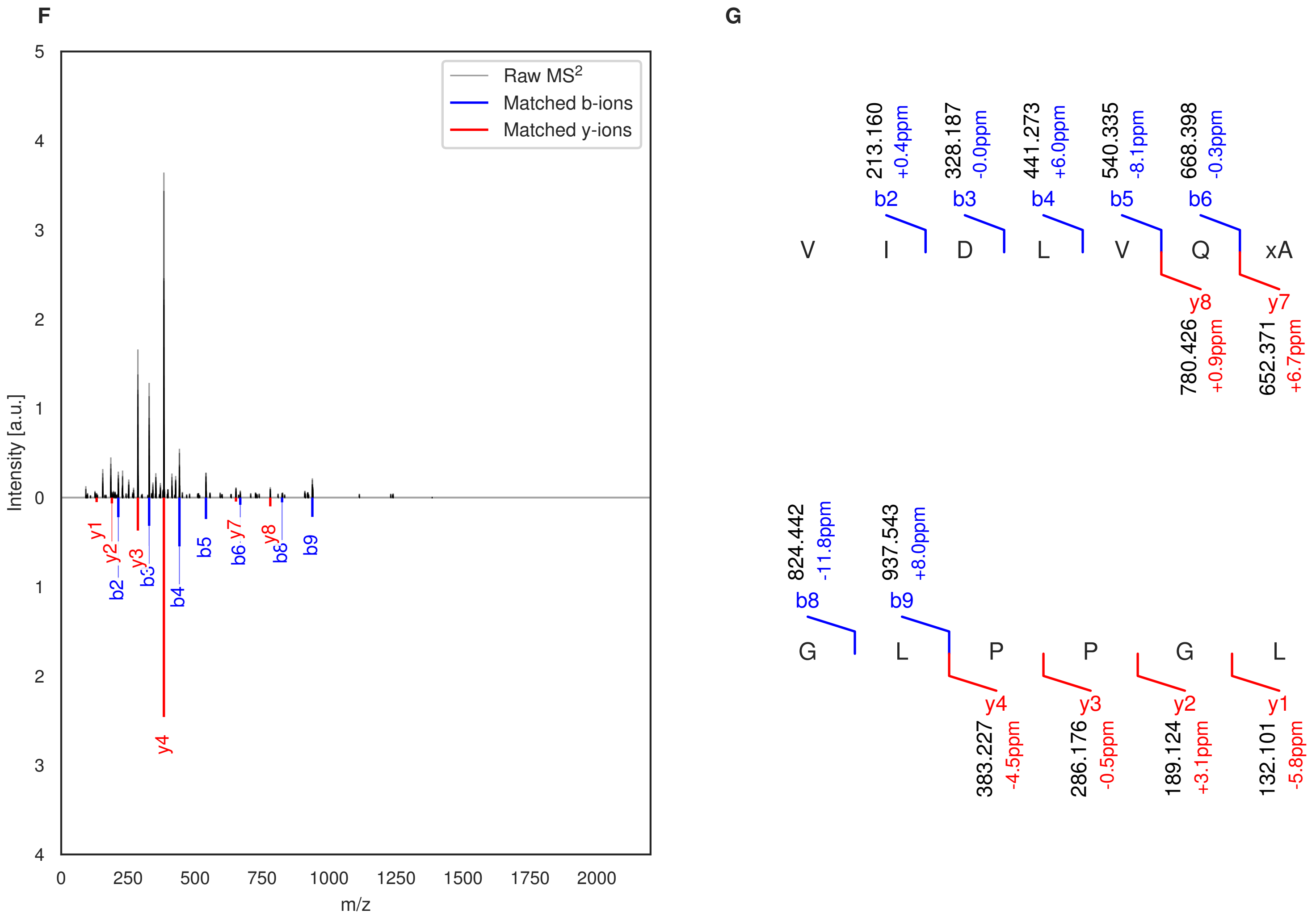

**Figure S14: Modification of the TJ-C-618 precursor peptide by HypXY.** LC-MS analysis of IMAC-purified precursor peptides from *E. coli* BL21(DE3)Tuner containing splicease-encoding plasmid TJ-C-689 (HypXY) and precursor peptide-encoding plasmid TJ-C-618 (Fh8-HypAc-HypA5) after expression at 24 °C for 22 h in 30 mL cultures after digestion with trypsin. Data from n=3 independent biological replicates. Spliced amino acids have an x as a prefix. Spliced refers to a single tyramine excision. A representative experimental replicate is shown in panels **A, B.**

**A** Close-up plot of the representative extracted ion chromatogram (EIC) of the various peptide states; the blue trace corresponds to the Unmodified peptide; the orange trace corresponds to the Spliced peptide. **B** Close-up plot of the m/z values of the peptides detected by the mass spectrometer, with the shaded colored areas corresponding to where the expected m/z values for the peptide m/z traces should occur, with the observed spectrum plotted in black. The Unmodified, Spliced peptide(s) were found in the mass spectrum. **C** Bar plot showing the relative abundance of the various peptide states at charge z=2; the bars represent the average relative abundances normalized to total peak areas, with each individual experimental replicate value plotted as a black dot. **D** Mirrored $MS^2$ plot for the Unmodified peptide. In this mirror plot, the full fragmentation spectrum is plotted in black pointing upward, and the matched fragment b ions (blue) and y ions (red) are plotted pointing downward, mirrored against it, so that each matched fragment can be compared directly with the measured spectrum. **E** $MS^2$ spectrum of the fragments corresponding to the Unmodified peptide sequence, with matched b (blue, above) and y ions (red, below) plotted. **F** Mirrored $MS^2$ plot for the Spliced peptide. In this mirror plot, the full fragmentation spectrum is plotted in black pointing upward, and the matched fragment b ions (blue) and y ions (red) are plotted pointing downward, mirrored against it, so that each matched fragment can be compared directly with the measured spectrum. **G** $MS^2$ spectrum of the fragments corresponding to the Spliced peptide sequence, with matched b (blue, above) and y ions (red, below) plotted.

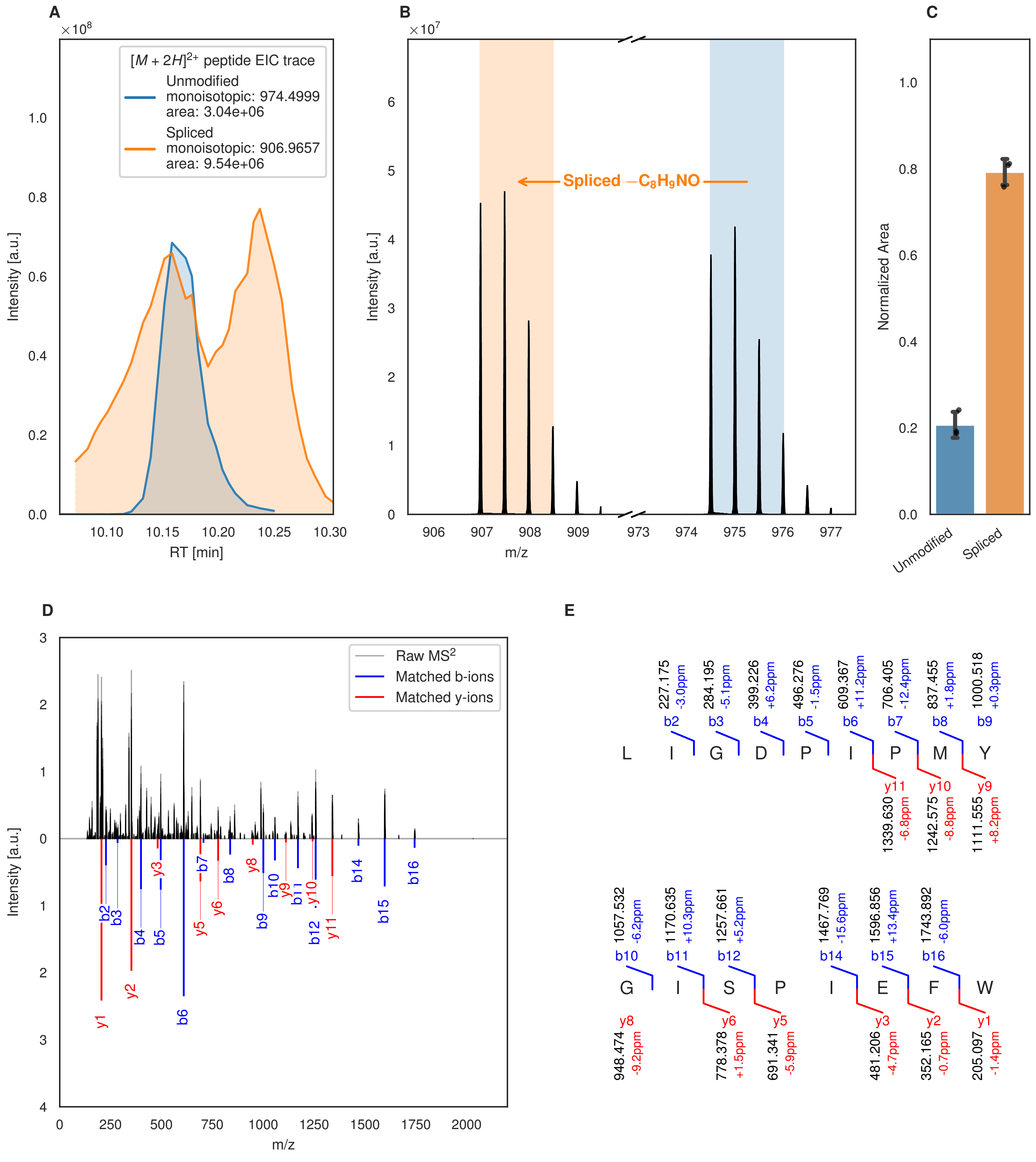
A
B
C
D
E
[M + 2H]2+ peptide EIC trace
Unmodified
monoisotopic: 974.4999
area: 3.04e+06
Spliced
monoisotopic: 906.9657
area: 9.54e+06
Intensity [a.u.]
RT [min]
Spliced −C8H9NO
m/z
Normalized Area
Unmodified
Spliced
Raw MS2
Matched b-ions
Matched y-ions
L I G D P I P M Y
G I S P I E F W
b2 227.175 -3.0ppm
b3 284.195 -5.1ppm
b4 399.226 +6.2ppm
b5 496.276 -1.5ppm
b6 609.367 +11.2ppm
b7 706.405 -12.4ppm
b8 837.455 +1.8ppm
b9 1000.518 +0.3ppm
y11 1339.630 -6.8ppm
y10 1242.575 -8.8ppm
y9 1111.555 +8.2ppm
b10 1057.532 -6.2ppm
b11 1170.635 +10.3ppm
b12 1257.661 +5.2ppm
b14 1467.769 -15.6ppm
b15 1596.856 +13.4ppm
b16 1743.892 -6.0ppm
y8 948.474 -9.2ppm
y6 778.378 +1.5ppm
y5 691.341 -5.9ppm
y3 481.206 -4.7ppm
y2 352.165 -0.7ppm
y1 205.097 -1.4ppm

Figure S15 (figure continued from the previous page).

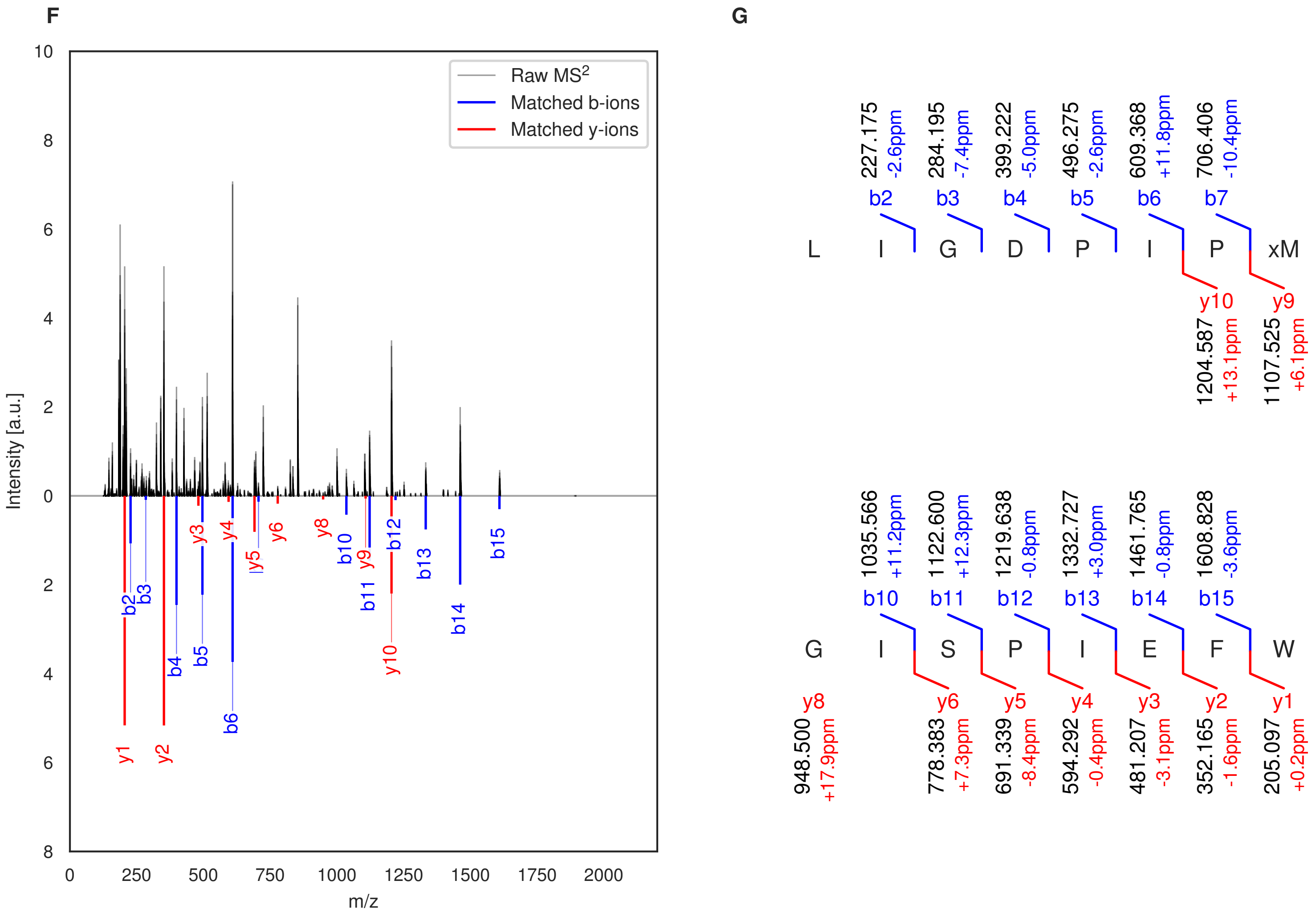

**Figure S15: Modification of the TJ-C-619 precursor peptide by HypXY.** LC-MS analysis of IMAC-purified precursor peptides from *E. coli* BL21(DE3)Tuner containing splicease-encoding plasmid TJ-C-689 (HypXY) and precursor peptide-encoding plasmid TJ-C-619 (Fh8-HypAc-HypA6) after expression at 24 °C for 22 h in 30 mL cultures after digestion with trypsin. Data from n=3 independent biological replicates. Spliced amino acids have an x as a prefix. Spliced refers to a single tyramine excision. A representative experimental replicate is shown in panels **A, B.**

**A** Close-up plot of the representative extracted ion chromatogram (EIC) of the various peptide states; the blue trace corresponds to the Unmodified peptide; the orange trace corresponds to the Spliced peptide. **B** Close-up plot of the m/z values of the peptides detected by the mass spectrometer, with the shaded colored areas corresponding to where the expected m/z values for the peptide m/z traces should occur, with the observed spectrum plotted in black. The Unmodified, Spliced peptide(s) were found in the mass spectrum. **C** Bar plot showing the relative abundance of the various peptide states at charge z=2; the bars represent the average relative abundances normalized to total peak areas, with each individual experimental replicate value plotted as a black dot. **D** Mirrored MS$^2$ plot for the Unmodified peptide. In this mirror plot, the full fragmentation spectrum is plotted in black pointing upward, and the matched fragment b ions (blue) and y ions (red) are plotted pointing downward, mirrored against it, so that each matched fragment can be compared directly with the measured spectrum. **E** MS$^2$ spectrum of the fragments corresponding to the Unmodified peptide sequence, with matched b (blue, above) and y ions (red, below) plotted. **F** Mirrored MS$^2$ plot for the Spliced peptide. In this mirror plot, the full fragmentation spectrum is plotted in black pointing upward, and the matched fragment b ions (blue) and y ions (red) are plotted pointing downward, mirrored against it, so that each matched fragment can be compared directly with the measured spectrum. **G** MS$^2$ spectrum of the fragments corresponding to the Spliced peptide sequence, with matched b (blue, above) and y ions (red, below) plotted.

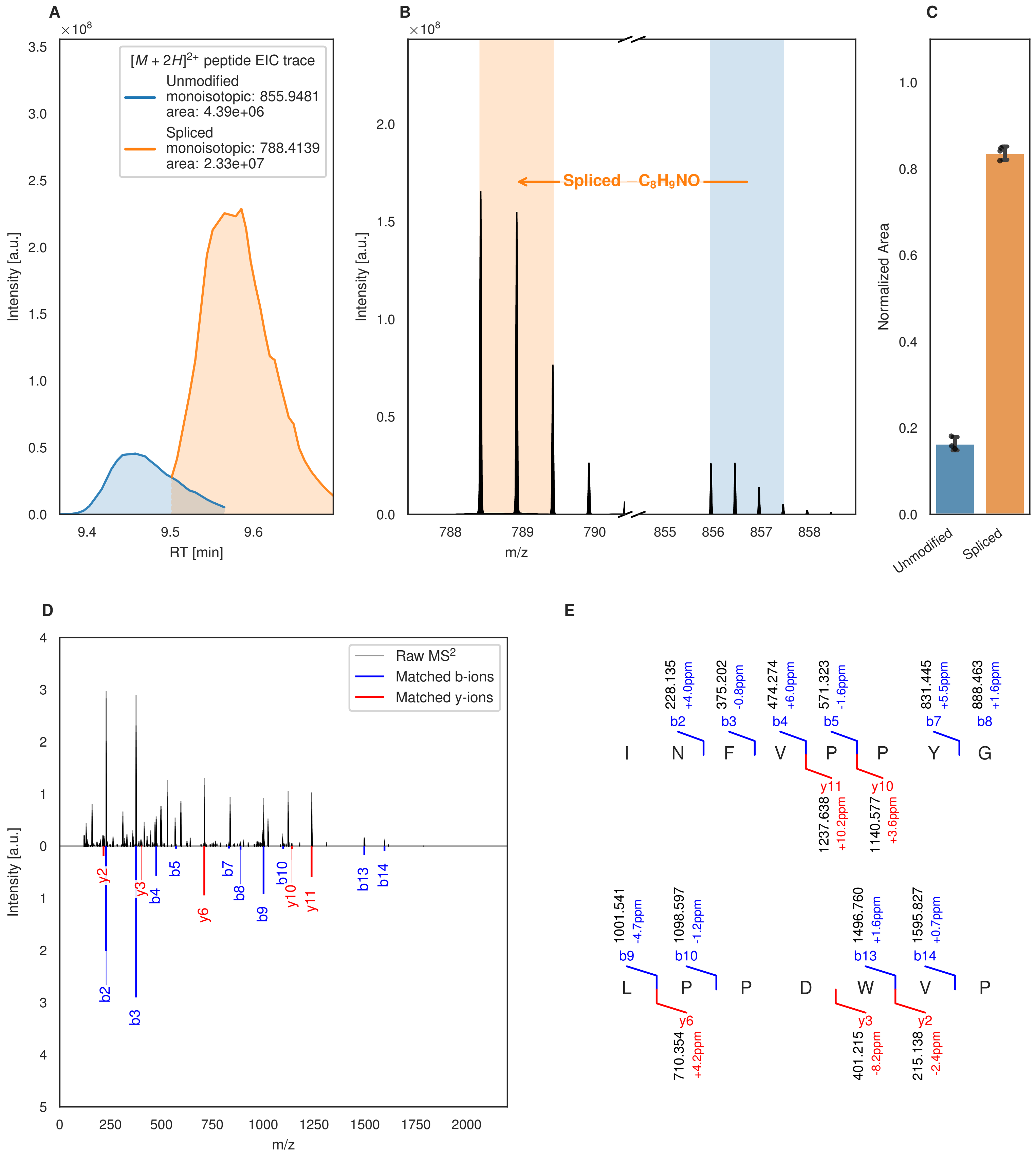
A
B
C
D
E
$[M+2H]^{2+}$ peptide EIC trace
Unmodified
monoisotopic: 855.9481
area: 4.39e+06
Spliced
monoisotopic: 788.4139
area: 2.33e+07
Intensity [a.u.]
RT [min]
Spliced $-C_8H_9NO$
m/z
Normalized Area
Unmodified
Spliced
Raw $MS^2$
Matched b-ions
Matched y-ions
I N F V P P Y G
L P P D W V P
b2 228.135 +4.0ppm
b3 375.202 -0.8ppm
b4 474.274 +6.0ppm
b5 571.323 -1.6ppm
b7 831.445 +5.5ppm
b8 888.463 +1.6ppm
y11 1237.638 +10.2ppm
y10 1140.577 +3.6ppm
b9 1001.541 -4.7ppm
b10 1098.597 -1.2ppm
b13 1496.760 +1.6ppm
b14 1595.827 +0.7ppm
y6 710.354 +4.2ppm
y3 401.215 -8.2ppm
y2 215.138 -2.4ppm

Figure S16 (figure continued from the previous page).

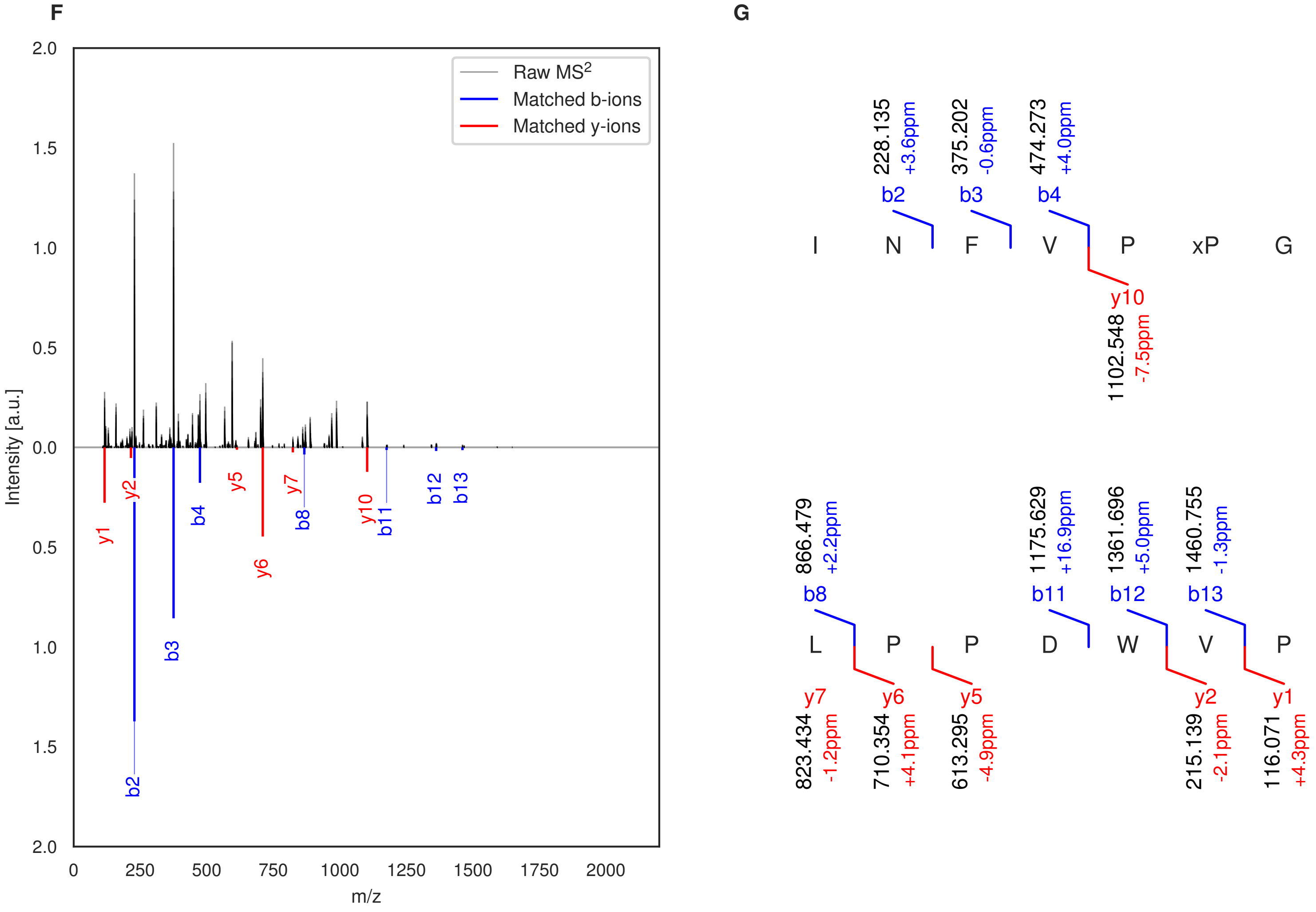

**Figure S16: Modification of the TJ-C-620 precursor peptide by HypXY.** LC-MS analysis of IMAC-purified precursor peptides from *E. coli* BL21(DE3)Tuner containing splicease-encoding plasmid TJ-C-689 (HypXY) and precursor peptide-encoding plasmid TJ-C-620 (Fh8-HypAc-HypA7) after expression at 24 °C for 22 h in 30 mL cultures after digestion with trypsin. Data from n=3 independent biological replicates. Spliced amino acids have an x as a prefix. Spliced refers to a single tyramine excision. A representative experimental replicate is shown in panels **A, B.**

**A** Close-up plot of the representative extracted ion chromatogram (EIC) of the various peptide states; the blue trace corresponds to the Unmodified peptide; the orange trace corresponds to the Spliced peptide. **B** Close-up plot of the m/z values of the peptides detected by the mass spectrometer, with the shaded colored areas corresponding to where the expected m/z values for the peptide m/z traces should occur, with the observed spectrum plotted in black. The Unmodified, Spliced peptide(s) were found in the mass spectrum. **C** Bar plot showing the relative abundance of the various peptide states at charge z=2; the bars represent the average relative abundances normalized to total peak areas, with each individual experimental replicate value plotted as a black dot. **D** Mirrored $MS^2$ plot for the Unmodified peptide. In this mirror plot, the full fragmentation spectrum is plotted in black pointing upward, and the matched fragment b ions (blue) and y ions (red) are plotted pointing downward, mirrored against it, so that each matched fragment can be compared directly with the measured spectrum. **E** $MS^2$ spectrum of the fragments corresponding to the Unmodified peptide sequence, with matched b (blue, above) and y ions (red, below) plotted. **F** Mirrored $MS^2$ plot for the Spliced peptide. In this mirror plot, the full fragmentation spectrum is plotted in black pointing upward, and the matched fragment b ions (blue) and y ions (red) are plotted pointing downward, mirrored against it, so that each matched fragment can be compared directly with the measured spectrum. **G** $MS^2$ spectrum of the fragments corresponding to the Spliced peptide sequence, with matched b (blue, above) and y ions (red, below) plotted.

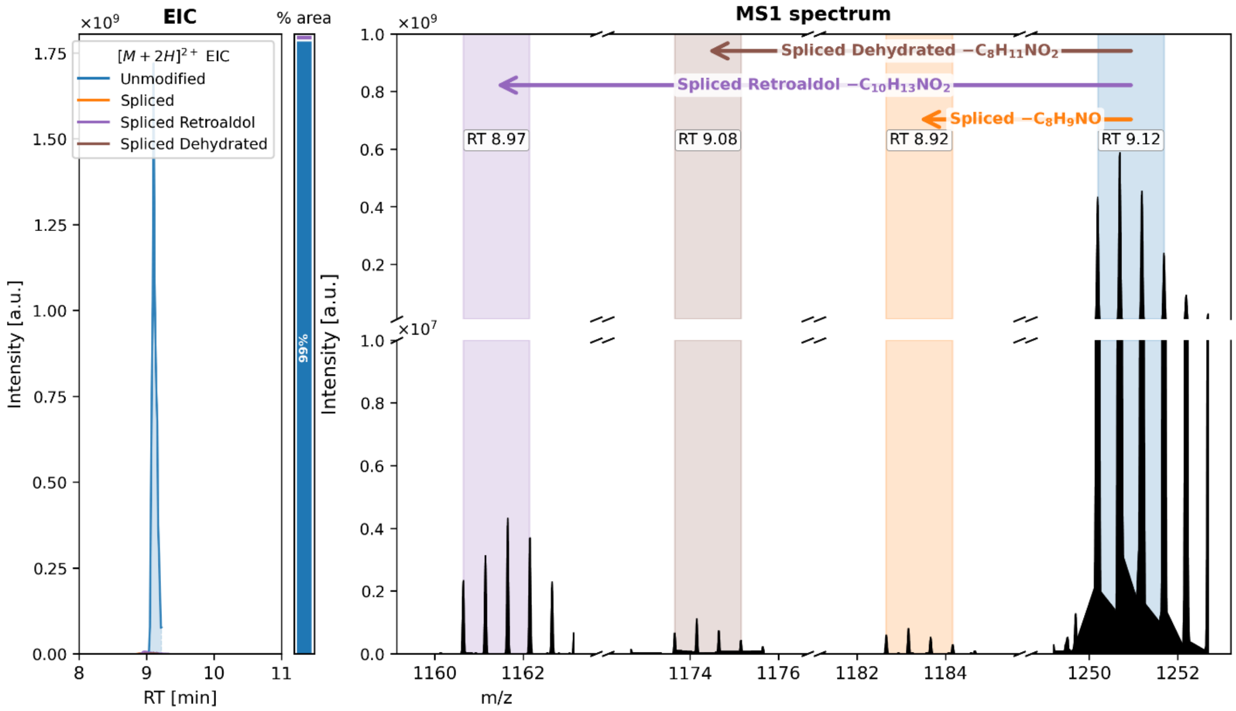

**Figure S17: HypA4 splicing by HypXY assessed by GluC digestion and LC-MS.** Left: Extracted Ion Chromatogram (EIC) of the GluC-digested peptide fragments with charge z=2 for the different modification states. The trace is dominated by the unmodified fragment, as it is much more abundant than the modified products. Right: MS spectra corresponding to the peak apex for the various EICs, with the corresponding retention time range indicated in the shaded area as text in a white box. The data show a fragment corresponding to the unmodified precursor peptide HypA4 with a small amount of fragments matching several modified products. The label "Spliced" refers to a single tyramine excision; "Spliced Dehydrated" refers to a dehydrobutyrine-derived ketoamide residue; "Spliced Retroaldol" refers to a glycine-derived ketoamide.

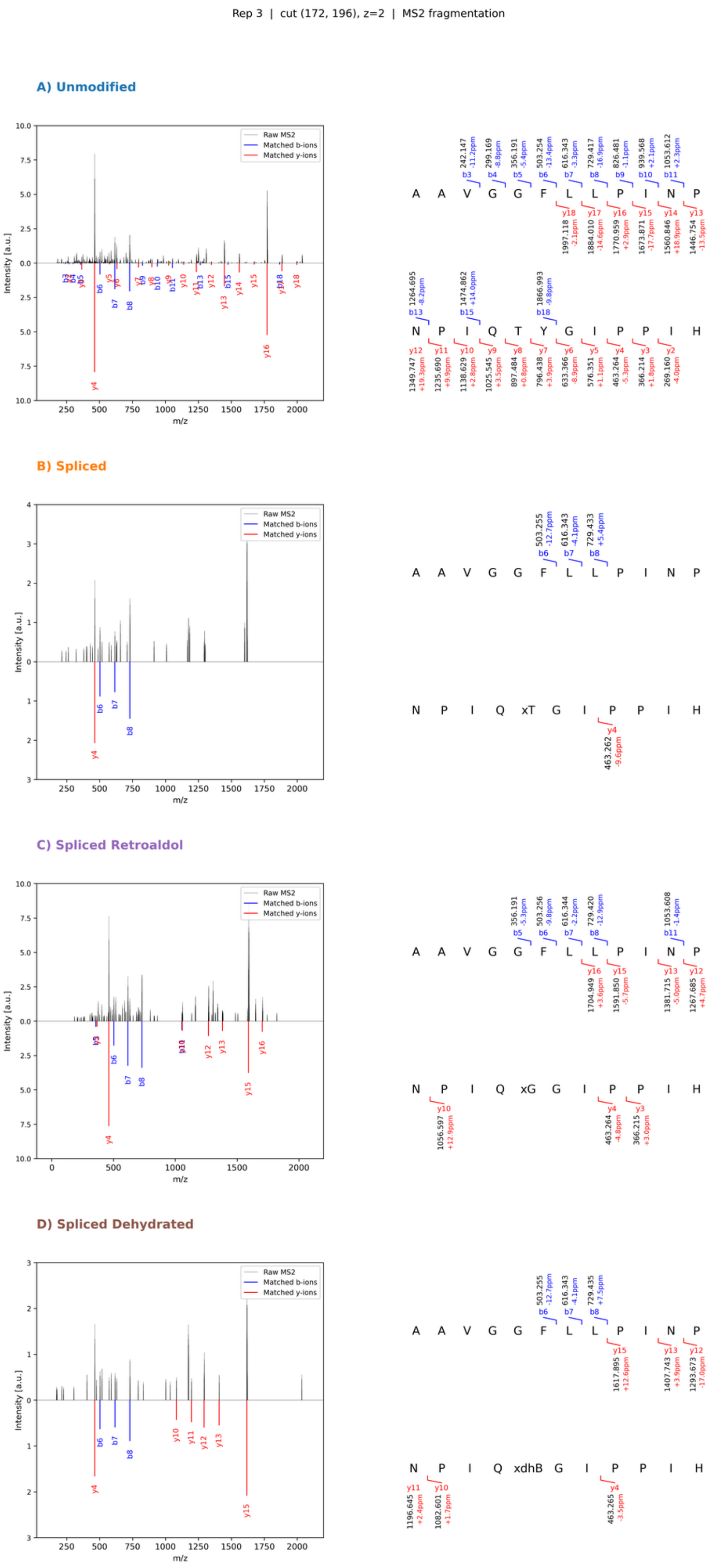

**Figure S18: HypA4 splicing by HypXY assessed via GluC digestion and MS$^2$ analysis.** MS$^2$ fragmentation corresponding to the unmodified precursor HypA4 with a small amount of fragments matching several modified products. Left: mirror plot showing the fragments resulting from isolating the precursor m/z matching the fragment, with data plotted in black on the upper half of the plot. The lower half of the plot shows the b and y ions matching the fragment from the overall fragment pool. Right: the fragment sequence and matching b and y ions. Non-canonical amino acids are represented via lower case letters followed by a single capital letter. The prefix x refers to a spliced amino acid, and represents the extra keto group added to the amino acid. The non-canonical α-keto-β-enamino-dehydrobutyrine is represented by xdhB. The label “Spliced” refers to a single tyramine excision, “Spliced Dehydrated” refers to an α-keto-β-enamino-dehydrobutyrine, “Spliced Retroaldol” refers to an α-keto-glycine.

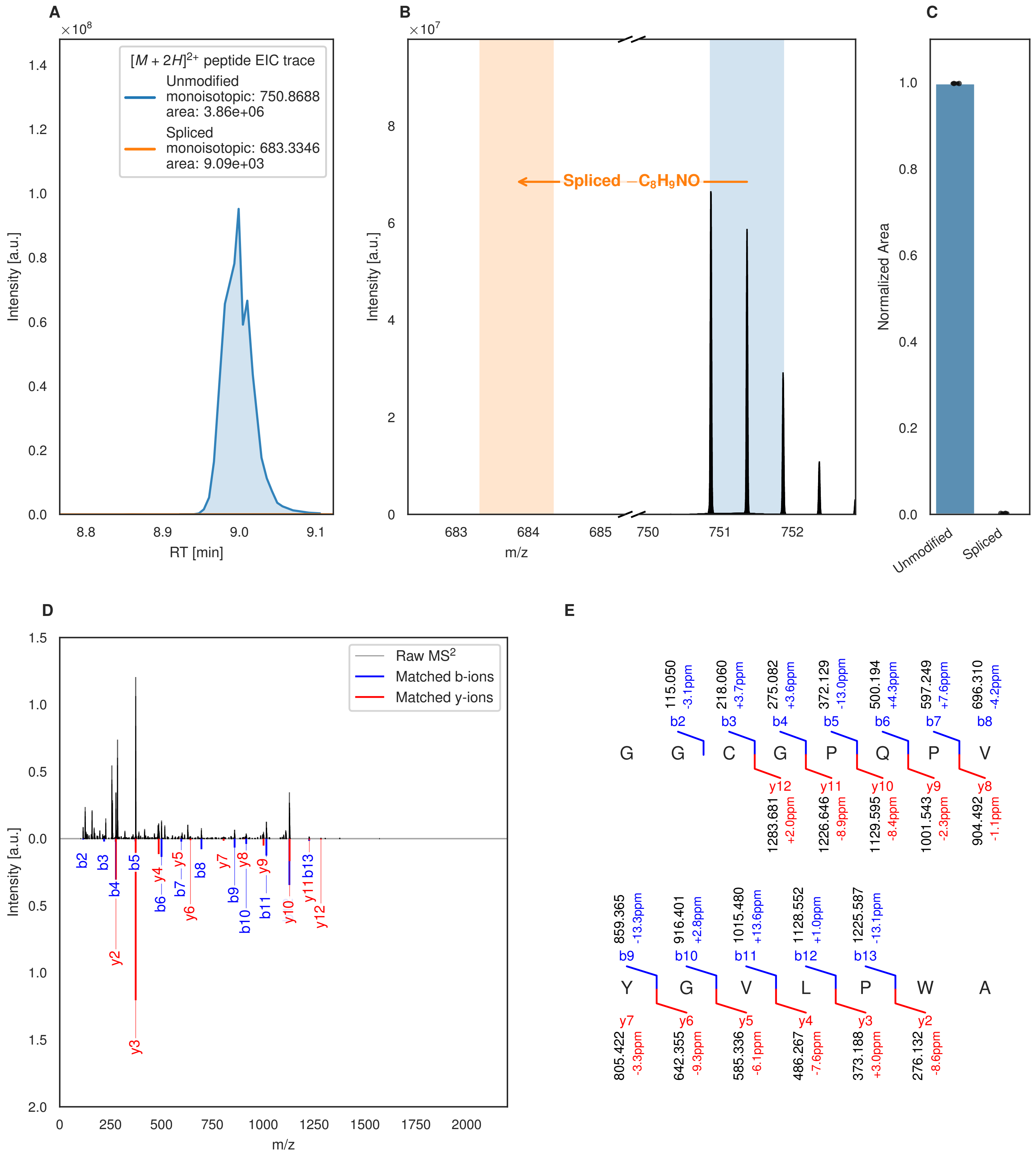

A
×10^8
[M + 2H]2+ peptide EIC trace
Unmodified
monoisotopic: 750.8688
area: 3.86e+06
Spliced
monoisotopic: 683.3346
area: 9.09e+03
Intensity [a.u.]
RT [min]
B
×10^7
Spliced −C8H9NO
Intensity [a.u.]
m/z
C
Normalized Area
Unmodified
Spliced
D
Raw MS2
Matched b-ions
Matched y-ions
Intensity [a.u.]
m/z
E
G G C G P Q P V
Y G V L P W A

Figure S19 (figure continued from the previous page).

**F**

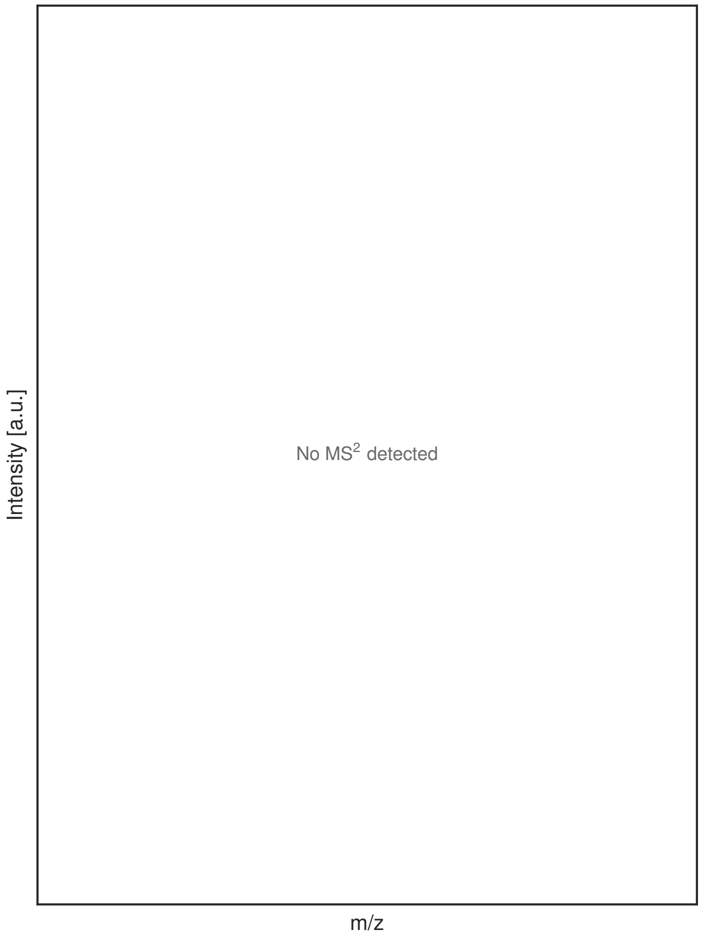

**G**

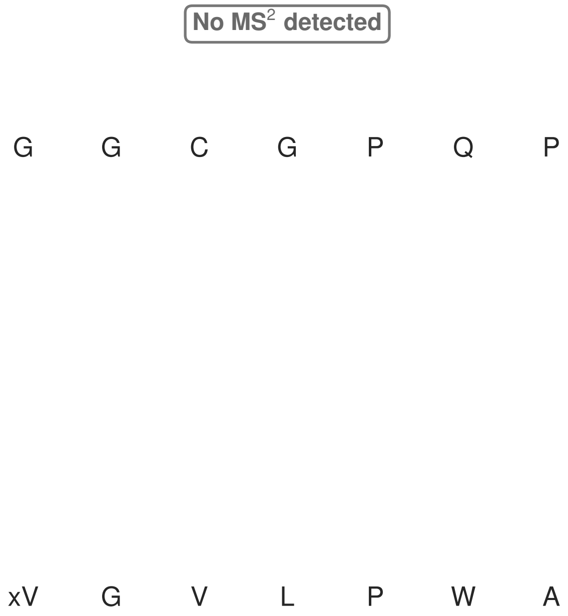

**Figure S19: Modification of the TJ-C-693 precursor peptide by HypXY.** LC-MS analysis of IMAC-purified precursor peptides from *E. coli* BL21(DE3)Tuner containing splicease-encoding plasmid TJ-C-560 (HypXY) and precursor peptide-encoding plasmid TJ-C-693 (Fh8-HypAc-TEV-Artificial-1) after expression at 24 °C for 22 h in 30 mL cultures after digestion with TEV. Data from n=2 independent biological replicates. Spliced amino acids have an x as a prefix. Spliced refers to a single tyramine excision. A representative experimental replicate is shown in panels **A, B.**

**A** Close-up plot of the representative extracted ion chromatogram (EIC) of the various peptide states; the blue trace corresponds to the Unmodified peptide; the orange trace corresponds to the Spliced peptide. **B** Close-up plot of the m/z values of the peptides detected by the mass spectrometer, with the shaded colored areas corresponding to where the expected m/z values for the peptide m/z traces should occur, with the observed spectrum plotted in black. The Unmodified, Spliced peptide(s) were found in the mass spectrum. **C** Bar plot showing the relative abundance of the various peptide states at charge z=2; the bars represent the average relative abundances normalized to total peak areas, with each individual experimental replicate value plotted as a black dot. **D** Mirrored $MS^2$ plot for the Unmodified peptide. In this mirror plot, the full fragmentation spectrum is plotted in black pointing upward, and the matched fragment b ions (blue) and y ions (red) are plotted pointing downward, mirrored against it, so that each matched fragment can be compared directly with the measured spectrum. **E** $MS^2$ spectrum of the fragments corresponding to the Unmodified peptide sequence, with matched b (blue, above) and y ions (red, below) plotted. **F** Mirrored $MS^2$ plot for the Spliced peptide. In this mirror plot, the full fragmentation spectrum is plotted in black pointing upward, and the matched fragment b ions (blue) and y ions (red) are plotted pointing downward, mirrored against it, so that each matched fragment can be compared directly with the measured spectrum. **G** $MS^2$ spectrum of the fragments corresponding to the Spliced peptide sequence, with matched b (blue, above) and y ions (red, below) plotted.

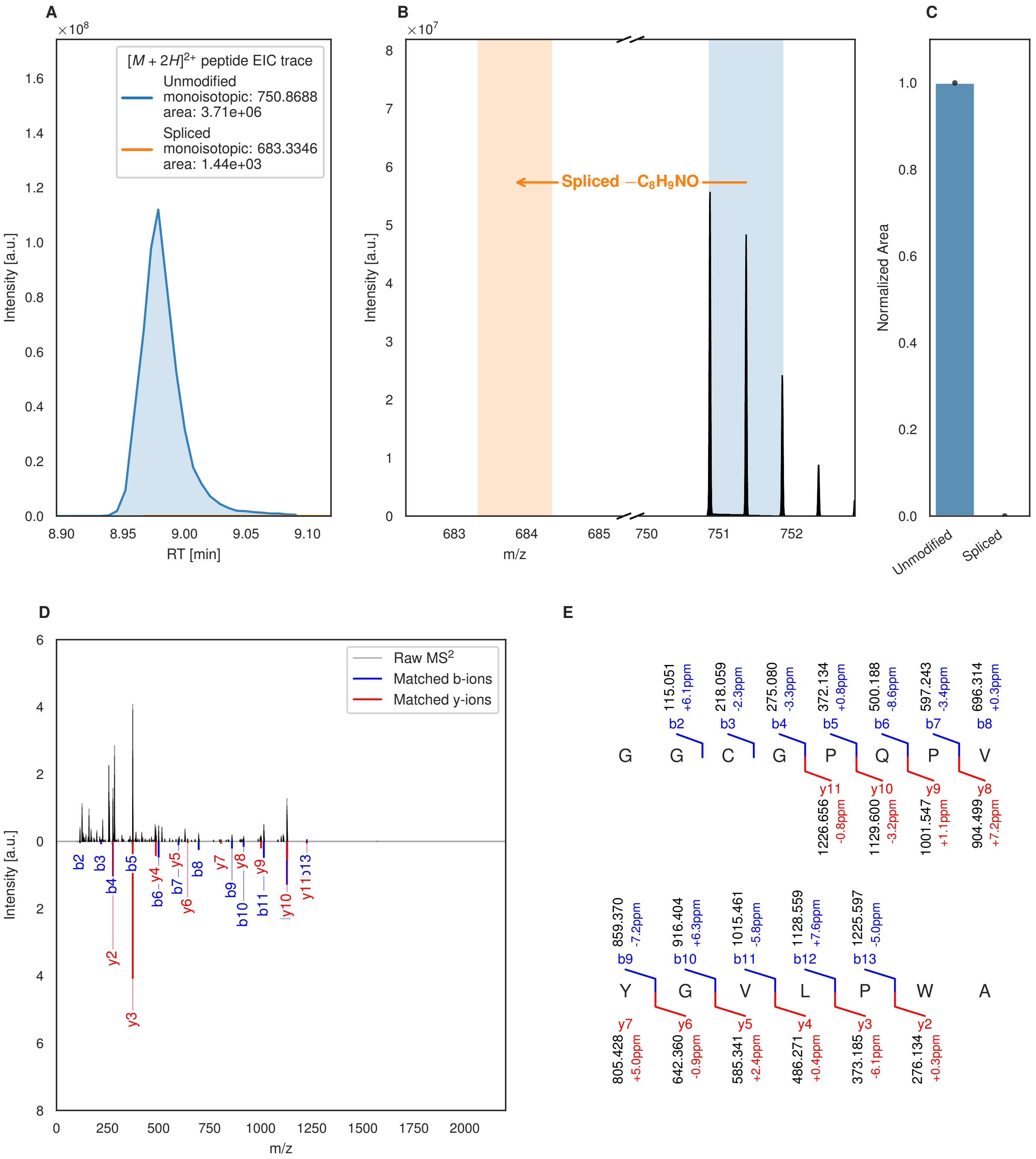

A
×10^8
[M + 2H]^2+ peptide EIC trace
Unmodified
monoisotopic: 750.8688
area: 3.71e+06
Spliced
monoisotopic: 683.3346
area: 1.44e+03
Intensity [a.u.]
RT [min]
B
×10^7
Spliced −C8H9NO
Intensity [a.u.]
m/z
C
Normalized Area
Unmodified
Spliced
D
Raw MS2
Matched b-ions
Matched y-ions
Intensity [a.u.]
m/z
E
G G C G P Q P V
b2 115.051 +6.1ppm
b3 218.059 -2.3ppm
b4 275.080 -3.3ppm
b5 372.134 +0.8ppm
b6 500.188 -8.6ppm
b7 597.243 -3.4ppm
b8 696.314 +0.3ppm
y11 1226.656 -0.8ppm
y10 1129.600 -3.2ppm
y9 1001.547 +1.1ppm
y8 904.499 +7.2ppm
Y G V L P W A
b9 859.370 -7.2ppm
b10 916.404 +6.3ppm
b11 1015.461 -5.8ppm
b12 1128.559 +7.6ppm
b13 1225.597 -5.0ppm
y7 805.428 +5.0ppm
y6 642.360 -0.9ppm
y5 585.341 +2.4ppm
y4 486.271 +0.4ppm
y3 373.185 -6.1ppm
y2 276.134 +0.3ppm

Figure S20 (figure continued from the previous page).

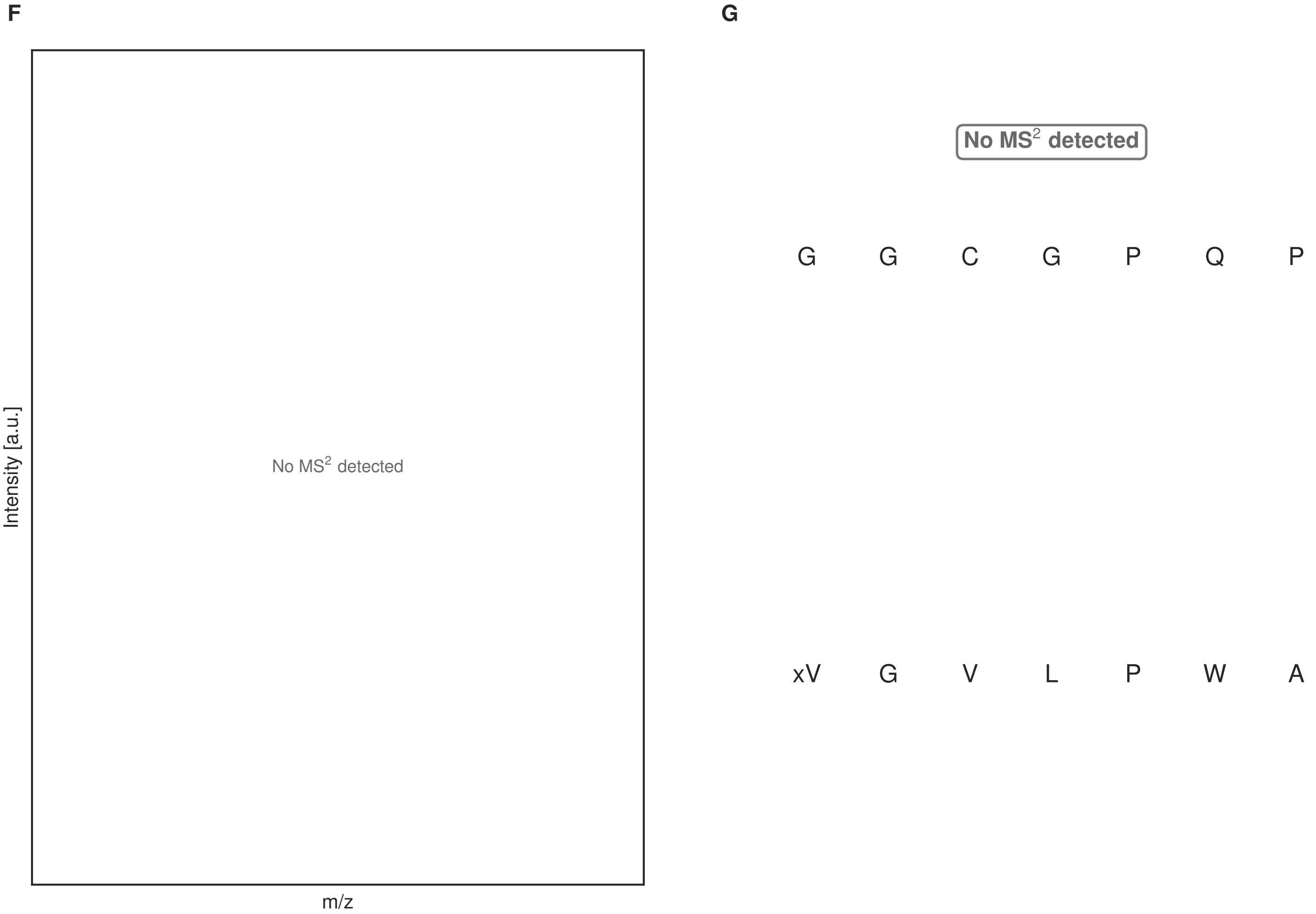

**Figure S20: Modification of the TJ-C-693 precursor peptide by HypXY.** LC-MS analysis of IMAC-purified precursor peptides from *E. coli* BL21(DE3)Tuner containing splicease-encoding plasmid TJ-C-689 (HypXY) and precursor peptide-encoding plasmid TJ-C-693 (Fh8-HypAc-TEV-Artificial-1) after expression at 24 °C for 22 h in 30 mL cultures after digestion with TEV. Data from n=1 independent biological replicates. Spliced amino acids have an x as a prefix. Spliced refers to a single tyramine excision. A representative experimental replicate is shown in panels **A, B.**

**A** Close-up plot of the representative extracted ion chromatogram (EIC) of the various peptide states; the blue trace corresponds to the Unmodified peptide; the orange trace corresponds to the Spliced peptide. **B** Close-up plot of the m/z values of the peptides detected by the mass spectrometer, with the shaded colored areas corresponding to where the expected m/z values for the peptide m/z traces should occur, with the observed spectrum plotted in black. The Unmodified, Spliced peptide(s) were found in the mass spectrum. **C** Bar plot showing the relative abundance of the various peptide states at charge z=2; the bars represent the average relative abundances normalized to total peak areas, with each individual experimental replicate value plotted as a black dot. **D** Mirrored $MS^2$ plot for the Unmodified peptide. In this mirror plot, the full fragmentation spectrum is plotted in black pointing upward, and the matched fragment b ions (blue) and y ions (red) are plotted pointing downward, mirrored against it, so that each matched fragment can be compared directly with the measured spectrum. **E** $MS^2$ spectrum of the fragments corresponding to the Unmodified peptide sequence, with matched b (blue, above) and y ions (red, below) plotted. **F** Mirrored $MS^2$ plot for the Spliced peptide. In this mirror plot, the full fragmentation spectrum is plotted in black pointing upward, and the matched fragment b ions (blue) and y ions (red) are plotted pointing downward, mirrored against it, so that each matched fragment can be compared directly with the measured spectrum. **G** $MS^2$ spectrum of the fragments corresponding to the Spliced peptide sequence, with matched b (blue, above) and y ions (red, below) plotted.

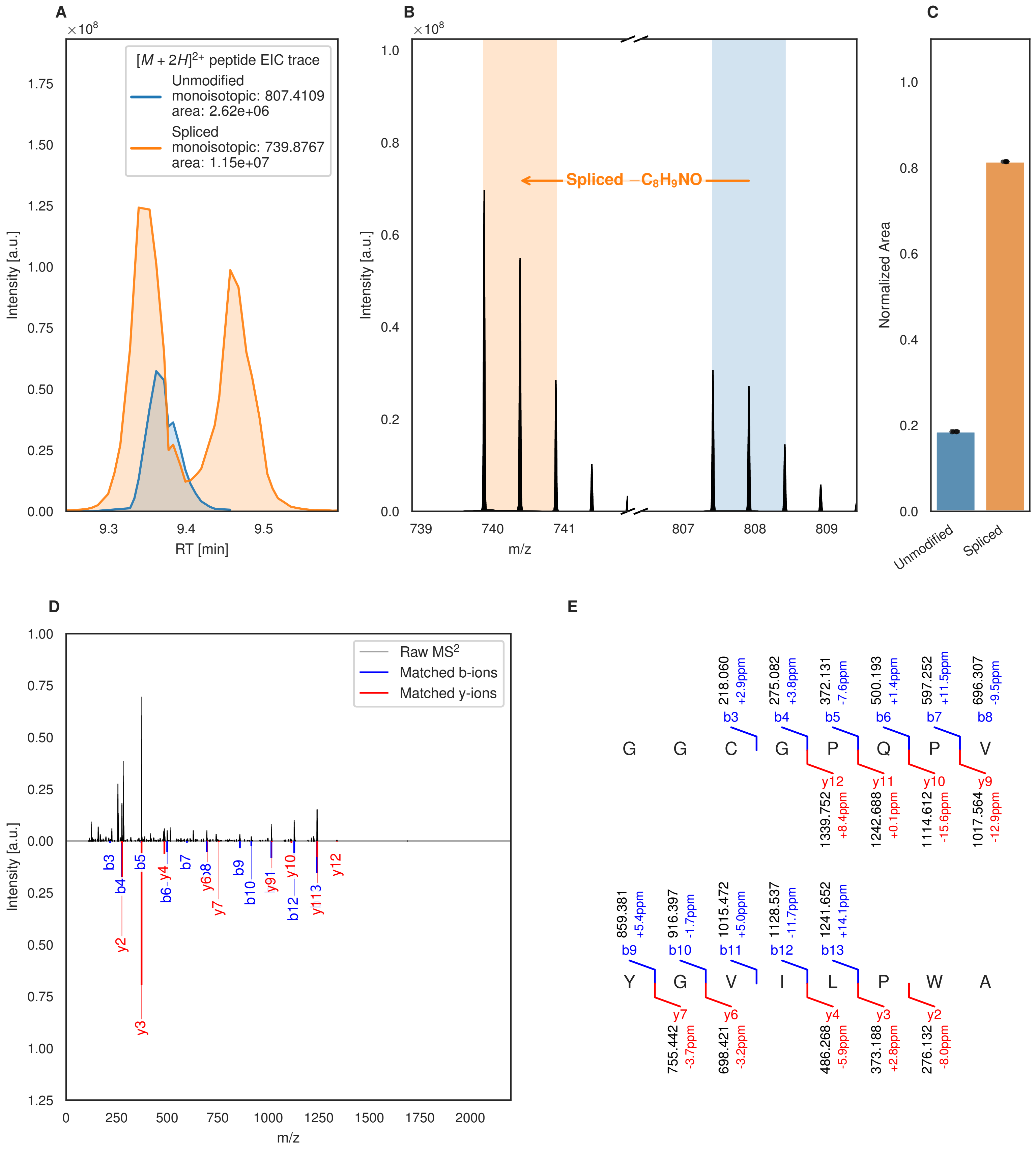
A
×10^8
[M + 2H]2+ peptide EIC trace
Unmodified
monoisotopic: 807.4109
area: 2.62e+06
Spliced
monoisotopic: 739.8767
area: 1.15e+07
Intensity [a.u.]
RT [min]
B
×10^8
Spliced −C8H9NO
Intensity [a.u.]
m/z
C
Normalized Area
Unmodified
Spliced
D
Raw MS2
Matched b-ions
Matched y-ions
Intensity [a.u.]
m/z
E
G G C G P Q P V
Y G V I L P W A
218.060 +2.9ppm b3
275.082 +3.8ppm b4
372.131 -7.6ppm b5
500.193 +1.4ppm b6
597.252 +11.5ppm b7
696.307 -9.5ppm b8
y12 1339.752 +8.4ppm
y11 1242.688 +0.1ppm
y10 1114.612 -15.6ppm
y9 1017.564 -12.9ppm
859.381 +5.4ppm b9
916.397 -1.7ppm b10
1015.472 +5.0ppm b11
1128.537 -11.7ppm b12
1241.652 +14.1ppm b13
y7 755.442 -3.7ppm
y6 698.421 -3.2ppm
y4 486.268 -5.9ppm
y3 373.188 +2.8ppm
y2 276.132 -8.0ppm

Figure S21 (figure continued from the previous page).

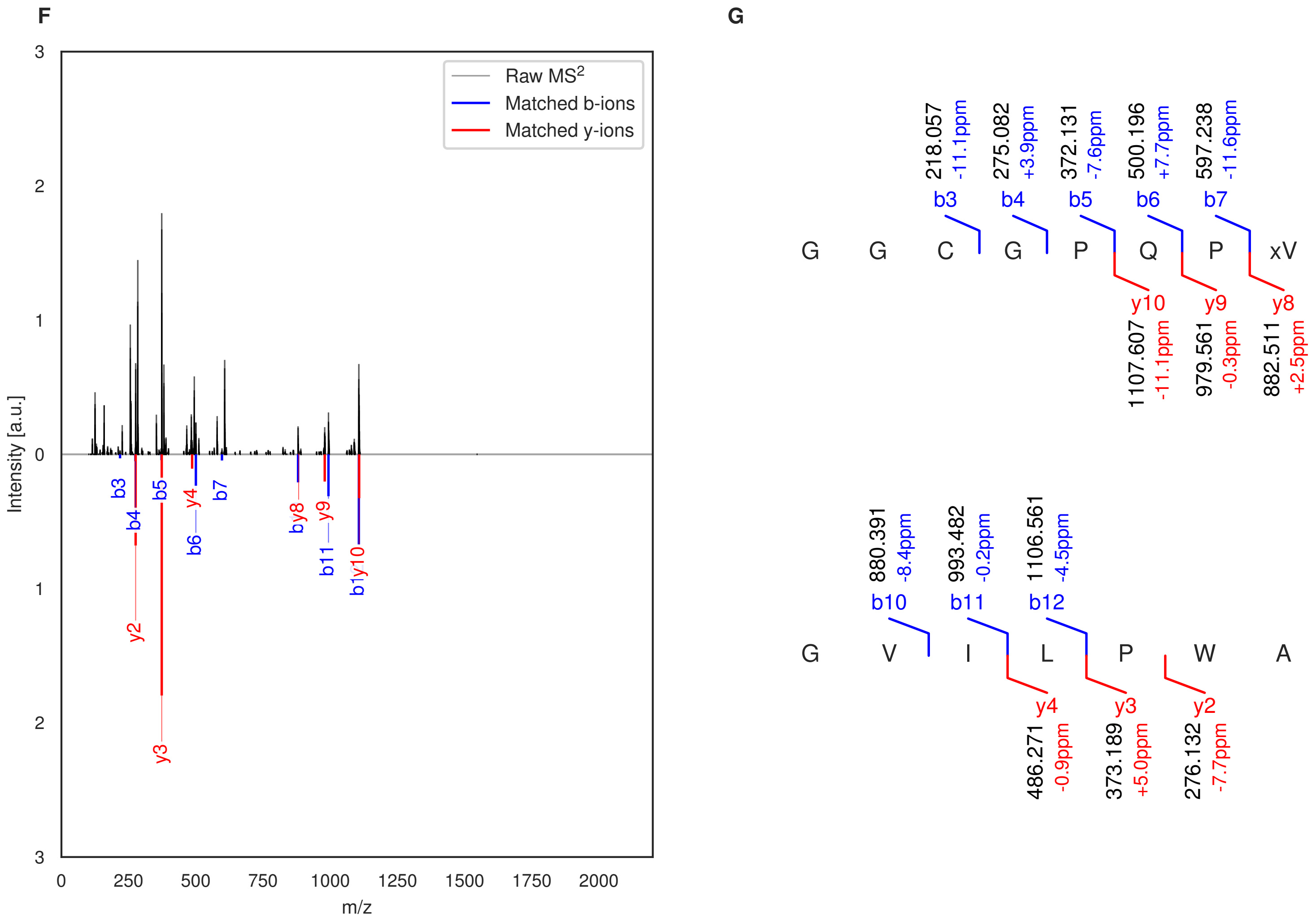

**Figure S21: Modification of the TJ-C-694 precursor peptide by HypXY.** LC-MS analysis of IMAC-purified precursor peptides from *E. coli* BL21(DE3)Tuner containing splicease-encoding plasmid TJ-C-560 (HypXY) and precursor peptide-encoding plasmid TJ-C-694 (Fh8-HypAc-TEV-Artificial-2) after expression at 24 °C for 22 h in 30 mL cultures after digestion with TEV. Data from n=2 independent biological replicates. Spliced amino acids have an x as a prefix. Spliced refers to a single tyramine excision. A representative experimental replicate is shown in panels **A, B.**

**A** Close-up plot of the representative extracted ion chromatogram (EIC) of the various peptide states; the blue trace corresponds to the Unmodified peptide; the orange trace corresponds to the Spliced peptide. **B** Close-up plot of the m/z values of the peptides detected by the mass spectrometer, with the shaded colored areas corresponding to where the expected m/z values for the peptide m/z traces should occur, with the observed spectrum plotted in black. The Unmodified, Spliced peptide(s) were found in the mass spectrum. **C** Bar plot showing the relative abundance of the various peptide states at charge z=2; the bars represent the average relative abundances normalized to total peak areas, with each individual experimental replicate value plotted as a black dot. **D** Mirrored MS$^2$ plot for the Unmodified peptide. In this mirror plot, the full fragmentation spectrum is plotted in black pointing upward, and the matched fragment b ions (blue) and y ions (red) are plotted pointing downward, mirrored against it, so that each matched fragment can be compared directly with the measured spectrum. **E** MS$^2$ spectrum of the fragments corresponding to the Unmodified peptide sequence, with matched b (blue, above) and y ions (red, below) plotted. **F** Mirrored MS$^2$ plot for the Spliced peptide. In this mirror plot, the full fragmentation spectrum is plotted in black pointing upward, and the matched fragment b ions (blue) and y ions (red) are plotted pointing downward, mirrored against it, so that each matched fragment can be compared directly with the measured spectrum. **G** MS$^2$ spectrum of the fragments corresponding to the Spliced peptide sequence, with matched b (blue, above) and y ions (red, below) plotted.

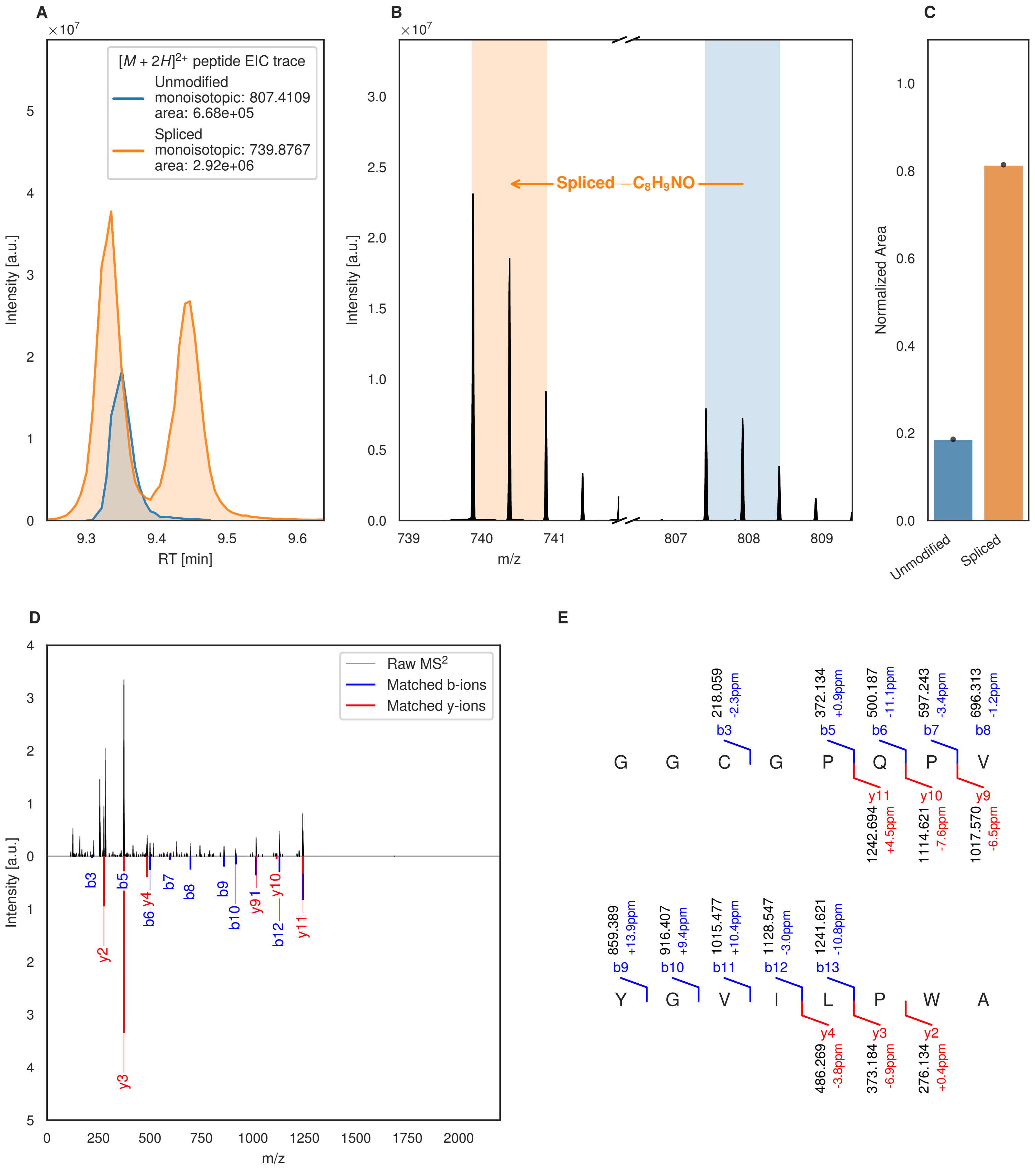
A
×10^7
[M + 2H]2+ peptide EIC trace
Unmodified
monoisotopic: 807.4109
area: 6.68e+05
Spliced
monoisotopic: 739.8767
area: 2.92e+06
Intensity [a.u.]
RT [min]
B
×10^7
Spliced −C8H9NO
Intensity [a.u.]
m/z
C
Normalized Area
Unmodified
Spliced
D
Raw MS2
Matched b-ions
Matched y-ions
Intensity [a.u.]
m/z
b3
b5
b6
y4
b7
b8
b9
b10
y91
y10
b12
y11
y2
y3
E
218.059
-2.3ppm
372.134
+0.9ppm
500.187
-11.1ppm
597.243
-3.4ppm
696.313
-1.2ppm
b3
b5
b6
b7
b8
G G C G P Q P V
y11
y10
y9
1242.694
+4.5ppm
1114.621
-7.6ppm
1017.570
-6.5ppm
859.389
+13.9ppm
916.407
+9.4ppm
1015.477
+10.4ppm
1128.547
-3.0ppm
1241.621
-10.8ppm
b9
b10
b11
b12
b13
Y G V I L P W A
y4
y3
y2
486.269
-3.8ppm
373.184
-6.9ppm
276.134
+0.4ppm

Figure S22 (figure continued from the previous page).

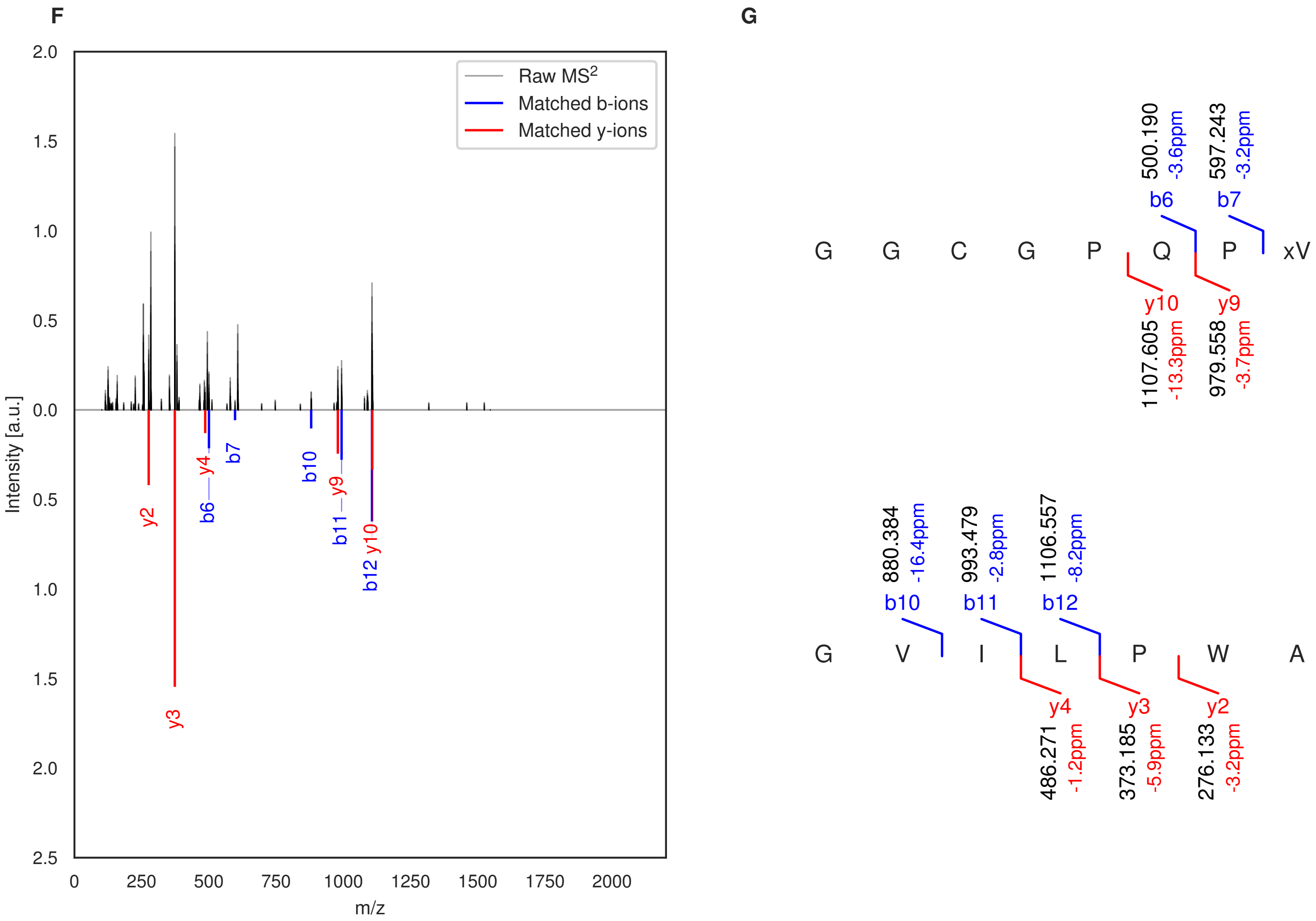

**Figure S22: Modification of the TJ-C-694 precursor peptide by HypXY.** LC-MS analysis of IMAC-purified precursor peptides from *E. coli* BL21(DE3)Tuner containing splicease-encoding plasmid TJ-C-689 (HypXY) and precursor peptide-encoding plasmid TJ-C-694 (Fh8-HypAc-TEV-Artificial-2) after expression at 24 °C for 22 h in 30 mL cultures after digestion with TEV. Data from n=1 independent biological replicates. Spliced amino acids have an x as a prefix. Spliced refers to a single tyramine excision. A representative experimental replicate is shown in panels **A, B.**

**A** Close-up plot of the representative extracted ion chromatogram (EIC) of the various peptide states; the blue trace corresponds to the Unmodified peptide; the orange trace corresponds to the Spliced peptide. **B** Close-up plot of the m/z values of the peptides detected by the mass spectrometer, with the shaded colored areas corresponding to where the expected m/z values for the peptide m/z traces should occur, with the observed spectrum plotted in black. The Unmodified, Spliced peptide(s) were found in the mass spectrum. **C** Bar plot showing the relative abundance of the various peptide states at charge z=2; the bars represent the average relative abundances normalized to total peak areas, with each individual experimental replicate value plotted as a black dot. **D** Mirrored $MS^2$ plot for the Unmodified peptide. In this mirror plot, the full fragmentation spectrum is plotted in black pointing upward, and the matched fragment b ions (blue) and y ions (red) are plotted pointing downward, mirrored against it, so that each matched fragment can be compared directly with the measured spectrum. **E** $MS^2$ spectrum of the fragments corresponding to the Unmodified peptide sequence, with matched b (blue, above) and y ions (red, below) plotted. **F** Mirrored $MS^2$ plot for the Spliced peptide. In this mirror plot, the full fragmentation spectrum is plotted in black pointing upward, and the matched fragment b ions (blue) and y ions (red) are plotted pointing downward, mirrored against it, so that each matched fragment can be compared directly with the measured spectrum. **G** $MS^2$ spectrum of the fragments corresponding to the Spliced peptide sequence, with matched b (blue, above) and y ions (red, below) plotted.

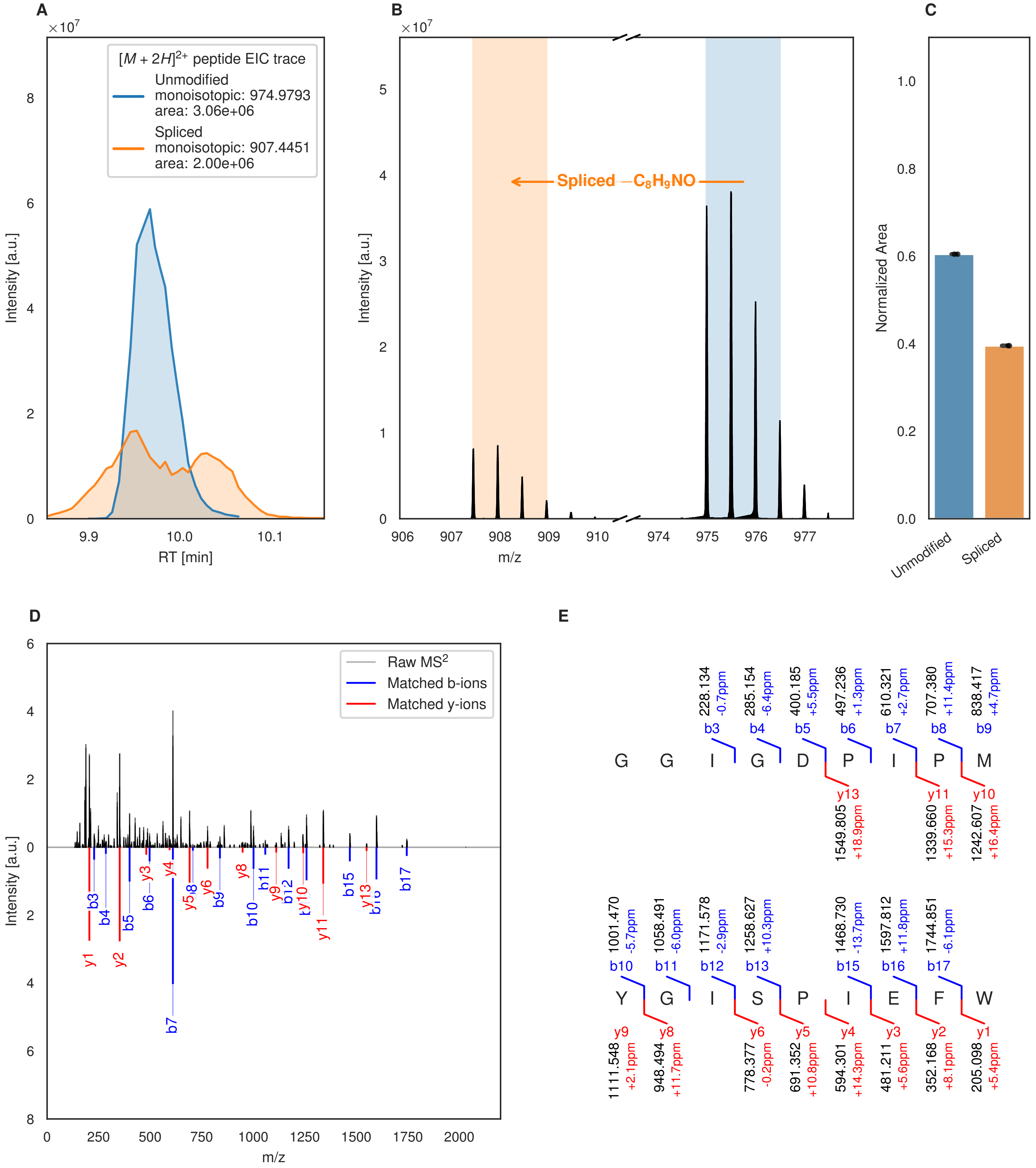

A
×10^7
[M + 2H]^2+ peptide EIC trace
Unmodified
monoisotopic: 974.9793
area: 3.06e+06
Spliced
monoisotopic: 907.4451
area: 2.00e+06
Intensity [a.u.]
RT [min]
B
×10^7
Spliced −C8H9NO
Intensity [a.u.]
m/z
C
Normalized Area
Unmodified
Spliced
D
Raw MS2
Matched b-ions
Matched y-ions
Intensity [a.u.]
m/z
E
G G I G D P I P M
228.134 -0.7ppm b3
285.154 -6.4ppm b4
400.185 +5.5ppm b5
497.236 +1.3ppm b6
610.321 +2.7ppm b7
707.380 +11.4ppm b8
838.417 +4.7ppm b9
y13 1549.805 +18.9ppm
y11 1339.660 +15.3ppm
y10 1242.607 +16.4ppm
Y G I S P I E F W
1001.470 -5.7ppm b10
1058.491 -6.0ppm b11
1171.578 -2.9ppm b12
1258.627 +10.3ppm b13
1468.730 -13.7ppm b15
1597.812 +11.8ppm b16
1744.851 -6.1ppm b17
y9 1111.548 +2.1ppm
y8 948.494 +11.7ppm
y6 778.377 -0.2ppm
y5 691.352 +10.8ppm
y4 594.301 +14.3ppm
y3 481.211 +5.6ppm
y2 352.168 +8.1ppm
y1 205.098 +5.4ppm

Figure S23 (figure continued from the previous page).

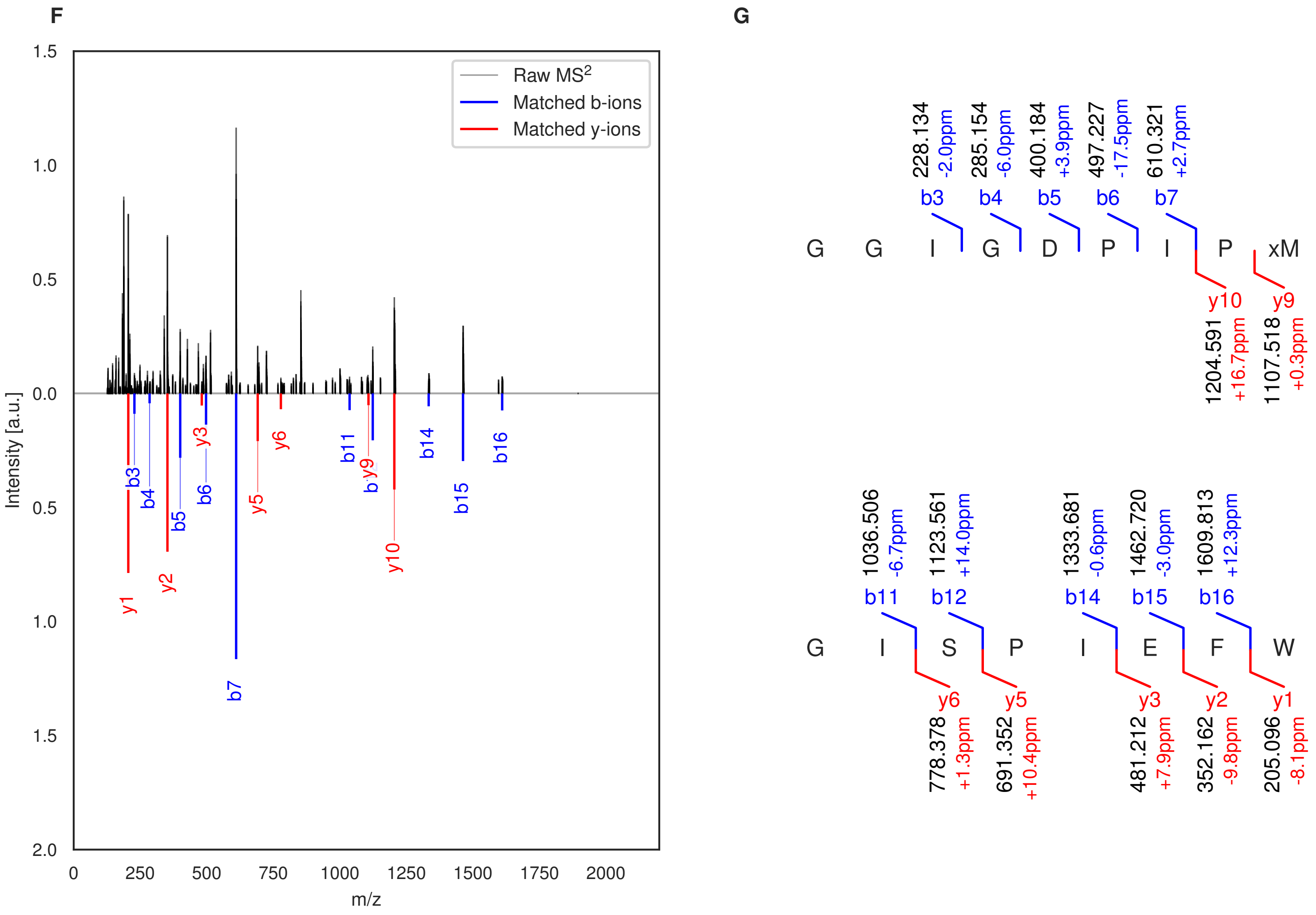

**Figure S23: Modification of the TJ-C-695 precursor peptide by HypXY.** LC-MS analysis of IMAC-purified precursor peptides from *E. coli* BL21(DE3)Tuner containing splicease-encoding plasmid TJ-C-560 (HypXY) and precursor peptide-encoding plasmid TJ-C-695 (Fh8-HypAc-TEV-Artificial-3) after expression at 24 °C for 22 h in 30 mL cultures after digestion with TEV. Data from n=2 independent biological replicates. Spliced amino acids have an x as a prefix. Spliced refers to a single tyramine excision. A representative experimental replicate is shown in panels **A, B.**

**A** Close-up plot of the representative extracted ion chromatogram (EIC) of the various peptide states; the blue trace corresponds to the Unmodified peptide; the orange trace corresponds to the Spliced peptide. **B** Close-up plot of the m/z values of the peptides detected by the mass spectrometer, with the shaded colored areas corresponding to where the expected m/z values for the peptide m/z traces should occur, with the observed spectrum plotted in black. The Unmodified, Spliced peptide(s) were found in the mass spectrum. **C** Bar plot showing the relative abundance of the various peptide states at charge z=2; the bars represent the average relative abundances normalized to total peak areas, with each individual experimental replicate value plotted as a black dot. **D** Mirrored MS$^2$ plot for the Unmodified peptide. In this mirror plot, the full fragmentation spectrum is plotted in black pointing upward, and the matched fragment b ions (blue) and y ions (red) are plotted pointing downward, mirrored against it, so that each matched fragment can be compared directly with the measured spectrum. **E** MS$^2$ spectrum of the fragments corresponding to the Unmodified peptide sequence, with matched b (blue, above) and y ions (red, below) plotted. **F** Mirrored MS$^2$ plot for the Spliced peptide. In this mirror plot, the full fragmentation spectrum is plotted in black pointing upward, and the matched fragment b ions (blue) and y ions (red) are plotted pointing downward, mirrored against it, so that each matched fragment can be compared directly with the measured spectrum. **G** MS$^2$ spectrum of the fragments corresponding to the Spliced peptide sequence, with matched b (blue, above) and y ions (red, below) plotted.

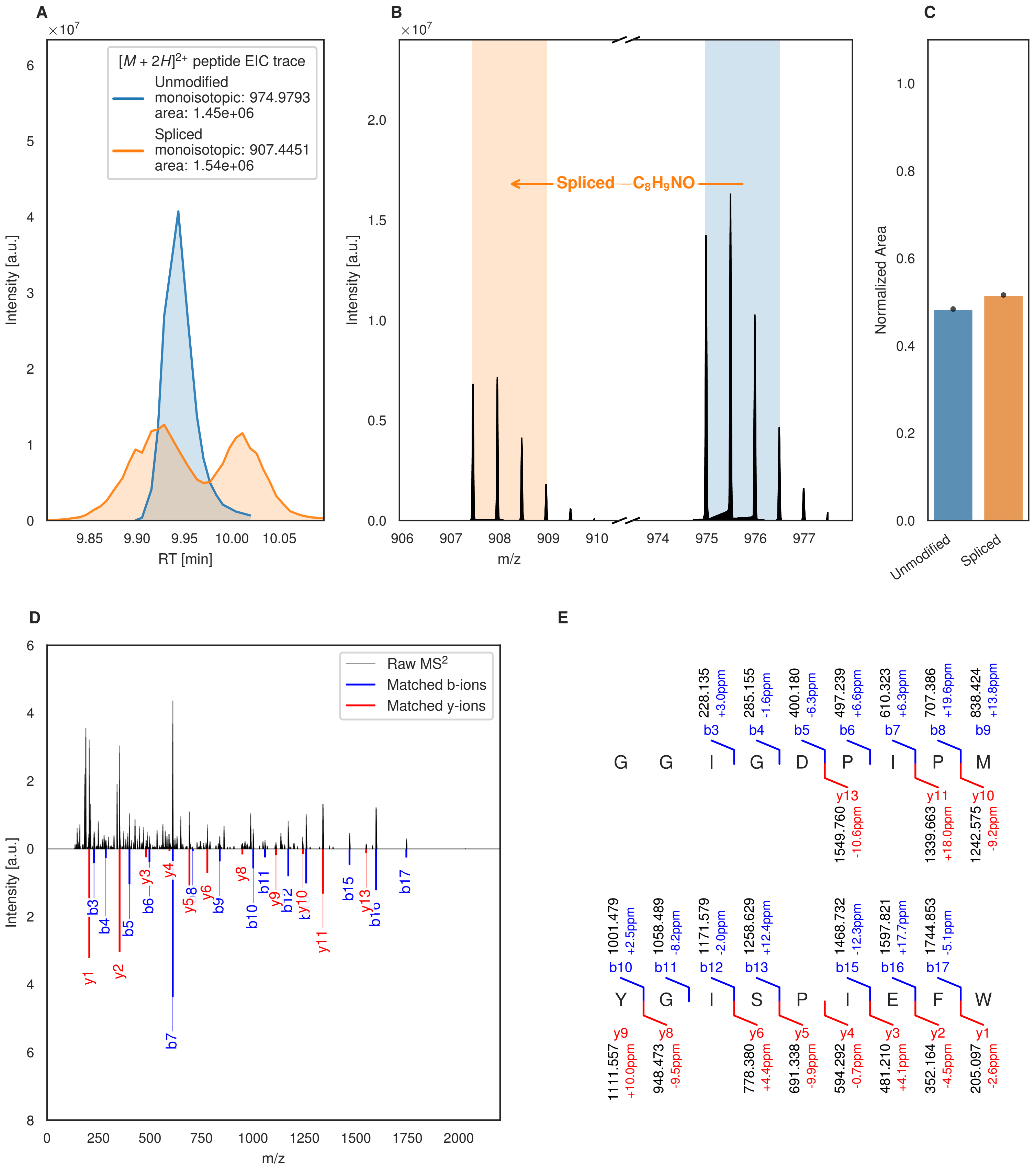
A
[M + 2H]2+ peptide EIC trace
Unmodified
monoisotopic: 974.9793
area: 1.45e+06
Spliced
monoisotopic: 907.4451
area: 1.54e+06
Intensity [a.u.]
RT [min]
B
Spliced −C8H9NO
Intensity [a.u.]
m/z
C
Normalized Area
Unmodified
Spliced
D
Raw MS2
Matched b-ions
Matched y-ions
Intensity [a.u.]
m/z
E
G G I G D P I P M
Y G I S P I E F W

Figure S24 (figure continued from the previous page).

F

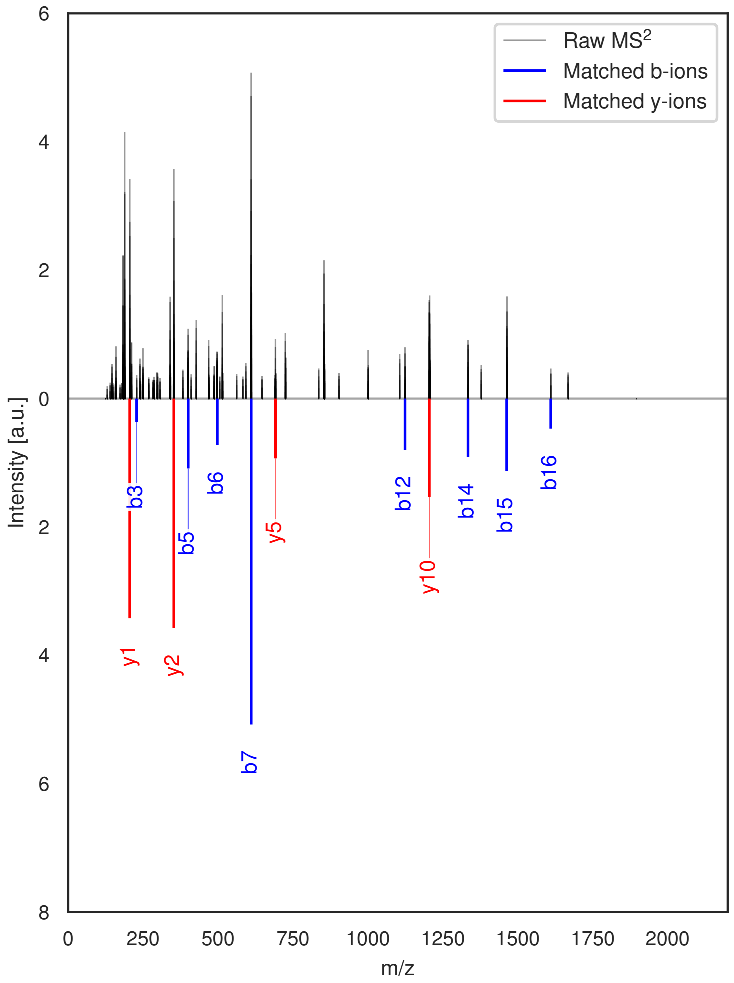

G

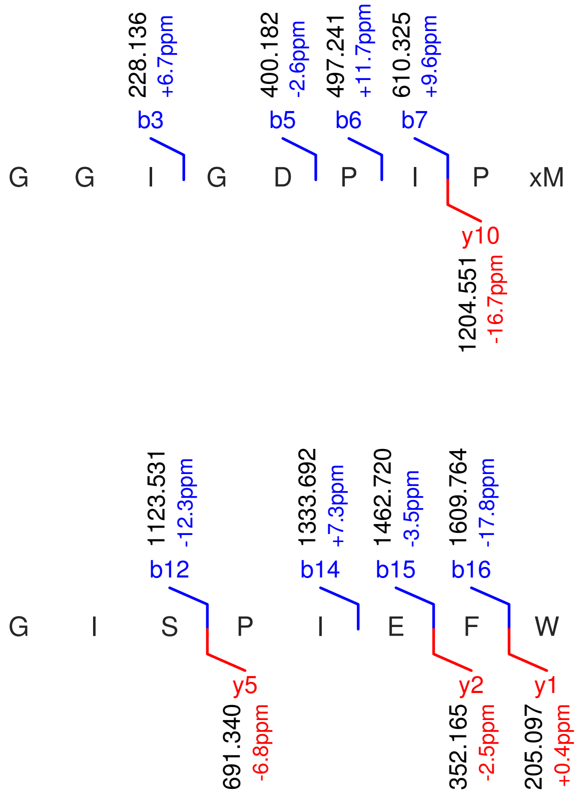

**Figure S24: Modification of the TJ-C-695 precursor peptide by HypXY.** LC-MS analysis of IMAC-purified precursor peptides from *E. coli* BL21(DE3)Tuner containing splicease-encoding plasmid TJ-C-689 (HypXY) and precursor peptide-encoding plasmid TJ-C-695 (Fh8-HypAc-TEV-Artificial-3) after expression at 24 °C for 22 h in 30 mL cultures after digestion with TEV. Data from n=1 independent biological replicates. Spliced amino acids have an x as a prefix. Spliced refers to a single tyramine excision. A representative experimental replicate is shown in panels **A, B.**

**A** Close-up plot of the representative extracted ion chromatogram (EIC) of the various peptide states; the blue trace corresponds to the Unmodified peptide; the orange trace corresponds to the Spliced peptide. **B** Close-up plot of the m/z values of the peptides detected by the mass spectrometer, with the shaded colored areas corresponding to where the expected m/z values for the peptide m/z traces should occur, with the observed spectrum plotted in black. The Unmodified, Spliced peptide(s) were found in the mass spectrum. **C** Bar plot showing the relative abundance of the various peptide states at charge z=2; the bars represent the average relative abundances normalized to total peak areas, with each individual experimental replicate value plotted as a black dot. **D** Mirrored MS$^2$ plot for the Unmodified peptide. In this mirror plot, the full fragmentation spectrum is plotted in black pointing upward, and the matched fragment b ions (blue) and y ions (red) are plotted pointing downward, mirrored against it, so that each matched fragment can be compared directly with the measured spectrum. **E** MS$^2$ spectrum of the fragments corresponding to the Unmodified peptide sequence, with matched b (blue, above) and y ions (red, below) plotted. **F** Mirrored MS$^2$ plot for the Spliced peptide. In this mirror plot, the full fragmentation spectrum is plotted in black pointing upward, and the matched fragment b ions (blue) and y ions (red) are plotted pointing downward, mirrored against it, so that each matched fragment can be compared directly with the measured spectrum. **G** MS$^2$ spectrum of the fragments corresponding to the Spliced peptide sequence, with matched b (blue, above) and y ions (red, below) plotted.

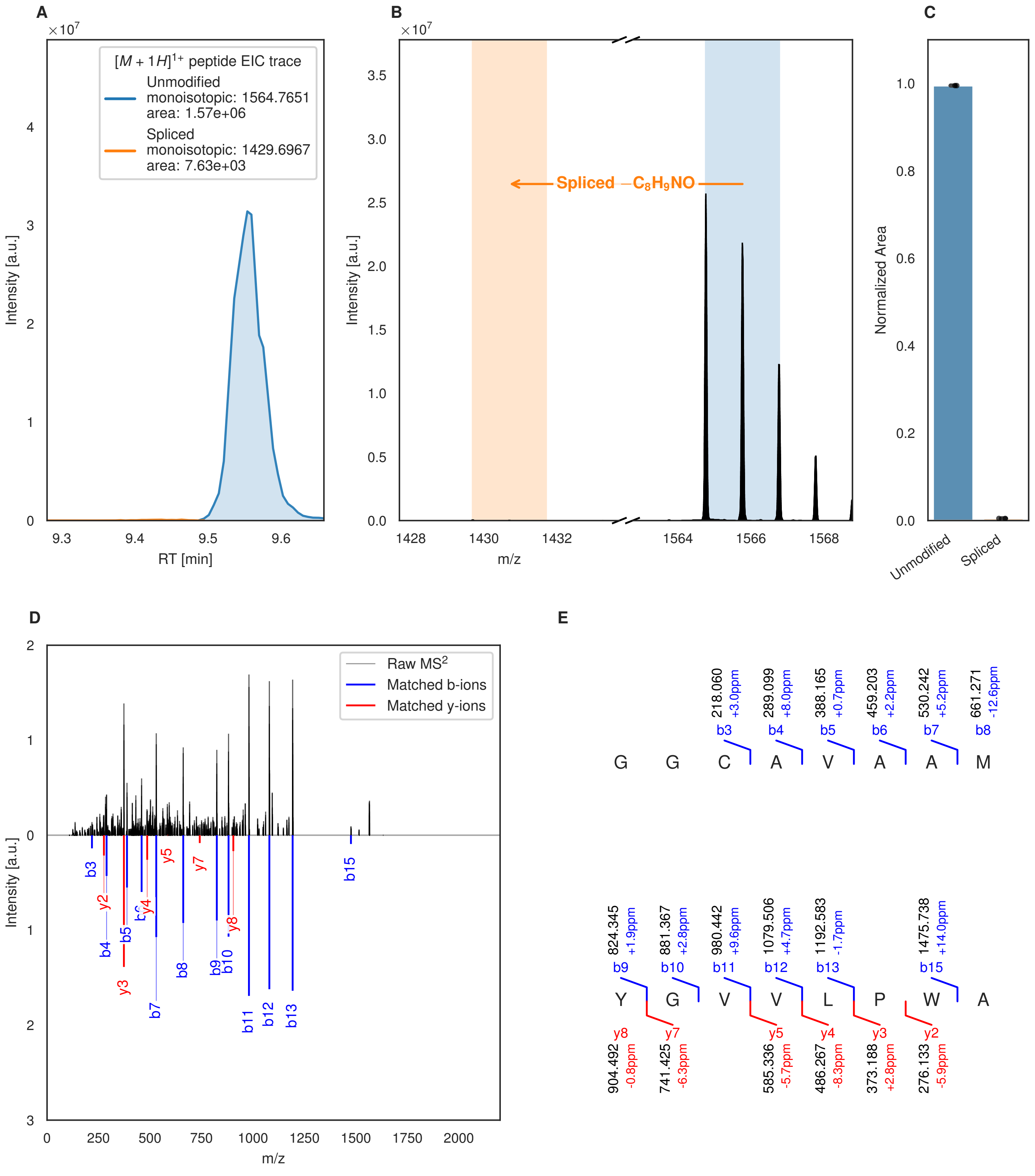
A
B
C
D
E
[M + 1H]1+ peptide EIC trace
Unmodified
monoisotopic: 1564.7651
area: 1.57e+06
Spliced
monoisotopic: 1429.6967
area: 7.63e+03
Intensity [a.u.]
RT [min]
Spliced −C8H9NO
m/z
Normalized Area
Unmodified
Spliced
Raw MS2
Matched b-ions
Matched y-ions
G G C A V A A M
Y G V V L P W A
218.060 +3.0ppm b3
289.099 +8.0ppm b4
388.165 +0.7ppm b5
459.203 +2.2ppm b6
530.242 +5.2ppm b7
661.271 -12.6ppm b8
824.345 +1.9ppm b9
881.367 +2.8ppm b10
980.442 +9.6ppm b11
1079.506 +4.7ppm b12
1192.583 -1.7ppm b13
1475.738 +14.0ppm b15
y8 904.492 -0.8ppm
y7 741.425 -6.3ppm
y5 585.336 -5.7ppm
y4 486.267 -8.3ppm
y3 373.188 +2.8ppm
y2 276.133 -5.9ppm

Figure S25 (figure continued from the previous page).

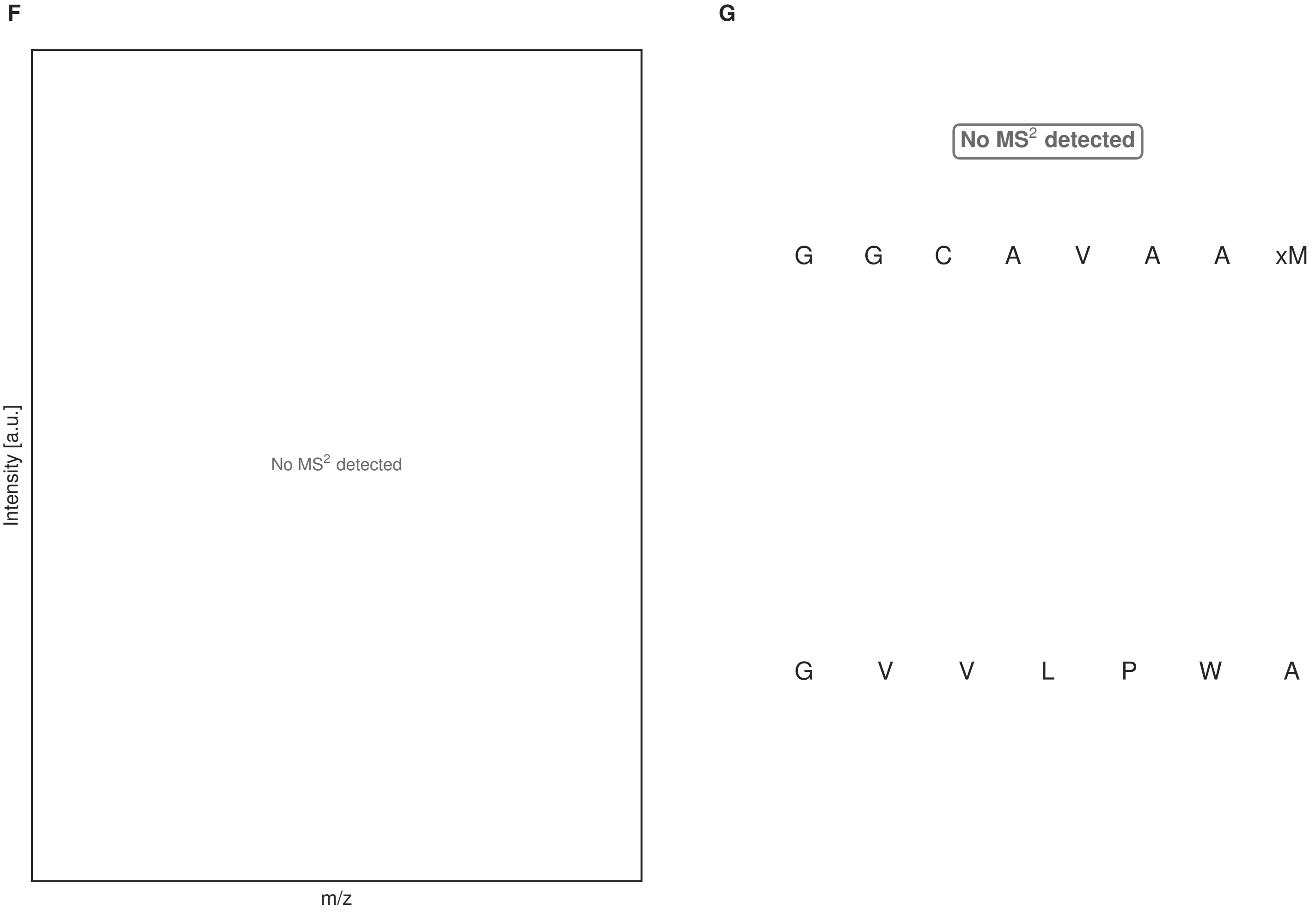

**Figure S25: Modification of the TJ-C-696 precursor peptide by HypXY.** LC-MS analysis of IMAC-purified precursor peptides from *E. coli* BL21(DE3)Tuner containing splicease-encoding plasmid TJ-C-560 (HypXY) and precursor peptide-encoding plasmid TJ-C-696 (Fh8-HypAc-TEV-Artificial-4) after expression at 24 °C for 22 h in 30 mL cultures after digestion with TEV. Data from n=2 independent biological replicates. Spliced amino acids have an x as a prefix. Spliced refers to a single tyramine excision. A representative experimental replicate is shown in panels **A, B.**

**A** Close-up plot of the representative extracted ion chromatogram (EIC) of the various peptide states; the blue trace corresponds to the Unmodified peptide; the orange trace corresponds to the Spliced peptide. **B** Close-up plot of the m/z values of the peptides detected by the mass spectrometer, with the shaded colored areas corresponding to where the expected m/z values for the peptide m/z traces should occur, with the observed spectrum plotted in black. The Unmodified, Spliced peptide(s) were found in the mass spectrum. **C** Bar plot showing the relative abundance of the various peptide states at charge z=1; the bars represent the average relative abundances normalized to total peak areas, with each individual experimental replicate value plotted as a black dot. **D** Mirrored MS$^2$ plot for the Unmodified peptide. In this mirror plot, the full fragmentation spectrum is plotted in black pointing upward, and the matched fragment b ions (blue) and y ions (red) are plotted pointing downward, mirrored against it, so that each matched fragment can be compared directly with the measured spectrum. **E** MS$^2$ spectrum of the fragments corresponding to the Unmodified peptide sequence, with matched b (blue, above) and y ions (red, below) plotted. **F** Mirrored MS$^2$ plot for the Spliced peptide. In this mirror plot, the full fragmentation spectrum is plotted in black pointing upward, and the matched fragment b ions (blue) and y ions (red) are plotted pointing downward, mirrored against it, so that each matched fragment can be compared directly with the measured spectrum. **G** MS$^2$ spectrum of the fragments corresponding to the Spliced peptide sequence, with matched b (blue, above) and y ions (red, below) plotted.

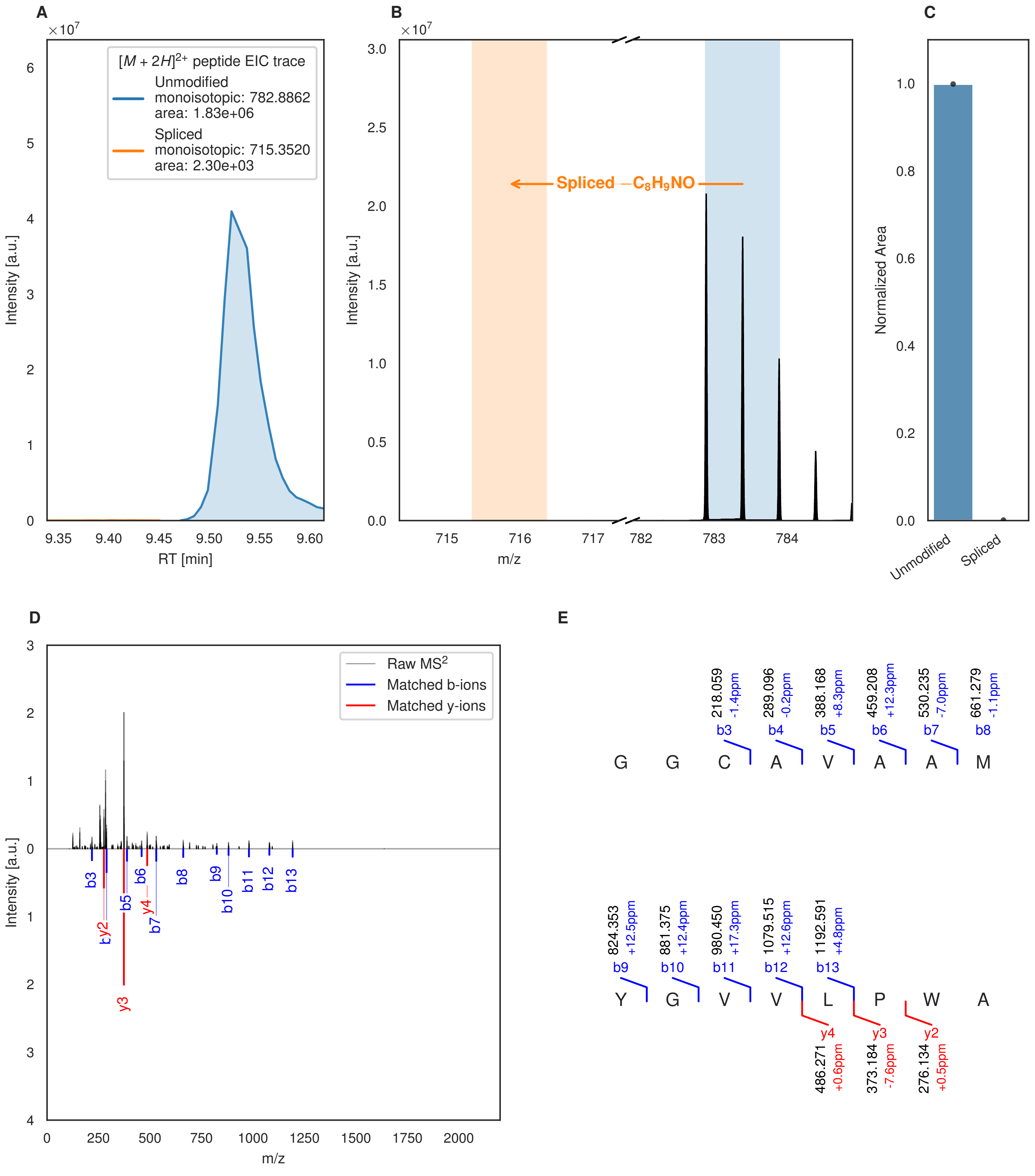

A
[M + 2H]2+ peptide EIC trace
Unmodified
monoisotopic: 782.8862
area: 1.83e+06
Spliced
monoisotopic: 715.3520
area: 2.30e+03
Intensity [a.u.]
RT [min]
B
Spliced −C8H9NO
m/z
C
Normalized Area
Unmodified
Spliced
D
Raw MS2
Matched b-ions
Matched y-ions
E
218.059 -1.4ppm b3
289.096 -0.2ppm b4
388.168 +8.3ppm b5
459.208 +12.3ppm b6
530.235 -7.0ppm b7
661.279 -1.1ppm b8
G G C A V A A M
824.353 +12.5ppm b9
881.375 +12.4ppm b10
980.450 +17.3ppm b11
1079.515 +12.6ppm b12
1192.591 +4.8ppm b13
Y G V V L P W A
y4 486.271 +0.6ppm
y3 373.184 -7.6ppm
y2 276.134 +0.5ppm

Figure S26 (figure continued from the previous page).

**F**

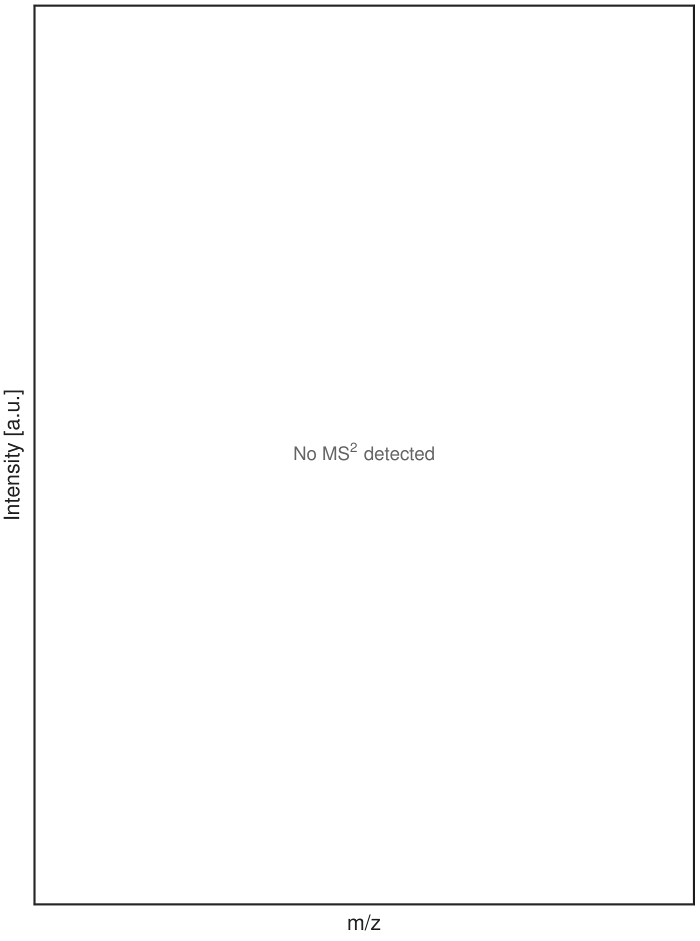

**G**

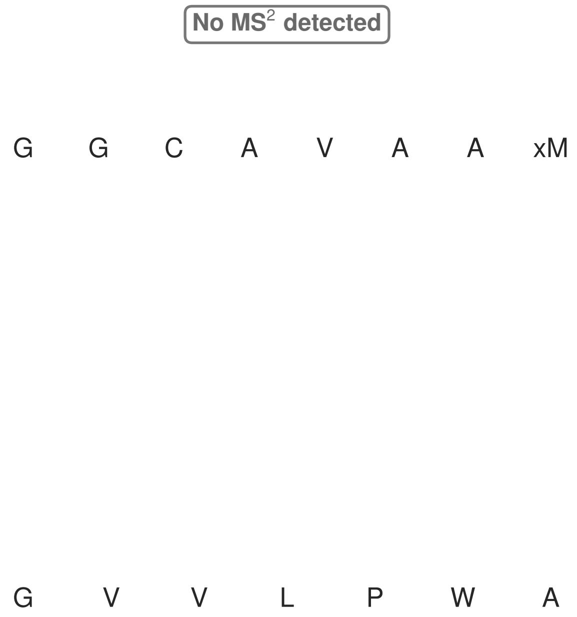

**Figure S26: Modification of the TJ-C-696 precursor peptide by HypXY.** LC-MS analysis of IMAC-purified precursor peptides from *E. coli* BL21(DE3)Tuner containing splicease-encoding plasmid TJ-C-689 (HypXY) and precursor peptide-encoding plasmid TJ-C-696 (Fh8-HypAc-TEV-Artificial-4) after expression at 24 °C for 22 h in 30 mL cultures after digestion with TEV. Data from n=1 independent biological replicates. Spliced amino acids have an x as a prefix. Spliced refers to a single tyramine excision. A representative experimental replicate is shown in panels **A, B.**

**A** Close-up plot of the representative extracted ion chromatogram (EIC) of the various peptide states; the blue trace corresponds to the Unmodified peptide; the orange trace corresponds to the Spliced peptide. **B** Close-up plot of the m/z values of the peptides detected by the mass spectrometer, with the shaded colored areas corresponding to where the expected m/z values for the peptide m/z traces should occur, with the observed spectrum plotted in black. The Unmodified, Spliced peptide(s) were found in the mass spectrum. **C** Bar plot showing the relative abundance of the various peptide states at charge z=2; the bars represent the average relative abundances normalized to total peak areas, with each individual experimental replicate value plotted as a black dot. **D** Mirrored $MS^2$ plot for the Unmodified peptide. In this mirror plot, the full fragmentation spectrum is plotted in black pointing upward, and the matched fragment b ions (blue) and y ions (red) are plotted pointing downward, mirrored against it, so that each matched fragment can be compared directly with the measured spectrum. **E** $MS^2$ spectrum of the fragments corresponding to the Unmodified peptide sequence, with matched b (blue, above) and y ions (red, below) plotted. **F** Mirrored $MS^2$ plot for the Spliced peptide. In this mirror plot, the full fragmentation spectrum is plotted in black pointing upward, and the matched fragment b ions (blue) and y ions (red) are plotted pointing downward, mirrored against it, so that each matched fragment can be compared directly with the measured spectrum. **G** $MS^2$ spectrum of the fragments corresponding to the Spliced peptide sequence, with matched b (blue, above) and y ions (red, below) plotted.

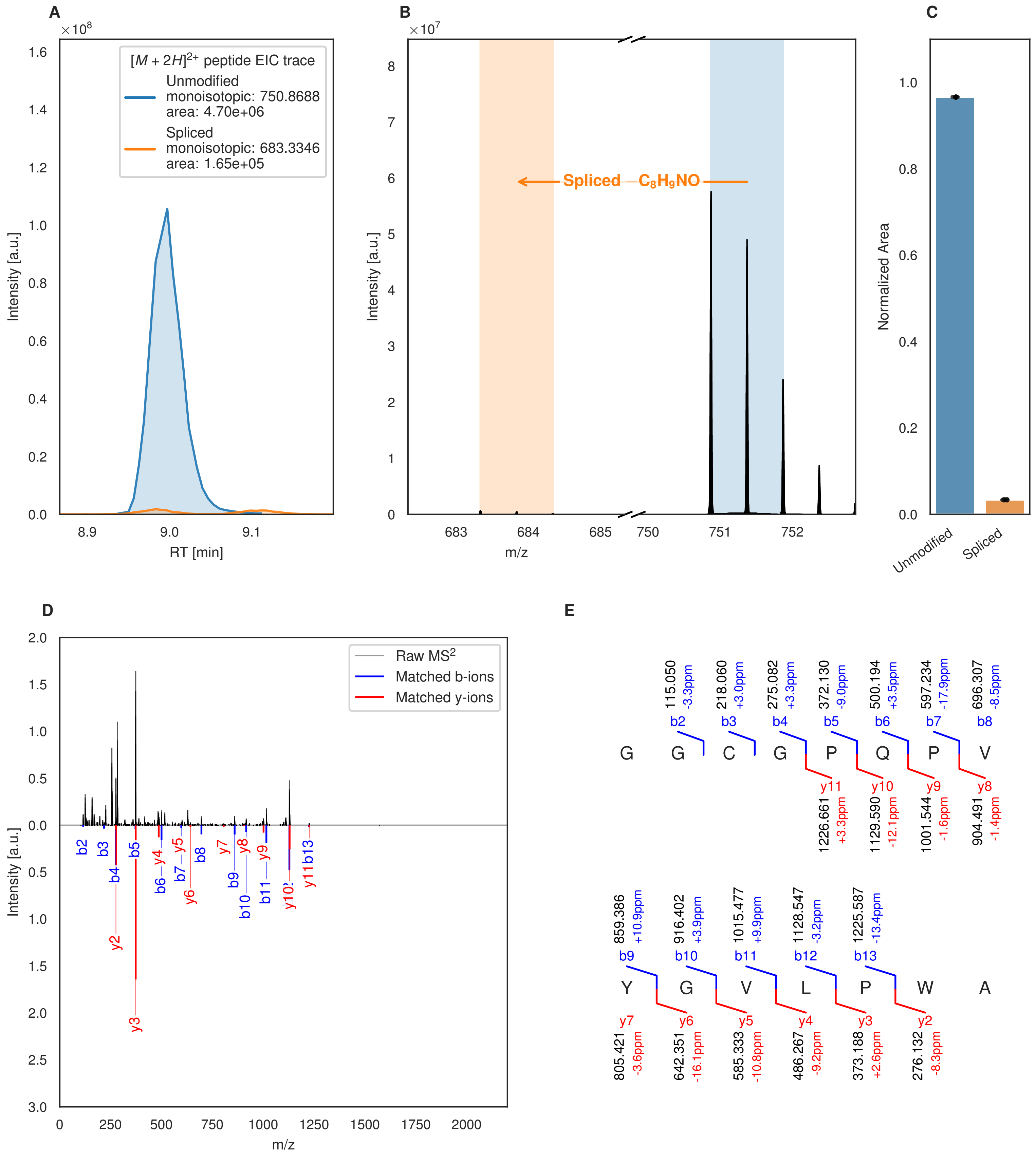
A
B
C
D
E
$[M+2H]^{2+}$ peptide EIC trace
Unmodified
monoisotopic: 750.8688
area: 4.70e+06
Spliced
monoisotopic: 683.3346
area: 1.65e+05
Intensity [a.u.]
RT [min]
Spliced $-C_8H_9NO$
m/z
Normalized Area
Unmodified
Spliced
Raw $MS^2$
Matched b-ions
Matched y-ions
G G C G P Q P V
Y G V L P W A
115.050 -3.3ppm
218.060 +3.0ppm
275.082 +3.3ppm
372.130 -9.0ppm
500.194 +3.5ppm
597.234 -17.9ppm
696.307 -8.5ppm
1226.661 +3.3ppm
1129.590 -12.1ppm
1001.544 -1.6ppm
904.491 -1.4ppm
859.386 +10.9ppm
916.402 +3.9ppm
1015.477 +9.9ppm
1128.547 -3.2ppm
1225.587 -13.4ppm
805.421 -3.6ppm
642.351 -16.1ppm
585.333 -10.8ppm
486.267 -9.2ppm
373.188 +2.6ppm
276.132 -8.3ppm

Figure S27 (figure continued from the previous page).

**F**

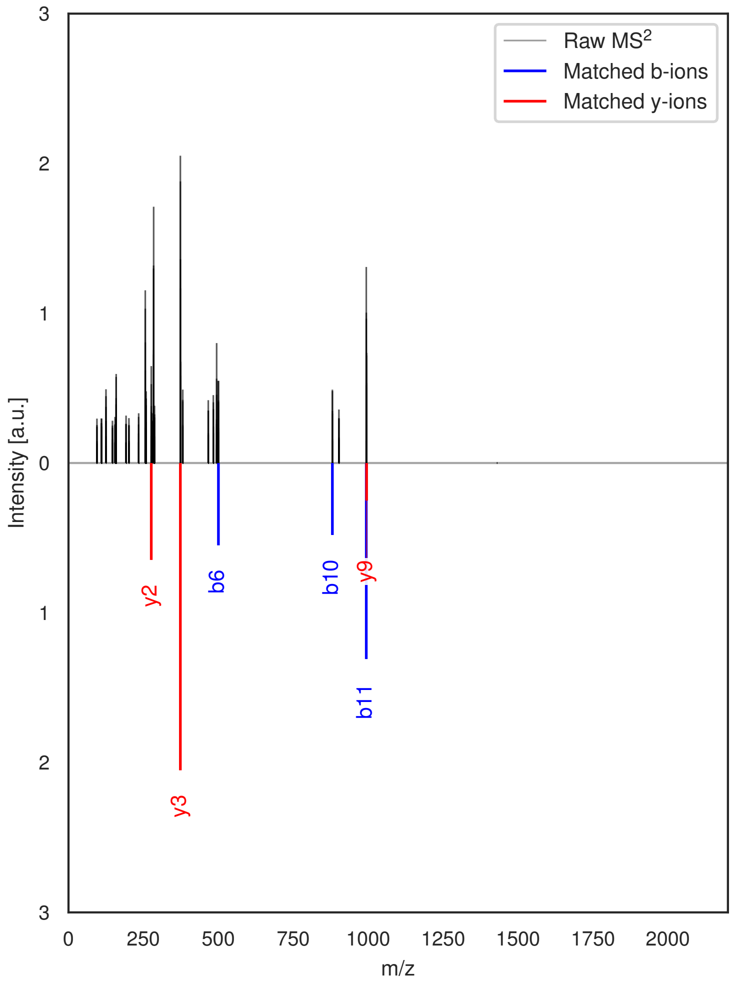

**G**

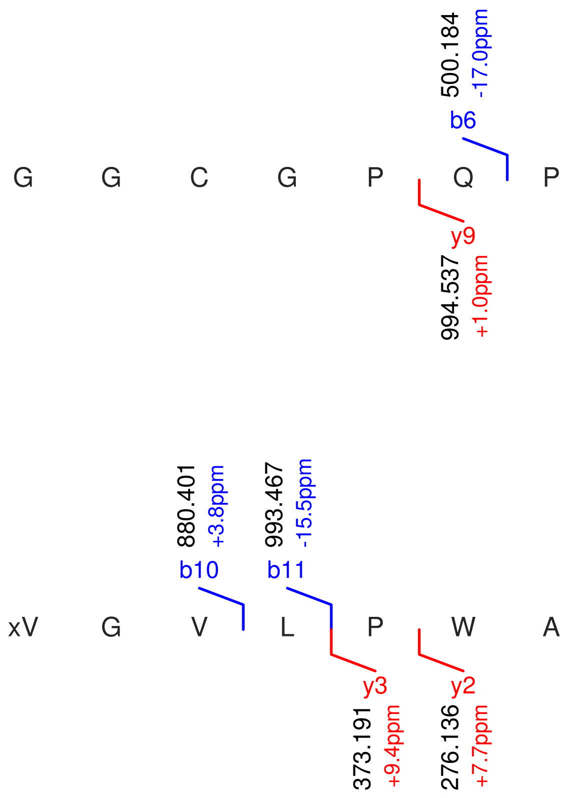

**Figure S27: Modification of the TJ-C-697 precursor peptide by PlpXY.** LC-MS analysis of IMAC-purified precursor peptides from *E. coli* BL21(DE3)Tuner containing splicease-encoding plasmid TJ-C-559 (PlpXY) and precursor peptide-encoding plasmid TJ-C-697 (Fh8-PlpA3-TEV-Artificial-1) after expression at 24 °C for 22 h in 30 mL cultures after digestion with TEV. Data from n=2 independent biological replicates. Spliced amino acids have an x as a prefix. Spliced refers to a single tyramine excision. A representative experimental replicate is shown in panels **A, B.**

**A** Close-up plot of the representative extracted ion chromatogram (EIC) of the various peptide states; the blue trace corresponds to the Unmodified peptide; the orange trace corresponds to the Spliced peptide. **B** Close-up plot of the m/z values of the peptides detected by the mass spectrometer, with the shaded colored areas corresponding to where the expected m/z values for the peptide m/z traces should occur, with the observed spectrum plotted in black. The Unmodified, Spliced peptide(s) were found in the mass spectrum. **C** Bar plot showing the relative abundance of the various peptide states at charge z=2; the bars represent the average relative abundances normalized to total peak areas, with each individual experimental replicate value plotted as a black dot. **D** Mirrored MS$^2$ plot for the Unmodified peptide. In this mirror plot, the full fragmentation spectrum is plotted in black pointing upward, and the matched fragment b ions (blue) and y ions (red) are plotted pointing downward, mirrored against it, so that each matched fragment can be compared directly with the measured spectrum. **E** MS$^2$ spectrum of the fragments corresponding to the Unmodified peptide sequence, with matched b (blue, above) and y ions (red, below) plotted. **F** Mirrored MS$^2$ plot for the Spliced peptide. In this mirror plot, the full fragmentation spectrum is plotted in black pointing upward, and the matched fragment b ions (blue) and y ions (red) are plotted pointing downward, mirrored against it, so that each matched fragment can be compared directly with the measured spectrum. **G** MS$^2$ spectrum of the fragments corresponding to the Spliced peptide sequence, with matched b (blue, above) and y ions (red, below) plotted.

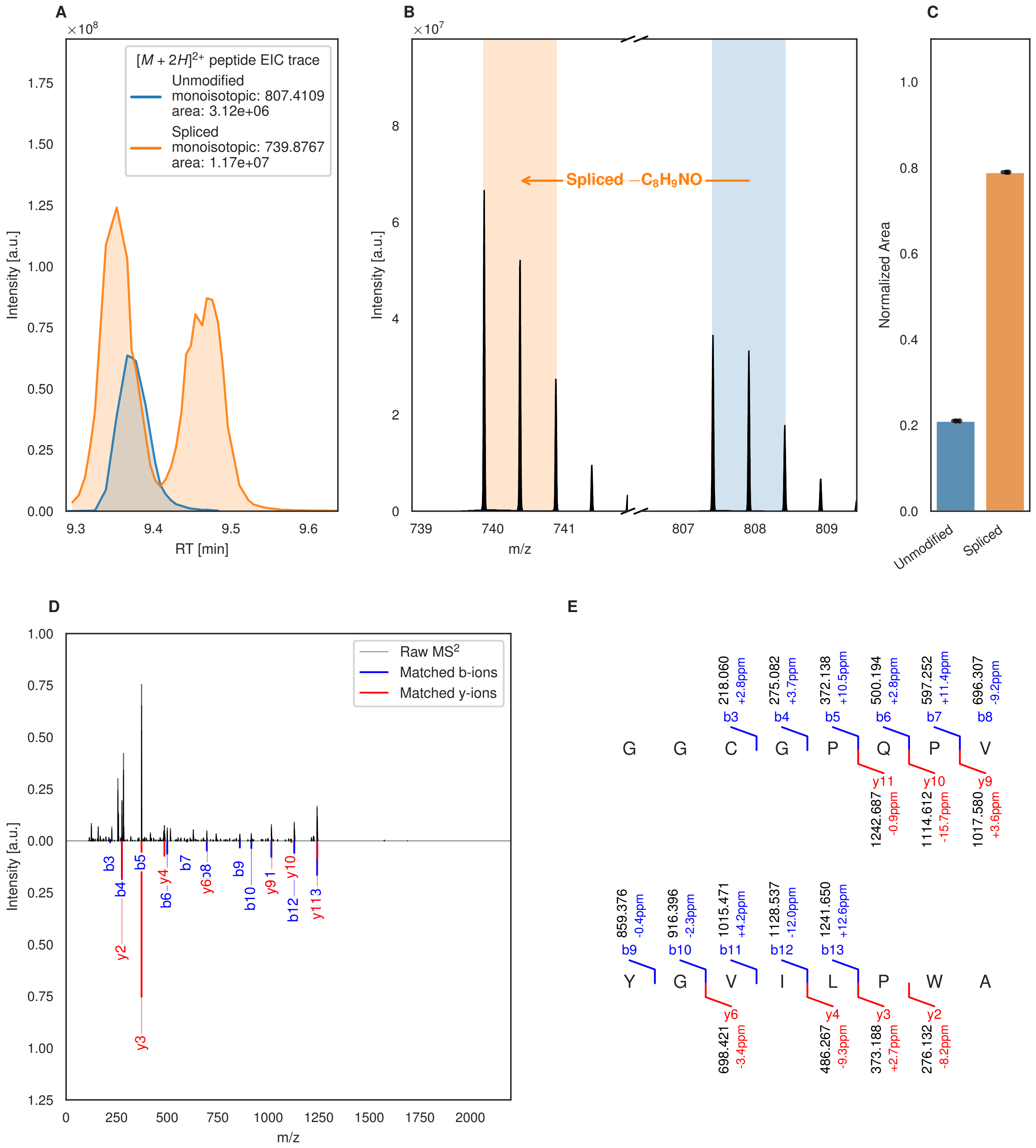
A
B
C
D
E
$[M+2H]^{2+}$ peptide EIC trace
Unmodified
monoisotopic: 807.4109
area: 3.12e+06
Spliced
monoisotopic: 739.8767
area: 1.17e+07
Intensity [a.u.]
RT [min]
Spliced $-C_8H_9NO$
m/z
Normalized Area
Unmodified
Spliced
Raw MS$^2$
Matched b-ions
Matched y-ions
G G C G P Q P V
Y G V I L P W A
b3 218.060 +2.8ppm
b4 275.082 +3.7ppm
b5 372.138 +10.5ppm
b6 500.194 +2.8ppm
b7 597.252 +11.4ppm
b8 696.307 -9.2ppm
y11 1242.687 -0.9ppm
y10 1114.612 -15.7ppm
y9 1017.580 +3.6ppm
b9 859.376 -0.4ppm
b10 916.396 -2.3ppm
b11 1015.471 +4.2ppm
b12 1128.537 -12.0ppm
b13 1241.650 +12.6ppm
y6 698.421 -3.4ppm
y4 486.267 -9.3ppm
y3 373.188 +2.7ppm
y2 276.132 -8.2ppm

Figure S28 (figure continued from the previous page).

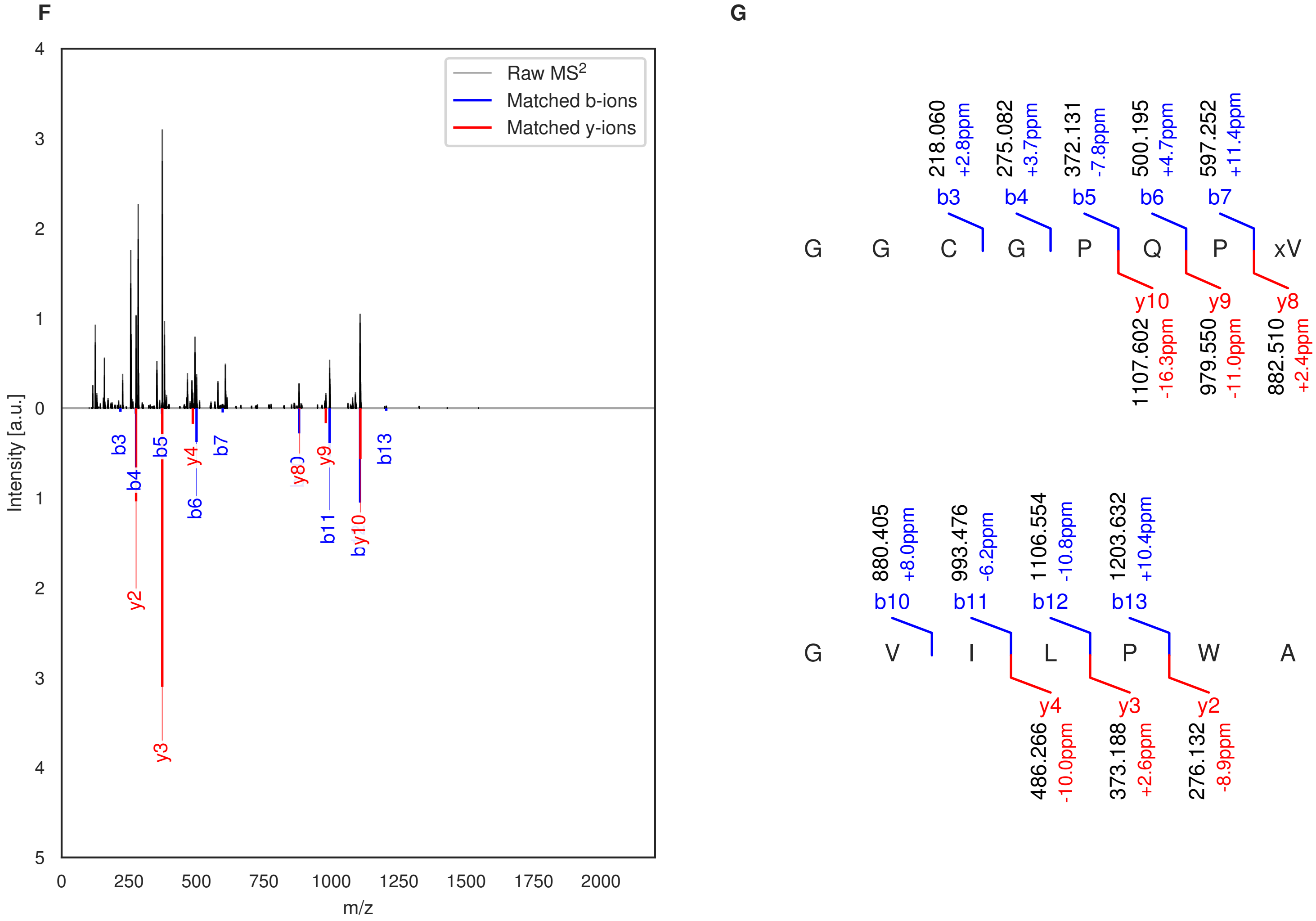

**Figure S28: Modification of the TJ-C-698 precursor peptide by PlpXY.** LC-MS analysis of IMAC-purified precursor peptides from *E. coli* BL21(DE3)Tuner containing splicease-encoding plasmid TJ-C-559 (PlpXY) and precursor peptide-encoding plasmid TJ-C-698 (Fh8-PlpA3-TEV-Artificial-2) after expression at 24 °C for 22 h in 30 mL cultures after digestion with TEV. Data from n=2 independent biological replicates. Spliced amino acids have an x as a prefix. Spliced refers to a single tyramine excision. A representative experimental replicate is shown in panels **A, B.**

**A** Close-up plot of the representative extracted ion chromatogram (EIC) of the various peptide states; the blue trace corresponds to the Unmodified peptide; the orange trace corresponds to the Spliced peptide. **B** Close-up plot of the m/z values of the peptides detected by the mass spectrometer, with the shaded colored areas corresponding to where the expected m/z values for the peptide m/z traces should occur, with the observed spectrum plotted in black. The Unmodified, Spliced peptide(s) were found in the mass spectrum. **C** Bar plot showing the relative abundance of the various peptide states at charge z=2; the bars represent the average relative abundances normalized to total peak areas, with each individual experimental replicate value plotted as a black dot. **D** Mirrored $MS^2$ plot for the Unmodified peptide. In this mirror plot, the full fragmentation spectrum is plotted in black pointing upward, and the matched fragment b ions (blue) and y ions (red) are plotted pointing downward, mirrored against it, so that each matched fragment can be compared directly with the measured spectrum. **E** $MS^2$ spectrum of the fragments corresponding to the Unmodified peptide sequence, with matched b (blue, above) and y ions (red, below) plotted. **F** Mirrored $MS^2$ plot for the Spliced peptide. In this mirror plot, the full fragmentation spectrum is plotted in black pointing upward, and the matched fragment b ions (blue) and y ions (red) are plotted pointing downward, mirrored against it, so that each matched fragment can be compared directly with the measured spectrum. **G** $MS^2$ spectrum of the fragments corresponding to the Spliced peptide sequence, with matched b (blue, above) and y ions (red, below) plotted.

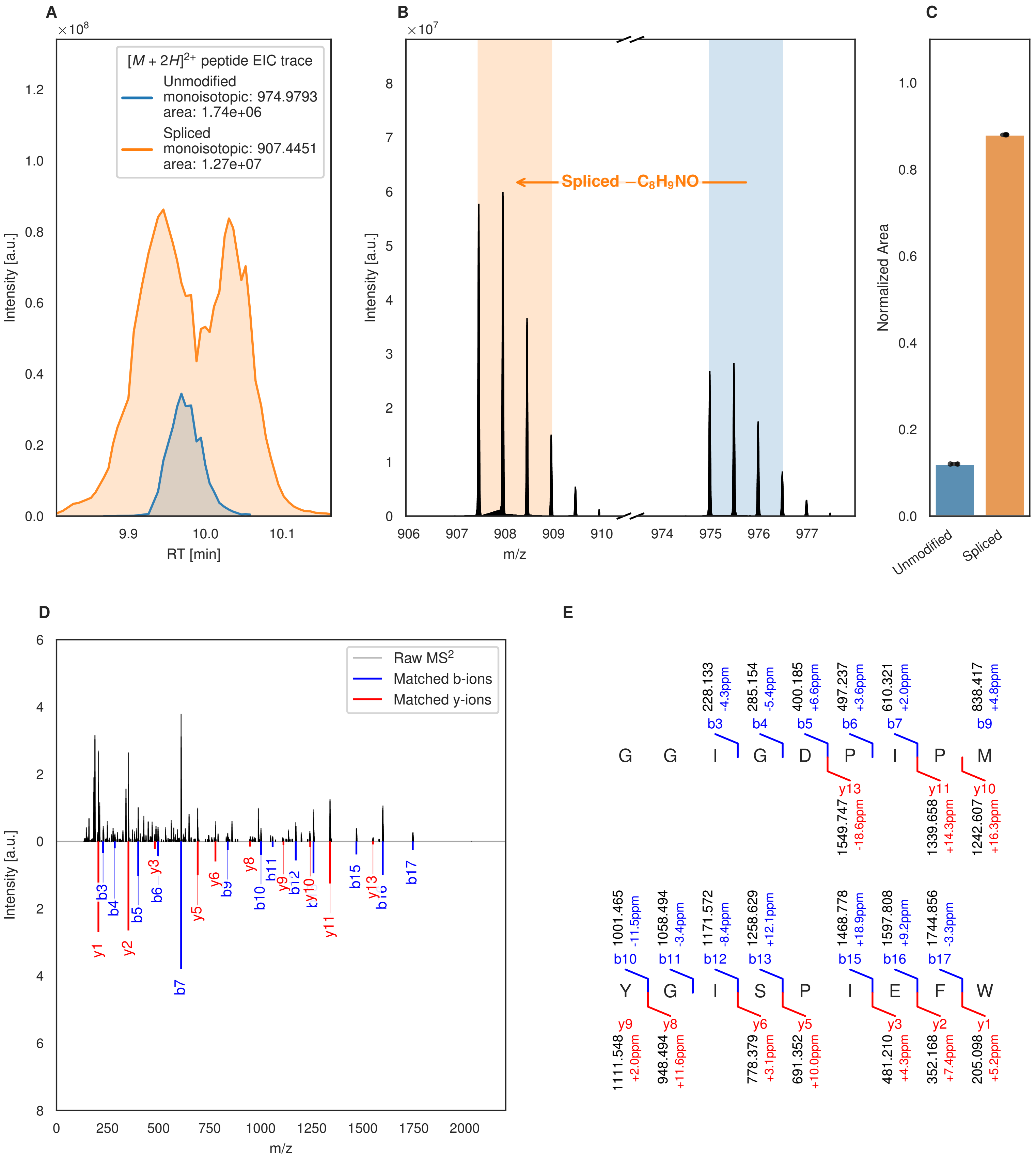

A
[M + 2H]2+ peptide EIC trace
Unmodified
monoisotopic: 974.9793
area: 1.74e+06
Spliced
monoisotopic: 907.4451
area: 1.27e+07
Intensity [a.u.]
RT [min]
B
Spliced −C8H9NO
Intensity [a.u.]
m/z
C
Normalized Area
Unmodified
Spliced
D
Raw MS2
Matched b-ions
Matched y-ions
Intensity [a.u.]
m/z
E
G G I G D P I P M
228.133 -4.3ppm b3
285.154 -5.4ppm b4
400.185 +6.6ppm b5
497.237 +3.6ppm b6
610.321 +2.0ppm b7
838.417 +4.8ppm b9
y13 1549.747 -18.6ppm
y11 1339.658 +14.3ppm
y10 1242.607 +16.3ppm
Y G I S P I E F W
1001.465 -11.5ppm b10
1058.494 -3.4ppm b11
1171.572 -8.4ppm b12
1258.629 +12.1ppm b13
1468.778 +18.9ppm b15
1597.808 +9.2ppm b16
1744.856 -3.3ppm b17
y9 1111.548 +2.0ppm
y8 948.494 +11.6ppm
y6 778.379 +3.1ppm
y5 691.352 +10.0ppm
y3 481.210 +4.3ppm
y2 352.168 +7.4ppm
y1 205.098 +5.2ppm

Figure S29 (figure continued from the previous page).

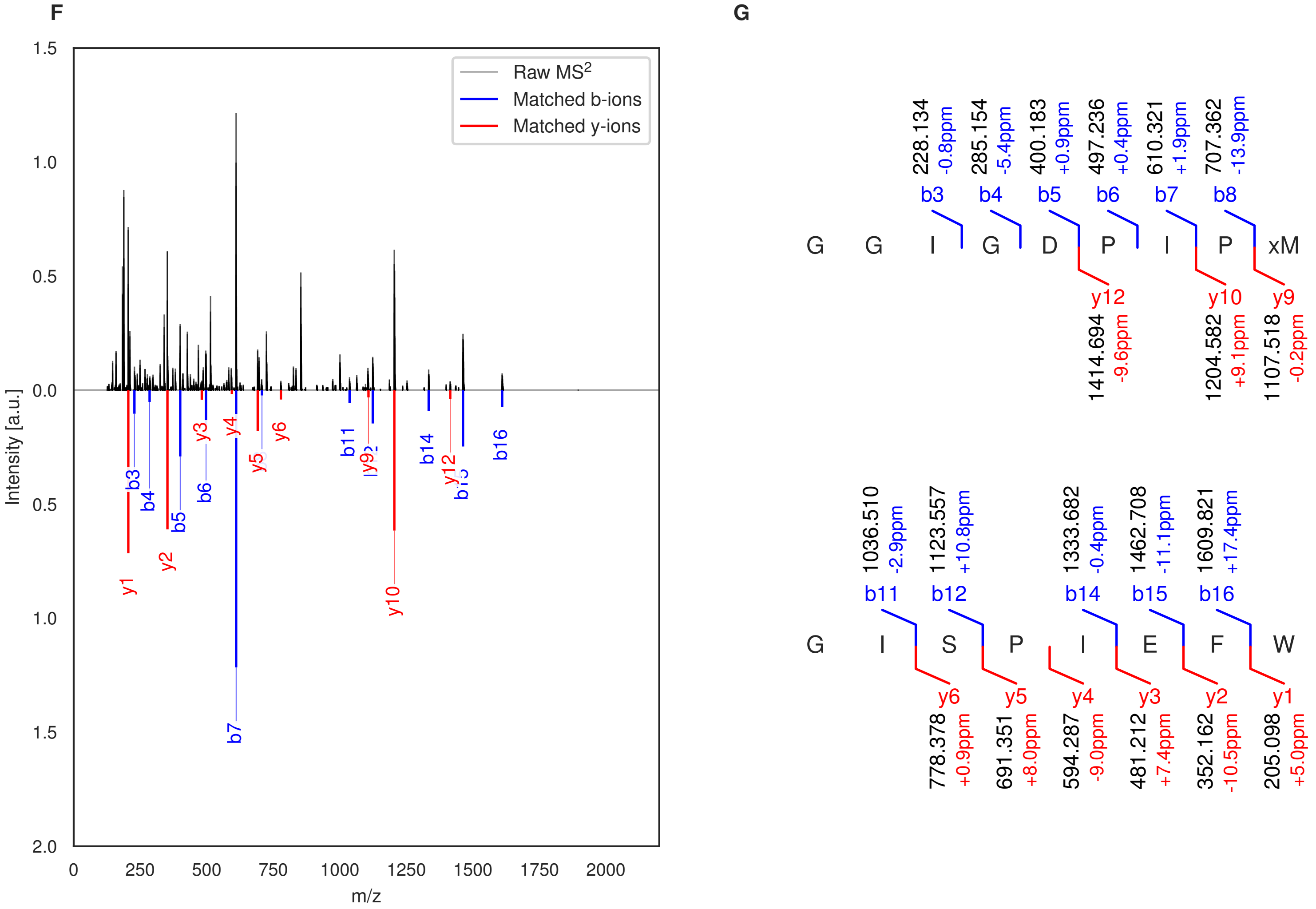

**Figure S29: Modification of the TJ-C-699 precursor peptide by PlpXY.** LC-MS analysis of IMAC-purified precursor peptides from *E. coli* BL21(DE3)Tuner containing splicease-encoding plasmid TJ-C-559 (PlpXY) and precursor peptide-encoding plasmid TJ-C-699 (Fh8-PlpA3-TEV-Artificial-3) after expression at 24 °C for 22 h in 30 mL cultures after digestion with TEV. Data from n=2 independent biological replicates. Spliced amino acids have an x as a prefix. Spliced refers to a single tyramine excision. A representative experimental replicate is shown in panels **A, B.**

**A** Close-up plot of the representative extracted ion chromatogram (EIC) of the various peptide states; the blue trace corresponds to the Unmodified peptide; the orange trace corresponds to the Spliced peptide. **B** Close-up plot of the m/z values of the peptides detected by the mass spectrometer, with the shaded colored areas corresponding to where the expected m/z values for the peptide m/z traces should occur, with the observed spectrum plotted in black. The Unmodified, Spliced peptide(s) were found in the mass spectrum. **C** Bar plot showing the relative abundance of the various peptide states at charge z=2; the bars represent the average relative abundances normalized to total peak areas, with each individual experimental replicate value plotted as a black dot. **D** Mirrored MS$^2$ plot for the Unmodified peptide. In this mirror plot, the full fragmentation spectrum is plotted in black pointing upward, and the matched fragment b ions (blue) and y ions (red) are plotted pointing downward, mirrored against it, so that each matched fragment can be compared directly with the measured spectrum. **E** MS$^2$ spectrum of the fragments corresponding to the Unmodified peptide sequence, with matched b (blue, above) and y ions (red, below) plotted. **F** Mirrored MS$^2$ plot for the Spliced peptide. In this mirror plot, the full fragmentation spectrum is plotted in black pointing upward, and the matched fragment b ions (blue) and y ions (red) are plotted pointing downward, mirrored against it, so that each matched fragment can be compared directly with the measured spectrum. **G** MS$^2$ spectrum of the fragments corresponding to the Spliced peptide sequence, with matched b (blue, above) and y ions (red, below) plotted.

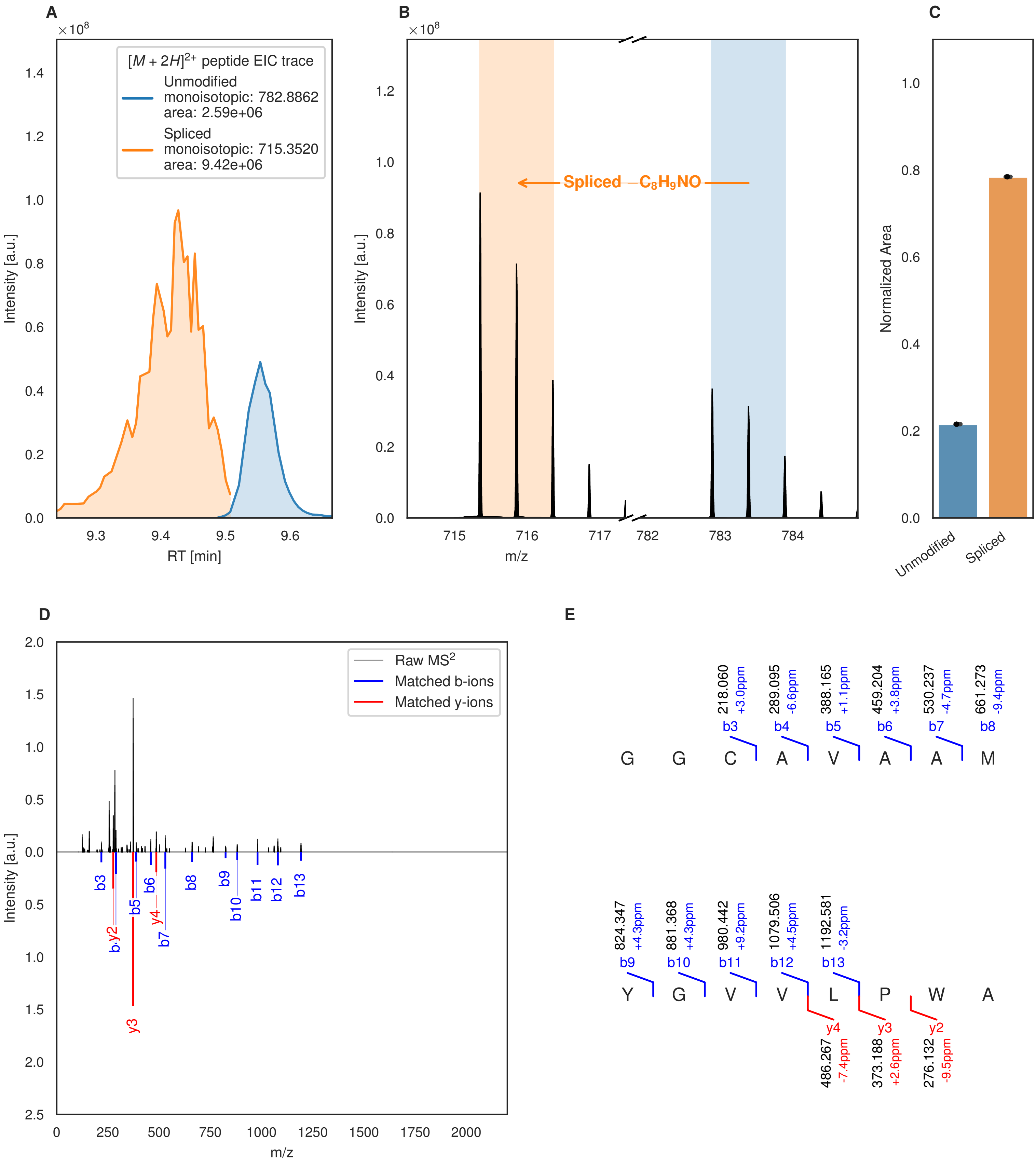
A
×10^8
[M + 2H]2+ peptide EIC trace
Unmodified
monoisotopic: 782.8862
area: 2.59e+06
Spliced
monoisotopic: 715.3520
area: 9.42e+06
Intensity [a.u.]
RT [min]
B
×10^8
Spliced −C8H9NO
Intensity [a.u.]
m/z
C
Normalized Area
Unmodified
Spliced
D
Raw MS2
Matched b-ions
Matched y-ions
Intensity [a.u.]
m/z
b3
b.y2
b5
b6
y4
b7
b8
b9
b10
b11
b12
b13
y3
E
218.060 +3.0ppm
289.095 -6.6ppm
388.165 +1.1ppm
459.204 +3.8ppm
530.237 -4.7ppm
661.273 -9.4ppm
b3 b4 b5 b6 b7 b8
G G C A V A A M
824.347 +4.3ppm
881.368 +4.3ppm
980.442 +9.2ppm
1079.506 +4.5ppm
1192.581 -3.2ppm
b9 b10 b11 b12 b13
Y G V V L P W A
y4 y3 y2
486.267 -7.4ppm
373.188 +2.6ppm
276.132 -9.5ppm

Figure S30 (figure continued from the previous page).

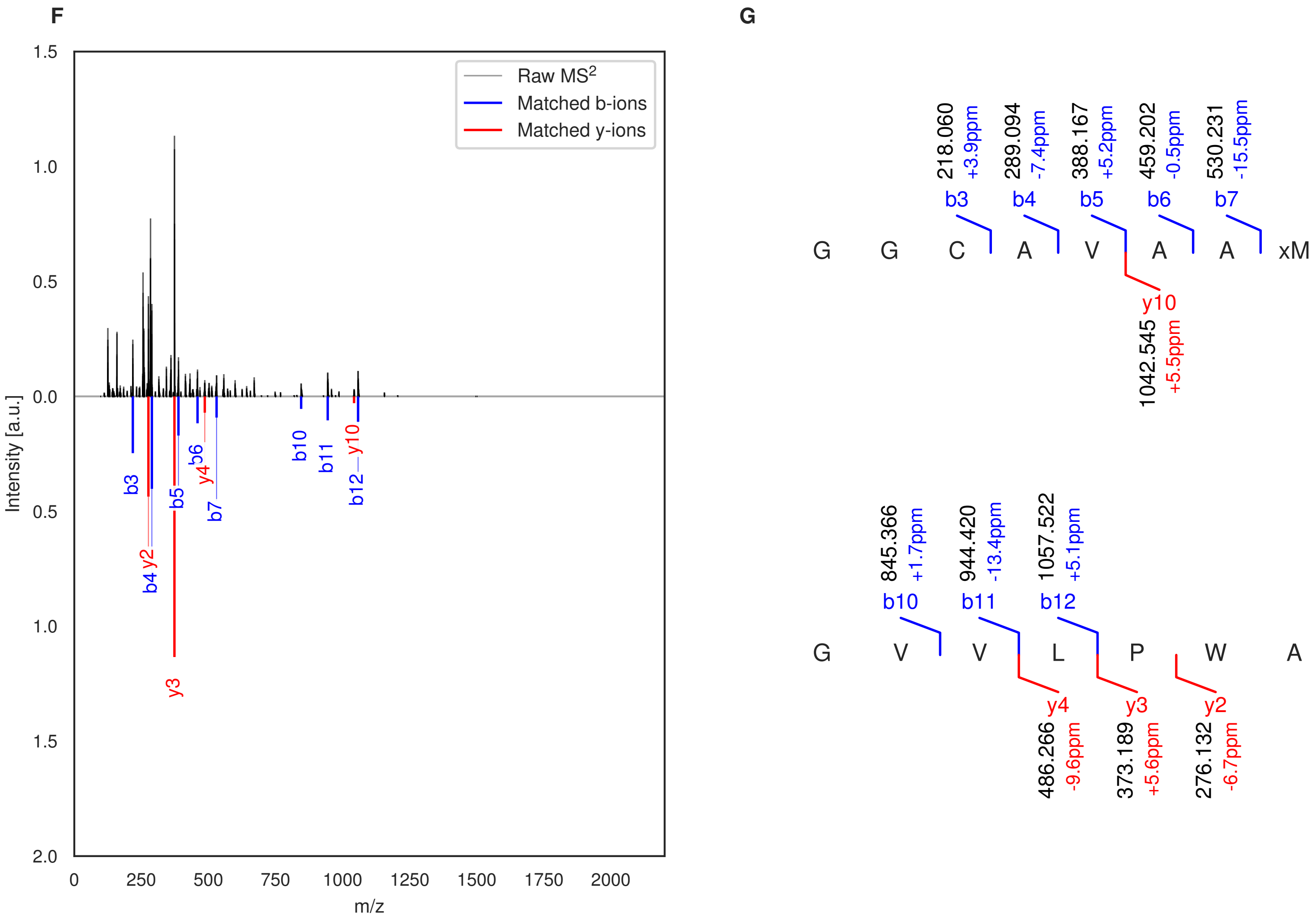

**Figure S30: Modification of the TJ-C-700 precursor peptide by PlpXY.** LC-MS analysis of IMAC-purified precursor peptides from *E. coli* BL21(DE3)Tuner containing splicease-encoding plasmid TJ-C-559 (PlpXY) and precursor peptide-encoding plasmid TJ-C-700 (Fh8-PlpA3-TEV-Artificial-4) after expression at 24 °C for 22 h in 30 mL cultures after digestion with TEV. Data from n=2 independent biological replicates. Spliced amino acids have an x as a prefix. Spliced refers to a single tyramine excision. A representative experimental replicate is shown in panels **A, B.**

**A** Close-up plot of the representative extracted ion chromatogram (EIC) of the various peptide states; the blue trace corresponds to the Unmodified peptide; the orange trace corresponds to the Spliced peptide. **B** Close-up plot of the m/z values of the peptides detected by the mass spectrometer, with the shaded colored areas corresponding to where the expected m/z values for the peptide m/z traces should occur, with the observed spectrum plotted in black. The Unmodified, Spliced peptide(s) were found in the mass spectrum. **C** Bar plot showing the relative abundance of the various peptide states at charge z=2; the bars represent the average relative abundances normalized to total peak areas, with each individual experimental replicate value plotted as a black dot. **D** Mirrored MS$^2$ plot for the Unmodified peptide. In this mirror plot, the full fragmentation spectrum is plotted in black pointing upward, and the matched fragment b ions (blue) and y ions (red) are plotted pointing downward, mirrored against it, so that each matched fragment can be compared directly with the measured spectrum. **E** MS$^2$ spectrum of the fragments corresponding to the Unmodified peptide sequence, with matched b (blue, above) and y ions (red, below) plotted. **F** Mirrored MS$^2$ plot for the Spliced peptide. In this mirror plot, the full fragmentation spectrum is plotted in black pointing upward, and the matched fragment b ions (blue) and y ions (red) are plotted pointing downward, mirrored against it, so that each matched fragment can be compared directly with the measured spectrum. **G** MS$^2$ spectrum of the fragments corresponding to the Spliced peptide sequence, with matched b (blue, above) and y ions (red, below) plotted.

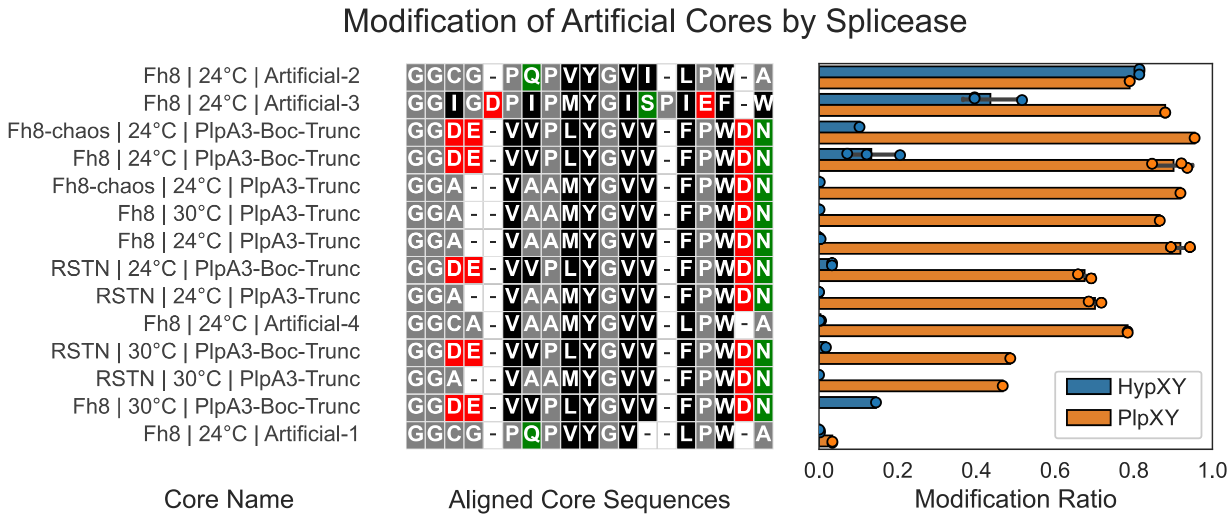

**Figure S31: Effect of temperature and solubility tag on the splicing efficiency.** The labels describe the solubility tag used (RSTN, Fh8, or Fh8-chaos), the expression temperature used, and the name of the core sequence. The bars on the right show the relative modification efficiencies of the HypX and PlpX spliceases on three different core sequences. Artificial-2 is the same core sequence used for screening the point mutants. The leader used for HypX is the consensus leader HypAc-TEV, and for PlpX it is PlpA3-TEV. In shake flasks, the modification efficiency of the RSTN-tagged precursor is slightly lower compared to the Fh8 tag, which we attribute to increased overall expression levels. The Fh8-chaos tag (a Fh8 tag with a C-terminal asparagine-rich chaotropic linker) behaves similarly to the Fh8 tag. The PlpA3-Boc-Trunc peptide is derived from the original splicease paper from Morinaka et al.[1] This particular core is modified much more effectively by the PlpX splicease compared to HypX. Bars show the mean ± SD of n = 1-3 biological replicates; points are individual biological replicates.

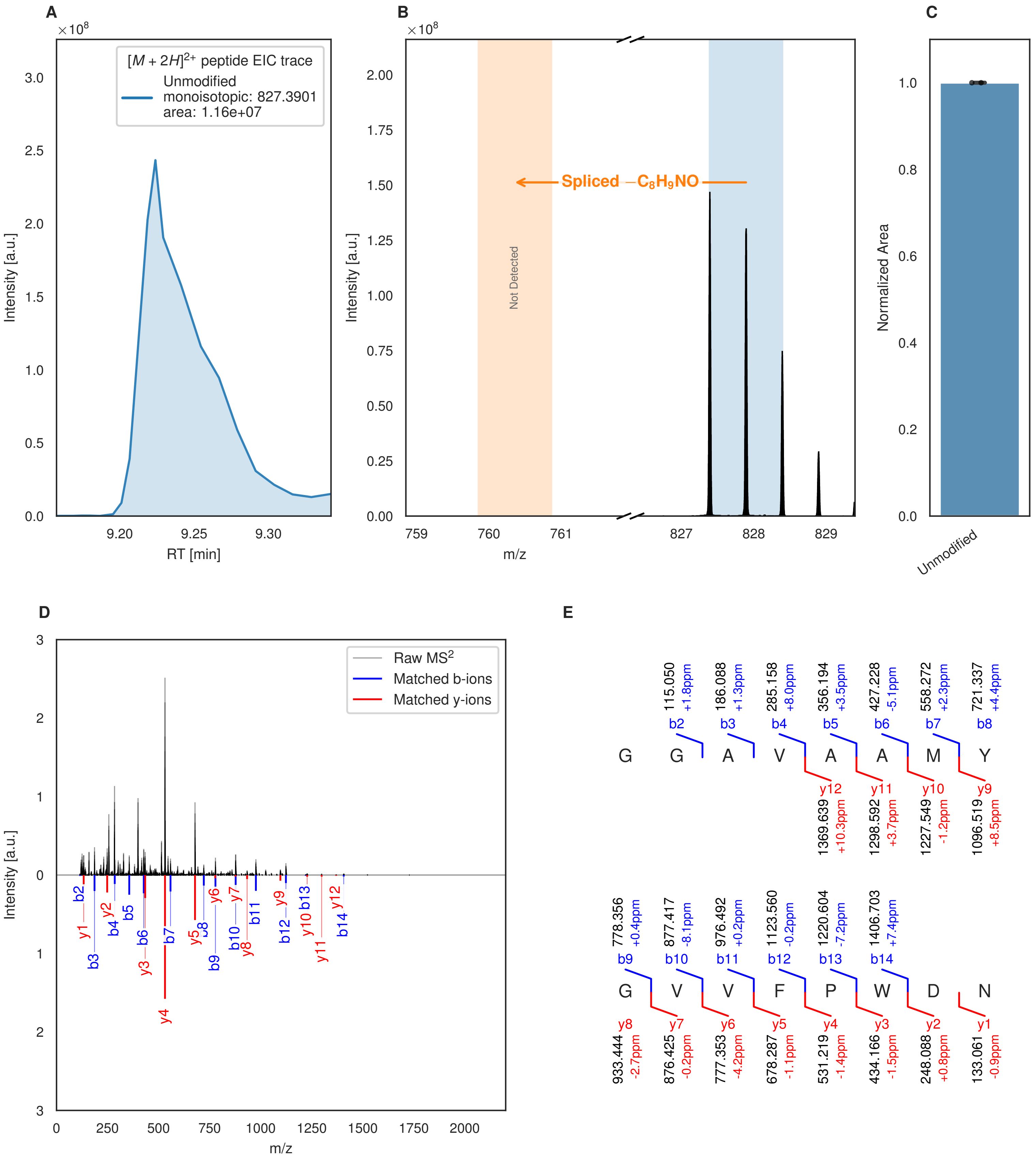

**Figure S32: Modification of the TJ-C-674 precursor peptide by HypXY.** LC-MS analysis of IMAC-purified precursor peptides from *E. coli* BL21(DE3)Tuner containing splicease-encoding plasmid TJ-C-560 (HypXY) and precursor peptide-encoding plasmid TJ-C-674 (RSTN1-HypAc-TEV-PlpA3-Trunc) after expression at 24 °C for 22 h in 30 mL cultures after digestion with TEV. Data from n=2 independent biological replicates. Spliced amino acids have an x as a prefix. Spliced refers to a single tyramine excision. A representative experimental replicate is shown in panels **A, B.**

**A** Close-up plot of the representative extracted ion chromatogram (EIC) of the various peptide states; the blue trace corresponds to the Unmodified peptide. **B** Close-up plot of the m/z values of the peptides detected by the mass spectrometer, with the shaded colored areas corresponding to where the expected m/z values for the peptide m/z

(Caption continued on the next page.)

traces should occur, with the observed spectrum plotted in black. The Unmodified peptide(s) were found in the mass spectrum and the Spliced peptide(s) were not detected. **C** Bar plot showing the relative abundance of the various peptide states at charge z=2; the bars represent the average relative abundances normalized to total peak areas, with each individual experimental replicate value plotted as a black dot. **D** Mirrored $MS^2$ plot for the Unmodified peptide. In this mirror plot, the full fragmentation spectrum is plotted in black pointing upward, and the matched fragment b ions (blue) and y ions (red) are plotted pointing downward, mirrored against it, so that each matched fragment can be compared directly with the measured spectrum. **E** $MS^2$ spectrum of the fragments corresponding to the Unmodified peptide sequence, with matched b (blue, above) and y ions (red, below) plotted.

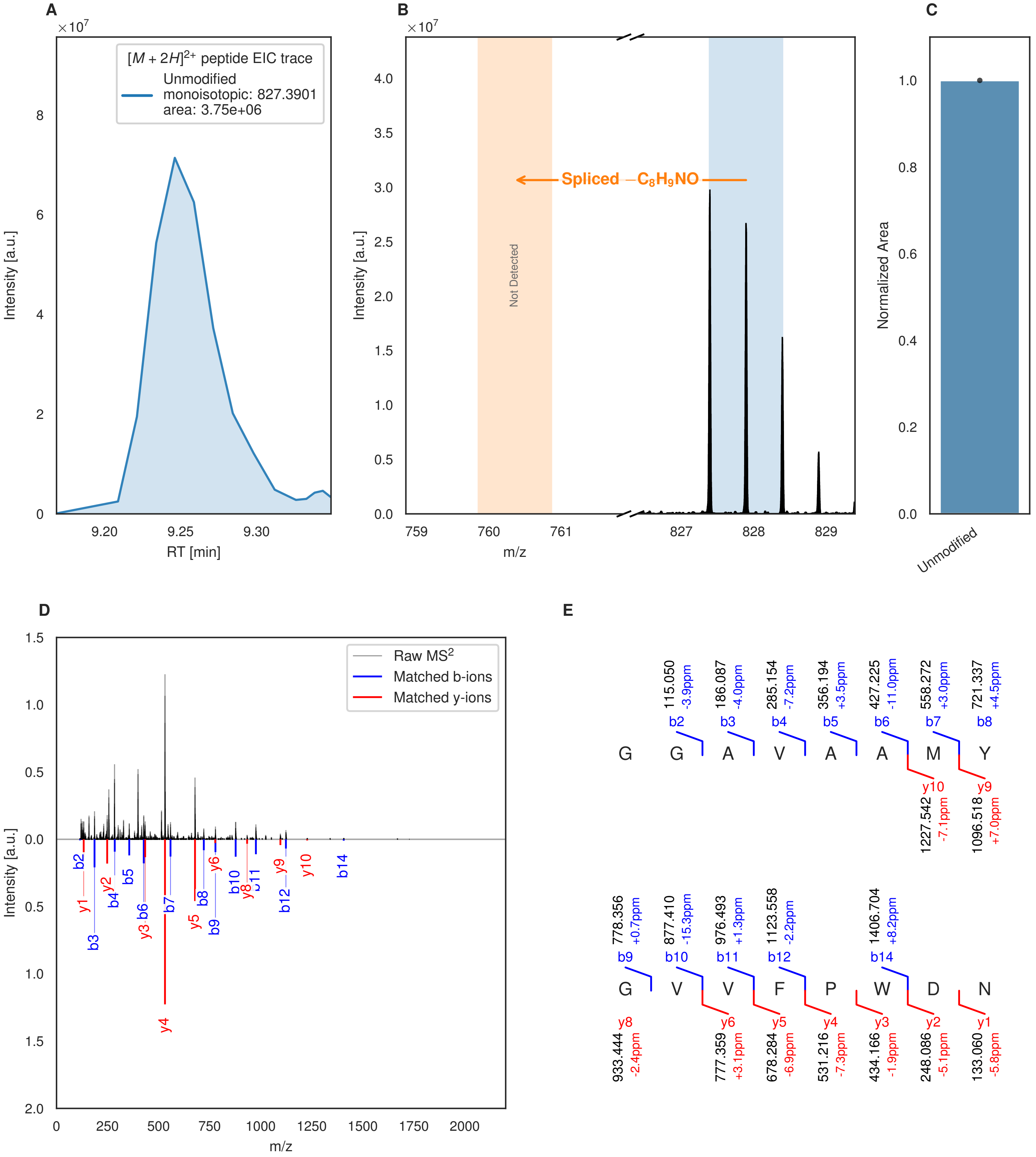

**Figure S33: Modification of the TJ-C-674 precursor peptide by HypXY.** LC-MS analysis of IMAC-purified precursor peptides from *E. coli* BL21(DE3)Tuner containing splicease-encoding plasmid TJ-C-560 (HypXY) and precursor peptide-encoding plasmid TJ-C-674 (RSTN1-HypAc-TEV-PlpA3-Trunc) after expression at 30 °C for 22 h in 30 mL cultures after digestion with TEV. Data from n=1 independent biological replicates. Spliced amino acids have an x as a prefix. Spliced refers to a single tyramine excision. A representative experimental replicate is shown in panels **A, B.**

**A** Close-up plot of the representative extracted ion chromatogram (EIC) of the various peptide states; the blue trace corresponds to the Unmodified peptide. **B** Close-up plot of the m/z values of the peptides detected by the mass spectrometer, with the shaded colored areas corresponding to where the expected m/z values for the peptide m/z

(Caption continued on the next page.)

traces should occur, with the observed spectrum plotted in black. The Unmodified peptide(s) were found in the mass spectrum and the Spliced peptide(s) were not detected. **C** Bar plot showing the relative abundance of the various peptide states at charge z=2; the bars represent the average relative abundances normalized to total peak areas, with each individual experimental replicate value plotted as a black dot. **D** Mirrored $MS^2$ plot for the Unmodified peptide. In this mirror plot, the full fragmentation spectrum is plotted in black pointing upward, and the matched fragment b ions (blue) and y ions (red) are plotted pointing downward, mirrored against it, so that each matched fragment can be compared directly with the measured spectrum. **E** $MS^2$ spectrum of the fragments corresponding to the Unmodified peptide sequence, with matched b (blue, above) and y ions (red, below) plotted.

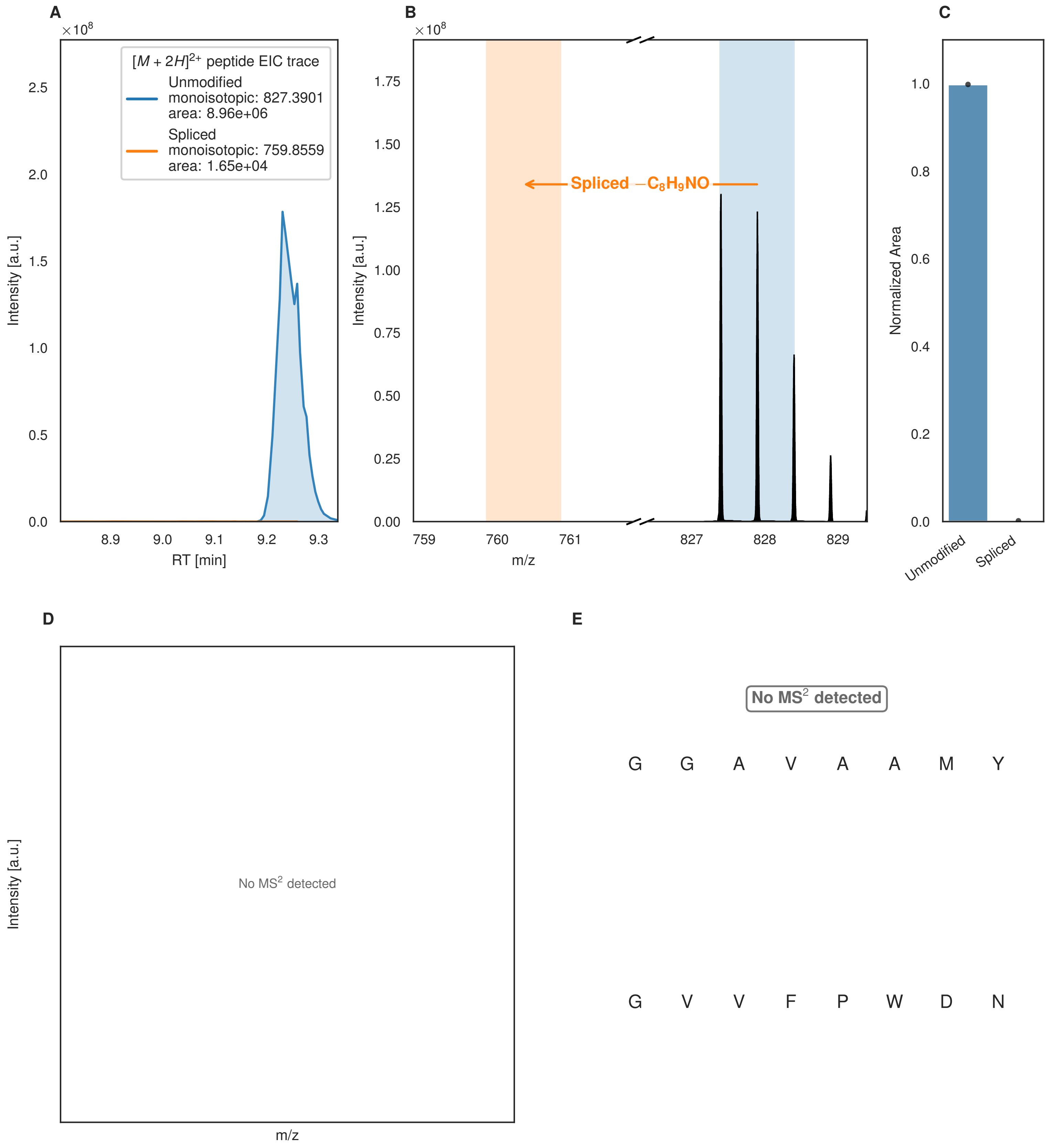

A
×10^8
[M + 2H]2+ peptide EIC trace
Unmodified
monoisotopic: 827.3901
area: 8.96e+06
Spliced
monoisotopic: 759.8559
area: 1.65e+04
Intensity [a.u.]
RT [min]
0.0
0.5
1.0
1.5
2.0
2.5
8.9
9.0
9.1
9.2
9.3
B
×10^8
Spliced −C8H9NO
Intensity [a.u.]
m/z
0.00
0.25
0.50
0.75
1.00
1.25
1.50
1.75
759
760
761
827
828
829
C
Normalized Area
0.0
0.2
0.4
0.6
0.8
1.0
Unmodified
Spliced
D
No MS2 detected
Intensity [a.u.]
m/z
E
No MS2 detected
G G A V A A M Y
G V V F P W D N

Figure S34 (figure continued from the previous page).

F

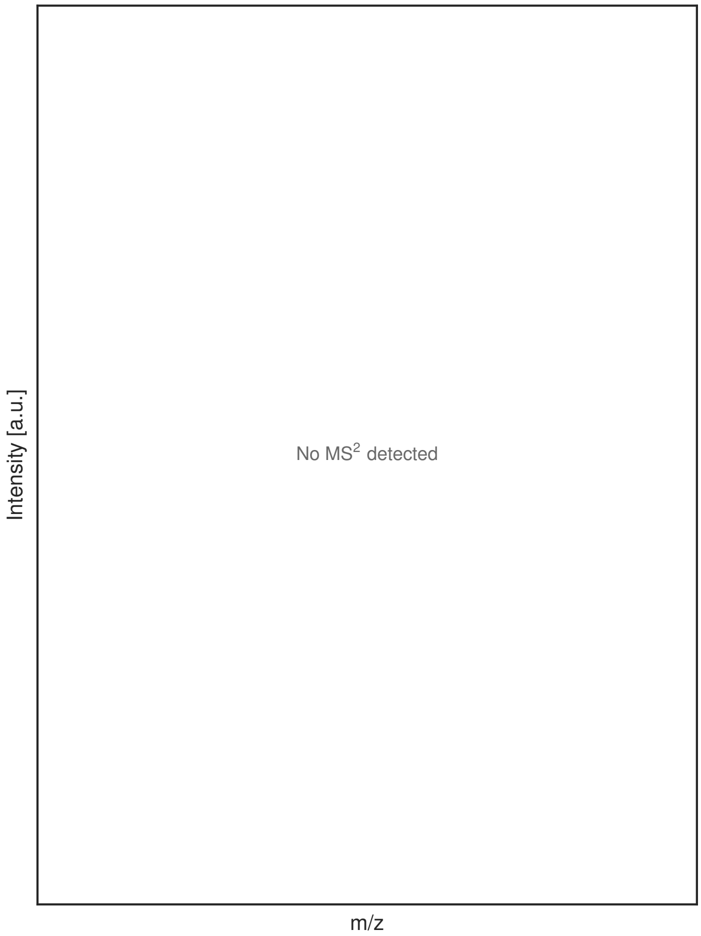

G

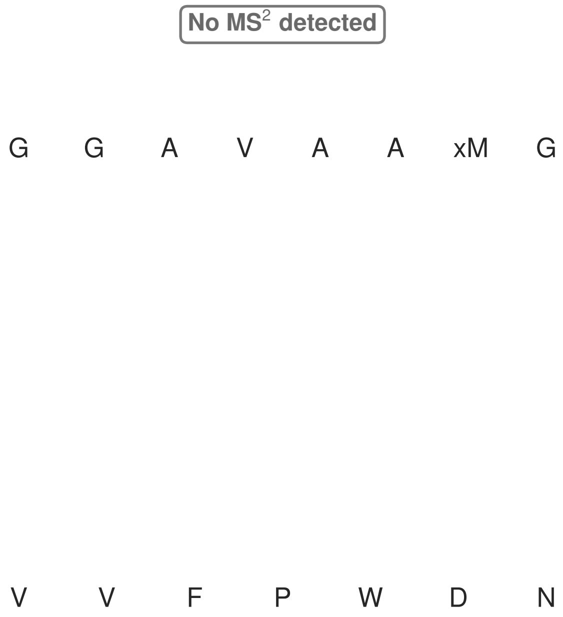

**Figure S34: Modification of the TJ-C-675 precursor peptide by HypXY.** LC-MS analysis of IMAC-purified precursor peptides from *E. coli* BL21(DE3)Tuner containing splicease-encoding plasmid TJ-C-560 (HypXY) and precursor peptide-encoding plasmid TJ-C-675 (Fh8-chaos-HypAc-TEV-PlpA3-Trunc) after expression at 24 °C for 22 h in 30 mL cultures after digestion with TEV. Data from n=1 independent biological replicates. Spliced amino acids have an x as a prefix. Spliced refers to a single tyramine excision. A representative experimental replicate is shown in panels **A, B.**

**A** Close-up plot of the representative extracted ion chromatogram (EIC) of the various peptide states; the blue trace corresponds to the Unmodified peptide; the orange trace corresponds to the Spliced peptide. **B** Close-up plot of the m/z values of the peptides detected by the mass spectrometer, with the shaded colored areas corresponding to where the expected m/z values for the peptide m/z traces should occur, with the observed spectrum plotted in black. The Unmodified, Spliced peptide(s) were found in the mass spectrum. **C** Bar plot showing the relative abundance of the various peptide states at charge z=2; the bars represent the average relative abundances normalized to total peak areas, with each individual experimental replicate value plotted as a black dot. **D** Mirrored MS$^2$ plot for the Unmodified peptide. In this mirror plot, the full fragmentation spectrum is plotted in black pointing upward, and the matched fragment b ions (blue) and y ions (red) are plotted pointing downward, mirrored against it, so that each matched fragment can be compared directly with the measured spectrum. **E** MS$^2$ spectrum of the fragments corresponding to the Unmodified peptide sequence, with matched b (blue, above) and y ions (red, below) plotted. **F** Mirrored MS$^2$ plot for the Spliced peptide. In this mirror plot, the full fragmentation spectrum is plotted in black pointing upward, and the matched fragment b ions (blue) and y ions (red) are plotted pointing downward, mirrored against it, so that each matched fragment can be compared directly with the measured spectrum. **G** MS$^2$ spectrum of the fragments corresponding to the Spliced peptide sequence, with matched b (blue, above) and y ions (red, below) plotted.

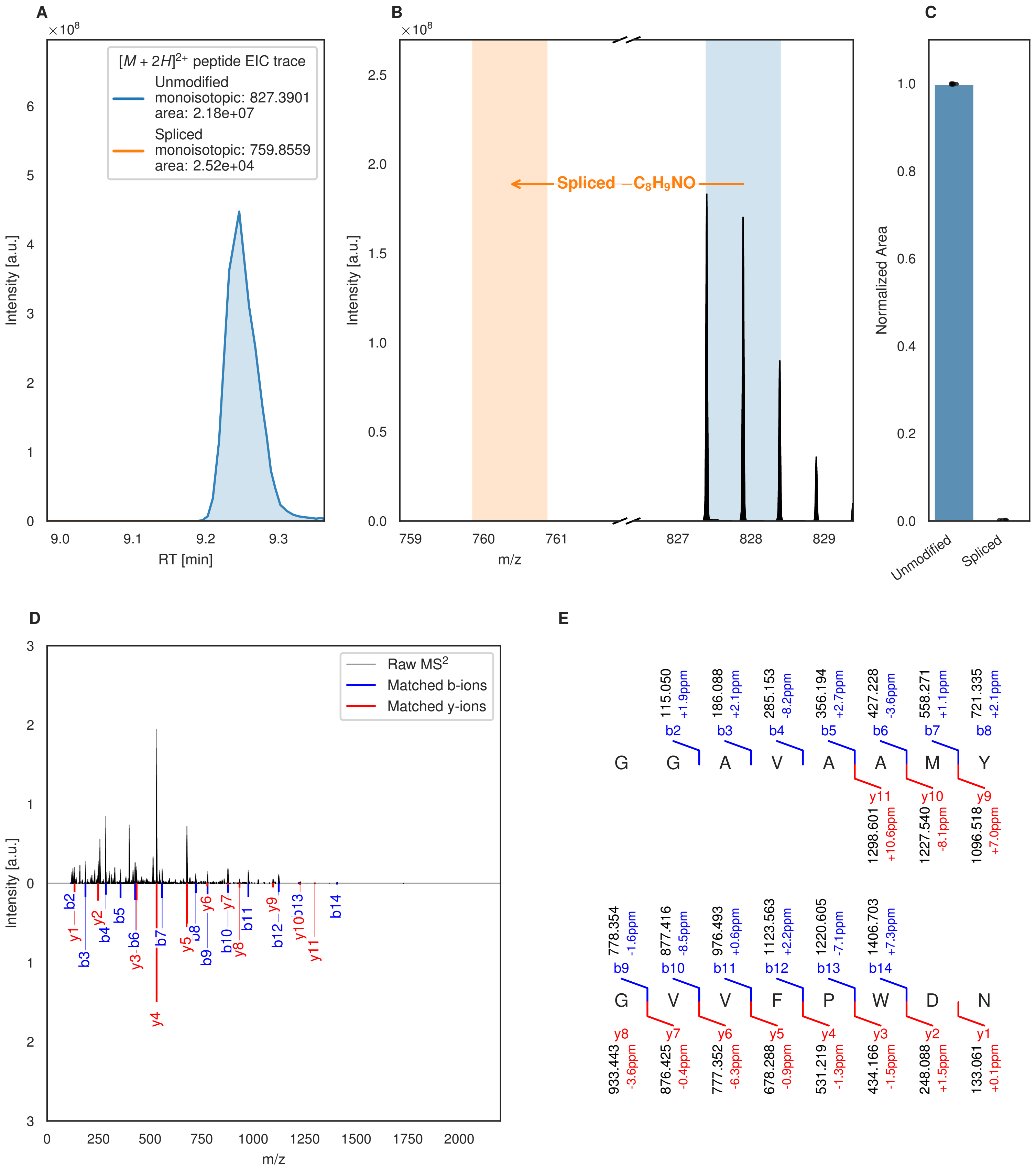
A
×10^8
[M + 2H]2+ peptide EIC trace
Unmodified
monoisotopic: 827.3901
area: 2.18e+07
Spliced
monoisotopic: 759.8559
area: 2.52e+04
Intensity [a.u.]
RT [min]
B
×10^8
Spliced −C8H9NO
Intensity [a.u.]
m/z
C
Normalized Area
Unmodified
Spliced
D
Raw MS2
Matched b-ions
Matched y-ions
Intensity [a.u.]
m/z
E
115.050 +1.9ppm b2
186.088 +2.1ppm b3
285.153 -8.2ppm b4
356.194 +2.7ppm b5
427.228 -3.6ppm b6
558.271 +1.1ppm b7
721.335 +2.1ppm b8
G G A V A A M Y
y11 1298.601 +10.6ppm
y10 1227.540 -8.1ppm
y9 1096.518 +7.0ppm
778.354 -1.6ppm b9
877.416 -8.5ppm b10
976.493 +0.6ppm b11
1123.563 +2.2ppm b12
1220.605 -7.1ppm b13
1406.703 +7.3ppm b14
G V V F P W D N
y8 933.443 -3.6ppm
y7 876.425 -0.4ppm
y6 777.352 -6.3ppm
y5 678.288 -0.9ppm
y4 531.219 -1.3ppm
y3 434.166 -1.5ppm
y2 248.088 +1.5ppm
y1 133.061 +0.1ppm

Figure S35 (figure continued from the previous page).

F

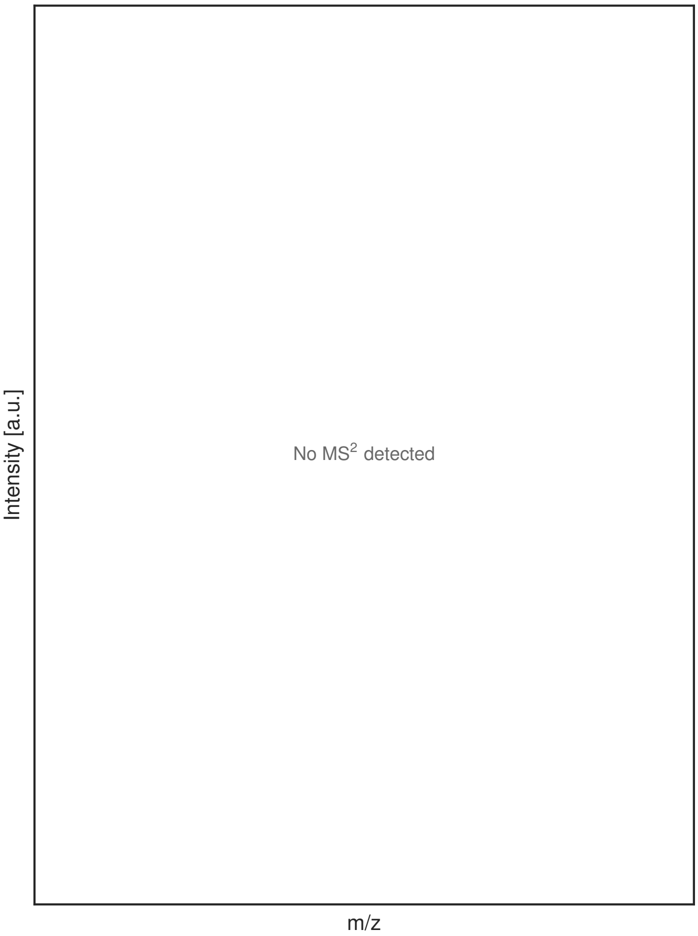

G

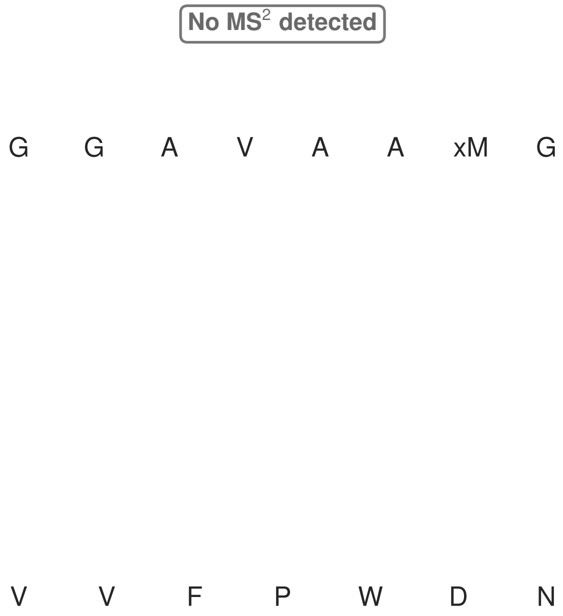

**Figure S35: Modification of the TJ-C-676 precursor peptide by HypXY.** LC-MS analysis of IMAC-purified precursor peptides from *E. coli* BL21(DE3)Tuner containing splicease-encoding plasmid TJ-C-560 (HypXY) and precursor peptide-encoding plasmid TJ-C-676 (Fh8-HypAc-TEV-PlpA3-Trunc) after expression at 24 °C for 22 h in 30 mL cultures after digestion with TEV. Data from n=2 independent biological replicates. Spliced amino acids have an x as a prefix. Spliced refers to a single tyramine excision. A representative experimental replicate is shown in panels **A, B.**

**A** Close-up plot of the representative extracted ion chromatogram (EIC) of the various peptide states; the blue trace corresponds to the Unmodified peptide; the orange trace corresponds to the Spliced peptide. **B** Close-up plot of the m/z values of the peptides detected by the mass spectrometer, with the shaded colored areas corresponding to where the expected m/z values for the peptide m/z traces should occur, with the observed spectrum plotted in black. The Unmodified, Spliced peptide(s) were found in the mass spectrum. **C** Bar plot showing the relative abundance of the various peptide states at charge z=2; the bars represent the average relative abundances normalized to total peak areas, with each individual experimental replicate value plotted as a black dot. **D** Mirrored MS$^2$ plot for the Unmodified peptide. In this mirror plot, the full fragmentation spectrum is plotted in black pointing upward, and the matched fragment b ions (blue) and y ions (red) are plotted pointing downward, mirrored against it, so that each matched fragment can be compared directly with the measured spectrum. **E** MS$^2$ spectrum of the fragments corresponding to the Unmodified peptide sequence, with matched b (blue, above) and y ions (red, below) plotted. **F** Mirrored MS$^2$ plot for the Spliced peptide. In this mirror plot, the full fragmentation spectrum is plotted in black pointing upward, and the matched fragment b ions (blue) and y ions (red) are plotted pointing downward, mirrored against it, so that each matched fragment can be compared directly with the measured spectrum. **G** MS$^2$ spectrum of the fragments corresponding to the Spliced peptide sequence, with matched b (blue, above) and y ions (red, below) plotted.

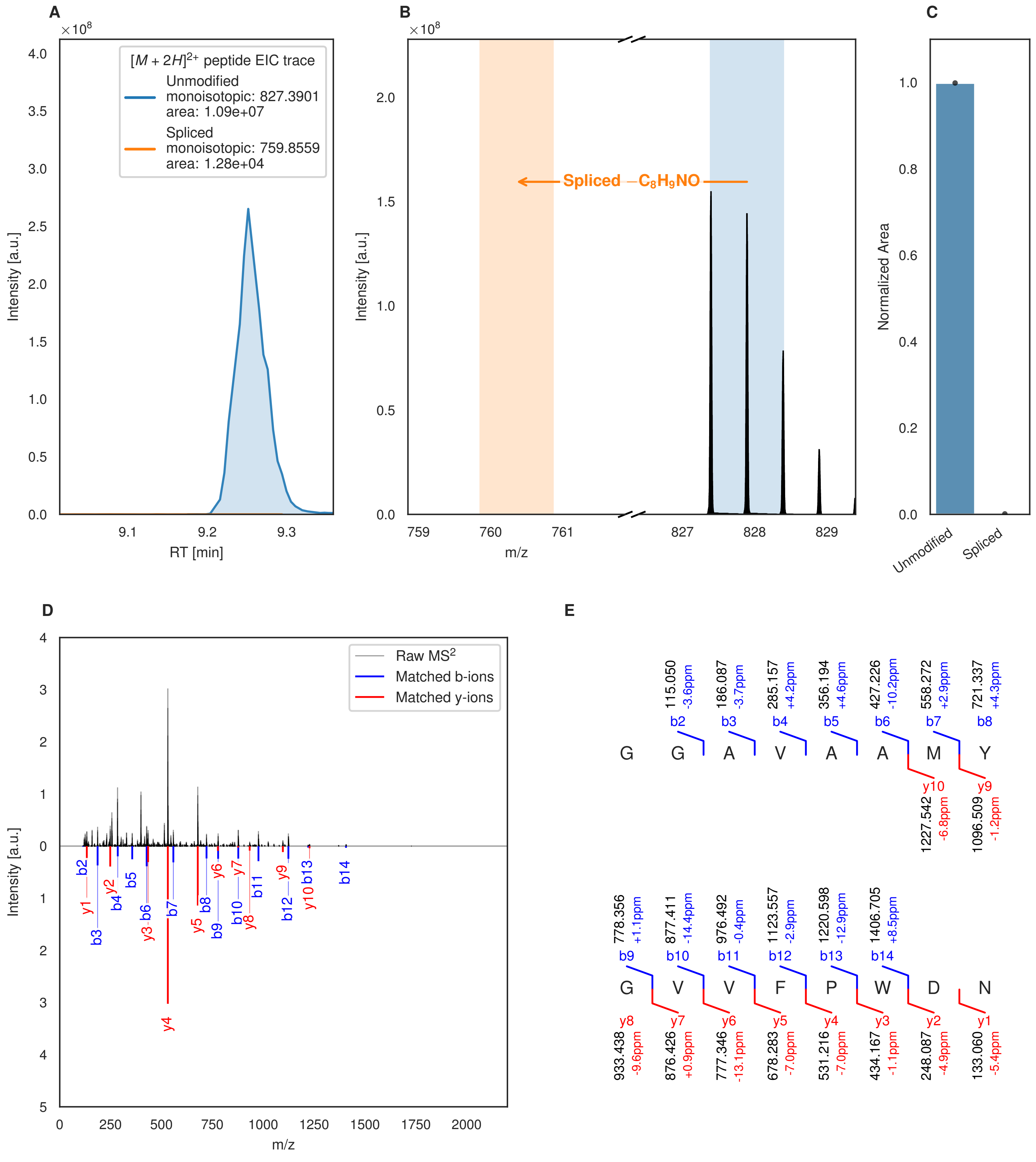
A
[M + 2H]2+ peptide EIC trace
Unmodified
monoisotopic: 827.3901
area: 1.09e+07
Spliced
monoisotopic: 759.8559
area: 1.28e+04
Intensity [a.u.]
RT [min]
B
Spliced −C8H9NO
Intensity [a.u.]
m/z
C
Normalized Area
Unmodified
Spliced
D
Raw MS2
Matched b-ions
Matched y-ions
Intensity [a.u.]
m/z
E
G G A V A A M Y
G V V F P W D N

Figure S36 (figure continued from the previous page).

**F**

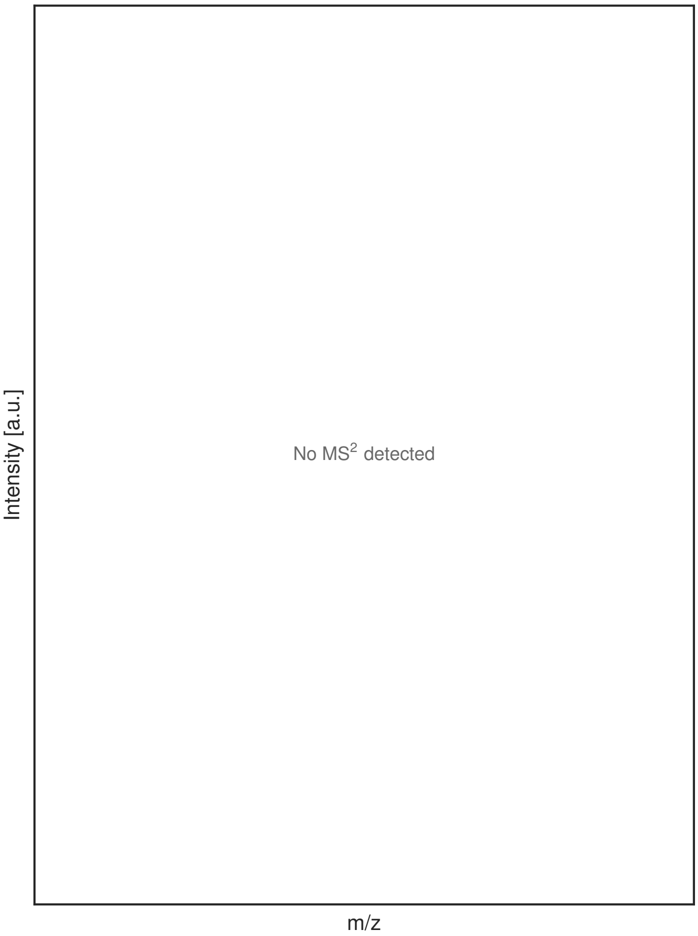

**G**

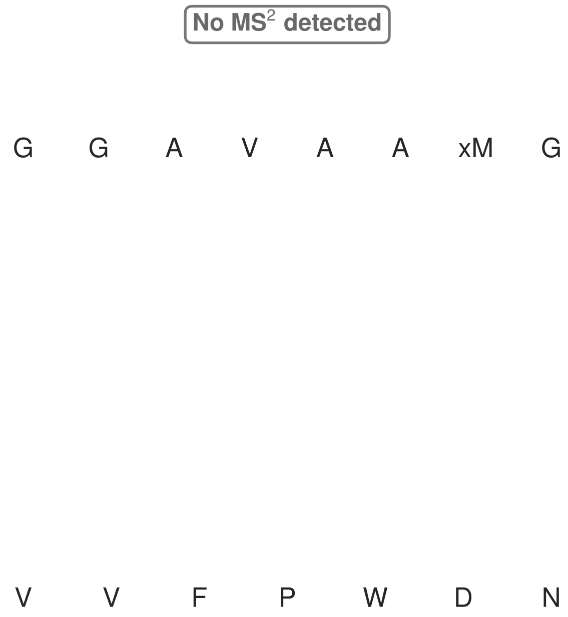

**Figure S36: Modification of the TJ-C-676 precursor peptide by HypXY.** LC-MS analysis of IMAC-purified precursor peptides from *E. coli* BL21(DE3)Tuner containing splicease-encoding plasmid TJ-C-560 (HypXY) and precursor peptide-encoding plasmid TJ-C-676 (Fh8-HypAc-TEV-PlpA3-Trunc) after expression at 30 °C for 22 h in 30 mL cultures after digestion with TEV. Data from n=1 independent biological replicates. Spliced amino acids have an x as a prefix. Spliced refers to a single tyramine excision. A representative experimental replicate is shown in panels **A, B.**
**A** Close-up plot of the representative extracted ion chromatogram (EIC) of the various peptide states; the blue trace corresponds to the Unmodified peptide; the orange trace corresponds to the Spliced peptide. **B** Close-up plot of the m/z values of the peptides detected by the mass spectrometer, with the shaded colored areas corresponding to where the expected m/z values for the peptide m/z traces should occur, with the observed spectrum plotted in black. The Unmodified, Spliced peptide(s) were found in the mass spectrum. **C** Bar plot showing the relative abundance of the various peptide states at charge z=2; the bars represent the average relative abundances normalized to total peak areas, with each individual experimental replicate value plotted as a black dot. **D** Mirrored $MS^2$ plot for the Unmodified peptide. In this mirror plot, the full fragmentation spectrum is plotted in black pointing upward, and the matched fragment b ions (blue) and y ions (red) are plotted pointing downward, mirrored against it, so that each matched fragment can be compared directly with the measured spectrum. **E** $MS^2$ spectrum of the fragments corresponding to the Unmodified peptide sequence, with matched b (blue, above) and y ions (red, below) plotted. **F** Mirrored $MS^2$ plot for the Spliced peptide. In this mirror plot, the full fragmentation spectrum is plotted in black pointing upward, and the matched fragment b ions (blue) and y ions (red) are plotted pointing downward, mirrored against it, so that each matched fragment can be compared directly with the measured spectrum. **G** $MS^2$ spectrum of the fragments corresponding to the Spliced peptide sequence, with matched b (blue, above) and y ions (red, below) plotted.

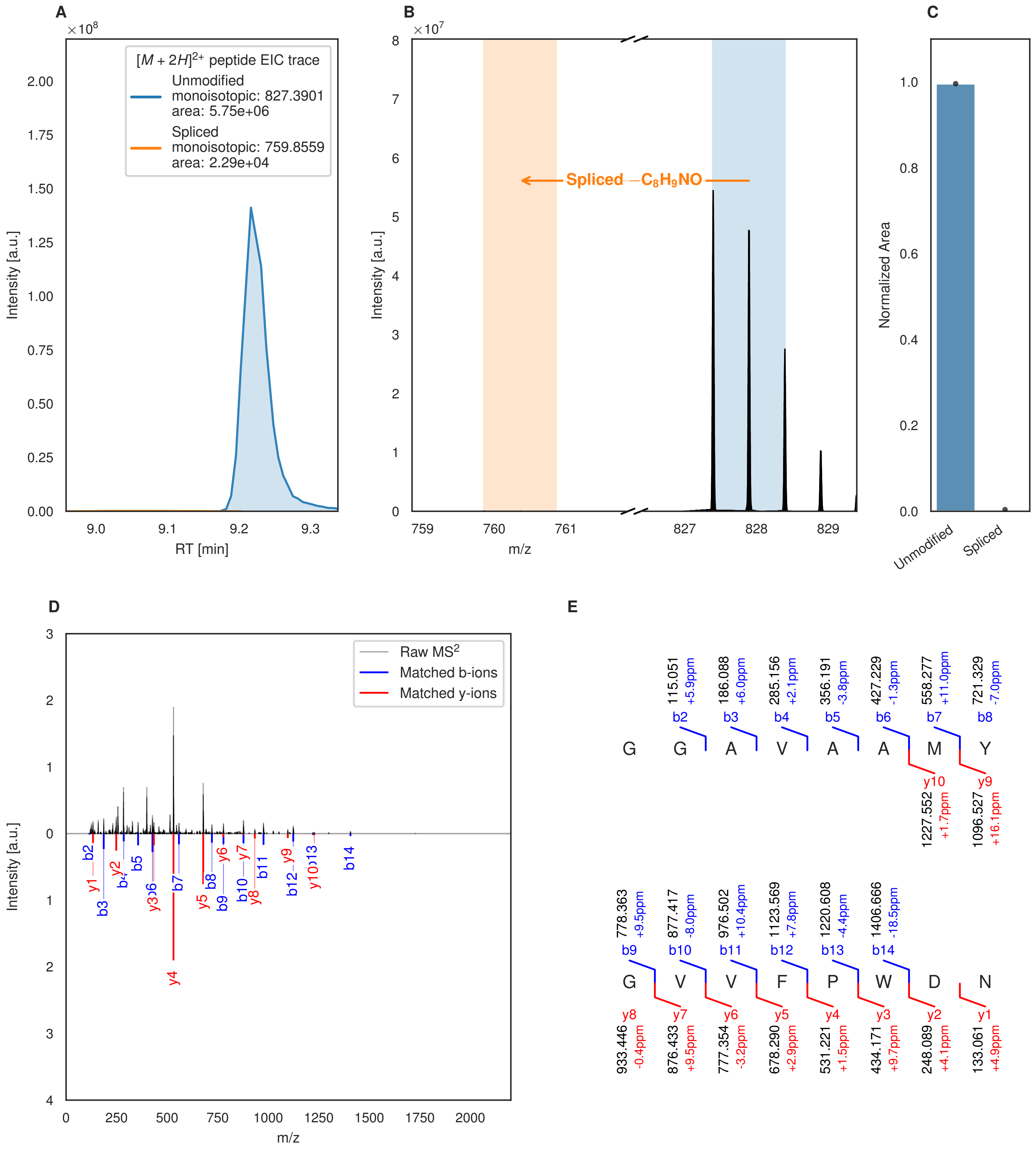
A
B
C
D
E
$[M+2H]^{2+}$ peptide EIC trace
Unmodified
monoisotopic: 827.3901
area: 5.75e+06
Spliced
monoisotopic: 759.8559
area: 2.29e+04
Intensity [a.u.]
RT [min]
Spliced $-C_8H_9NO$
m/z
Normalized Area
Unmodified
Spliced
Raw MS²
Matched b-ions
Matched y-ions
G G A V A A M Y
G V V F P W D N
b2 115.051 +5.9ppm
b3 186.088 +6.0ppm
b4 285.156 +2.1ppm
b5 356.191 -3.8ppm
b6 427.229 -1.3ppm
b7 558.277 +11.0ppm
b8 721.329 -7.0ppm
y10 1227.552 +1.7ppm
y9 1096.527 +16.1ppm
b9 778.363 +9.5ppm
b10 877.417 -8.0ppm
b11 976.502 +10.4ppm
b12 1123.569 +7.8ppm
b13 1220.608 -4.4ppm
b14 1406.666 -18.5ppm
y8 933.446 -0.4ppm
y7 876.433 +9.5ppm
y6 777.354 -3.2ppm
y5 678.290 +2.9ppm
y4 531.221 +1.5ppm
y3 434.171 +9.7ppm
y2 248.089 +4.1ppm
y1 133.061 +4.9ppm

Figure S37 (figure continued from the previous page).

F

G

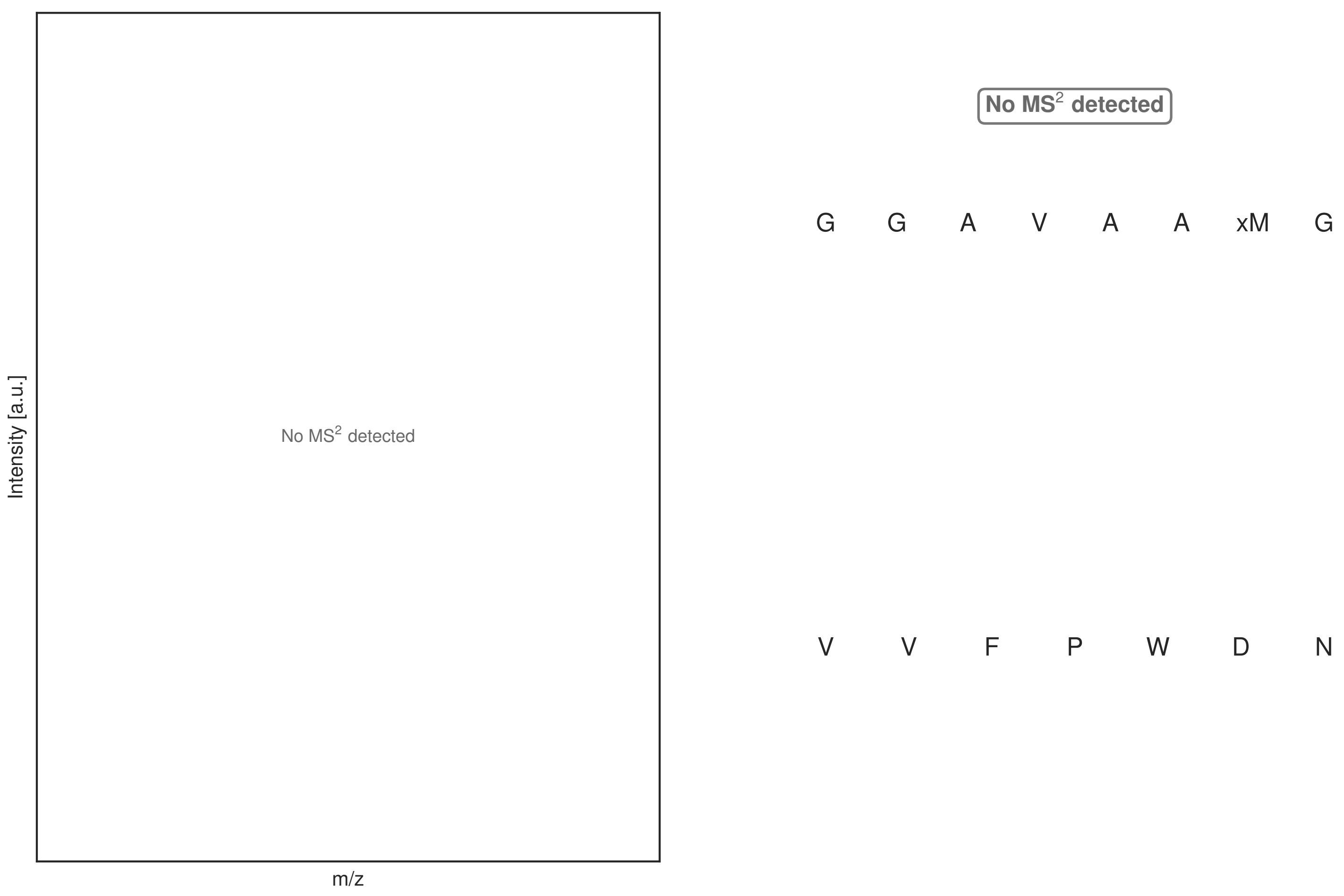

**Figure S37: Modification of the TJ-C-676 precursor peptide by HypXY.** LC-MS analysis of IMAC-purified precursor peptides from *E. coli* BL21(DE3)Tuner containing splicease-encoding plasmid TJ-C-689 (HypXY) and precursor peptide-encoding plasmid TJ-C-676 (Fh8-HypAc-TEV-PlpA3-Trunc) after expression at 24 °C for 22 h in 30 mL cultures after digestion with TEV. Data from n=1 independent biological replicates. Spliced amino acids have an x as a prefix. Spliced refers to a single tyramine excision. A representative experimental replicate is shown in panels **A, B.**
**A** Close-up plot of the representative extracted ion chromatogram (EIC) of the various peptide states; the blue trace corresponds to the Unmodified peptide; the orange trace corresponds to the Spliced peptide. **B** Close-up plot of the m/z values of the peptides detected by the mass spectrometer, with the shaded colored areas corresponding to where the expected m/z values for the peptide m/z traces should occur, with the observed spectrum plotted in black. The Unmodified, Spliced peptide(s) were found in the mass spectrum. **C** Bar plot showing the relative abundance of the various peptide states at charge z=2; the bars represent the average relative abundances normalized to total peak areas, with each individual experimental replicate value plotted as a black dot. **D** Mirrored $MS^2$ plot for the Unmodified peptide. In this mirror plot, the full fragmentation spectrum is plotted in black pointing upward, and the matched fragment b ions (blue) and y ions (red) are plotted pointing downward, mirrored against it, so that each matched fragment can be compared directly with the measured spectrum. **E** $MS^2$ spectrum of the fragments corresponding to the Unmodified peptide sequence, with matched b (blue, above) and y ions (red, below) plotted. **F** Mirrored $MS^2$ plot for the Spliced peptide. In this mirror plot, the full fragmentation spectrum is plotted in black pointing upward, and the matched fragment b ions (blue) and y ions (red) are plotted pointing downward, mirrored against it, so that each matched fragment can be compared directly with the measured spectrum. **G** $MS^2$ spectrum of the fragments corresponding to the Spliced peptide sequence, with matched b (blue, above) and y ions (red, below) plotted.

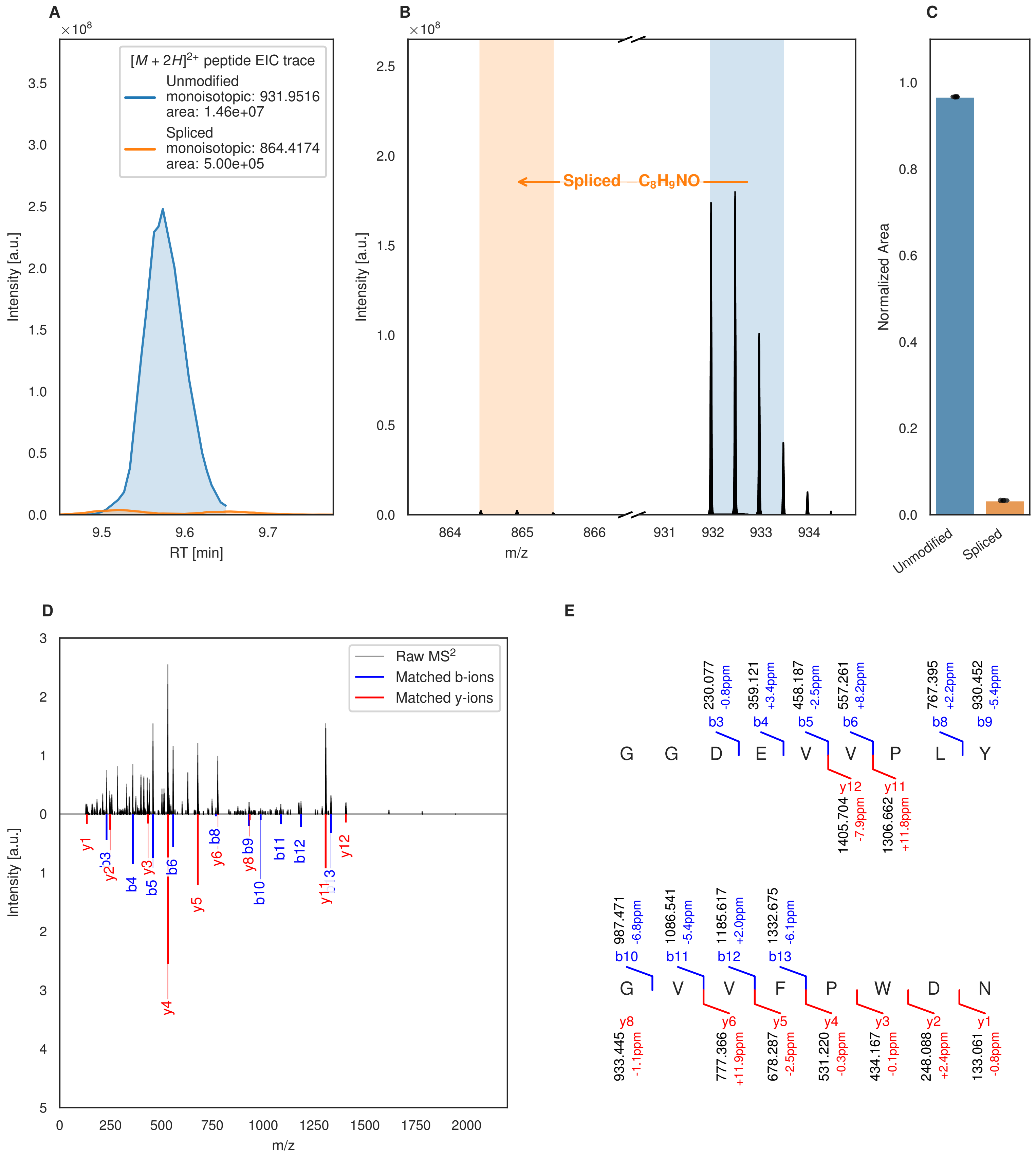

A
×10^8
[M + 2H]2+ peptide EIC trace
Unmodified
monoisotopic: 931.9516
area: 1.46e+07
Spliced
monoisotopic: 864.4174
area: 5.00e+05
Intensity [a.u.]
RT [min]
B
×10^8
Spliced −C8H9NO
Intensity [a.u.]
m/z
C
Normalized Area
Unmodified
Spliced
D
Raw MS2
Matched b-ions
Matched y-ions
Intensity [a.u.]
m/z
E
230.077 -0.8ppm b3
359.121 +3.4ppm b4
458.187 -2.5ppm b5
557.261 +8.2ppm b6
767.395 +2.2ppm b8
930.452 -5.4ppm b9
G G D E V V P L Y
y12 1405.704 -7.9ppm
y11 1306.662 +11.8ppm
987.471 -6.8ppm b10
1086.541 -5.4ppm b11
1185.617 +2.0ppm b12
1332.675 -6.1ppm b13
G V V F P W D N
y8 933.445 -1.1ppm
y6 777.366 +11.9ppm
y5 678.287 -2.5ppm
y4 531.220 -0.3ppm
y3 434.167 -0.1ppm
y2 248.088 +2.4ppm
y1 133.061 -0.8ppm

Figure S38 (figure continued from the previous page).

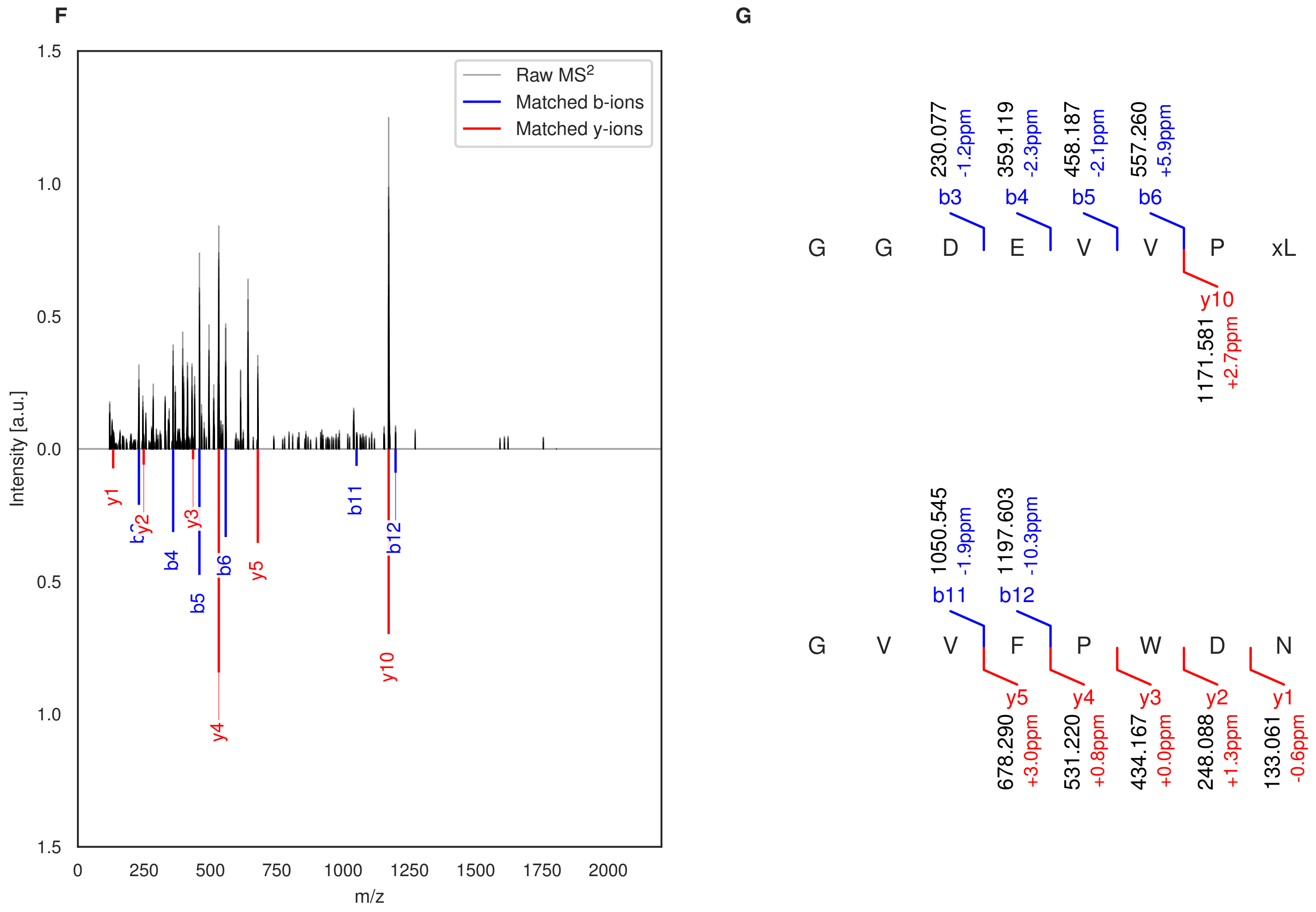

**Figure S38: Modification of the TJ-C-677 precursor peptide by HypXY.** LC-MS analysis of IMAC-purified precursor peptides from *E. coli* BL21(DE3)Tuner containing splicease-encoding plasmid TJ-C-560 (HypXY) and precursor peptide-encoding plasmid TJ-C-677 (RSTN1-HypAc-TEV-PlpA3-Boc-Trunc) after expression at 24 °C for 22 h in 30 mL cultures after digestion with TEV. Data from n=2 independent biological replicates. Spliced amino acids have an x as a prefix. Spliced refers to a single tyramine excision. A representative experimental replicate is shown in panels **A, B.**

**A** Close-up plot of the representative extracted ion chromatogram (EIC) of the various peptide states; the blue trace corresponds to the Unmodified peptide; the orange trace corresponds to the Spliced peptide. **B** Close-up plot of the m/z values of the peptides detected by the mass spectrometer, with the shaded colored areas corresponding to where the expected m/z values for the peptide m/z traces should occur, with the observed spectrum plotted in black. The Unmodified, Spliced peptide(s) were found in the mass spectrum. **C** Bar plot showing the relative abundance of the various peptide states at charge z=2; the bars represent the average relative abundances normalized to total peak areas, with each individual experimental replicate value plotted as a black dot. **D** Mirrored $MS^2$ plot for the Unmodified peptide. In this mirror plot, the full fragmentation spectrum is plotted in black pointing upward, and the matched fragment b ions (blue) and y ions (red) are plotted pointing downward, mirrored against it, so that each matched fragment can be compared directly with the measured spectrum. **E** $MS^2$ spectrum of the fragments corresponding to the Unmodified peptide sequence, with matched b (blue, above) and y ions (red, below) plotted. **F** Mirrored $MS^2$ plot for the Spliced peptide. In this mirror plot, the full fragmentation spectrum is plotted in black pointing upward, and the matched fragment b ions (blue) and y ions (red) are plotted pointing downward, mirrored against it, so that each matched fragment can be compared directly with the measured spectrum. **G** $MS^2$ spectrum of the fragments corresponding to the Spliced peptide sequence, with matched b (blue, above) and y ions (red, below) plotted.

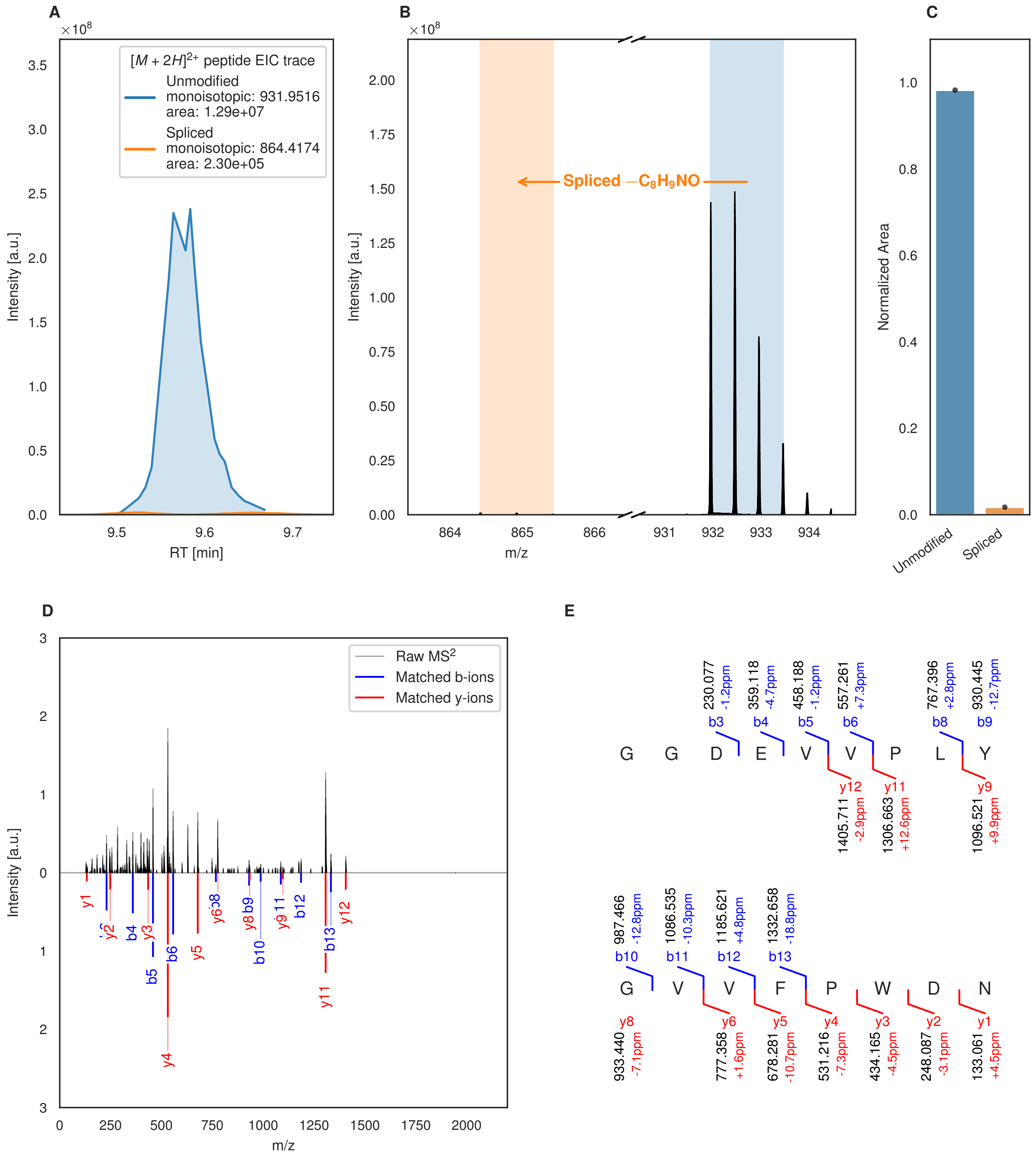
A
B
C
D
E
$[M+2H]^{2+}$ peptide EIC trace
Unmodified
monoisotopic: 931.9516
area: 1.29e+07
Spliced
monoisotopic: 864.4174
area: 2.30e+05
Intensity [a.u.]
RT [min]
Spliced $-C_8H_9NO$
m/z
Normalized Area
Unmodified
Spliced
Raw $MS^2$
Matched b-ions
Matched y-ions
G G D E V V P L Y
G V V F P W D N

Figure S39 (figure continued from the previous page).

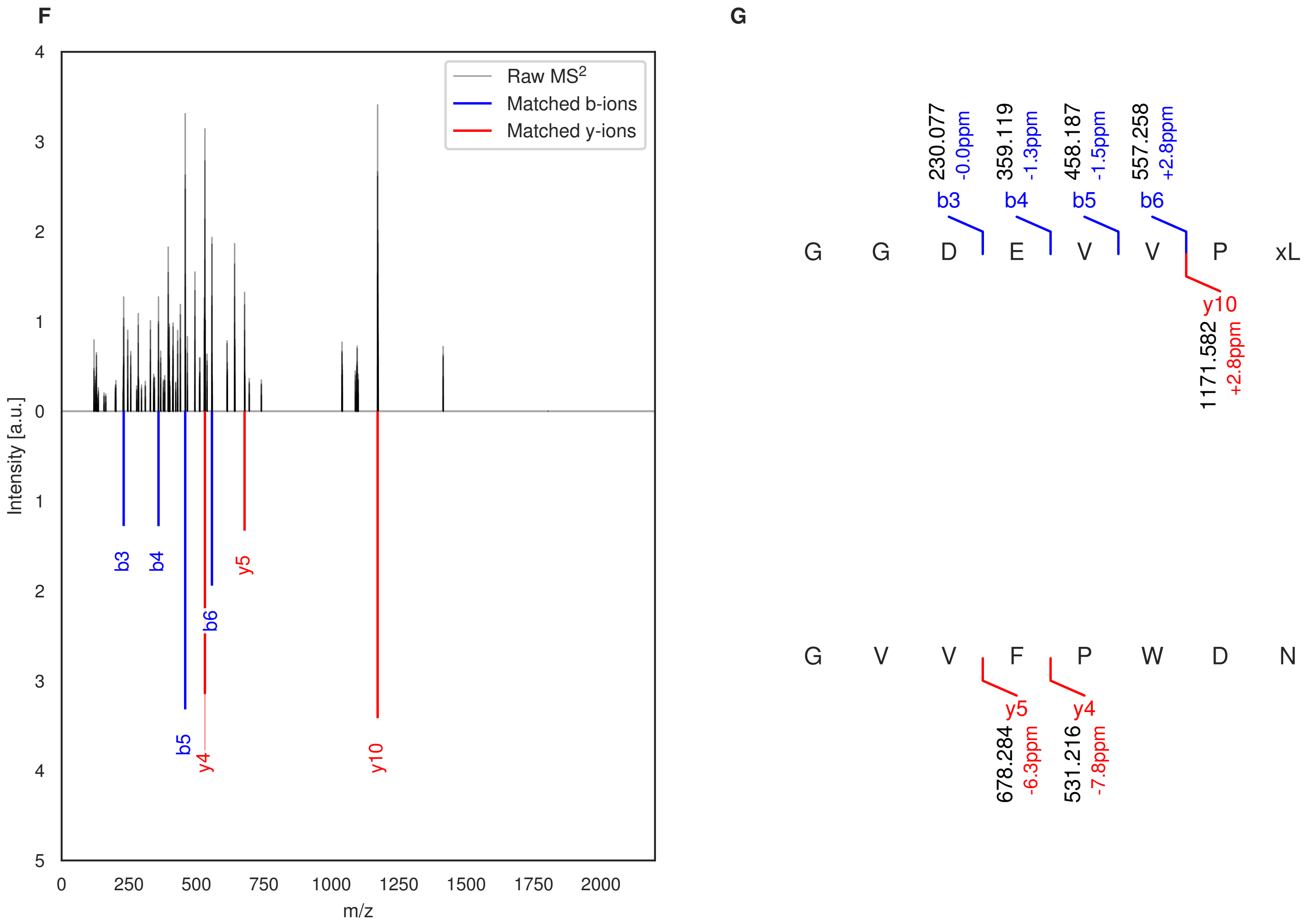

**Figure S39: Modification of the TJ-C-677 precursor peptide by HypXY.** LC-MS analysis of IMAC-purified precursor peptides from *E. coli* BL21(DE3)Tuner containing splicease-encoding plasmid TJ-C-560 (HypXY) and precursor peptide-encoding plasmid TJ-C-677 (RSTN1-HypAc-TEV-PlpA3-Boc-Trunc) after expression at 30 °C for 22 h in 30 mL cultures after digestion with TEV. Data from n=1 independent biological replicates. Spliced amino acids have an x as a prefix. Spliced refers to a single tyramine excision. A representative experimental replicate is shown in panels **A, B.**

**A** Close-up plot of the representative extracted ion chromatogram (EIC) of the various peptide states; the blue trace corresponds to the Unmodified peptide; the orange trace corresponds to the Spliced peptide. **B** Close-up plot of the m/z values of the peptides detected by the mass spectrometer, with the shaded colored areas corresponding to where the expected m/z values for the peptide m/z traces should occur, with the observed spectrum plotted in black. The Unmodified, Spliced peptide(s) were found in the mass spectrum. **C** Bar plot showing the relative abundance of the various peptide states at charge z=2; the bars represent the average relative abundances normalized to total peak areas, with each individual experimental replicate value plotted as a black dot. **D** Mirrored MS$^2$ plot for the Unmodified peptide. In this mirror plot, the full fragmentation spectrum is plotted in black pointing upward, and the matched fragment b ions (blue) and y ions (red) are plotted pointing downward, mirrored against it, so that each matched fragment can be compared directly with the measured spectrum. **E** MS$^2$ spectrum of the fragments corresponding to the Unmodified peptide sequence, with matched b (blue, above) and y ions (red, below) plotted. **F** Mirrored MS$^2$ plot for the Spliced peptide. In this mirror plot, the full fragmentation spectrum is plotted in black pointing upward, and the matched fragment b ions (blue) and y ions (red) are plotted pointing downward, mirrored against it, so that each matched fragment can be compared directly with the measured spectrum. **G** MS$^2$ spectrum of the fragments corresponding to the Spliced peptide sequence, with matched b (blue, above) and y ions (red, below) plotted.

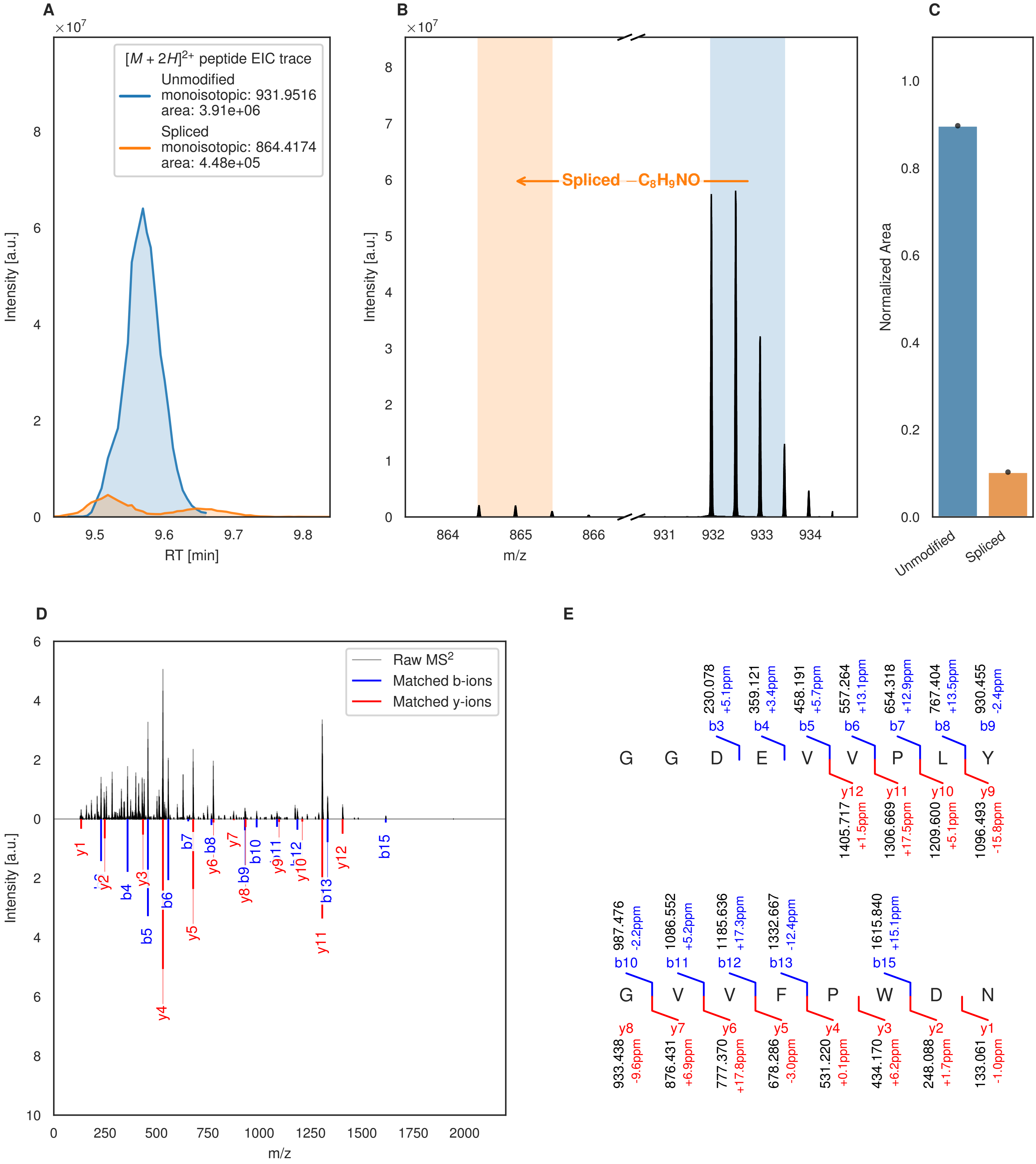
A
×10^7
[M + 2H]^2+ peptide EIC trace
Unmodified
monoisotopic: 931.9516
area: 3.91e+06
Spliced
monoisotopic: 864.4174
area: 4.48e+05
Intensity [a.u.]
RT [min]
B
×10^7
Spliced −C8H9NO
Intensity [a.u.]
m/z
C
Normalized Area
Unmodified
Spliced
D
Raw MS2
Matched b-ions
Matched y-ions
Intensity [a.u.]
m/z
E
G G D E V V P L Y
b3 230.078 +5.1ppm
b4 359.121 +3.4ppm
b5 458.191 +5.7ppm
b6 557.264 +13.1ppm
b7 654.318 +12.9ppm
b8 767.404 +13.5ppm
b9 930.455 -2.4ppm
y12 1405.717 +1.5ppm
y11 1306.669 +17.5ppm
y10 1209.600 +5.1ppm
y9 1096.493 -15.8ppm
G V V F P W D N
b10 987.476 -2.2ppm
b11 1086.552 +5.2ppm
b12 1185.636 +17.3ppm
b13 1332.667 -12.4ppm
b15 1615.840 +15.1ppm
y8 933.438 -9.6ppm
y7 876.431 +6.9ppm
y6 777.370 +17.8ppm
y5 678.286 -3.0ppm
y4 531.220 +0.1ppm
y3 434.170 +6.2ppm
y2 248.088 +1.7ppm
y1 133.061 -1.0ppm

Figure S40 (figure continued from the previous page).

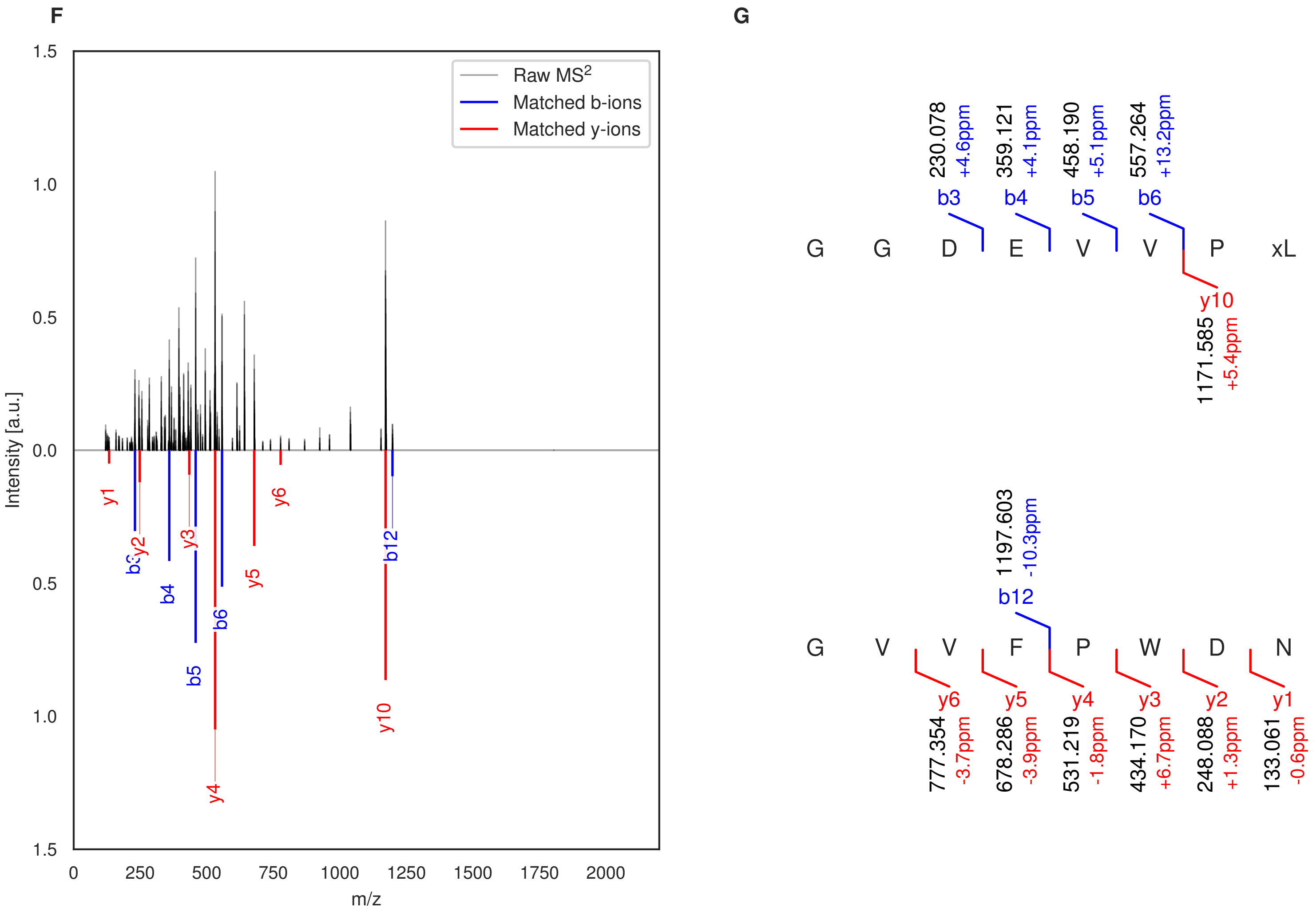

**Figure S40: Modification of the TJ-C-678 precursor peptide by HypXY.** LC-MS analysis of IMAC-purified precursor peptides from *E. coli* BL21(DE3)Tuner containing splicease-encoding plasmid TJ-C-560 (HypXY) and precursor peptide-encoding plasmid TJ-C-678 (Fh8-chaos-HypAc-TEV-PlpA3-Boc-Trunc) after expression at 24 °C for 22 h in 30 mL cultures after digestion with TEV. Data from n=1 independent biological replicates. Spliced amino acids have an x as a prefix. Spliced refers to a single tyramine excision. A representative experimental replicate is shown in panels **A, B.**

**A** Close-up plot of the representative extracted ion chromatogram (EIC) of the various peptide states; the blue trace corresponds to the Unmodified peptide; the orange trace corresponds to the Spliced peptide. **B** Close-up plot of the m/z values of the peptides detected by the mass spectrometer, with the shaded colored areas corresponding to where the expected m/z values for the peptide m/z traces should occur, with the observed spectrum plotted in black. The Unmodified, Spliced peptide(s) were found in the mass spectrum. **C** Bar plot showing the relative abundance of the various peptide states at charge z=2; the bars represent the average relative abundances normalized to total peak areas, with each individual experimental replicate value plotted as a black dot. **D** Mirrored $MS^2$ plot for the Unmodified peptide. In this mirror plot, the full fragmentation spectrum is plotted in black pointing upward, and the matched fragment b ions (blue) and y ions (red) are plotted pointing downward, mirrored against it, so that each matched fragment can be compared directly with the measured spectrum. **E** $MS^2$ spectrum of the fragments corresponding to the Unmodified peptide sequence, with matched b (blue, above) and y ions (red, below) plotted. **F** Mirrored $MS^2$ plot for the Spliced peptide. In this mirror plot, the full fragmentation spectrum is plotted in black pointing upward, and the matched fragment b ions (blue) and y ions (red) are plotted pointing downward, mirrored against it, so that each matched fragment can be compared directly with the measured spectrum. **G** $MS^2$ spectrum of the fragments corresponding to the Spliced peptide sequence, with matched b (blue, above) and y ions (red, below) plotted.

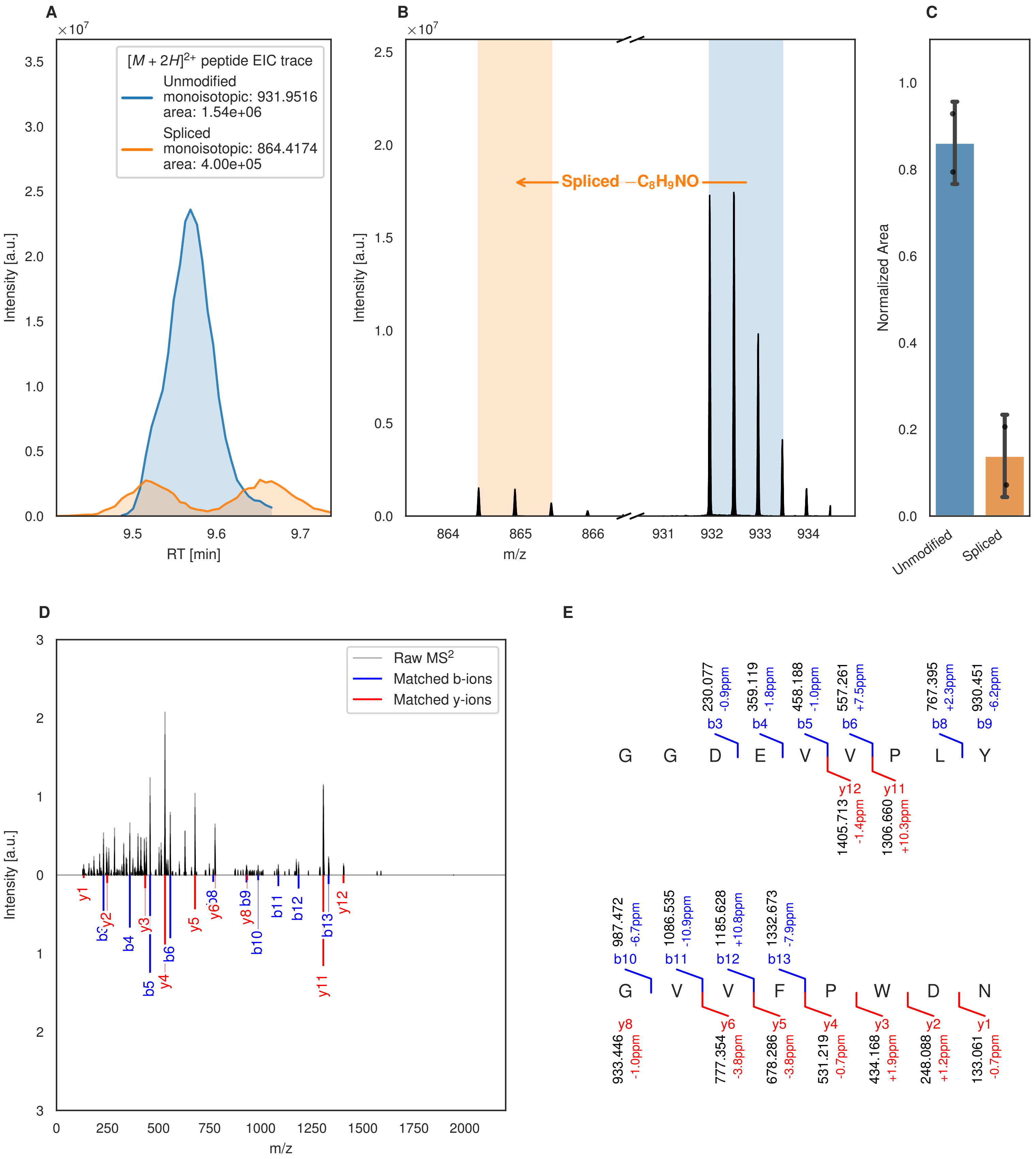

A
B
C
D
E
[M + 2H]2+ peptide EIC trace
Unmodified
monoisotopic: 931.9516
area: 1.54e+06
Spliced
monoisotopic: 864.4174
area: 4.00e+05
Intensity [a.u.]
RT [min]
Spliced −C8H9NO
m/z
Normalized Area
Unmodified
Spliced
Raw MS2
Matched b-ions
Matched y-ions
G G D E V V P L Y
G V V F P W D N

Figure S41 (figure continued from the previous page).

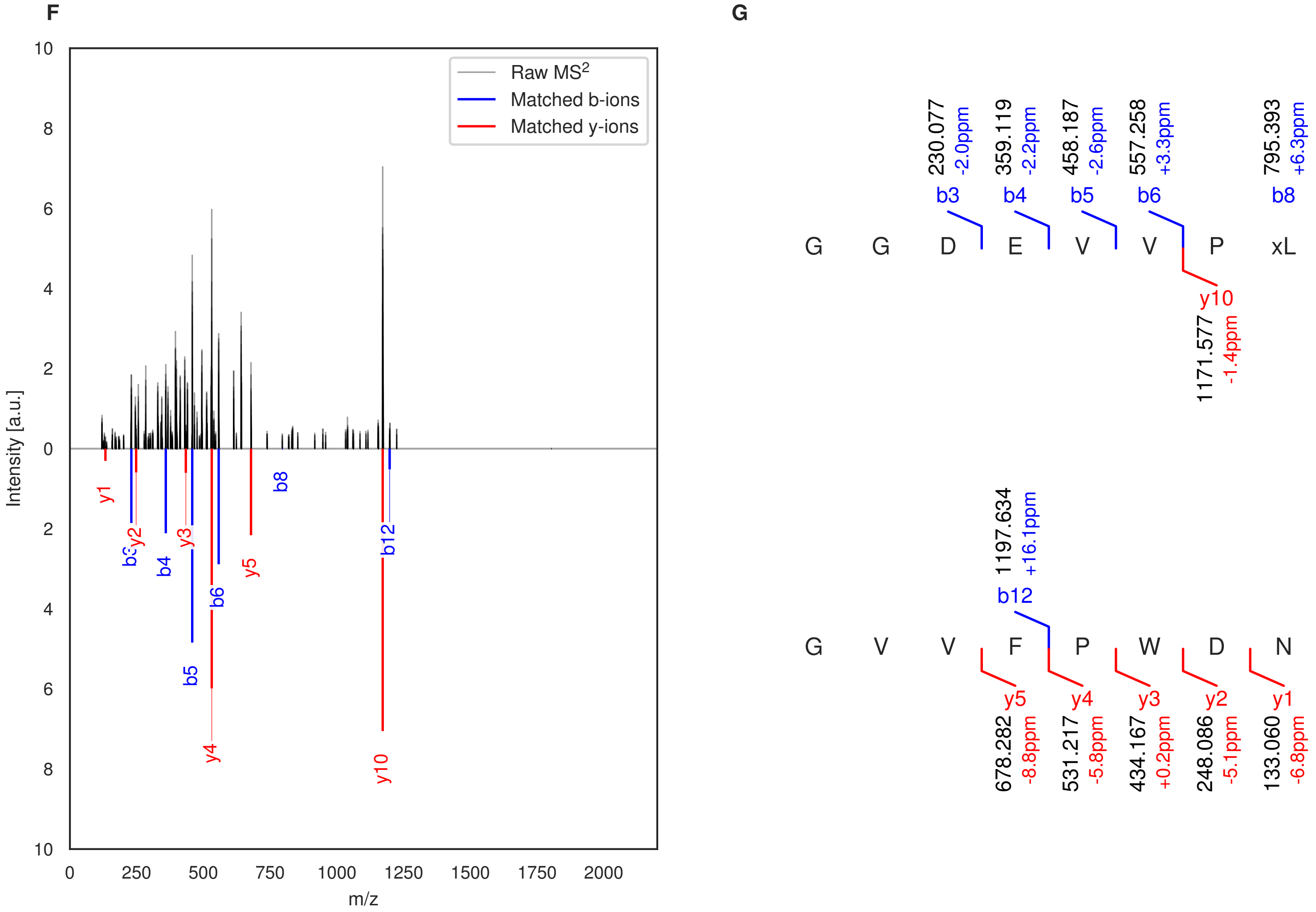

**Figure S41: Modification of the TJ-C-679 precursor peptide by HypXY.** LC-MS analysis of IMAC-purified precursor peptides from *E. coli* BL21(DE3)Tuner containing splicease-encoding plasmid TJ-C-560 (HypXY) and precursor peptide-encoding plasmid TJ-C-679 (Fh8-HypAc-TEV-PlpA3-Boc-Trunc) after expression at 24 °C for 22 h in 30 mL cultures after digestion with TEV. Data from n=2 independent biological replicates. Spliced amino acids have an x as a prefix. Spliced refers to a single tyramine excision. A representative experimental replicate is shown in panels **A, B.**

**A** Close-up plot of the representative extracted ion chromatogram (EIC) of the various peptide states; the blue trace corresponds to the Unmodified peptide; the orange trace corresponds to the Spliced peptide. **B** Close-up plot of the m/z values of the peptides detected by the mass spectrometer, with the shaded colored areas corresponding to where the expected m/z values for the peptide m/z traces should occur, with the observed spectrum plotted in black. The Unmodified, Spliced peptide(s) were found in the mass spectrum. **C** Bar plot showing the relative abundance of the various peptide states at charge z=2; the bars represent the average relative abundances normalized to total peak areas, with each individual experimental replicate value plotted as a black dot. **D** Mirrored $MS^2$ plot for the Unmodified peptide. In this mirror plot, the full fragmentation spectrum is plotted in black pointing upward, and the matched fragment b ions (blue) and y ions (red) are plotted pointing downward, mirrored against it, so that each matched fragment can be compared directly with the measured spectrum. **E** $MS^2$ spectrum of the fragments corresponding to the Unmodified peptide sequence, with matched b (blue, above) and y ions (red, below) plotted. **F** Mirrored $MS^2$ plot for the Spliced peptide. In this mirror plot, the full fragmentation spectrum is plotted in black pointing upward, and the matched fragment b ions (blue) and y ions (red) are plotted pointing downward, mirrored against it, so that each matched fragment can be compared directly with the measured spectrum. **G** $MS^2$ spectrum of the fragments corresponding to the Spliced peptide sequence, with matched b (blue, above) and y ions (red, below) plotted.

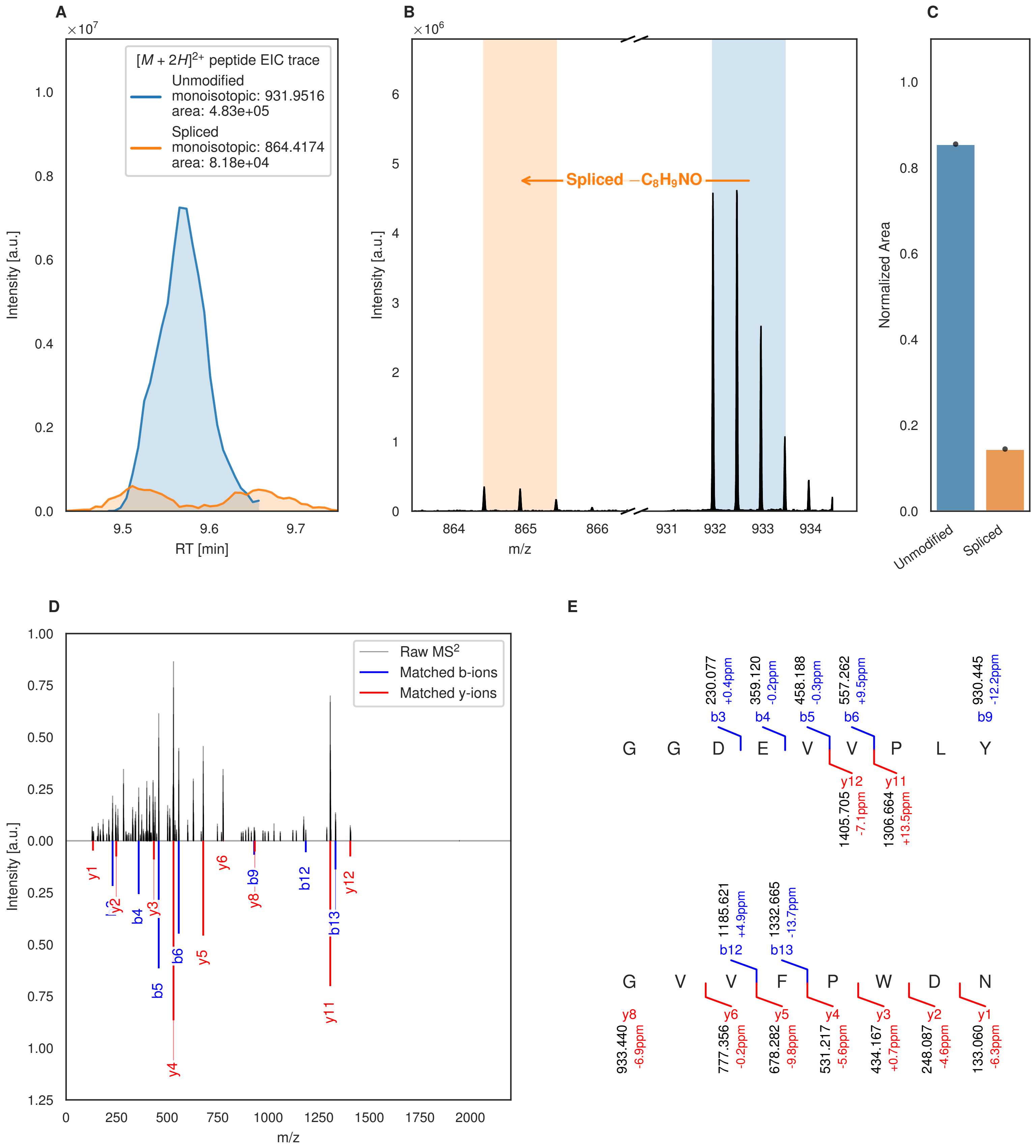

A
B
C
D
E
[M + 2H]2+ peptide EIC trace
Unmodified
monoisotopic: 931.9516
area: 4.83e+05
Spliced
monoisotopic: 864.4174
area: 8.18e+04
Intensity [a.u.]
RT [min]
Spliced −C8H9NO
m/z
Normalized Area
Unmodified
Spliced
Raw MS2
Matched b-ions
Matched y-ions
G G D E V V P L Y
230.077 +0.4ppm b3
359.120 -0.2ppm b4
458.188 -0.3ppm b5
557.262 +9.5ppm b6
930.445 -12.2ppm b9
y12 1405.705 -7.1ppm
y11 1306.664 +13.5ppm
G V V F P W D N
1185.621 +4.9ppm b12
1332.665 -13.7ppm b13
y8 933.440 -6.9ppm
y6 777.356 -0.2ppm
y5 678.282 -9.8ppm
y4 531.217 -5.6ppm
y3 434.167 +0.7ppm
y2 248.087 -4.6ppm
y1 133.060 -6.3ppm

Figure S42 (figure continued from the previous page).

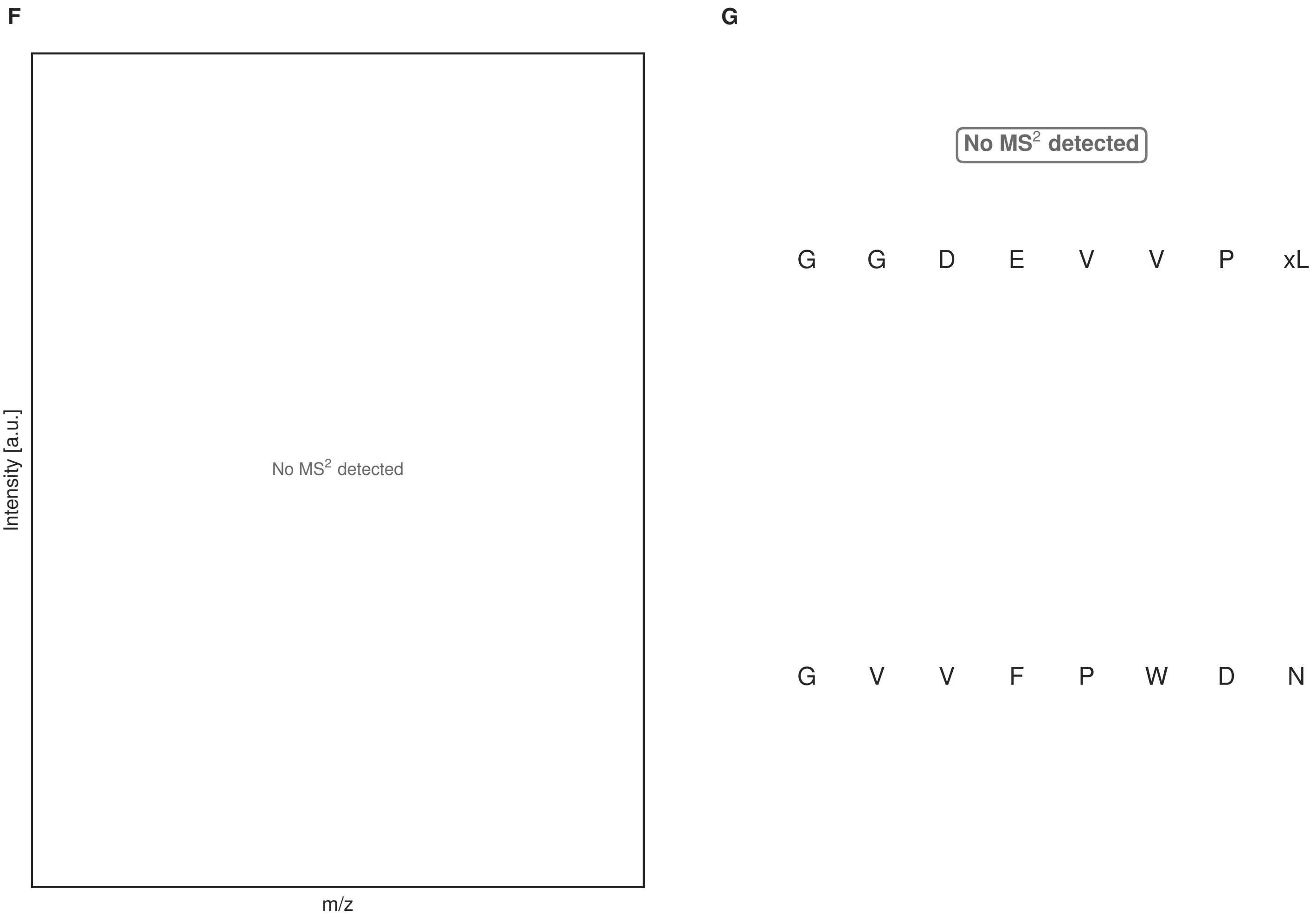

**Figure S42: Modification of the TJ-C-679 precursor peptide by HypXY.** LC-MS analysis of IMAC-purified precursor peptides from *E. coli* BL21(DE3)Tuner containing splicease-encoding plasmid TJ-C-560 (HypXY) and precursor peptide-encoding plasmid TJ-C-679 (Fh8-HypAc-TEV-PlpA3-Boc-Trunc) after expression at 30 °C for 22 h in 30 mL cultures after digestion with TEV. Data from n=1 independent biological replicates. Spliced amino acids have an x as a prefix. Spliced refers to a single tyramine excision. A representative experimental replicate is shown in panels **A, B.**

**A** Close-up plot of the representative extracted ion chromatogram (EIC) of the various peptide states; the blue trace corresponds to the Unmodified peptide; the orange trace corresponds to the Spliced peptide. **B** Close-up plot of the m/z values of the peptides detected by the mass spectrometer, with the shaded colored areas corresponding to where the expected m/z values for the peptide m/z traces should occur, with the observed spectrum plotted in black. The Unmodified, Spliced peptide(s) were found in the mass spectrum. **C** Bar plot showing the relative abundance of the various peptide states at charge z=2; the bars represent the average relative abundances normalized to total peak areas, with each individual experimental replicate value plotted as a black dot. **D** Mirrored $MS^2$ plot for the Unmodified peptide. In this mirror plot, the full fragmentation spectrum is plotted in black pointing upward, and the matched fragment b ions (blue) and y ions (red) are plotted pointing downward, mirrored against it, so that each matched fragment can be compared directly with the measured spectrum. **E** $MS^2$ spectrum of the fragments corresponding to the Unmodified peptide sequence, with matched b (blue, above) and y ions (red, below) plotted. **F** Mirrored $MS^2$ plot for the Spliced peptide. In this mirror plot, the full fragmentation spectrum is plotted in black pointing upward, and the matched fragment b ions (blue) and y ions (red) are plotted pointing downward, mirrored against it, so that each matched fragment can be compared directly with the measured spectrum. **G** $MS^2$ spectrum of the fragments corresponding to the Spliced peptide sequence, with matched b (blue, above) and y ions (red, below) plotted.

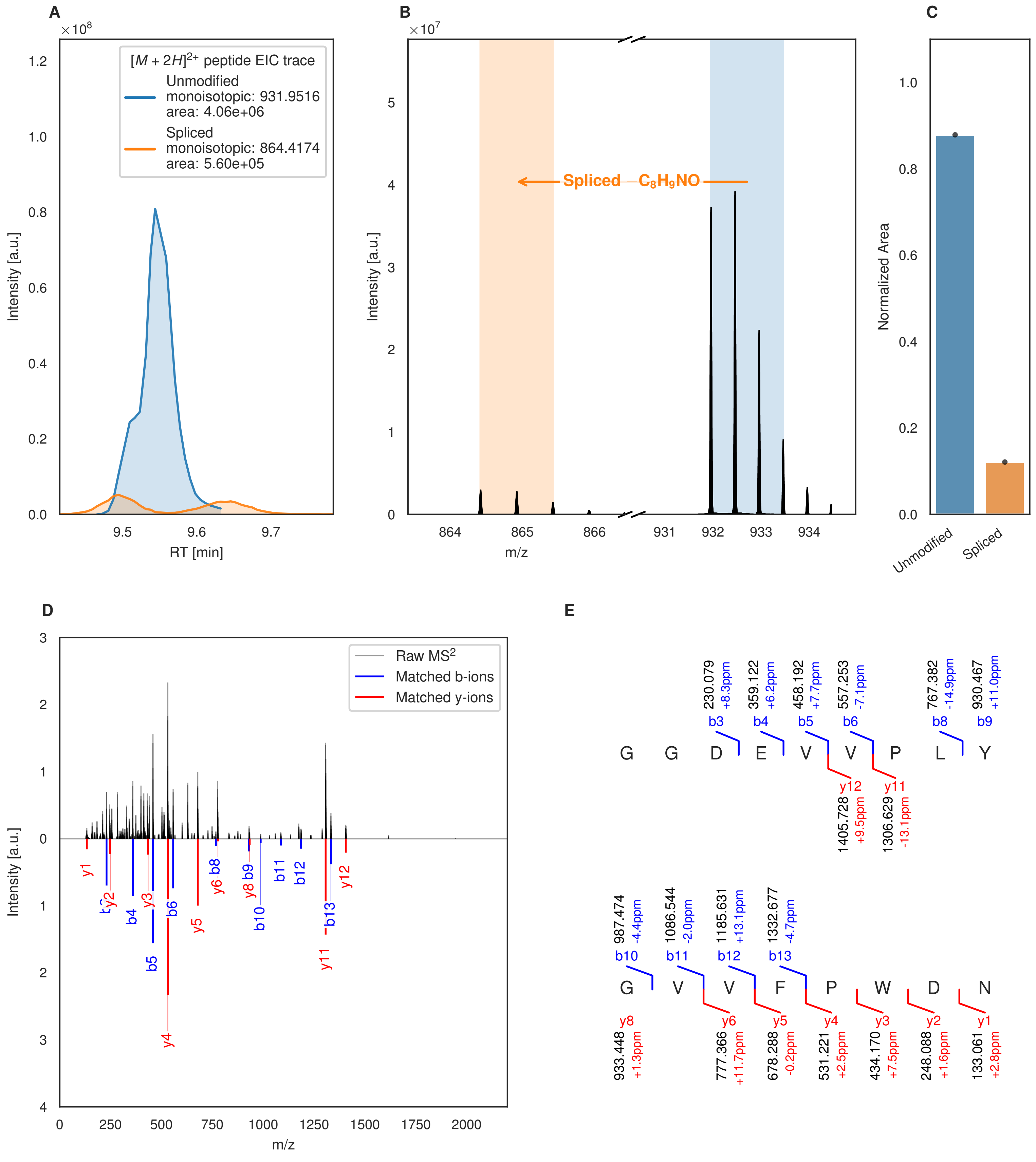
A
[M + 2H]2+ peptide EIC trace
Unmodified
monoisotopic: 931.9516
area: 4.06e+06
Spliced
monoisotopic: 864.4174
area: 5.60e+05
Intensity [a.u.]
RT [min]
B
Spliced −C8H9NO
Intensity [a.u.]
m/z
C
Normalized Area
Unmodified
Spliced
D
Raw MS2
Matched b-ions
Matched y-ions
Intensity [a.u.]
m/z
E
G G D E V V P L Y
G V V F P W D N

Figure S43 (figure continued from the previous page).

F

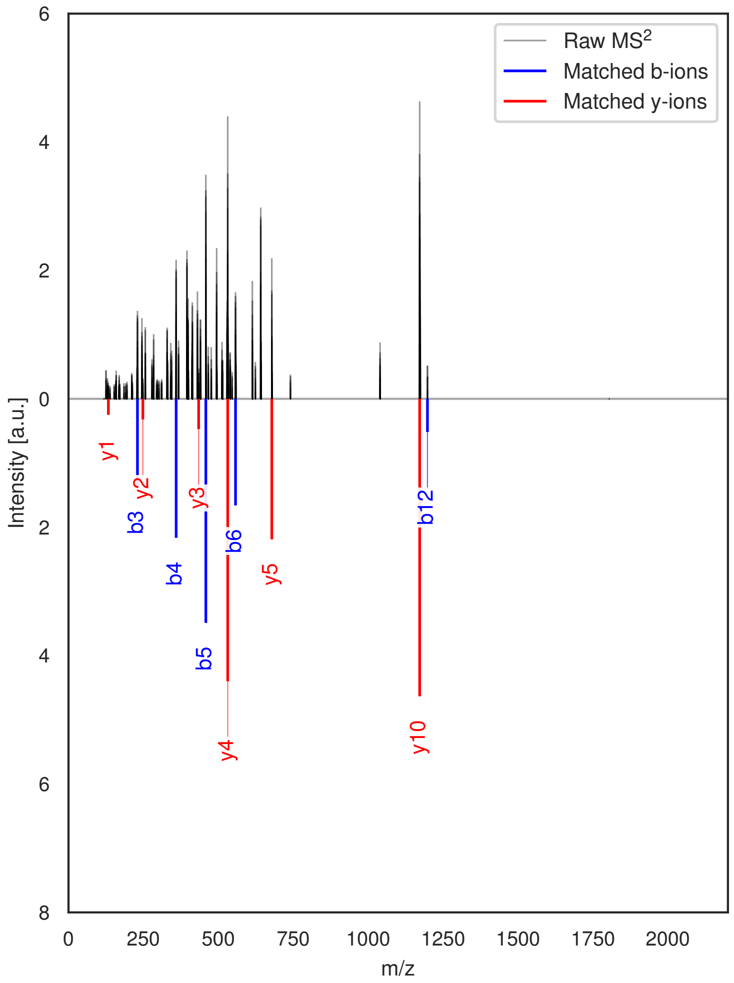

G

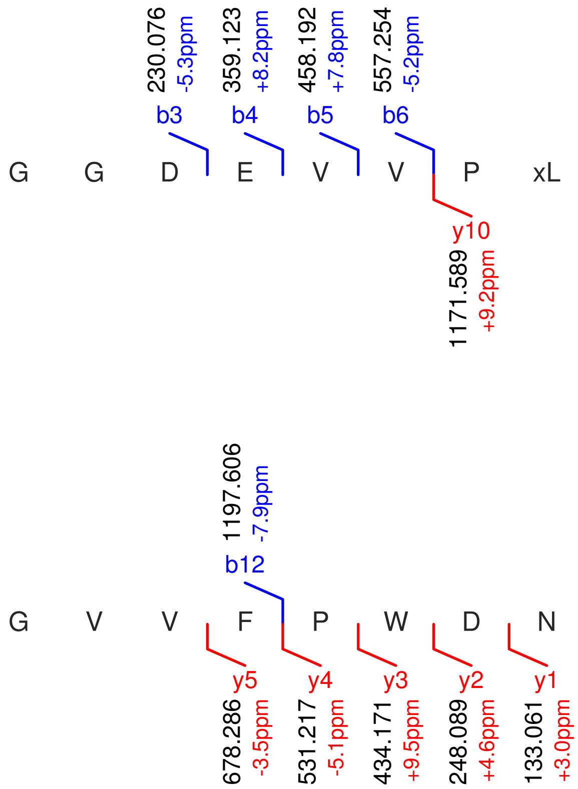

**Figure S43: Modification of the TJ-C-679 precursor peptide by HypXY.** LC-MS analysis of IMAC-purified precursor peptides from *E. coli* BL21(DE3)Tuner containing splicease-encoding plasmid TJ-C-689 (HypXY) and precursor peptide-encoding plasmid TJ-C-679 (Fh8-HypAc-TEV-PlpA3-Boc-Trunc) after expression at 24 °C for 22 h in 30 mL cultures after digestion with TEV. Data from n=5 independent biological replicates. Spliced amino acids have an x as a prefix. Spliced refers to a single tyramine excision. A representative experimental replicate is shown in panels **A, B.**

**A** Close-up plot of the representative extracted ion chromatogram (EIC) of the various peptide states; the blue trace corresponds to the Unmodified peptide; the orange trace corresponds to the Spliced peptide. **B** Close-up plot of the m/z values of the peptides detected by the mass spectrometer, with the shaded colored areas corresponding to where the expected m/z values for the peptide m/z traces should occur, with the observed spectrum plotted in black. The Unmodified, Spliced peptide(s) were found in the mass spectrum. **C** Bar plot showing the relative abundance of the various peptide states at charge z=2; the bars represent the average relative abundances normalized to total peak areas, with each individual experimental replicate value plotted as a black dot. **D** Mirrored MS$^2$ plot for the Unmodified peptide. In this mirror plot, the full fragmentation spectrum is plotted in black pointing upward, and the matched fragment b ions (blue) and y ions (red) are plotted pointing downward, mirrored against it, so that each matched fragment can be compared directly with the measured spectrum. **E** MS$^2$ spectrum of the fragments corresponding to the Unmodified peptide sequence, with matched b (blue, above) and y ions (red, below) plotted. **F** Mirrored MS$^2$ plot for the Spliced peptide. In this mirror plot, the full fragmentation spectrum is plotted in black pointing upward, and the matched fragment b ions (blue) and y ions (red) are plotted pointing downward, mirrored against it, so that each matched fragment can be compared directly with the measured spectrum. **G** MS$^2$ spectrum of the fragments corresponding to the Spliced peptide sequence, with matched b (blue, above) and y ions (red, below) plotted.

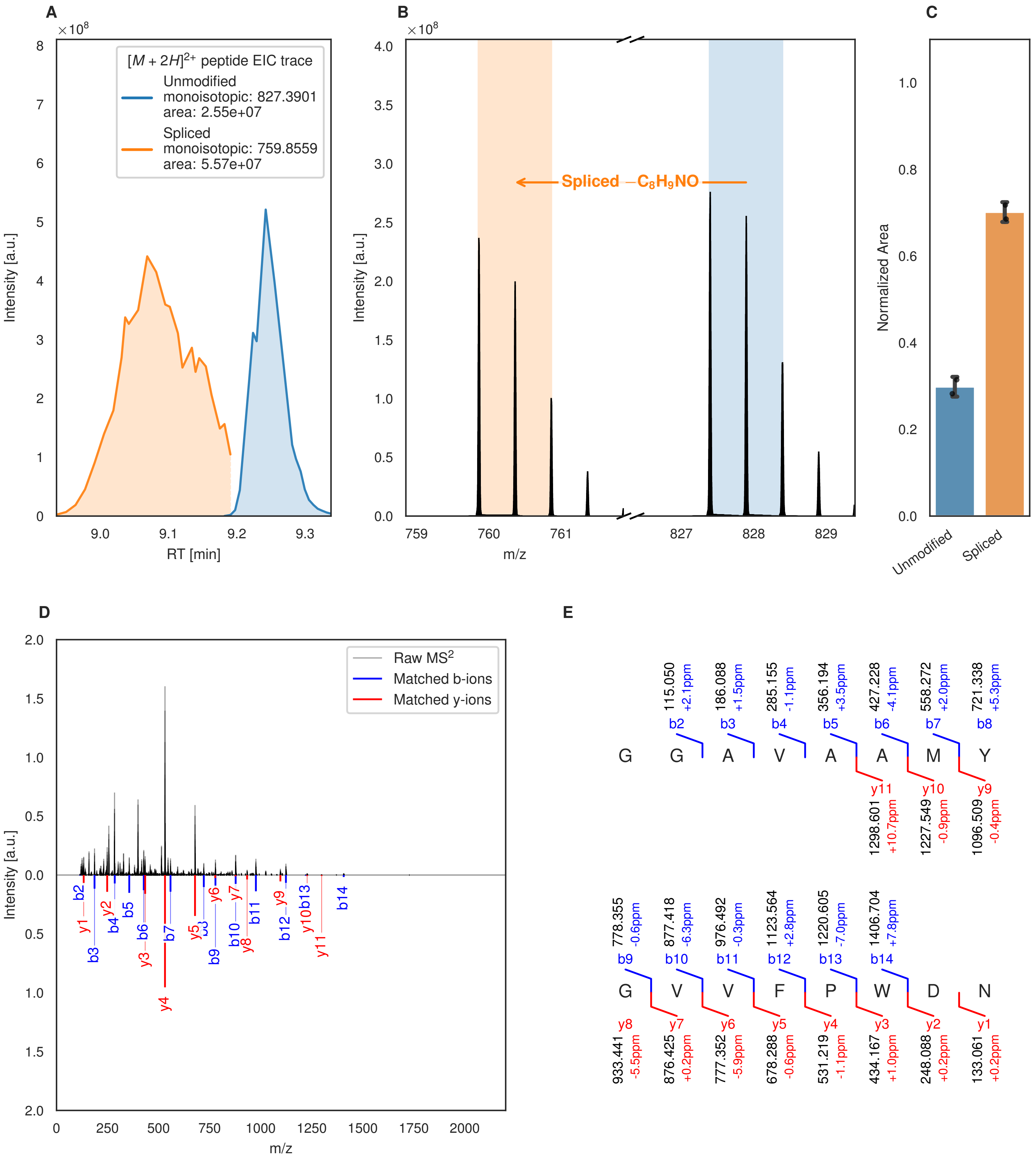

A
B
C
D
E
[M + 2H]2+ peptide EIC trace
Unmodified
monoisotopic: 827.3901
area: 2.55e+07
Spliced
monoisotopic: 759.8559
area: 5.57e+07
Intensity [a.u.]
RT [min]
Spliced −C8H9NO
m/z
Normalized Area
Unmodified
Spliced
Raw MS2
Matched b-ions
Matched y-ions
G G A V A A M Y
G V V F P W D N

Figure S44 (figure continued from the previous page).

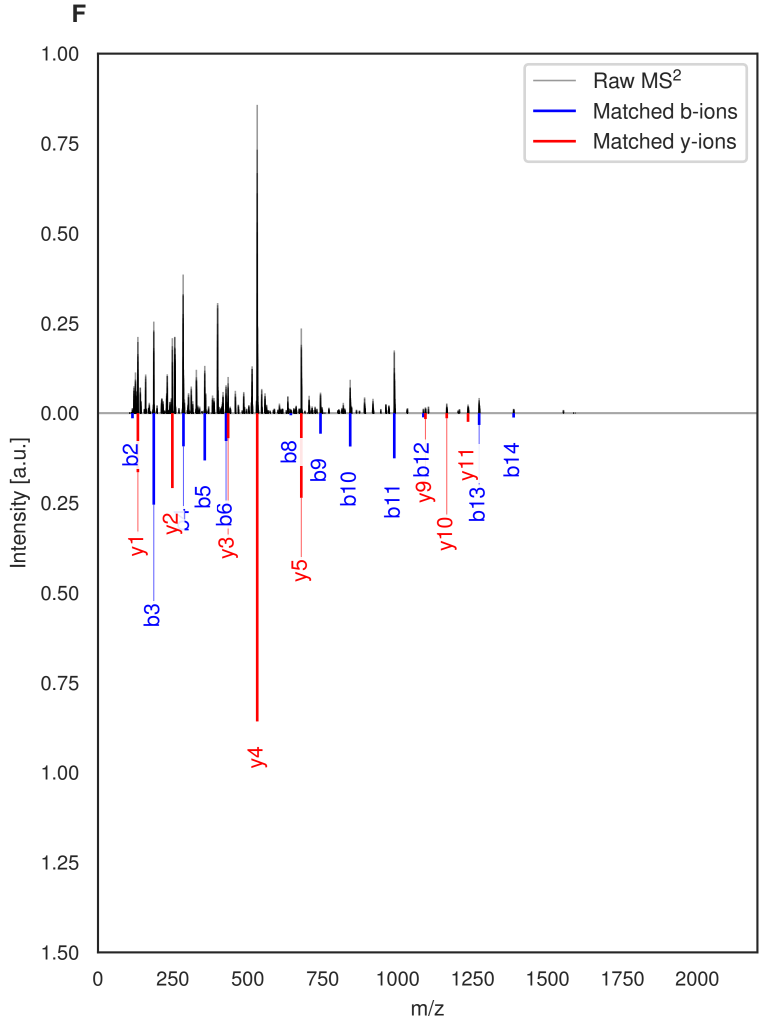

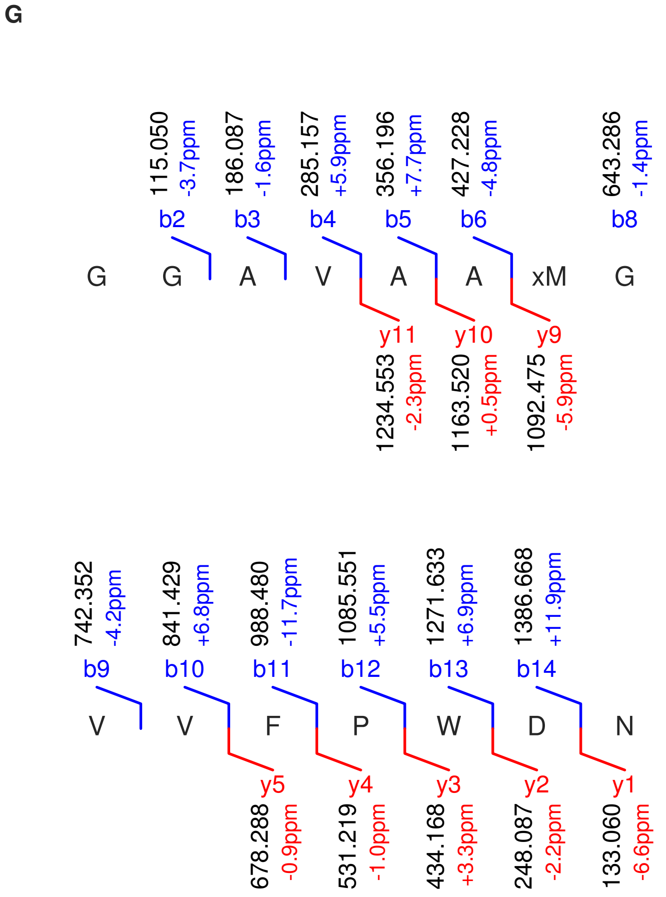

**Figure S44: Modification of the TJ-C-680 precursor peptide by PlpXY.** LC-MS analysis of IMAC-purified precursor peptides from *E. coli* BL21(DE3)Tuner containing splicease-encoding plasmid TJ-C-559 (PlpXY) and precursor peptide-encoding plasmid TJ-C-680 (RSTN1-PlpA3-TEV-PlpA3-Trunc) after expression at 24 °C for 22 h in 30 mL cultures after digestion with TEV. Data from n=2 independent biological replicates. Spliced amino acids have an x as a prefix. Spliced refers to a single tyramine excision. A representative experimental replicate is shown in panels **A, B.**

**A** Close-up plot of the representative extracted ion chromatogram (EIC) of the various peptide states; the blue trace corresponds to the Unmodified peptide; the orange trace corresponds to the Spliced peptide. **B** Close-up plot of the m/z values of the peptides detected by the mass spectrometer, with the shaded colored areas corresponding to where the expected m/z values for the peptide m/z traces should occur, with the observed spectrum plotted in black. The Unmodified, Spliced peptide(s) were found in the mass spectrum. **C** Bar plot showing the relative abundance of the various peptide states at charge z=2; the bars represent the average relative abundances normalized to total peak areas, with each individual experimental replicate value plotted as a black dot. **D** Mirrored $MS^2$ plot for the Unmodified peptide. In this mirror plot, the full fragmentation spectrum is plotted in black pointing upward, and the matched fragment b ions (blue) and y ions (red) are plotted pointing downward, mirrored against it, so that each matched fragment can be compared directly with the measured spectrum. **E** $MS^2$ spectrum of the fragments corresponding to the Unmodified peptide sequence, with matched b (blue, above) and y ions (red, below) plotted. **F** Mirrored $MS^2$ plot for the Spliced peptide. In this mirror plot, the full fragmentation spectrum is plotted in black pointing upward, and the matched fragment b ions (blue) and y ions (red) are plotted pointing downward, mirrored against it, so that each matched fragment can be compared directly with the measured spectrum. **G** $MS^2$ spectrum of the fragments corresponding to the Spliced peptide sequence, with matched b (blue, above) and y ions (red, below) plotted.

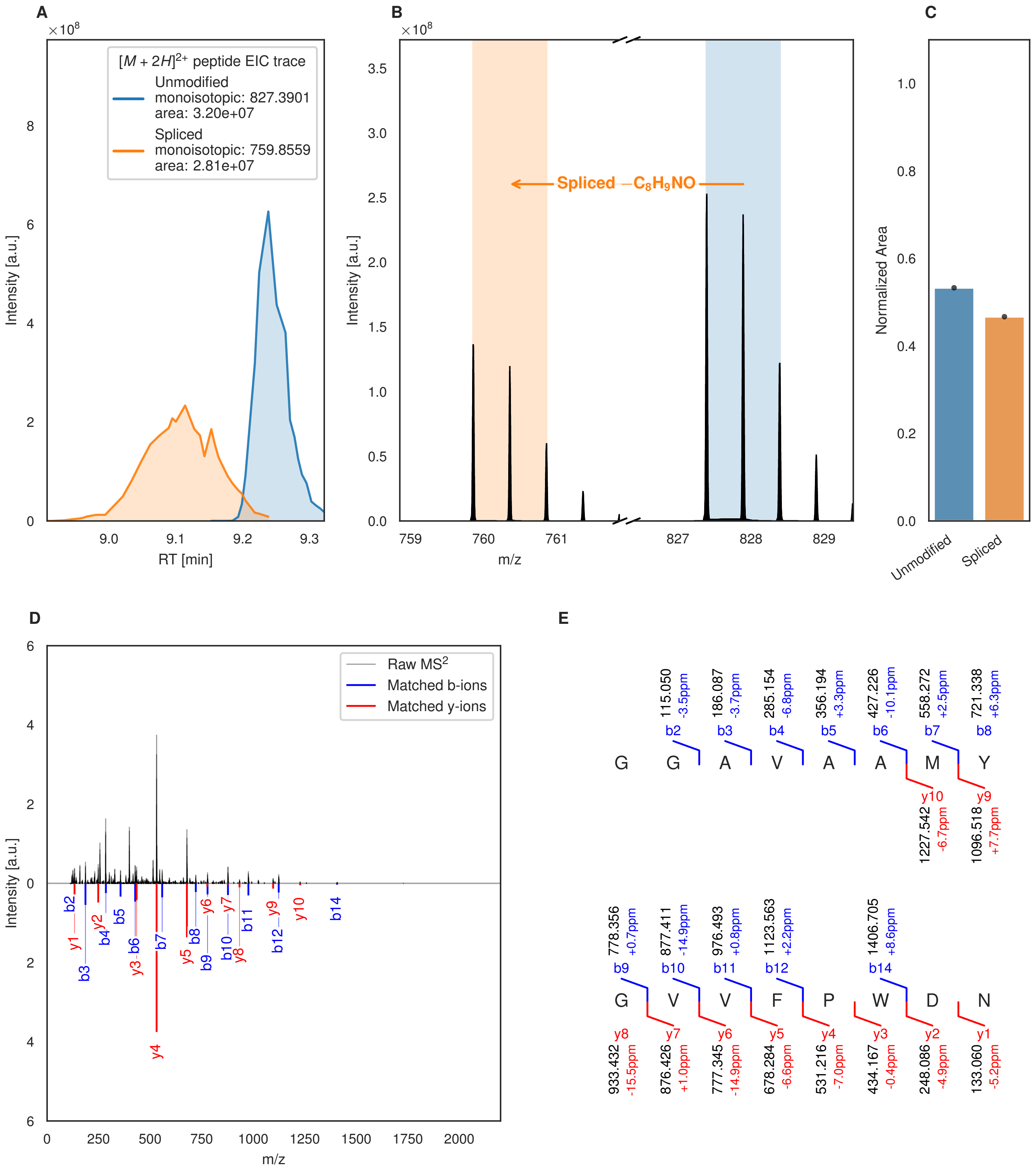
A
×10^8
[M + 2H]2+ peptide EIC trace
Unmodified
monoisotopic: 827.3901
area: 3.20e+07
Spliced
monoisotopic: 759.8559
area: 2.81e+07
Intensity [a.u.]
RT [min]
B
×10^8
Spliced −C8H9NO
Intensity [a.u.]
m/z
C
Normalized Area
Unmodified
Spliced
D
Raw MS2
Matched b-ions
Matched y-ions
Intensity [a.u.]
m/z
E
115.050 -3.5ppm b2
186.087 -3.7ppm b3
285.154 -6.8ppm b4
356.194 +3.3ppm b5
427.226 -10.1ppm b6
558.272 +2.5ppm b7
721.338 +6.3ppm b8
G G A V A A M Y
y10 1227.542 -6.7ppm
y9 1096.518 +7.7ppm
778.356 +0.7ppm b9
877.411 -14.9ppm b10
976.493 +0.8ppm b11
1123.563 +2.2ppm b12
1406.705 +8.6ppm b14
G V V F P W D N
y8 933.432 -15.5ppm
y7 876.426 +1.0ppm
y6 777.345 -14.9ppm
y5 678.284 -6.6ppm
y4 531.216 -7.0ppm
y3 434.167 -0.4ppm
y2 248.086 -4.9ppm
y1 133.060 -5.2ppm

Figure S45 (figure continued from the previous page).

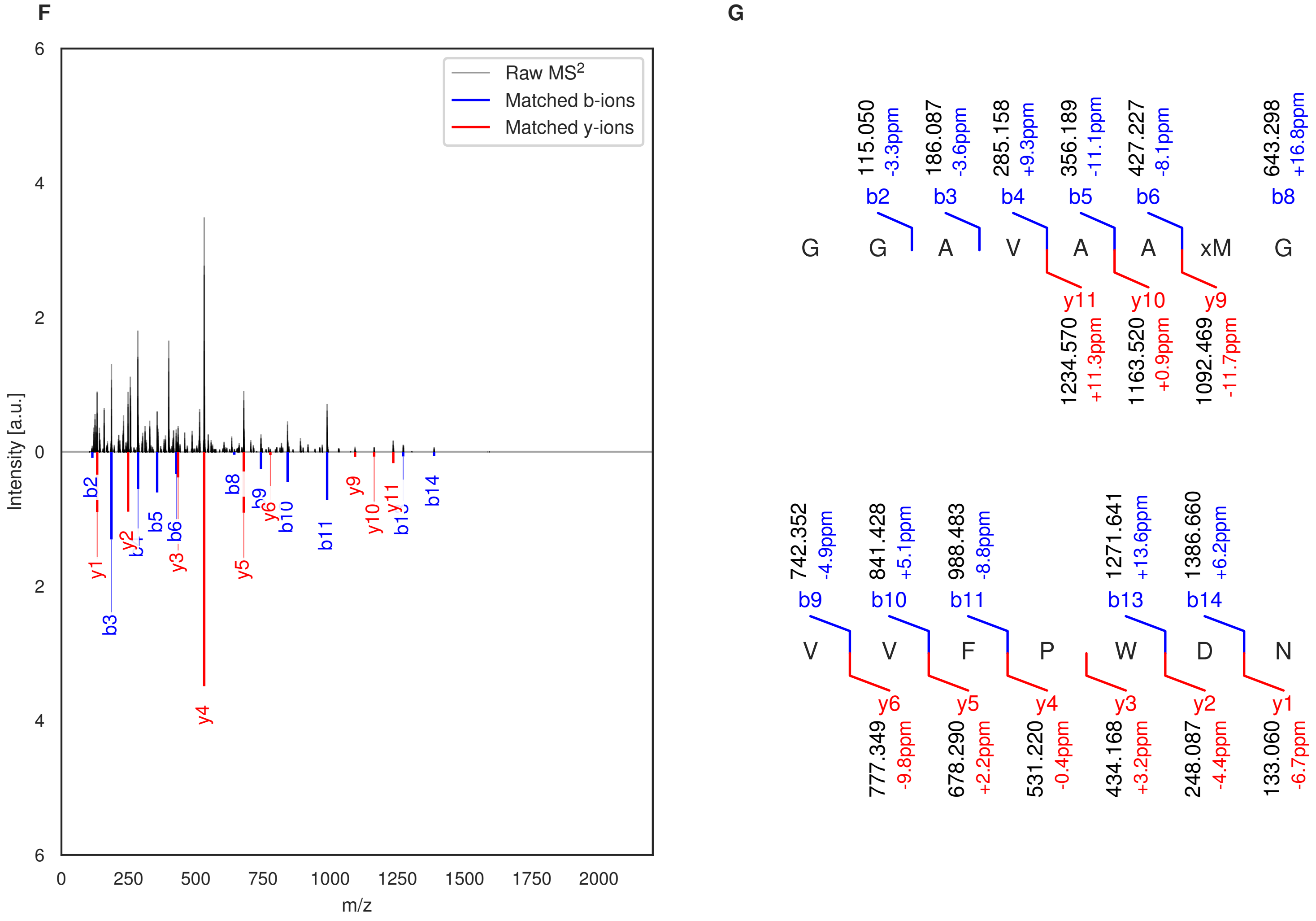

**Figure S45: Modification of the TJ-C-680 precursor peptide by PlpXY.** LC-MS analysis of IMAC-purified precursor peptides from *E. coli* BL21(DE3)Tuner containing splicease-encoding plasmid TJ-C-559 (PlpXY) and precursor peptide-encoding plasmid TJ-C-680 (RSTN1-PlpA3-TEV-PlpA3-Trunc) after expression at 30 °C for 22 h in 30 mL cultures after digestion with TEV. Data from n=1 independent biological replicates. Spliced amino acids have an x as a prefix. Spliced refers to a single tyramine excision. A representative experimental replicate is shown in panels **A, B.**

**A** Close-up plot of the representative extracted ion chromatogram (EIC) of the various peptide states; the blue trace corresponds to the Unmodified peptide; the orange trace corresponds to the Spliced peptide. **B** Close-up plot of the m/z values of the peptides detected by the mass spectrometer, with the shaded colored areas corresponding to where the expected m/z values for the peptide m/z traces should occur, with the observed spectrum plotted in black. The Unmodified, Spliced peptide(s) were found in the mass spectrum. **C** Bar plot showing the relative abundance of the various peptide states at charge z=2; the bars represent the average relative abundances normalized to total peak areas, with each individual experimental replicate value plotted as a black dot. **D** Mirrored MS$^2$ plot for the Unmodified peptide. In this mirror plot, the full fragmentation spectrum is plotted in black pointing upward, and the matched fragment b ions (blue) and y ions (red) are plotted pointing downward, mirrored against it, so that each matched fragment can be compared directly with the measured spectrum. **E** MS$^2$ spectrum of the fragments corresponding to the Unmodified peptide sequence, with matched b (blue, above) and y ions (red, below) plotted. **F** Mirrored MS$^2$ plot for the Spliced peptide. In this mirror plot, the full fragmentation spectrum is plotted in black pointing upward, and the matched fragment b ions (blue) and y ions (red) are plotted pointing downward, mirrored against it, so that each matched fragment can be compared directly with the measured spectrum. **G** MS$^2$ spectrum of the fragments corresponding to the Spliced peptide sequence, with matched b (blue, above) and y ions (red, below) plotted.

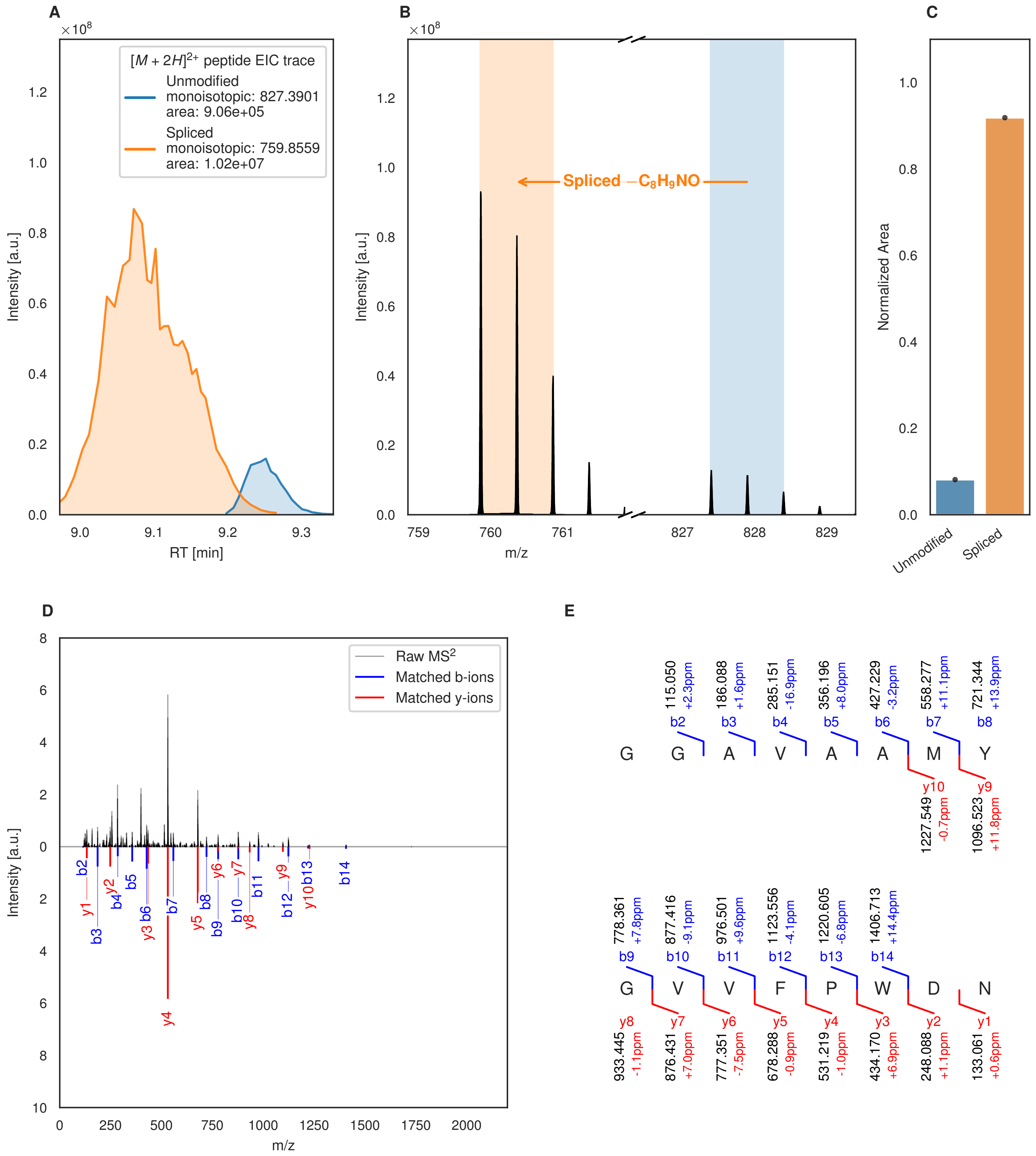
A
[M + 2H]2+ peptide EIC trace
Unmodified
monoisotopic: 827.3901
area: 9.06e+05
Spliced
monoisotopic: 759.8559
area: 1.02e+07
Intensity [a.u.]
RT [min]
B
Spliced −C8H9NO
Intensity [a.u.]
m/z
C
Normalized Area
Unmodified
Spliced
D
Raw MS2
Matched b-ions
Matched y-ions
Intensity [a.u.]
m/z
E
G G A V A A M Y
G V V F P W D N

Figure S46 (figure continued from the previous page).

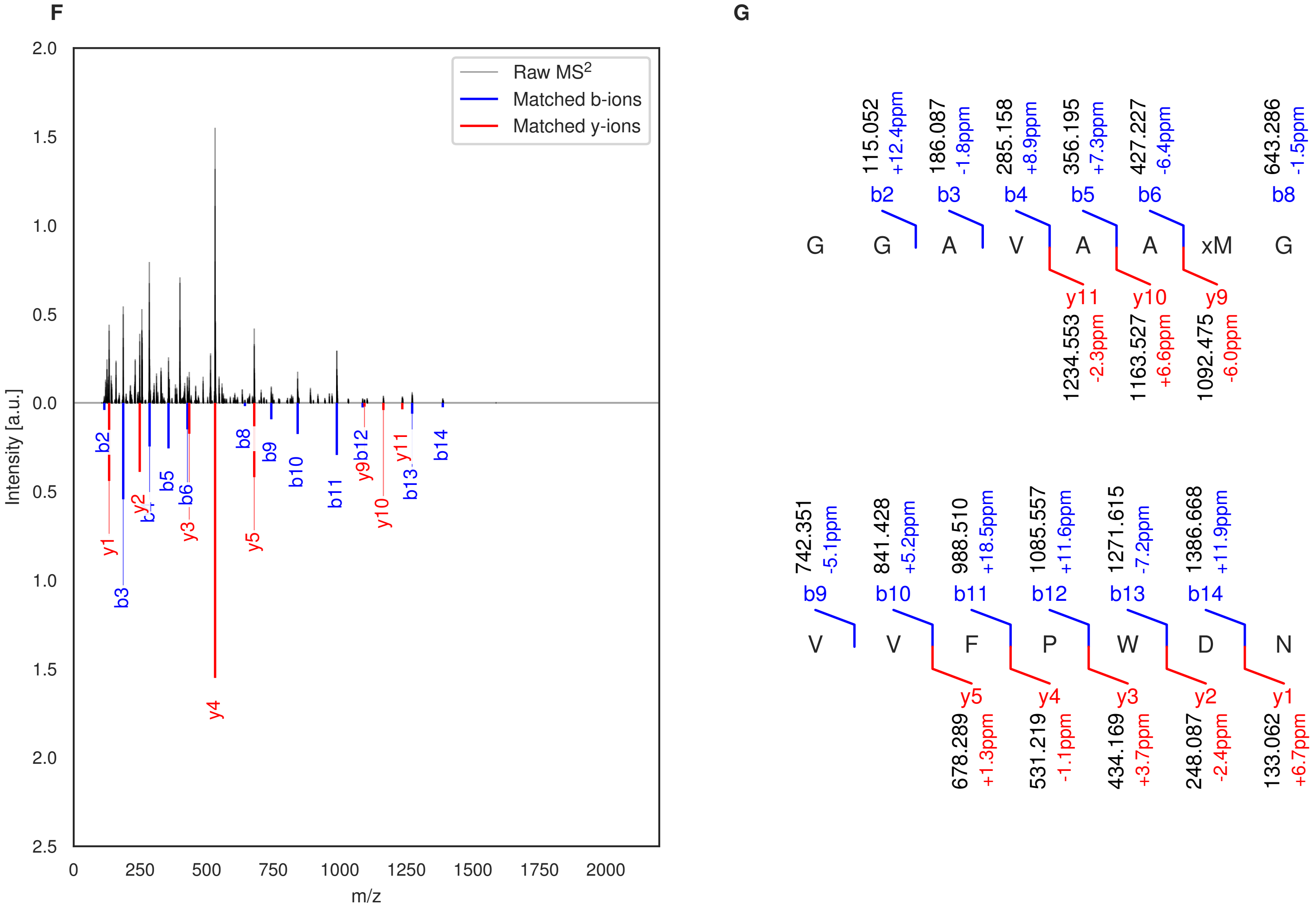

**Figure S46: Modification of the TJ-C-681 precursor peptide by PlpXY.** LC-MS analysis of IMAC-purified precursor peptides from *E. coli* BL21(DE3)Tuner containing splicease-encoding plasmid TJ-C-559 (PlpXY) and precursor peptide-encoding plasmid TJ-C-681 (Fh8-chaos-PlpA3-TEV-PlpA3-Trunc) after expression at 24 °C for 22 h in 30 mL cultures after digestion with TEV. Data from n=1 independent biological replicates. Spliced amino acids have an x as a prefix. Spliced refers to a single tyramine excision. A representative experimental replicate is shown in panels **A, B.**

**A** Close-up plot of the representative extracted ion chromatogram (EIC) of the various peptide states; the blue trace corresponds to the Unmodified peptide; the orange trace corresponds to the Spliced peptide. **B** Close-up plot of the m/z values of the peptides detected by the mass spectrometer, with the shaded colored areas corresponding to where the expected m/z values for the peptide m/z traces should occur, with the observed spectrum plotted in black. The Unmodified, Spliced peptide(s) were found in the mass spectrum. **C** Bar plot showing the relative abundance of the various peptide states at charge z=2; the bars represent the average relative abundances normalized to total peak areas, with each individual experimental replicate value plotted as a black dot. **D** Mirrored MS$^2$ plot for the Unmodified peptide. In this mirror plot, the full fragmentation spectrum is plotted in black pointing upward, and the matched fragment b ions (blue) and y ions (red) are plotted pointing downward, mirrored against it, so that each matched fragment can be compared directly with the measured spectrum. **E** MS$^2$ spectrum of the fragments corresponding to the Unmodified peptide sequence, with matched b (blue, above) and y ions (red, below) plotted. **F** Mirrored MS$^2$ plot for the Spliced peptide. In this mirror plot, the full fragmentation spectrum is plotted in black pointing upward, and the matched fragment b ions (blue) and y ions (red) are plotted pointing downward, mirrored against it, so that each matched fragment can be compared directly with the measured spectrum. **G** MS$^2$ spectrum of the fragments corresponding to the Spliced peptide sequence, with matched b (blue, above) and y ions (red, below) plotted.

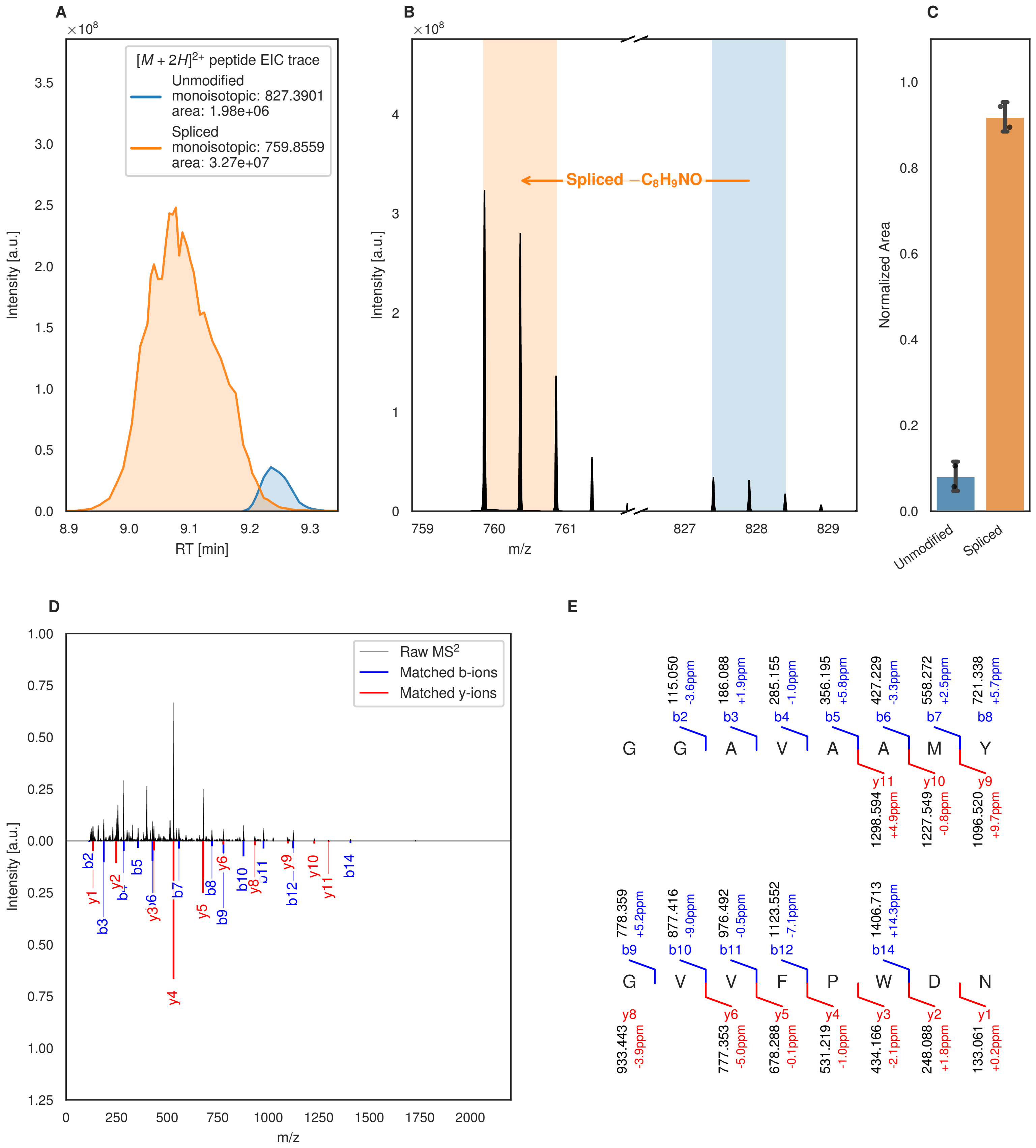
A
B
C
D
E
[M + 2H]2+ peptide EIC trace
Unmodified
monoisotopic: 827.3901
area: 1.98e+06
Spliced
monoisotopic: 759.8559
area: 3.27e+07
Intensity [a.u.]
RT [min]
Spliced −C8H9NO
m/z
Normalized Area
Unmodified
Spliced
Raw MS2
Matched b-ions
Matched y-ions
G G A V A A M Y
G V V F P W D N

Figure S47 (figure continued from the previous page).

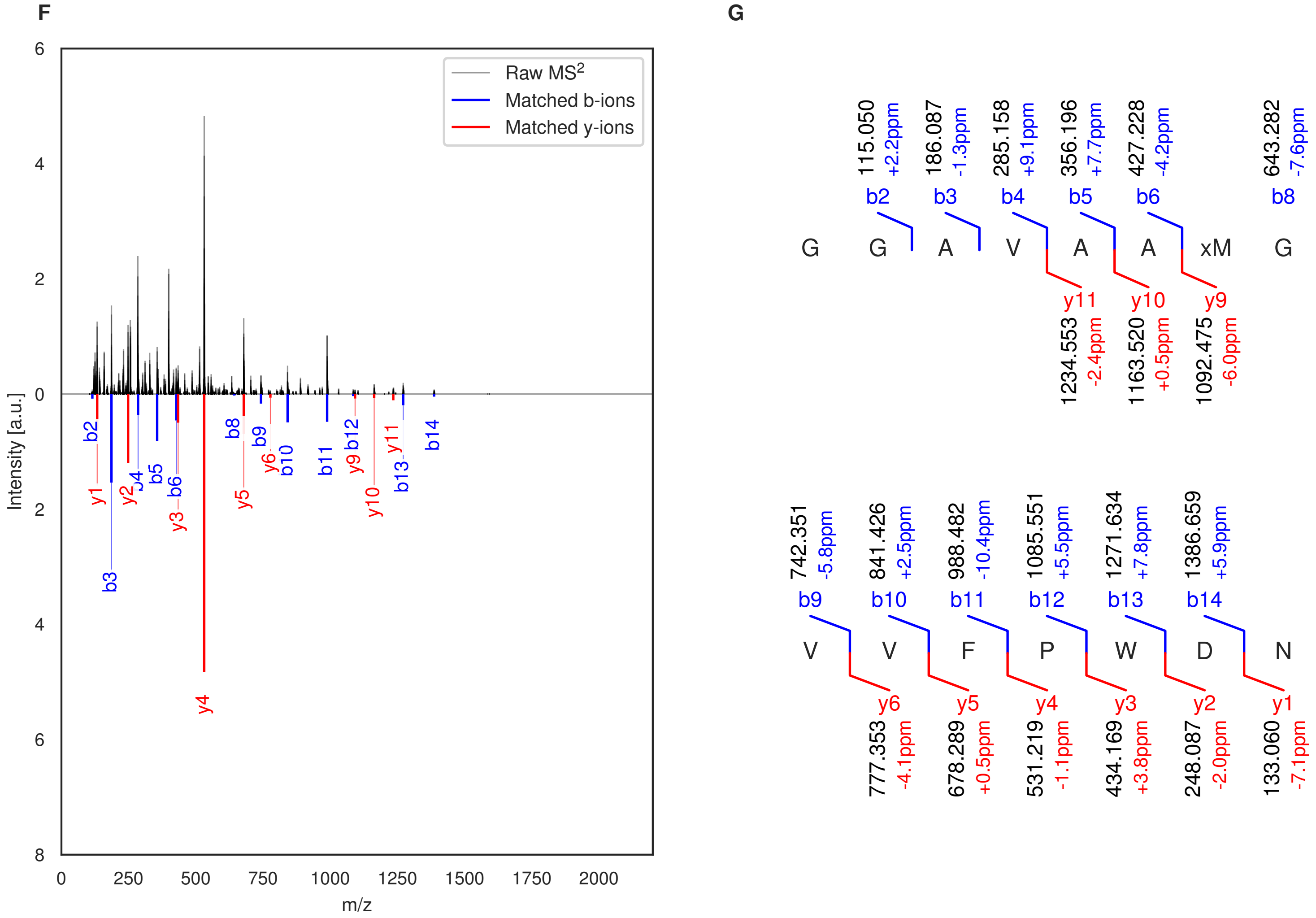

**Figure S47: Modification of the TJ-C-682 precursor peptide by PlpXY.** LC-MS analysis of IMAC-purified precursor peptides from *E. coli* BL21(DE3)Tuner containing splicease-encoding plasmid TJ-C-559 (PlpXY) and precursor peptide-encoding plasmid TJ-C-682 (Fh8-PlpA3-TEV-PlpA3-Trunc) after expression at 24 °C for 22 h in 30 mL cultures after digestion with TEV. Data from n=2 independent biological replicates. Spliced amino acids have an x as a prefix. Spliced refers to a single tyramine excision. A representative experimental replicate is shown in panels **A, B.**

**A** Close-up plot of the representative extracted ion chromatogram (EIC) of the various peptide states; the blue trace corresponds to the Unmodified peptide; the orange trace corresponds to the Spliced peptide. **B** Close-up plot of the m/z values of the peptides detected by the mass spectrometer, with the shaded colored areas corresponding to where the expected m/z values for the peptide m/z traces should occur, with the observed spectrum plotted in black. The Unmodified, Spliced peptide(s) were found in the mass spectrum. **C** Bar plot showing the relative abundance of the various peptide states at charge z=2; the bars represent the average relative abundances normalized to total peak areas, with each individual experimental replicate value plotted as a black dot. **D** Mirrored MS$^2$ plot for the Unmodified peptide. In this mirror plot, the full fragmentation spectrum is plotted in black pointing upward, and the matched fragment b ions (blue) and y ions (red) are plotted pointing downward, mirrored against it, so that each matched fragment can be compared directly with the measured spectrum. **E** MS$^2$ spectrum of the fragments corresponding to the Unmodified peptide sequence, with matched b (blue, above) and y ions (red, below) plotted. **F** Mirrored MS$^2$ plot for the Spliced peptide. In this mirror plot, the full fragmentation spectrum is plotted in black pointing upward, and the matched fragment b ions (blue) and y ions (red) are plotted pointing downward, mirrored against it, so that each matched fragment can be compared directly with the measured spectrum. **G** MS$^2$ spectrum of the fragments corresponding to the Spliced peptide sequence, with matched b (blue, above) and y ions (red, below) plotted.

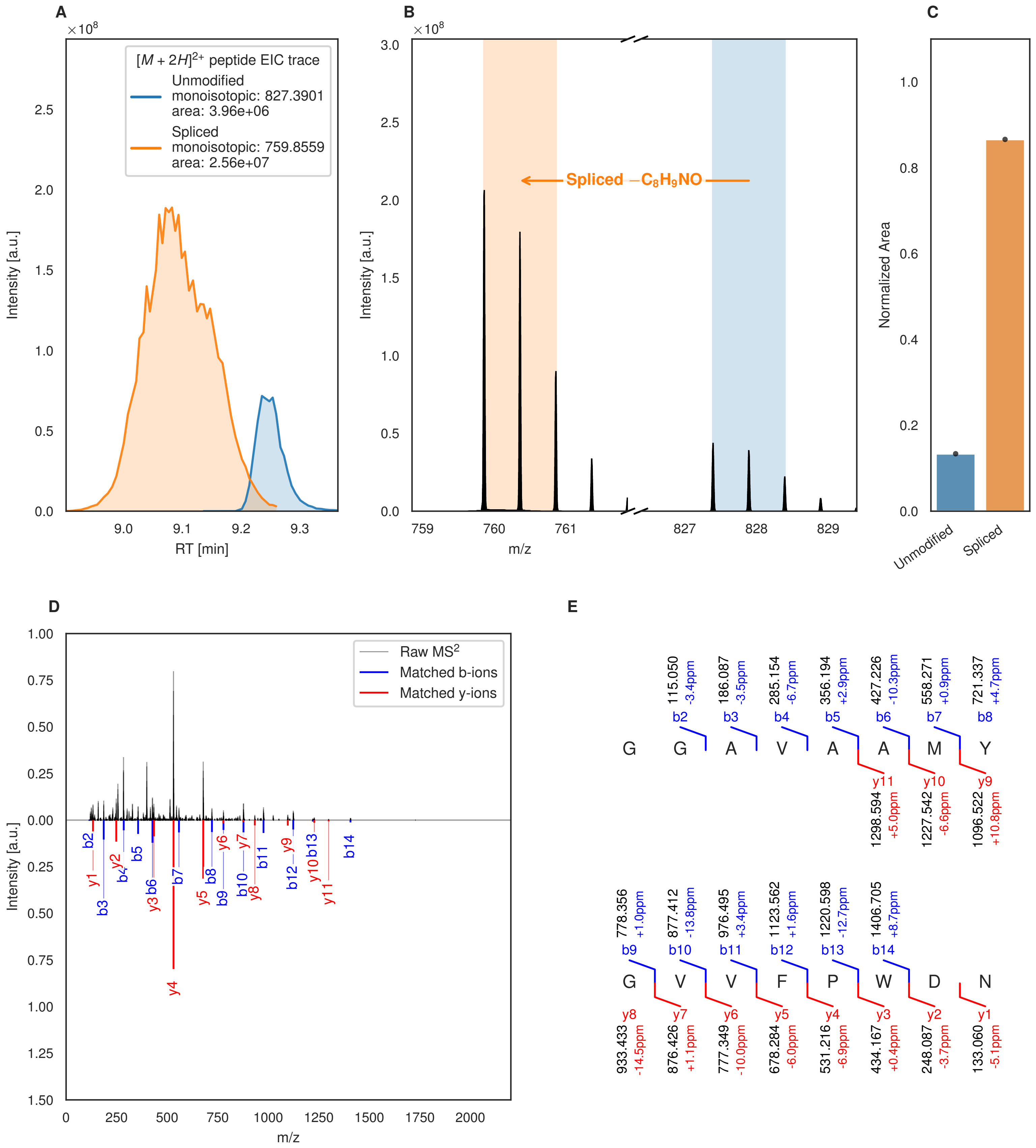
A
B
C
D
E
$[M+2H]^{2+}$ peptide EIC trace
Unmodified
monoisotopic: 827.3901
area: 3.96e+06
Spliced
monoisotopic: 759.8559
area: 2.56e+07
Intensity [a.u.]
RT [min]
Spliced $-C_8H_9NO$
m/z
Normalized Area
Unmodified
Spliced
Raw MS$^2$
Matched b-ions
Matched y-ions
G G A V A A M Y
G V V F P W D N
115.050 -3.4ppm b2
186.087 -3.5ppm b3
285.154 -6.7ppm b4
356.194 +2.9ppm b5
427.226 -10.3ppm b6
558.271 +0.9ppm b7
721.337 +4.7ppm b8
y11 1298.594 +5.0ppm
y10 1227.542 -6.6ppm
y9 1096.522 +10.8ppm
778.356 +1.0ppm b9
877.412 -13.8ppm b10
976.495 +3.4ppm b11
1123.562 +1.6ppm b12
1220.598 -12.7ppm b13
1406.705 +8.7ppm b14
y8 933.433 -14.5ppm
y7 876.426 +1.1ppm
y6 777.349 -10.0ppm
y5 678.284 -6.0ppm
y4 531.216 -6.9ppm
y3 434.167 +0.4ppm
y2 248.087 -3.7ppm
y1 133.060 -5.1ppm

Figure S48 (figure continued from the previous page).

F

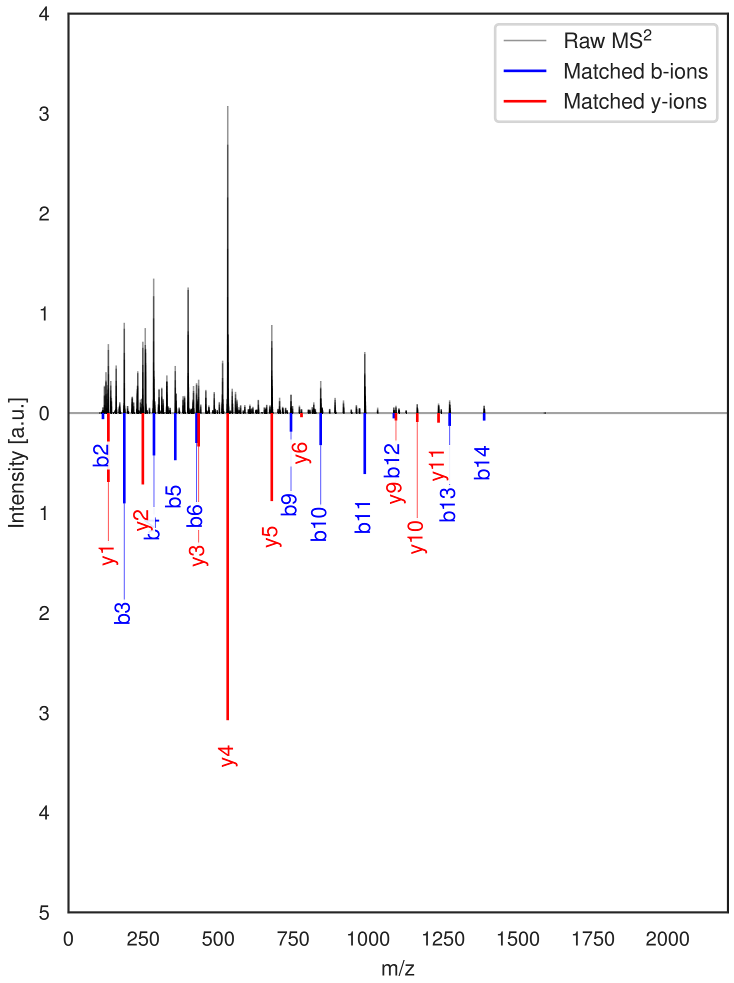

G

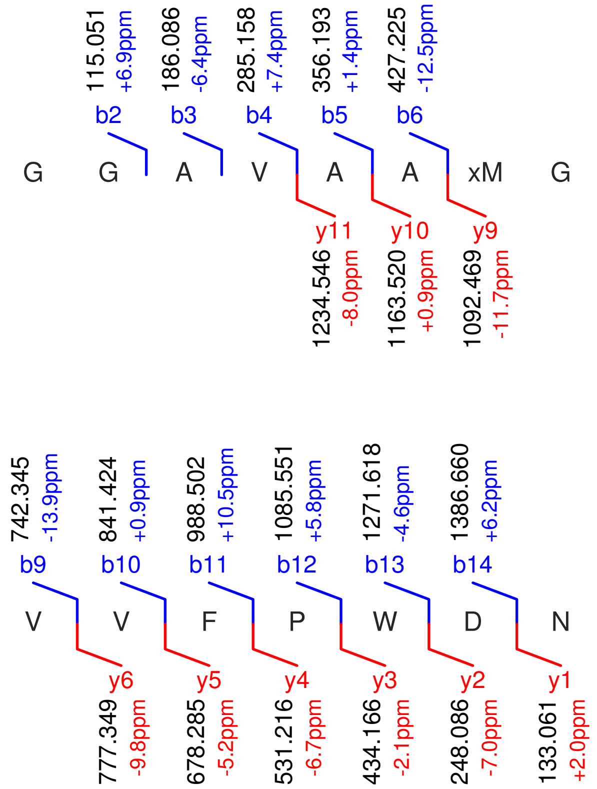

**Figure S48: Modification of the TJ-C-682 precursor peptide by PlpXY.** LC-MS analysis of IMAC-purified precursor peptides from *E. coli* BL21(DE3)Tuner containing splicease-encoding plasmid TJ-C-559 (PlpXY) and precursor peptide-encoding plasmid TJ-C-682 (Fh8-PlpA3-TEV-PlpA3-Trunc) after expression at 30 °C for 22 h in 30 mL cultures after digestion with TEV. Data from n=1 independent biological replicates. Spliced amino acids have an x as a prefix. Spliced refers to a single tyramine excision. A representative experimental replicate is shown in panels **A, B.**

**A** Close-up plot of the representative extracted ion chromatogram (EIC) of the various peptide states; the blue trace corresponds to the Unmodified peptide; the orange trace corresponds to the Spliced peptide. **B** Close-up plot of the m/z values of the peptides detected by the mass spectrometer, with the shaded colored areas corresponding to where the expected m/z values for the peptide m/z traces should occur, with the observed spectrum plotted in black. The Unmodified, Spliced peptide(s) were found in the mass spectrum. **C** Bar plot showing the relative abundance of the various peptide states at charge z=2; the bars represent the average relative abundances normalized to total peak areas, with each individual experimental replicate value plotted as a black dot. **D** Mirrored MS$^2$ plot for the Unmodified peptide. In this mirror plot, the full fragmentation spectrum is plotted in black pointing upward, and the matched fragment b ions (blue) and y ions (red) are plotted pointing downward, mirrored against it, so that each matched fragment can be compared directly with the measured spectrum. **E** MS$^2$ spectrum of the fragments corresponding to the Unmodified peptide sequence, with matched b (blue, above) and y ions (red, below) plotted. **F** Mirrored MS$^2$ plot for the Spliced peptide. In this mirror plot, the full fragmentation spectrum is plotted in black pointing upward, and the matched fragment b ions (blue) and y ions (red) are plotted pointing downward, mirrored against it, so that each matched fragment can be compared directly with the measured spectrum. **G** MS$^2$ spectrum of the fragments corresponding to the Spliced peptide sequence, with matched b (blue, above) and y ions (red, below) plotted.

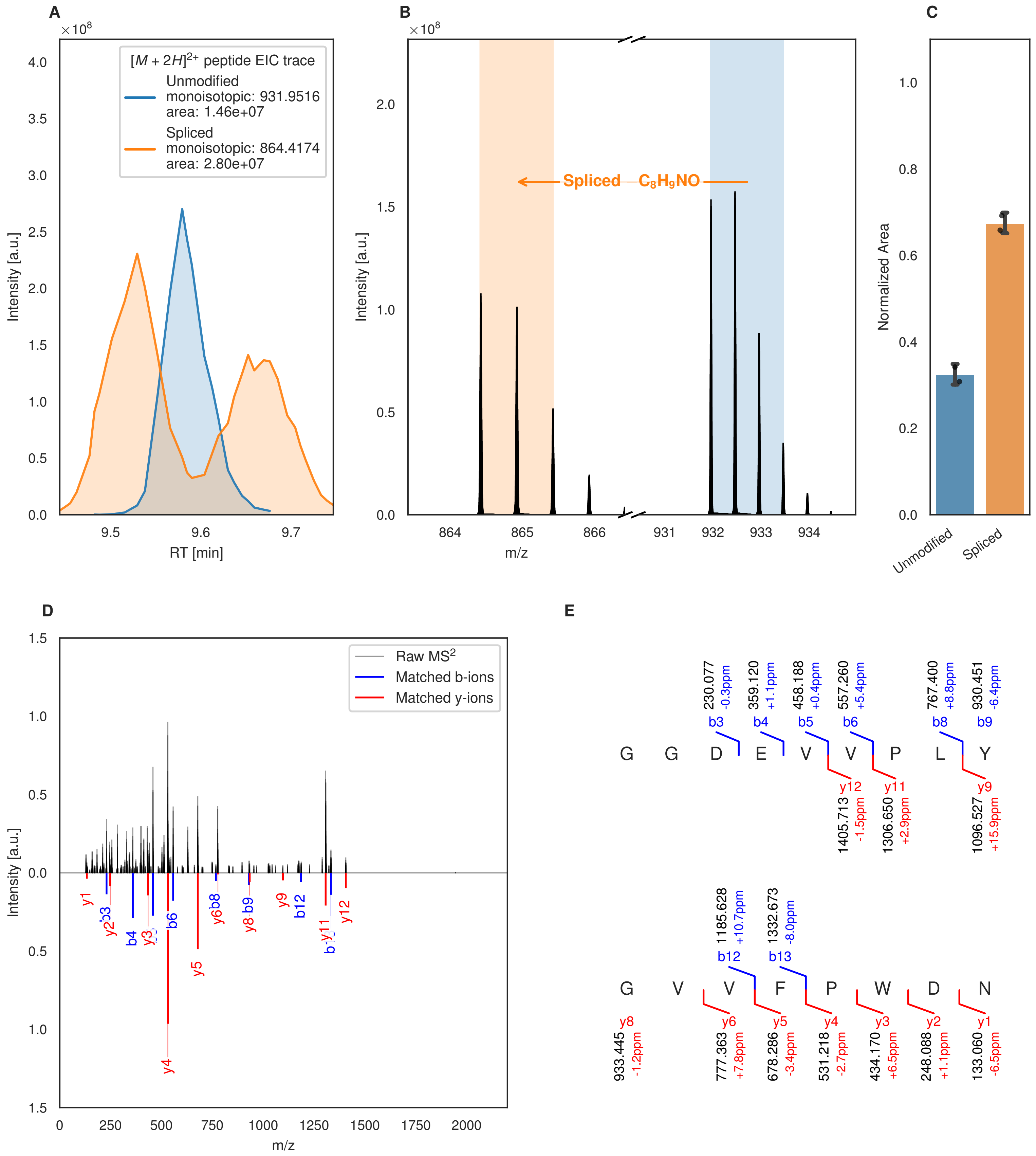

A
B
C
D
E
[M + 2H]2+ peptide EIC trace
Unmodified
monoisotopic: 931.9516
area: 1.46e+07
Spliced
monoisotopic: 864.4174
area: 2.80e+07
Intensity [a.u.]
RT [min]
Spliced −C8H9NO
m/z
Normalized Area
Unmodified
Spliced
Raw MS2
Matched b-ions
Matched y-ions
G G D E V V P L Y
b3 b4 b5 b6 b8 b9
230.077 -0.3ppm
359.120 +1.1ppm
458.188 +0.4ppm
557.260 +5.4ppm
767.400 +8.8ppm
930.451 -6.4ppm
y12 y11 y9
1405.713 -1.5ppm
1306.650 +2.9ppm
1096.527 +15.9ppm
G V V F P W D N
b12 b13
1185.628 +10.7ppm
1332.673 -8.0ppm
y8 y6 y5 y4 y3 y2 y1
933.445 -1.2ppm
777.363 +7.8ppm
678.286 -3.4ppm
531.218 -2.7ppm
434.170 +6.5ppm
248.088 +1.1ppm
133.060 -6.5ppm

Figure S49 (figure continued from the previous page).

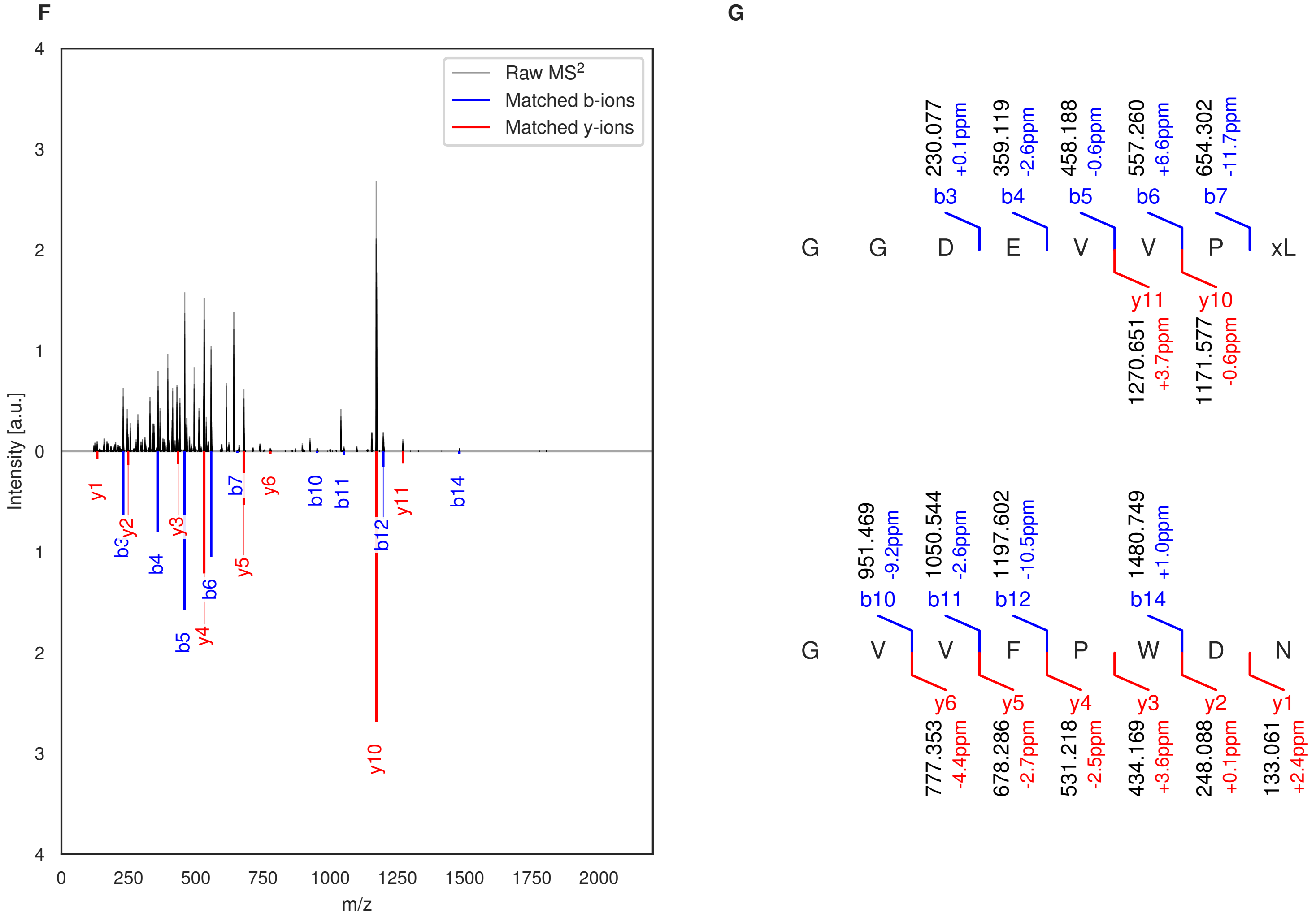

**Figure S49: Modification of the TJ-C-683 precursor peptide by PlpXY.** LC-MS analysis of IMAC-purified precursor peptides from *E. coli* BL21(DE3)Tuner containing splicease-encoding plasmid TJ-C-559 (PlpXY) and precursor peptide-encoding plasmid TJ-C-683 (RSTN1-PlpA3-TEV-PlpA3-Boc-Trunc) after expression at 24 °C for 22 h in 30 mL cultures after digestion with TEV. Data from n=2 independent biological replicates. Spliced amino acids have an x as a prefix. Spliced refers to a single tyramine excision. A representative experimental replicate is shown in panels **A, B.**

**A** Close-up plot of the representative extracted ion chromatogram (EIC) of the various peptide states; the blue trace corresponds to the Unmodified peptide; the orange trace corresponds to the Spliced peptide. **B** Close-up plot of the m/z values of the peptides detected by the mass spectrometer, with the shaded colored areas corresponding to where the expected m/z values for the peptide m/z traces should occur, with the observed spectrum plotted in black. The Unmodified, Spliced peptide(s) were found in the mass spectrum. **C** Bar plot showing the relative abundance of the various peptide states at charge z=2; the bars represent the average relative abundances normalized to total peak areas, with each individual experimental replicate value plotted as a black dot. **D** Mirrored MS$^2$ plot for the Unmodified peptide. In this mirror plot, the full fragmentation spectrum is plotted in black pointing upward, and the matched fragment b ions (blue) and y ions (red) are plotted pointing downward, mirrored against it, so that each matched fragment can be compared directly with the measured spectrum. **E** MS$^2$ spectrum of the fragments corresponding to the Unmodified peptide sequence, with matched b (blue, above) and y ions (red, below) plotted. **F** Mirrored MS$^2$ plot for the Spliced peptide. In this mirror plot, the full fragmentation spectrum is plotted in black pointing upward, and the matched fragment b ions (blue) and y ions (red) are plotted pointing downward, mirrored against it, so that each matched fragment can be compared directly with the measured spectrum. **G** MS$^2$ spectrum of the fragments corresponding to the Spliced peptide sequence, with matched b (blue, above) and y ions (red, below) plotted.

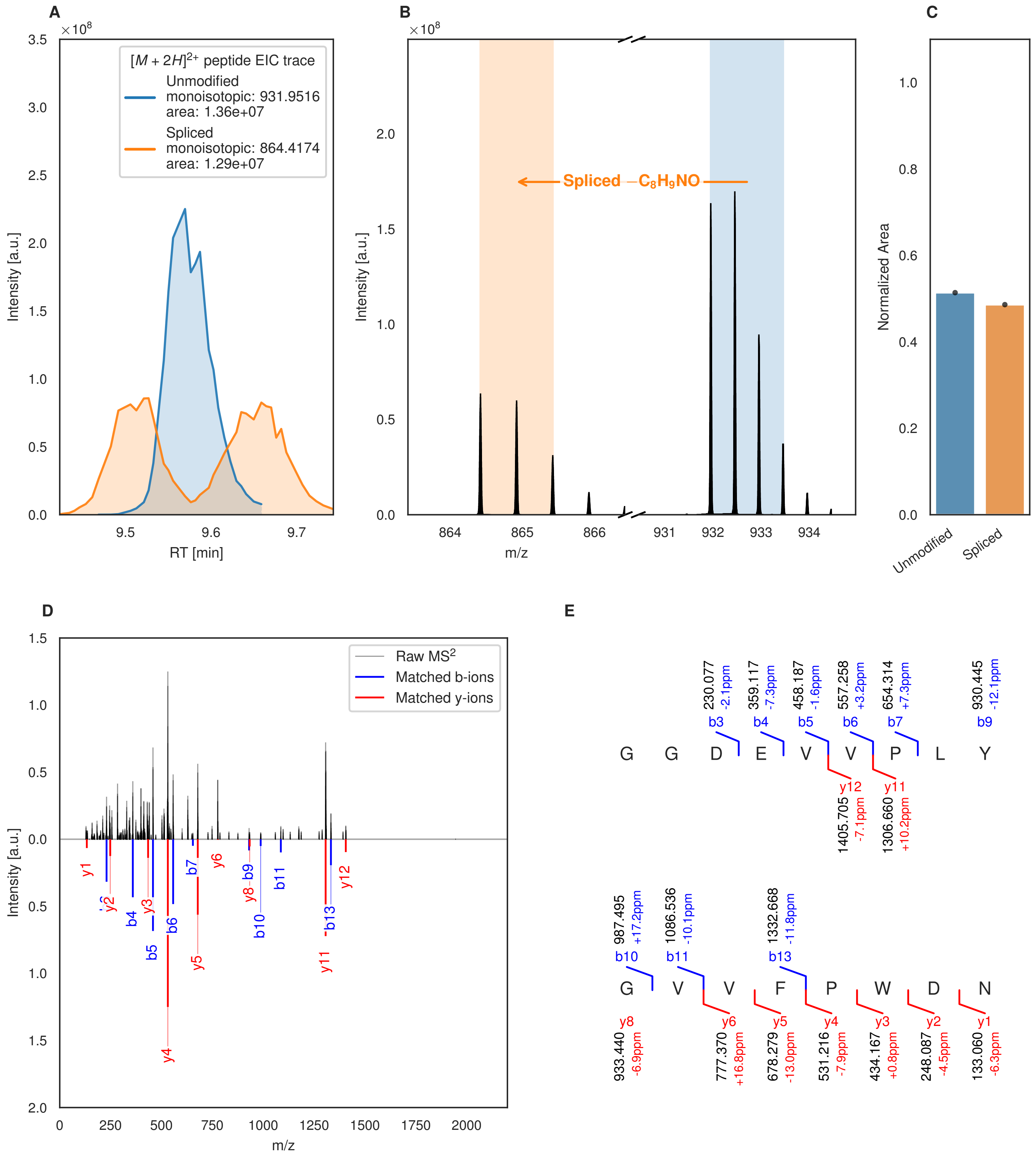

A
B
C
D
E
[M + 2H]2+ peptide EIC trace
Unmodified
monoisotopic: 931.9516
area: 1.36e+07
Spliced
monoisotopic: 864.4174
area: 1.29e+07
Intensity [a.u.]
RT [min]
Spliced −C8H9NO
m/z
Normalized Area
Unmodified
Spliced
Raw MS2
Matched b-ions
Matched y-ions
G G D E V V P L Y
G V V F P W D N

Figure S50 (figure continued from the previous page).

F

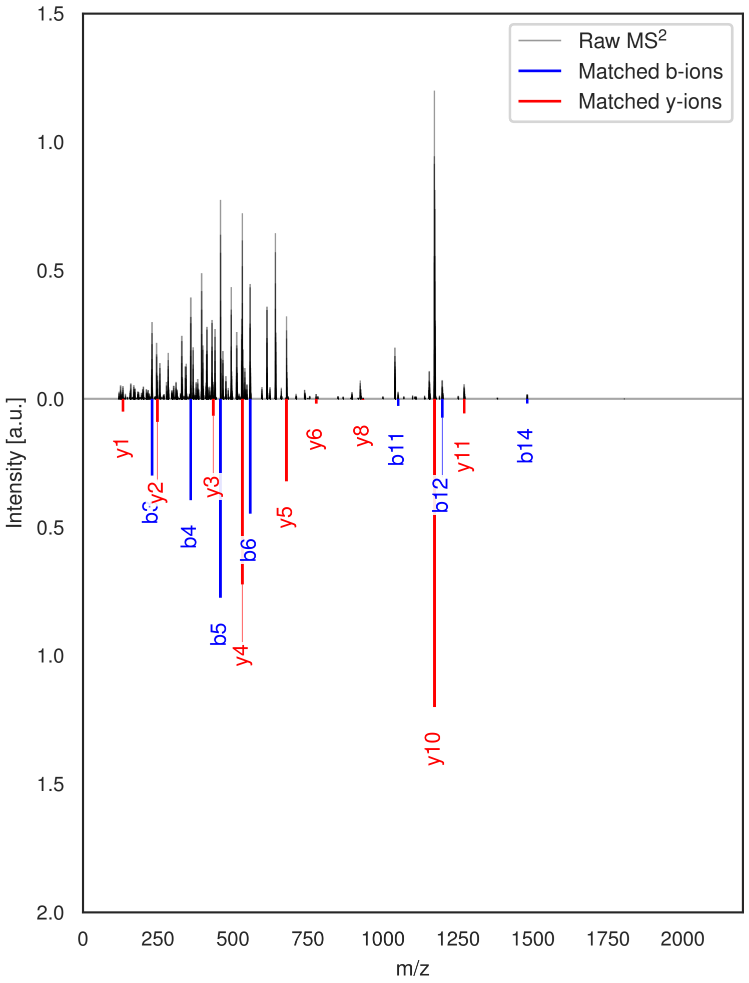

G

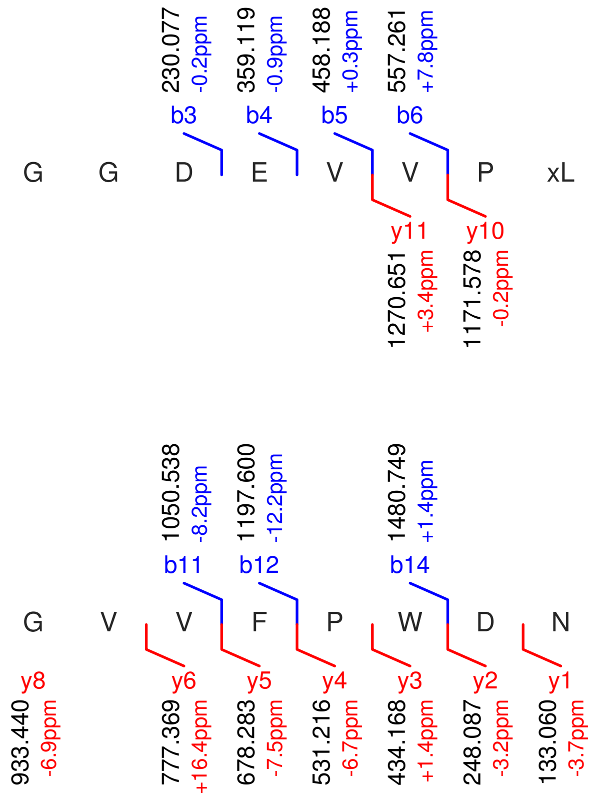

**Figure S50: Modification of the TJ-C-683 precursor peptide by PlpXY.** LC-MS analysis of IMAC-purified precursor peptides from *E. coli* BL21(DE3)Tuner containing splicease-encoding plasmid TJ-C-559 (PlpXY) and precursor peptide-encoding plasmid TJ-C-683 (RSTN1-PlpA3-TEV-PlpA3-Boc-Trunc) after expression at 30 °C for 22 h in 30 mL cultures after digestion with TEV. Data from n=1 independent biological replicates. Spliced amino acids have an x as a prefix. Spliced refers to a single tyramine excision. A representative experimental replicate is shown in panels **A, B.**

**A** Close-up plot of the representative extracted ion chromatogram (EIC) of the various peptide states; the blue trace corresponds to the Unmodified peptide; the orange trace corresponds to the Spliced peptide. **B** Close-up plot of the m/z values of the peptides detected by the mass spectrometer, with the shaded colored areas corresponding to where the expected m/z values for the peptide m/z traces should occur, with the observed spectrum plotted in black. The Unmodified, Spliced peptide(s) were found in the mass spectrum. **C** Bar plot showing the relative abundance of the various peptide states at charge z=2; the bars represent the average relative abundances normalized to total peak areas, with each individual experimental replicate value plotted as a black dot. **D** Mirrored $MS^2$ plot for the Unmodified peptide. In this mirror plot, the full fragmentation spectrum is plotted in black pointing upward, and the matched fragment b ions (blue) and y ions (red) are plotted pointing downward, mirrored against it, so that each matched fragment can be compared directly with the measured spectrum. **E** $MS^2$ spectrum of the fragments corresponding to the Unmodified peptide sequence, with matched b (blue, above) and y ions (red, below) plotted. **F** Mirrored $MS^2$ plot for the Spliced peptide. In this mirror plot, the full fragmentation spectrum is plotted in black pointing upward, and the matched fragment b ions (blue) and y ions (red) are plotted pointing downward, mirrored against it, so that each matched fragment can be compared directly with the measured spectrum. **G** $MS^2$ spectrum of the fragments corresponding to the Spliced peptide sequence, with matched b (blue, above) and y ions (red, below) plotted.

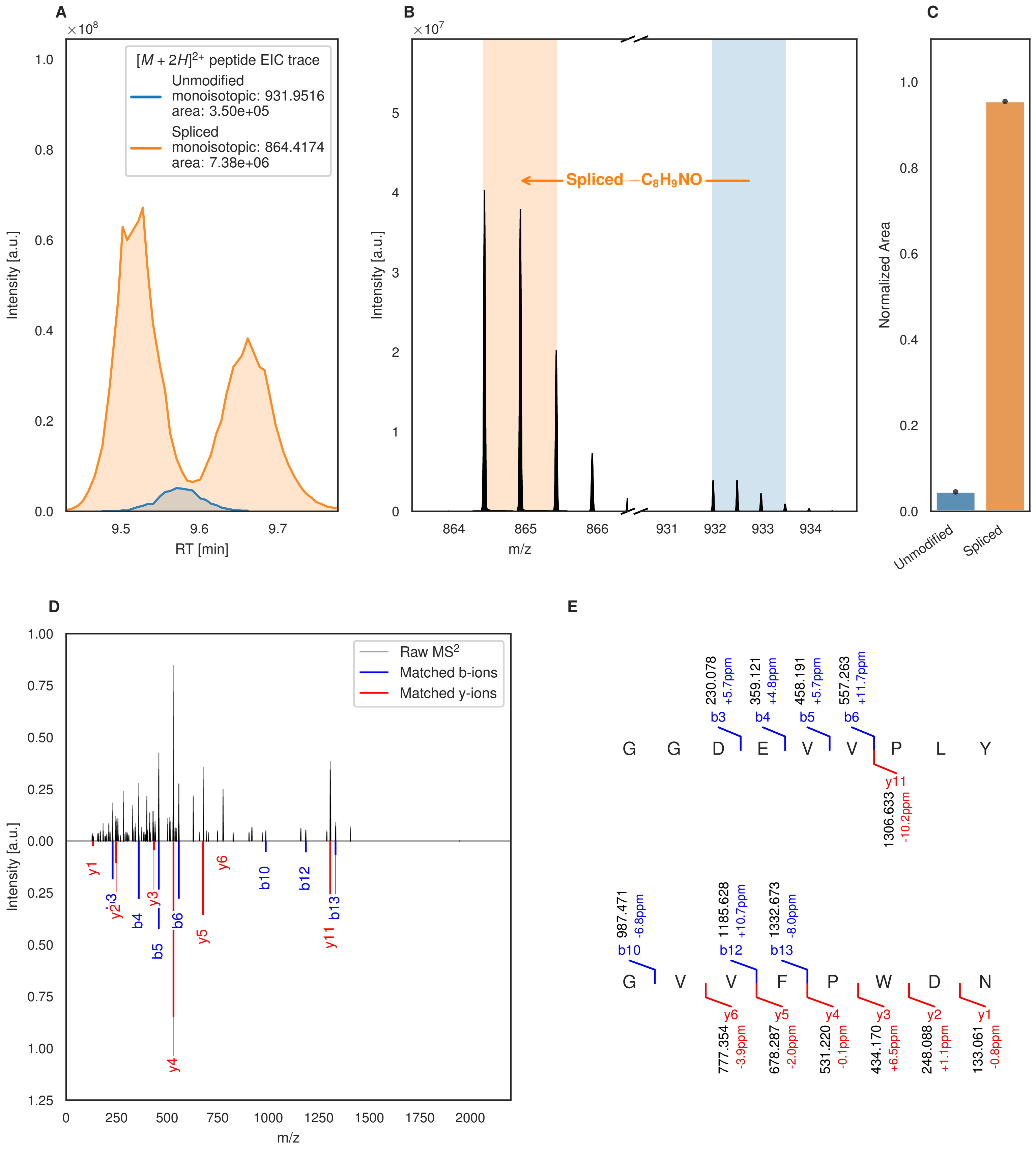
A
×10^8
[M + 2H]2+ peptide EIC trace
Unmodified
monoisotopic: 931.9516
area: 3.50e+05
Spliced
monoisotopic: 864.4174
area: 7.38e+06
Intensity [a.u.]
RT [min]
B
×10^7
Spliced −C8H9NO
Intensity [a.u.]
m/z
C
Normalized Area
Unmodified
Spliced
D
Raw MS2
Matched b-ions
Matched y-ions
Intensity [a.u.]
m/z
E
230.078 +5.7ppm b3
359.121 +4.8ppm b4
458.191 +5.7ppm b5
557.263 +11.7ppm b6
G G D E V V P L Y
y11 1306.633 -10.2ppm
987.471 -6.8ppm b10
1185.628 +10.7ppm b12
1332.673 -8.0ppm b13
G V V F P W D N
y6 777.354 -3.9ppm
y5 678.287 -2.0ppm
y4 531.220 -0.1ppm
y3 434.170 +6.5ppm
y2 248.088 +1.1ppm
y1 133.061 -0.8ppm

Figure S51 (figure continued from the previous page).

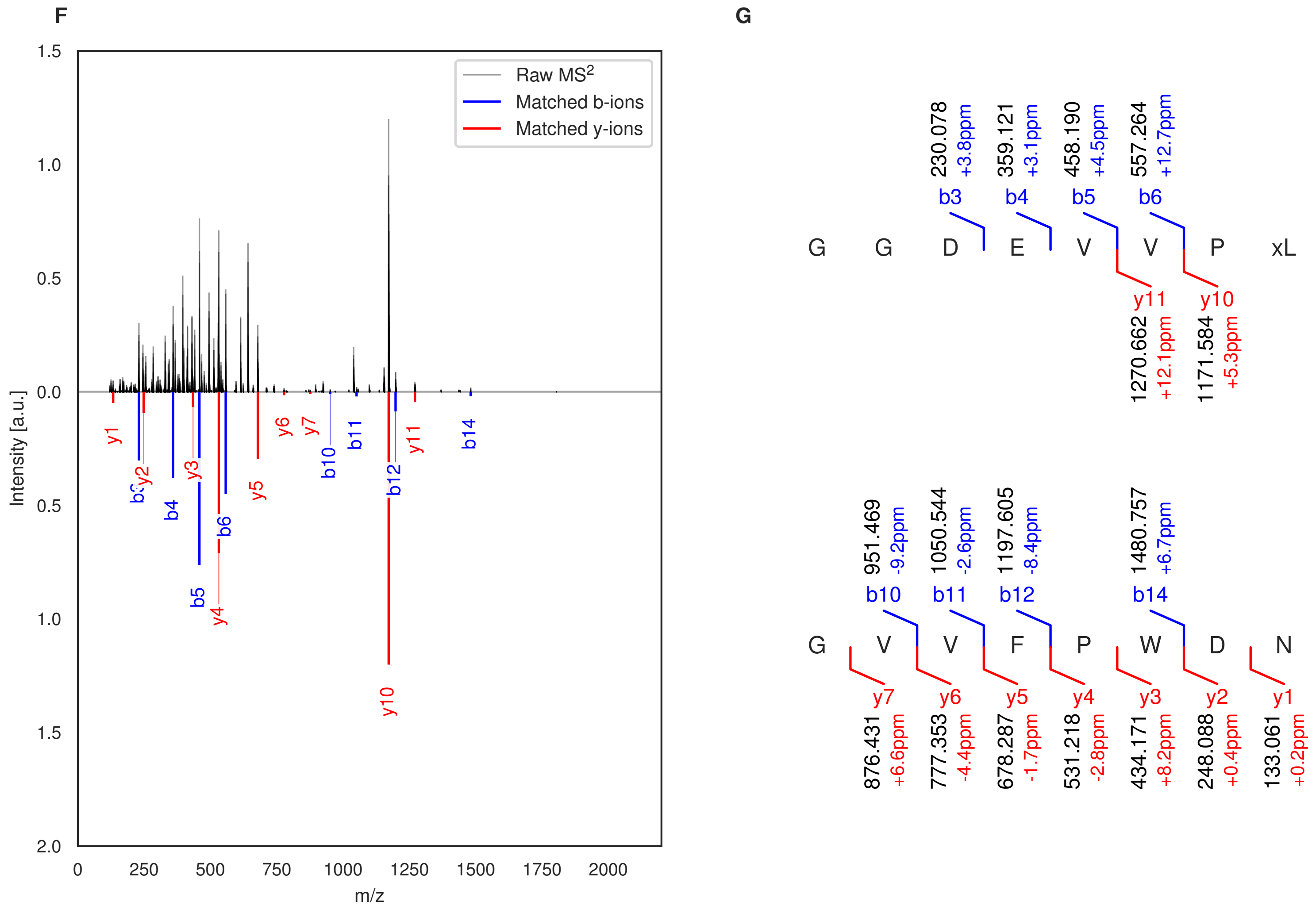

**Figure S51: Modification of the TJ-C-684 precursor peptide by PlpXY.** LC-MS analysis of IMAC-purified precursor peptides from *E. coli* BL21(DE3)Tuner containing splicease-encoding plasmid TJ-C-559 (PlpXY) and precursor peptide-encoding plasmid TJ-C-684 (Fh8-chaos-PlpA3-TEV-PlpA3-Boc-Trunc) after expression at 24 °C for 22 h in 30 mL cultures after digestion with TEV. Data from n=1 independent biological replicates. Spliced amino acids have an x as a prefix. Spliced refers to a single tyramine excision. A representative experimental replicate is shown in panels **A, B.**

**A** Close-up plot of the representative extracted ion chromatogram (EIC) of the various peptide states; the blue trace corresponds to the Unmodified peptide; the orange trace corresponds to the Spliced peptide. **B** Close-up plot of the m/z values of the peptides detected by the mass spectrometer, with the shaded colored areas corresponding to where the expected m/z values for the peptide m/z traces should occur, with the observed spectrum plotted in black. The Unmodified, Spliced peptide(s) were found in the mass spectrum. **C** Bar plot showing the relative abundance of the various peptide states at charge z=2; the bars represent the average relative abundances normalized to total peak areas, with each individual experimental replicate value plotted as a black dot. **D** Mirrored $MS^2$ plot for the Unmodified peptide. In this mirror plot, the full fragmentation spectrum is plotted in black pointing upward, and the matched fragment b ions (blue) and y ions (red) are plotted pointing downward, mirrored against it, so that each matched fragment can be compared directly with the measured spectrum. **E** $MS^2$ spectrum of the fragments corresponding to the Unmodified peptide sequence, with matched b (blue, above) and y ions (red, below) plotted. **F** Mirrored $MS^2$ plot for the Spliced peptide. In this mirror plot, the full fragmentation spectrum is plotted in black pointing upward, and the matched fragment b ions (blue) and y ions (red) are plotted pointing downward, mirrored against it, so that each matched fragment can be compared directly with the measured spectrum. **G** $MS^2$ spectrum of the fragments corresponding to the Spliced peptide sequence, with matched b (blue, above) and y ions (red, below) plotted.

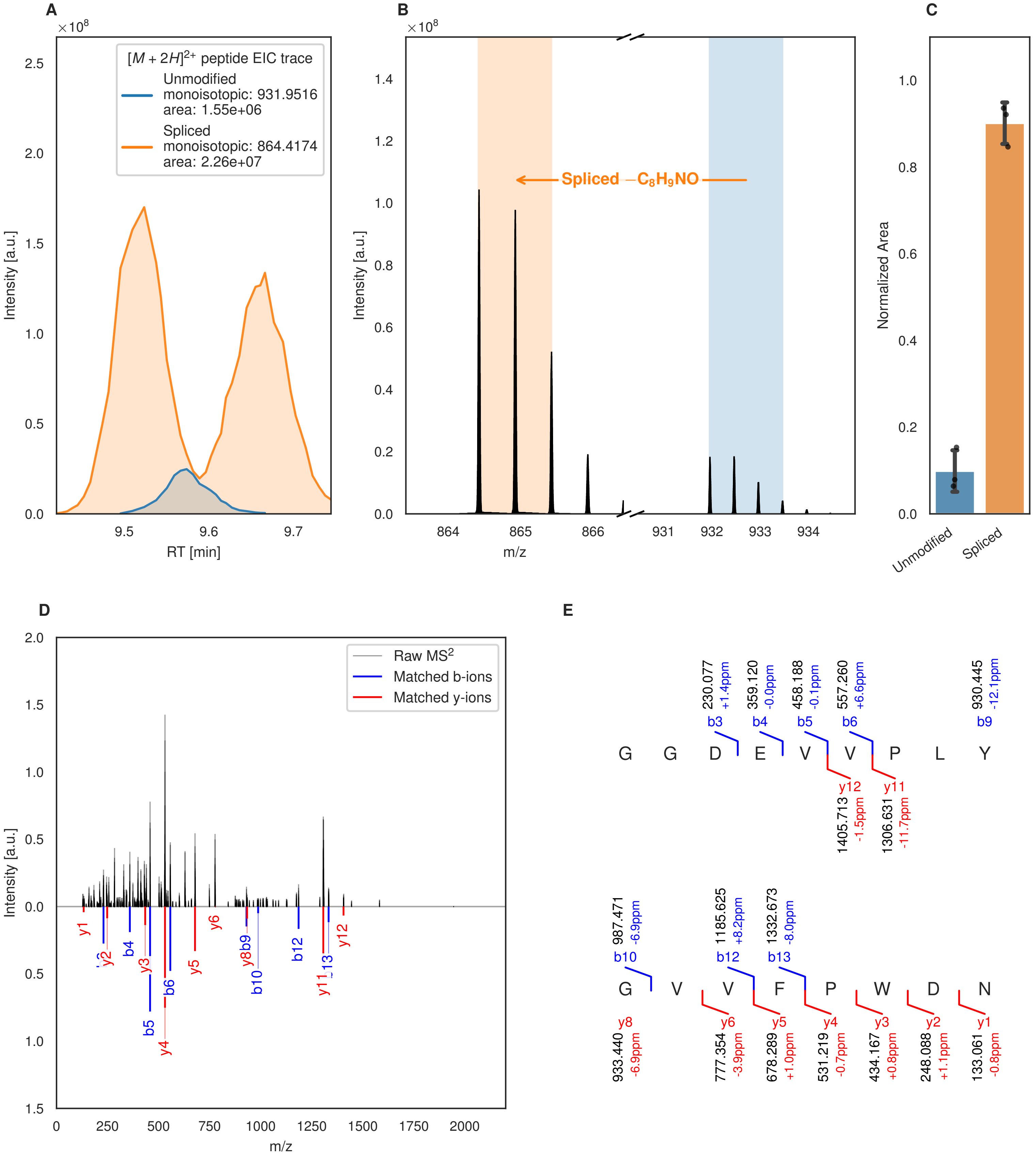

A
B
C
D
E
[M + 2H]2+ peptide EIC trace
Unmodified
monoisotopic: 931.9516
area: 1.55e+06
Spliced
monoisotopic: 864.4174
area: 2.26e+07
Intensity [a.u.]
RT [min]
Spliced −C8H9NO
m/z
Normalized Area
Unmodified
Spliced
Raw MS2
Matched b-ions
Matched y-ions
G G D E V V P L Y
G V V F P W D N

Figure S52 (figure continued from the previous page).

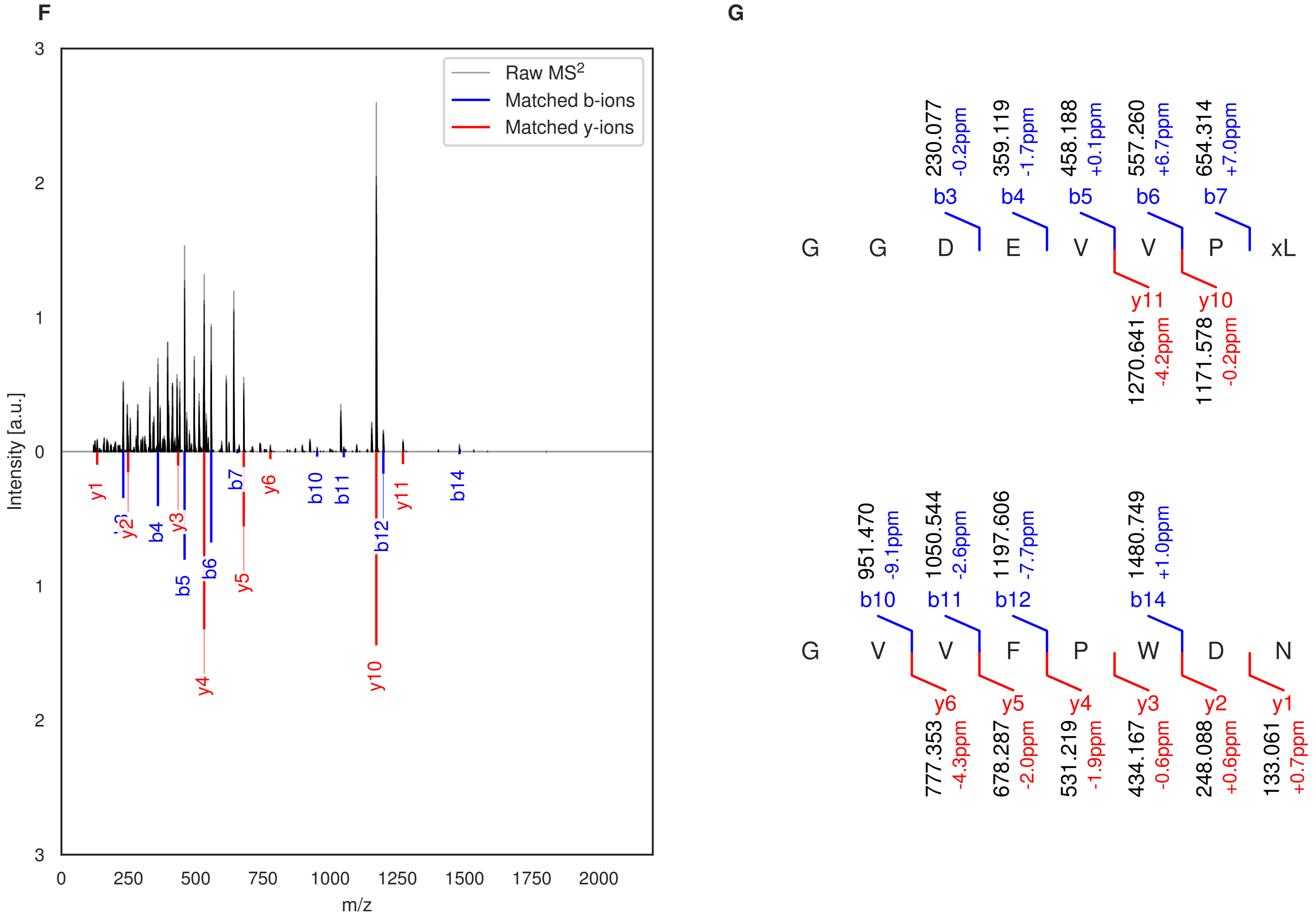

**Figure S52: Modification of the TJ-C-685 precursor peptide by PlpXY.** LC-MS analysis of IMAC-purified precursor peptides from *E. coli* BL21(DE3)Tuner containing splicease-encoding plasmid TJ-C-559 (PlpXY) and precursor peptide-encoding plasmid TJ-C-685 (Fh8-PlpA3-TEV-PlpA3-Boc-Trunc) after expression at 24 °C for 22 h in 30 mL cultures after digestion with TEV. Data from n=3 independent biological replicates. Spliced amino acids have an x as a prefix. Spliced refers to a single tyramine excision. A representative experimental replicate is shown in panels **A, B.**

**A** Close-up plot of the representative extracted ion chromatogram (EIC) of the various peptide states; the blue trace corresponds to the Unmodified peptide; the orange trace corresponds to the Spliced peptide. **B** Close-up plot of the m/z values of the peptides detected by the mass spectrometer, with the shaded colored areas corresponding to where the expected m/z values for the peptide m/z traces should occur, with the observed spectrum plotted in black. The Unmodified, Spliced peptide(s) were found in the mass spectrum. **C** Bar plot showing the relative abundance of the various peptide states at charge z=2; the bars represent the average relative abundances normalized to total peak areas, with each individual experimental replicate value plotted as a black dot. **D** Mirrored $MS^2$ plot for the Unmodified peptide. In this mirror plot, the full fragmentation spectrum is plotted in black pointing upward, and the matched fragment b ions (blue) and y ions (red) are plotted pointing downward, mirrored against it, so that each matched fragment can be compared directly with the measured spectrum. **E** $MS^2$ spectrum of the fragments corresponding to the Unmodified peptide sequence, with matched b (blue, above) and y ions (red, below) plotted. **F** Mirrored $MS^2$ plot for the Spliced peptide. In this mirror plot, the full fragmentation spectrum is plotted in black pointing upward, and the matched fragment b ions (blue) and y ions (red) are plotted pointing downward, mirrored against it, so that each matched fragment can be compared directly with the measured spectrum. **G** $MS^2$ spectrum of the fragments corresponding to the Spliced peptide sequence, with matched b (blue, above) and y ions (red, below) plotted.

### 2. Parts, plasmids, and oligonucleotides used in this study

**Table S1: Overhangs used for EcoFlex[2] compatible Golden Gate assembly.** Note that the last nucleotides of γ and δ overhangs (lower case letters, underlined) are the first nucleotides of the ATG start codon of the protein sequence encoded. The CGGA overhang has the last three nucleotides in frame, encoding a GG scar that is common between leader and core peptides in RiPPs. In subsequent part descriptions, the overhangs are not included in the sequence except for the initial A nucleotide in the case of start codons.

| **Overhang name** | **4 bp overhang sequence** | **Source** |
|---|---|---|
| α | CTAT | EcoFlex standard |
| β | GTAC | EcoFlex standard |
| γ | TAA<u>a</u> | EcoFlex standard |
| δ | CAT<u>a</u> | EcoFlex standard |
| δ’ | CGGA | Custom |
| ε | TCGA | EcoFlex standard |
| ζ | TGTT | EcoFlex standard |

**Table S2: Promoters and composite parts used in this work**. Insulators are labeled in green, tags labeled in blue.

| Part name | Part sequence | Ends | Source |
|---|---|---|---|
| T7-lacO-RiboJ53 | TAATACGACTCACTATAGGGAGGAATTGTGAGCGGATA<br>ACAATTCCAGCGGTCAACGCATGTGCTTTGCGTTCTGA<br>TGAGACAGTGATGTCGAAACCGCCTCTACAAATAATTT<br>TGTTTAA | α-β | Custom |
| T7-lacO-RBS-His6-Thr | CCCGCGAAATTAATACGACTCACTATAGGGGAATTGTG<br>AGCGGATAACAATTCCCCTCTAGAAATAATTTTGTTTA<br>ACTTTAAGAAGGAGATATACCATGGGCAGCAGCCATCA<br>TCATCATCATCACAGCAGCGGCCTGGTGCCGCGCGGCA<br>GC | α-δ | EcoFlex kit[2] |
| T7-lacO-PP1-RSTN1 | TAATACGACTCACTATAGGGAGGAATTGTGAGCGGATA<br>ACAATTACCCAACACCACCAGCAAGCCTAAGGAGGAGA<br>AATATGTCATATTACCACCATCACCATCATCACGGGTC<br>CCTGCAGGACTCAGAAGTCAATCAAGAAGCTAAGCCAG<br>AGGTCAAGCCAGAAGTCAAGCCTGAGACTCACATCAAT<br>TTAAAGGTGTCCGATGGATCTTCAGAGATCTTCTTCAA<br>GATCAAAAAGACCACTCCTTTAAGAAGGCTGATGGAAG<br>CGTTCGCTAAAAGACAGGGTAAGGAAATGGACTCCTTA<br>AGATTCTTGTACGACGGTATTAGAATTCAAGCTGATCA<br>GGCCCCTGAAGATTTGGACATGGAGGATAACGATATTA<br>TTGAGGCTCACCGCGAACAGATTGGAGGTCATCACCAT<br>CACCACCATGGATATGATATTAGCACAGGT | α-δ | Voigt lab, modified[3] |
| T7-lacO-ElvJ-100k-Fh8-chaos | TAATACGACTCACTATAGGGAGGAATTGTGAGCGGATA<br>ACAATTAGCCCCATAGGGTGGTGTGTACCACCCCTGAT<br>GAGTCCAAAAGGACGAAATGGGGCCTCTACAAATAATT<br>TTGTTTAAAACTGAGTAAGGAGGTCTTAATGAGTTACT<br>ACCATCACCACCATCATCATGGTTCGCCCAGCGTACAG<br>GAAGTAGAAAAACTCCTACATGTACTGGATAGAAATGG<br>TGACGGAAAAGTTAGCGCGGAAGAACTGAAGGCATTCG<br>CGGATGACAGCAAATGCCCGTTAGATTCGAATAAGATT<br>AAAGCGTTCATCAAGGAACACGACAAAAATAAAGACGG<br>CAAACTCGATTTAAAAGAGCTGGTGAGCATCCTTTCCA<br>GTTCGTCCAGCAATAACAACAACAATAATAATAATAAT<br>AACGAATTTGAGAATCTGTACTTTCAAGGTTGC | α-δ | Custom, RBS calc,[4–9] Kim et al[10] |
| T7-lacO-ElvJ-100k-Fh8 | TAATACGACTCACTATAGGGAGGAATTGTGAGCGGATA<br>ACAATTAGCCCCATAGGGTGGTGTGTACCACCCCTGAT<br>GAGTCCAAAAGGACGAAATGGGGCCTCTACAAATAATT<br>TTGTTTAAAACTGAGTAAGGAGGTCTTAATGAGTTACT | α-δ | Custom, RBS calc,[4–9] |

| | | | |
|---|---|---|---|
| | ACCATCACCACCATCATCATGGTTCGCCCAGCGTACAG<br>GAAGTAGAAAAACTCCTACATGTACTGGATAGAAATGG<br>TGACGGAAAAGTTAGCGCGGAAGAACTGAAGGCATTCG<br>CGGATGACAGCAAATGCCCGTTAGATTCGAATAAGATT<br>AAAGCGTTCATCAAGGAACACGACAAAAATAAAGACGG<br>CAAACTCGATTTAAAAGAGCTGGTGAGCATCCTTTCCA<br>GTCATCACCACCATCATCATGGTTCGTTTGAGAATCTG<br>TACTTTCAAGGTTGC | | Costa et al[11–13] |
| T7-lacO-ElvJ-30k-Pfu-MBP | TAATACGACTCACTATAGGGAGGAATTGTGAGCGGATA<br>ACAATTAGCCCCATAGGGTGGTGTGTACCACCCCTGAT<br>GAGTCCAAAAGGACGAAATGGGGCCTCTACAAATAATT<br>TTGTTTAAAAATATAGTCTAAGAAGGTTTTAATGAGTT<br>ACTACCATCACCACCATCATCATGGTAAGATTGAGGAA<br>GGGAAAGTTGTTATTTGGCATGCAATGCAACCGAATGA<br>ACTTGAGGTTTTCCAGTCCTTAGCCGAGGAATACATGG<br>CCTTGTGTCCCGAAGTCGAGATCGTGTTCGAGCAGAAA<br>CCCAATCTGGAAGATGCCTTGAAAGCAGCCATCCCTAC<br>AGGCCAGGGTCCGGATCTGTTCATCTGGGCTCACGACT<br>GGATAGGTAAATTCGCTGAAGCTGGACTGCTGGAACCG<br>ATTGATGAATACGTTACCGAGGACCTGTTAAATGAATT<br>CGCGCCCATGGCCCAAGACGCTATGCAGTACAAAGGTC<br>ACTACTACGCCCTGCCTTTCGCGGCTGAAACCGTGGCC<br>ATCATCTATAACAAGGAGATGGTTTCTGAGCCACCGAA<br>AACGTTCGACGAAATGAAAGCAATTATGGAAAAGTATT<br>ACGACCCTGCTAACGAGAAATACGGCATCGCCTGGCCC<br>ATTAACGCTTACTTTATTAGCGCGATAGCCCAGGCCTT<br>TGGAGGCTACTACTTTGACGATAAAACCGAACAACCTG<br>GCCTGGATAAGCCGGAAACAATCGAGGGCTTTAAATTT<br>TTCTTCACCGAGATCTGGCCGTACATGGCTCCCACCGG<br>TGATTACAACACCCAACAGTCCATCTTCCTGGAAGGTC<br>GAGCTCCCATGATGGTTAACGGTCCGTGGAGCATTAAT<br>GATGTAAAAAAAGCAGGAATTAATTTCGGCGTTGTACC<br>CTTACCGCCCATCATCAAAGACGGAAAAGAATACTGGC<br>CACGTCCGTACGGAGGCGTGAAGCTGATCTACTTTGCG<br>GCCGGTATTAAAAACAAAGATGCCGCTTGGAAATTCGC<br>TAAATGGTTAACCACCTCGGAGGAGAGTATTAAAACCT<br>TAGCACTGGAACTGGGTTATATCCCTGTGCTCACGAAA<br>GTATTAGACGATCCTGAAATTAAAAACGACCCTGTCAT<br>TTACGGATTTGGTCAGGCAGTTCAGCATGCCTATTTGA<br>TGCCCAAATCTCCGAAAATGTCCGCTGTGTGGGGTGGC | α-δ | Custom, RBS calc, Luke et al[14] |

| | |
|---|---|
| | GTAGACGGAGCGATTAACGAAATCCTTCAGGACCCTCA<br>GAATGCAGATATTGAGGGCATTCTTAAAAAGTACCAGC<br>AGGAAATTCTCAATAATATGCAGGGCGGTTCGTTTGAG<br>AATCTGTACTTTCAAGGTTGC |

**Table S3: Maturase genes used in this work.** In the case of non-basic parts, the maturases are colored in magenta and the accessory RRE protein in cyan.

| Part name | Part sequence | Ends | Source |
|---|---|---|---|
| HypX | ATGAAGAAGTACCGACGGATATCATATGCGGTATGGGA<br>GATAACACTCAAATGCAACCTGGCCTGCCAGCACTGCG<br>GCAGCCGGGCCGGAGATGCACGTGCAGAGGAACTGTCA<br>ACAGAAGAGGCGCTGGATATGGTTAAGCAGTTGGCTGA<br>GGTTGGTATAAAAGAGGTGACACTGATAGGCGGTGAGG<br>CGTTCCTTCGACCCGATTGGCTGGAGATAGCACAAGCC<br>ATCACGGAAGCCGGCATGTTGTGCGGAATGACAACGGG<br>CGGCTTCGGCATCTCTTTAGACACCGCCCGCCGCATGA<br>AAGAGGCCGGTATCAAGGTGGTCAGCGTATCTGTGGAC<br>GGATTAGAGGAAACGCACAACCGTCTGCGAGGCAAGCA<br>GGGAAGTTGGCAGTGGGCCTTCAAGACGATGTCGCACT<br>TAAAAGAGGTTGGAATCCGCTTCGGGTGCAACACCCAG<br>ATCAATCGGCTGAGCGCTCCCGAGTTCCCGTTGATTTA<br>TGAGCGAATTCGTGATGCGGGCATCTTTGCGTGGCAAA<br>TCCAACTGACTGTTCCTATGGGGAACGCCGCCGACAAC<br>AGCGAAATCCTTCTCCAGCCCTACGAGCTGTTGGACCT<br>GTACCCTATGATAGCGCGCGTGGTCAAGCGTGCCCGGA<br>AAGAGGGCGTACAGGTGCAGCCAGGTAACAACATCGGC<br>TACTACGGTCCGTACGAGCGACTGTTGCGTGGCCAAGG<br>GGAAGCTTGGAAGTTCTGGCAAGGGTGTACCGCGGGCT<br>TGAGCAGTCTTGGGATAGAGGCCGACGGTTCTATCAAG<br>GGTTGCCCGTCGCTGCCCACGAGCGCGTATACCGGCGG<br>GAACATTCGTGACCACTCGCTGAGAAAGATTGTTGAGG<br>AAACTGAGGAGTTACGGTTCAACCTCGGCGCGGACACT<br>GCGAAGGGTACTGAGCACTTGTGGGGCTTCTGCAAAAC<br>ATGTGAGTTTGCCGAGTTGTGCAGAGGCGGATGCACGT<br>GGACAGCCCATGTGTTCTTCAATAAACGCGGGAATAAC<br>CCGTACTGCCATCATCGTGCGTTAGCACAGGAGAAGCA<br>GGGCATTCGCGAGCGCGTCGTACTGAAGTACAAAGCTG<br>AAGGTAACCCGTTCGATAACGGGGAATTCTCTCTGATC<br>GAGGAGCCTATCAACGCGTGCTGGCCTGAGAATGACCC<br>ACTGCACTTCACGGCGGATAGAATACAGTGGCCGGAGT<br>CGTGGCAAGAGGAACAGGAACCGGTACCGACGTCTATC<br>GCAACGTCTAACTAATAA | δ-ε | Custom, optimized |
| HypY | ATGCTGGCGAACAGTAATTCACCGGAAGATCGCCCCGT<br>GGCCGACGGCCAGACCCTGAACTCGGAAAGCAGCGAGT<br>ATAACAGCTCGAATCGCGAAAATTCCTCAAGCCCGCTT | δ-ε | Custom, optimized |

| | | | |
|---|---|---|---|
| | CTGCCACGAAGTCCGTGGAATTCTAAACTGGCTTTTCT<br>CCGTGCGATCTTCAAGGCAAAGAAAGCCTTAGATCGTA<br>TTGAGAACTTGGAACCTAAAAAACATTAATAA | | |
| PlpX | ATGACTAAAAAATACAGACGAGTTAGTTATGCAGTTTG<br>GGAAATTACCTTGAAATGCAATCTAGCTTGTAGTCACT<br>GTGGTTCGAGAGCAGGGCAGGCAAGAACCAAGGAACTA<br>TCTACAGAAGAAGCTTTTAATCTGGTTCGGCAACTAGC<br>CGATGTAGGAATCAAAGAGGTTACTCTAATCGGTGGCG<br>AAGCCTTTATGCGCTCTGATTGGCTAGAAATTGCCAAG<br>GCTGTTACTGAGGCAGGGATGATCTGCGGTATGACTAC<br>AGGTGGATTTGGTGTCAGTTTGGAAACTGCCAGAAAAA<br>TGAAAGAAGCTGGAATTAAAACAGTTTCTGTATCTATC<br>GATGGTGGCATACCAGAAACCCACGATCGCCAGCGAGG<br>GAAAAAAGGTGCTTGGCATTCTGCTTTTAGAACCATGA<br>GCCATCTAAAAGAAGTCGGCATCTATTTTGGCTGCAAT<br>ACCCAGATAAACCGTTTATCTGCCTCTGAATTCCCAAT<br>AATTTACGAACGAATAAGGGACGCTGGAGCAAGAGCTT<br>GGCAGATTCAATTAACTGTACCTATGGGTAATGCGGCA<br>GATAATGCAGACATGTTATTGCAACCATACGAACTATT<br>AGATATTTATCCCATGTTAGCTCGTGTTGCTAAACGAG<br>CTAAACAGGAAGGTGTTCGCATACAGGCGGGAAATAAT<br>ATTGGCTATTATGGCCCTTATGAAAGACTGCTGCGTGG<br>TAGTGATGAATGGACATTTTGGCAGGGTTGCGGAGCGG<br>GTTTAAATACCTTGGGTATCGAAGCTGATGGCAAAATT<br>AAAGGTTGTCCTTCTTTACCTACGGCTGCTTATACGGG<br>CGGTAATATCCGCGATCGCCCTTTAAGAGAAATAGTCG<br>AACAGACTGAAGAGCTTAAATTTAATCTGAAGGCTGGG<br>ACTGAACAGGGCACAGACCACATGTGGGGATTTTGTAA<br>AACCTGTGAATTTGCTGAACTCTGTCGAGGTGGTTGTT<br>CTTGGACGGCTCATGTCTTCTTTGATCGCCGTGGGAAT<br>AATCCCTACTGCCATCATCGTGCTTTGAAACAGGCACA<br>AAAAGACATCAGAGAAAGATTCTATTTAAAAGTAAAAG<br>CAAAAGGGAATCCTTTTGATAATGGGGAATTTGTCATT<br>ATAGAAGAACCTTTCAACGCACCTTTGCCAGAGAACGA<br>TTTGCTTCATTTTAATAGCGATCACATTCAGTGGCCAG<br>AAAACTGGCAAAATTCTGAATCTGCTTACGCTTTAGCA<br>AAGTAATAA | δ-ε | Morinaka et al.[1] |
| PlpY | ATGAACTCTAATCAAATACCAAATAAAGTAGCCACAGC<br>AGCTCAAAAATCAGATGATTCCAGCTCGGTTTTACCTC<br>GTCAGGGTTGGCAAGACAAGCAAGCTTTTATCAAAGCA | δ-ε | Morinaka et al.[1] |

| | | | |
|---|---|---|---|
| | TTAATTAAAGCAAAACAAAGTTTAGAAATTGCTGAAAT<br>TAGCAATTTTCTGACATAATAA | | |
| 20k-HypX-B34-HypY | AGTTAATACCCTATCGCTAGGAGGAAGAAATGAAGAAG<br>TACCGACGGATATCATATGCGGTATGGGAGATAACACT<br>CAAATGCAACCTGGCCTGCCAGCACTGCGGCAGCCGGG<br>CCGGAGATGCACGTGCAGAGGAACTGTCAACAGAAGAG<br>GCGCTGGATATGGTTAAGCAGTTGGCTGAGGTTGGTAT<br>AAAAGAGGTGACACTGATAGGCGGTGAGGCGTTCCTTC<br>GACCCGATTGGCTGGAGATAGCACAAGCCATCACGGAA<br>GCCGGCATGTTGTGCGGAATGACAACGGGCGGCTTCGG<br>CATCTCTTTAGACACCGCCCGCCGCATGAAAGAGGCCG<br>GTATCAAGGTGGTCAGCGTATCTGTGGACGGATTAGAG<br>GAAACGCACAACCGTCTGCGAGGCAAGCAGGGAAGTTG<br>GCAGTGGGCCTTCAAGACGATGTCGCACTTAAAAGAGG<br>TTGGAATCCGCTTCGGGTGCAACACCCAGATCAATCGG<br>CTGAGCGCTCCCGAGTTCCCGTTGATTTATGAGCGAAT<br>TCGTGATGCGGGCATCTTTGCGTGGCAAATCCAACTGA<br>CTGTTCCTATGGGGAACGCCGCCGACAACAGCGAAATC<br>CTTCTCCAGCCCTACGAGCTGTTGGACCTGTACCCTAT<br>GATAGCGCGCGTGGTCAAGCGTGCCCGGAAAGAGGGCG<br>TACAGGTGCAGCCAGGTAACAACATCGGCTACTACGGT<br>CCGTACGAGCGACTGTTGCGTGGCCAAGGGGAAGCTTG<br>GAAGTTCTGGCAAGGGTGTACCGCGGGCTTGAGCAGTC<br>TTGGGATAGAGGCCGACGGTTCTATCAAGGGTTGCCCG<br>TCGCTGCCCACGAGCGCGTATACCGGCGGGAACATTCG<br>TGACCACTCGCTGAGAAAGATTGTTGAGGAAACTGAGG<br>AGTTACGGTTCAACCTCGGCGCGGACACTGCGAAGGGT<br>ACTGAGCACTTGTGGGGCTTCTGCAAAACATGTGAGTT<br>TGCCGAGTTGTGCAGAGGCGGATGCACGTGGACAGCCC<br>ATGTGTTCTTCAATAAACGCGGGAATAACCCGTACTGC<br>CATCATCGTGCGTTAGCACAGGAGAAGCAGGGCATTCG<br>CGAGCGCGTCGTACTGAAGTACAAAGCTGAAGGTAACC<br>CGTTCGATAACGGGGAATTCTCTCTGATCGAGGAGCCT<br>ATCAACGCGTGCTGGCCTGAGAATGACCCACTGCACTT<br>CACGGCGGATAGAATACAGTGGCCGGAGTCGTGGCAAG<br>AGGAACAGGAACCGGTACCGACGTCTATCGCAACGTCT<br>AACTAATAAGTACAGAAAGAGGAGAAATACCATATGCT<br>GGCGAACAGTAATTCACCGGAAGATCGCCCCGTGGCCG<br>ACGGCCAGACCCTGAACTCGGAAAGCAGCGAGTATAAC<br>AGCTCGAATCGCGAAAATTCCTCAAGCCCGCTTCTGCC | β-ε | Custom, RBS calc, optimized |

| | | | |
|---|---|---|---|
| | ACGAAGTCCGTGGAATTCTAAACTGGCTTTTCTCCGTG CGATCTTCAAGGCAAAGAAAGCCTTAGATCGTATTGAG AACTTGGAACCTAAAAAACATTAATAA | | |
| 20k-PlpX-B34-PlpY | ACAGCAGAAGATTAAGGAGTTCAAATAATGACTAAAAA ATACAGACGAGTTAGTTATGCAGTTTGGGAAATTACCT TGAAATGCAATCTAGCTTGTAGTCACTGTGGTTCGAGA GCAGGGCAGGCAAGAACCAAGGAACTATCTACAGAAGA AGCTTTTAATCTGGTTCGGCAACTAGCCGATGTAGGAA TCAAAGAGGTTACTCTAATCGGTGGCGAAGCCTTTATG CGCTCTGATTGGCTAGAAATTGCCAAGGCTGTTACTGA GGCAGGGATGATCTGCGGTATGACTACAGGTGGATTTG GTGTCAGTTTGGAAACTGCCAGAAAAATGAAAGAAGCT GGAATTAAAACAGTTTCTGTATCTATCGATGGTGGCAT ACCAGAAACCCACGATCGCCAGCGAGGGAAAAAAGGTG CTTGGCATTCTGCTTTTAGAACCATGAGCCATCTAAAA GAAGTCGGCATCTATTTTGGCTGCAATACCCAGATAAA CCGTTTATCTGCCTCTGAATTCCCAATAATTTACGAAC GAATAAGGGACGCTGGAGCAAGAGCTTGGCAGATTCAA TTAACTGTACCTATGGGTAATGCGGCAGATAATGCAGA CATGTTATTGCAACCATACGAACTATTAGATATTTATC CCATGTTAGCTCGTGTTGCTAAACGAGCTAAACAGGAA GGTGTTCGCATACAGGCGGGAAATAATATTGGCTATTA TGGCCCTTATGAAAGACTGCTGCGTGGTAGTGATGAAT GGACATTTTGGCAGGGTTGCGGAGCGGGTTTAAATACC TTGGGTATCGAAGCTGATGGCAAAATTAAAGGTTGTCC TTCTTTACCTACGGCTGCTTATACGGGCGGTAATATCC GCGATCGCCCTTTAAGAGAAATAGTCGAACAGACTGAA GAGCTTAAATTTAATCTGAAGGCTGGGACTGAACAGGG CACAGACCACATGTGGGGATTTTGTAAAACCTGTGAAT TTGCTGAACTCTGTCGAGGTGGTTGTTCTTGGACGGCT CATGTCTTCTTTGATCGCCGTGGGAATAATCCCTACTG CCATCATCGTGCTTTGAAACAGGCACAAAAAGACATCA GAGAAAGATTCTATTTAAAAGTAAAAGCAAAAGGGAAT CCTTTTGATAATGGGGAATTTGTCATTATAGAAGAACC TTTCAACGCACCTTTGCCAGAGAACGATTTGCTTCATT TTAATAGCGATCACATTCAGTGGCCAGAAAACTGGCAA AATTCTGAATCTGCTTACGCTTTAGCAAAGTAATAAGT ACAGAAAGAGGAGAAATACCATATGAACTCTAATCAAA TACCAAATAAAGTAGCCACAGCAGCTCAAAAATCAGAT GATTCCAGCTCGGTTTTACCTCGTCAGGGTTGGCAAGA | β-ε | Custom, RBS calc |

| | | | |
|---|---|---|---|
| | CAAGCAAGCTTTTATCAAAGCATTAATTAAAGCAAAAC<br>AAAGTTTAGAAATTGCTGAAATTAGCAATTTTCTGACA<br>TAATAA | | |
| 20k-HypX-<br>45k-HypY | AGTTAATACCCTATCGCTAGGAGGAAGAAATGAAGAAG<br>TACCGACGGATATCATATGCGGTATGGGAGATAACACT<br>CAAATGCAACCTGGCCTGCCAGCACTGCGGCAGCCGGG<br>CCGGAGATGCACGTGCAGAGGAACTGTCAACAGAAGAG<br>GCGCTGGATATGGTTAAGCAGTTGGCTGAGGTTGGTAT<br>AAAAGAGGTGACACTGATAGGCGGTGAGGCGTTCCTTC<br>GACCCGATTGGCTGGAGATAGCACAAGCCATCACGGAA<br>GCCGGCATGTTGTGCGGAATGACAACGGGCGGCTTCGG<br>CATCTCTTTAGACACCGCCCGCCGCATGAAAGAGGCCG<br>GTATCAAGGTGGTCAGCGTATCTGTGGACGGATTAGAG<br>GAAACGCACAACCGTCTGCGAGGCAAGCAGGGAAGTTG<br>GCAGTGGGCCTTCAAGACGATGTCGCACTTAAAAGAGG<br>TTGGAATCCGCTTCGGGTGCAACACCCAGATCAATCGG<br>CTGAGCGCTCCCGAGTTCCCGTTGATTTATGAGCGAAT<br>TCGTGATGCGGGCATCTTTGCGTGGCAAATCCAACTGA<br>CTGTTCCTATGGGGAACGCCGCCGACAACAGCGAAATC<br>CTTCTCCAGCCCTACGAGCTGTTGGACCTGTACCCTAT<br>GATAGCGCGCGTGGTCAAGCGTGCCCGGAAAGAGGGCG<br>TACAGGTGCAGCCAGGTAACAACATCGGCTACTACGGT<br>CCGTACGAGCGACTGTTGCGTGGCCAAGGGGAAGCTTG<br>GAAGTTCTGGCAAGGGTGTACCGCGGGCTTGAGCAGTC<br>TTGGGATAGAGGCCGACGGTTCTATCAAGGGTTGCCCG<br>TCGCTGCCCACGAGCGCGTATACCGGCGGGAACATTCG<br>TGACCACTCGCTGAGAAAGATTGTTGAGGAAACTGAGG<br>AGTTACGGTTCAACCTCGGCGCGGACACTGCGAAGGGT<br>ACTGAGCACTTGTGGGGCTTCTGCAAAACATGTGAGTT<br>TGCCGAGTTGTGCAGAGGCGGATGCACGTGGACAGCCC<br>ATGTGTTCTTCAATAAACGCGGGAATAACCCGTACTGC<br>CATCATCGTGCGTTAGCACAGGAGAAGCAGGGCATTCG<br>CGAGCGCGTCGTACTGAAGTACAAAGCTGAAGGTAACC<br>CGTTCGATAACGGGGAATTCTCTCTGATCGAGGAGCCT<br>ATCAACGCGTGCTGGCCTGAGAATGACCCACTGCACTT<br>CACGGCGGATAGAATACAGTGGCCGGAGTCGTGGCAAG<br>AGGAACAGGAACCGGTACCGACGTCTATCGCAACGTCT<br>AACTAATAACAGCGACTATATAAGGAGGTAATTTAATG<br>CTGGCGAACAGTAATTCACCGGAAGATCGCCCCGTGGC<br>CGACGGCCAGACCCTGAACTCGGAAAGCAGCGAGTATA | β-ε | Custom,<br>RBS calc,<br>optimized |

| | |
|---|---|
| | ACAGCTCGAATCGCGAAAATTCCTCAAGCCCGCTTCTG<br>CCACGAAGTCCGTGGAATTCTAAACTGGCTTTTCTCCG<br>TGCGATCTTCAAGGCAAAGAAAGCCTTAGATCGTATTG<br>AGAACTTGGAACCTAAAAAACATTAATAA |

**Table S4: Precursor peptide genes used in this work.** Leader-encoding regions are <u>underlined</u>, including the conserved **Gly-Gly** sequence separating the leader from the core that is indicated in bold font.

| Part name | Part sequence | Ends | Source |
|---|---|---|---|
| HypA1 | <u>ATGTCTCAAGAACAAGTTAAATCCTTTTATGAAAGATT</u><br><u>GGCACAAGATGAAGTTTTCAAAGCTCAAATTGAAGGAG</u><br><u>CTGAAAGTCCACAGGAATGCAGCCAAATTATGCGAGAA</u><br><u>GCTGGCTATGATTTTACTCAGGAAGAATTTGAAGCATT</u><br><u>TACAGCTCAGTTATTAGATTCTGATGATGCCAAAAATA</u><br><u>GTTTTGACGAGTTAAGCGAAAAGACGCTAGAAGCAGTT</u><br><u>TTG**GGCGGA**</u>TGCGGCGATCCTCAACCGGTGTATGGTGT<br>TATTCAGTAATAA | δ-ε | This work |
| HypA2 | <u>ATGTCTCAAGAACAAGTTAAAGCCATTTACAAAAGACT</u><br><u>GGCACAAGATGAAGTTTTCAAAGCTCAAATTGAAGGAG</u><br><u>CTGAAAGTCCACAGGAATGCAGCCAAATTATACGAGAA</u><br><u>GCTGGCTATGATTTCACTCAGGAAGAATTTGAAGCATT</u><br><u>TACAGCCCAGTTATTAGATTCTAGTGATGCCGAAAATA</u><br><u>GTTTTGAAGGGTTAAGCGAAAAGACACTAGAAGCAGTT</u><br><u>TTCGGGGTA**GGCGGA**</u>TCGGCGCAGGTTTATGGCACCAC<br>AACTAGCACCACGGTGCTGTTTTAATAA | δ-ε | This work |
| HypA3 | <u>ATGTCTCAAGAACACGTTAAAGCCTTTTACAAAAGACT</u><br><u>AGGACAAGATGAAGTTTTCAAAGCTCAAATTGAAGGAG</u><br><u>CTGAAAGTCCACAGGAATGTAGCCAAATTATACGAGAA</u><br><u>GCTGGCTATGATTTCACTCAAGAAGAATTTGAAGTATT</u><br><u>TACAGCCCAGTTATTAGATTCTAGTGATGCCAAAAATA</u><br><u>ATTTTGAAGAGTTAAGCGAAAAGGCACTAGAAGCAGTT</u><br><u>TTC**GGCGGA**</u>CGCCCGACCTACCAGTGCTATGGCGTGAT<br>TGAATAATAA | δ-ε | This work |
| HypA4 | <u>ATGTCTCAAAAAGCCGTTAAAGCCTTTTACGAAAGACT</u><br><u>GGCACAAGATGAAGTTTTTAAAGCTCAAATCGAAAAAG</u><br><u>CCAAAACTAAACAAGAATGTAGTCAAATTGTACGAGAA</u><br><u>GCGGGATACATTTTCACTCAAGAAGAATTTGAAGCATT</u><br><u>TACAGCTCAGTTATTAGATTCTAGTGCTGAATTGAATT</u><br><u>CGGAACAATTGGGTGAAGCCGAGCTGGAAGCTGCTGTT</u><br><u>**GGCGGA**</u>TTTCTGTTACCGATTAATCCGAACCCTATCCA<br>GACCTATGGCATTCCACCGATCCATTAATAA | δ-ε | This work |
| HypA5 | <u>ATGTCTCAAGAACACGTTAAAGCCTTTTACGAAAGACT</u><br><u>GGCACAAGATGAAATTTTTAAGGCTCAAATTGAAAAAG</u><br><u>CCAAAACTAAACAAGAATGTAGTCAAATTGTACGAGAA</u> | δ-ε | This work |

| | | | |
|---|---|---|---|
| | GCGGGATACATTTTCACTCAAGAAGAATTTGAAGCATT<br>TACAGCTCAGTTATTAGATTCTAGTGCTGAATTGAATT<br>CGGAACAATTGGGTGAAGCCGAGCTGGAAGCTGCTGTT<br>GGCGGATTTCTTCTGGAAAAAGTGATTGATTTGGTTCA<br>GGCGTATGGTCTGCCGCCTGGCTTATAATAA | | |
| HypA6 | ATGTCACAAGAACACGTTAAAGCTTTTTACGAAAGATT<br>GTCACAAGATGAAGTTTTTAAAGCTCAAATTGAAAAAG<br>CCAAAACTAAACAAGAATGTAGTCAAATCGTGCGAGAA<br>GCTGGTTATATCTTTACTCAAGAAGAATGGGAAGCATT<br>TACGGCTCAGTTATTAGATTCTAGTGCTGAATTCAATT<br>CAGGACAATTGGGTGAAGCCGAGCTGGAAGCTGCTGTT<br>**GGCGGA**TTCCTGAAAGAAAAGTTAATTGGTGATCCGAT<br>TCCTATGTATGGCATCAGCCCGATCGAGTTTTGGTAAT<br>AA | δ-ε | This work |
| HypA7 | ATGACCATAGCAAATGTTGAAGCCTTCTATAAAAGGCT<br>GGCAGTTGATGAAGCTTTTCGCTCCAAAATTCAAAAAG<br>CTGGAAGCAAGCAAAAATGTCGCCAAATCGTGCGAGAG<br>GCTGGTTATGATTTCACTCAAGAAGAATTTGAAGCATT<br>TACATCTCAGTTATTGGATTCAAATACTGAAGTAAATC<br>TGGGAGAATTAAGCGAAATAGAGCTTGAAGCTGCTGTT<br>**GGCGGA**ATTAACGCGTTATTCCAGGAAAATCGCATCAA<br>CTTTGTGCCGCCTTATGGCCTGCCGCCAGATTGGGTTC<br>CGTAATAA | δ-ε | This work |
| HypAc | ATGAGCCAGGAACATGTCAAAGCGTTCTATGAACGTCT<br>TGCGCAGGATGAGGTTTTTAAAGCGCAAATCGAGAAAG<br>CTGAATCAAAGCAGGAATGCAGCCAGATTGTGCGCGAG<br>GCCGGCTACGACTTTACGCAGGAGGAATTCGAGGCGTT<br>TACCGCACAACTGTTAGATAGTTCCGCCGAAAAAAACT<br>CGTTCGAAGAATTGTCTGAAGCCGAACTGGAAGCAGCC<br>GTA**GGCGGA** | δ – δ’ | This work |
| HypAc-TEV | ATGAGCCAGGAACATGTCAAAGCGTTCTATGAACGTCT<br>TGCGCAGGATGAGGTTTTTAAAGCGCAAATCGAGAAAG<br>CTGAATCAAAGCAGGAATGCAGCCAGATTGTGCGCGAG<br>GCCGGCTACGACTTTACGCAGGAGGAATTCGAGGCGTT<br>TACCGCACAACTGTTAGATAGTTCCGCCGAAAAAAACT<br>CGTTCGAAGAATTGTCTGAAGCCGAACTGGAAGCAGCC<br>GTAGAGAACTTGTACTTTCAA**GGCGGA** | δ – δ’ | Codon optimized |
| PlpA3-TEV | ATGTCTATCGAAAGTGCAAAGGCGTTTTACCAACGCAT<br>GACCGATGATGCCTCGTTTCGTACCCCGTTCGAAGCCG<br>AATTAAGCAAAGAAGAACGCCAGCAACTGATTAAAGAC | δ – δ’ | This work |

| | | | |
|---|---|---|---|
| | TCAGGCTATGATTTCACTGCCGAGGAATGGCAGCAGGCGATGACGGAAATTCAGGCGGCTCGTAGCAACGAGGAACTGAATGAGGAAGAACTTGAGGCGATCGCAGAGAACTTGTACTTTCAA**GGCGGA** | | |
| PlpA2-core | GTGGACTTGAGCATTTTTGAGCTGCTTGATGAAGAACCGCTGTTCCCTATCCGCCCGTTATATGGCCTGCCAATTTAATAA | δ'- ε | This work |
| PlpA3-core | GCCGTCGCAGCGATGTATGGAGTTGTGTTTCCGTGGGATAACGAATTCCCTTGGCCGCGCTGGGGCGGTTAATAA | δ'- ε | This work |
| PlpA3-trunc-core | GCCGTCGCAGCGATGTATGGAGTTGTGTTTCCGTGGGATAACTAATAA | δ'- ε | This work |
| PlpA3-bocep | GATGAAGTTGTGCCGCTGTATGGAGTTGTGTTTCCGTGGGATAACGAATTCCCTTGGCCGCGCTGGGGCGGTTAATAA | δ'- ε | This work |
| PlpA3-bocep-trunc | GATGAAGTTGTGCCGCTGTATGGAGTTGTGTTTCCGTGGGATAACTAATAA | δ'- ε | This work |
| Artificial-1 | TGCGGACCACAGCCTGTTTACGGCGTTTTGCCATGGGCTTAATAA | δ'- ε | This work |
| Artificial-2 | TGCGGACCACAGCCTGTTTACGGCGTTATTTTGCCATGGGCTTAATAA | δ'- ε | This work |
| Artificial-3 | ATTGGTGATCCGATTCCTATGTATGGCATCAGCCCGATCGAGTTTTGGTAATAA | δ'- ε | This work |
| Artificial-4 | TGCGCCGTCGCAGCGATGTACGGCGTTGTGTTGCCATGGGCTTAATAA | δ'- ε | This work |

**Table S5:** Terminators used in this work.

| Part name | Part sequence | Ends | Source |
|---|---|---|---|
| BBa_B0015 | CCAGGCATCAAATAAAACGAAAGGCTCAGTCGAAAGACTGGGCCTTTCGTTTTATCTGTTGTTTGTCGGTGAACGCTCTCTACTAGAGTCACACTGGCTCACCTTCGGGTGGGCCTTTCTGCGTTTATA | ε-ζ | EcoFlex kit |
| BBa_B0012 | TCACACTGGCTCACCTTCGGGTGGGCCTTTCTGCGTTTATA | ε-ζ | EcoFlex kit |

**Table S6: Plasmid backbones used in this work.** Dropout marker in red corresponds to a constitutively expressed sfGFP cassette and a high copy pUC ori to increase copy number.

| Part name | Part sequence | AB | Source |
|---|---|---|---|
| pCDF-Spec-LacI | GCCAATAGGCGTATCACGAGGCAGAATTTCAGATAAAA<br>AAAATCCTTAGCTTTCGCTAAGGATGATTTCTGGGCGT<br>TGACAATGCGAAGTCAGTTCTATAGAGACCTAATAAGC<br>CAGGAACCGTAAAAAGGCCGCGTTGCTGGCGTTTTTCC<br>ATAGGCTCCGCCCCCCTGACGAGCATCACAAAAATCGA<br>CGCTCAAGTCAGAGGTGGCGAAACCCGACAGGACTATA<br>AAGATACCAGGCGTTTCCCCCTGGAAGCTCCCTCGTGC<br>GCTCTCCTGTTCCGACCCTGCCGCTTACCGGATACCTG<br>TCCGCCTTTCTCCCTTCGGGAAGCGTGGCGCTTTCTCA<br>TAGCTCACGCTGTAGGTATCTCAGTTCGGTGTAGGTCG<br>TTCGCTCCAAGCTGGGCTGTGTGCACGAACCCCCCGTT<br>CAGCCCGACCGCTGCGCCTTATCCGGTAACTATCGTCT<br>TGAGTCCAACCCGGTAAGACACGACTTATCGCCACTGG<br>CAGCAGCCACTGGTAACAGGATTAGCAGAGCGAGGTAT<br>GTAGGCGGTGCTACAGAGTTCTTGAAGTGGTGGCCTAA<br>CTACGGCTACACTAGAAGGACAGTATTTGGTATCTGCG<br>CTCTGCTGAAGCCAGTTACCTTCGGAAAAAGAGTTGGT<br>AGCTCTTGATCCGGCAAACAAACCACCGCTGGTAGCGG<br>TGGTTTTTTTGTTTGCAAGCAGCAGATTACGCGCAGAA<br>AAAAAGGATCTCAAGAAGATCCTTTGATCTTTTCTACG<br>GGGTCTGACGCTCAGTGGAACGAAAACTCACGTTAAGG<br>GATTTTGGTCATGAGATTATCAAAAAGGATCTTCACCT<br>AGATCCTTTTAAATTAAAAATGAAGTTTTAAATCAATC<br>TAAAGTATATATGAGTAAACTTGGTCTGTAATAACTCG<br>GTACCAAATTCCAGAAAAGAGGCCTCCCGAAAGGGGGG<br>CCTTTTTTCGTTTTGGTCCTAATAATTGACGGCTAGCT<br>CAGTCCTAGGTACAGTGCTAGCGTACTTTAACTTTAAG<br>AAGGAGATATACATATGAGCAAAGGAGAAGAACTTTTC<br>ACTGGAGTTGTCCCAATTCTTGTTGAATTAGATGGTGA<br>TGTTAATGGGCACAAATTTTCTGTCCGTGGAGAGGGTG<br>AAGGTGATGCTACAAACGGAAAACTCACCCTTAAATTT<br>ATTTGCACTACTGGAAAACTACCTGTTCCGTGGCCAAC<br>ACTTGTCACTACTCTGACCTATGGTGTTCAATGCTTTT<br>CCCGTTATCCGGATCACATGAAACGGCATGACTTTTTC<br>AAGAGTGCCATGCCCGAAGGTTATGTACAGGAACGCAC<br>TATATCTTTCAAAGATGACGGGACCTACAAGACGCGTG<br>CTGAAGTCAAGTTTGAAGGTGATACCCTTGTTAATCGT | Spec 50 | Custom, adapted from Duet System (Novagen)[15] |

| | |
|---|---|
| | ATCGAGTTAAAGGGTATTGATTTTAAAGAAGATGGAAA<br>CATTCTTGGACACAAACTCGAGTACAACTTTAACTCAC<br>ACAATGTATACATCACGGCAGACAAACAAAAGAATGGA<br>ATCAAAGCTAACTTCAAAATTCGCCACAACGTTGAAGA<br>TGGTTCCGTTCAACTAGCAGACCATTATCAACAAAATA<br>CTCCAATTGGCGATGGCCCTGTCCTTTTACCAGACAAC<br>CATTACCTGTCGACACAATCTGTCCTTTCGAAAGATCC<br>CAACGAAAAGCGTGACCACATGGTCCTTCTTGAGTTTG<br>TAACTGCTGCTGGGATTACACATGGCATGGATGAGCTC<br>TACAAATAATAATCGACCAGGCATCAAATAAAACGAAA<br>GGCTCAGTCGAAAGACTGGGCCTTTCGTTTTATCTGTT<br>GTTTGTCGGTGAACGCTCTCTACTAGAGTCACACTGGC<br>TCACCTTCGGGTGGGCCTTTCTGCGTTTATAGGTCTCA<br>TGTTGGTCGATAGGGACATCTGTCGAGCCGGCAAACGC<br>TTACTAGTAGCGGCCGCTGCAGTCCGGCAAAAAAGGGC<br>AAGGTGTCACCACCCTGCCCTTTTTCTTTAAAACCGAA<br>AAGATTACTTCGCGTTATGCAGGCTTCCTCGCTCACTG<br>ACTCGCTGCGCTCGGTCGTTCGGCTGCGGCGAGCGGTA<br>TCAGCTCACTCAAAGGCGGTAATGACATCAGGCATTTG<br>AGAAGCACACGGTCACACTGCTTCCGGTAGTCAATAAA<br>CCGGTAAACCAGCAATAGACATAAGCGGCTATTTAACG<br>ACCCTGCCCTGAACCGACGACCGGGTCATCGTGGCCGG<br>ATCTTGCGGCCCCTCGGCTTGAACGAATTGTTAGACAT<br>TATTTGCCGACTACCTTGGTGATCTCGCCTTTCACGTA<br>GTGGACAAATTCTTCCAACTGATCTGCGCGCGAGGCCA<br>AGCGATCTTCTTCTTGTCCAAGATAAGCCTGTCTAGCT<br>TCAAGTATGACGGGCTGATACTGGGCCGGCAGGCGCTC<br>CATTGCCCAGTCGGCAGCGACATCCTTCGGCGCGATTT<br>TGCCGGTTACTGCGCTGTACCAAATGCGGGACAACGTA<br>AGCACTACATTTCGCTCATCGCCAGCCCAGTCGGGCGG<br>CGAGTTCCATAGCGTTAAGGTTTCATTTAGCGCCTCAA<br>ATAGATCCTGTTCAGGAACCGGATCAAAGAGTTCCTCC<br>GCCGCTGGACCTACCAAGGCAACGCTATGTTCTCTTGC<br>TTTTGTCAGCAAGATAGCCAGATCAATGTCGATCGTGG<br>CTGGCTCGAAGATACCTGCAAGAATGTCATTGCGCTGC<br>CATTCTCCAAATTGCAGTTCGCGCTTAGCTGGATAACG<br>CCACGGAATGATGTCGTCGTGCACAACAATGGTGACTT<br>CTACAGCGCGGAGAATCTCGCTCTCTCCAGGGGAAGCC<br>GAAGTTTCCAAAAGGTCGTTGATCAAAGCTCGCCGCGT<br>TGTTTCATCAAGCCTTACGGTCACCGTAACCAGCAAAT |

| | |
|---|---|
| | CAATATCACTGTGTGGCTTCAGGCCGCCATCCACTGCG<br>GAGCCGTACAAATGTACGGCCAGCAACGTCGGTTCGAG<br>ATGGCGCTCGATGACGCCAACTACCTCTGATAGTTGAG<br>TCGATACTTCGGCGATCACCGCTTCCCTCATACTCTTC<br>CTTTTTCAATATTATTGAAGCATTTATCAGGGTTATTG<br>TCTCATGAGCGGATACATATTTGAATGTATTTAGAAAA<br>ATAAACAAATAGCTAGCTCACTCGGTCGCTACGCTCCG<br>GGCGTGAGACTGCGGCGGGCGCTGCGGACACATACAAA<br>GTTACCCACAGATTCCGTGGATAAGCAGGGGACTAACA<br>TGTGAGGCAAAACAGCAGGGCCGCGCCGGTGGCGTTTT<br>TCCATAGGCTCCGCCCTCCTGCCAGAGTTCACATAAAC<br>AGACGCTTTTCCGGTGCATCTGTGGGAGCCGTGAGGCT<br>CAACCATGAATCTGACAGTACGGGCGAAACCCGACAGG<br>ACTTAAAGATCCCCACCGTTTCCGGCGGGTCGCTCCCT<br>CTTGCGCTCTCCTGTTCCGACCCTGCCGTTTACCGGAT<br>ACCTGTTCCGCCTTTCTCCCTTACGGGAAGTGTGGCGC<br>TTTCTCATAGCTCACACACTGGTATCTCGGCTCGGTGT<br>AGGTCGTTCGCTCCAAGCTGGGCTGTAAGCAAGAACTC<br>CCCGTTCAGCCCGACTGCTGCGCCTTATCCGGTAACTG<br>TTCACTTGAGTCCAACCCGGAAAAGCACGGTAAAACGC<br>CACTGGCAGCAGCCATTGGTAACTGGGAGTTCGCAGAG<br>GATTTGTTTAGCTAAACACGCGGTTGCTCTTGAAGTGT<br>GCGCCAAAGTCCGGCTACACTGGAAGGACAGATTTGGT<br>TGCTGTGCTCTGCGAAAGCCAGTTACCACGGTTAAGCA<br>GTTCCCCAACTGACTTAACCTTCGATCAAACCACCTCC<br>CCAGGTGGTTTTTTCGTTTACAGGGCAAAAGATTACGC<br>GCAGAAAAAAAGGATCTCAAGAAGATCCTTTGATCTTT<br>TCTACTGAACCGCTCTAGATTTCAGTGCAATTTATCTC<br>TTCAAATGTAGCACCTGAAGTCAGCCCCATACGATATA<br>AGTTGTAATTCTCATGTTAGTCATGCCCCGCGCCCACC<br>GGAAGGAGCTGACTGGGTTGAAGGCTCTCAAGGGCATC<br>GGTCGAGATCCCGGTGCCTAATGAGTGAGCTAACAAAA<br>GTGCCACCTGACGTCTAAGAAACCATTATTATCATGAC<br>ATTAACCTATAAAACCGGCTTATCGGTCAGTTTCACCT<br>GATTTACGTAAAAACCCGCTTCGGCGGGTTTTTGCTTT<br>TGGAGGGGCAGAAAGATGAATGACTGTCCACGACGCTA<br>TACCCAAAAGAAAAAAAAAAAACCCCGCCCCTGACAGG<br>GCGGGGTTTTTTTTCGTCCTATAAGTCTTCCATATGCA<br>CGCATGCGAAGACAAGTACGAATTCGCGGCCGCTTCTA<br>GAGAACAGATAAAATATTTGCTCATGAGCCCGAAGTGG |

| | | | |
|---|---|---|---|
| | CGAGCCCGATCTTCCCCATCGGTGATGTCGGCGATATA<br>GGCGCCAGCAACCGCACCTGTGGCGCCGGTGATGCCGG<br>CCACGATGCGTCCGGCGTAGAGGATCTGCTCATGTTTG<br>ACAGCTTATCATCGATGCATAATGTGCCTGTCAAATGG<br>ACGAAGCAGGGATTCTGCAAACCCTATGCTACTCCCTC<br>GAGCCGTCAATTGTCTGATTCGTTACCAATTAGGATCC<br>TTATCACTGCCCGCTTTCCAGTCGGGAAACCTGTCGTG<br>CCAGCTGCATTAATGAATCGGCCAACGCGCGGGGAGAG<br>GCGGTTTGCGTATTGGGCGCCAGGGTGGTTTTTCTTTT<br>CACCAGTGAGACTGGCAACAGCTGATTGCCCTTCACCG<br>CCTGGCCCTGAGAGAGTTGCAGCAAGCGGTCCACGCTG<br>GTTTGCCCCAGCAGGCGAAAATCCTGTTTGATGGTGGT<br>TAACGGCGGGATATAACATGAGCTATCTTCGGTATCGT<br>CGTATCCCACTACCGAGATATCCGCACCAACGCGCAGC<br>CCGGACTCGGTAATGGCGCGCATTGCGCCCAGCGCCAT<br>CTGATCGTTGGCAACCAGCATCGCAGTGGGAACGATGC<br>CCTCATTCAGCATTTGCATGGTTTGTTGAAAACCGGAC<br>ATGGCACTCCAGTCGCCTTCCCGTTCCGCTATCGGCTG<br>AATTTGATTGCGAGTGAGATATTTATGCCAGCCAGCCA<br>GACGCAGACGCGCCGAGACAGAACTTAATGGGCCCGCT<br>AACAGCGCGATTTGCTGGTGACCCAATGCGACCAGATG<br>CTCCACGCCCAGTCGCGTACCGTCCTCATGGGAGTAAA<br>TAATACTGTTGATGGGTGTCTGGTCAGAGACATCAAGA<br>AATAACGCCGGAACATTAGTGCAGGCAGCTTCCACAGC<br>AATGGCATCCTGGTCATCCAGCGGATAGTTAATGATCA<br>GCCCACTGACGCGTTGCGCGAGAAGATTGTGCACCGCC<br>GCTTTACAGGCTTCGACGCCGCTTCGTTCTACCATCGA<br>CACCACCACGCTGGCACCCAGTTGATCGGCGCGAGATT<br>TAATCGCCGCGACAATTTGCGACGGCGCGTGCAGGGCC<br>AGACTGGAGGTGGCAACGCCAATCAGCAACGACTGTTT<br>GCCCGCCAGTTGTTGTGCCACGCGGTTGGGAATGTAAT<br>TCAGCTCCACCATCGCCGCTTCCACTTTTTCCCGCGTT<br>TTCGCAGAAACGTGGCTGGCCTGGTTCACCACGCGGGA<br>AACGGTCATATAAGAGACACCGGCATACTCTGCGACAT<br>CGTATAACGTTACTGGTTTCATATTCACCACCCTGAAT<br>TGACTCTCTTCCGGGCGCTATCATGCCATACCGCGAAA<br>GGTTTTGCGCCATTCGATGGCGCGCC | | |
| pACYC-Kan-LacI | GCCAATAGGCGTATCACGAGGCAGAATTTCAGATAAAA<br>AAAATCCTTAGCTTTCGCTAAGGATGATTTCTGGGCGT<br>TGACAATGCGAAGTCAGTTCTATAGAGACCTAATAAGC | Kan 50 | Custom, adapted from Duet |

| | | | |
|---|---|---|---|
| | CAGGAACCGTAAAAAGGCCGCGTTGCTGGCGTTTTTCC<br>ATAGGCTCCGCCCCCCTGACGAGCATCACAAAAATCGA<br>CGCTCAAGTCAGAGGTGGCGAAACCCGACAGGACTATA<br>AAGATACCAGGCGTTTCCCCCTGGAAGCTCCCTCGTGC<br>GCTCTCCTGTTCCGACCCTGCCGCTTACCGGATACCTG<br>TCCGCCTTTCTCCCTTCGGGAAGCGTGGCGCTTTCTCA<br>TAGCTCACGCTGTAGGTATCTCAGTTCGGTGTAGGTCG<br>TTCGCTCCAAGCTGGGCTGTGTGCACGAACCCCCCGTT<br>CAGCCCGACCGCTGCGCCTTATCCGGTAACTATCGTCT<br>TGAGTCCAACCCGGTAAGACACGACTTATCGCCACTGG<br>CAGCAGCCACTGGTAACAGGATTAGCAGAGCGAGGTAT<br>GTAGGCGGTGCTACAGAGTTCTTGAAGTGGTGGCCTAA<br>CTACGGCTACACTAGAAGGACAGTATTTGGTATCTGCG<br>CTCTGCTGAAGCCAGTTACCTTCGGAAAAAGAGTTGGT<br>AGCTCTTGATCCGGCAAACAAACCACCGCTGGTAGCGG<br>TGGTTTTTTTGTTTGCAAGCAGCAGATTACGCGCAGAA<br>AAAAAGGATCTCAAGAAGATCCTTTGATCTTTTCTACG<br>GGGTCTGACGCTCAGTGGAACGAAAACTCACGTTAAGG<br>GATTTTGGTCATGAGATTATCAAAAAGGATCTTCACCT<br>AGATCCTTTTAAATTAAAAATGAAGTTTTAAATCAATC<br>TAAAGTATATATGAGTAAACTTGGTCTGTAATAACTCG<br>GTACCAAATTCCAGAAAAGAGGCCTCCCGAAAGGGGGG<br>CCTTTTTTCGTTTTGGTCCTAATAATTGACGGCTAGCT<br>CAGTCCTAGGTACAGTGCTAGCGTACTTTAACTTTAAG<br>AAGGAGATATACATATGAGCAAAGGAGAAGAACTTTTC<br>ACTGGAGTTGTCCCAATTCTTGTTGAATTAGATGGTGA<br>TGTTAATGGGCACAAATTTTCTGTCCGTGGAGAGGGTG<br>AAGGTGATGCTACAAACGGAAAACTCACCCTTAAATTT<br>ATTTGCACTACTGGAAAACTACCTGTTCCGTGGCCAAC<br>ACTTGTCACTACTCTGACCTATGGTGTTCAATGCTTTT<br>CCCGTTATCCGGATCACATGAAACGGCATGACTTTTTC<br>AAGAGTGCCATGCCCGAAGGTTATGTACAGGAACGCAC<br>TATATCTTTCAAAGATGACGGGACCTACAAGACGCGTG<br>CTGAAGTCAAGTTTGAAGGTGATACCCTTGTTAATCGT<br>ATCGAGTTAAAGGGTATTGATTTTAAAGAAGATGGAAA<br>CATTCTTGGACACAAACTCGAGTACAACTTTAACTCAC<br>ACAATGTATACATCACGGCAGACAAACAAAAGAATGGA<br>ATCAAAGCTAACTTCAAAATTCGCCACAACGTTGAAGA<br>TGGTTCCGTTCAACTAGCAGACCATTATCAACAAAATA<br>CTCCAATTGGCGATGGCCCTGTCCTTTTACCAGACAAC | | System (Novagen)[15] |

| | |
|---|---|
| | CATTACCTGTCGACACAATCTGTCCTTTCGAAAGATCC<br>CAACGAAAAGCGTGACCACATGGTCCTTCTTGAGTTTG<br>TAACTGCTGCTGGGATTACACATGGCATGGATGAGCTC<br>TACAAATAATAATCGACCAGGCATCAAATAAAACGAAA<br>GGCTCAGTCGAAAGACTGGGCCTTTCGTTTTATCTGTT<br>GTTTGTCGGTGAACGCTCTCTACTAGAGTCACACTGGC<br>TCACCTTCGGGTGGGCCTTTCTGCGTTTATAGGTCTCA<br>TGTTGGTCGATAGGGACATCTGTCGAGCCGGCAAACGC<br>TTACTAGTAGCGGCCGCTGCAGTCCGGCAAAAAAGGGC<br>AAGGTGTCACCACCCTGCCCTTTTTCTTTAAAACCGAA<br>AAGATTACTTCGCGTTATGCAGGCTTCCTCGCTCACTG<br>ACTCGCTGCGCTCGGTCGTTCGGCTGCGGCGAGCGGTA<br>TCAGCTCACTCAAAGGCGGTAATGACATCCTCGCTCAC<br>TGACTCGCTGCACGAGGCAGACCTCAGCGCTAGCGGAG<br>TGTATACTGGCTTACTATGTTGGCACTGATGAGGGTGT<br>CAGTGAAGTGCTTCATGTGGCAGGAGAAAAAAGGCTGC<br>ACCGGTGCGTCAGCAGAATATGTGATACAGGATATATT<br>CCGCTTCCTCGCTCACTGACTCGCTACGCTCGGTCGTT<br>CGACTGCGGCGAGCGGAAATGGCTTACGAACGGGGCGG<br>AGATTTCCTGGAAGATGCCAGGAAGATACTTAACAGGG<br>AAGTGAGAGGGCCGCGGCAAAGCCGTTTTTCCATAGGC<br>TCCGCCCCCCTGACAAGCATCACGAAATCTGACGCTCA<br>AATCAGTGGTGGCGAAACCCGACAGGACTATAAAGATA<br>CCAGGCGTTTCCCCCTGGCGGCTCCCTCGTGCGCTCTC<br>CTGTTCCTGCCTTTCGGTTTACCGGTGTCATTCCGCTG<br>TTATGGCCGCGTTTGTCTCATTCCACGCCTGACACTCA<br>GTTCCGGGTAGGCAGTTCGCTCCAAGCTGGACTGTATG<br>CACGAACCCCCCGTTCAGTCCGACCGCTGCGCCTTATC<br>CGGTAACTATCGTCTTGAGTCCAACCCGGAAAGACATG<br>CAAAAGCACCACTGGCAGCAGCCACTGGTAATTGATTT<br>AGAGGAGTTAGTCTTGAAGTCATGCGCCGGTTAAGGCT<br>AAACTGAAAGGACAAGTTTTGGTGACTGCGCTCCTCCA<br>AGCCAGTTACCTCGGTTCAAAGAGTTGGTAGCTCAGAG<br>AACCTTCGAAAAACCGCCCCGCAAGGCGGTTTTTTCGT<br>TTTCAGAGCAAGAGATTACGCGCAGACCAAAACGATCT<br>CAAGAAGATCATCTTATTAAGGGGTCTGACGCTCAGTG<br>GAACGAAAAATCAATCTAAAGTATATATGAGTAAACTT<br>GGTCTGACAGTTACCTTAGAAAAACTCATCGAGCATCA<br>AATGAAACTGCAATTTATTCATATCAGGATTATCAATA<br>CCATATTTTTGAAAAAGCCGTTTCTGTAATGAAGGAGA |

| | |
|---|---|
| | AAACTCACCGAGGCAGTTCCATAGGATGGCAAGATCCT<br>GGTATCGGTCTGCGATTCCGACTCGTCCAACATCAATA<br>CAACCTATTAATTTCCCCTCGTCAAAAATAAGGTTATC<br>AAGTGAGAAATCACCATGAGTGACGACTGAATCCGGTG<br>AGAATGGCAAAAGCTTATGCATTTCTTTCCAGACTTGT<br>TCAACAGGCCAGCCATTACGCTCGTCATCAAAATCACT<br>CGCATCAACCAAACCGTTATTCATTCGTGATTGCGCCT<br>GAGCTAGACGAAATACGCGATCGCTGTTAAAAGGACAA<br>TTACAAACAGGAATCGAATGCAACCGGCGCAGGAACAC<br>TGCCAGCGCATCAACAATATTTTCACCTGAATCAGGAT<br>ATTCTTCTAATACCTGGAATGCTGTTTTCCCGGGGATC<br>GCAGTGGTGAGTAACCATGCATCATCAGGAGTACGGAT<br>AAAATGCTTGATGGTCGGAAGAGGCATAAATTCCGTCA<br>GCCAGTTTAGTCTGACCATCTCATCTGTAACATCATTG<br>GCAACGCTACCTTTGCCATGTTTCAGAAACAACTCTGG<br>CGCATCGGGCTTCCCATACAATCGATAGATTGTCGCAC<br>CTGATTGCCCGACATTATCGCGAGCCCATTTATACCCA<br>TATAAATCAGCATCCATGTTGGAATTTAATCGCGGCCT<br>GGAGCAAGACGTTTCCCGTTGAATATGGCTCATAACAC<br>CCCTTGTATTACTGTTTATGTAAGCAGACCAAAAGTGC<br>CACCTGACGTCTAAGAAACCATTATTATCATGACATTA<br>ACCTATAAAAACCGGCTTATCGGTCAGTTTCACCTGATT<br>TACGTAAAAACCCGCTTCGGCGGGTTTTTGCTTTTGGA<br>GGGGCAGAAAGATGAATGACTGTCCACGACGCTATACC<br>CAAAAGAAAAAAAAAAAACCCCGCCCCTGACAGGGCGG<br>GGTTTTTTTTCGTCCTATAAGTCTTCCATATGCACGCA<br>TGCGAAGACAAGTACGAATTCGCGGCCGCTTCTAGAGA<br>ACAGATAAAATATTTGCTCATGAGCCCGAAGTGGCGAG<br>CCCGATCTTCCCCATCGGTGATGTCGGCGATATAGGCG<br>CCAGCAACCGCACCTGTGGCGCCGGTGATGCCGGCCAC<br>GATGCGTCCGGCGTAGAGGATCTGCTCATGTTTGACAG<br>CTTATCATCGATGCATAATGTGCCTGTCAAATGGACGA<br>AGCAGGGATTCTGCAAACCCTATGCTACTCCCTCGAGC<br>CGTCAATTGTCTGATTCGTTACCAATTAGGATCCTTAT<br>CACTGCCCGCTTTCCAGTCGGGAAACCTGTCGTGCCAG<br>CTGCATTAATGAATCGGCCAACGCGCGGGGAGAGGCGG<br>TTTGCGTATTGGGCGCCAGGGTGGTTTTTCTTTTCACC<br>AGTGAGACTGGCAACAGCTGATTGCCCTTCACCGCCTG<br>GCCCTGAGAGAGTTGCAGCAAGCGGTCCACGCTGGTTT<br>GCCCCAGCAGGCGAAAATCCTGTTTGATGGTGGTTAAC |

| | |
|---|---|
| | GGCGGGATATAACATGAGCTATCTTCGGTATCGTCGTA<br>TCCCACTACCGAGATATCCGCACCAACGCGCAGCCCGG<br>ACTCGGTAATGGCGCGCATTGCGCCCAGCGCCATCTGA<br>TCGTTGGCAACCAGCATCGCAGTGGGAACGATGCCCTC<br>ATTCAGCATTTGCATGGTTTGTTGAAAACCGGACATGG<br>CACTCCAGTCGCCTTCCCGTTCCGCTATCGGCTGAATT<br>TGATTGCGAGTGAGATATTTATGCCAGCCAGCCAGACG<br>CAGACGCGCCGAGACAGAACTTAATGGGCCCGCTAACA<br>GCGCGATTTGCTGGTGACCCAATGCGACCAGATGCTCC<br>ACGCCCAGTCGCGTACCGTCCTCATGGGAGTAAATAAT<br>ACTGTTGATGGGTGTCTGGTCAGAGACATCAAGAAATA<br>ACGCCGGAACATTAGTGCAGGCAGCTTCCACAGCAATG<br>GCATCCTGGTCATCCAGCGGATAGTTAATGATCAGCCC<br>ACTGACGCGTTGCGCGAGAAGATTGTGCACCGCCGCTT<br>TACAGGCTTCGACGCCGCTTCGTTCTACCATCGACACC<br>ACCACGCTGGCACCCAGTTGATCGGCGCGAGATTTAAT<br>CGCCGCGACAATTTGCGACGGCGCGTGCAGGGCCAGAC<br>TGGAGGTGGCAACGCCAATCAGCAACGACTGTTTGCCC<br>GCCAGTTGTTGTGCCACGCGGTTGGGAATGTAATTCAG<br>CTCCACCATCGCCGCTTCCACTTTTTCCCGCGTTTTCG<br>CAGAAACGTGGCTGGCCTGGTTCACCACGCGGGAAACG<br>GTCATATAAGAGACACCGGCATACTCTGCGACATCGTA<br>TAACGTTACTGGTTTCATATTCACCACCCTGAATTGAC<br>TCTCTTCCGGGCGCTATCATGCCATACCGCGAAAGGTT<br>TTGCGCCATTCGATGGCGCGCC |

**Table S7: Plasmids used in this work and the parts used to assemble them**. The assemblies were all performed with BsaI-HFv2 and T4 DNA Ligase as described in the methods section.

| Plasmid name | Full plasmid name | Parts used |
|---|---|---|
| TJ-C-559 | pCDF-T7-lacO-RiboJ53-20k-PlpX-B34-PlpY | pCDF-Spec-LacI<br>T7-lacO-RiboJ53<br>20k-PlpX-B34-PlpY<br>Bba_B0015 |
| TJ-C-560 | pCDF-T7-lacO-RiboJ53-20k-HypX-B34-HypY | pCDF-Spec-LacI<br>T7-lacO-RiboJ53<br>20k-HypX-B34-HypY<br>Bba_B0015 |
| TJ-C-689 | pCDF-T7-lacO-RiboJ53-20k-HypX-45k-HypY | pCDF-Spec-LacI<br>T7-lacO-RiboJ53<br>20k-HypX-45k-HypY<br>Bba_B0015 |
| TJ-C-593 | pACYC-T7-lacO-ElvJ-100k-Fh8-chaos-HypA1-HypA1 | pACYC-Kan-LacI<br>T7-lacO-ElvJ-100k-Fh8-chaos<br>HypA1<br>BBa_B0015 |
| TJ-C-594 | pACYC-T7-lacO-ElvJ-100k-Fh8-chaos-HypA2-HypA2 | pACYC-Kan-LacI<br>T7-lacO-ElvJ-100k-Fh8-chaos<br>HypA2<br>BBa_B0015 |
| TJ-C-595 | pACYC-T7-lacO-ElvJ-100k-Fh8-chaos-HypA3-HypA3 | pACYC-Kan-LacI<br>T7-lacO-ElvJ-100k-Fh8-chaos<br>HypA3<br>BBa_B0015 |
| TJ-C-596 | pACYC-T7-lacO-ElvJ-100k-Fh8-chaos-HypA4-HypA4 | pACYC-Kan-LacI<br>T7-lacO-ElvJ-100k-Fh8-chaos<br>HypA4<br>BBa_B0015 |
| TJ-C-597 | pACYC-T7-lacO-ElvJ-100k-Fh8-chaos-HypA5-HypA5 | pACYC-Kan-LacI<br>T7-lacO-ElvJ-100k-Fh8-chaos<br>HypA5<br>BBa_B0015 |

| | | |
|---|---|---|
| TJ-C-598 | pACYC-T7-lacO-ElvJ-100k-Fh8-chaos-HypA6-HypA6 | pACYC-Kan-LacI<br>T7-lacO-ElvJ-100k-Fh8-chaos<br>HypA6<br>BBa_B0015 |
| TJ-C-599 | pACYC-T7-lacO-ElvJ-100k-Fh8-chaos-HypA7-HypA7 | pACYC-Kan-LacI<br>T7-lacO-ElvJ-100k-Fh8-chaos<br>HypA7<br>BBa_B0015 |
| TJ-C-614 | pACYC-T7-lacO-ElvJ-100k-Fh8-HypAc-HypA1 | pACYC-Kan-LacI<br>T7-lacO-ElvJ-100k-Fh8<br>HypAc leader<br>HypA1 core<br>BBa_B0015 |
| TJ-C-615 | pACYC-T7-lacO-ElvJ-100k-Fh8-HypAc-HypA2 | pACYC-Kan-LacI<br>T7-lacO-ElvJ-100k-Fh8<br>HypAc leader<br>HypA2 core<br>BBa_B0015 |
| TJ-C-616 | pACYC-T7-lacO-ElvJ-100k-Fh8-HypAc-HypA3 | pACYC-Kan-LacI<br>T7-lacO-ElvJ-100k-Fh8<br>HypAc leader<br>HypA3 core<br>BBa_B0015 |
| TJ-C-617 | pACYC-T7-lacO-ElvJ-100k-Fh8-HypAc-HypA4 | pACYC-Kan-LacI<br>T7-lacO-ElvJ-100k-Fh8<br>HypAc leader<br>HypA4 core<br>BBa_B0015 |
| TJ-C-618 | pACYC-T7-lacO-ElvJ-100k-Fh8-HypAc-HypA5 | pACYC-Kan-LacI<br>T7-lacO-ElvJ-100k-Fh8<br>HypAc leader<br>HypA5 core<br>BBa_B0015 |
| TJ-C-619 | pACYC-T7-lacO-ElvJ-100k-Fh8-HypAc-HypA6 | pACYC-Kan-LacI<br>T7-lacO-ElvJ-100k-Fh8<br>HypAc leader<br>HypA6 core<br>BBa_B0015 |

| | | |
|---|---|---|
| TJ-C-620 | pACYC-T7-lacO-ElvJ-100k-Fh8-HypAc-HypA7 | pACYC-Kan-LacI<br>T7-lacO-ElvJ-100k-Fh8<br>HypAc leader<br>HypA7 core<br>BBa_B0015 |
| TJ-C-693 | pACYC-K-T7-lacO-Fh8-HypAc-TEV-Artificial-1 | pACYC-Kan-LacI<br>T7-lacO-ElvJ-100k-Fh8<br>HypAc-TEV leader<br>Artificial-1 core<br>BBa_B0015 |
| TJ-C-694 | pACYC-K-T7-lacO-Fh8-HypAc-TEV-Artificial-2 | pACYC-Kan-LacI<br>T7-lacO-ElvJ-100k-Fh8<br>HypAc-TEV leader<br>Artificial-2 core<br>BBa_B0015 |
| TJ-C-695 | pACYC-K-T7-lacO-Fh8-HypAc-TEV-Artificial-3 | pACYC-Kan-LacI<br>T7-lacO-ElvJ-100k-Fh8<br>HypAc-TEV leader<br>Artificial-3 core<br>BBa_B0015 |
| TJ-C-696 | pACYC-K-T7-lacO-Fh8-HypAc-TEV-Artificial-4 | pACYC-Kan-LacI<br>T7-lacO-ElvJ-100k-Fh8<br>HypAc-TEV leader<br>Artificial-4 core<br>BBa_B0015 |
| TJ-C-697 | pACYC-K-T7-lacO-Fh8-PlpA3-TEV-Artificial-1 | pACYC-Kan-LacI<br>T7-lacO-ElvJ-100k-Fh8<br>PlpA3-TEV leader<br>Artificial-1 core<br>BBa_B0015 |
| TJ-C-698 | pACYC-K-T7-lacO-Fh8-PlpA3-TEV-Artificial-2 | pACYC-Kan-LacI<br>T7-lacO-ElvJ-100k-Fh8<br>PlpA3-TEV leader<br>Artificial-2 core<br>BBa_B0015 |
| TJ-C-699 | pACYC-K-T7-lacO-Fh8-PlpA3-TEV-Artificial-3 | pACYC-Kan-LacI<br>T7-lacO-ElvJ-100k-Fh8<br>PlpA3-TEV leader<br>Artificial-3 core |

| | | |
|---|---|---|
| | | BBa_B0015 |
| TJ-C-700 | pACYC-K-T7-lacO-Fh8-PlpA3-TEV-Artificial-4 | pACYC-Kan-LacI<br>T7-lacO-ElvJ-100k-Fh8<br>PlpA3-TEV leader<br>Artificial-4 core<br>BBa_B0015 |
| TJ-C-706 | pACYC-K-T7-lacO-PP1-RSTN-HypAc-TEV-Artificial-2 | pACYC-Kan-LacI<br>T7-lacO-PP1-RSTN1<br>HypAc-TEV leader<br>Artificial-2 core<br>BBa_B0015 |
| TJ-C-710 | pACYC-K-T7-lacO-PP1-RSTN-PlpA3-TEV-Artificial-2 | pACYC-Kan-LacI<br>T7-lacO-PP1-RSTN1<br>PlpA3-TEV leader<br>Artificial-2 core<br>BBa_B0015 |
| TJ-C-708 | pACYC-K-T7-lacO-ElvJ-Pfu-MBP-HypAc-TEV-Artificial-2 | pACYC-Kan-LacI<br>T7-lacO-ElvJ-30k-Pfu-MBP<br>HypAc-TEV leader<br>Artificial-2 core<br>BBa_B0015 |
| TJ-C-712 | pACYC-K-T7-lacO-ElvJ-Pfu-MBP-PlpA3-TEV-Artificial-2 | pACYC-Kan-LacI<br>T7-lacO-ElvJ-30k-Pfu-MBP<br>PlpA3-TEV leader<br>Artificial-2 core<br>BBa_B0015 |

**Table S8: Primers used to amplify TJ-C-706 or TJ-C-710 to produce the point mutants.** The point mutants were then used in the 96-well plate screen via blunt end cloning post-PCR.

| Primer | Sequence | Description |
|---|---|---|
| TJ-O-1091 | TGGTCCGCATCCGTG | mutates the AA Q in the Artificial 1 core |
| TJ-O-1142 | GATCCTGTTTACGGCGTTATTTTG | use with 1091 for making D, mutant-1 |
| TJ-O-1143 | GGTCCTGTTTACGGCGTTATTTTG | use with 1091 for making G, mutant-2 |
| TJ-O-1144 | CCTCCTGTTTACGGCGTTATTTTG | use with 1091 for making P, mutant-3 |
| TJ-O-1145 | TCTCCTGTTTACGGCGTTATTTTG | use with 1091 for making S, mutant-4 |
| TJ-O-1146 | CATCCTGTTTACGGCGTTATTTTG | use with 1091 for making H, mutant-5 |
| TJ-O-1147 | AAACCTGTTTACGGCGTTATTTTG | use with 1091 for making K, mutant-6 |
| TJ-O-1148 | TTTCCTGTTTACGGCGTTATTTTG | use with 1091 for making F, mutant-7 |
| TJ-O-1149 | ATGCCTGTTTACGGCGTTATTTTG | use with 1091 for making M, mutant-8 |
| TJ-O-1150 | ATTCCTGTTTACGGCGTTATTTTG | use with 1091 for making I, mutant-9 |
| TJ-O-1151 | GTTCCTGTTTACGGCGTTATTTTG | use with 1091 for making V, mutant-10 |
| TJ-O-1102 | CTGTGGTCCGCATCCG | mutates the AA P in the Artificial 2 core |
| TJ-O-1103 | GATGTTTACGGCGTTATTTTGCC | use with 1102 for making D, mutant-11 |
| TJ-O-1104 | GGTGTTTACGGCGTTATTTTGCC | use with 1102 for making G, mutant-12 |
| TJ-O-1105 | ACTGTTTACGGCGTTATTTTGCC | use with 1102 for making T, mutant-13 |
| TJ-O-1106 | CAAGTTTACGGCGTTATTTTGCC | use with 1102 for making Q, mutant-14 |
| TJ-O-1107 | TCTGTTTACGGCGTTATTTTGCC | use with 1102 for making S, mutant-15 |

| | | |
|---|---|---|
| TJ-O-1108 | CATGTTTACGGCGTTATTTTGCC | use with 1102 for making H, mutant-16 |
| TJ-O-1109 | AAAGTTTACGGCGTTATTTTGCC | use with 1102 for making K, mutant-17 |
| TJ-O-1110 | TTTGTTTACGGCGTTATTTTGCC | use with 1102 for making F, mutant-18 |
| TJ-O-1111 | ATGGTTTACGGCGTTATTTTGCC | use with 1102 for making M, mutant-19 |
| TJ-O-1112 | ATTGTTTACGGCGTTATTTTGCC | use with 1102 for making I, mutant-20 |
| TJ-O-1113 | AGGCTGTGGTCCGCATC | mutates the AA V in the Artificial 3 core |
| TJ-O-1114 | GATTACGGCGTTATTTTGCCATG | use with 1113 for making D, mutant-21 |
| TJ-O-1115 | GGTTACGGCGTTATTTTGCCATG | use with 1113 for making G, mutant-22 |
| TJ-O-1116 | CCTTACGGCGTTATTTTGCCATG | use with 1113 for making P, mutant-23 |
| TJ-O-1117 | CAATACGGCGTTATTTTGCCATG | use with 1113 for making Q, mutant-24 |
| TJ-O-1118 | TCTTACGGCGTTATTTTGCCATG | use with 1113 for making S, mutant-25 |
| TJ-O-1119 | CATTACGGCGTTATTTTGCCATG | use with 1113 for making H, mutant-26 |
| TJ-O-1120 | AAATACGGCGTTATTTTGCCATG | use with 1113 for making K, mutant-27 |
| TJ-O-1121 | TTTTACGGCGTTATTTTGCCATG | use with 1113 for making F, mutant-28 |
| TJ-O-1122 | ATGTACGGCGTTATTTTGCCATG | use with 1113 for making M, mutant-29 |
| TJ-O-1123 | ATTTACGGCGTTATTTTGCCATG | use with 1113 for making I, mutant-30 |
| TJ-O-1124 | GCCGTAAACAGGCTGTGGT | mutates the AA after the glycine splice site |
| TJ-O-1125 | GATATTTTGCCATGGGCTTAATAATC | use with 1124 for making D, mutant-31 |
| TJ-O-1126 | GGTATTTTGCCATGGGCTTAATAATC | use with 1124 for making G, mutant-32 |

| | | |
|---|---|---|
| TJ-O-1127 | CCTATTTTGCCATGGGCTTAATAATC | use with 1124 for making P, mutant-33 |
| TJ-O-1128 | CAAATTTTGCCATGGGCTTAATAATC | use with 1124 for making Q, mutant-34 |
| TJ-O-1129 | TCTATTTTGCCATGGGCTTAATAATC | use with 1124 for making S, mutant-35 |
| TJ-O-1130 | CATATTTTGCCATGGGCTTAATAATC | use with 1124 for making H, mutant-36 |
| TJ-O-1131 | AAAATTTTGCCATGGGCTTAATAATC | use with 1124 for making K, mutant-37 |
| TJ-O-1132 | TTTATTTTGCCATGGGCTTAATAATC | use with 1124 for making F, mutant-38 |
| TJ-O-1133 | ATGATTTTGCCATGGGCTTAATAATC | use with 1124 for making M, mutant-39 |
| TJ-O-1134 | ATTATTTTGCCATGGGCTTAATAATC | use with 1124 for making I, mutant-40 |

**Table S9: Primers used to amplify TJ-C-710 to produce the Rosetta-designed spliceotides**. Precursor peptide sequences were generated for the 96-well plate screen via blunt end cloning post-PCR.

| Primer | Sequence | Description |
|---|---|---|
| TJ-O-1268 | TCCGCATCCGCCTTG | use on TJ-C-710 to make the mutants, all other primers used alongside this one |
| TJ-O-1269 | GAACGTCCTGCTTATGGTACTGTTTTGCCATGGGCTTAATAATCG | GGCGERPAYGTVLPWA, design-1 |
| TJ-O-1270 | GAACGTCCTGCTTATGGTGTTGTTTTGCCATGGGCTTAATAATCG | GGCGERPAYGVVLPWA, design-2 |
| TJ-O-1271 | GAATATCCTTCTTATGGTGTTATTTTGCCATGGGCTTAATAATCG | GGCGEYPSYGVILPWA, design-3 |
| TJ-O-1272 | GAACGTCCTGCTTATGGTGTTACTTTGCCATGGGCTTAATAATCG | GGCGERPAYGVTLPWA, design-4 |
| TJ-O-1273 | GAATTTCCTGCTTATGGTGTTGTTTTGCCATGGGCTTAATAATCG | GGCGEFPAYGVVLPWA, design-5 |
| TJ-O-1274 | CCTTATCCTGCTTATGGTGTTGTTTTGCCATGGGCTTAATAATCG | GGCGPYPAYGVVLPWA, design-6 |
| TJ-O-1275 | ACTACTCCTGCTTATGGTGTTGTTTTGCCATGGGCTTAATAATCG | GGCGTTPAYGVVLPWA, design-7 |
| TJ-O-1276 | ACTTATCCTGCTTATGGTGTTATTTTGCCATGGGCTTAATAATCG | GGCGTYPAYGVILPWA, design-8 |
| TJ-O-1277 | GTTTATCCTGCTTATGGTGTTGTTTTGCCATGGGCTTAATAATCG | GGCGVYPAYGVVLPWA, design-9 |
| TJ-O-1278 | ACTTATCCTGCTTATGGTACTGTTTTGCCATGGGCTTAATAATCG | GGCGTYPAYGTVLPWA, design-10 |
| TJ-O-1279 | GCTTTTCCTTCTTATGGTGTTGTTTTGCCATGGGCTTAATAATCG | GGCGAFPSYGVVLPWA, design-11 |
| TJ-O-1280 | ACTATGCCTGCTTATGGTGTTGTTTTGCCATGGGCTTAATAATCG | GGCGTMPAYGVVLPWA, design-12 |
| TJ-O-1281 | TCTTATCCTGAATATGGTGTTGTTTTGCCATGGGCTTAATAATCG | GGCGSYPEYGVVLPWA, design-13 |
| TJ-O-1282 | ACTTATCCTGCTTATGGTTGTGTTTTGCCATGGGCTTAATAATCG | GGCGTYPAYGCVLPWA, design-14 |
| TJ-O-1283 | ACTACTCCTGCTTATGGTGTTATTTTGCCATGGGCTTAATAATCG | GGCGTTPAYGVILPWA, design-15 |

| | | |
|---|---|---|
| TJ-O-1284 | TCTACTCCTGCTTATGGTGTTGTTTTGCCATGGGCTTAATAATCG | GGCGSTPAYGVVLPWA, design-16 |
| TJ-O-1285 | ACTCGTCCTGAATATGGTACTATTTTGCCATGGGCTTAATAATCG | GGCGTRPEYGTILPWA, design-17 |
| TJ-O-1286 | TCTTATCCTGCTTATGGTGTTATTTTGCCATGGGCTTAATAATCG | GGCGSYPAYGVILPWA, design-18 |
| TJ-O-1287 | GCTTATCCTGCTTATGGTACTTGTTTGCCATGGGCTTAATAATCG | GGCGAYPAYGTCLPWA, design-19 |
| TJ-O-1288 | ACTTATCCTGCTTATGGTATTGTTTTGCCATGGGCTTAATAATCG | GGCGTYPAYGIVLPWA, design-20 |
| TJ-O-1289 | ACTTATCCTGAATATGGTGTTGTTTTGCCATGGGCTTAATAATCG | GGCGTYPEYGVVLPWA, design-21 |
| TJ-O-1290 | TCTATTCCTGTTTATGGTGTTGTTTTGCCATGGGCTTAATAATCG | GGCGSIPVYGVVLPWA, design-22 |
| TJ-O-1291 | ACTTATCCTGAATATGGTACTGTTTTGCCATGGGCTTAATAATCG | GGCGTYPEYGTVLPWA, design-23 |
| TJ-O-1292 | ACTTATCCTGCTTATGGTATTACTTTGCCATGGGCTTAATAATCG | GGCGTYPAYGITLPWA, design-24 |
| TJ-O-1293 | TCTTATCCTGCTTATGGTGATGTTTTGCCATGGGCTTAATAATCG | GGCGSYPAYGDVLPWA, design-25 |
| TJ-O-1294 | ACTTTTCCTTCTTATGGTACTTGTTTGCCATGGGCTTAATAATCG | GGCGTFPSYGTCLPWA, design-26 |
| TJ-O-1295 | ACTCATCCTTGTTATGGTGTTGTTTTGCCATGGGCTTAATAATCG | GGCGTHPCYGVVLPWA, design-27 |
| TJ-O-1296 | ACTTTACCTGAATATGGTGTTGTTTTGCCATGGGCTTAATAATCG | GGCGTLPEYGVVLPWA, design-28 |
| TJ-O-1297 | TCTTATCCTTCTTATGGTGTTGTTTTGCCATGGGCTTAATAATCG | GGCGSYPSYGVVLPWA, design-29 |
| TJ-O-1298 | TCTCGTGCTGCTTATGGTGTTGTTTTGCCATGGGCTTAATAATCG | GGCGSRAAYGVVLPWA, design-30 |
| TJ-O-1299 | ACTTTTCCTGCTTATGGTACTGTTTTGCCATGGGCTTAATAATCG | GGCGTFPAYGTVLPWA, design-31 |
| TJ-O-1300 | ACTCATCCTGAATATGGTGTTGTTTTGCCATGGGCTTAATAATCG | GGCGTHPEYGVVLPWA, design-32 |
| TJ-O-1301 | ACTCATCCTTCTTATGGTGTTGTTTTGCCATGGGCTTAATAATCG | GGCGTHPSYGVVLPWA, design-33 |
| TJ-O-1302 | ACTCGTCCTGCTTATGGTACTGTTTTGCCATGGGCTTAATAATCG | GGCGTRPAYGTVLPWA, design-34 |

| | | |
|---|---|---|
| TJ-O-1303 | ACTCGTCCTGGTTATGGTGTTGTTTTGCCATGGGCTTAATAATCG | GGCGTRPGYGVVLPWA, design-35 |
| TJ-O-1304 | TCTTTTCCTGCTTATGGTGCTACTTTGCCATGGGCTTAATAATCG | GGCGSFPAYGATLPWA, design-36 |
| TJ-O-1305 | TCTATTCCTTCTTATGGTGTTGTTTTGCCATGGGCTTAATAATCG | GGCGSIPSYGVVLPWA, design-37 |
| TJ-O-1306 | TCTTTTCCTGCTTATGGTGTTGCTTTGCCATGGGCTTAATAATCG | GGCGSFPAYGVALPWA, design-38 |
| TJ-O-1307 | TCTCGTCCTGCTTATGGTGTTATTTTGCCATGGGCTTAATAATCG | GGCGSRPAYGVILPWA, design-39 |
| TJ-O-1308 | ACTCGTCCTGCTTATGGTGTTGATTTGCCATGGGCTTAATAATCG | GGCGTRPAYGVDLPWA, design-40 |
| TJ-O-1309 | TCTCATCCTTCTTATGGTGTTACTTTGCCATGGGCTTAATAATCG | GGCGSHPSYGVTLPWA, design-41 |
| TJ-O-1310 | GCTTTTCCTGCTTATGGTCAAATTTTGCCATGGGCTTAATAATCG | GGCGAFPAYGQILPWA, design-42 |
| TJ-O-1311 | TCTTATCCTGAATATGGTACTACTTTGCCATGGGCTTAATAATCG | GGCGSYPEYGTTLPWA, design-43 |
| TJ-O-1312 | GGTATTCCTGCTTATGGTACTGTTTTGCCATGGGCTTAATAATCG | GGCGGIPAYGTVLPWA, design-44 |
| TJ-O-1313 | ACTATGCCTGCTTATGGTACTGTTTTGCCATGGGCTTAATAATCG | GGCGTMPAYGTVLPWA, design-45 |
| TJ-O-1314 | TCTATGCCTGCTTATGGTGTTGTTTTGCCATGGGCTTAATAATCG | GGCGSMPAYGVVLPWA, design-46 |
| TJ-O-1315 | GTTTATCCTGCTTATGGTGCTATTTTGCCATGGGCTTAATAATCG | GGCGVYPAYGAILPWA, design-47 |

#### 3. Code and data analysis

Extracted ion chromatograms (EICs) were processed with custom Python code, adapted in part from the analysis code of Voigt and co-workers.[3,16] For each precursor and charge state, the monoisotopic and isotope m/z windows were extracted across all replicates, and the apex MS spectrum was denoised with an adapted AutoPique algorithm:[17–19] the intensity threshold is raised until the autocorrelation of the thresholded spectrum has a narrow central peak, and the final threshold is set at twice the signal-to-noise level above the baseline. Chromatographic peaks were then fitted with an asymmetric (exponentially modified) Gaussian, which describes the tailing elution profile better than a symmetric Gaussian.[20] Fit quality ($R^2$) was retained and used in the peak-quality score alongside isotope-pattern agreement and $MS^2$ fragment matching. Candidate peaks were clustered across replicates by density-based clustering (DBSCAN), and replicates lacking a detectable peak were recorded as non-detections rather than set to zero. Peaks were then manually controlled and adjusted using an interactive reviewing workflow. The complete code can be found under https://github.com/thomashjordan/LCMS_RiPP.

(12) Costa, S. J.; Almeida, A.; Castro, A.; Domingues, L.; Besir, H. The Novel Fh8 and H Fusion Partners for Soluble Protein Expression in Escherichia Coli: A Comparison with

the Traditional Gene Fusion Technology. *Appl. Microbiol. Biotechnol.* **2013**, *97* (15), 6779–6791. https://doi.org/10.1007/s00253-012-4559-1
(13) Costa, S.; Almeida, A.; Castro, A.; Domingues, L. Fusion Tags for Protein Solubility, Purification and Immunogenicity in Escherichia Coli: The Novel Fh8 System. *Front. Microbiol.* **2014**, *5*, 63. https://doi.org/10.3389/fmicb.2014.00063
(14) Luke, J. M.; Carnes, A. E.; Sun, P.; Hodgson, C. P.; Waugh, D. S.; Williams, J. A. Thermostable Tag (TST) Protein Expression System: Engineering Thermotolerant Recombinant Proteins and Vaccines. *J. Biotechnol.* **2011**, *151* (3), 242–250. https://doi.org/10.1016/j.jbiotec.2010.12.011
(15) EMD Chemicals, Inc. *Duet Vectors User Protocol*; Technical Bulletin; Novagen / Merck KGaA, 2011. https://www.sigmaaldrich.com/deepweb/assets/sigmaaldrich/product/documents/221/590/tb340.pdf (accessed 2026-10-15)
(16) Glassey, E.; Zhang, Z.; King, A. M.; Niquille, D. L.; Voigt, C. A. De Novo Design of Ribosomally Synthesized and Post-Translationally Modified Peptides. *Nat. Chem.* **2025**, 1–13. https://doi.org/10.1038/s41557-024-01685-9
(17) Kilgour, D. P. A.; Hughes, S.; Kilgour, S. L.; Mackay, C. L.; Palmblad, M.; Tran, B. Q.; Goo, Y. A.; Ernst, R. K.; Clarke, D. J.; Goodlett, D. R. Autopiquer - a Robust and Reliable Peak Detection Algorithm for Mass Spectrometry. *J. Am. Soc. Mass Spectrom.* **2017**, *28* (2), 253–262. https://doi.org/10.1007/s13361-016-1549-z
(18) Du, P.; Kibbe, W. A.; Lin, S. M. Improved Peak Detection in Mass Spectrum by Incorporating Continuous Wavelet Transform-Based Pattern Matching. *Bioinformatics* **2006**, *22* (17), 2059–2065. https://doi.org/10.1093/bioinformatics/btl355
(19) Deng, F.; Li, H.; Wang, R.; Yue, H.; Zhao, Z.; Duan, Y. An Improved Peak Detection Algorithm in Mass Spectra Combining Wavelet Transform and Image Segmentation. *Int. J. Mass Spectrom.* **2021**, *465*, 116601. https://doi.org/10.1016/j.ijms.2021.116601
(20) Goodman, K. J.; Brenna, J. Thomas. Curve Fitting for Restoration of Accuracy for Overlapping Peaks in Gas Chromatography/Combustion Isotope Ratio Mass Spectrometry. *Anal. Chem.* **1994**, *66* (8), 1294–1301. https://doi.org/10.1021/ac00080a015
